# Novel method for the prediction of drug-drug Interaction based on Gene Expression profiles

**DOI:** 10.1101/2020.05.13.092718

**Authors:** Yh. Taguchi, Turki Turki

**Affiliations:** Department of Physics, Chuo University, Tokyo 112-8551, Japan; Department of Computer Science, King Abdulaziz University, Jeddah, 21589, Saudi Arabia

**Keywords:** Bioinformatics, Drug-drug interaction, Feature extraction, Gene expression, Tensor decomposition, Unsupervised learning

## Abstract

The accurate prediction of new interactions between drugs is important for avoiding unknown (mild or severe) adverse reactions to drug combinations. The development of effective *in silico* methods for evaluating drug interactions based on gene expression data requires an understanding of how various drugs alter gene expression. Current computational methods for the prediction of drug-drug interactions (DDIs) utilize data for known DDIs to predict unknown interactions. However, these methods are limited in the absence of known predictive DDIs. To improve DDIs’ interpretation, a recent study has demonstrated strong non-linear (i.e., dosedependent) effects of DDIs. In this study, we present a new unsupervised learning approach involving tensor decomposition (TD)-based unsupervised feature extraction (FE) in 3D. We utilize our approach to reanalyze available gene expression profiles for *Saccharomyces cerevisiae*. We found that non-linearity is possible, even for single drugs. Thus, non-linear dose-dependence cannot always be attributed to DDIs. Our analysis provides a basis for the design of effective methods for evaluating DDIs.

## 1. Introduction

Although *in silico* methods are thought to be effective strategies for improving the long, expensive process of drug discovery, *in silico* drug discovery is, at best, still under development (Rifaioglu et al., 2018; Vamathevan et al., 2019; Kazmi et al., 2019). In addition to the two main approaches for drug discovery, i.e., ligand-based drug discovery (Bacilieri and Moro, 2006; Pal et al., 2019; Robinson et al., 2020) and structure-based drug discovery (Batool et al., 2019; Taguchi, 2017; Lee et al., 2019), interest in gene expression profile-based drug discovery (Chengalvala et al., 2007) has recently increased. For this process, it is important to understand how drug treatments alter gene expression profiles. However, this is a complex issue owing to the huge number of gene expression alterations resulting from each treatment. The alterations are often non-linear, with non-monotonic dose-dependent effects. This non-linearity often prevents the selection of effective drugs, since it is difficult to determine if expression levels of individual genes are up- or downregulated by particular drug treatments. In drug discovery, analyses of drug-drug interactions (DDIs) are aimed at the prevention or reduction of possible reactions caused by therapeutic drug combinations (Celebi et al., 2019; Yao et al., 2020; Shi et al., 2018; Poleksic and Xie, 2019; Zhang et al., 2019; Langness and Everson, 2016; Masoudi-Sobhanzadeh et al., 2019). Celebi et al. (2019) has generated graph representation of DDI embedded into future vector representations with which three supervised learning, logistic regression, naive Bayes and random forest, were performed. Yao et al. (2020) has formatted DDI as vectors, from which predictors were generated. Shi et al. (2018) generated DDI matrices from known DDI and predicted DDI with a newly identified drug from the matricies. Poleksic and Xie (2019) generated two adverse effect (ADE) data bases, i.e. associations between ADE and individual drugs and those between ADE and drug pairs, from which they predicted unknown DDI. Zhang et al. (2019) validated DDI between two specific drugs experimentally. Langness and Everson (2016) evaluated DDIs of HCV patients experimentally. Masoudi-Sobhanzadeh et al. (2019) generadted DDI database from which they can predict new DDI. However, the above-mentioned methods are not capable of predicting unknown interactions if data for known DDIs are not available. Hence, Lukac̆is̆in and Bollenbach (2019) evaluated how DDIs affect gene expression profiles in a combinatorial manner; they found that DDIs can exhibit convex relationships with gene expression profiles. Although Lukac̆is̆in and Bollenbach (2019) identified that convex dose dependence was caused by DDI, it is unclear if it was really caused by DDI. To verify if it was indeed caused by DDI, we compared dose dependence between single dose and combinatorial dose; if convex doses can be observed even in a single dose, we can deny the possibility that convex dose dependence was primarily caused by DDI. To identify this point, we employed integrated analysis using tensor decomposition (TD) based unsupervised feature extraction (FE) as well as principal component analysis (PCA) based unsupervised feature extraction (FE).

## 2. Materials and Methods

### 2.1. Gene expression profiles

Gene expression profiles were downloaded from Gene Expression Omnibus (GEO) (Clough and Barrett, 2016) with GEO ID GSE138256. The processed file named “GSE138256_GeneExpression.csv.gz” was used. The file “GSE138256_SampleConditionsAndOrdering.csv.gz” was also downloaded for sample annotations. These dataset sets were composed of gene expression profiles of *Saccharomyces cerevisiae* treated with individual drugs or pairs of the following four drugs: myriocin, cycloheximide, LiCl, and rapamycin. When *S. cerevisiae* was treated with pairs of drugs, and the combinatorial dose was carefully tuned to ensure the same growth rate, to the greatest extent possible.

### 2.2. PCA

PCA was applied to individual pairs of drugs. For the *i*th gene expression level and *j*th dose, *x_ij_* ∈ ℝ^*N*×*M*^, where *N* is total number of genes (i.e., 6717) and *M* is total number of combinatorial doses for each pair of drugs (Table 1). *x_i__j_* is normalized as Σ*_i_ x_ij_* = 0 and 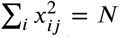. PCA was applied to such *x_ij_* that PC loadings and PC scores were attributed to samples and genes, respectively. Lowess smoothing was applied to PC loadings to reduce noise signals using the lowess command implemented in R (R Core Team, 2019).

**Table 1.**
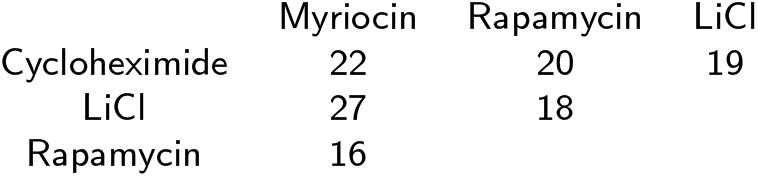
Number of doses tested for drug combinations.

### 2.3. TD-based unsupervised FE

TD-based unsupervised FE was applied to gene expression profiles. Gene expression profiles were formatted as a tensor, *x_ijk_* ∈ ℝ^*N*×16×6^, representing the expression of the *i*th gene and *j*th combinatorial dose of the *k*th pair of drugs. Since the number of combinatorial doses varied among pairs, the minimum number of combinatorial doses, 16, was employed. When more combinatorial doses were tested for specific pairs of drugs, some measurements were discarded, attempting to maintain equal intervals between doses. *x_ijk_* was normalized as Σ*_i_x_ijk_* = 0 and 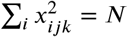. Higher order singular value decomposition (HOSVD) (Taguchi, 2019b) was applied to *x_ijk_* to obtain the following:

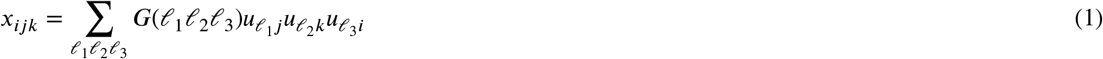

where *G*(*ℓ*_1_*ℓ*_2_*ℓ*_3_) ∈ ℝ^*N*×16×6^ is a core tensor, and 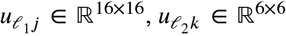, and 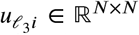 are the singular value vectors defined as the column vectors of orthogonal matrices. 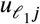 is attributed to the *j*th dose, 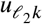 is attributed to the *k*th pair of drugs, and 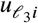 is attributed to the *i*th gene. Lowess smoothing was also applied to singular value vectors to reduce noise using the lowess command implemented in R (R Core Team, 2019). To select 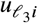 for gene selection in subsequent analyses, it was first necessary to determine which 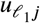 and 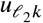 are biologically meaningful. After identifying such *ℓ*_1_ and *ℓ*_2_, it is necessary to identify the *ℓ*_3_ associated with *G*(*ℓ*_1_*ℓ*_2_*ℓ*_3_) having the largest absolute values given fixed *ℓ*_1_ and *ℓ*_2_. With the selected *ℓ*_3_, *P*-values, *P_i_*, were obtained for the *i*th gene as follows:

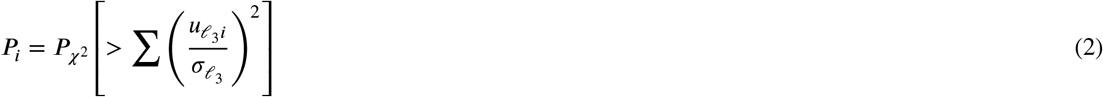

where 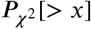 is the cumulative probability of the *χ*^2^ distribution with an argument larger than *x*. The summation is taken over only selected *ℓ*_3_s to compute *P*-values. *P_i_* was corrected with the BH criterion (Taguchi, 2019b) and genes with *P* < 0.01 were selected.

Gene expression levels in response to a single dose (Table 2) were also formatted as a three-mode tensor, *x_ijk_* ∈ ℝ^*N*×14×4^, which represents the *i*th gene expression level for the *j*th dose of the *k*th drug. Since the number of unique doses was 14, excluding cycloheximide, the total number of doses for cycloheximide was also set to 14, and two replicates were included for three doses. The same procedure employed for the analysis of combinatorial drug treatments was repeated, and genes were selected.

**Table 2.** Number of doses tested for individual drugs. Numbers in parentheses indicate unique doses.

| drug | Number of samples |
| --- | --- |
| Myriocin | 25 (14) |
| Cycloheximide | 23 (11) |
| LiCl | 28 (14) |
| Rapamycin | 30 (14) |

### 2.4. Enrichment Analysis

The gene symbols of selected genes were uploaded to YeastEnrichr, a yeast version of Enricher (Kuleshov et al., 2016; Chen et al., 2013) prepared for humans, as well as to g:profiler (Raudvere et al., 2019)

#### 2.4.1. KEGG pathway

We uploaded 157 gene symbols selected by TD based unsupervised FE for single dose treatment as well as 77 gene symbols selected by TD based unsupervised FE for single dose treatment to YeastEnrichr. Then, we referred to the KEGG pathway category of YeastEnrichr and downloaded the results as a text file.

#### 2.4.2. GO BP

We uploaded 157 gene symbols selected by TD based unsupervised FE for single dose treatment as well as 77 gene symbols selected by TD based unsupervised FE for single dose treatment to YeastEnrichr. Then, we referred to the GO BP category of YeastEnrichr and downloaded the results as a text file.

#### 2.4.3. GO CC

We uploaded 157 gene symbols selected by TD based unsupervised FE for single dose treatment as well as 77 gene symbols selected by TD based unsupervised FE for single dose treatment to YeastEnrichr. Then, we referred to the GO CC category of YeastEnrichr and downloaded the results as a text file.

#### 2.4.4. GO MP

We uploaded 157 gene symbols selected by TD based unsupervised FE for single dose treatment as well as 77 gene symbols selected by TD based unsupervised FE for single dose treatment to YeastEnrichr. Then, we referred to the GO MP category of YeastEnrichr and downloaded the results as a text file.

### 2.5. Flow Chart

Fig. 1 shows a diagram of the analyses performed in this study.

**Figure 1:**
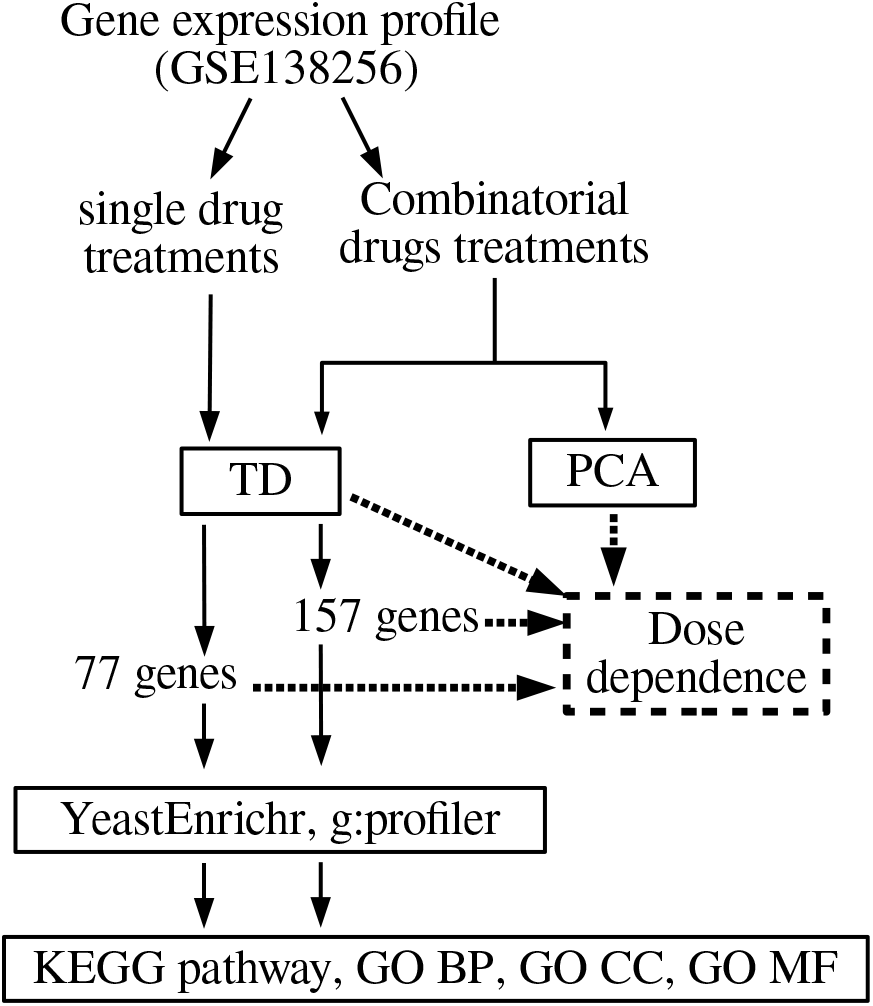
Flowchart of analyses performed in this study.

## 3. Results

We first applied PCA to gene expression levels, *x_ij_*, attributed to individual pairs of drug treatments to attempt to reproduce the previous observations (Lukac̆is̆in and Bollenbach, 2019). In the previous study (Lukac̆is̆in and Bollenbach, 2019), the first PC loading takes constant values, independent of dose, while the second and the third PC loadings exhibit linear and convex dose-dependence, regardless of pairs of drugs. In our analysis, the first PC also took constant values, irrespective of the drug combination (not shown). However, the second and third PC loadings behaved slightly differently (Fig. S1). For the combination of cycloheximide and LiCl, although the second and the third PC loadings behaved as expected, the fourth PC loading also showed concave or convex dose-dependence. Since the fourth PC loadings were not discussed in the previous study (Lukac̆is̆in and Bollenbach, 2019), it is possible that this observation was present in the previous research, but not reported. Nevertheless, for the combination of LiCl and rapamycin, the second PC loadings did not have linear dependence but instead showed stepwise dependence, which was not reported in the previous study (Lukac̆is̆in and Bollenbach, 2019). Additionally, for the remaining four combinatorial cases, the second and third PCs did not always have linear and concave or convex dose-dependence, respectively.

It is possible that the disagreement between the present study (in which the third PC did not always have linear and concave or convex dependence on the dose, respectively) and the previous study (Lukac̆is̆in and Bollenbach, 2019) could be explained by insufficient pre-processing of gene expression profiles. To evaluate this possibility, we applied HOSVD to the tensor, *x_ijk_*, generated from combinatorial drug treatments (for a summary of this analysis, see Fig. 2). It was evident that *u*_1*j*_ takes constant values independent of dose density, *u*_2*j*_ has a linear dependence on dose density, and *u*_3*j*_ has concave or convex dependence on dose density, as observed in the original study (Lukac̆is̆in and Bollenbach, 2019) (Fig. S2). As a result, we could identify singular value vectors that exhibit nonlinear dose dependence independent of individual drugs used using TD whereas PCA could not identify common PC loading that exhibits non-linearity independent of individual drugs used. This suggests the superiority of TD-based unsupervised FE to identify essential features, regardless of pre-processing.

**Figure 2:**
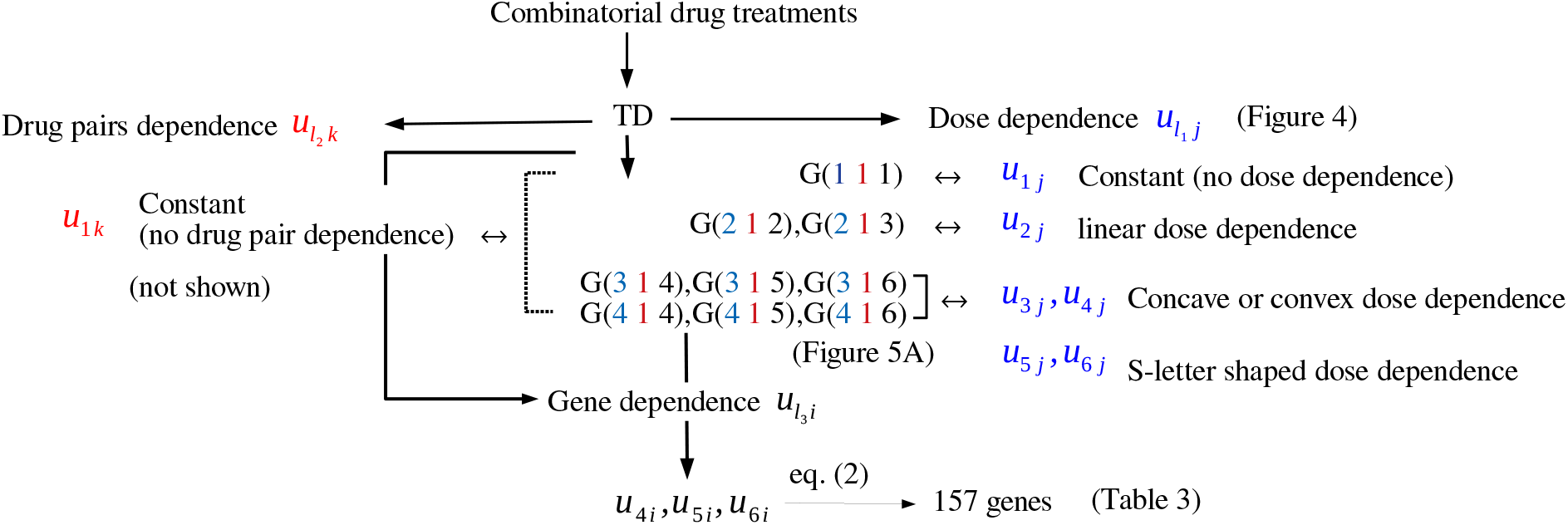
Summary of TD-based unsupervised FE applied to gene expression with combinatorial drug treatments. As denoted in Materials and Methods, we first need to identify which singular value vectors attributed to does dependence have non-linearity as well as which singular value vectors attributed to drugs have constant values independent of drugs used. Then we selected singular value vectors attributed to genes and associated with selected singular value vectors attributed to genes and drugs by investigating *G*s. Finally, we could selected genes based upon selected singular value vectors attributed to genes.

One might notice that *u*_4*j*_ is also concave or convex and *u*_5*j*_ and *u*_6*j*_ have more complex shapes (S-letter shaped). To see if these shapes are artifacts or reflect individual gene expression profiles, we focused on genes whose expression levels are likely coincident with these concave and convex shapes. Since we noticed that *u*_1*k*_ had constant values over combinatorial treatments, we searched for *G*(*ℓ*_1_ 1 *ℓ*_3_) with the largest contribution to *ℓ*_1_ = 3, 4 and relatively small contributions to *ℓ*_1_ = 1, 2, which are associated with constant or linear dependence (Fig. S3A). It is obvious that *G*(*ℓ*_1_ 1 1) had the largest contribution to *ℓ*_1_ = 1, i.e., a constant (or dose density-independent) profile, while *G*(*ℓ*_1_ 1 2) and *G*(*ℓ*_1_ 1 3) had the largest contribution to *ℓ*_1_ = 2, i.e., linearly dependent on dose density. Thus, to identify 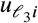 associated with profiles other than constant or linear profiles, we employed 4 ≤ *ℓ*_3_ ≤ 6 for gene selection. Based on *P*-values and correction as described in the Materials and Methods, we selected 157 genes (Table 3).

**Table 3.**
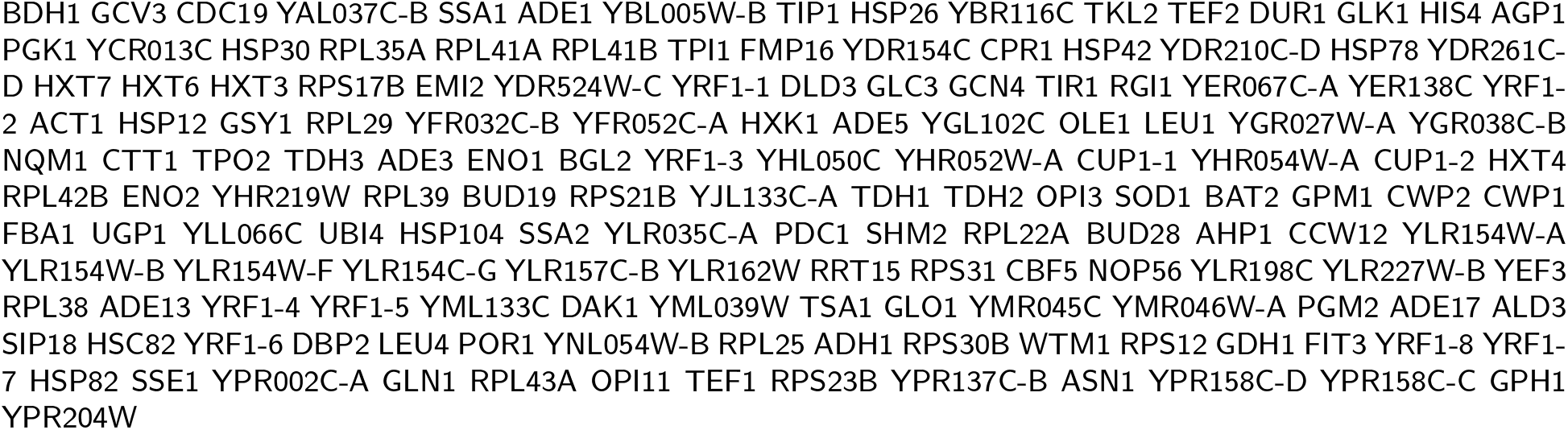
List of 157 genes selected by TD-based unsupervised FE toward combinatorial drugs treatments. These genes are associated with concave or convex dose-dependence, since they are expected to be associated with *u*_3*j*_ and *u*_4*j*_ (Fig. S2).

To see if the 157 genes selected in this analysis were associated with concave, convex, or the more complicated S-shaped pattern, we plotted Lowess-smoothed expression profiles of two representative genes, BDH1 and SSA1, as shown in Fig. 3 (note that gene expression profiles of other genes are available as supplementary materials). Gene expression profiles have distinct dose-dependence for drug combinations, although concave, convex, and S-shaped profiles were observed. Thus, the profiles shown in Fig. S2 were not artifacts but reflected the expression patterns of individual genes. TD-based unsupervised FE not only generated singular value vectors that represent constant, linear, concave, or convex dependence on dose density but also characterizes more complicated (S-letter shaped) profiles for individual genes. Thus, it is an advantageous strategy for analyzing gene expression profiles obtained under distinct conditions in an integrated manner.

**Figure 3:**
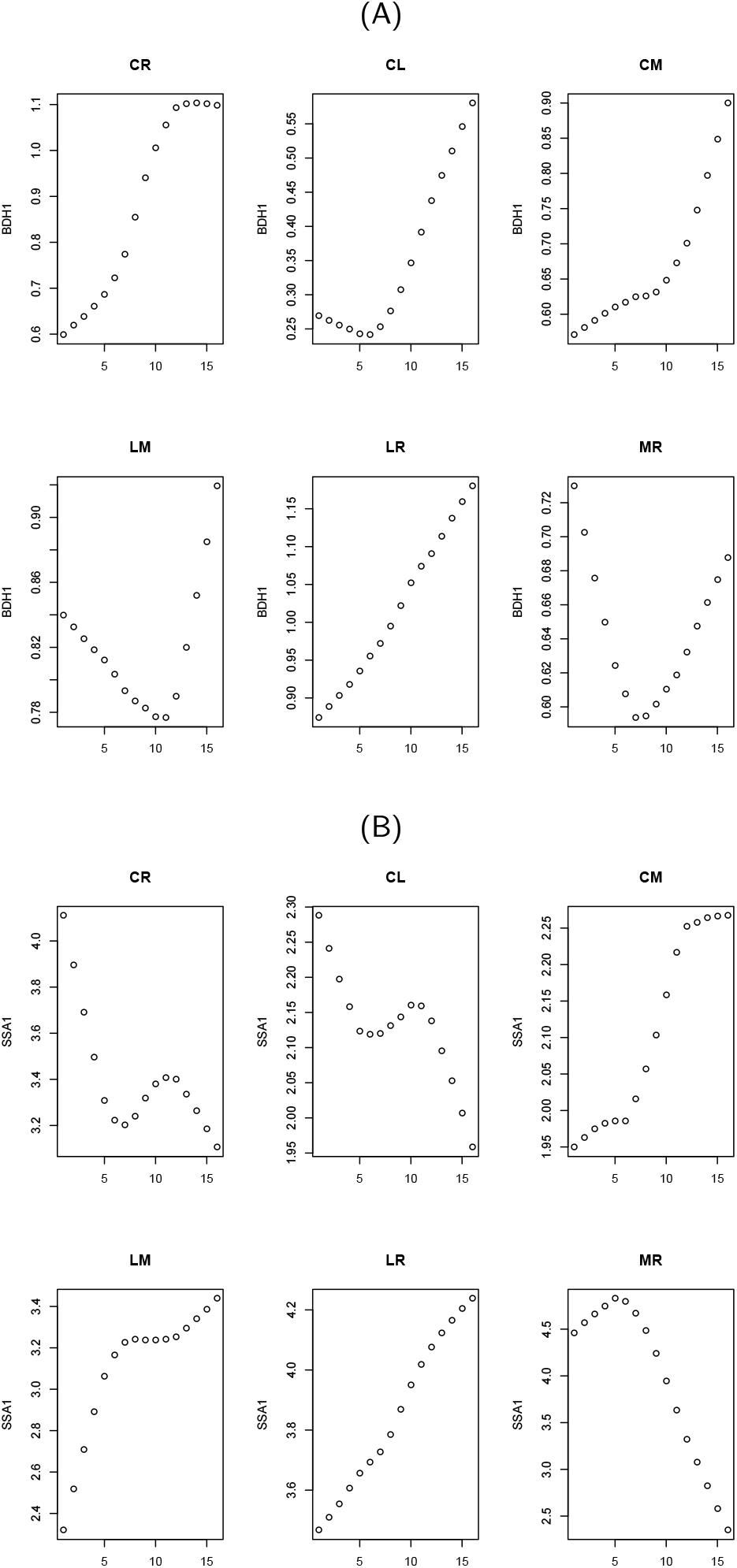
Lowess-smoothed gene expression profiles for BDH1 (A) and SSA1 (B). Two letters above each panel show the combinations of drugs: M: Myriocin C: Cycloheximide, L: LiCl, R: Rapamycin

Next, we validated the selected genes by evaluating their biological functions. We uploaded 157 genes to Yeast-Enrichr and found enrichment for numerous biological functions. In particularly, we detected 23 significant biological terms in the KEGG pathway analysis (see 10 top-ranked terms in Table 4), 91 terms in the GO Biological Process (BP) category (see 10 top-ranked terms in Table 5), 22 terms in the GO Cellular Component (CC) category (see 10 top-ranked terms in Table 6), and 35 terms in the GO Molecular Function (MF) category (see 10 top-ranked terms in Table 7). Thus, the selected genes had key biological functions. To confirm the observed enrichment, we also analyzed the genes using g:profiler. Although we obtained fewer significantly enriched terms, there were 219 biological terms, including KEGG pathways and GO BP, MF, and CC terms (lists of individual biological terms obtained using YeastEnrichr and g:profiler are available as supplementary materials). Thus, the biological significance of the selected genes is not database-dependent, supporting the robustness and reliability of the analysis.

**Table 4.**
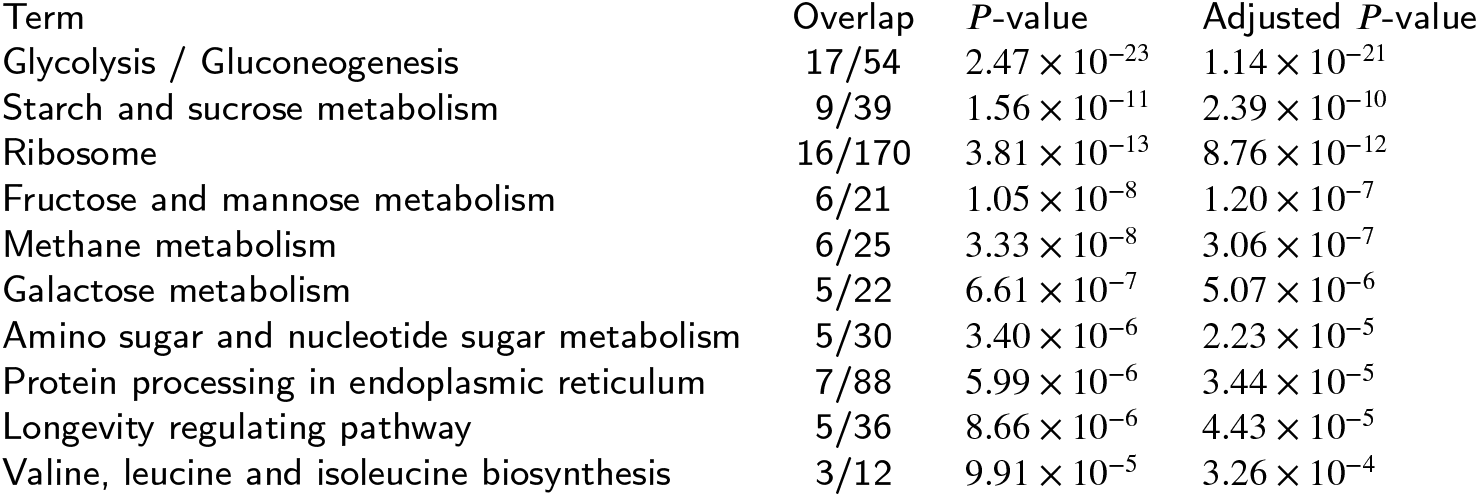
Ten top-ranked KEGG 2019 terms for 157 genes selected by TD-based unsupervised FE when combinatorial drug treatments were employed.

| Term | Overlap | *P*-value | Adjusted *P*-value |
| --- | --- | --- | --- |
| Glycolysis / Gluconeogenesis | 17/54 | $2.47 \times 10^{-23}$ | $1.14 \times 10^{-21}$ |
| Starch and sucrose metabolism | 9/39 | $1.56 \times 10^{-11}$ | $2.39 \times 10^{-10}$ |
| Ribosome | 16/170 | $3.81 \times 10^{-13}$ | $8.76 \times 10^{-12}$ |
| Fructose and mannose metabolism | 6/21 | $1.05 \times 10^{-8}$ | $1.20 \times 10^{-7}$ |
| Methane metabolism | 6/25 | $3.33 \times 10^{-8}$ | $3.06 \times 10^{-7}$ |
| Galactose metabolism | 5/22 | $6.61 \times 10^{-7}$ | $5.07 \times 10^{-6}$ |
| Amino sugar and nucleotide sugar metabolism | 5/30 | $3.40 \times 10^{-6}$ | $2.23 \times 10^{-5}$ |
| Protein processing in endoplasmic reticulum | 7/88 | $5.99 \times 10^{-6}$ | $3.44 \times 10^{-5}$ |
| Longevity regulating pathway | 5/36 | $8.66 \times 10^{-6}$ | $4.43 \times 10^{-5}$ |
| Valine, leucine and isoleucine biosynthesis | 3/12 | $9.91 \times 10^{-5}$ | $3.26 \times 10^{-4}$ |

**Table 5.** Ten top-ranked GO biological process (BP) 2018 terms for 157 genes selected by TD-based unsupervised FE when combinatorial drug treatments were employed.

| Term | Overlap | *P*-value | Adjusted *P*-value |
| --- | --- | --- | --- |
| glycolytic process (GO:0006096) | 13/20 | $1.92 \times 10^{-23}$ | $3.06 \times 10^{-21}$ |
| ATP generation from ADP (GO:0006757) | 13/20 | $1.92 \times 10^{-23}$ | $3.06 \times 10^{-21}$ |
| nicotinamide nucleotide metabolic process (GO:0046496) | 13/24 | $6.00 \times 10^{-22}$ | $6.38 \times 10^{-20}$ |
| pyruvate metabolic process (GO:0006090) | 14/33 | $1.35 \times 10^{-21}$ | $1.07 \times 10^{-19}$ |
| carbohydrate catabolic process (GO:0016052) | 13/28 | $8.77 \times 10^{-21}$ | $5.59 \times 10^{-19}$ |
| glucose metabolic process (GO:0006006) | 13/32 | $7.92 \times 10^{-20}$ | $4.21 \times 10^{-18}$ |
| gluconeogenesis (GO:0006094) | 9/16 | $9.80 \times 10^{-16}$ | $3.91 \times 10^{-14}$ |
| hexose biosynthetic process (GO:0019319) | 9/16 | $9.80 \times 10^{-16}$ | $3.91 \times 10^{-14}$ |
| cytoplasmic translation (GO:0002181) | 16/162 | $1.79 \times 10^{-13}$ | $6.35 \times 10^{-12}$ |
| translation (GO:0006412) | 20/297 | $2.22 \times 10^{-13}$ | $7.07 \times 10^{-12}$ |

**Table 6.** Ten top-ranked GO cellular component (CC) 2018 terms for 157 genes selected by TD-based unsupervised FE when combinatorial drug treatments were employed.

| Term | Overlap | *P*-value | Adjusted *P*-value |
| --- | --- | --- | --- |
| cytosolic part (GO:0044445) | 20/204 | $1.64 \times 10^{-16}$ | $5.64 \times 10^{-15}$ |
| cytosol (GO:0005829) | 32/676 | $1.82 \times 10^{-16}$ | $5.64 \times 10^{-15}$ |
| retrotransposon nucleocapsid (GO:0000943) | 15/91 | $4.40 \times 10^{-16}$ | $9.10 \times 10^{-15}$ |
| nucleus (GO:0005634) | 44/1599 | $7.80 \times 10^{-14}$ | $1.21 \times 10^{-12}$ |
| cytosolic ribosome (GO:0022626) | 17/185 | $1.00 \times 10^{-13}$ | $1.25 \times 10^{-12}$ |
| mitochondrion (GO:0005739) | 33/1063 | $8.36 \times 10^{-12}$ | $8.64 \times 10^{-11}$ |
| fungus-type cell wall (GO:0009277) | 12/132 | $5.56 \times 10^{-10}$ | $4.93 \times 10^{-9}$ |
| cytosolic large ribosomal subunit (GO:0022625) | 10/101 | $6.94 \times 10^{-9}$ | $5.38 \times 10^{-8}$ |
| large ribosomal subunit (GO:0015934) | 10/104 | $9.24 \times 10^{-9}$ | $6.36 \times 10^{-8}$ |
| cytosolic small ribosomal subunit (GO:0022627) | 6/71 | $2.00 \times 10^{-5}$ | $1.24 \times 10^{-4}$ |

**Table 7.** Ten top-ranked GO molecular function (MF) 2018 terms for 157 genes selected by TD-based unsupervised FE when combinatorial drug treatments were employed.

| Term | Overlap | *P*-value | Adjusted *P*-value |
| --- | --- | --- | --- |
| helicase activity (GO:0004386) | 12/39 | $1.16 \times 10^{-16}$ | $1.30 \times 10^{-14}$ |
| RNA-directed DNA polymerase activity (GO:0003964) | 12/48 | $1.95 \times 10^{-15}$ | $1.09 \times 10^{-13}$ |
| DNA-directed DNA polymerase activity (GO:0003887) | 12/59 | $2.91 \times 10^{-14}$ | $1.09 \times 10^{-12}$ |
| DNA polymerase activity (GO:0034061) | 12/61 | $4.47 \times 10^{-14}$ | $1.25 \times 10^{-12}$ |
| nuclease activity (GO:0004518) | 12/64 | $8.27 \times 10^{-14}$ | $1.85 \times 10^{-12}$ |
| ribonuclease activity (GO:0004540) | 12/66 | $1.22 \times 10^{-13}$ | $2.28 \times 10^{-12}$ |
| RNA binding (GO:0003723) | 24/477 | $4.39 \times 10^{-13}$ | $7.02 \times 10^{-12}$ |
| DNA helicase activity (GO:0003678) | 8/35 | $2.37 \times 10^{-10}$ | $3.32 \times 10^{-9}$ |
| purine ribonucleoside triphosphate binding (GO:0035639) | 7/78 | $2.66 \times 10^{-6}$ | $3.31 \times 10^{-5}$ |
| nucleoside-triphosphatase activity (GO:0017111) | 10/195 | $3.33 \times 10^{-6}$ | $3.74 \times 10^{-5}$ |

In this study, we partially reproduced the original observations (Lukac̆is̆in and Bollenbach, 2019) by PCA; however, TD-based unsupervised FE allowed us to obtain the same results more robustly and reliably. Based on our findings, the expression levels of some genes exhibit non-linear dependence on the dose density. However, non-linear dependence on the dose density was also observed for treatment with single drugs (see Additional file 2 (Taguchi, 2019a)). Thus, it is not clear whether the concave or convex dependence on the dose can be explained by DDIs. To further evaluate the ability of individual drugs to result in non-linear dose-dependence, we applied the newly developed TD-based unsupervised FE to the alternative tensor, *x_ijk_*, generated from gene expression profiles of *S. cerevisiae* treated with single drugs (see Materials and Methods and Fig. 4 for the summary of this analysis). Fig. S4 shows the Lowess-smoothed 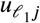, 1 ≤ *ℓ*_1_ ≤ 6 for single-drug treatments. Contrary to our expectation, non-linearity was substantially greater than that shown in Fig. S2 based on combinatorial treatments. Linear dependence was minimal, and an S-letter shaped pattern was observed prior to concave or convex patterns. To determine if the strong non-linearity is associated with individual gene expression profiles, we selected genes associated with singular value vectors that exhibit non-linearity, as shown in Fig. 3. We initially noticed that *u*_1*k*_ has constant values over four single-drug treatments, as in the case of combinatorial drug treatments (not shown). Thus we need to find ***G***(*ℓ*_1_ 1 *ℓ*_3_) with the largest contribution to 3 ≤ *ℓ*_1_ ≤ 6 and relatively small contributions to *ℓ*_1_ = 1, 2, indicating constant or linear dependence (Fig. S4).

**Figure 4:**
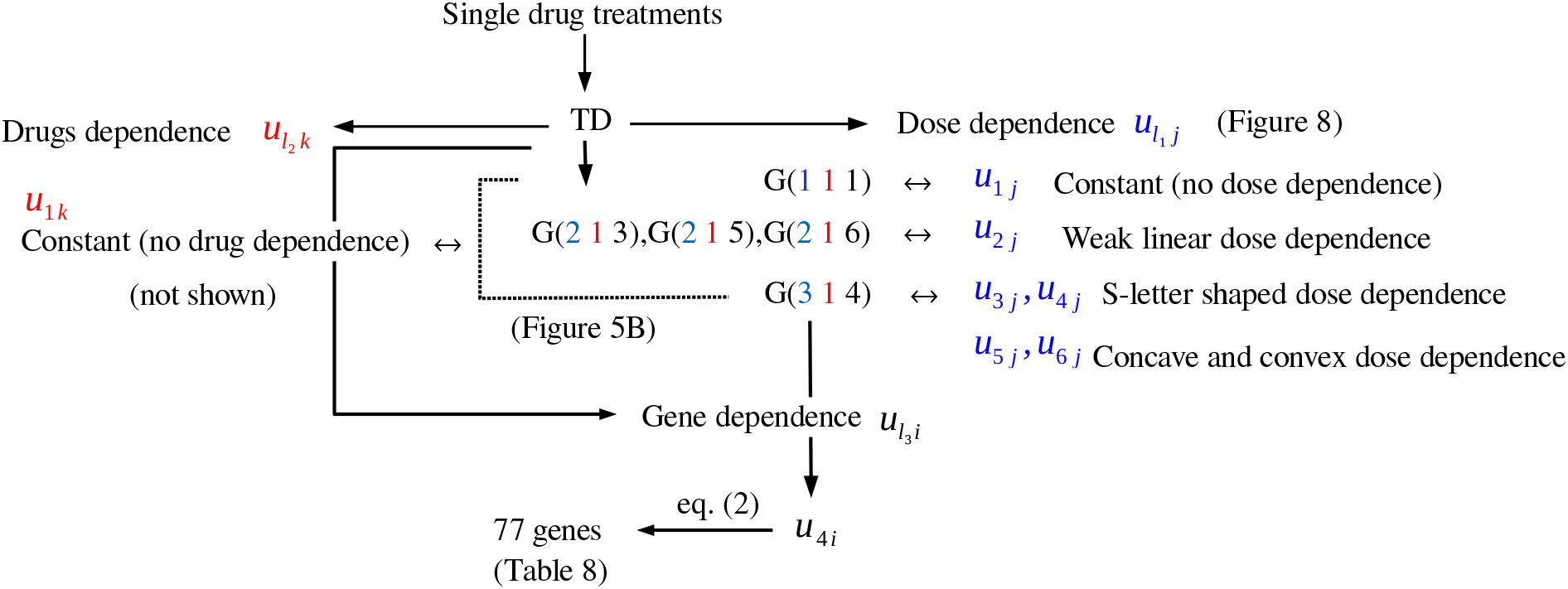
Summary of TD-based unsupervised FE applied to gene expression with single drug treatments

Observed patterns (Fig. S3B) exhibited greater non-linearity than those shown in Fig. S3A for combinatorial treatments. When drugs were treated in combinatorial manner, *G*(1 1 1) has the largest absolute values among *G*(1 1 *ℓ*_3_); this means that constant profiles are associated with the first singular value vector, *u_ℓi_*. *G*(2 1 3), *G*(2 1 5), and *G*(2 1 6) had the largest absolute values among *G*(2 1 *ℓ*_3_), indicating that linear profiles are associated with the second singular value vector, *u*_2*i*_, as well as the third singular value vector, *u*_3*i*_. Nevertheless, in Fig. S3B, although ***G***(1 1 1) had the largest absolute values among ***G***(1 1 *ℓ*_3_)s, *G*(2 1 *ℓ*_3_), 2 ≤ *ℓ*_3_ ≤ 6, had substantial contributions, indicating that there is no clear separation between genes whose expression profiles are associated with dose-dependence represented by *u*_2*i*_, which are most likely linear profiles, and those with dose-dependence represented by *u*_3*i*_ to *u*_6*i*_, likely representing non-linear profiles, i.e., concave, convex, and S-letter shaped profiles. Thus, to select genes with strong non-linear dependence on dose, we selected *ℓ*_3_ = 4, since *G*(3 1 4) had the largest absolute values among ***G***(3 1 *ℓ*_3_) and ultimately identified 77 significant genes (Table 8).

**Table 8.**
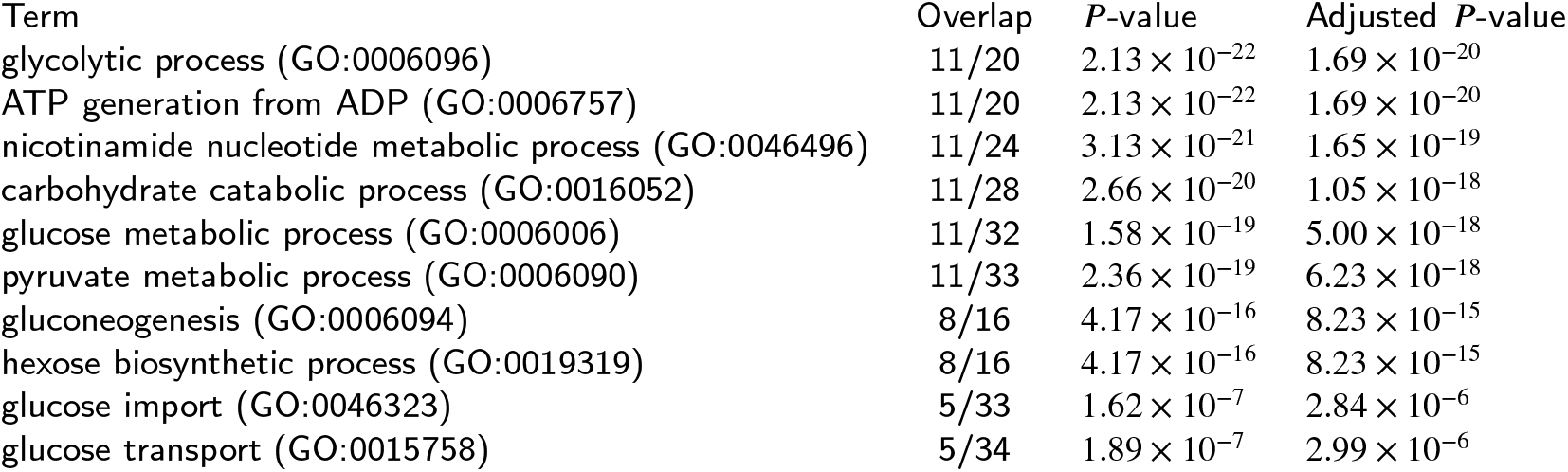
List of 77 genes selected by TD-based unsupervised FE toward single drugs treatments. These genes are associated with non-linear dose-dependence, since they are expected to be associated with *u*_3*j*_ (Fig. S4).

Fig. 5 shows Lowess-smoothed gene expression profiles of two representative genes whose expression levels are also shown in Fig. 3 with respect to combinatorial drug treatments (expression profiles of other genes are available as supplementary materials). Non-linearity of dose-dependence is not clearly reduced. Accordingly, the strong non-linearity of dose-dependence observed for combinatorial drug treatments may not reflect DDIs but rather the non-linearity on the dose-dependence of the expression of individual genes (as shown in Fig. 6, showing an extensive overlap of selected genes for single and combinatorial drug treatments). In conclusion, in our comparison of gene expression profiles between single and combinatorial drug treatments, we did not obtain clear evidence that the strong non-linearity between gene expression levels and dose can be directly attributed to DDIs.

**Figure 5:**
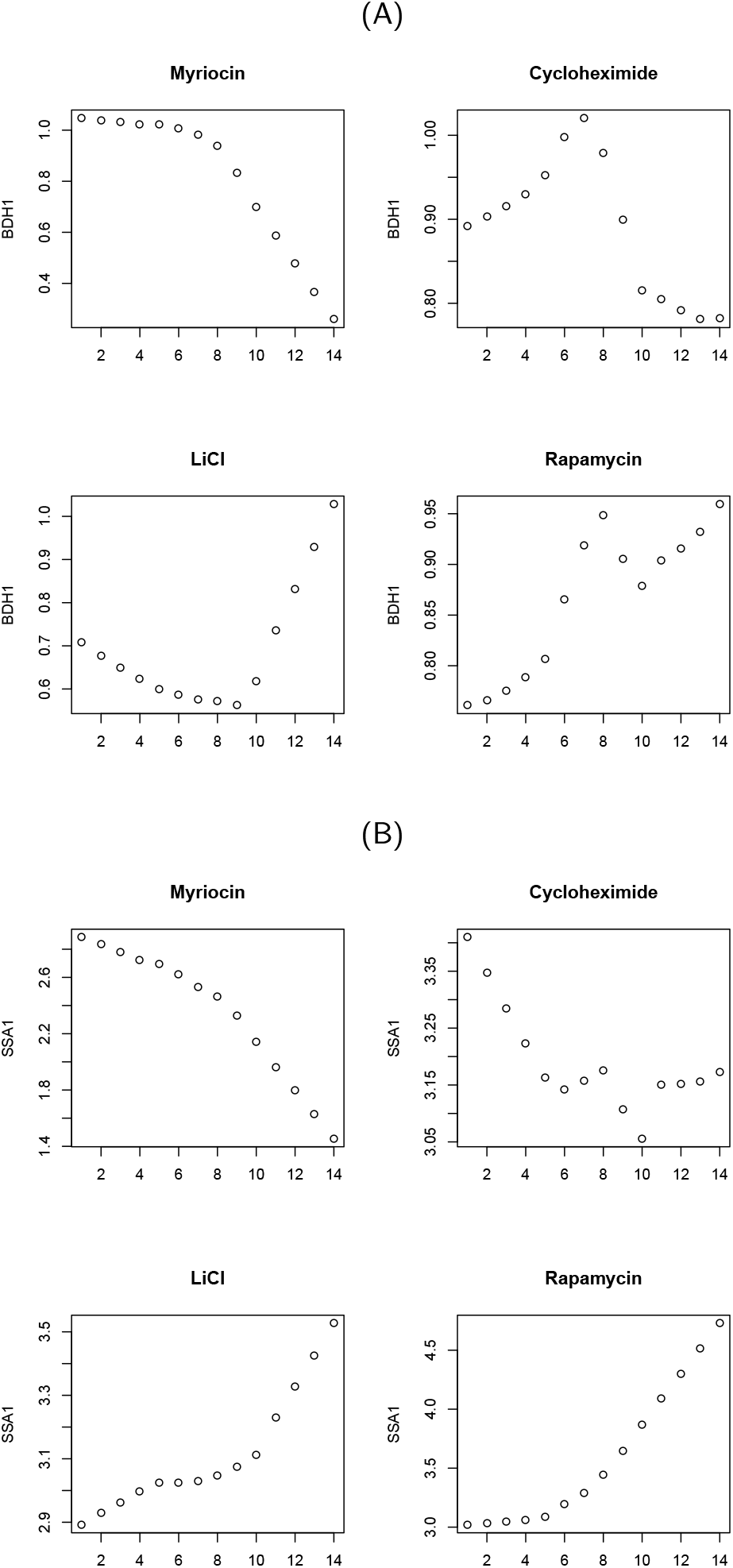
Lowess-smoothed gene expression profiles for BDH1 (A) and SSA1 (B). Two letters above each panel show the combinations of drugs: M: Myriocin C: Cycloheximide, L: LiCl, R: Rapamycin

**Figure 6:**
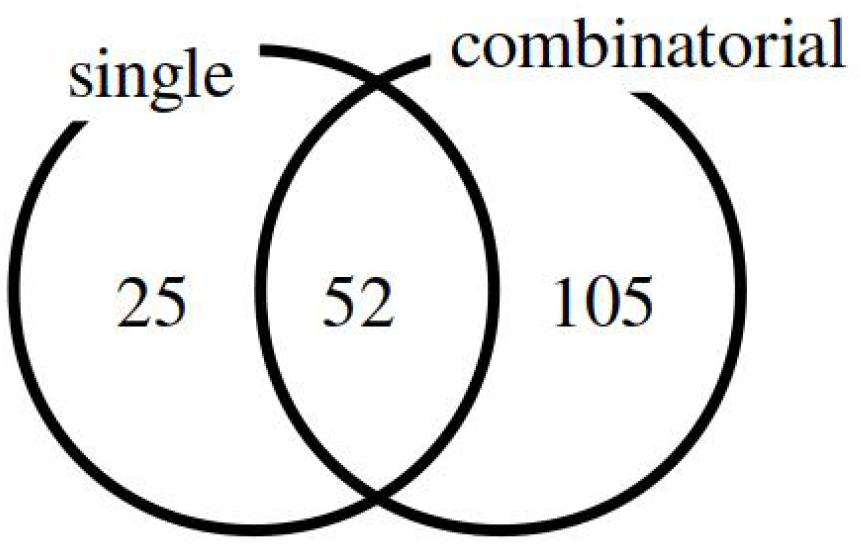
Venn diagram of genes selected for combinatorial and single drug treatments.

We further evaluated the biological significance of the 77 genes selected for treatment with single drugs. We identified a number of significant (adjusted *P*-values less than 0.05) KEGG pathways (Table 9) and GO terms in the BP (Table 10), CC (Table 11), and MF categories (Table 12). Thus, the selected genes were biologically relevant.

**Table 9.** Ten top-ranked KEGG 2019 terms for 77 genes selected by TD-based unsupervised FE when single drug treatments were employed.

| Term | Overlap | *P*-value | Adjusted *P*-value |
| --- | --- | --- | --- |
| Glycolysis / Gluconeogenesis | 14/54 | $1.30 \times 10^{-22}$ | $3.76 \times 10^{-21}$ |
| Starch and sucrose metabolism | 7/39 | $1.32 \times 10^{-10}$ | $1.92 \times 10^{-9}$ |
| Fructose and mannose metabolism | 5/21 | $1.44 \times 10^{-8}$ | $1.34 \times 10^{-7}$ |
| Galactose metabolism | 5/22 | $1.85 \times 10^{-8}$ | $1.34 \times 10^{-7}$ |
| Amino sugar and nucleotide sugar metabolism | 5/30 | $9.80 \times 10^{-8}$ | $5.68 \times 10^{-7}$ |
| Longevity regulating pathway | 5/36 | $2.55 \times 10^{-7}$ | $1.23 \times 10^{-6}$ |
| Methane metabolism | 3/25 | $1.19 \times 10^{-4}$ | $4.92 \times 10^{-4}$ |
| Protein processing in endoplasmic reticulum | 4/88 | $3.71 \times 10^{-4}$ | $1.34 \times 10^{-3}$ |
| Ribosome | 5/170 | $5.04 \times 10^{-4}$ | $1.63 \times 10^{-3}$ |
| Tyrosine metabolism | 2/14 | $1.29 \times 10^{-3}$ | $3.75 \times 10^{-3}$ |

**Table 10.**
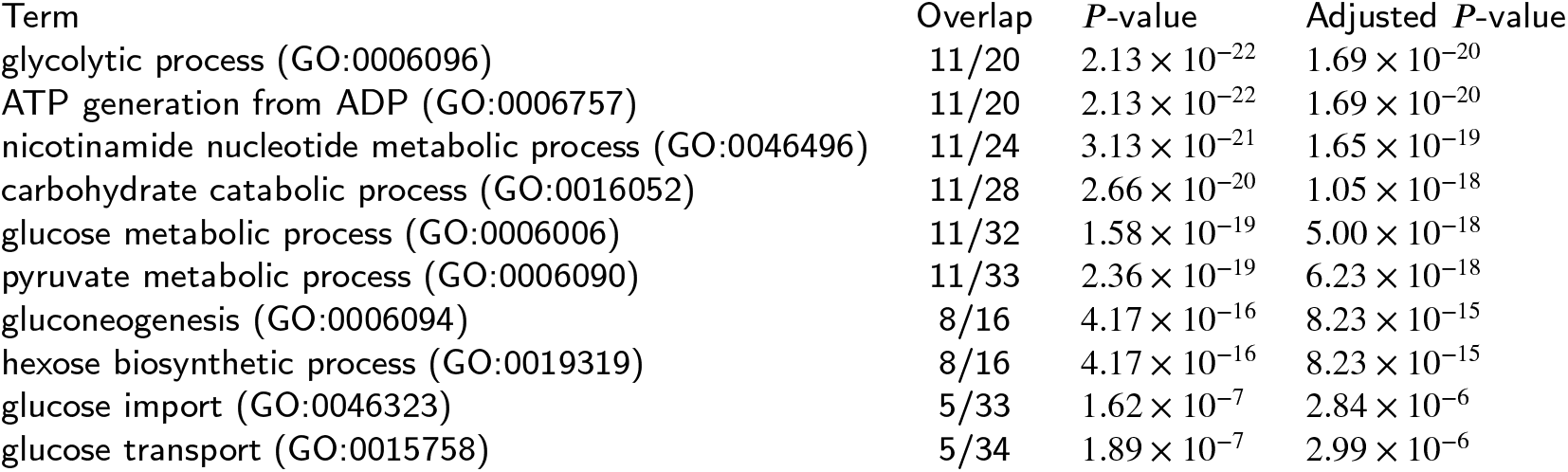
Ten top-ranked GO BP 2018 terms for 77 genes selected by TD-based unsupervised FE when single drug treatments were employed.

**Table 11.** Ten top-ranked GO CC 2018 terms for 77 genes selected by TD-based unsupervised FE when single drug treatments were employed.

| Term | Overlap | *P*-value | Adjusted *P*-value |
| --- | --- | --- | --- |
| retrotransposon nucleocapsid (GO:0000943) | 22/91 | $1.62 \times 10^{-34}$ | $6.30 \times 10^{-33}$ |
| nucleus (GO:0005634) | 34/1599 | $9.70 \times 10^{-18}$ | $1.89 \times 10^{-16}$ |
| cytosol (GO:0005829) | 17/676 | $5.94 \times 10^{-10}$ | $7.73 \times 10^{-9}$ |
| cytosolic part (GO:0044445) | 8/204 | $1.18 \times 10^{-6}$ | $1.15 \times 10^{-5}$ |
| cytosolic small ribosomal subunit (GO:0022627) | 5/71 | $7.92 \times 10^{-6}$ | $6.18 \times 10^{-5}$ |
| mitochondrion (GO:0005739) | 15/1063 | $1.10 \times 10^{-5}$ | $7.12 \times 10^{-5}$ |
| small ribosomal subunit (GO:0015935) | 5/79 | $1.34 \times 10^{-5}$ | $7.45 \times 10^{-5}$ |
| cytosolic ribosome (GO:0022626) | 6/185 | $7.94 \times 10^{-5}$ | $3.87 \times 10^{-4}$ |
| fungal-type cell wall (GO:0009277) | 5/132 | $1.57 \times 10^{-4}$ | $6.79 \times 10^{-4}$ |
| chaperonin-containing T-complex (GO:0005832) | 2/12 | $9.42 \times 10^{-4}$ | $3.67 \times 10^{-3}$ |

**Table 12.**
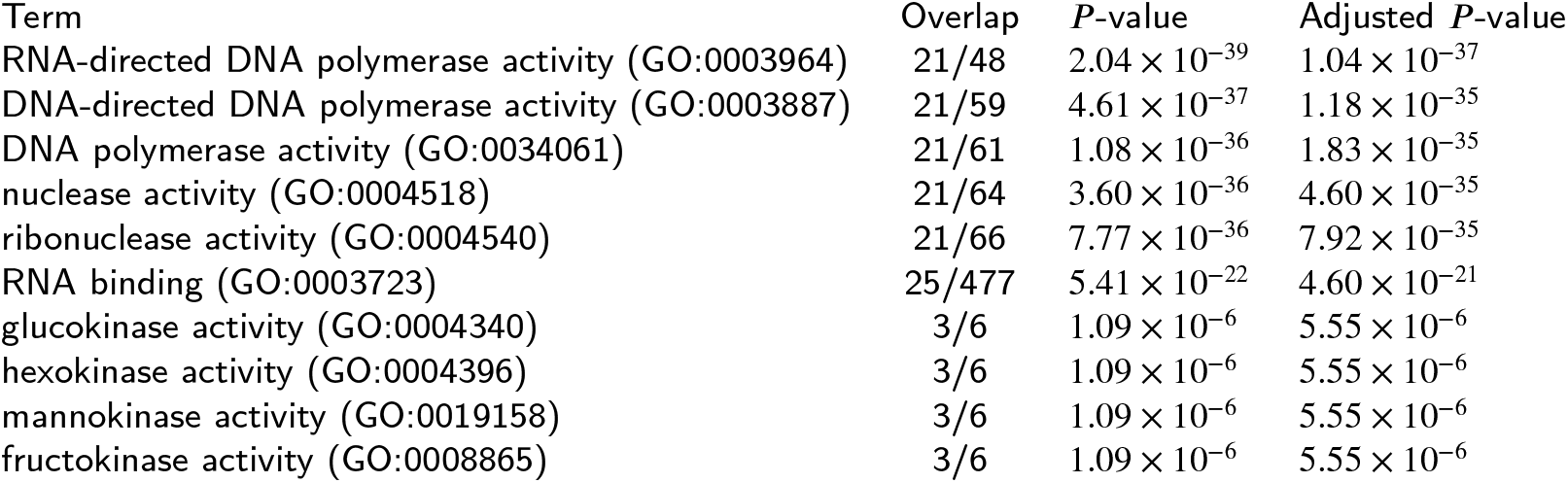
Ten top-ranked GO MF 2018 terms for 77 genes selected by TD-based unsupervised FE when single drug treatments were employed.

We confirmed the observed patterns of enrichment using g:profiler. In this analysis, we detected fewer significantly enriched terms overall but still observed enrichment for various KEGG pathways and GO terms. Thus, the biological significance of the selected genes did not depend on the database, and the analyses were robust and reliable (lists of individual biological terms obtained using YeastEnrichr and g:profiler are available as supplementary materials).

Our main contributions are summarized as follows. (1) We provide a method for the reliable interpretation of the effects of interactions between drugs on gene expression data; in particular, we propose a noevl unsupervised method involving tensor decomposition (TD)-based unsupervised feature extraction (FE) (Taguchi, 2019b) and apply this approach to datasets used in (Lukac̆is̆in and Bollenbach, 2019). (2) We demonstrate that our TD-based unsupervised FE can replicate the findings of Lukacišin and Bollenbach (Lukac̆is̆in and Bollenbach, 2019) based on a principal component analysis (PCA) (Jolliffe and Cadima, 2016). (3) Using the newly proposed TD-based unsupervised FE method, we show that convex dose-dependence can appear in single-drug treatments. Thus, our analysis improves our general understanding of DDIs in (Lukac̆is̆in and Bollenbach, 2019), especially when considering multi-drug effects. (4) Because our analysis provides detailed insights into interactions between drugs in the context of gene expression (Benet et al., 2019), it has practical implications for improving performance when designing computational methods to predict interactions between drugs accurately.

## 4. Discussion

To analyze and interpret the effects of drug interactions on gene expression, we propose a new unsupervised method, TD-based unsupervised FE in 3D, and applied it to gene expression profiles of *S. cerevisiae* treated with single or combinatorial drugs. Because strong non-linear dependence was observed for both treatments (separate and combined), our analysis demonstrates that these effects are unlikely to reflect DDIs.

One might wonder why we did not compare our methods with other methods. First of all, we compare our methods with PCA that was employed in (Lukac̆is̆in and Bollenbach, 2019). Thus, it is miss-interpretation that we did not compare ours with no other methods. Second, since our approach is an unsupervised method, to our knowledge, there are no other methods to achieve that we have had in this study; i. e., investigate what the representative gene expression profiles without assuming anything and compare the representative gene expression profiles with individual gene expression profiles. Thus, we could not find anything to be compared with our methods other than PCA.

One might also mind if the present study is novel enough to be published since I have even a whole book on this TD-based unsupervised FE. Nevertheless, TD-based unsupervised FE is too general to deny individuals because a dual application is nothing but an application of this method. TD-based unsupervised FE could be used to various topics ranging from biomarker identification, diseases causing genes identification, and drug discovery. Since Application to DDI is nothing but a new application of TD-based unsupervised FE, we do not think that the present study does not have enough novelty to be published.

The reason why we employed the yeast as a target organism is because we do not have data for other organisms. When there are data sets, we can apply our methods to these data sets.

At the moment, we do not have any limitations to our methods. We believe that we can apply our technique to any kind of gene expression profiles obtained by the combinatorial treatments of multiple drugs. We are waiting for data set which are publicly available.

Numerous researchers highlight the appearance of non-liner dose dependence. Bolger et al. (2009) proposed computational models when intestinal influx and efflux transporters are involved in gastrointestinal absorption and found non-linear dose dependence. Chen and Leung (2001) proposed the model of optically stimulated luminescence and thermoluminescence. Potashnikova et al. (2019) experimentally showed the non-linear dose response of lymphocyte cell lines to microtubule inhibitors. Lutz et al. (1978) observed a nonlinear dose-response relationship for the binding of the carcinogen benzo(a)pyrene to rat liver DNA *in vivo*. Cai et al. (2015) experimentally showed the nonlinear dose response for the protective effects of resveratrol in humans and mice. Although the number of studies is limited, non-linear dose dependence has been experimentally observed and computationally predicted even in single doses. Identification of whether non-linear dose dependence is caused by DDI is important. To identify adverse effects caused by combinatorial drug treatments, it is important to identify which genes are affected by DDI. If we would like to make use of non-linear dose dependence as an evidence of DDI, genes that exhibit non-linear dose dependence by single doses must be excluded. In this sense, TD based unsupervised FE that can deal with both single dose and combinatorial dose in an integrated manner is a very useful tool. In the feature, single as well as combinatorial doses should be analyzed by TD based unsupervised FE in an integrated manner prior to the identification of genes affected by DDI.

## 5. Conclusion

In this study, we show that convex dose dependence previously regarded as an evidence of DDI can be found in single dose treatment using integrated analysis using TD based unsupervised FE. To identify DDI based on the appearance of convex dose dependence, how single dose causes non-linear dose dependence is important because genes that exhibit non-linear dose dependence upon single dose must be excluded from the candidate genes affected by DDI. Thus, TD based unsupervised FE is an important tool to identify DDI based on the appearance of convex dose dependence of gene expression.

## Acknowledgment

The study was supported by KAKENHI, 19H05270, 20H04848, and 20K12067. This project was also funded by the Deanship of Scientific Research (DSR) at King Abdulaziz University, Jeddah, under grant no. KEP-8-611-38. The authors thank DSR for technical and financial support.

---

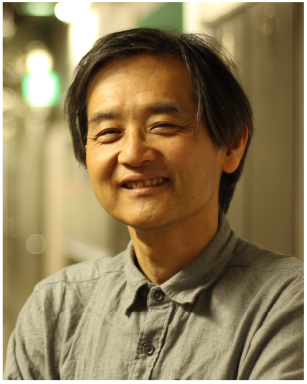

YH. Taguchi received a B.S. degree in physics from the Tokyo Institute of Technology and a Ph.D. degree in physics from the Tokyo Institute of Technology. He is currently a full professor with the Department of Physics, Chuo University, Japan. His works have been published in leading journals such as Physical Review Letters, Bioinformatics, and Scientific Reports. His research interests include bioinformatics, machine-learning, and non-linear physics. He is also an editorial board member of Frontiers in Genetics:RNA, PloS ONE, BMC Medical Genomics, Medicine (Lippincott Williams & Wilkins journal), BMC Research Notes, non-coding RNA (MDPI), and IPSJ Transaction on Bioinformatics.

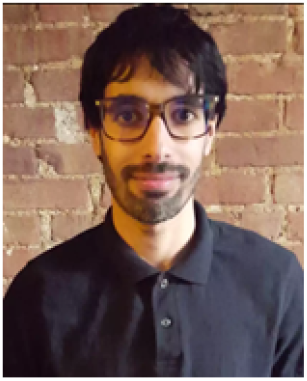

TURKI TURKI received a B.S. in computer science from King Abdulaziz University, an M.S. in computer science from NYU.POLY, and a Ph.D. in computer science from the New Jersey Institute of Technology. He is currently an assistant professor with the Department of Computer Science, King Abdulaziz University, Saudi Arabia. His research interests include artificial intelligence, machine learning, deep learning, data mining, data science, big data analytics, and bioinformatics. His research has been accepted and published in journals such as Frontiers in Genetics, BMC Genomics, BMC Systems Biology, Expert Systems with Applications, Computers in Biology and Medicine, and Current Pharmaceutical Design. He was awarded several distinction awards from the Deanship of Scientific Research at King Abdulaziz University. He is supported by King Abdulaziz University and is currently working on several biomedicine related projects. Dr. Turki has served on the program committees of several international conferences. Additionally, he is an editorial board member of Sustainable Computing: Informatics and Systems and Computers in Biology and Medicine.

