## Supplementary material for "Novel method for the prediction of drug-drug Interaction based on Gene Expression profiles": gene_exp_combinatorial.pdf

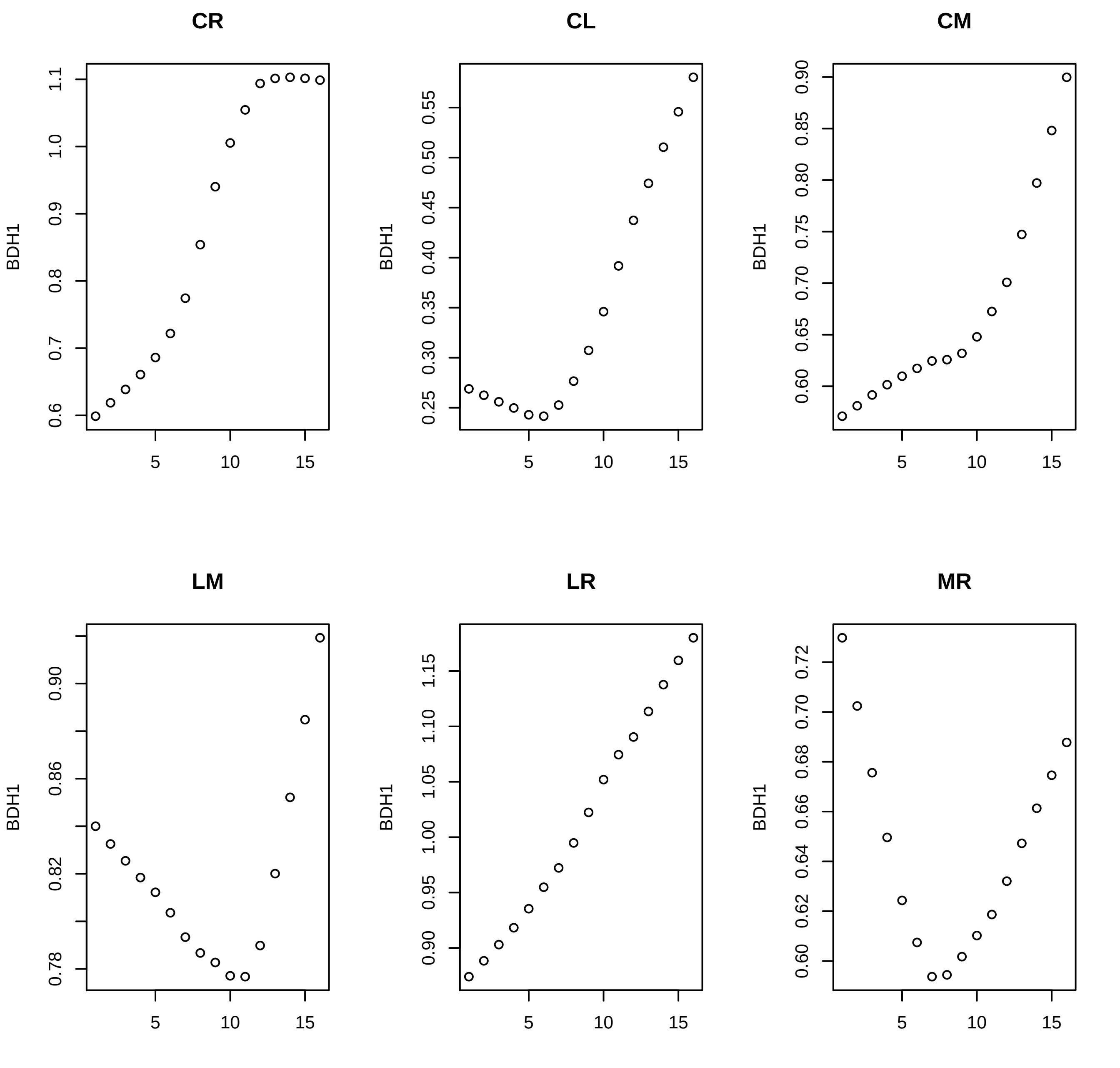
CR
CL
CM
LM
LR
MR
BDH1
5
10
15

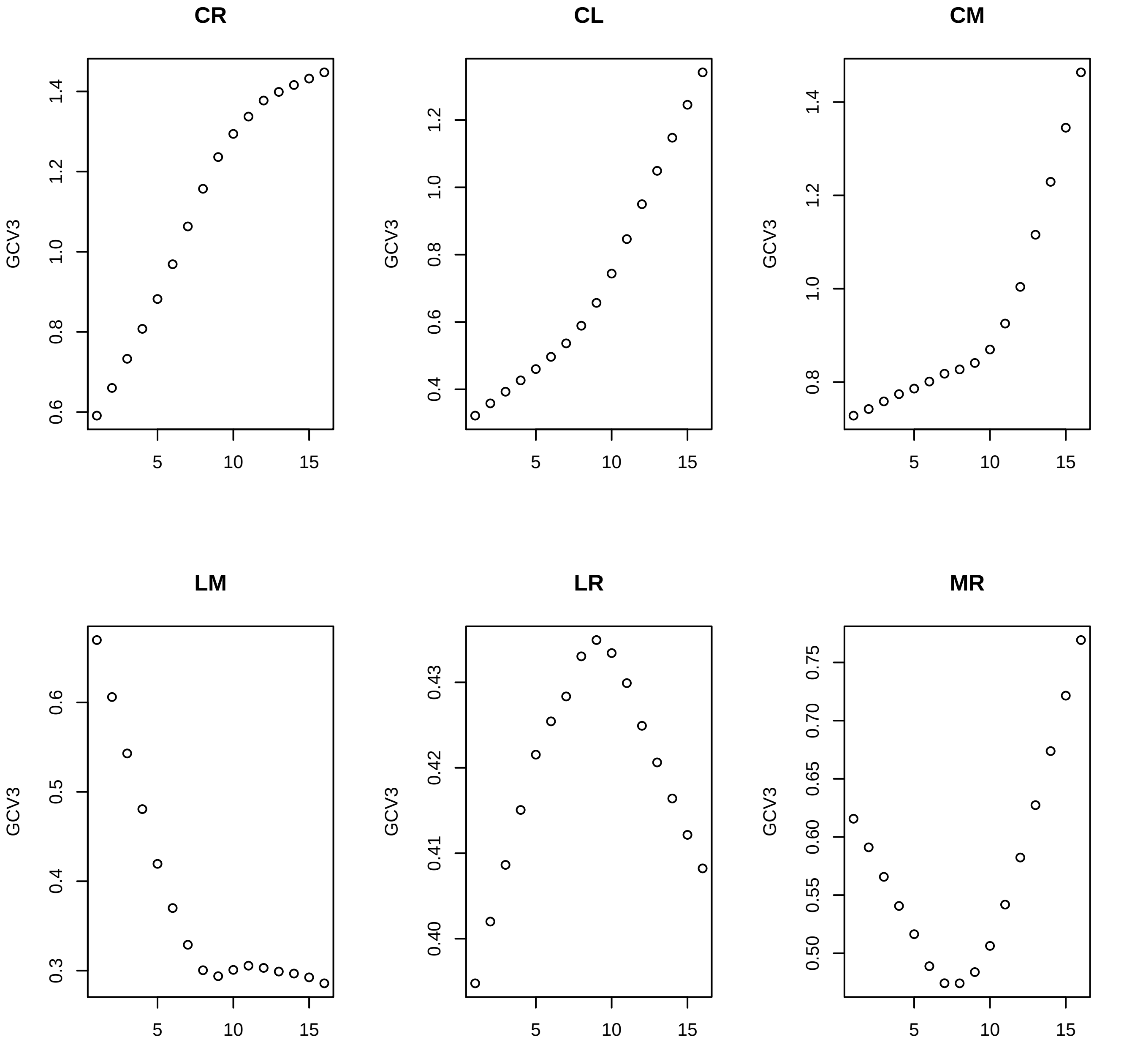

CR
CL
CM
LM
LR
MR
GCV3
5
10
15

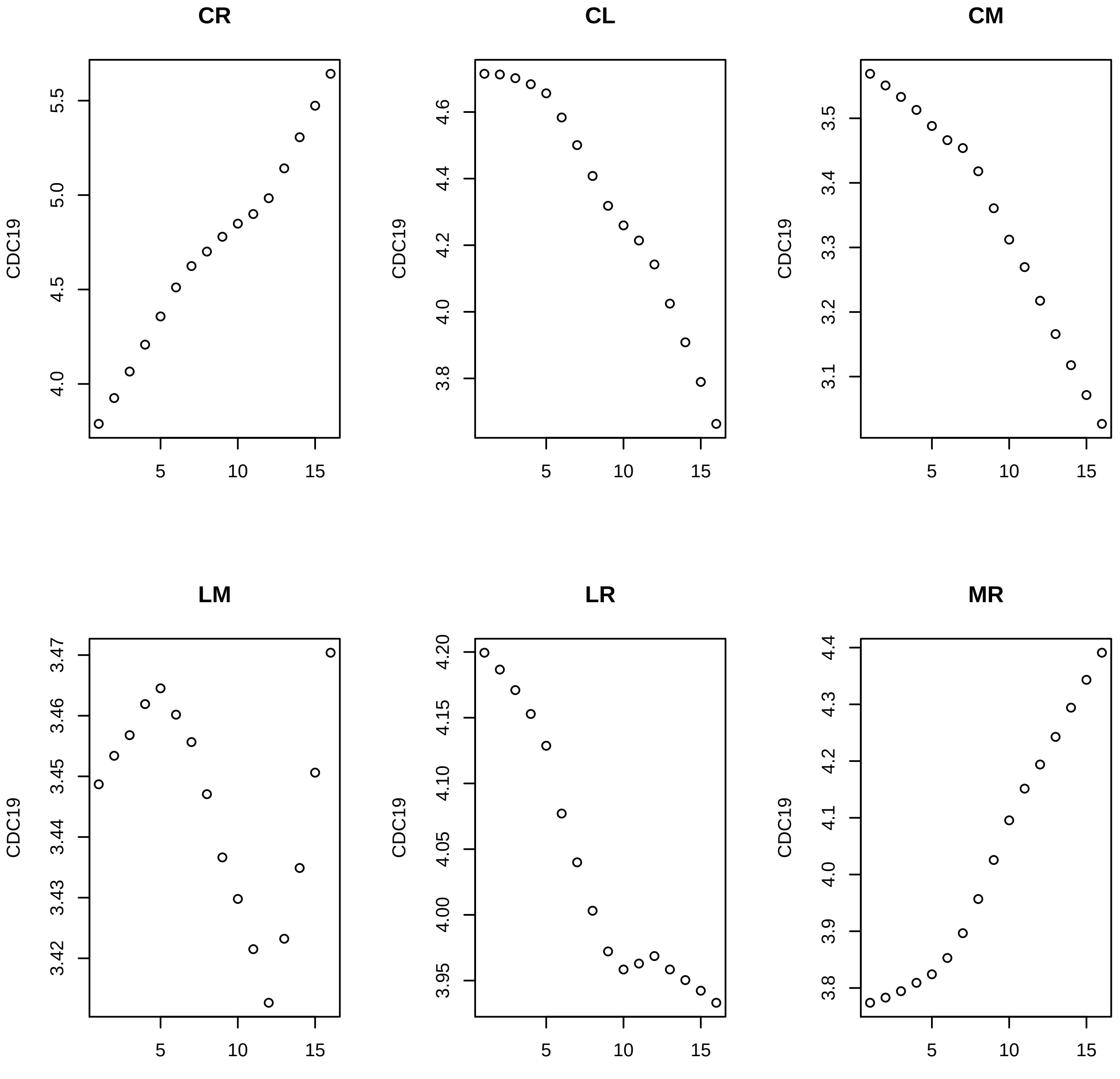
CR
CL
CM
LM
LR
MR
CDC19
5
10
15
4.0
4.5
5.0
5.5
3.8
4.0
4.2
4.4
4.6
3.1
3.2
3.3
3.4
3.5
3.42
3.43
3.44
3.45
3.46
3.47
3.95
4.00
4.05
4.10
4.15
4.20
3.8
3.9
4.0
4.1
4.2
4.3
4.4

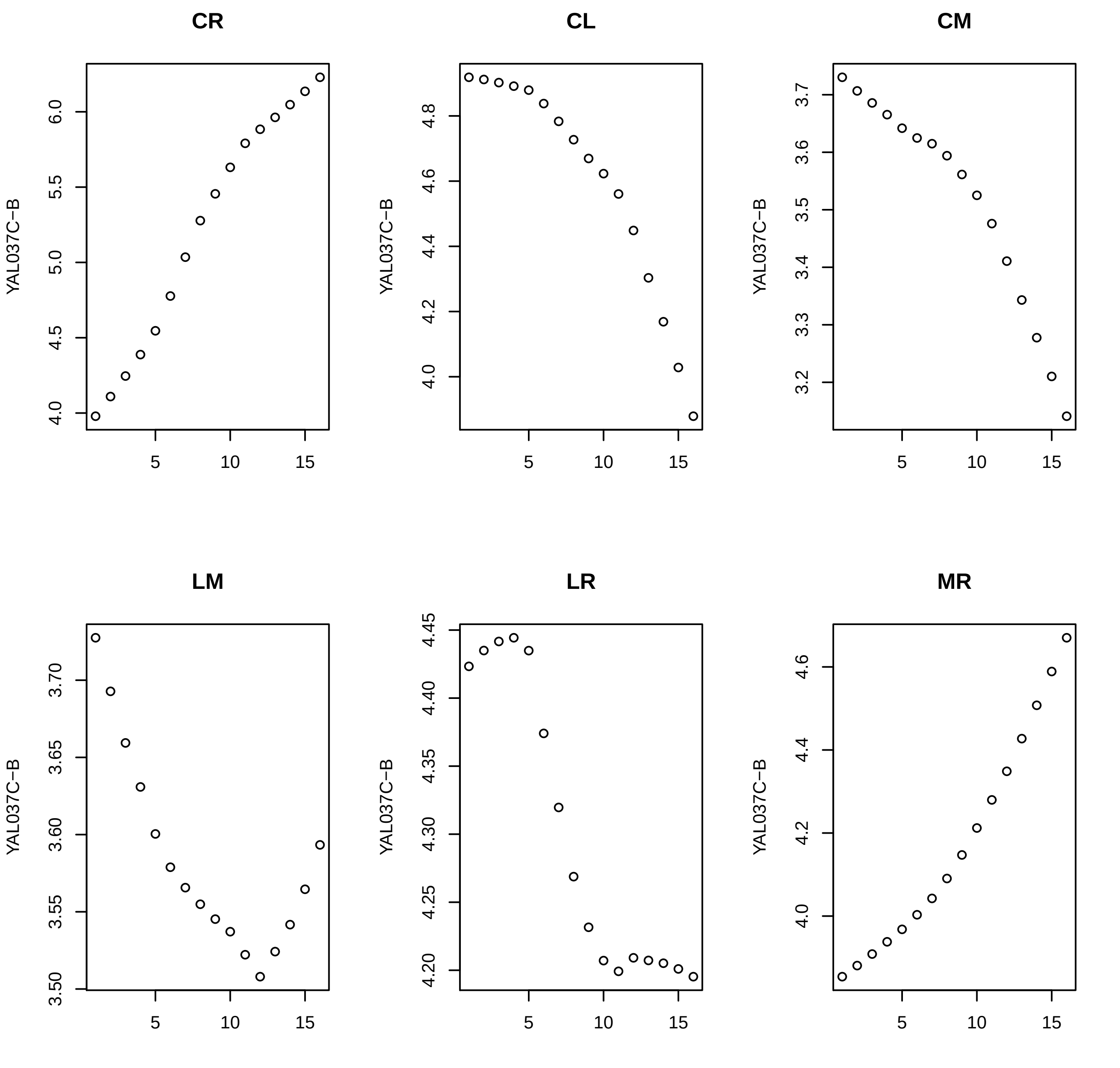
CR
CL
CM
LM
LR
MR
YAL037C−B
5
10
15

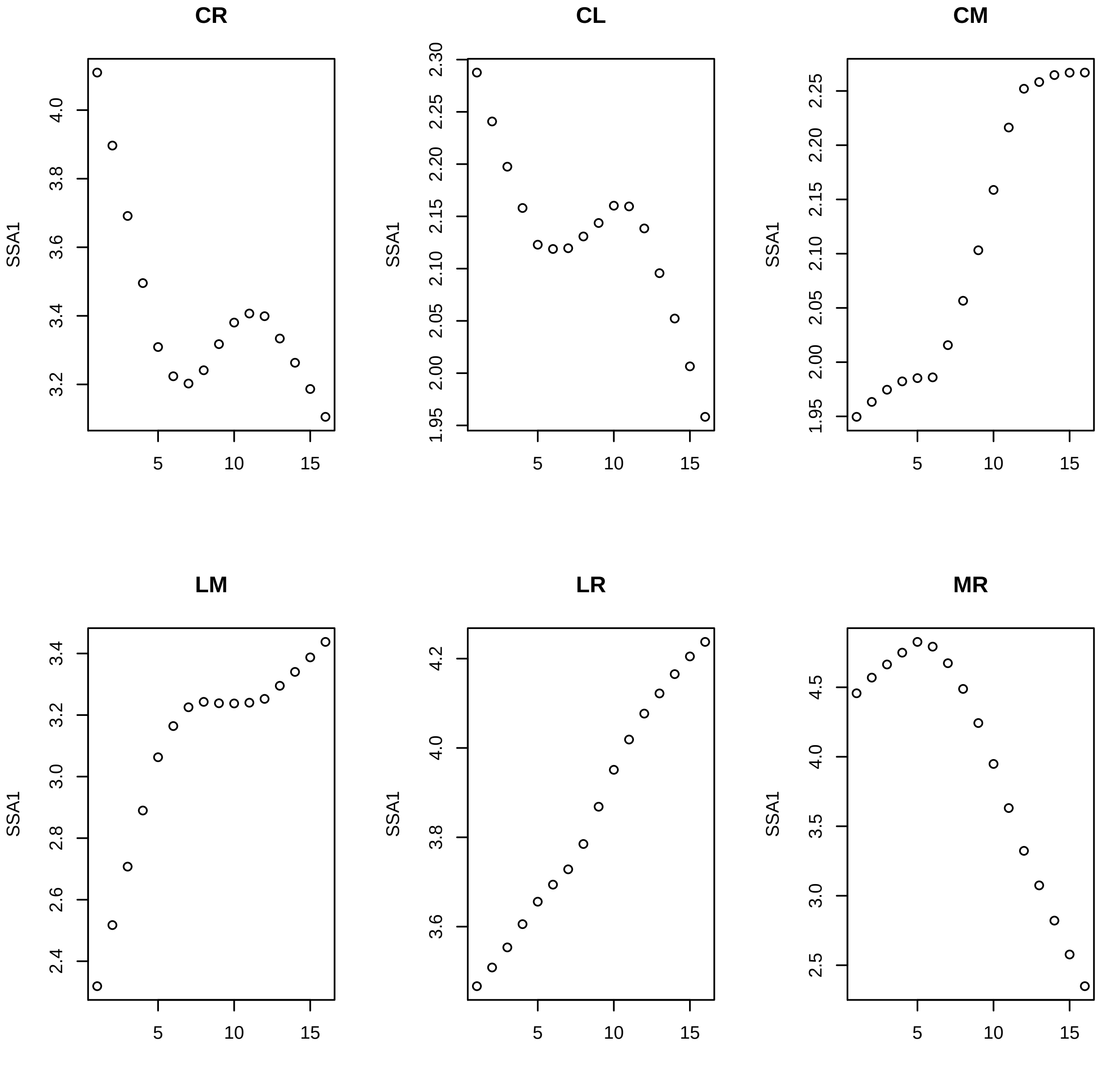
CR
CL
CM
LM
LR
MR
SSA1
3.2
3.4
3.6
3.8
4.0
1.95
2.00
2.05
2.10
2.15
2.20
2.25
2.30
2.4
2.6
2.8
3.0
4.2
2.5
3.5
4.5
5
10
15

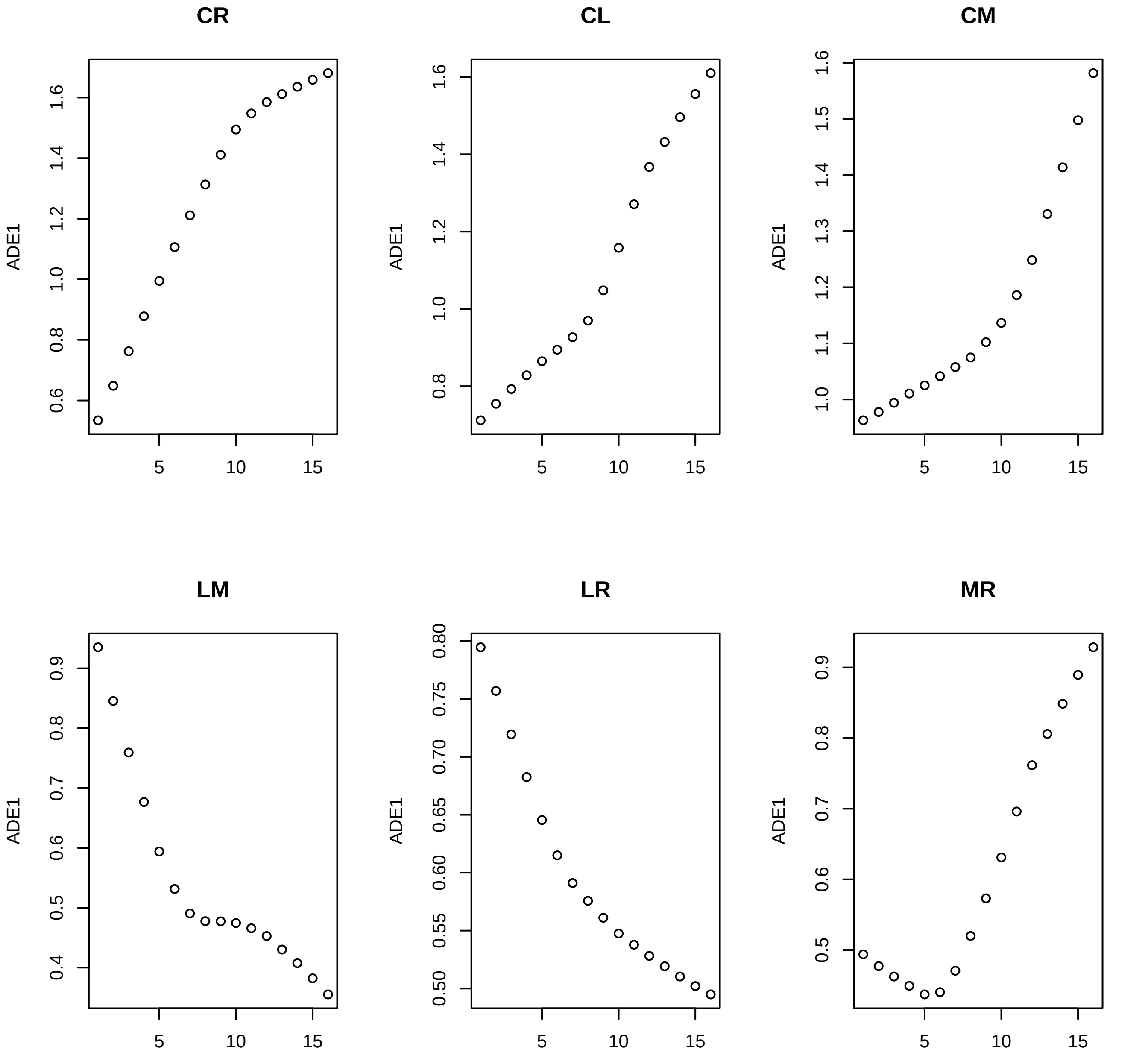
CR
ADE1
0.6
0.8
1.0
1.2
1.4
1.6
5
10
15
CL
ADE1
0.8
1.0
1.2
1.4
1.6
5
10
15
CM
ADE1
1.0
1.1
1.2
1.3
1.4
1.5
1.6
5
10
15
LM
ADE1
0.4
0.5
0.6
0.7
0.8
0.9
5
10
15
LR
ADE1
0.50
0.55
0.60
0.65
0.70
0.75
0.80
5
10
15
MR
ADE1
0.5
0.6
0.7
0.8
0.9
5
10
15

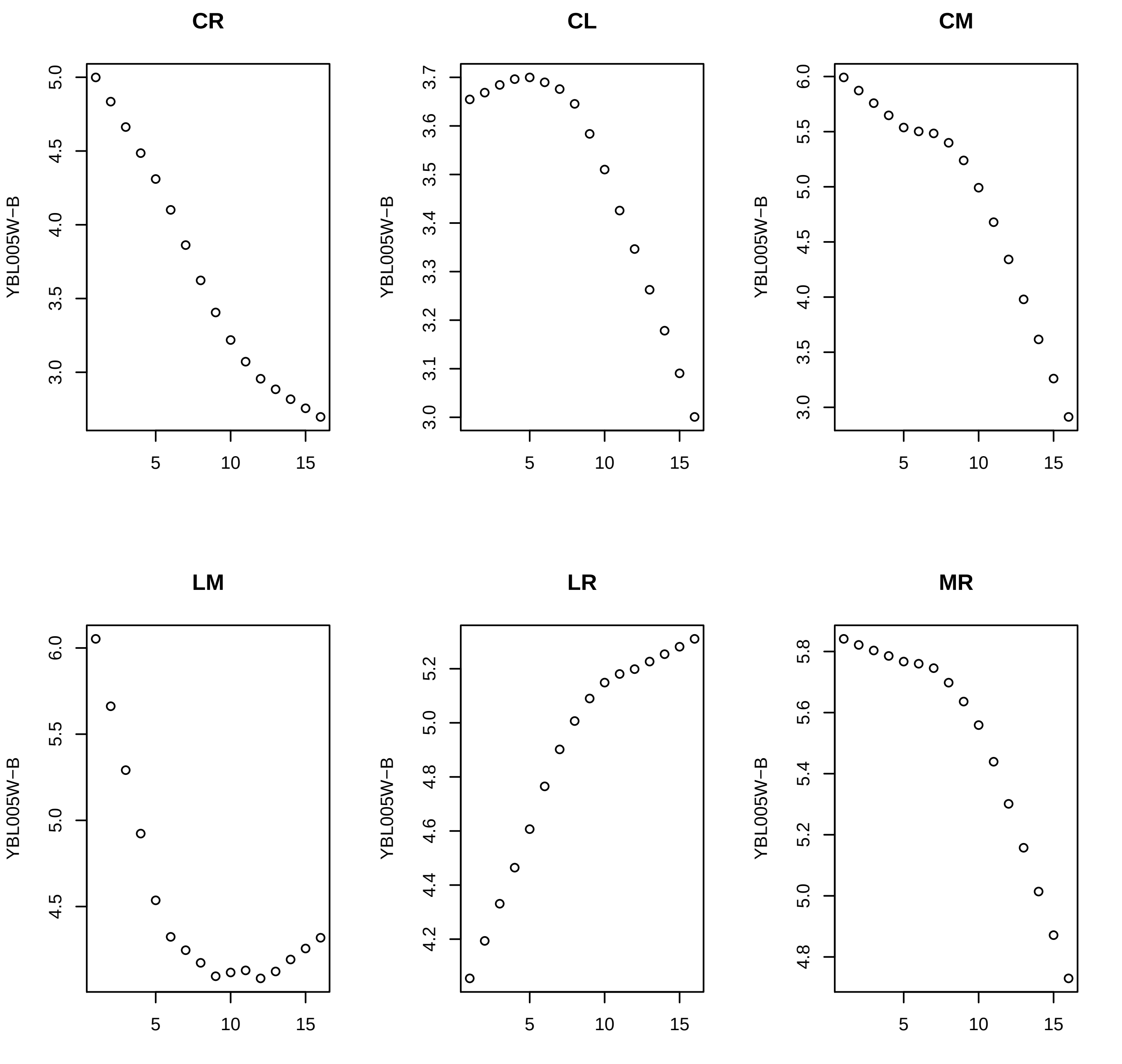

CR
CL
CM
LM
LR
MR
YBL005W−B

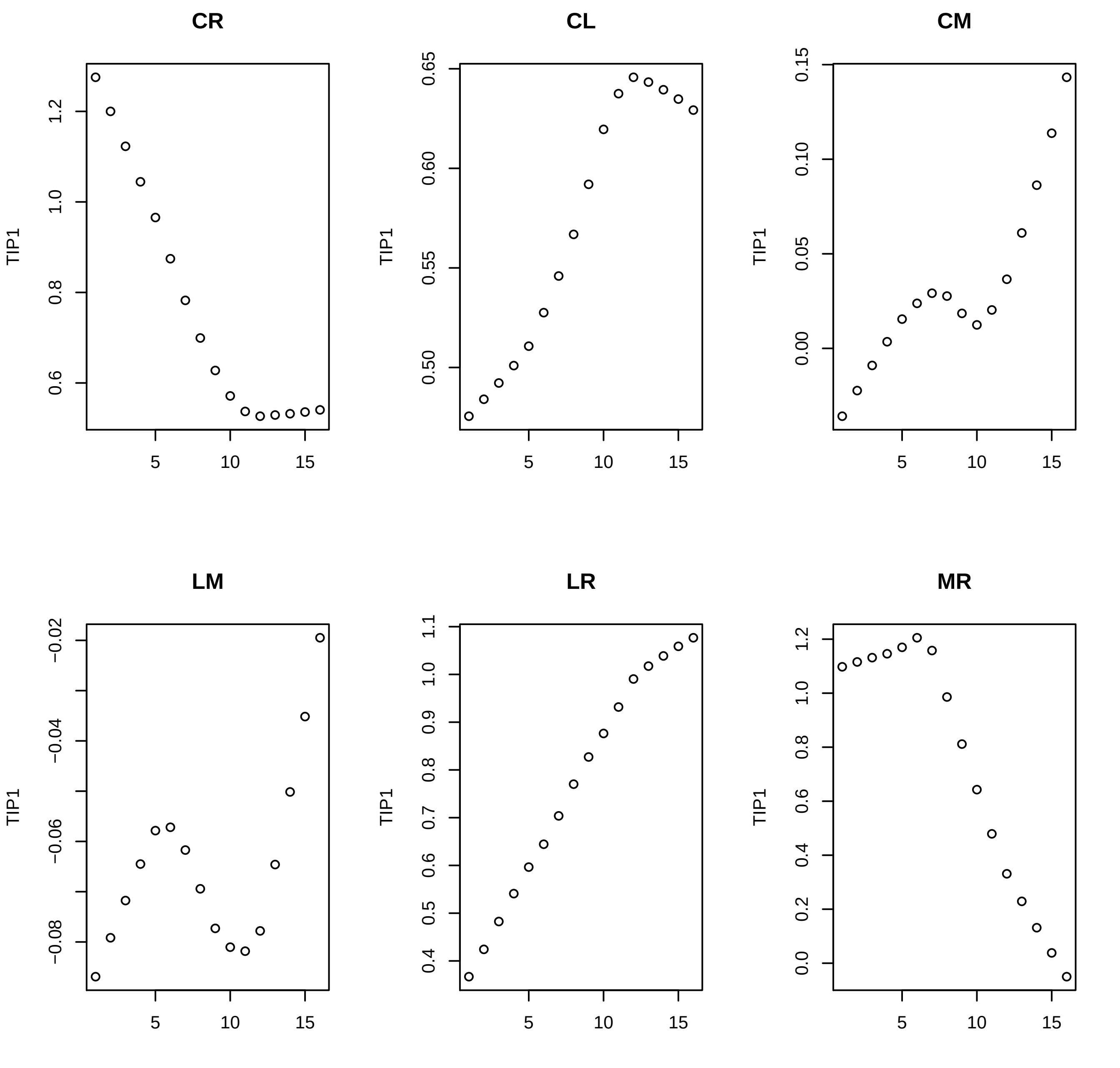
CR
CL
CM
LM
LR
MR
TIP1
5
10
15
1.2
1.0
0.8
0.6
0.65
0.60
0.55
0.50
0.15
0.10
0.05
0.00
−0.02
−0.04
−0.06
−0.08
1.1
1.0
0.9
0.8
0.7
0.6
0.5
0.4
1.2
1.0
0.8
0.6
0.4
0.2
0.0

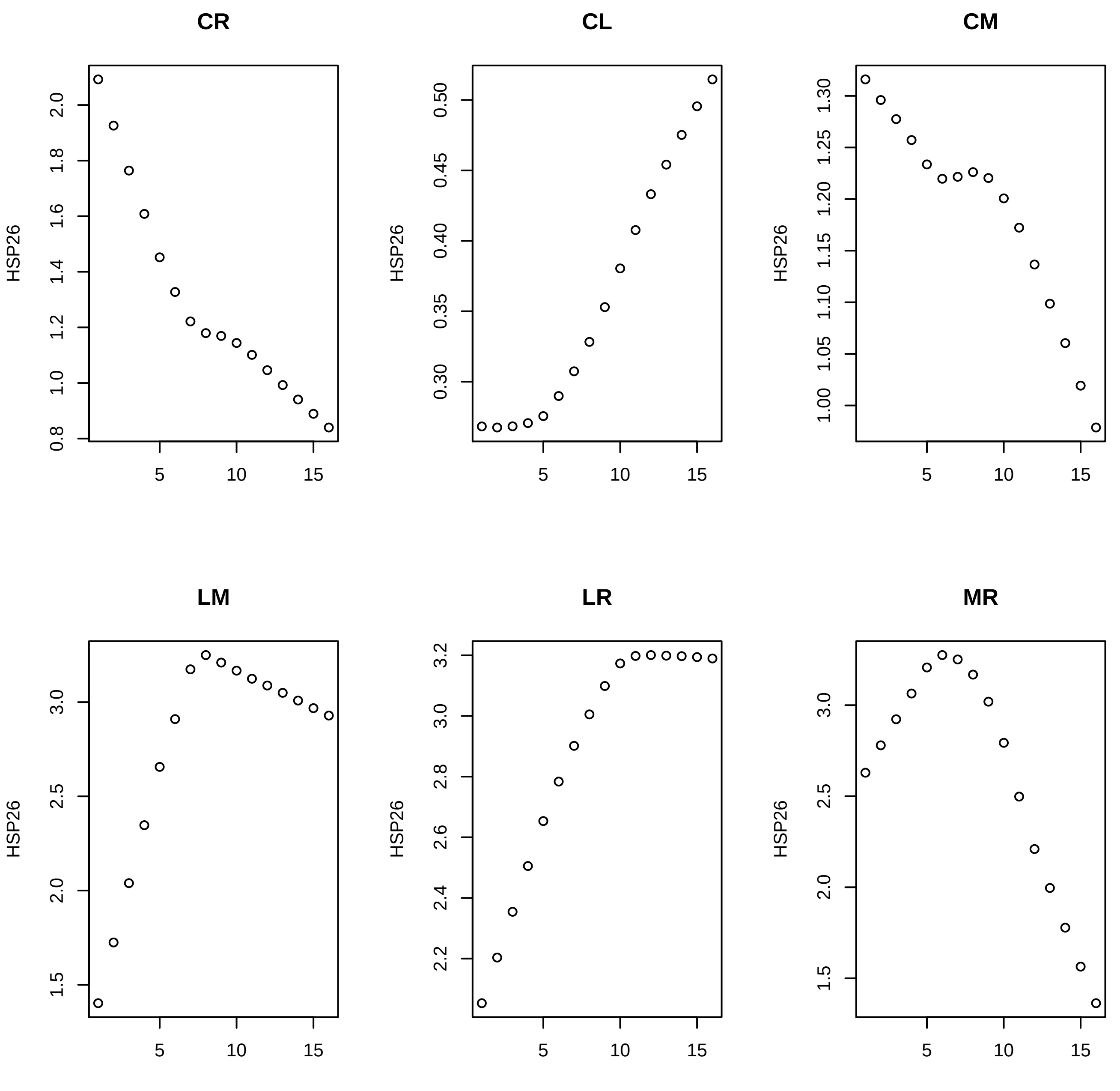
CR
CL
CM
LM
LR
MR
HSP26
0.8
1.0
1.2
1.4
1.6
1.8
2.0
0.30
0.35
0.40
0.45
0.50
1.00
1.05
1.10
1.15
1.20
1.25
1.30
1.5
2.0
2.5
3.0
2.2
2.4
2.6
2.8
3.2
5
10
15

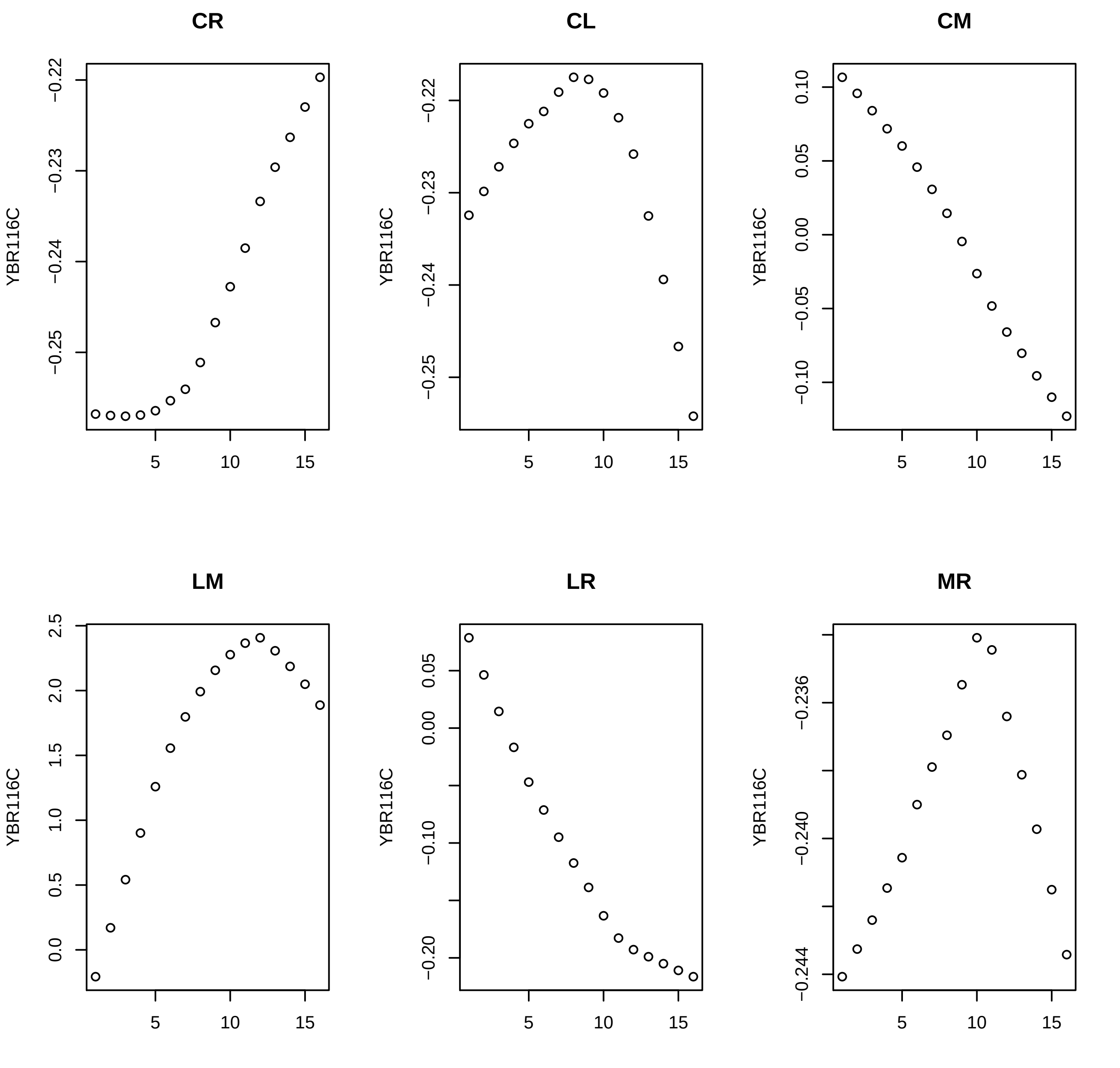
CR
CL
CM
LM
LR
MR
YBR116C
−0.22
−0.23
−0.24
−0.25
0.10
0.05
0.00
−0.05
−0.10
2.5
2.0
1.5
1.0
0.5
0.0
−0.20
−0.236
−0.240
−0.244
5
10
15

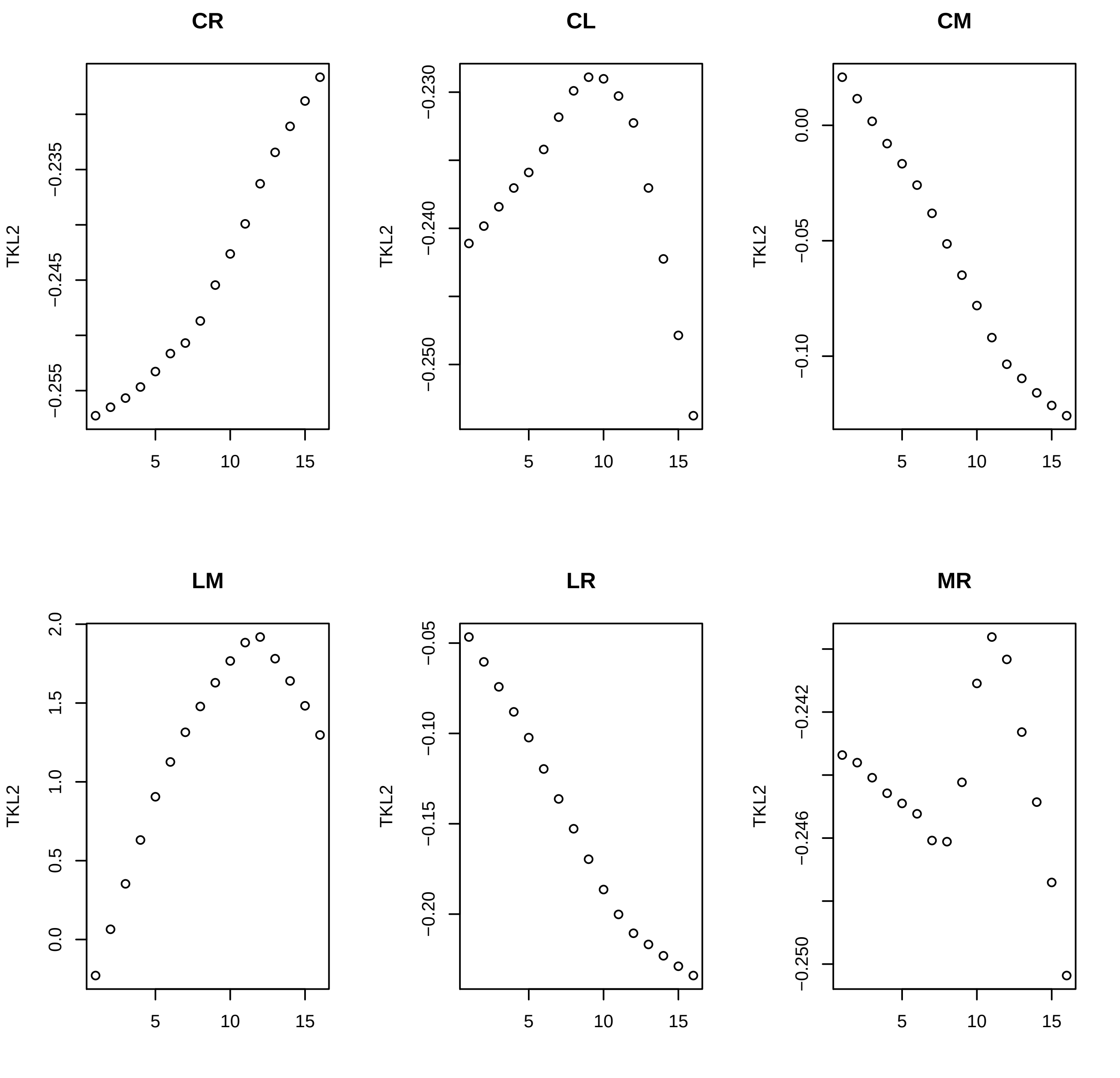

CR
CL
CM
LM
LR
MR
TKL2
−0.255
−0.245
−0.235
−0.250
−0.240
−0.230
−0.10
−0.05
0.00
0.0
0.5
1.0
1.5
2.0
−0.20
−0.15
−0.10
−0.05
−0.250
−0.246
−0.242
5
10
15

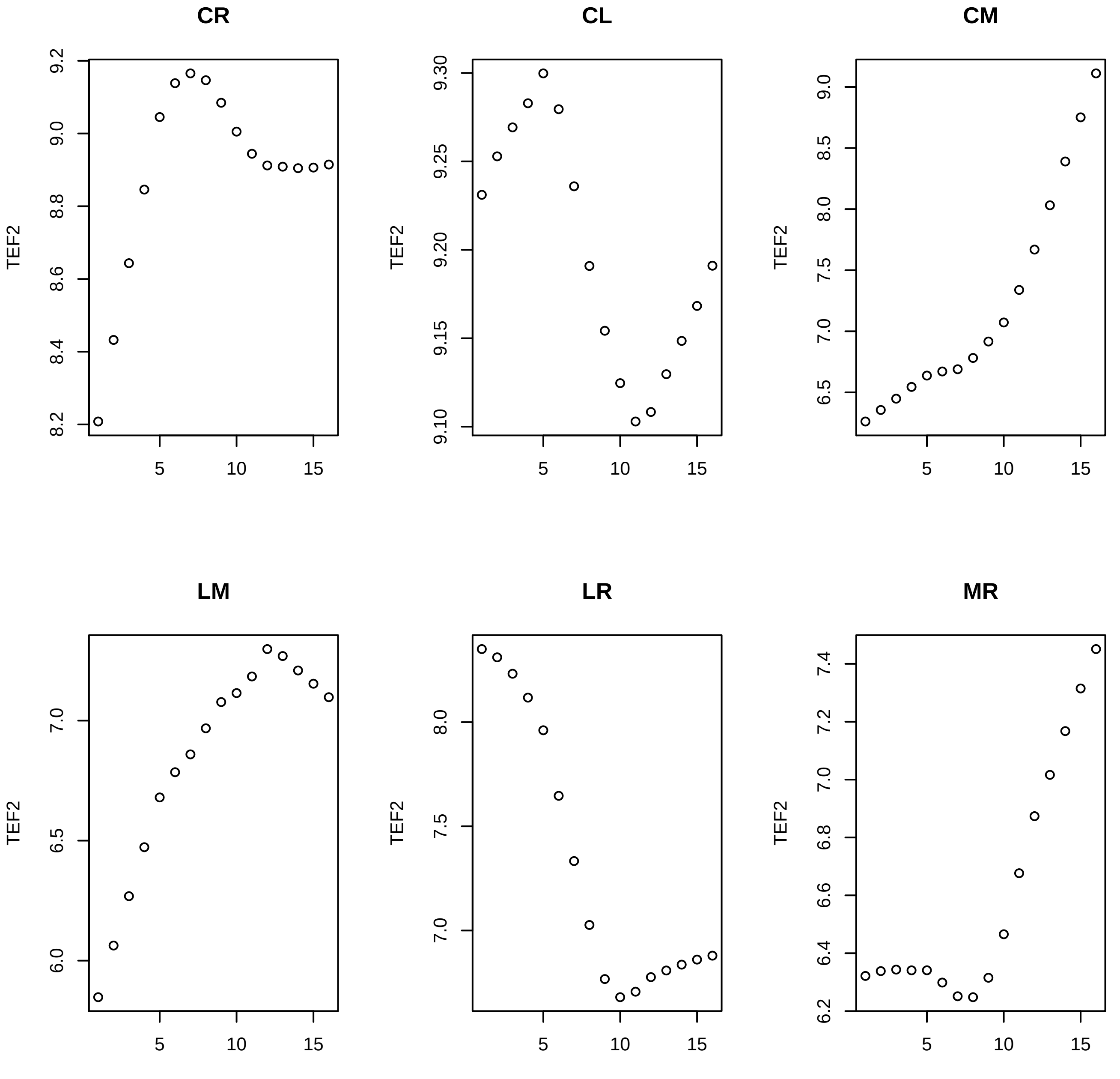
CR
CL
CM
LM
LR
MR
TEF2
5
10
15

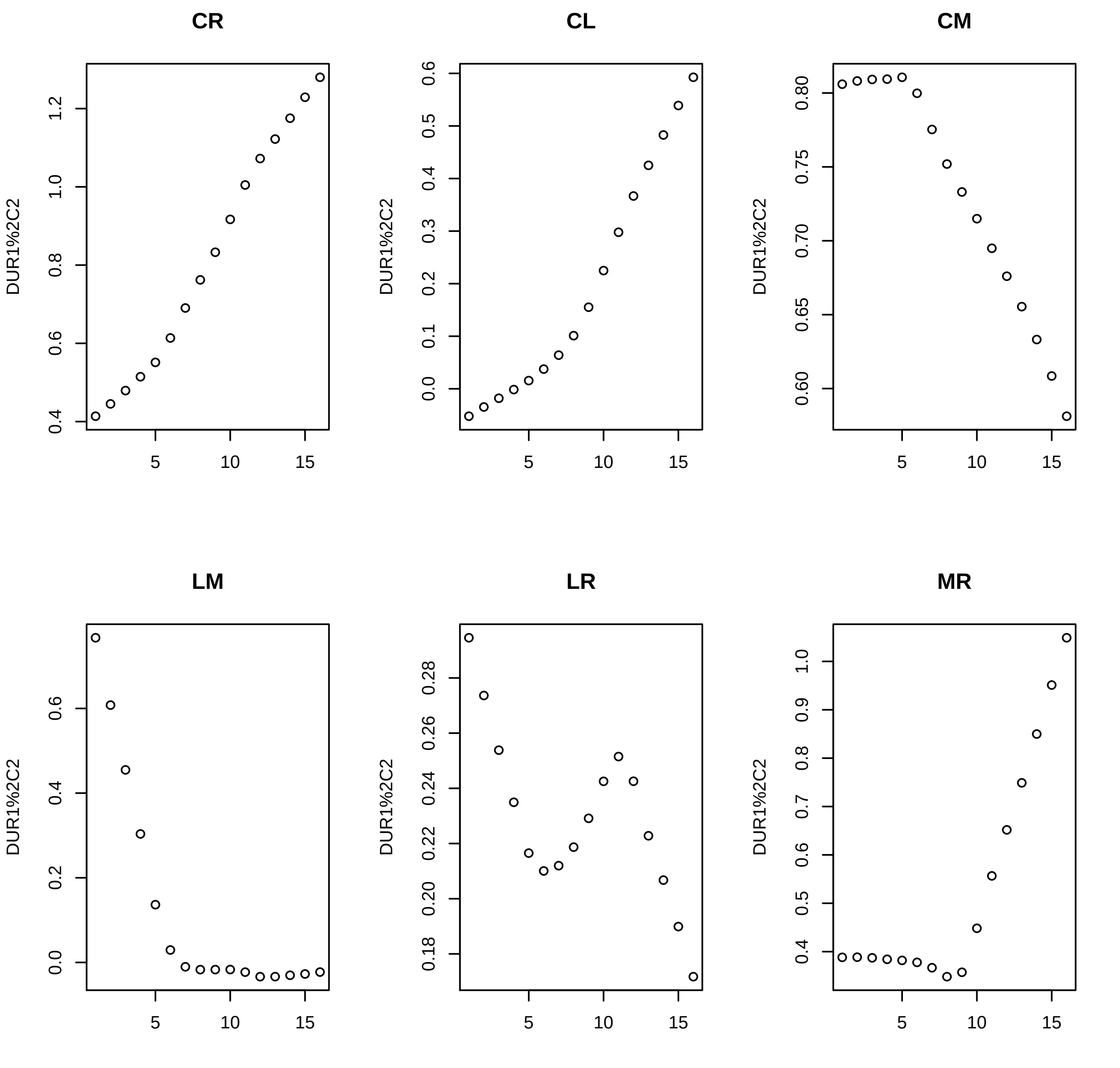

CR
CL
CM
LM
LR
MR
DUR1%2C2

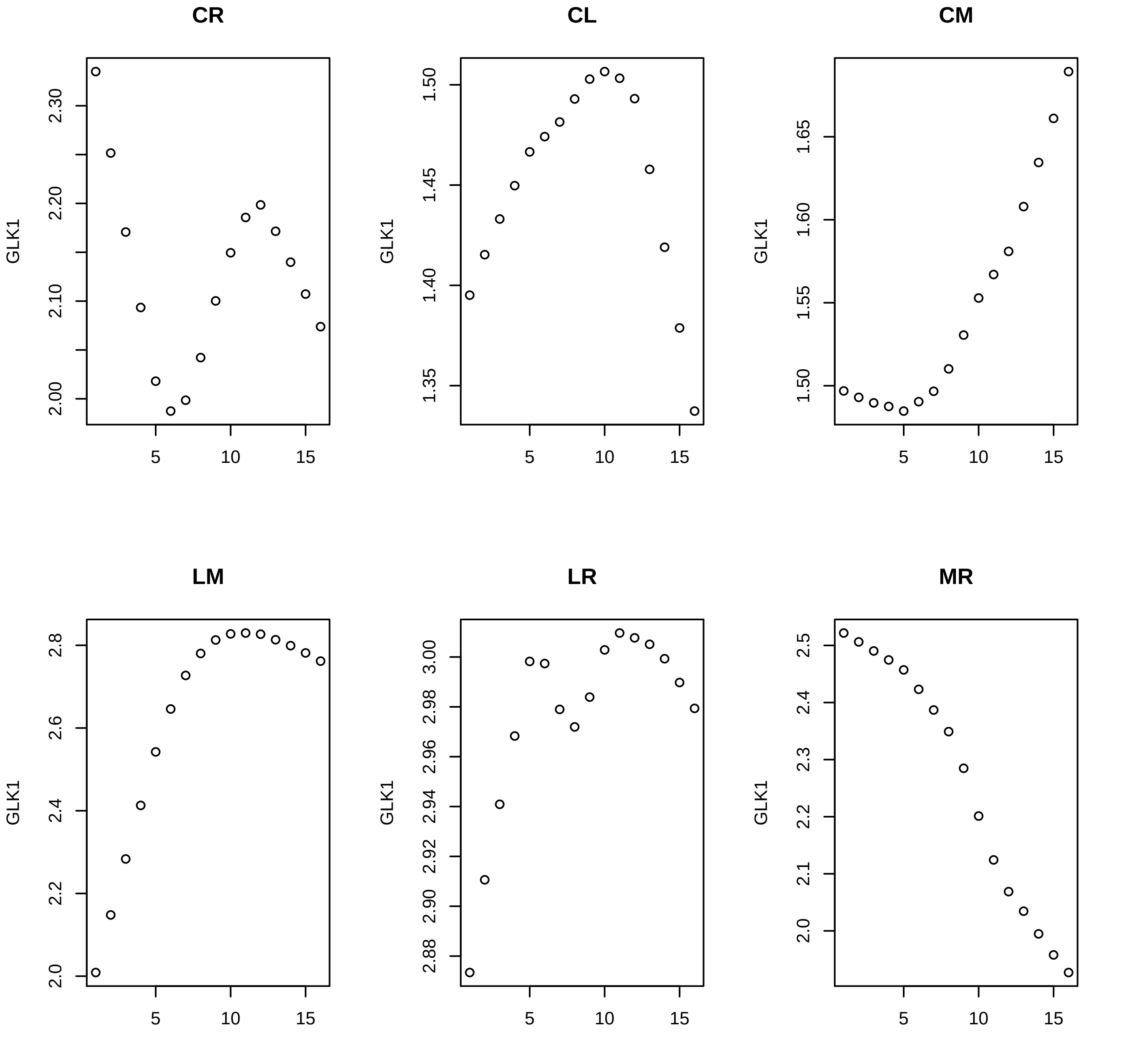

CR
CL
CM
LM
LR
MR
GLK1

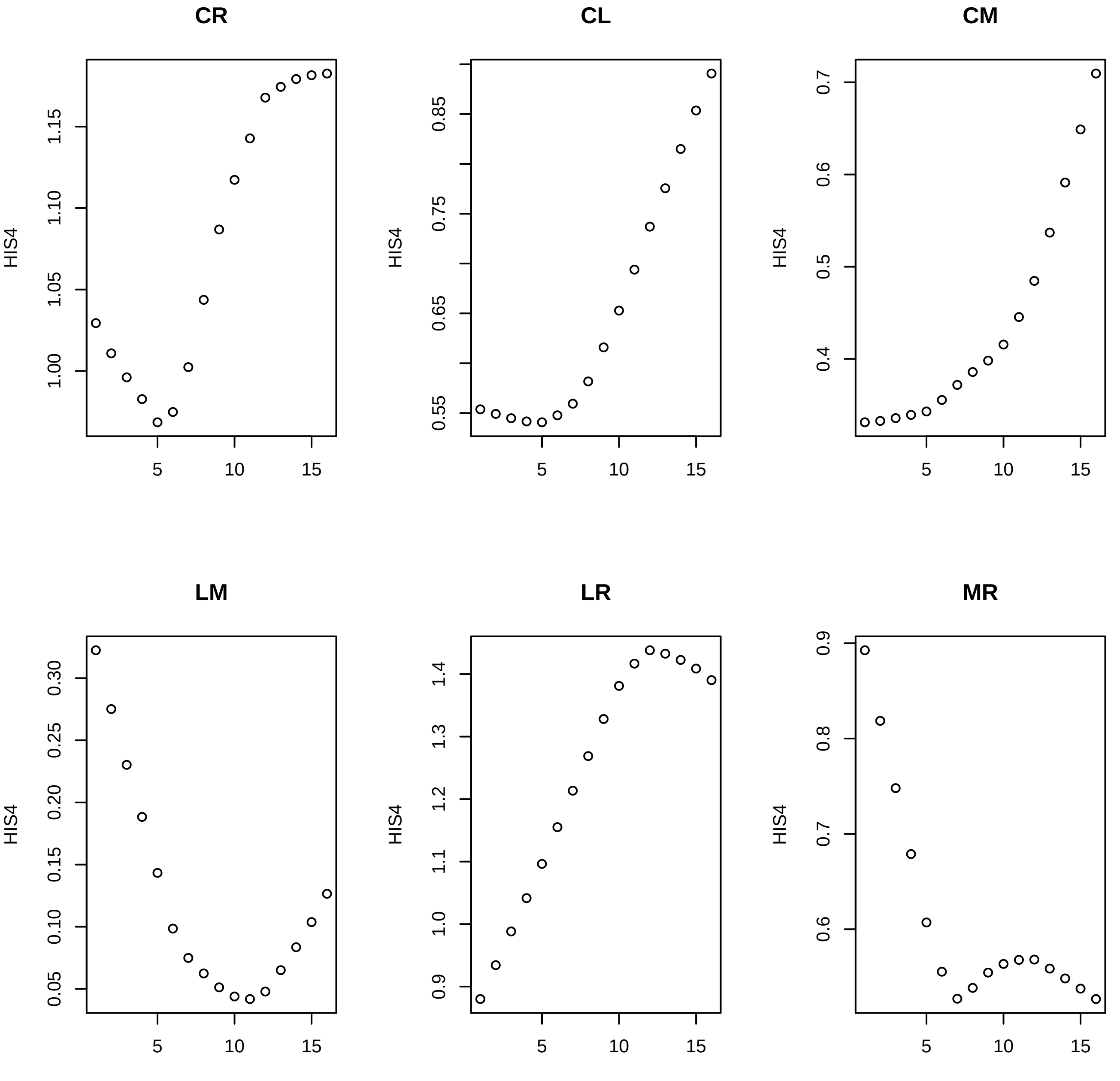

CR
CL
CM
LM
LR
MR
HIS4

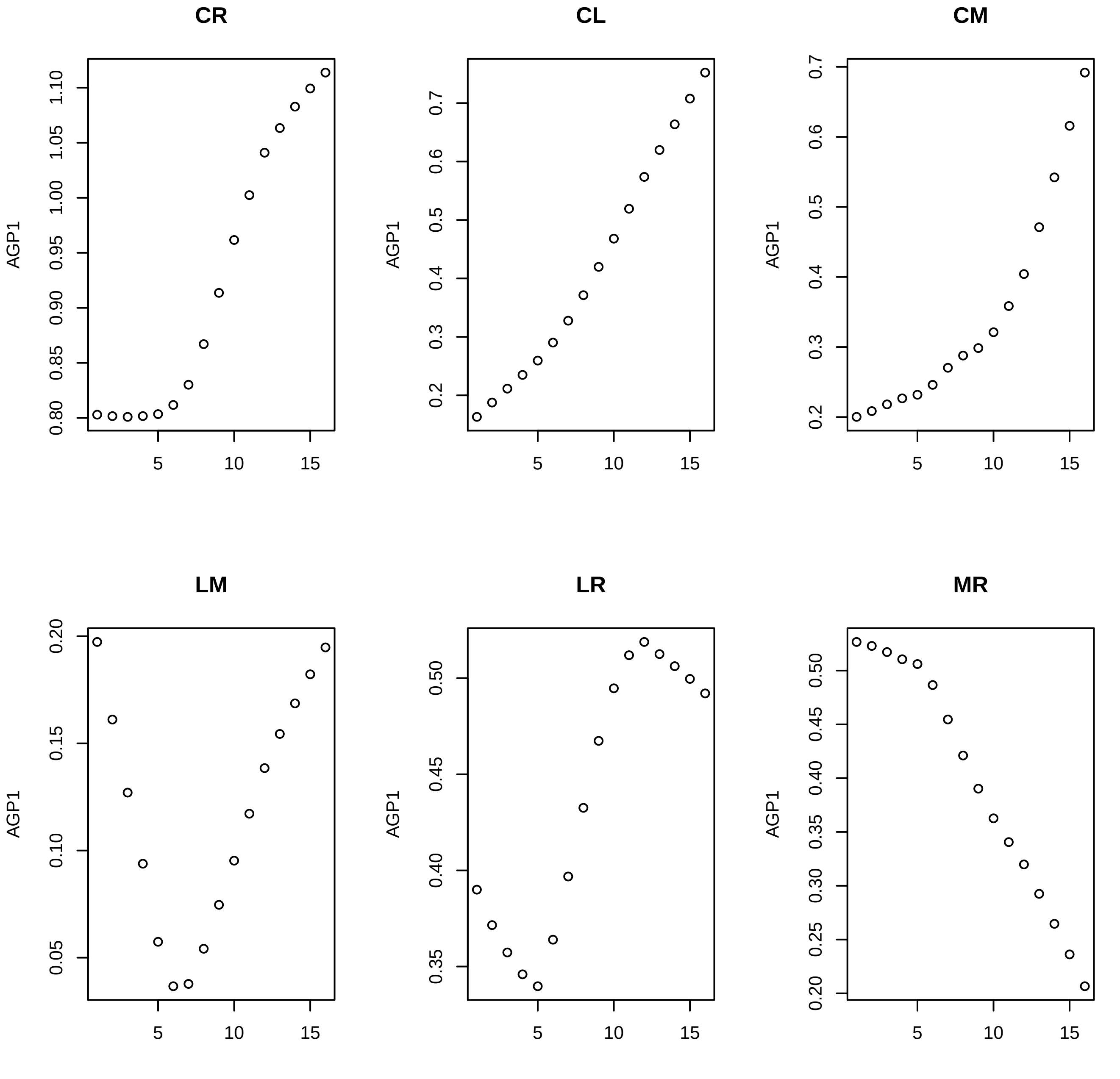
CR
AGP1
0.80
0.85
0.90
0.95
1.00
1.05
1.10
5
10
15
CL
AGP1
0.2
0.3
0.4
0.5
0.6
0.7
5
10
15
CM
AGP1
0.2
0.3
0.4
0.5
0.6
0.7
5
10
15
LM
AGP1
0.05
0.10
0.15
0.20
5
10
15
LR
AGP1
0.35
0.40
0.45
0.50
5
10
15
MR
AGP1
0.20
0.25
0.30
0.35
0.40
0.45
0.50
5
10
15

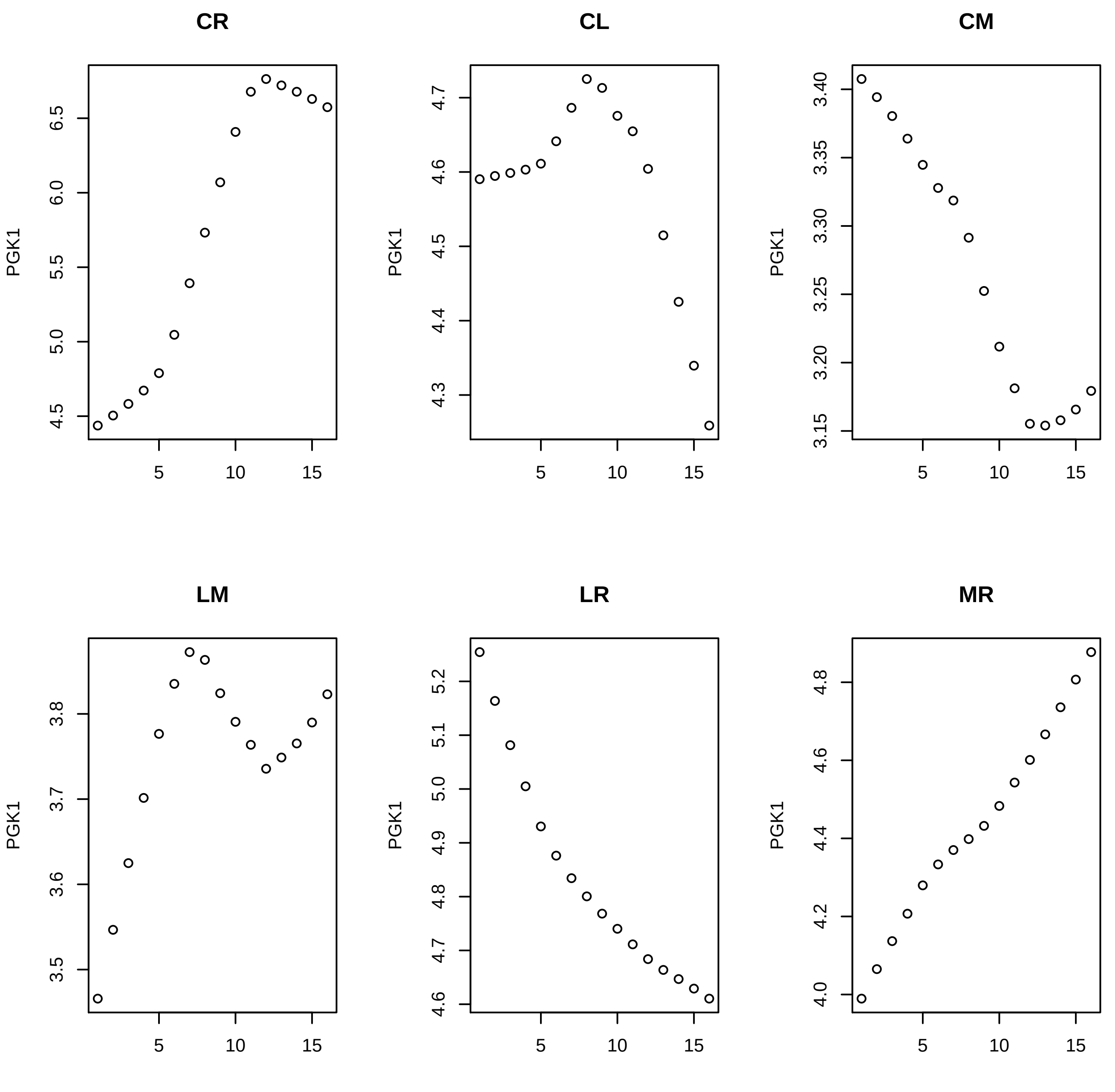
CR
CL
CM
LM
LR
MR
PGK1
6.5
6.0
5.5
5.0
4.5
4.7
4.6
4.5
4.4
4.3
3.40
3.35
3.30
3.25
3.20
3.15
3.8
3.7
3.6
3.5
5.2
5.1
5.0
4.9
4.8
4.7
4.6
4.8
4.6
4.4
4.2
4.0
5
10
15

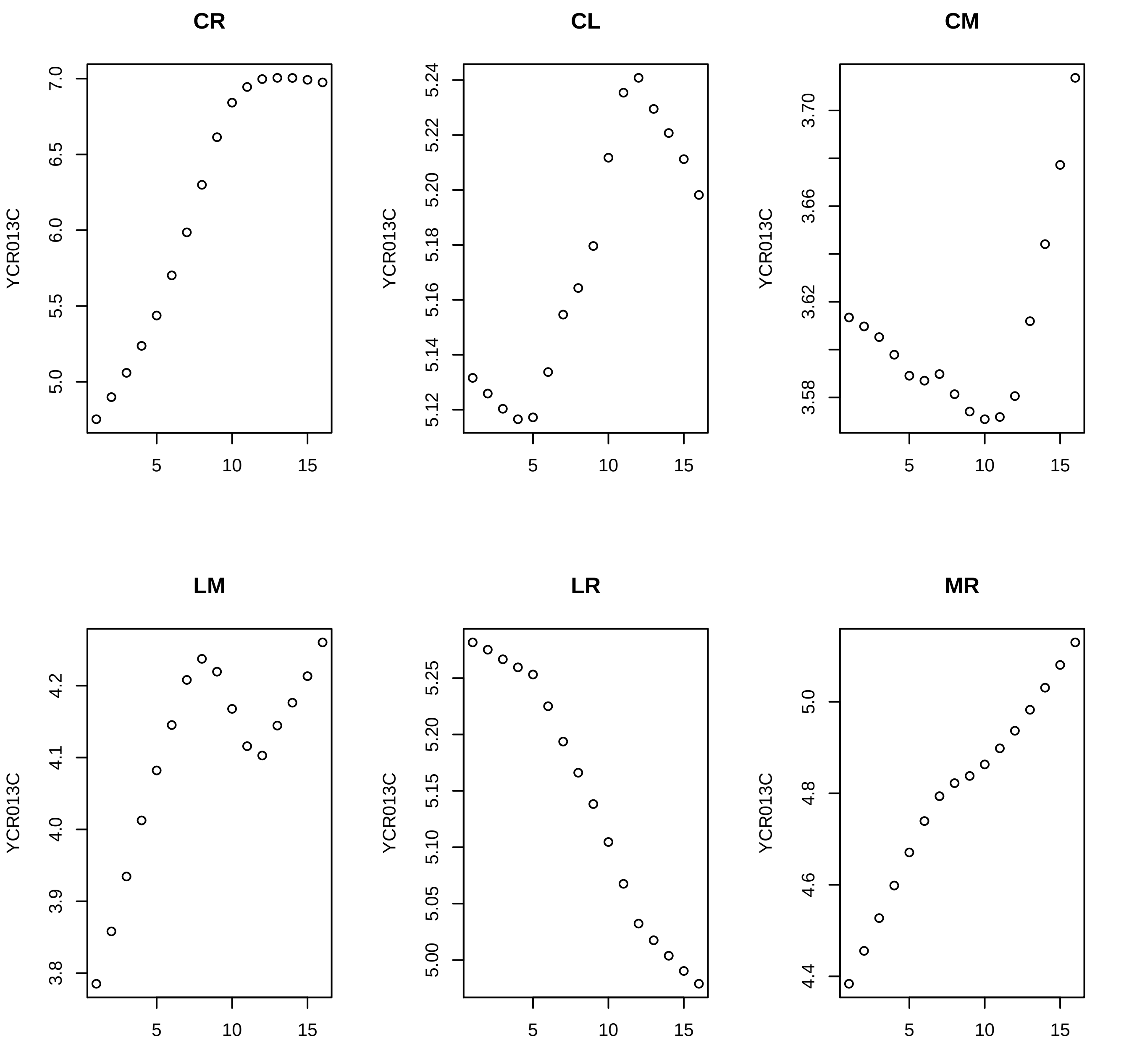

CR
CL
CM
LM
LR
MR
YCR013C

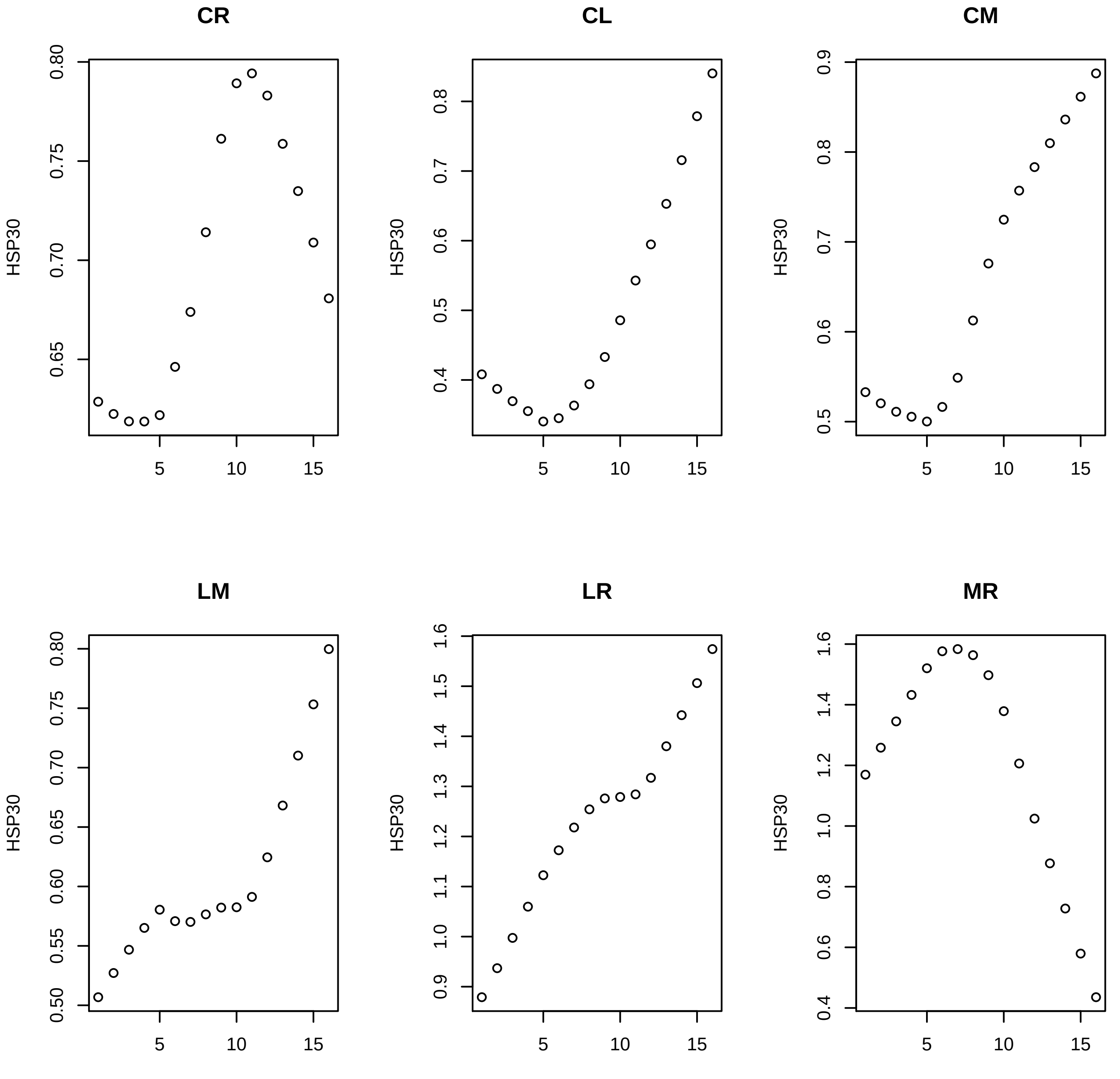
CR
CL
CM
LM
LR
MR
HSP30
0.65
0.70
0.75
0.80
0.4
0.5
0.6
0.7
0.8
0.9
0.50
0.55
0.60
0.9
1.0
1.1
1.2
1.3
1.4
1.5
1.6
0.4
0.6
0.8
1.0
1.2
1.4
5
10
15

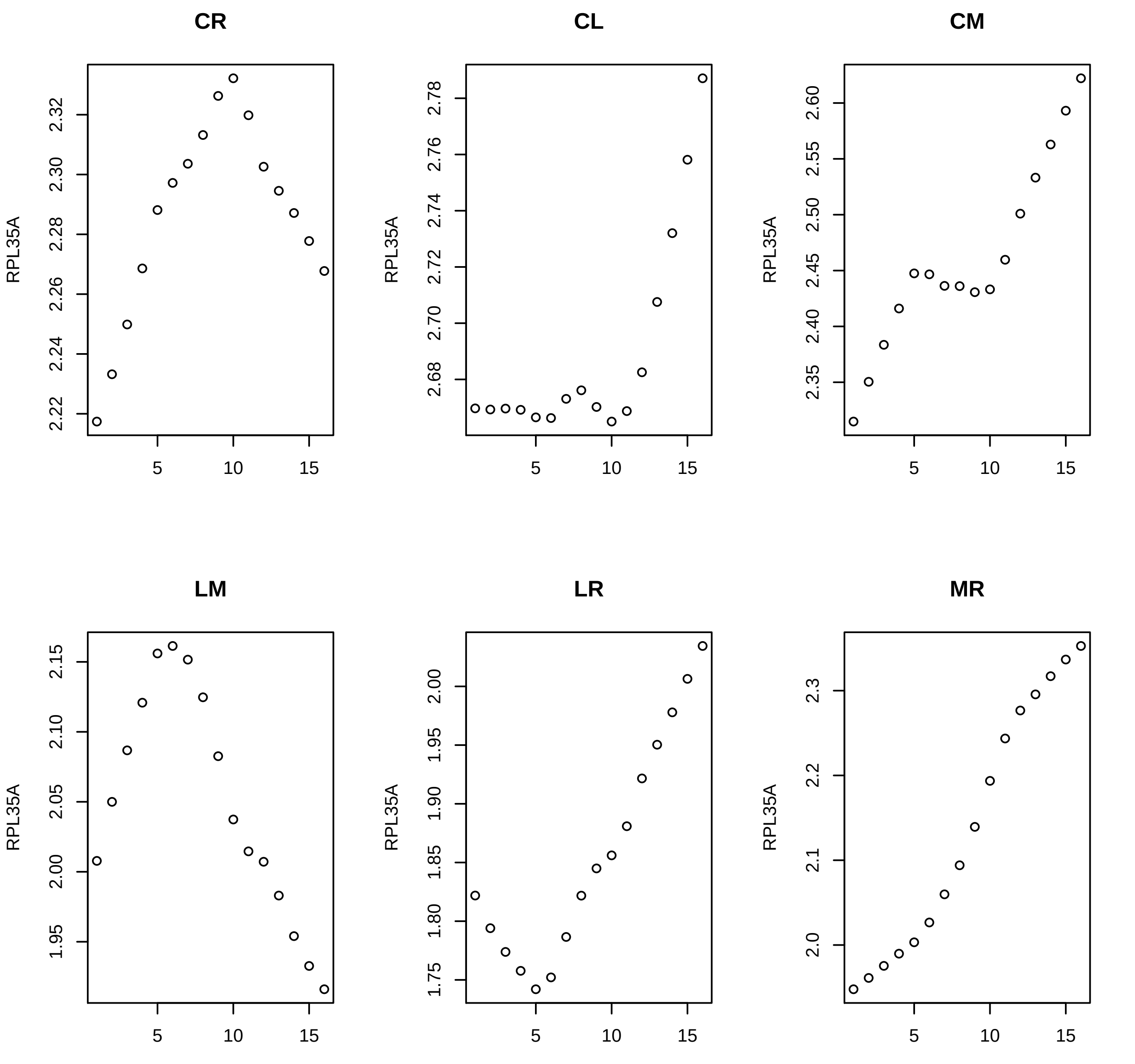
CR
CL
CM
LM
LR
MR
RPL35A
5
10
15

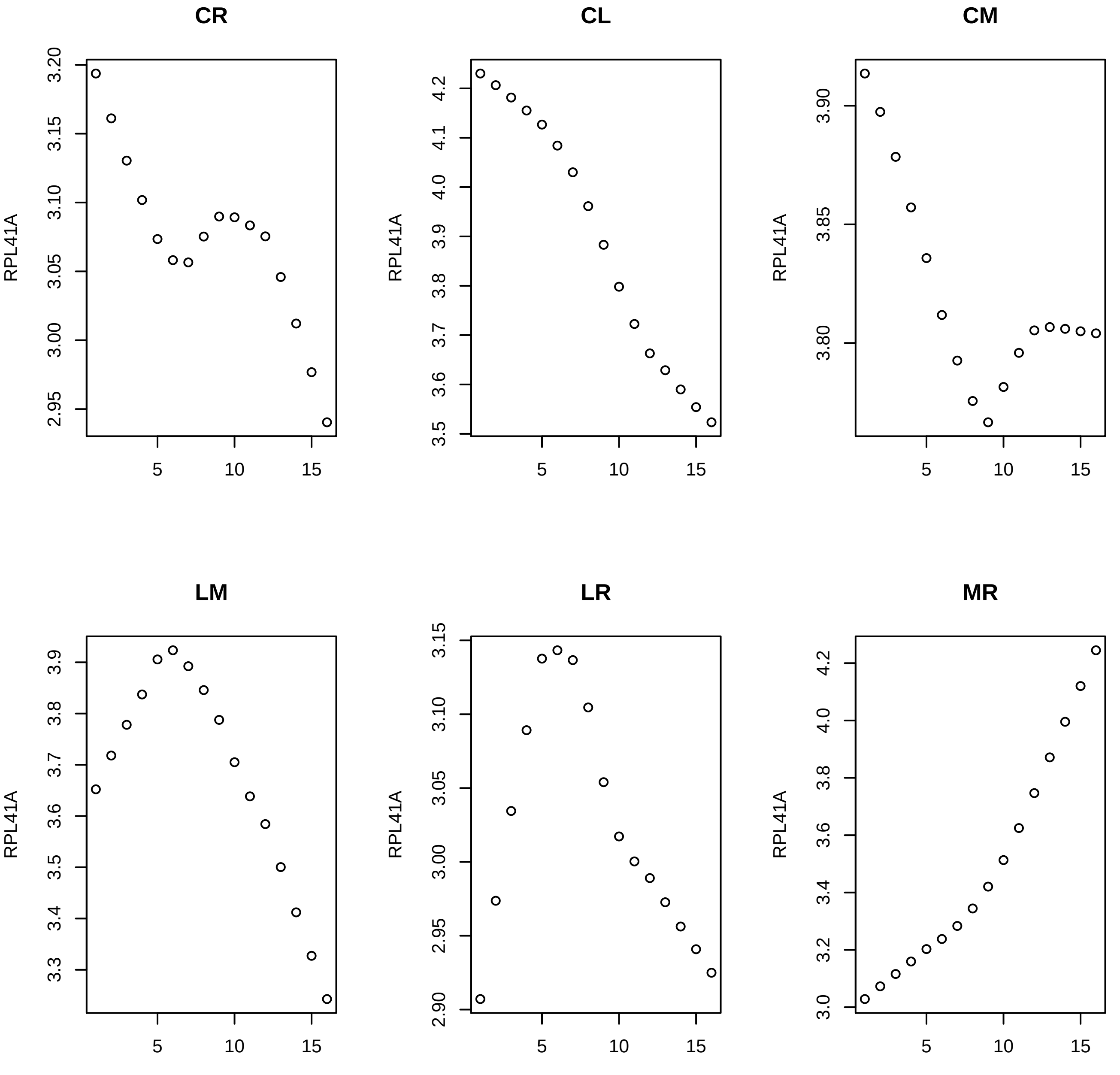

CR
CL
CM
LM
LR
MR
RPL41A

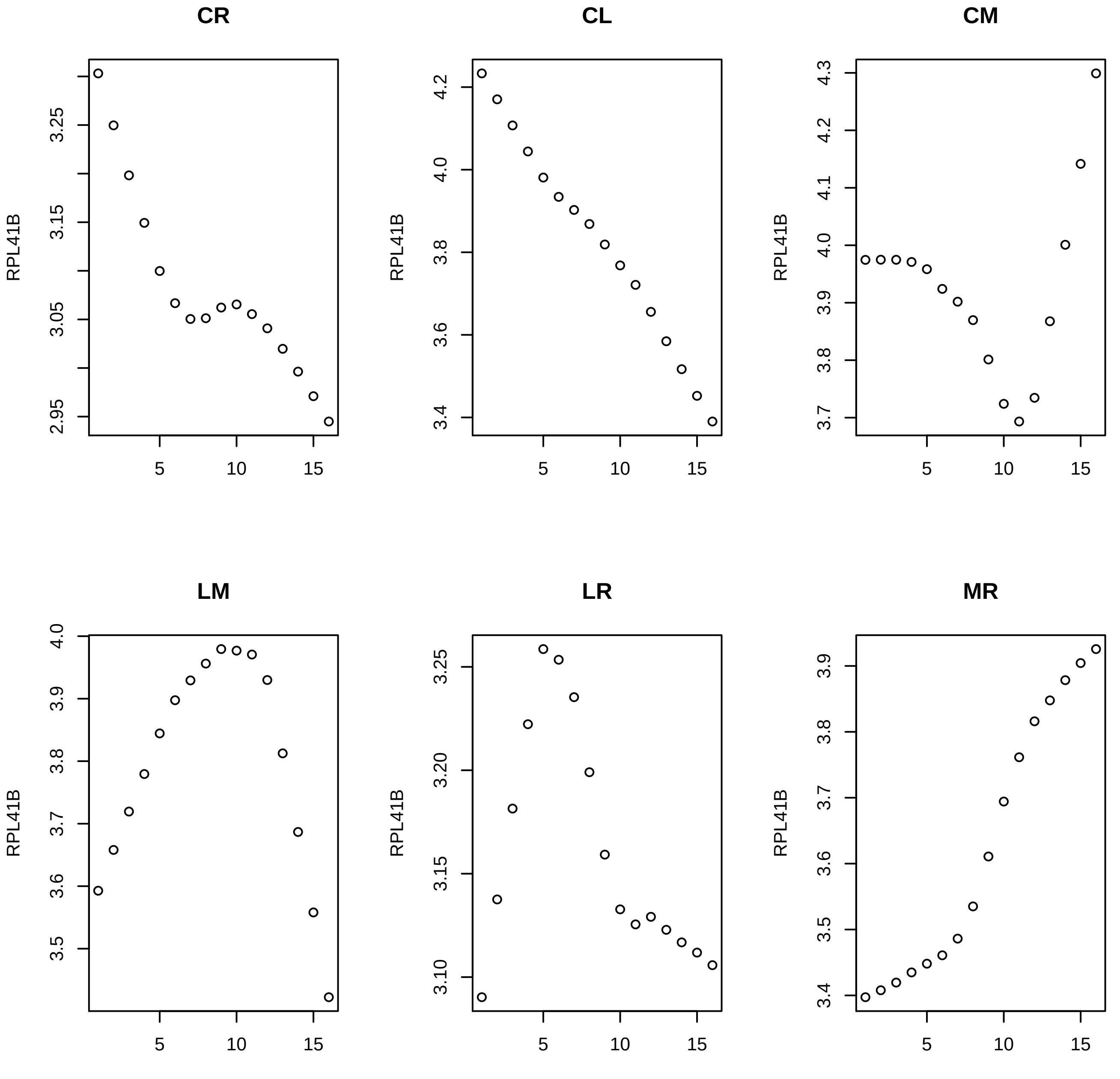
CR
CL
CM
LM
LR
MR
RPL41B

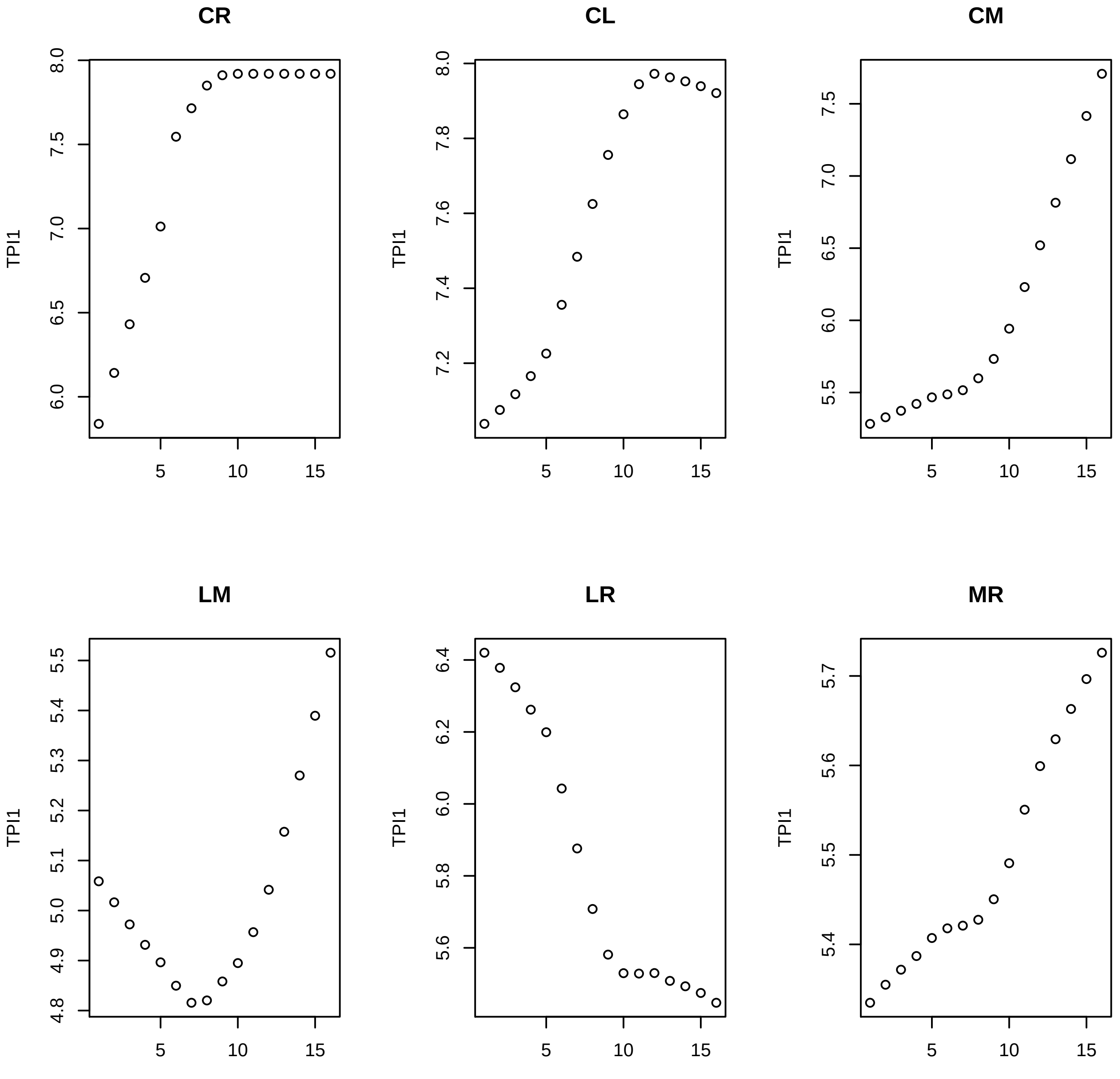
CR
CL
CM
LM
LR
MR
TPI1
5
10
15
6.0
6.5
7.0
7.5
8.0
7.2
7.4
7.6
7.8
5.5
4.8
4.9
5.0
5.1
5.2
5.3
5.4
5.6
5.8
6.2
6.4
5.7

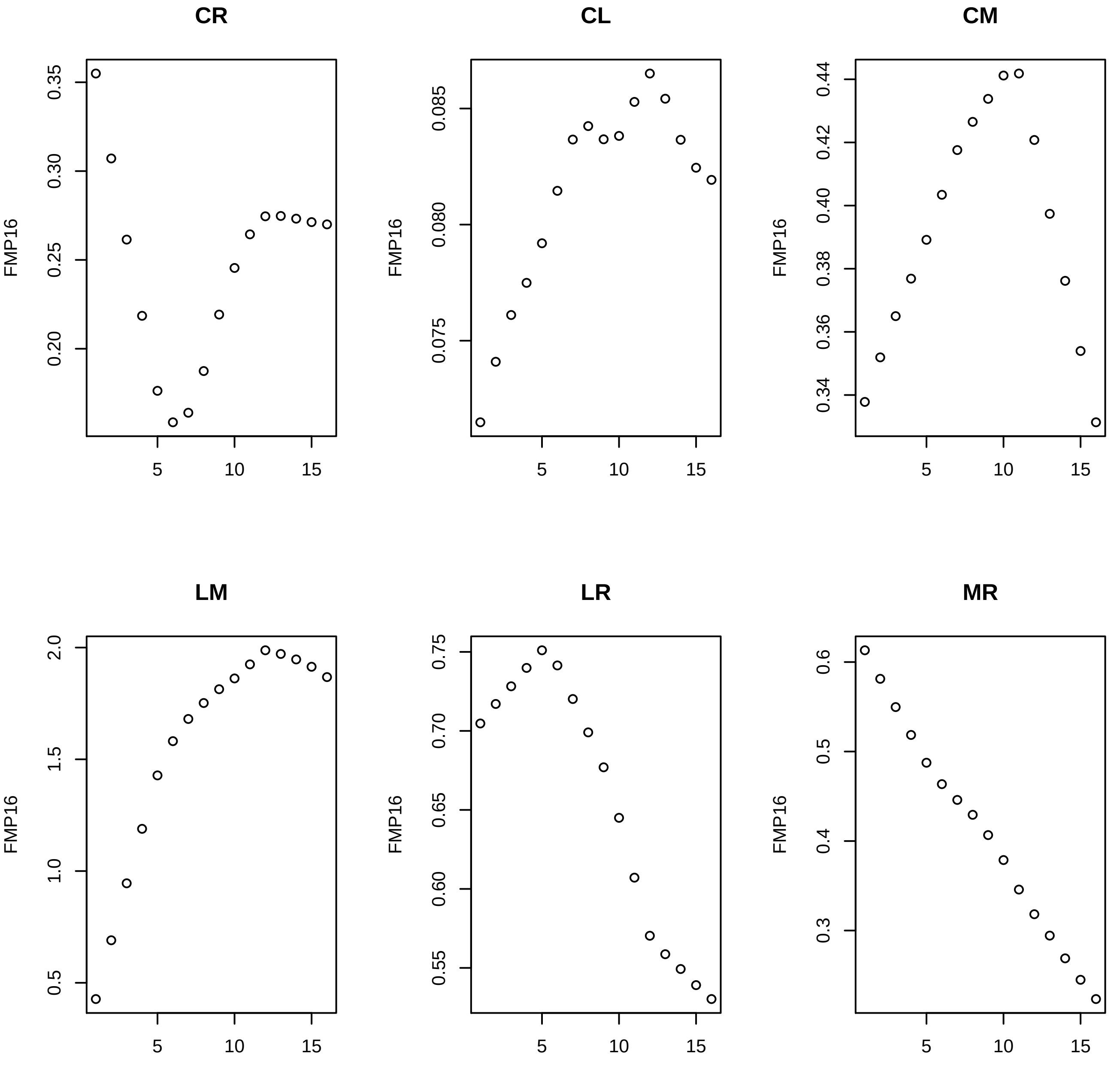
CR
CL
CM
LM
LR
MR
FMP16
0.35
0.30
0.25
0.20
0.085
0.080
0.075
0.44
0.42
0.40
0.38
0.36
0.34
2.0
1.5
1.0
0.5
0.75
0.70
0.65
0.60
0.55
0.6
0.5
0.4
0.3
5
10
15

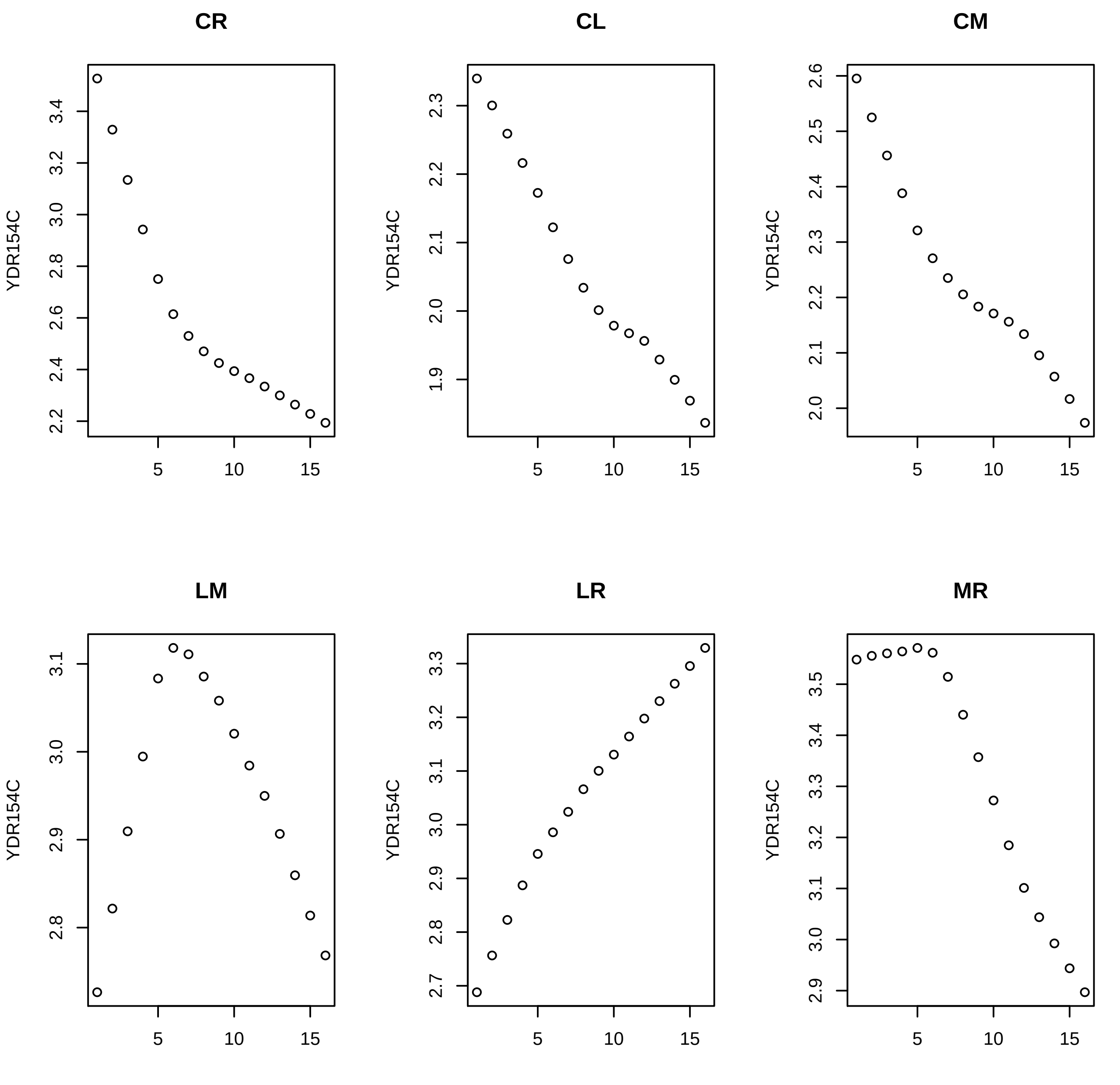
CR
CL
CM
LM
LR
MR
YDR154C

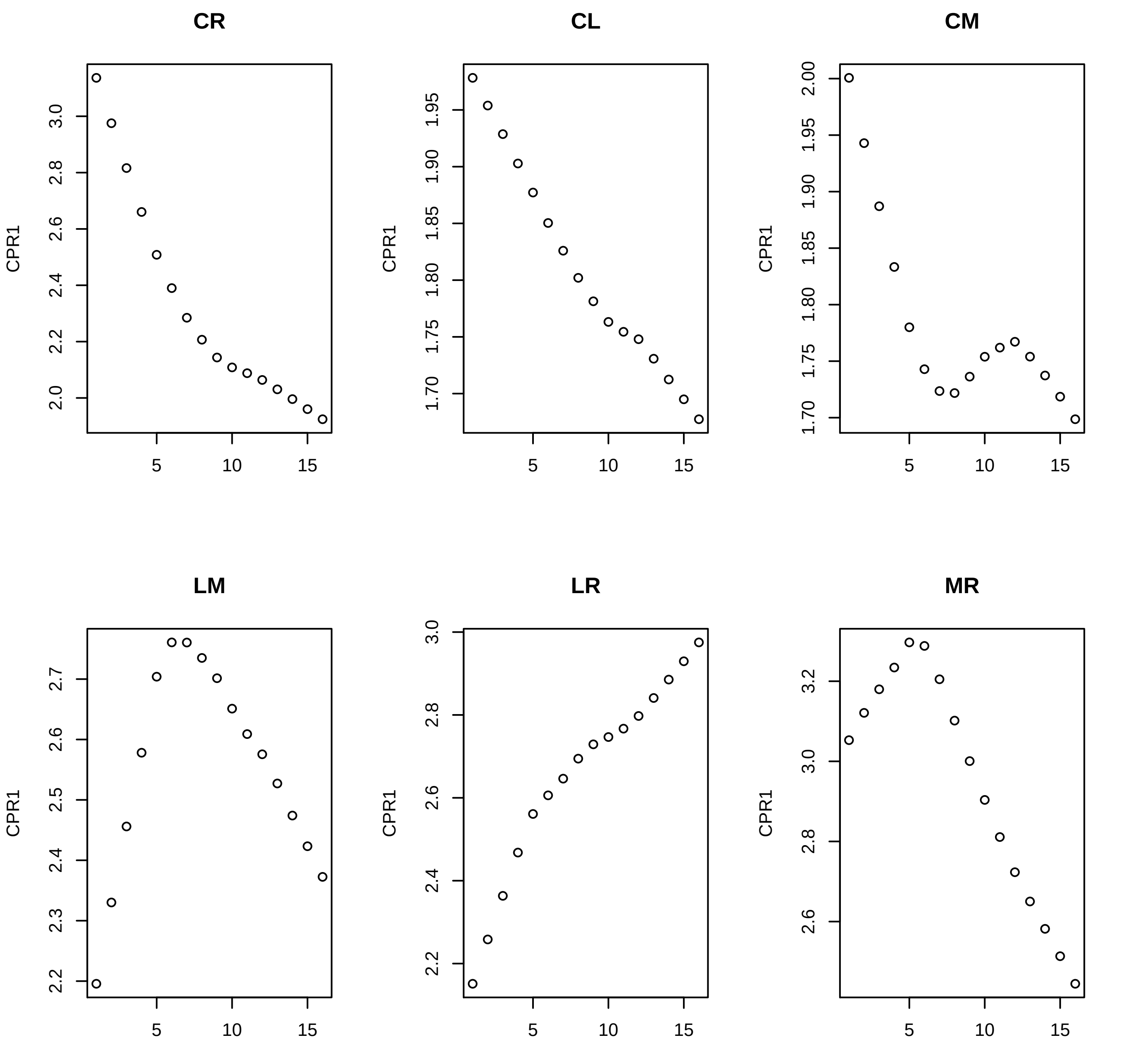

CR
CL
CM
LM
LR
MR
CPR1

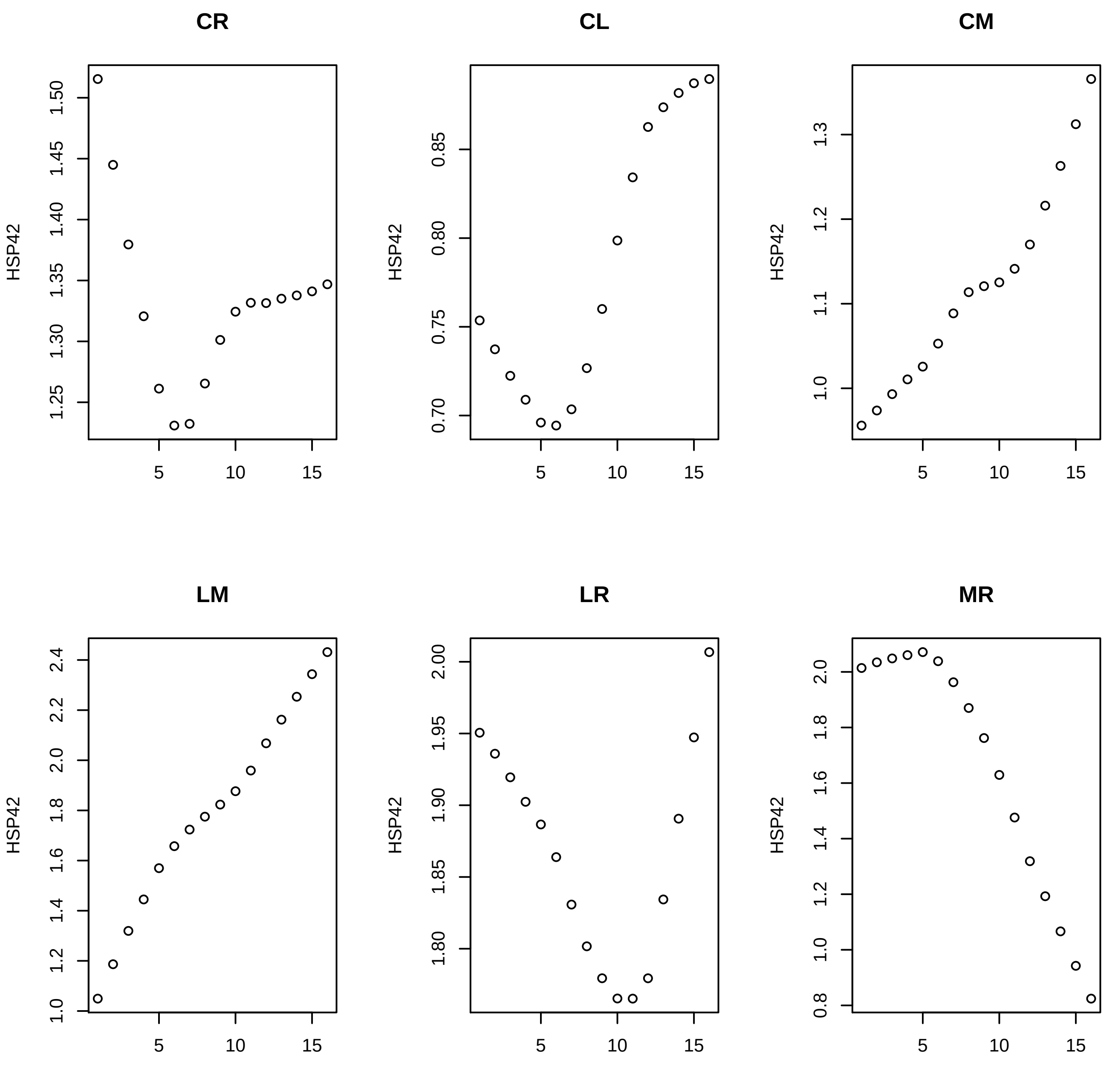

CR
CL
CM
LM
LR
MR
HSP42

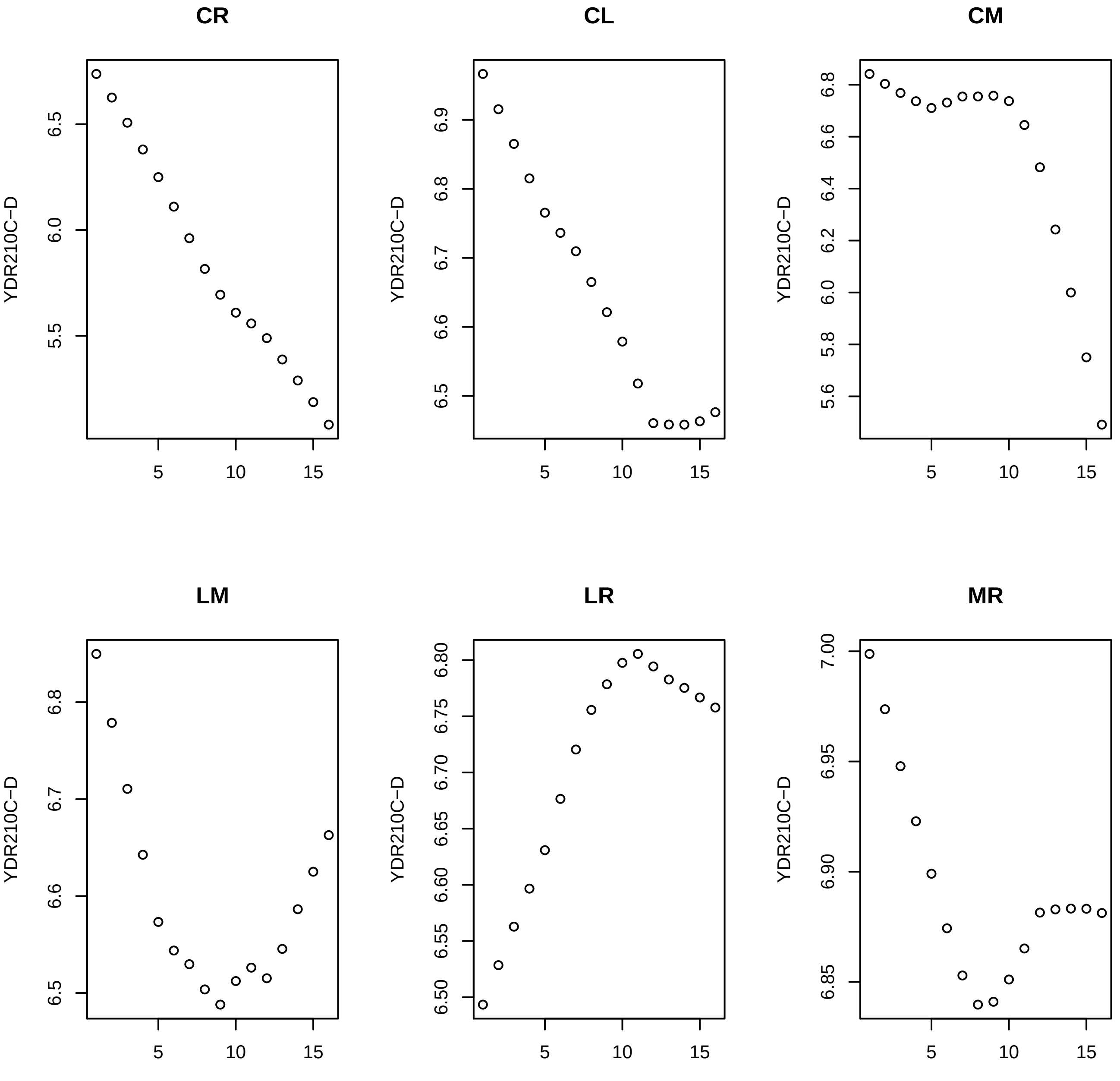

CR
CL
CM
LM
LR
MR
YDR210C−D

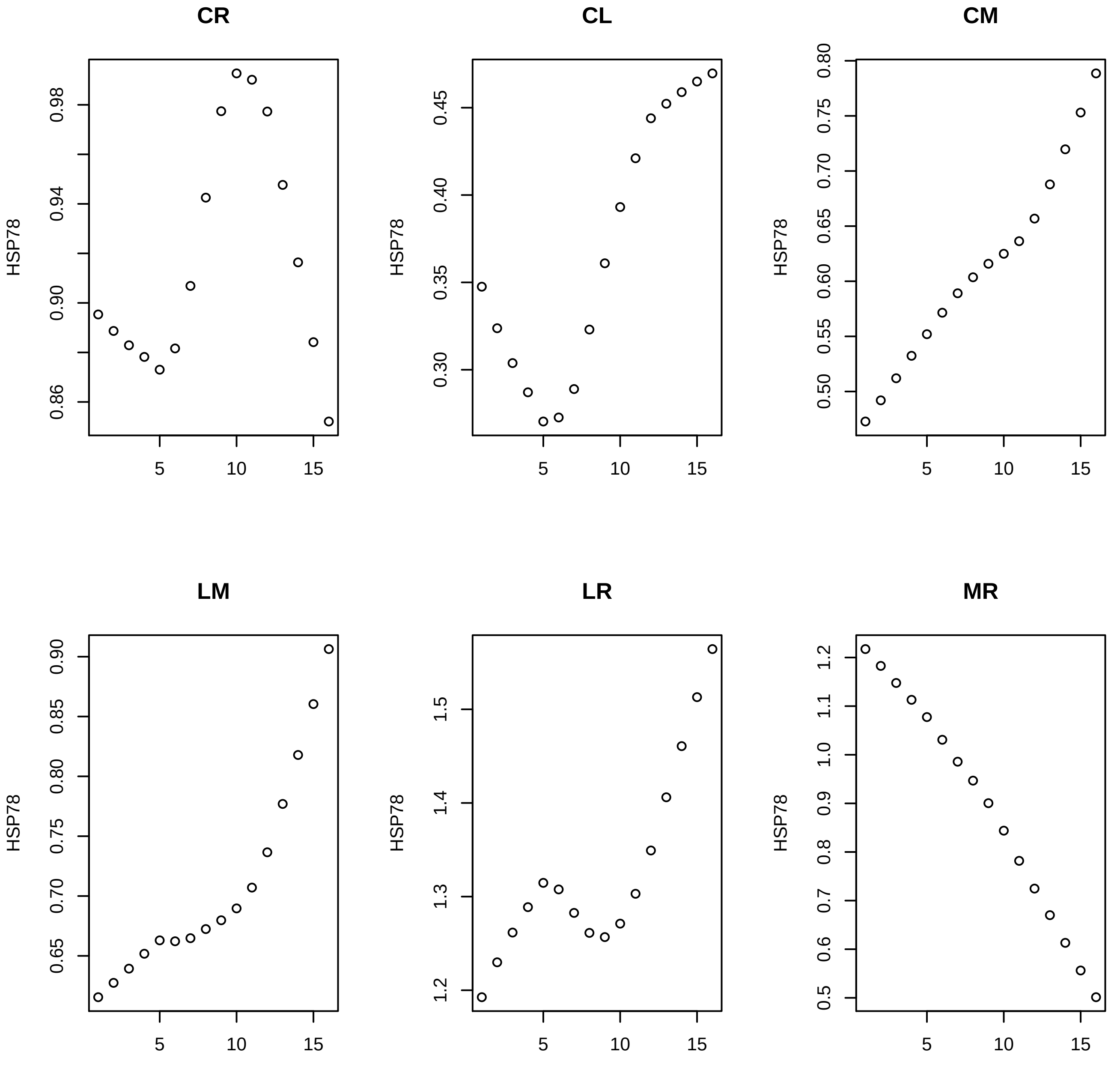
CR
CL
CM
LM
LR
MR
HSP78
0.86
0.90
0.94
0.98
0.30
0.35
0.40
0.45
0.50
0.55
0.60
0.65
0.70
0.75
0.80
0.85
0.90
1.2
1.3
1.4
1.5
0.5
0.6
0.7
0.8
0.9
1.0
1.1
5
10
15

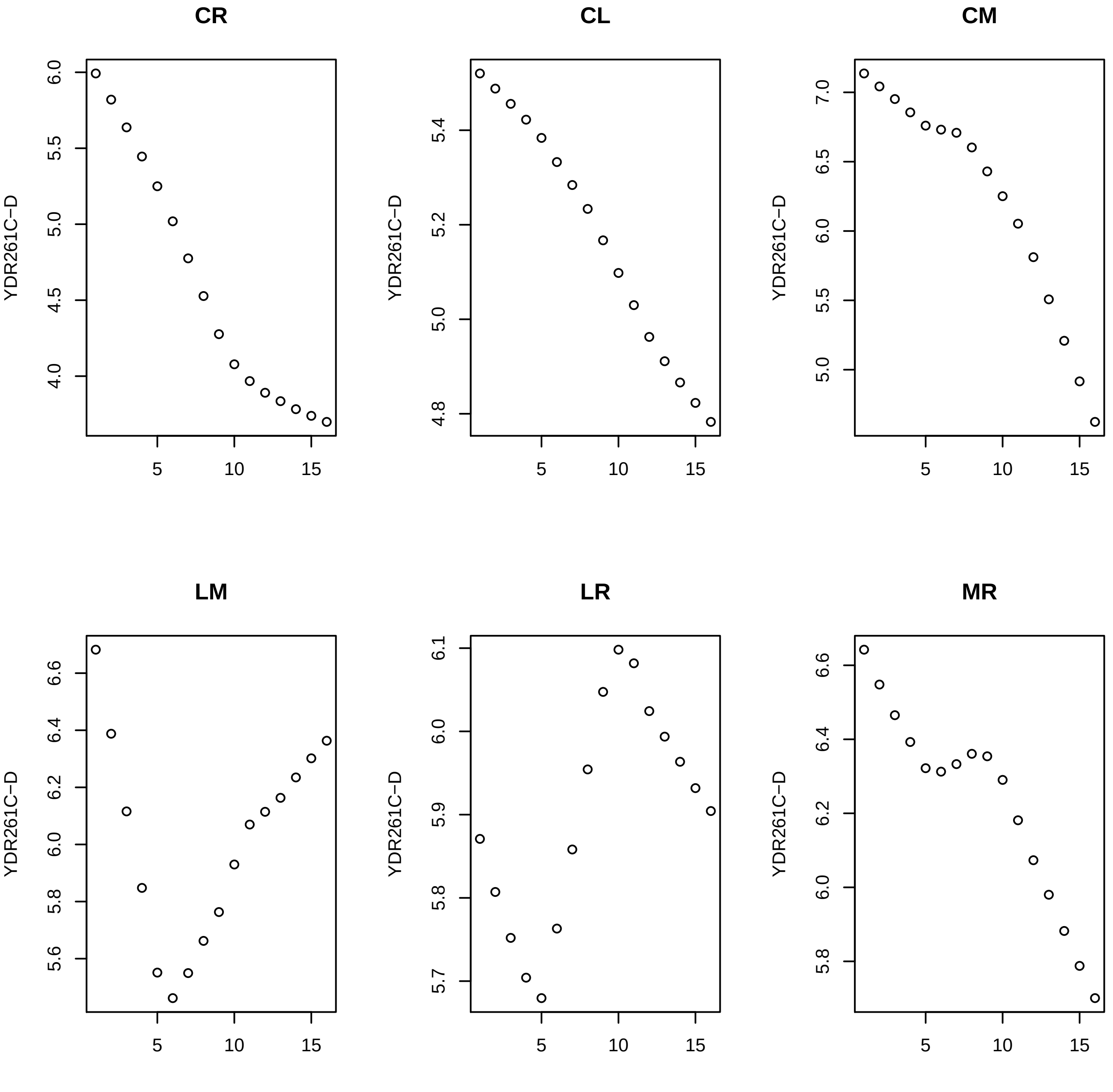
CR
CL
CM
LM
LR
MR
YDR261C−D

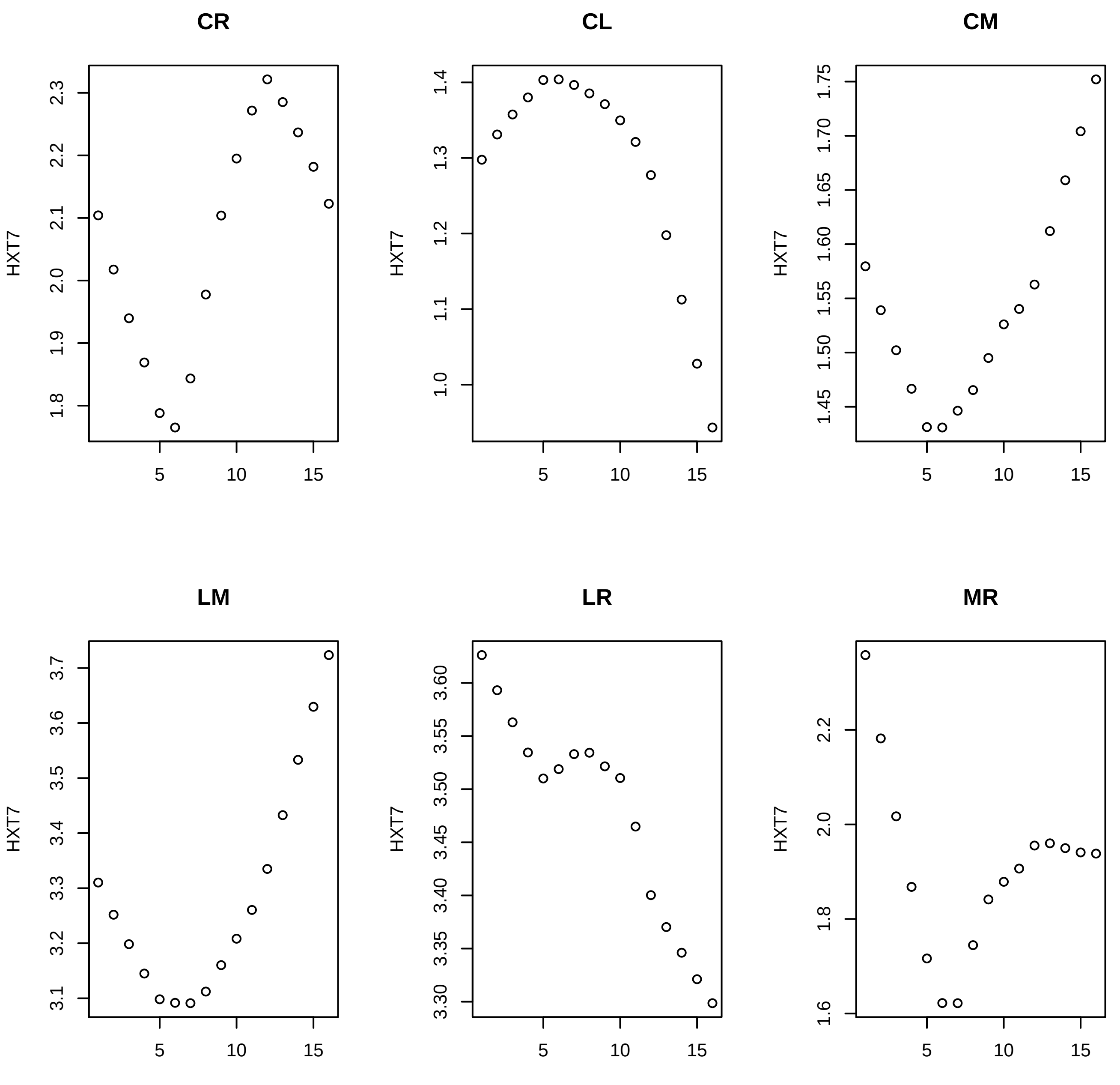

CR
CL
CM
LM
LR
MR
HXT7
5
10
15
2.3
2.2
2.1
2.0
1.9
1.8
1.4
1.3
1.2
1.1
1.0
1.75
1.70
1.65
1.60
1.55
1.50
1.45
3.7
3.6
3.5
3.4
3.3
3.2
3.1
3.60
3.55
3.50
3.45
3.40
3.35
3.30
2.2
2.0
1.8
1.6

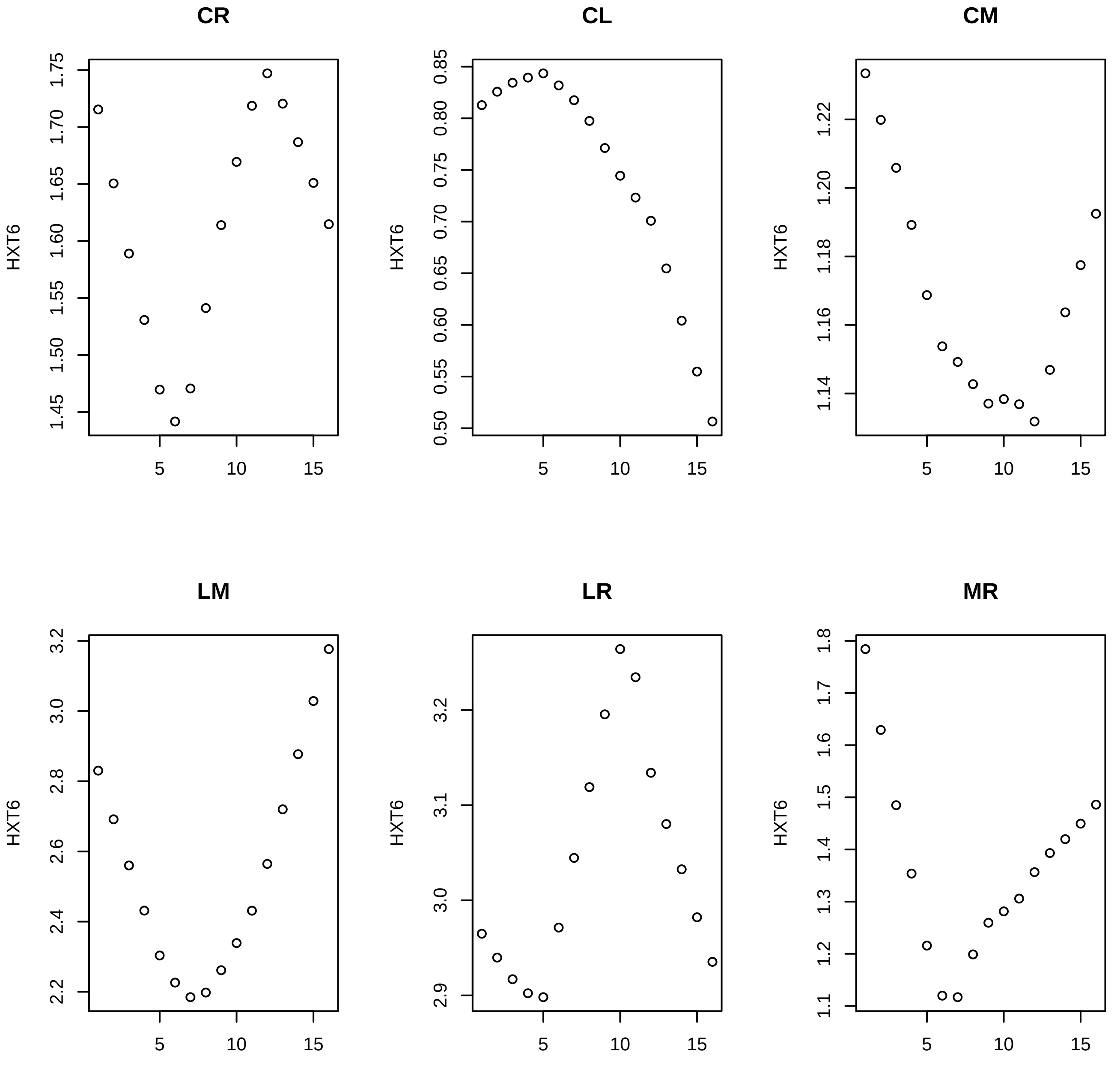
CR
CL
CM
LM
LR
MR
HXT6
1.45
1.50
1.55
1.60
1.65
1.70
1.75
0.50
0.55
0.60
0.65
0.70
0.75
0.80
0.85
1.14
1.16
1.18
1.20
1.22
2.2
2.4
2.6
2.8
3.0
3.2
2.9
3.0
3.1
3.2
1.1
1.2
1.3
1.4
1.5
1.6
1.7
1.8
5
10
15

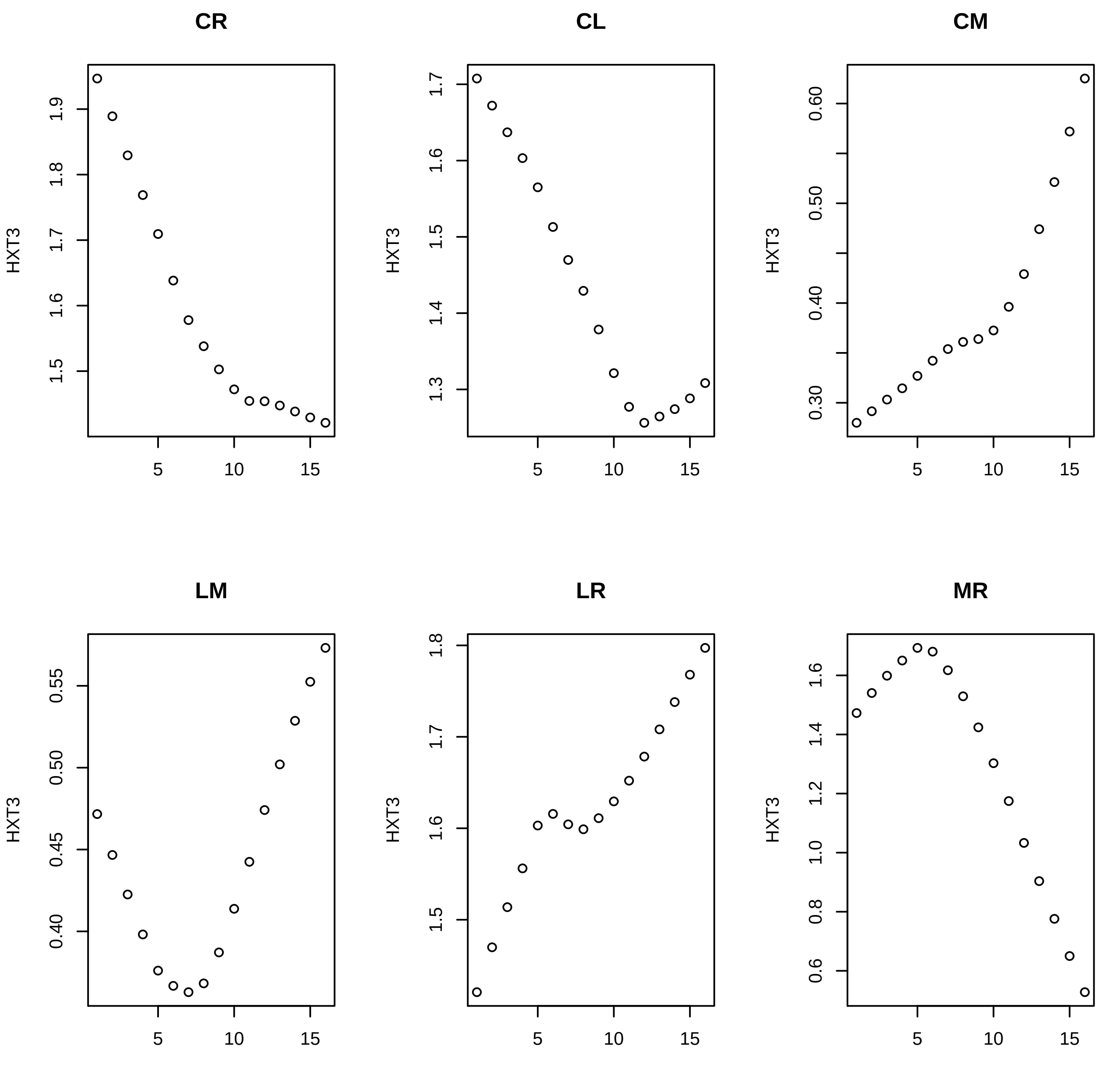
CR
CL
CM
LM
LR
MR
HXT3
1.9
1.8
1.7
1.6
1.5
1.4
1.3
0.60
0.50
0.40
0.30
0.55
0.45
1.2
1.0
0.8
0.6
5
10
15

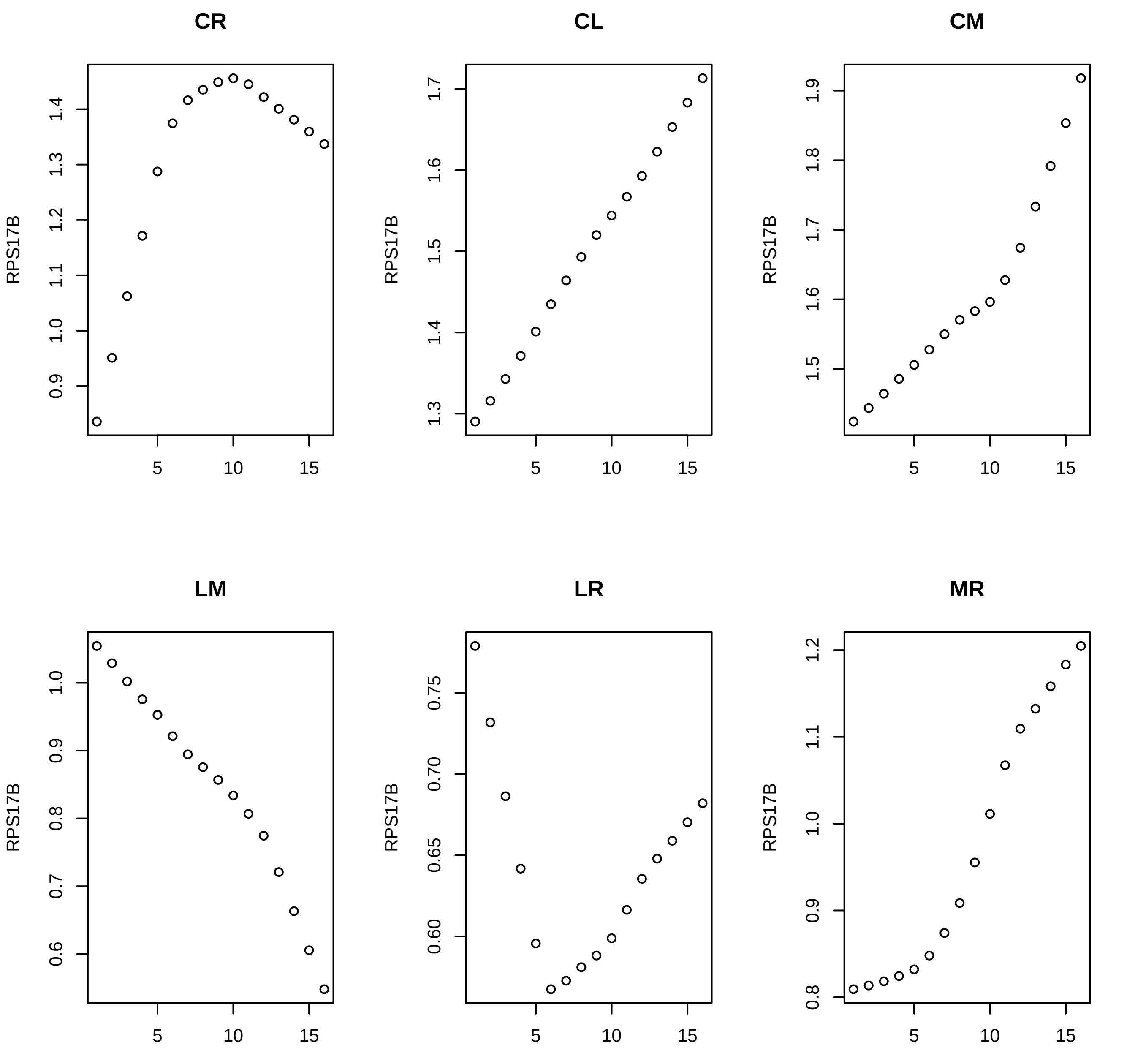

CR
CL
CM
LM
LR
MR
RPS17B

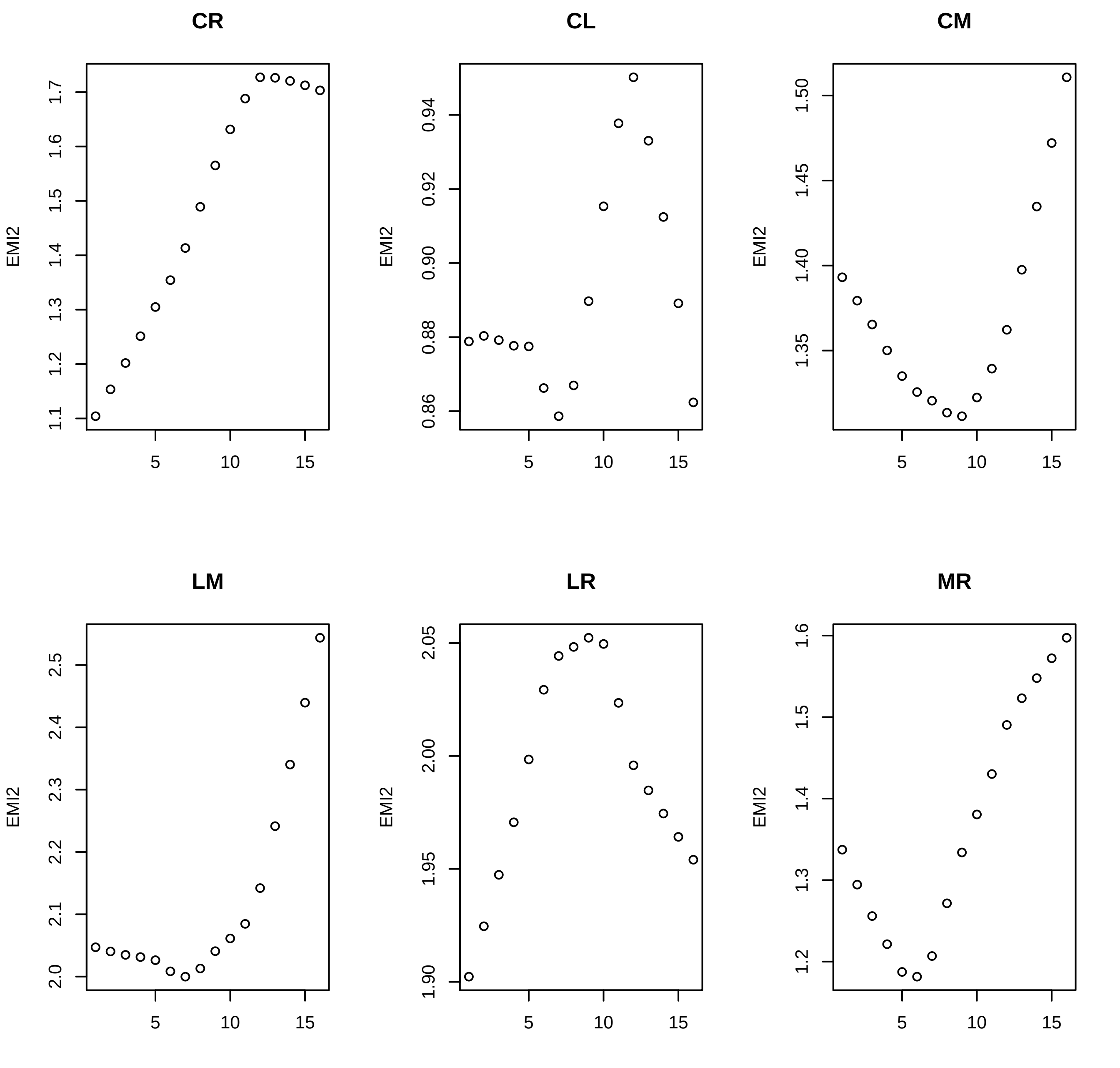
CR
CL
CM
LM
LR
MR
EMI2
1.1
1.2
1.3
1.4
1.5
1.6
1.7
0.86
0.88
0.90
0.92
0.94
1.35
1.40
1.45
1.50
2.0
2.1
2.2
2.3
2.4
2.5
1.90
1.95
2.00
2.05
1.2
1.3
1.4
1.5
1.6
5
10
15

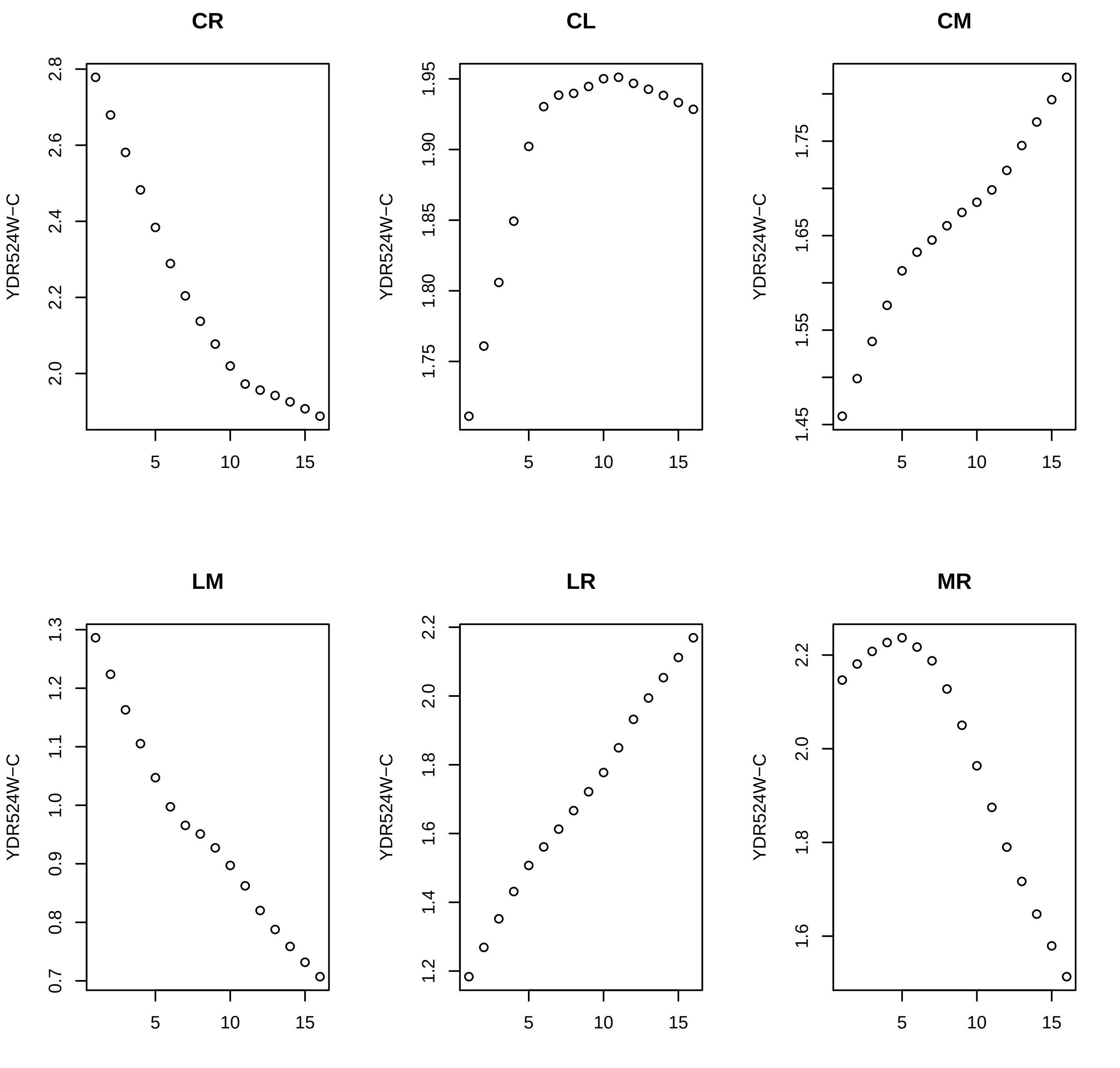
CR
CL
CM
LM
LR
MR
YDR524W−C
2.8
2.6
2.4
2.2
2.0
1.95
1.90
1.85
1.80
1.75
1.65
1.55
1.45
1.3
1.2
1.1
1.0
0.9
0.8
0.7
1.8
1.6
1.4
5
10
15

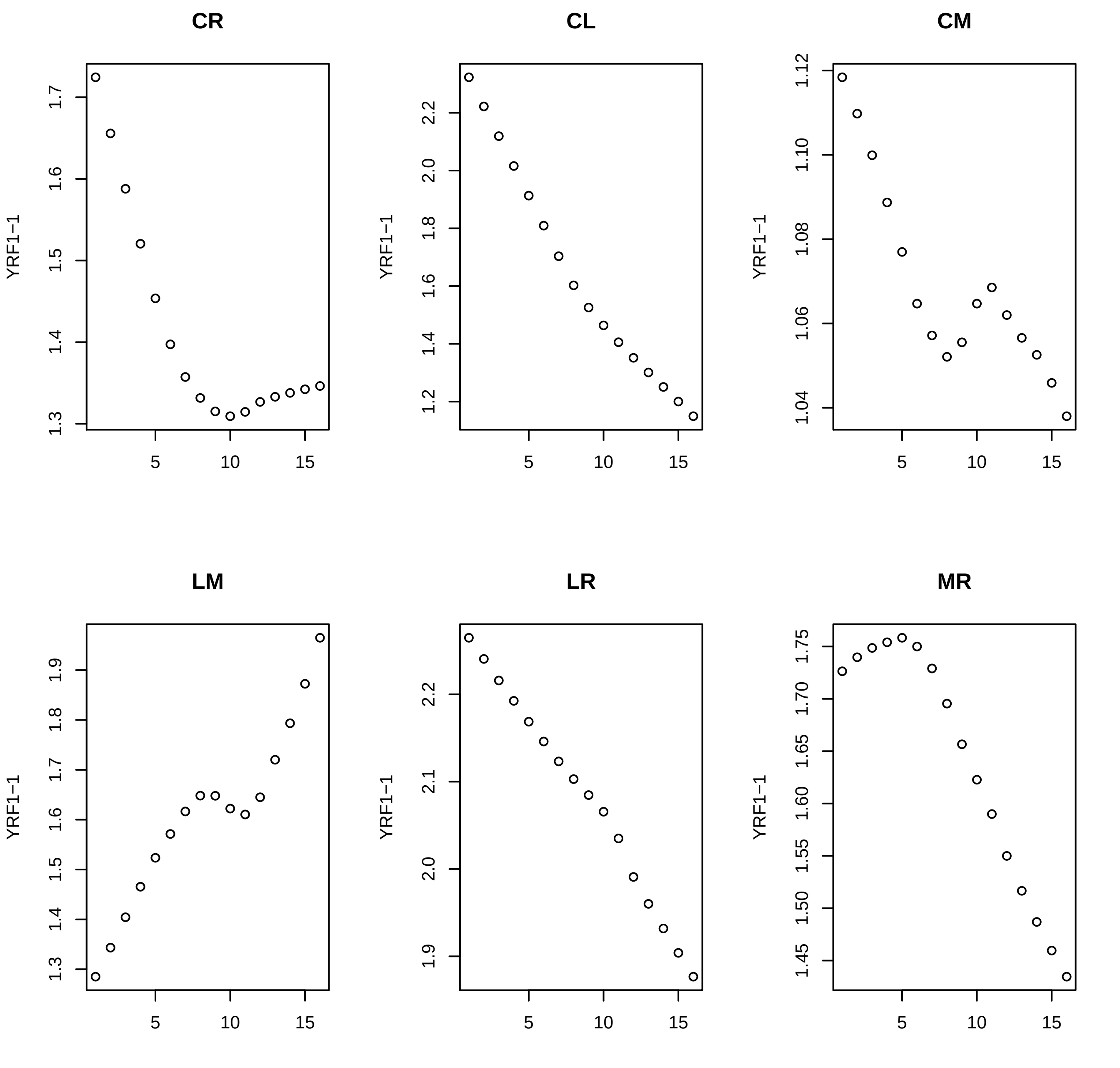
CR
CL
CM
LM
LR
MR
YRF1−1

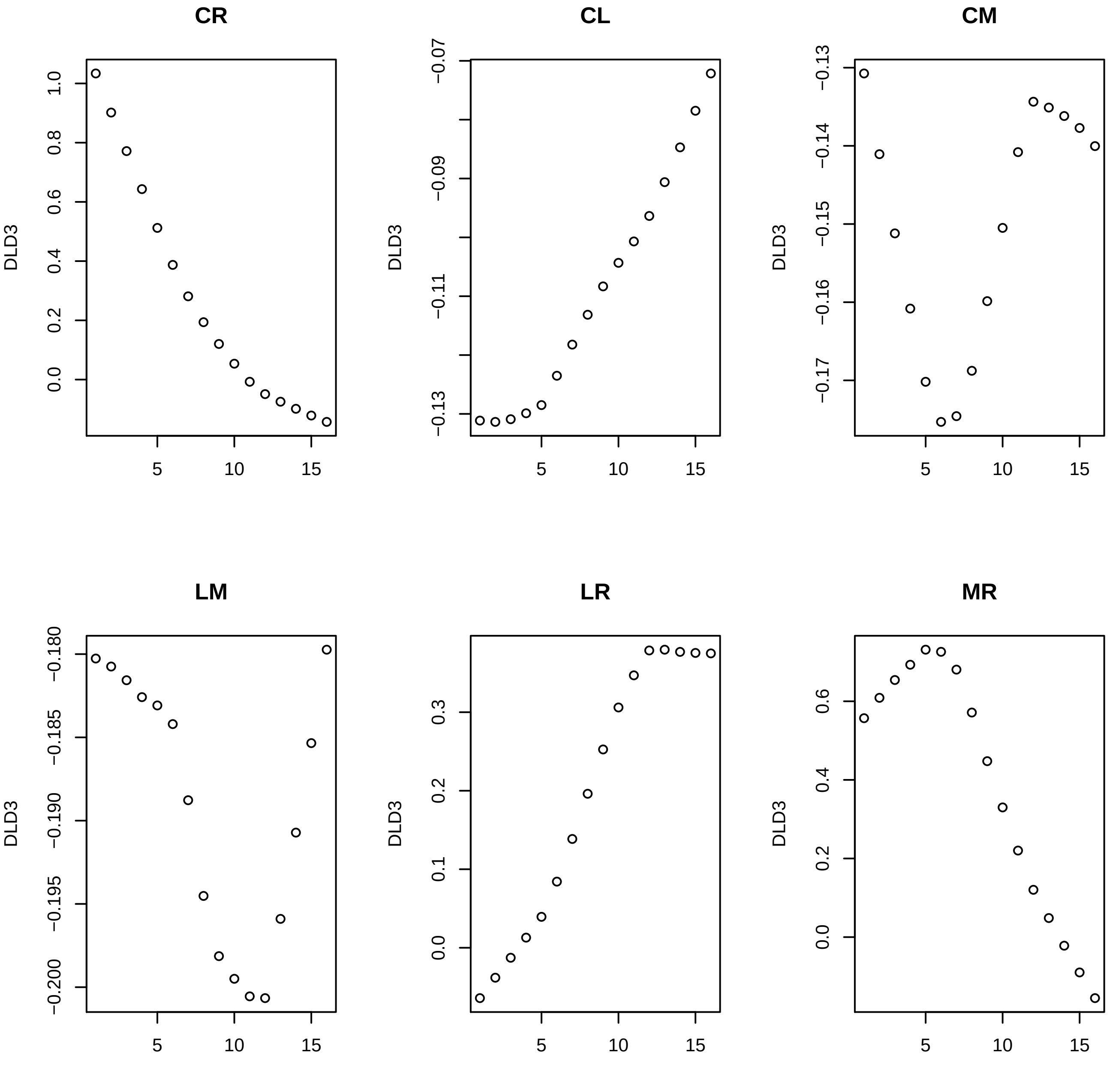
CR
CL
CM
LM
LR
MR
DLD3
5
10
15

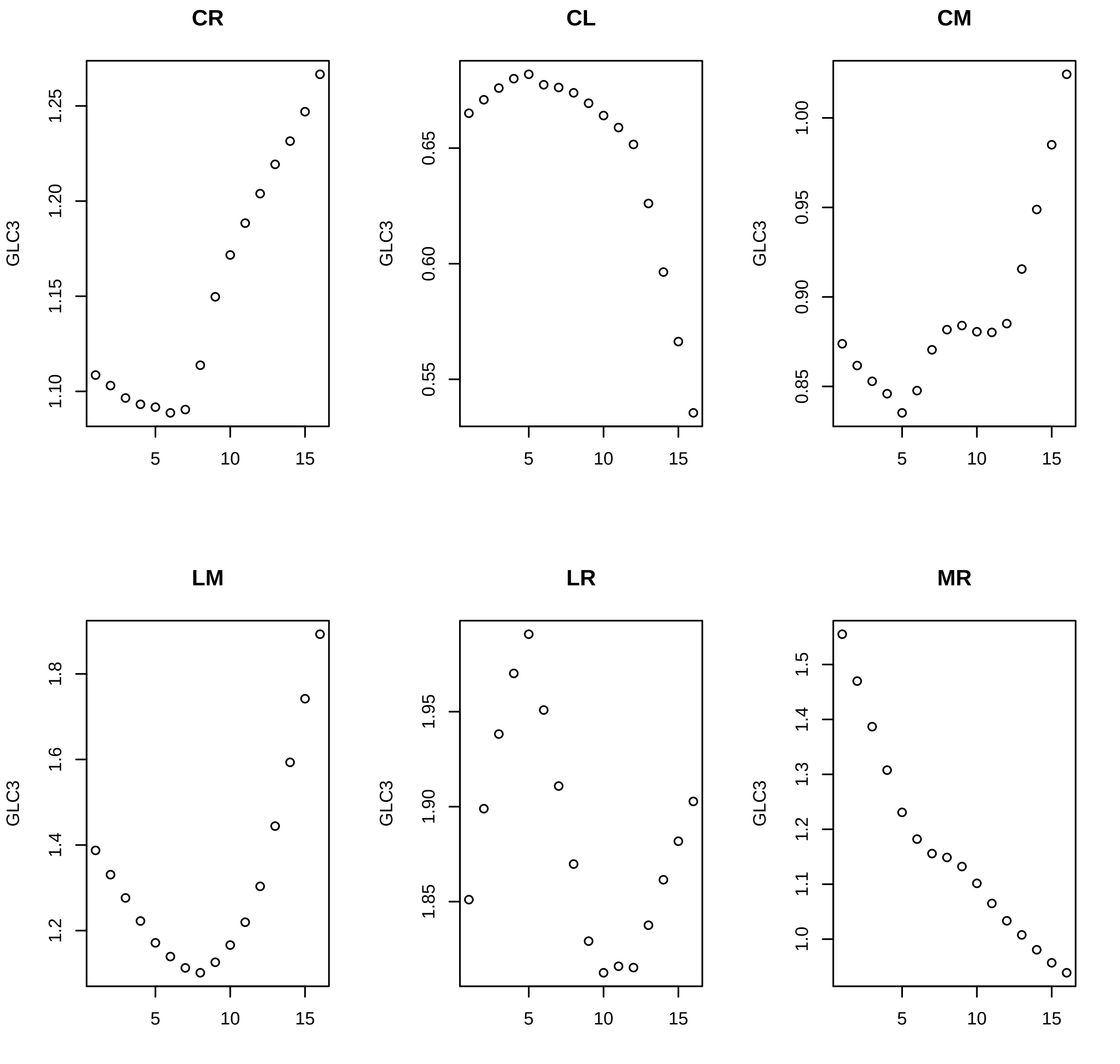
CR
CL
CM
LM
LR
MR
GLC3
1.10
1.15
1.20
1.25
0.55
0.60
0.65
0.85
0.90
0.95
1.00
1.2
1.4
1.6
1.8
1.85
1.90
1.95
1.0
1.1
1.2
1.3
1.4
1.5
5
10
15

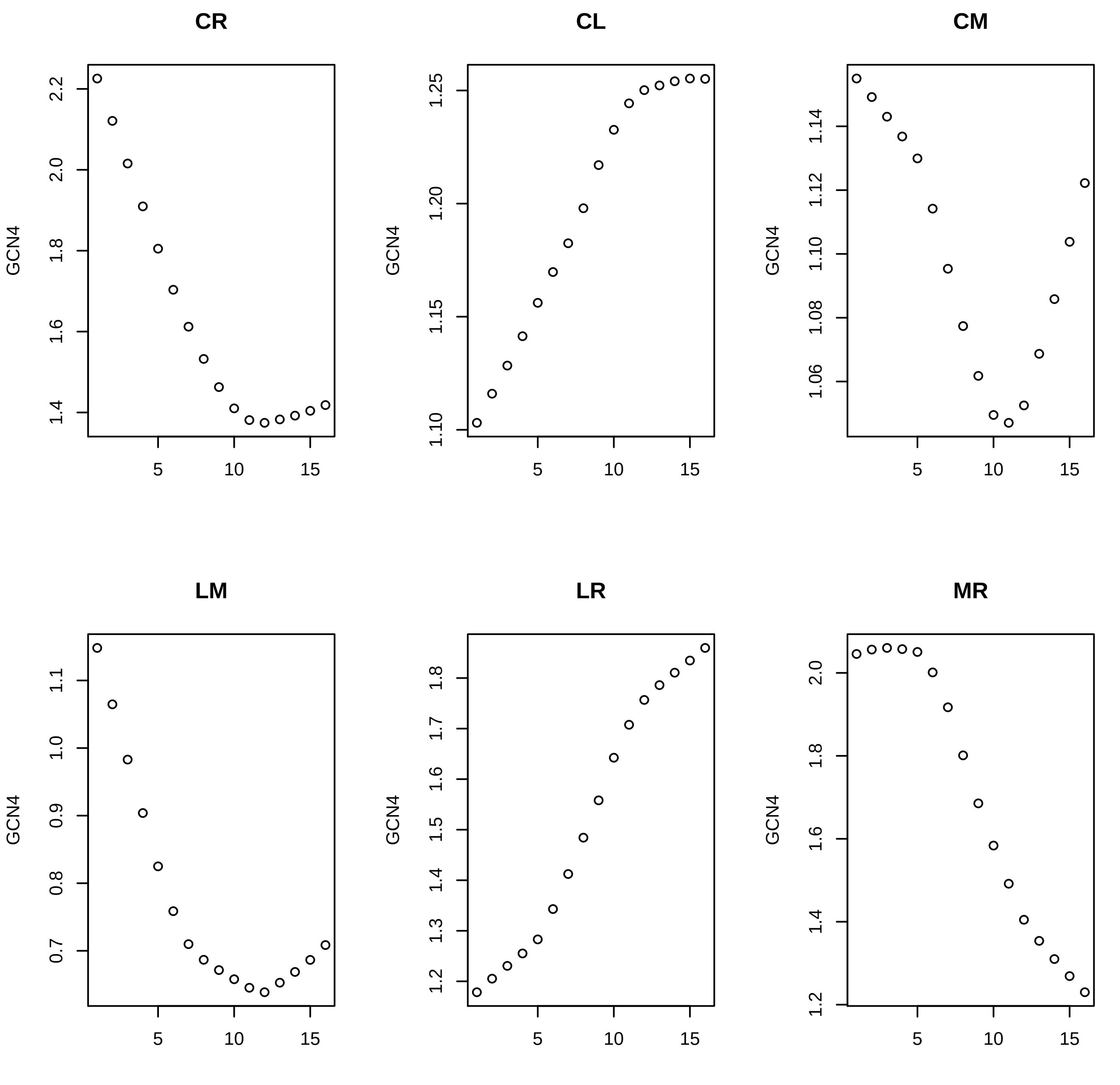

CR
CL
CM
LM
LR
MR
GCN4
2.2
2.0
1.8
1.6
1.4
1.25
1.20
1.15
1.10
1.14
1.12
1.10
1.08
1.06
1.1
1.0
0.9
0.8
0.7
1.8
1.7
1.6
1.5
1.4
1.3
1.2
2.0
1.8
1.6
1.4
1.2
5
10
15

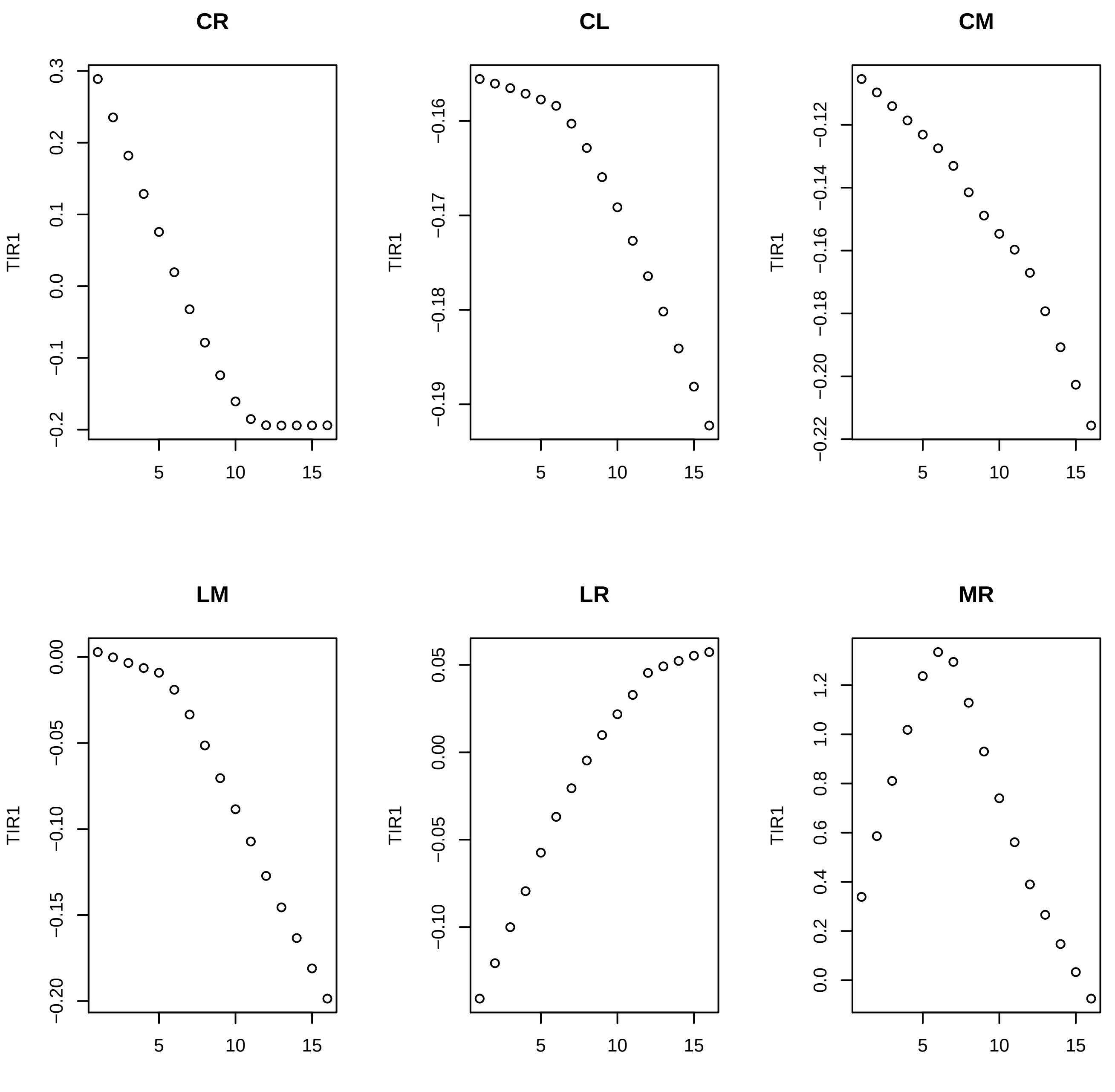

CR
TIR1
0.3
0.2
0.1
0.0
−0.1
−0.2
5
10
15
CL
TIR1
−0.16
−0.17
−0.18
−0.19
5
10
15
CM
TIR1
−0.12
−0.14
−0.16
−0.18
−0.20
−0.22
5
10
15
LM
TIR1
0.00
−0.05
−0.10
−0.15
−0.20
5
10
15
LR
TIR1
0.05
0.00
−0.05
−0.10
5
10
15
MR
TIR1
1.2
1.0
0.8
0.6
0.4
0.2
0.0
5
10
15

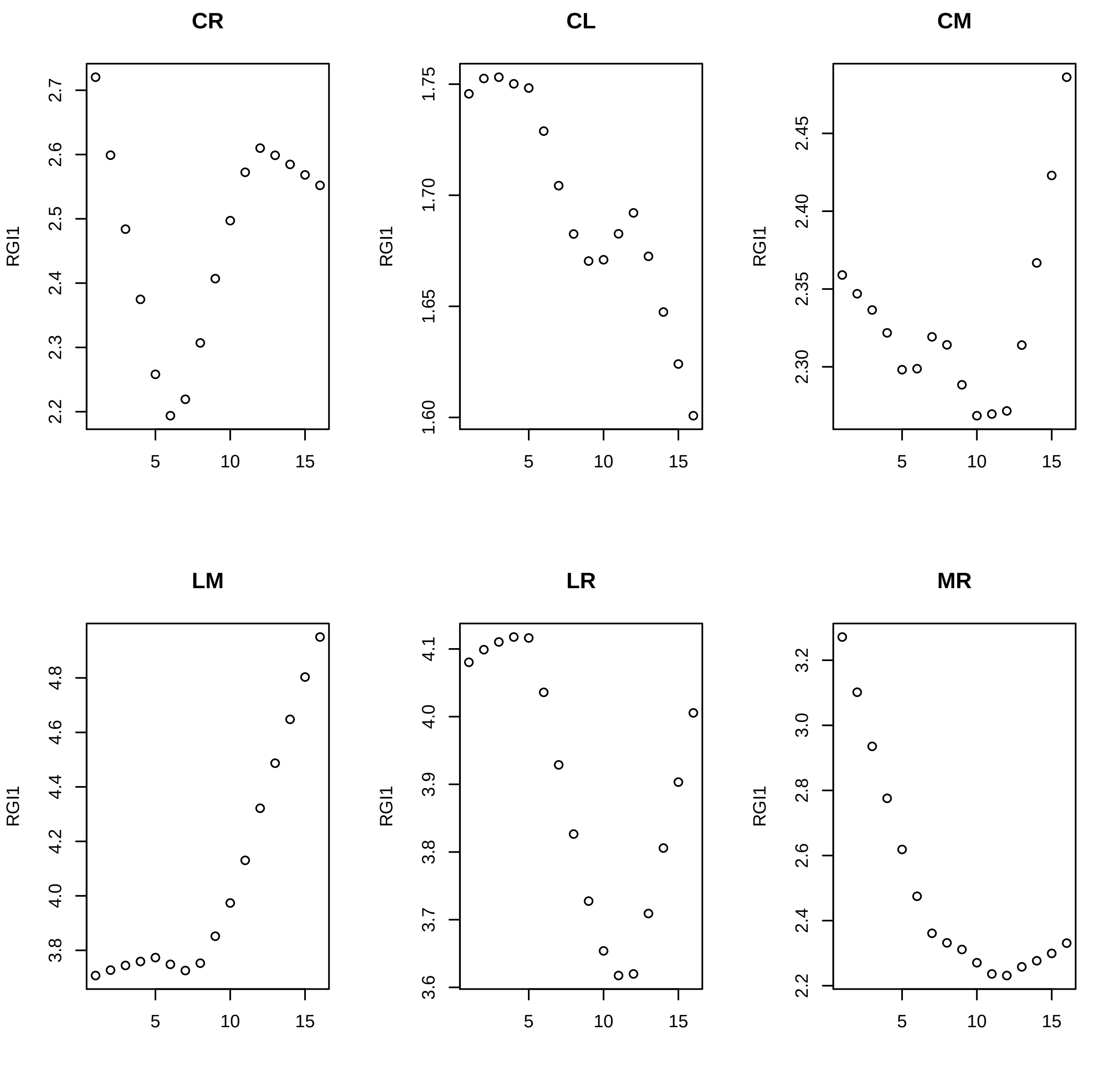
CR
CL
CM
LM
LR
MR
RGI1
5
10
15
2.2
2.3
2.4
2.5
2.6
2.7
1.60
1.65
1.70
1.75
2.30
2.35
2.40
2.45
3.8
4.0
4.2
4.4
4.6
4.8
3.6
3.7
3.8
3.9
4.0
4.1
2.2
2.4
2.6
2.8
3.0
3.2

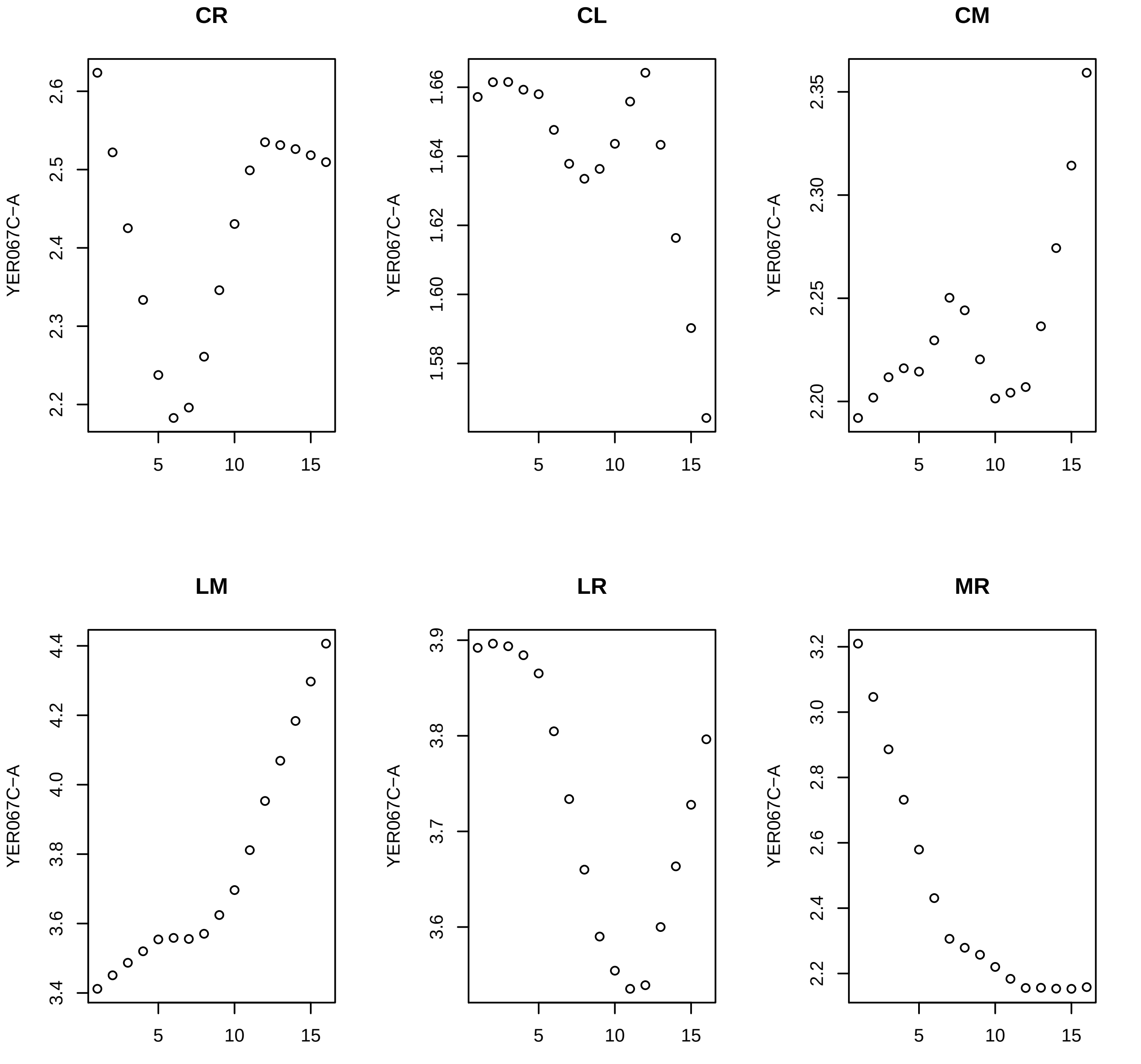

CR
CL
CM
LM
LR
MR
YER067C−A

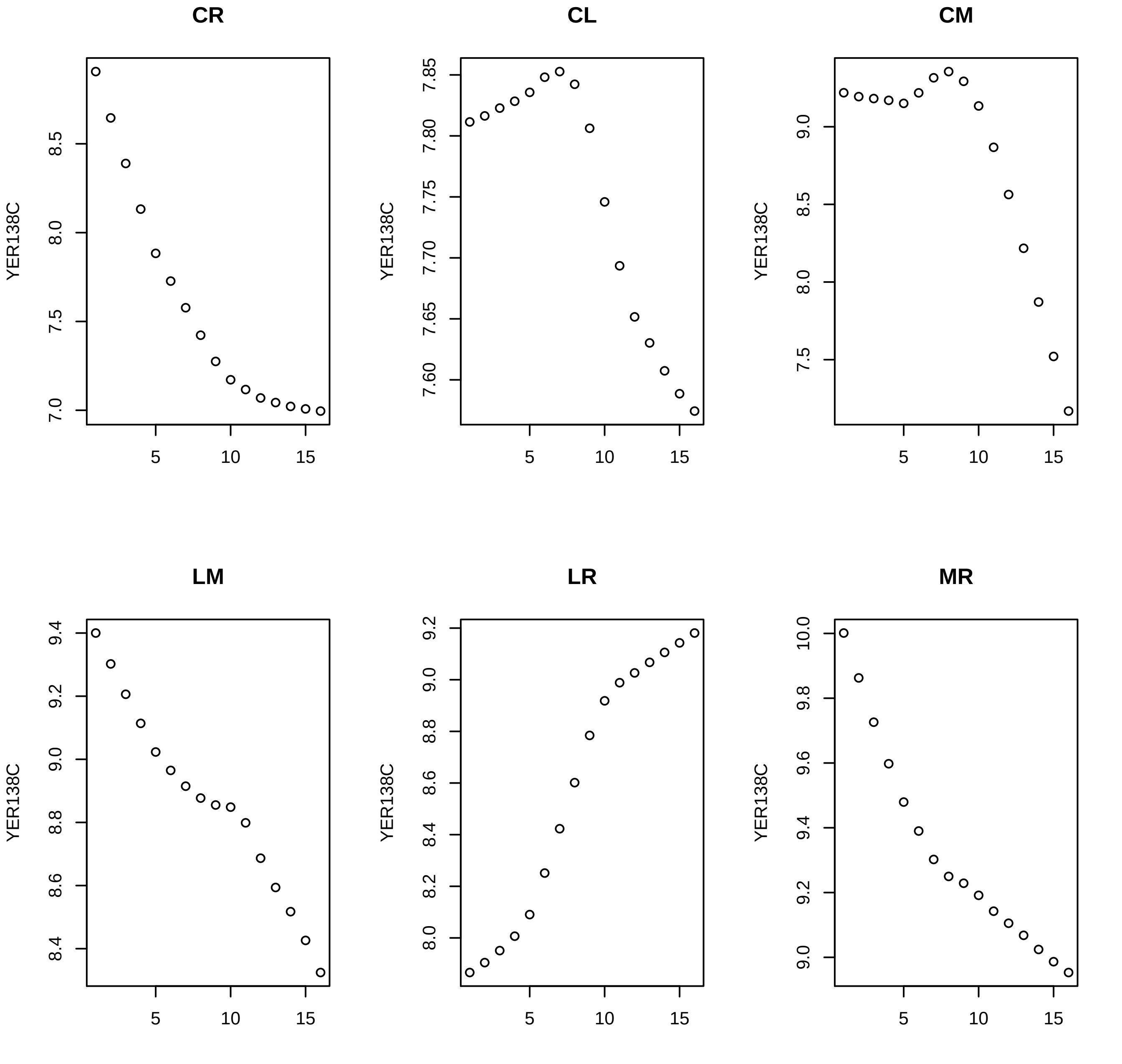
CR
CL
CM
LM
LR
MR
YER138C
8.5
8.0
7.5
7.0
7.85
7.80
7.75
7.70
7.65
7.60
9.0
9.4
9.2
8.8
8.6
8.4
8.2
10.0
9.8
9.6
5
10
15

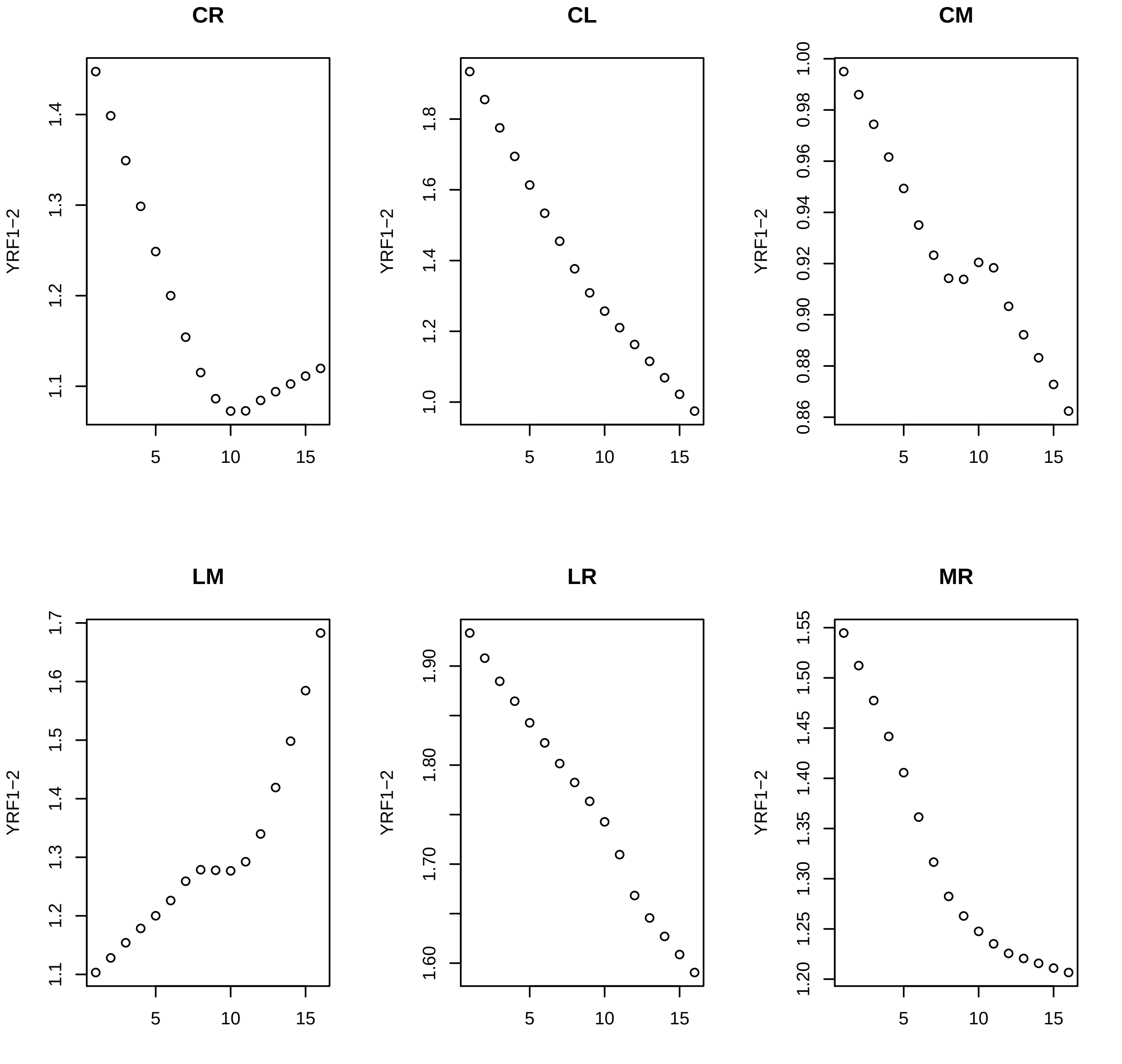

CR
CL
CM
LM
LR
MR
YRF1−2
5
10
15

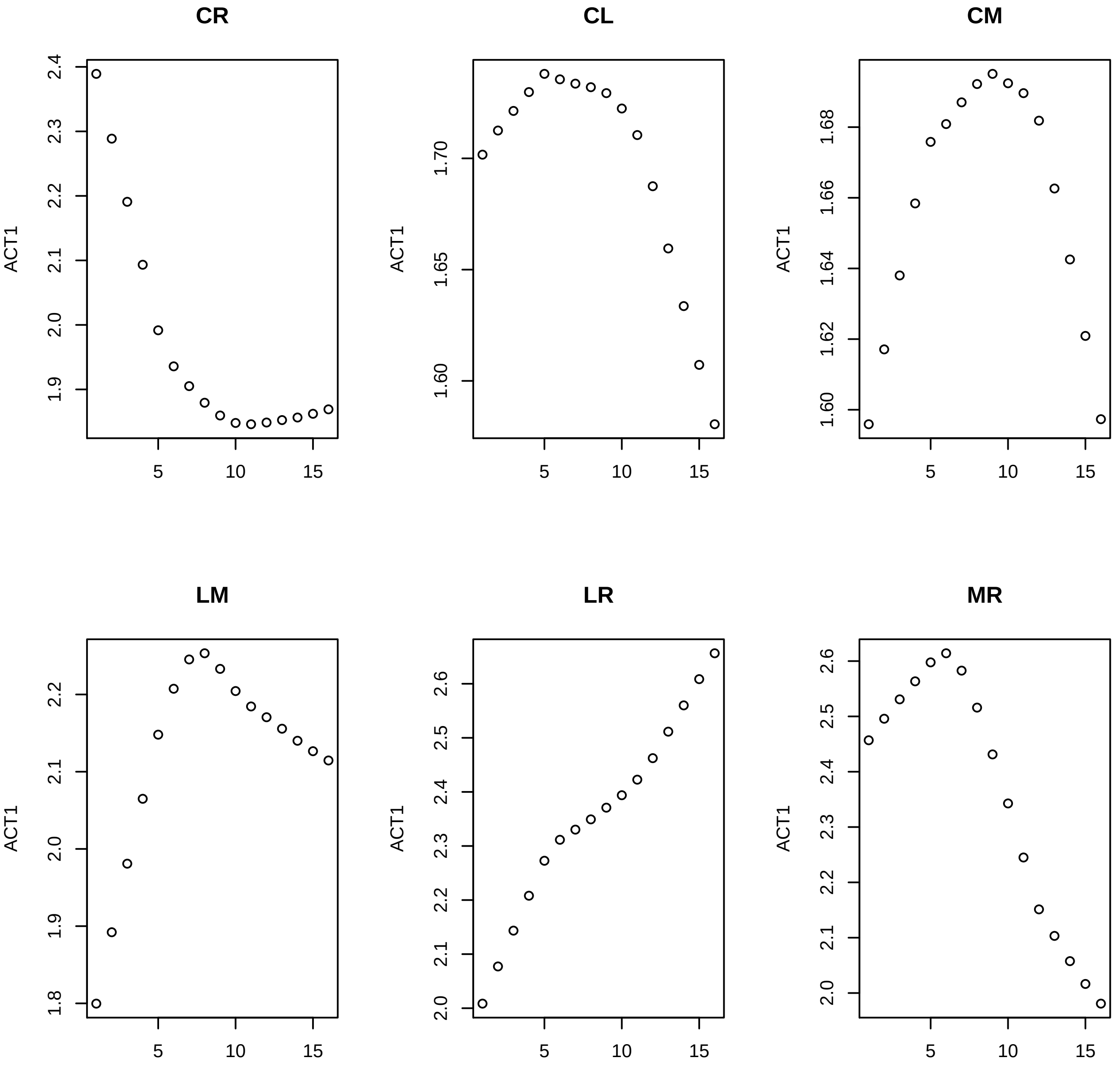
CR
ACT1
2.4
2.3
2.2
2.1
2.0
1.9
5
10
15
CL
ACT1
1.70
1.65
1.60
5
10
15
CM
ACT1
1.68
1.66
1.64
1.62
1.60
5
10
15
LM
ACT1
2.2
2.1
2.0
1.9
1.8
5
10
15
LR
ACT1
2.6
2.5
2.4
2.3
2.2
2.1
2.0
5
10
15
MR
ACT1
2.6
2.5
2.4
2.3
2.2
2.1
2.0
5
10
15

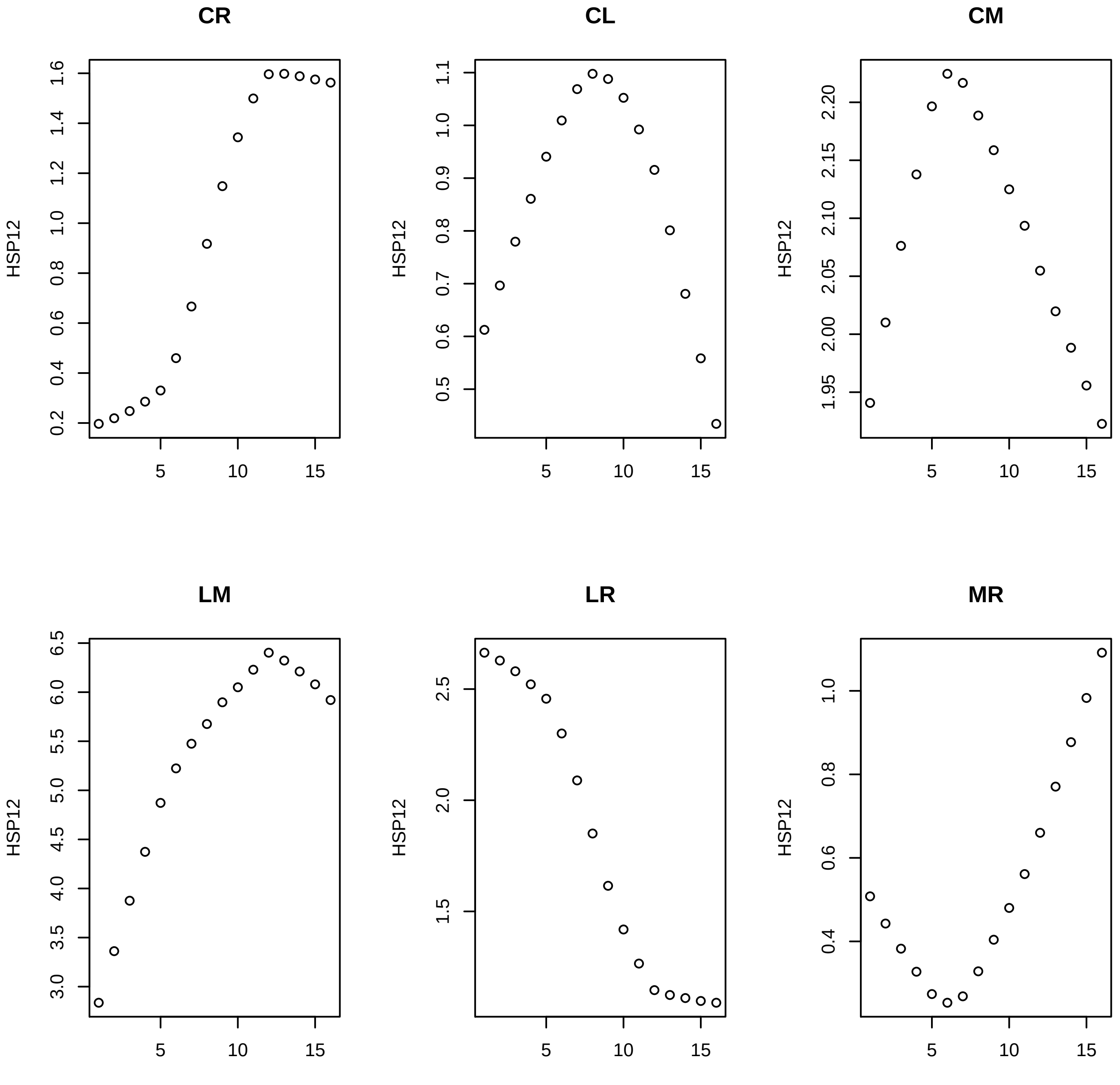
CR
CL
CM
LM
LR
MR
HSP12
5
10
15
0.2
0.4
0.6
0.8
1.0
1.2
1.4
1.6
0.5
0.6
0.7
0.8
0.9
1.0
1.1
1.95
2.00
2.05
2.10
2.15
2.20
3.0
3.5
4.0
4.5
5.0
5.5
6.0
6.5
1.5
2.0
2.5

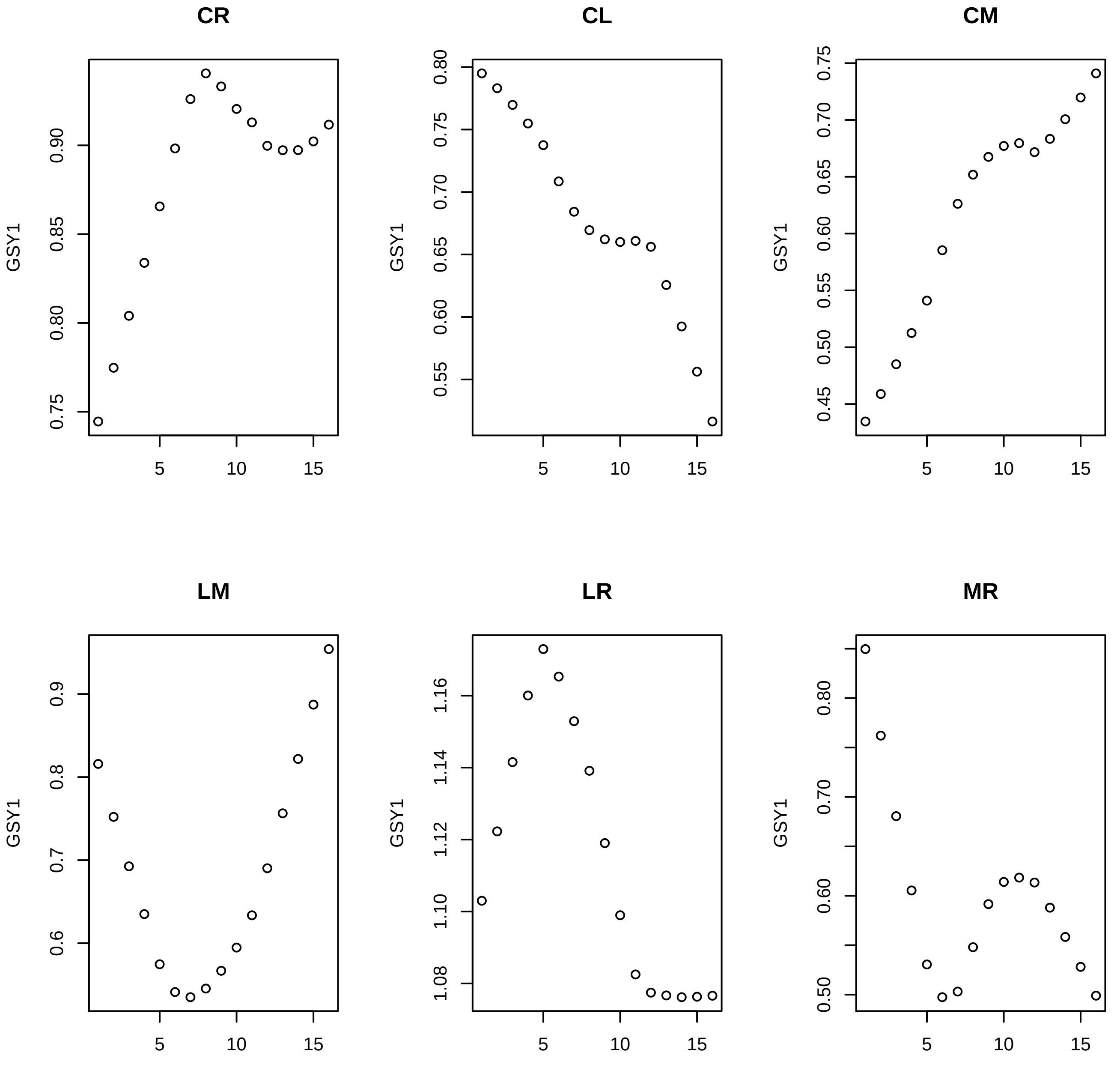
CR
GSY1
0.75
0.80
0.85
0.90
5
10
15
CL
GSY1
0.55
0.60
0.65
0.70
0.75
0.80
5
10
15
CM
GSY1
0.45
0.50
0.55
0.60
0.65
0.70
0.75
5
10
15
LM
GSY1
0.6
0.7
0.8
0.9
5
10
15
LR
GSY1
1.08
1.10
1.12
1.14
1.16
5
10
15
MR
GSY1
0.50
0.60
0.70
0.80
5
10
15

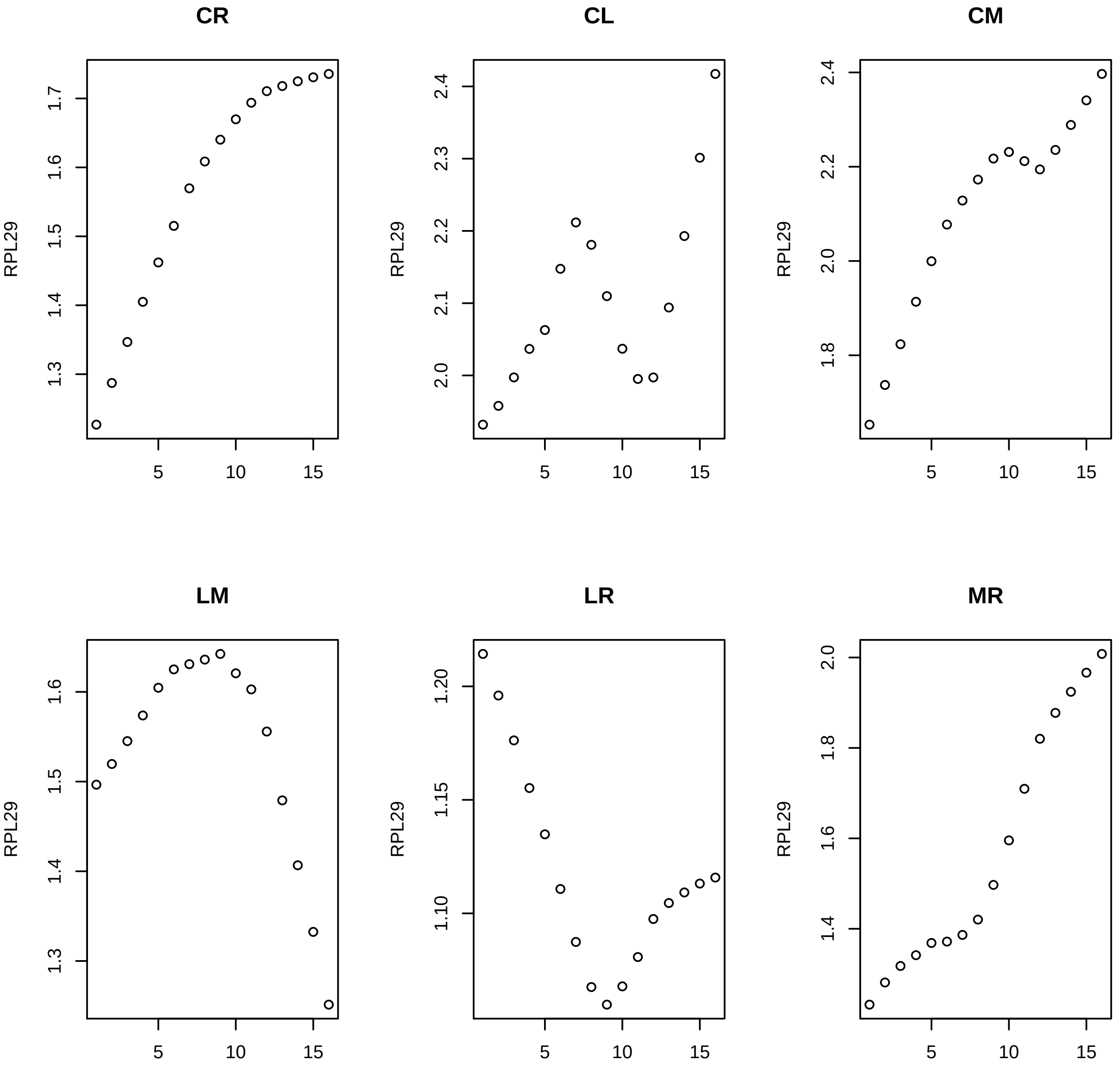

CR
CL
CM
LM
LR
MR
RPL29

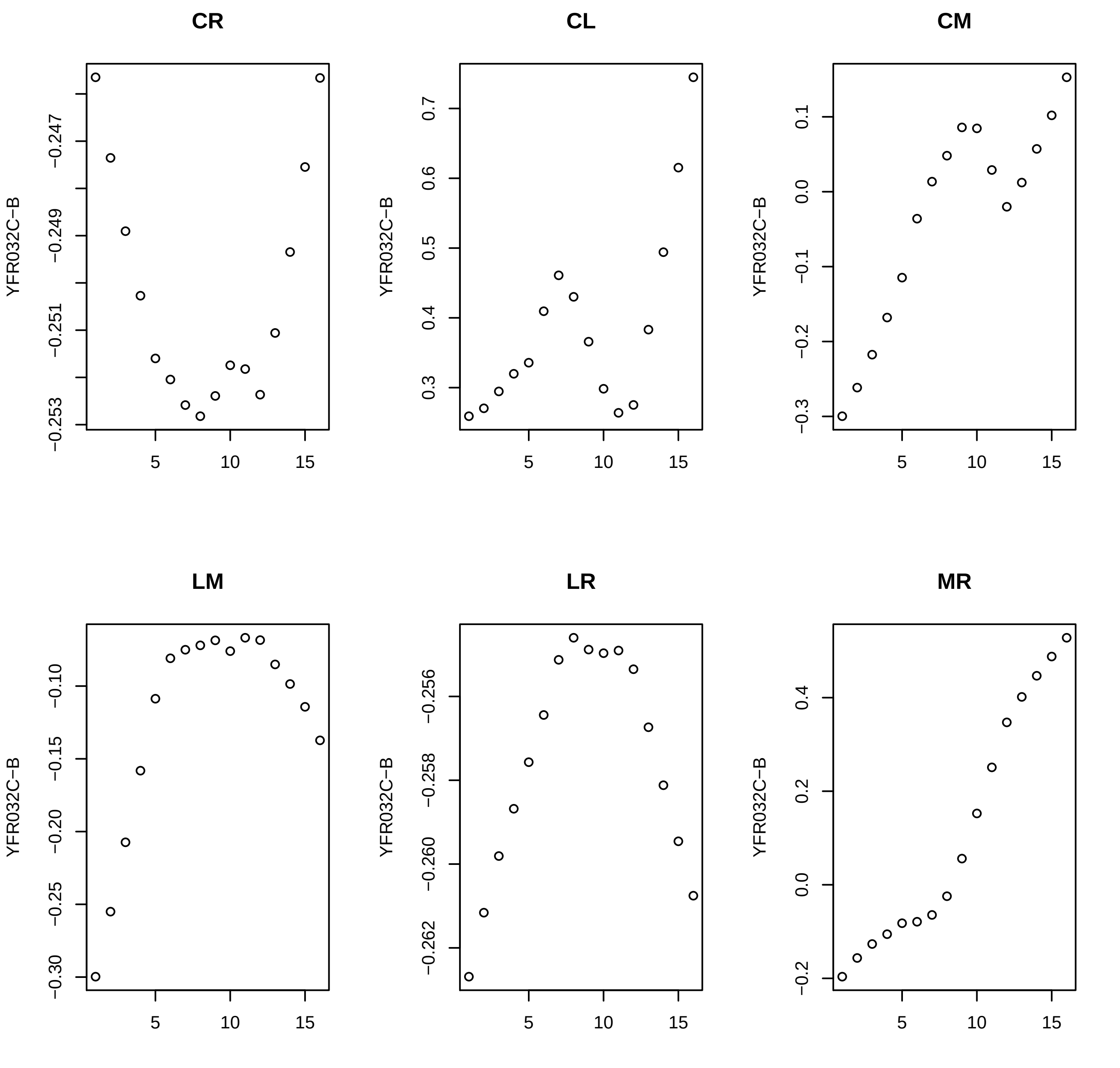

CR
CL
CM
LM
LR
MR
YFR032C−B
−0.247
−0.249
−0.251
−0.253
0.7
0.6
0.5
0.4
0.3
0.1
0.0
−0.1
−0.2
−0.3
−0.10
−0.15
−0.20
−0.25
−0.30
−0.256
−0.258
−0.260
−0.262
0.4
0.2
0.0
−0.2
5
10
15

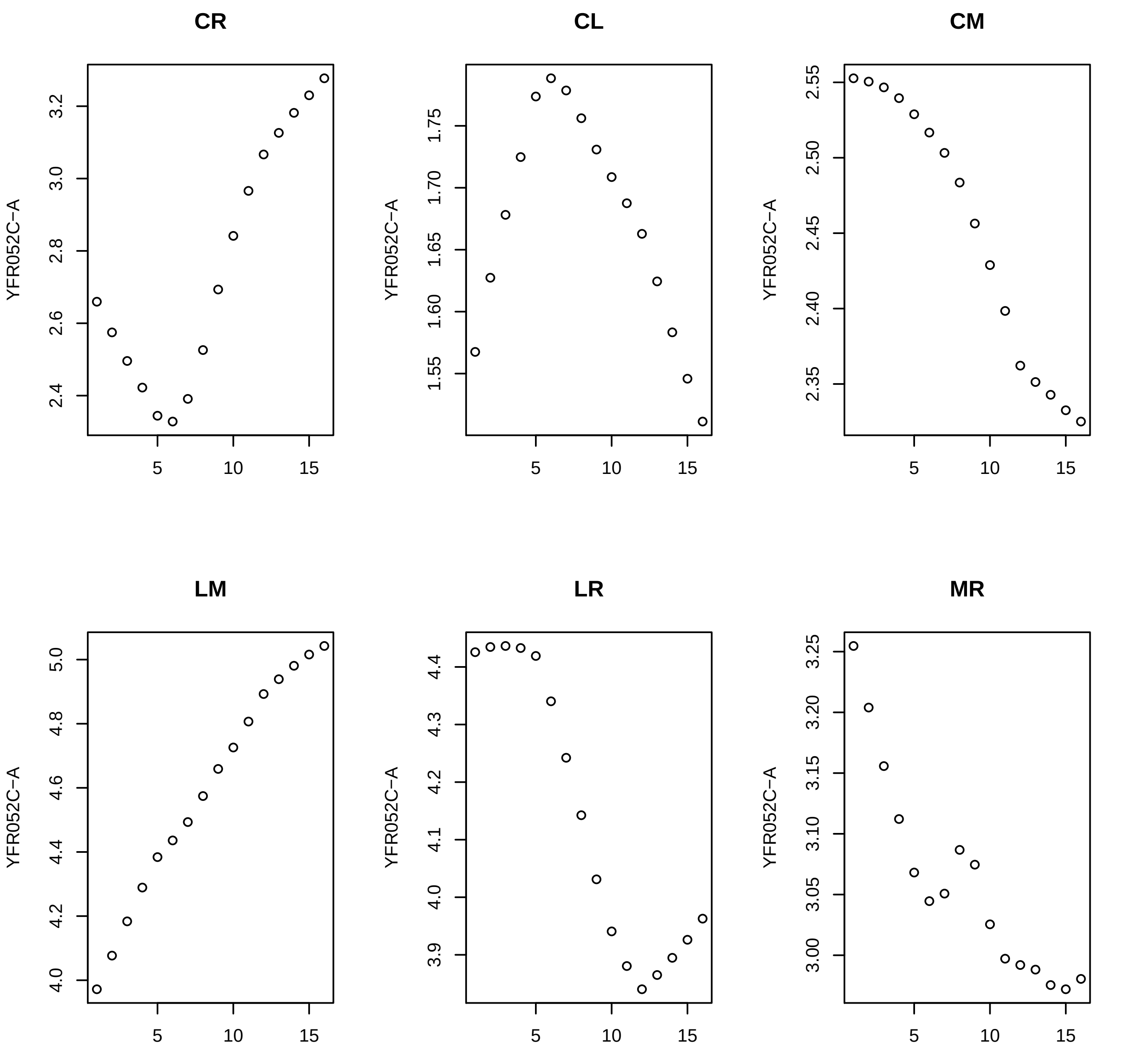
CR
CL
CM
LM
LR
MR
YFR052C−A
5
10
15

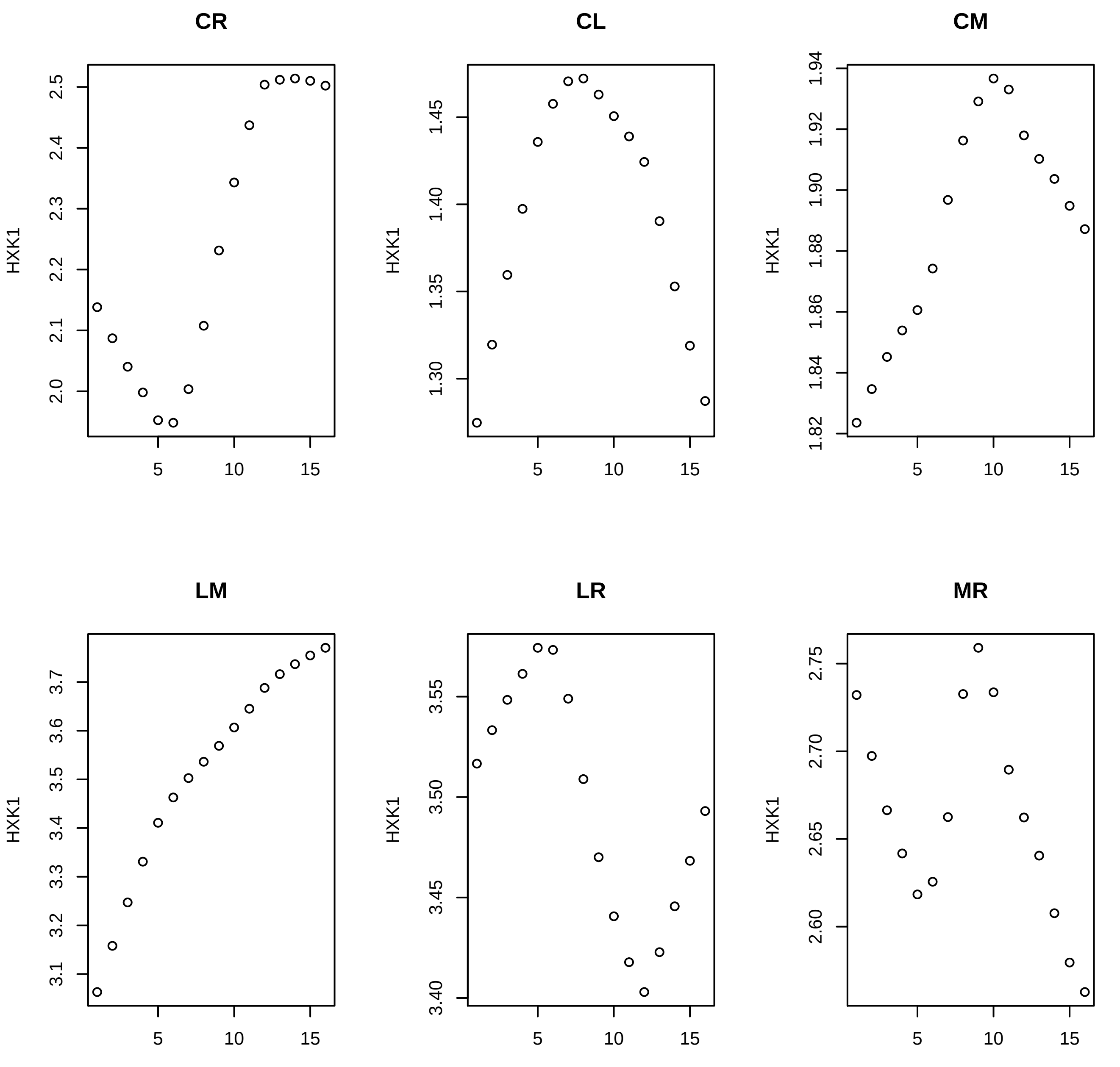
CR
CL
CM
LM
LR
MR
HXK1

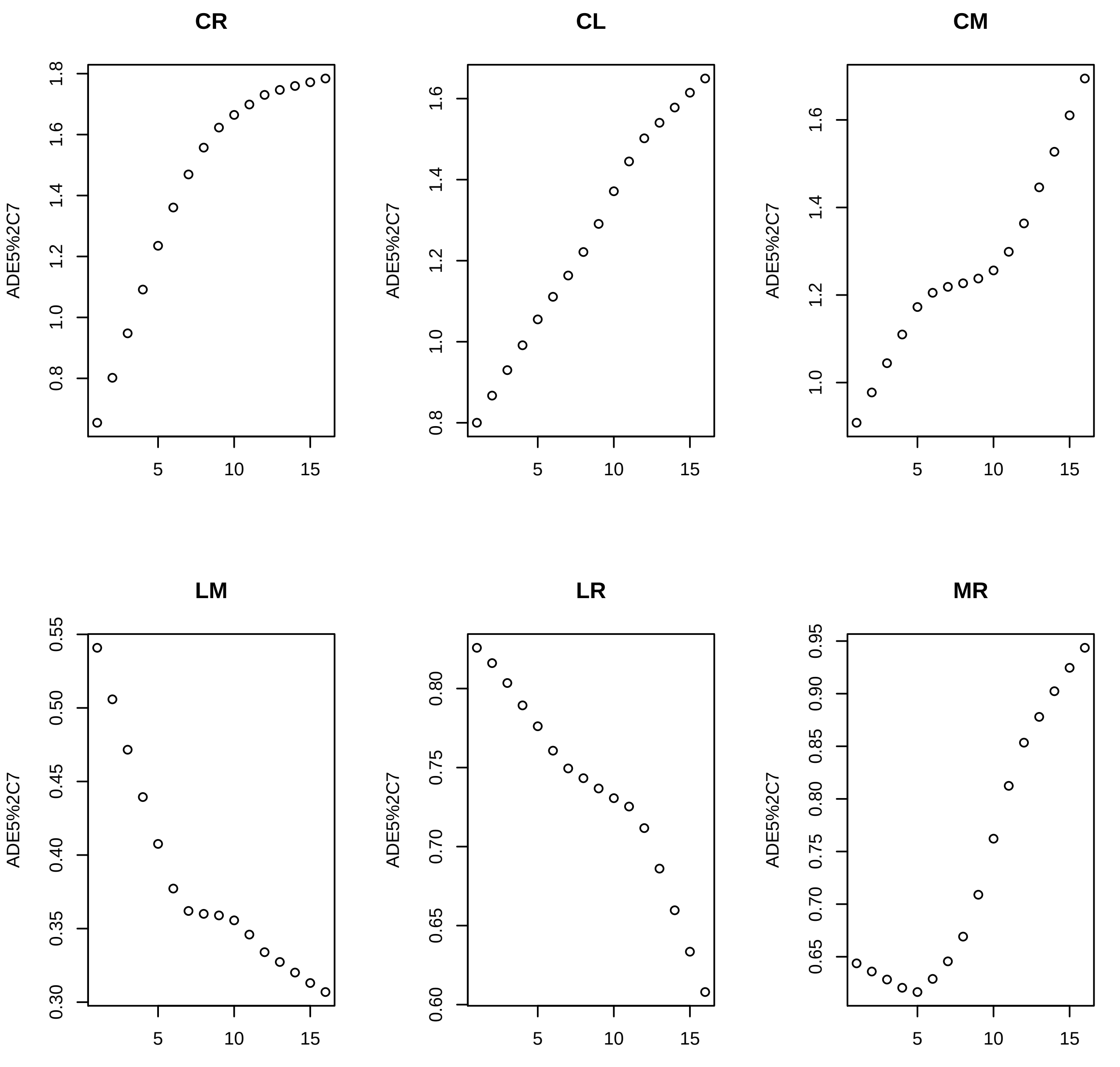
CR
CL
CM
LM
LR
MR
ADE5%2C7

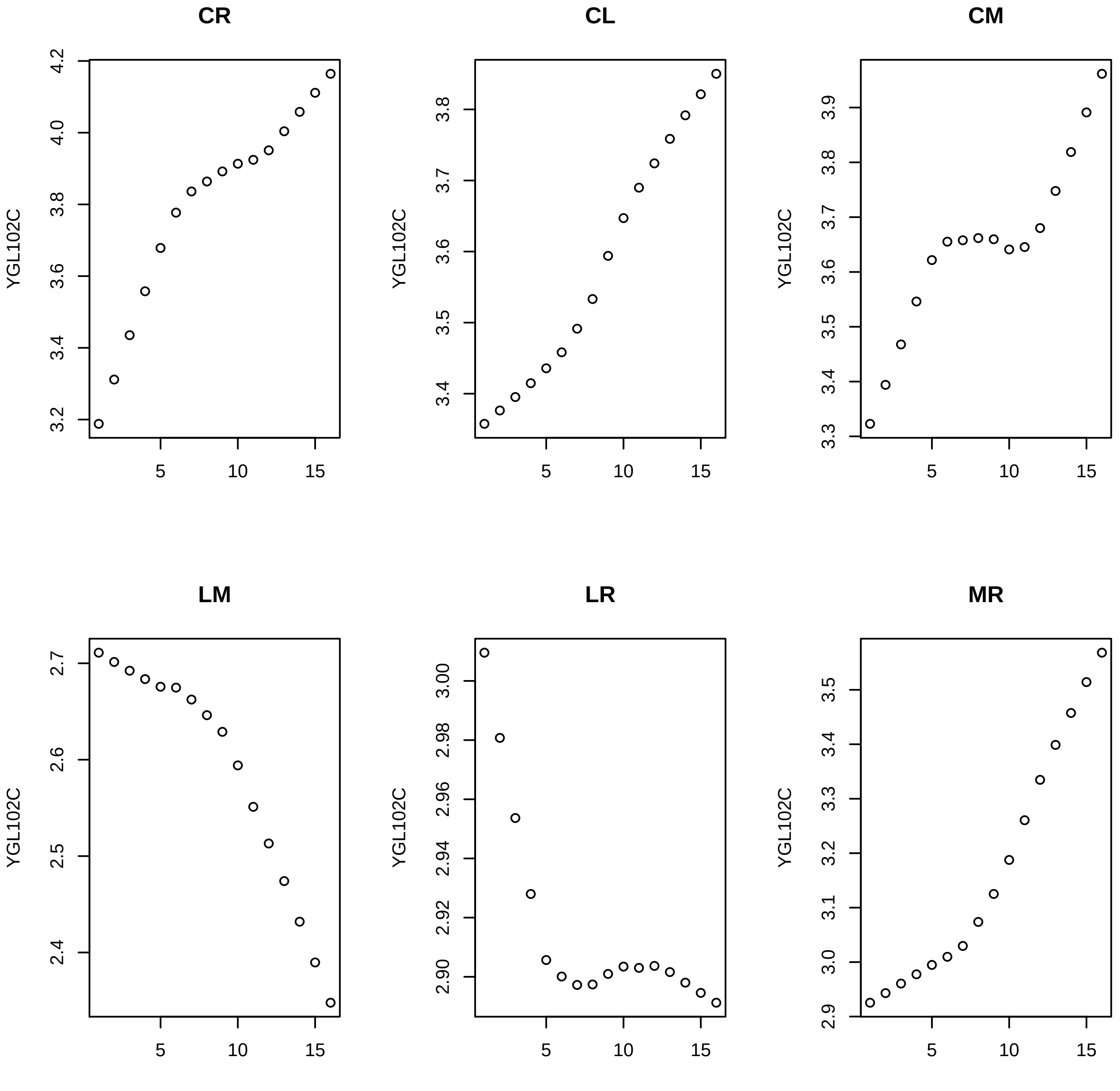
CR
CL
CM
LM
LR
MR
YGL102C
5
10
15
4.2
4.0
3.8
3.6
3.4
3.2
3.8
3.7
3.6
3.5
3.4
3.9
3.3
2.7
2.6
2.5
2.4
3.00
2.98
2.96
2.94
2.92
2.90
3.1
3.0
2.9

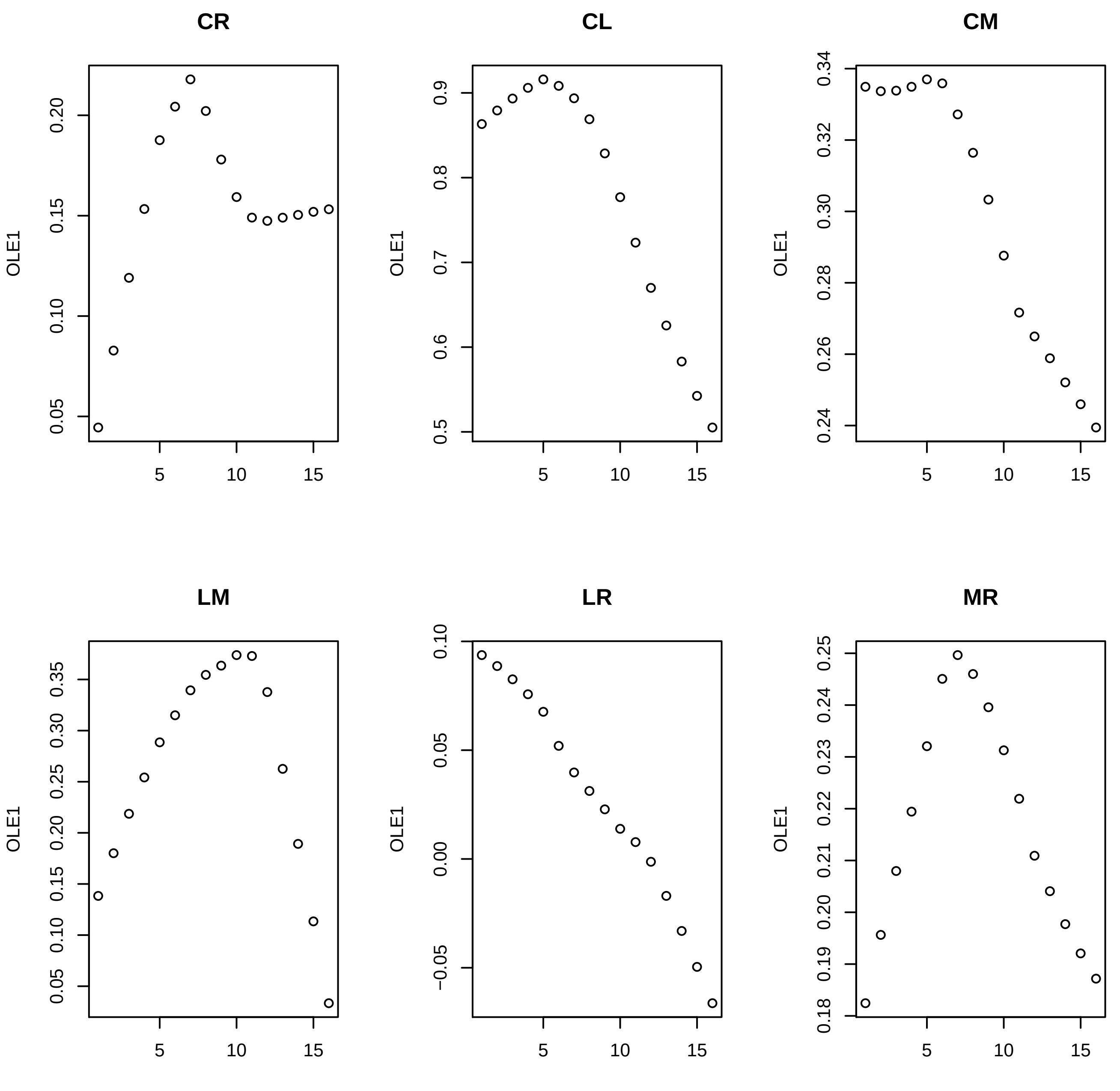
CR
CL
CM
LM
LR
MR
OLE1
OLE1
OLE1
OLE1
OLE1
OLE1
0.05
0.10
0.15
0.20
0.5
0.6
0.7
0.8
0.9
0.24
0.26
0.28
0.30
0.32
0.34
0.05
0.10
0.15
0.20
0.25
0.30
0.35
−0.05
0.00
0.05
0.10
0.18
0.19
0.20
0.21
0.22
0.23
0.24
0.25
5
10
15

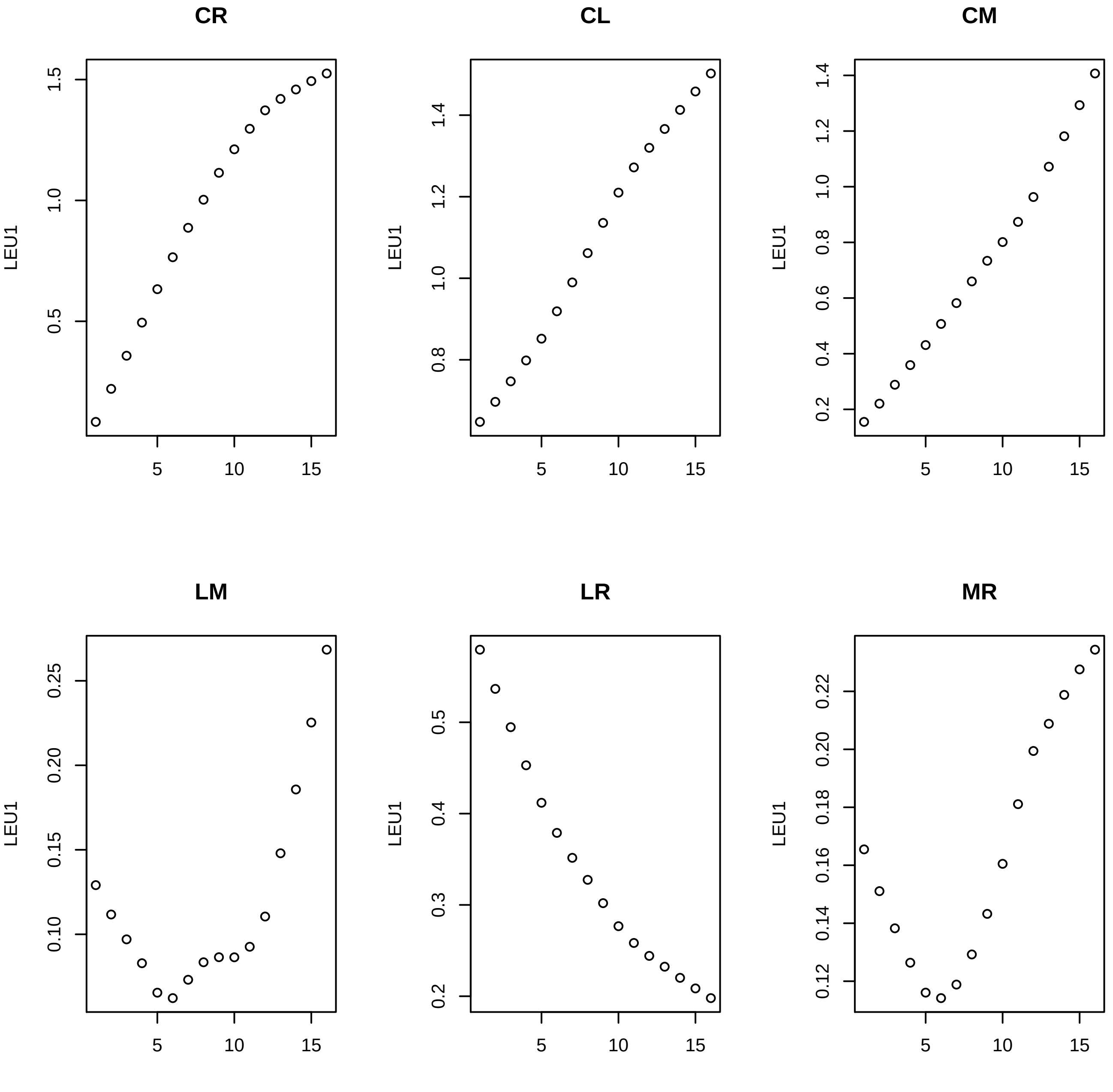

CR
CL
CM
LM
LR
MR
LEU1

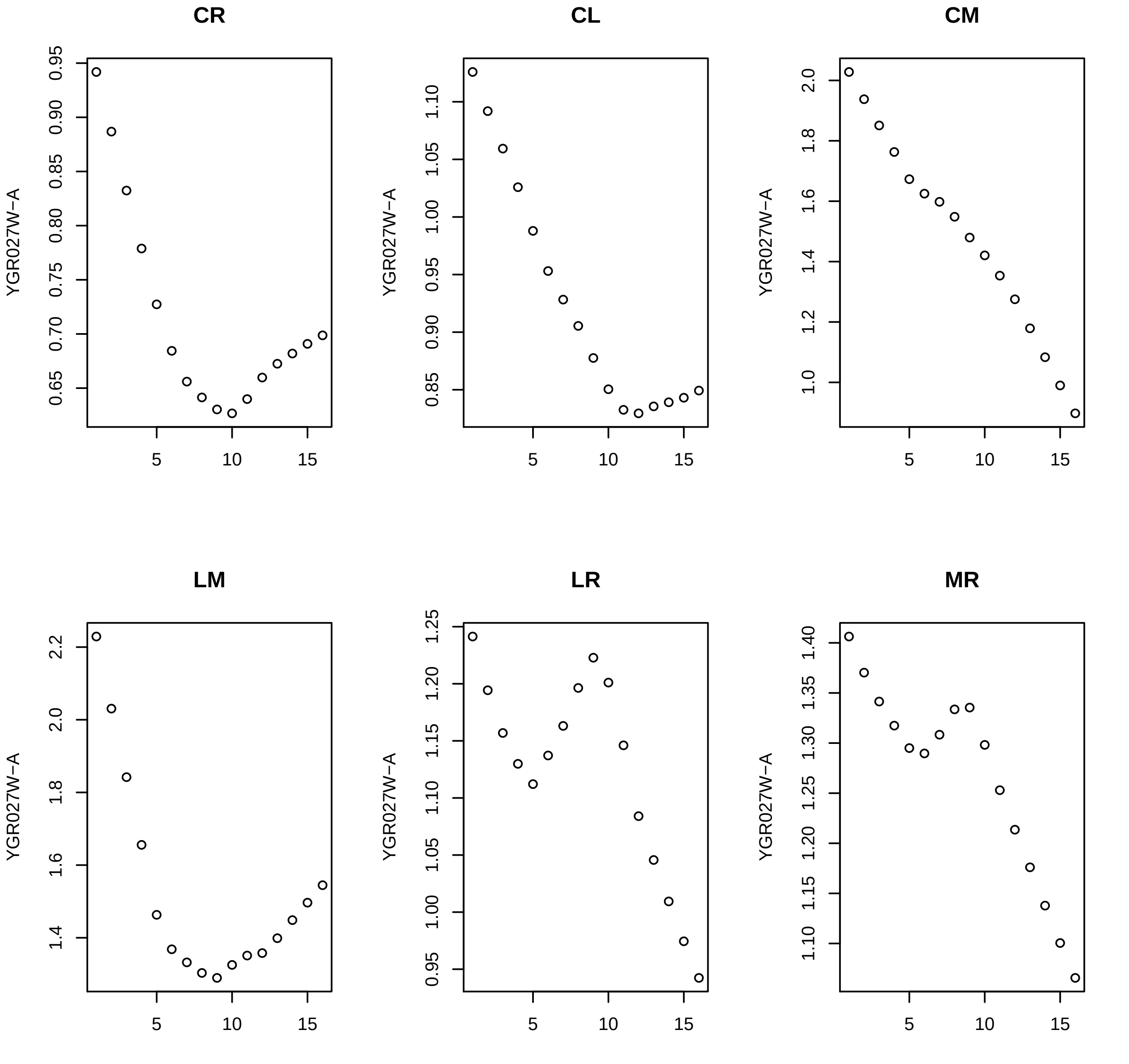

CR
CL
CM
LM
LR
MR
YGR027W−A

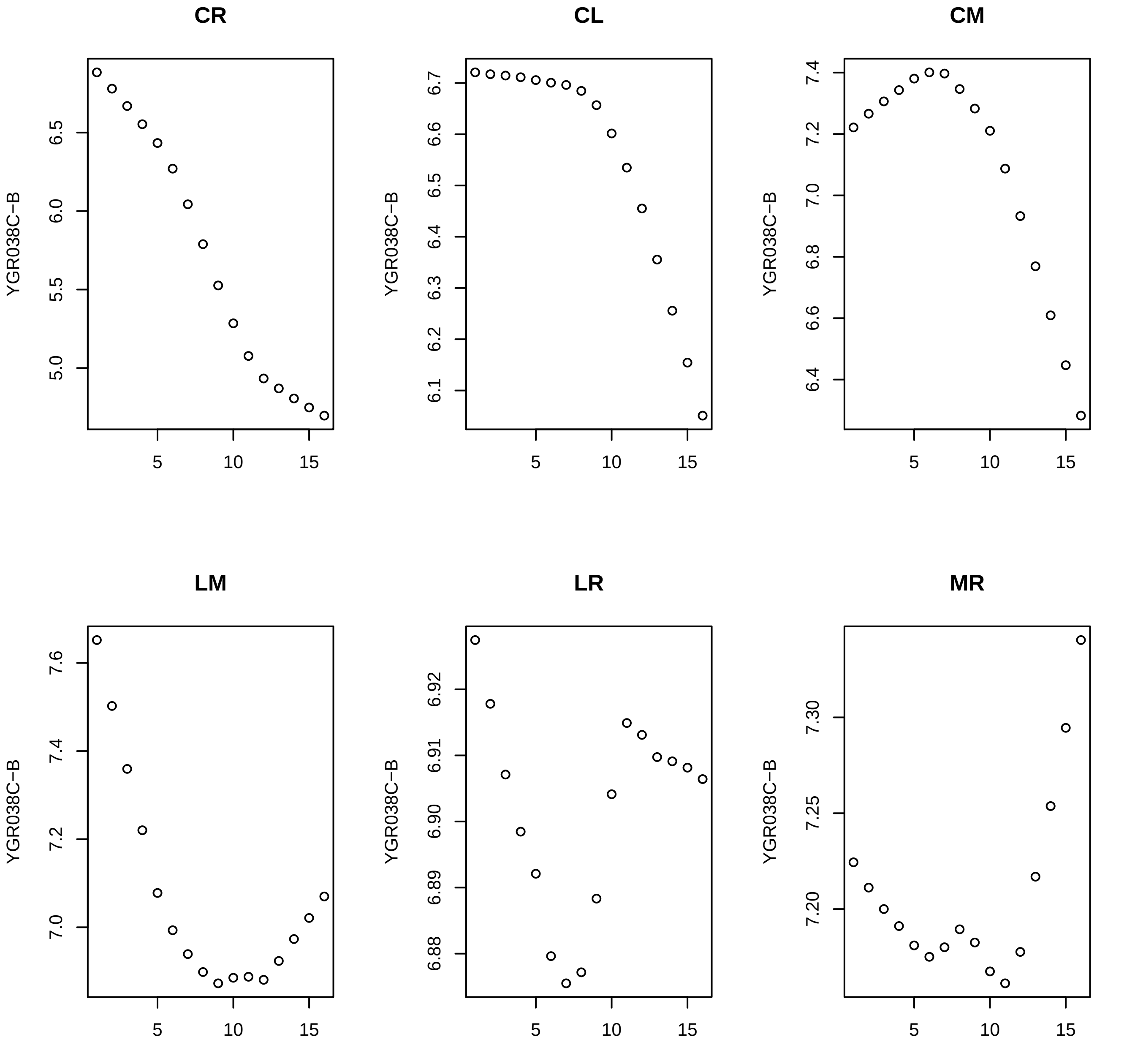

CR
CL
CM
LM
LR
MR
YGR038C−B

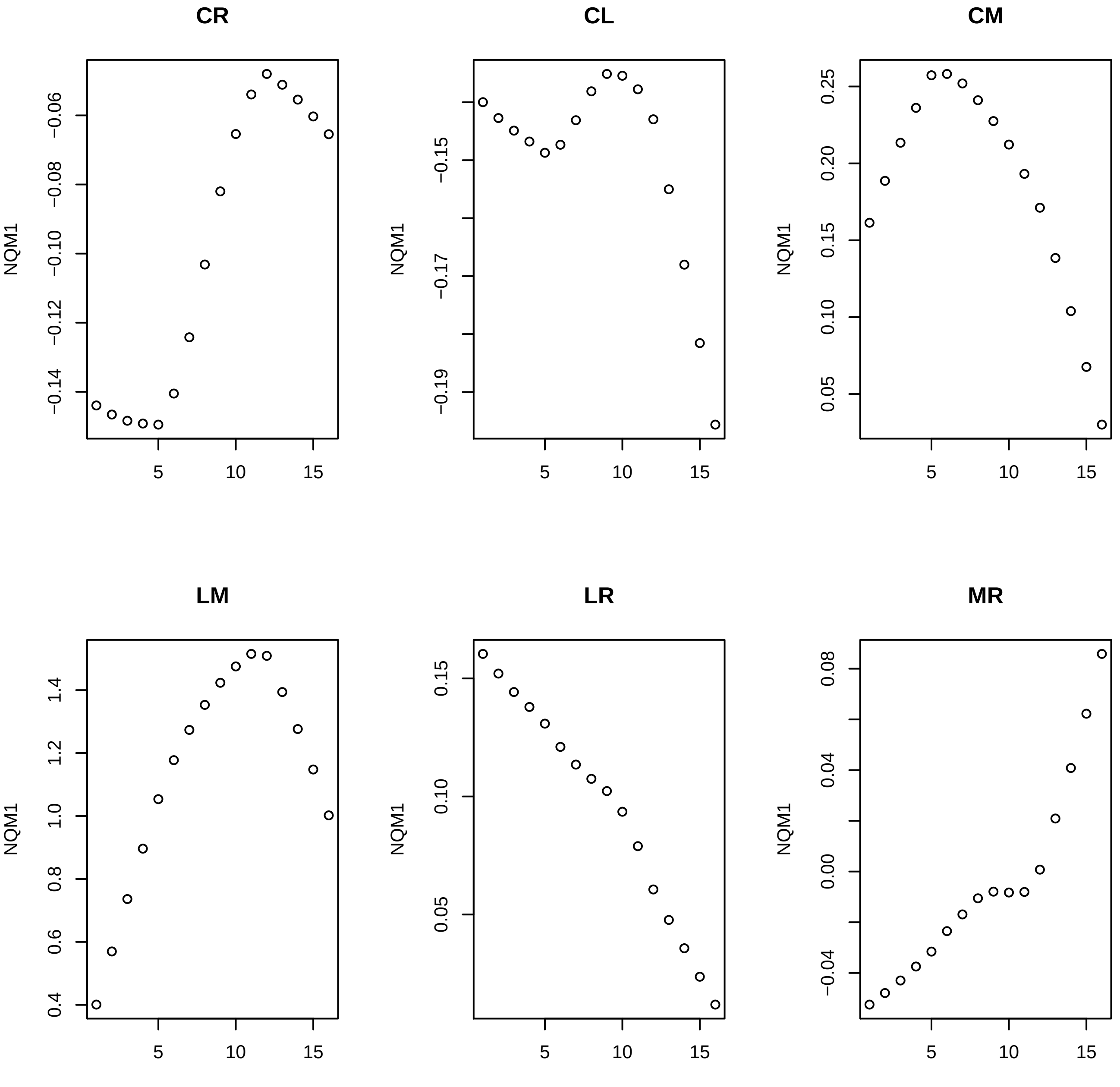
CR
CL
CM
LM
LR
MR
NQM1
−0.06
−0.08
−0.10
−0.12
−0.14
−0.15
−0.17
−0.19
0.25
0.20
0.15
0.10
0.05
1.4
1.2
1.0
0.8
0.6
0.4
0.15
0.10
0.05
0.08
0.04
0.00
−0.04
5
10
15

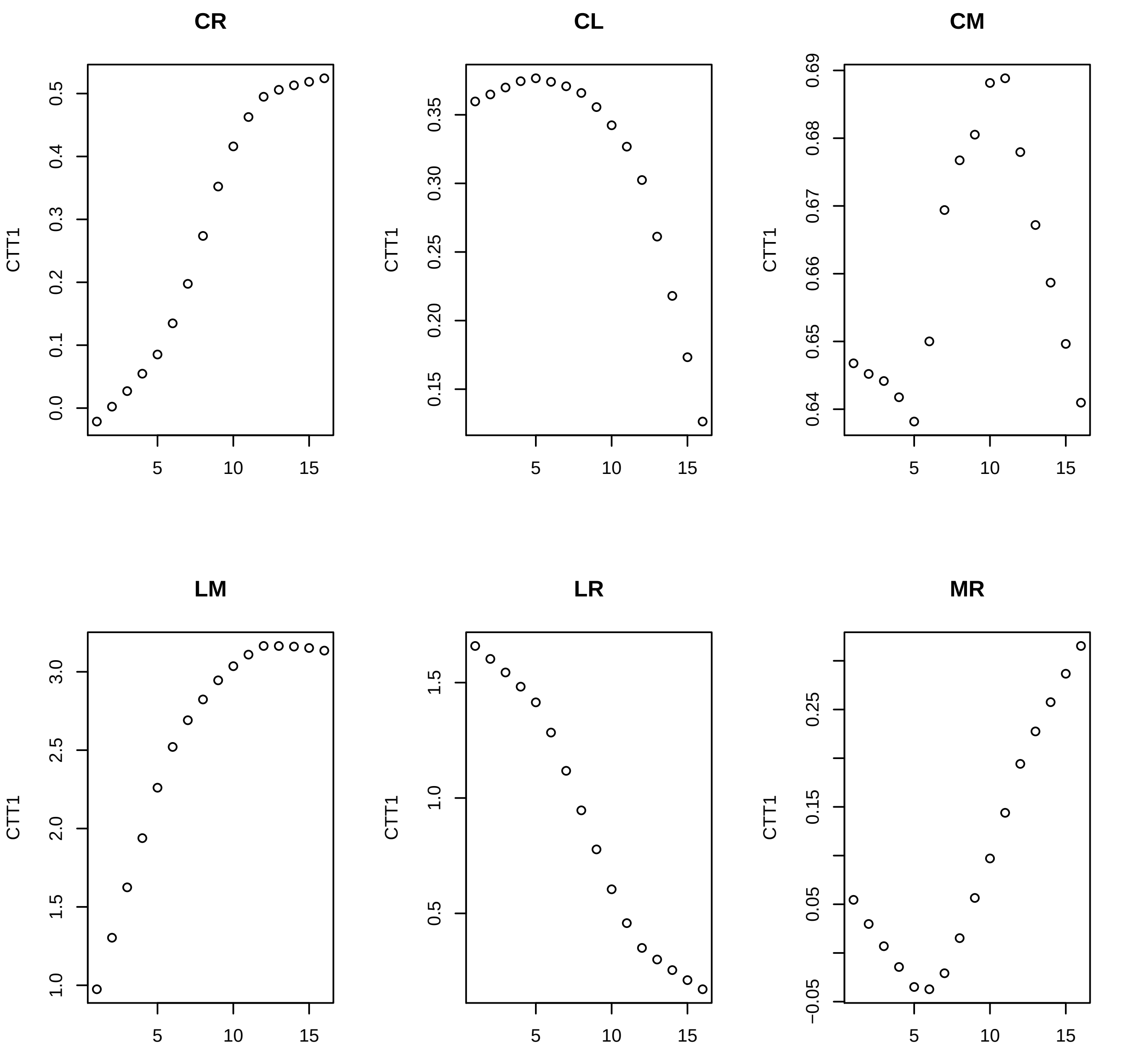
CR
CL
CM
LM
LR
MR
CTT1
0.0
0.1
0.2
0.3
0.4
0.5
0.15
0.20
0.25
0.30
0.35
0.64
0.65
0.66
0.67
0.68
0.69
1.0
1.5
2.0
2.5
3.0
0.5
1.0
1.5
−0.05
0.05
0.15
0.25
5
10
15

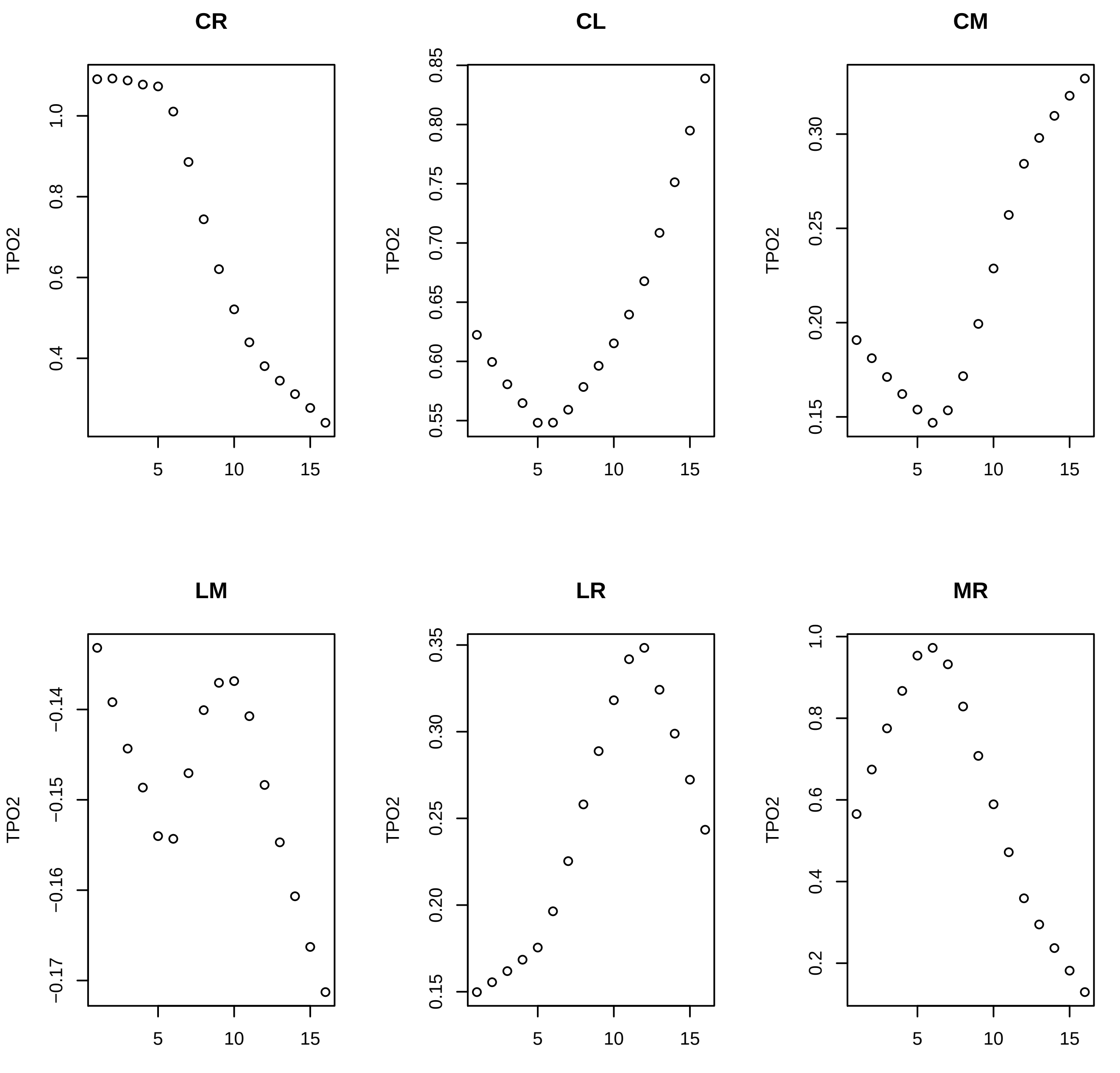

CR
CL
CM
LM
LR
MR
TPO2

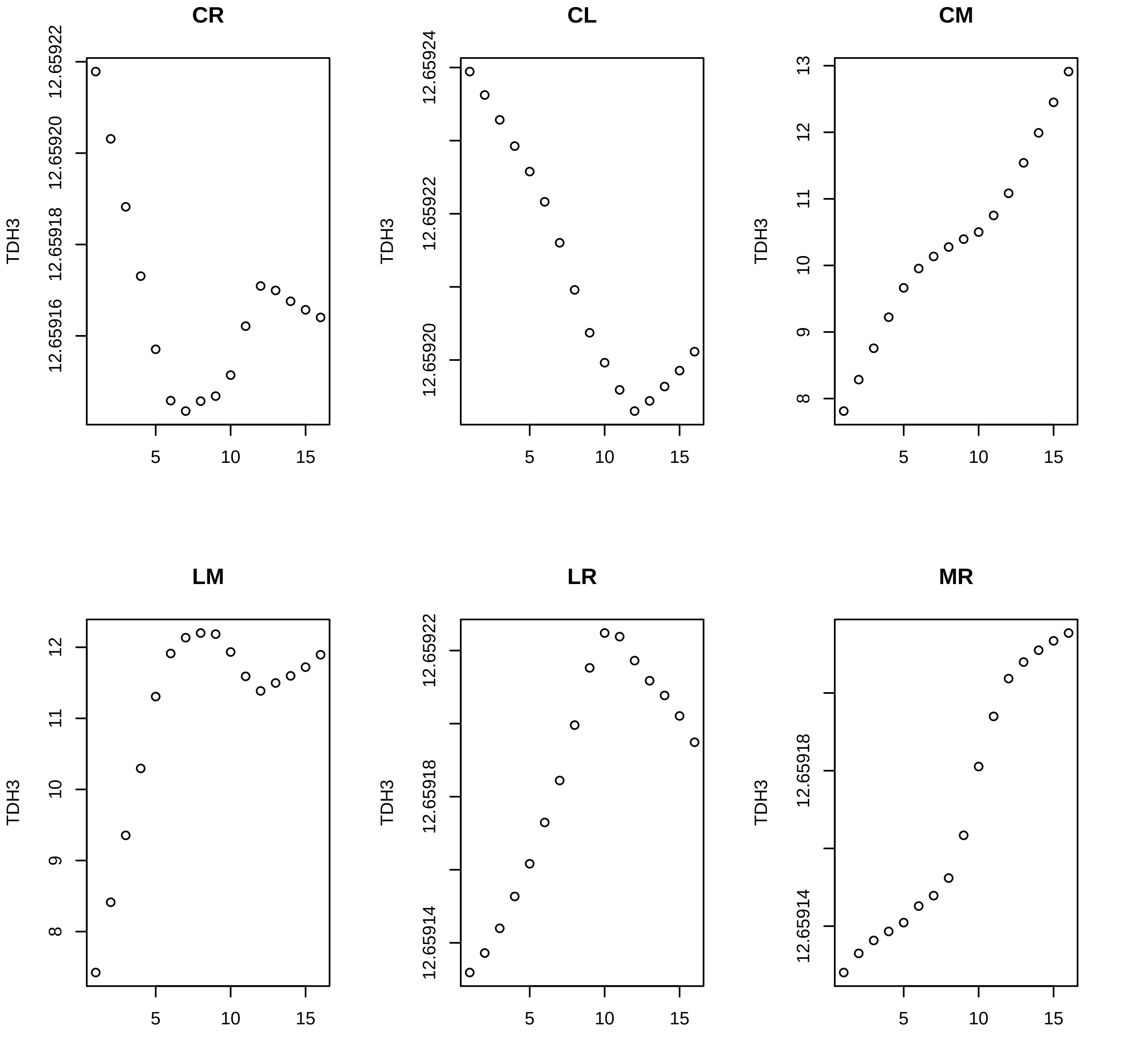

CR
CL
CM
LM
LR
MR
TDH3
12.65922
12.65920
12.65918
12.65916
12.65924
13
12
11
10
9
8
12.65914
5
10
15

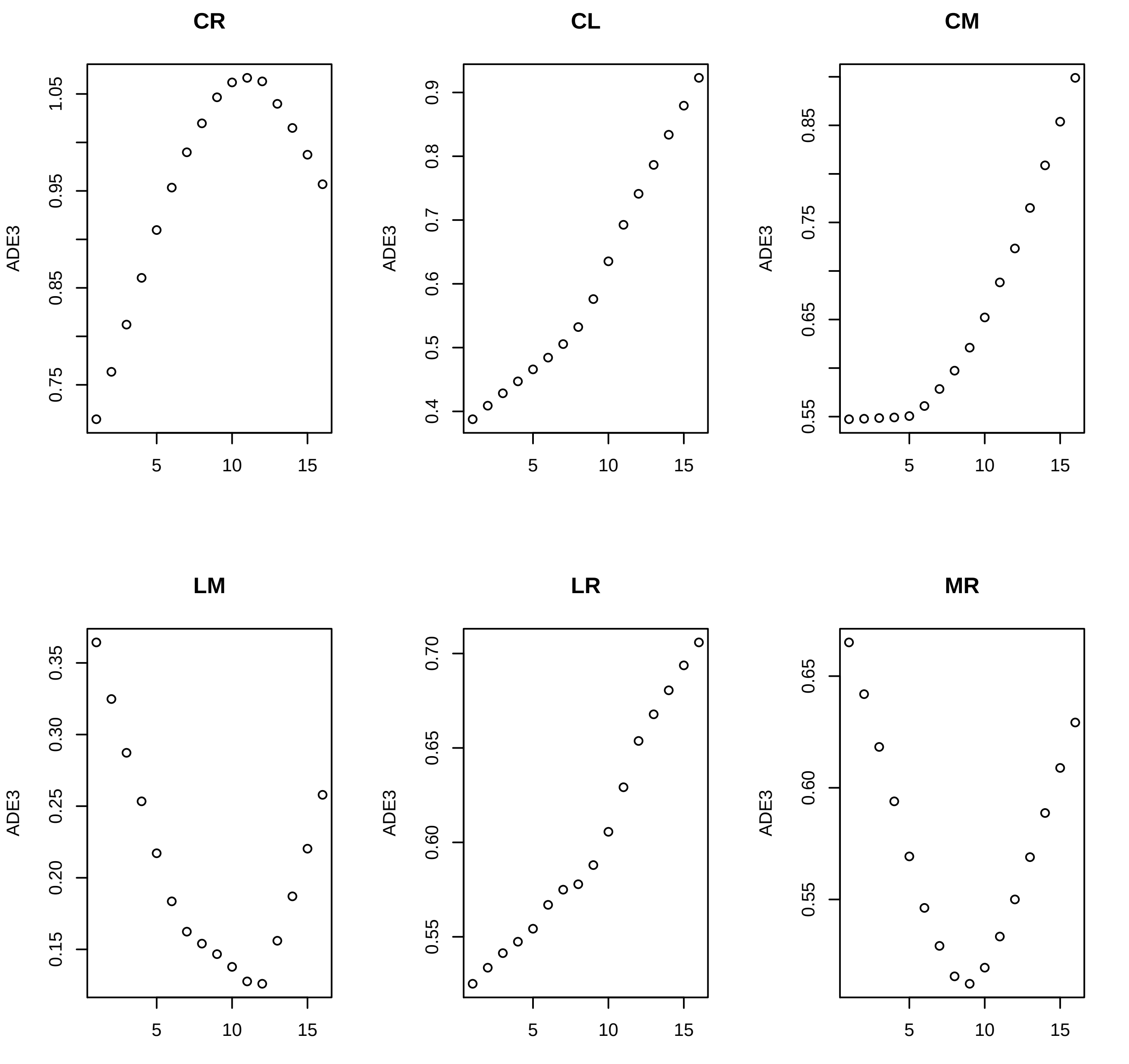
CR
CL
CM
LM
LR
MR
ADE3
5
10
15
0.75
0.85
0.95
1.05
0.4
0.5
0.6
0.7
0.8
0.9
0.55
0.65
0.75
0.85
0.15
0.20
0.25
0.30
0.35
0.55
0.60
0.65
0.70

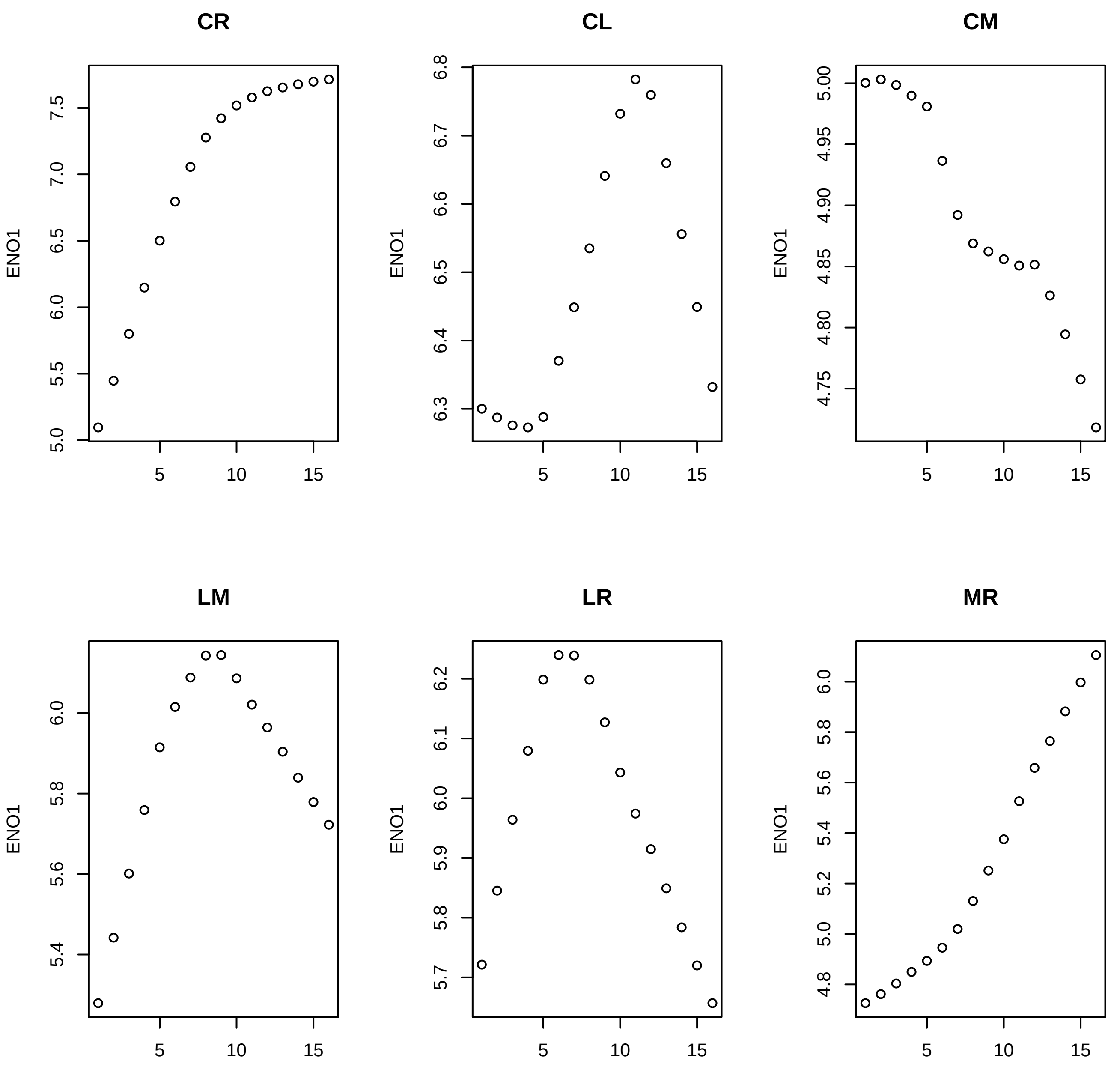

CR
CL
CM
LM
LR
MR
ENO1
5
10
15
5.0
5.5
6.0
6.5
7.0
7.5
6.3
6.4
6.5
6.6
6.7
6.8
4.75
4.80
4.85
4.90
4.95
5.00
5.4
5.6
5.8
6.0
5.7
5.8
5.9
6.0
6.1
6.2
4.8
5.0
5.2
5.4
5.6
5.8
6.0

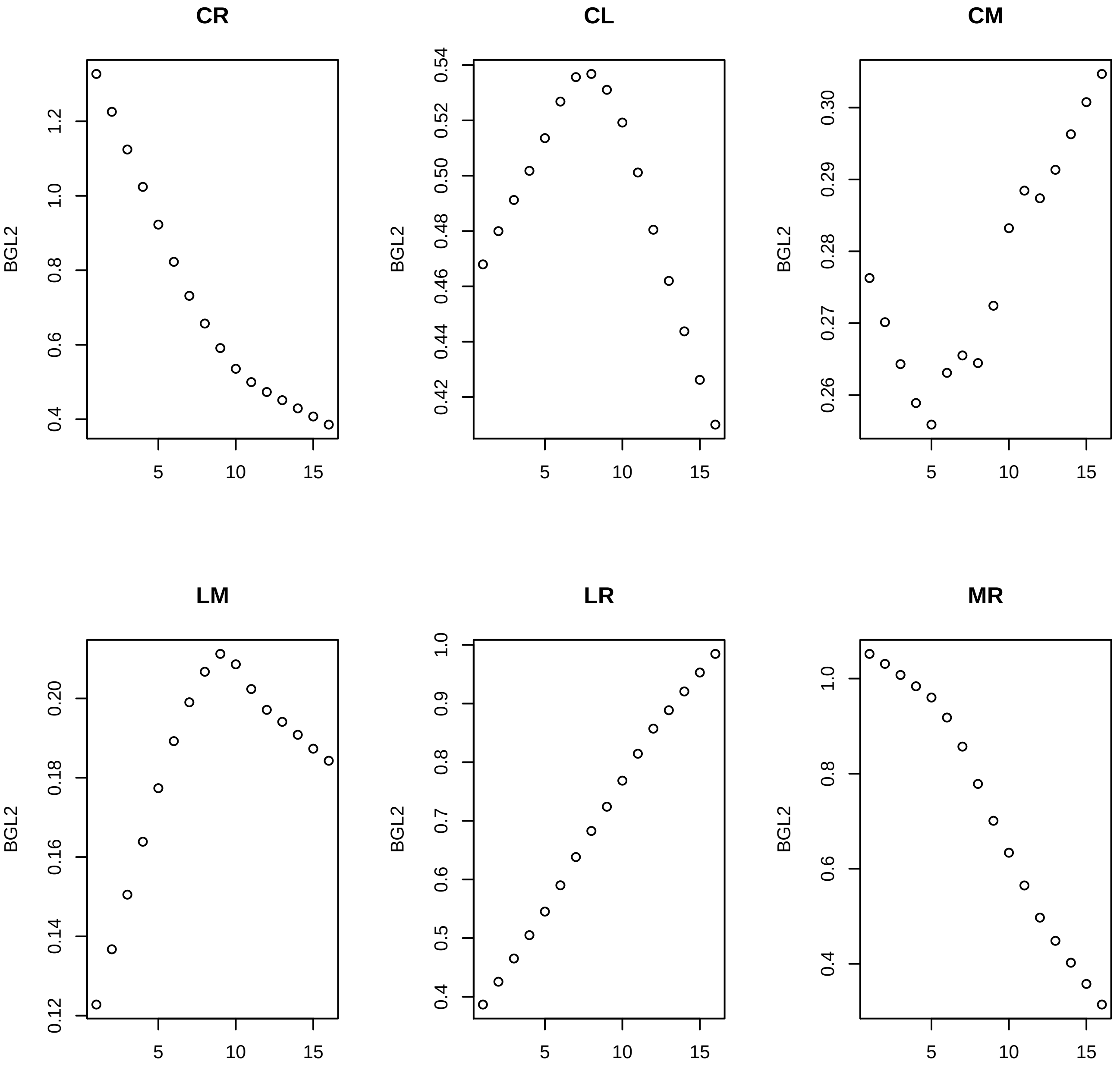
CR
CL
CM
LM
LR
MR
BGL2
5
10
15
1.2
1.0
0.8
0.6
0.4
0.54
0.52
0.50
0.48
0.46
0.44
0.42
0.30
0.29
0.28
0.27
0.26
0.20
0.18
0.16
0.14
0.12
0.9
0.7
0.5

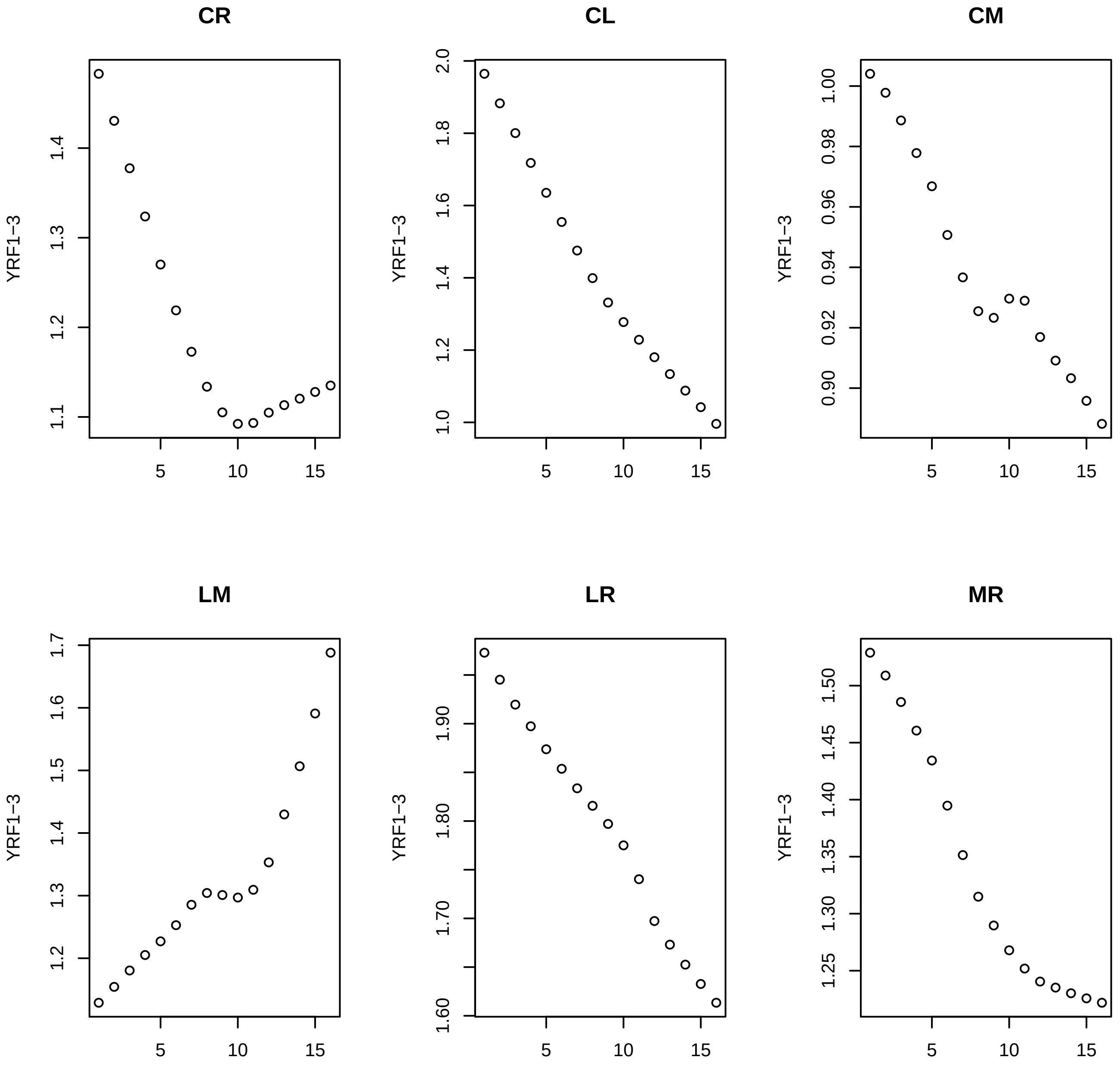

CR
CL
CM
LM
LR
MR
YRF1−3

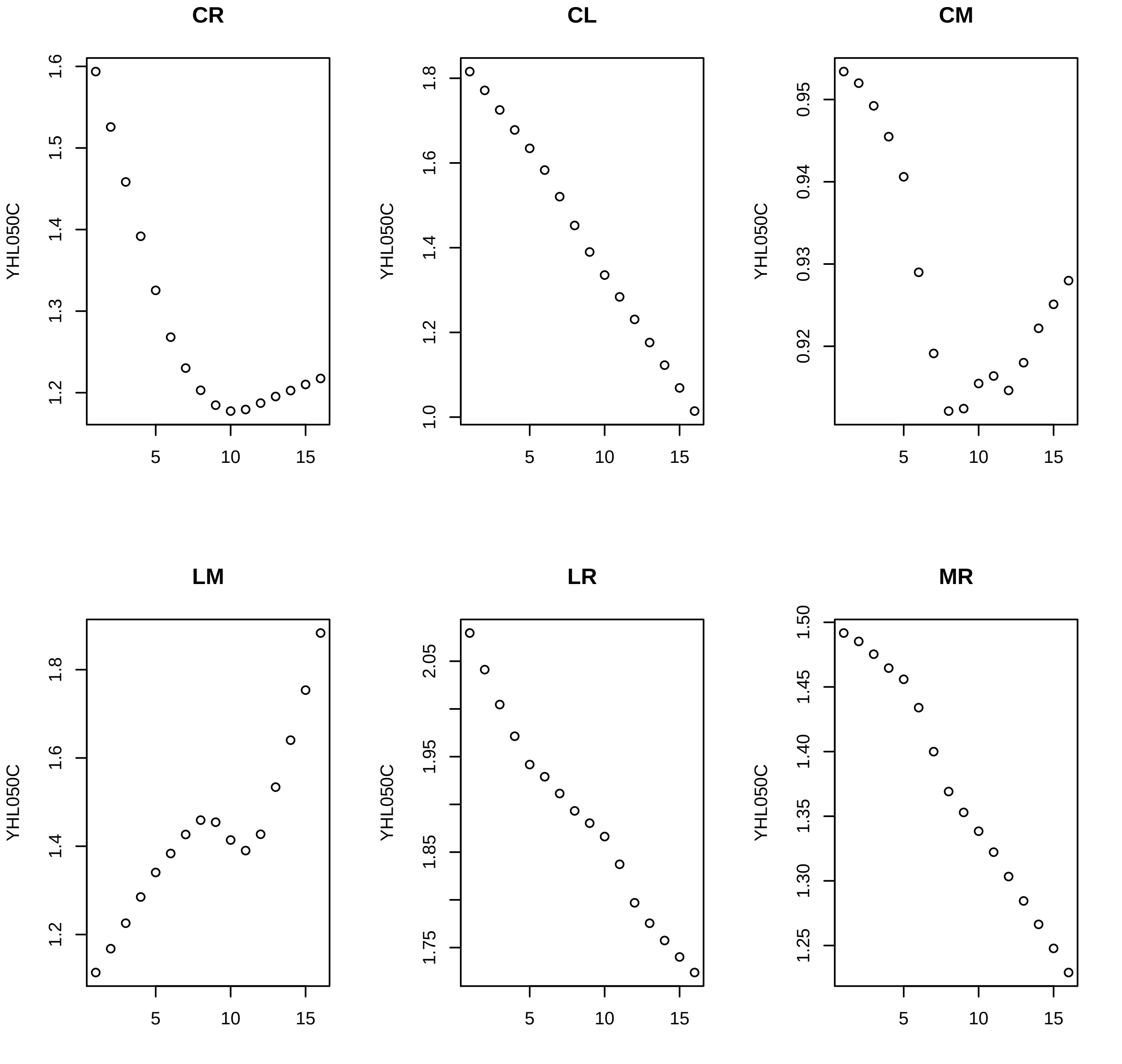
CR
CL
CM
LM
LR
MR
YHL050C

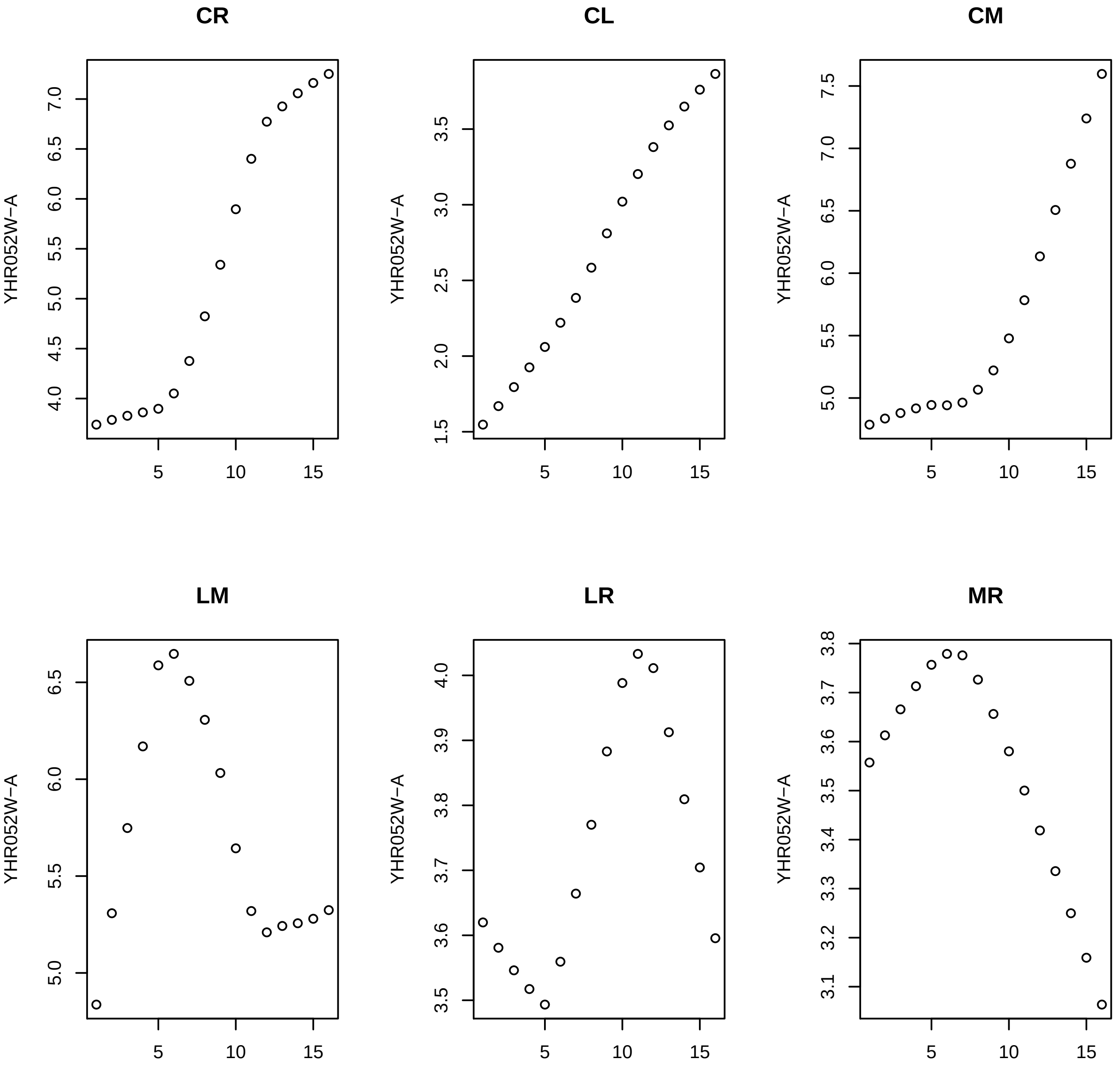
CR
CL
CM
LM
LR
MR
YHR052W−A

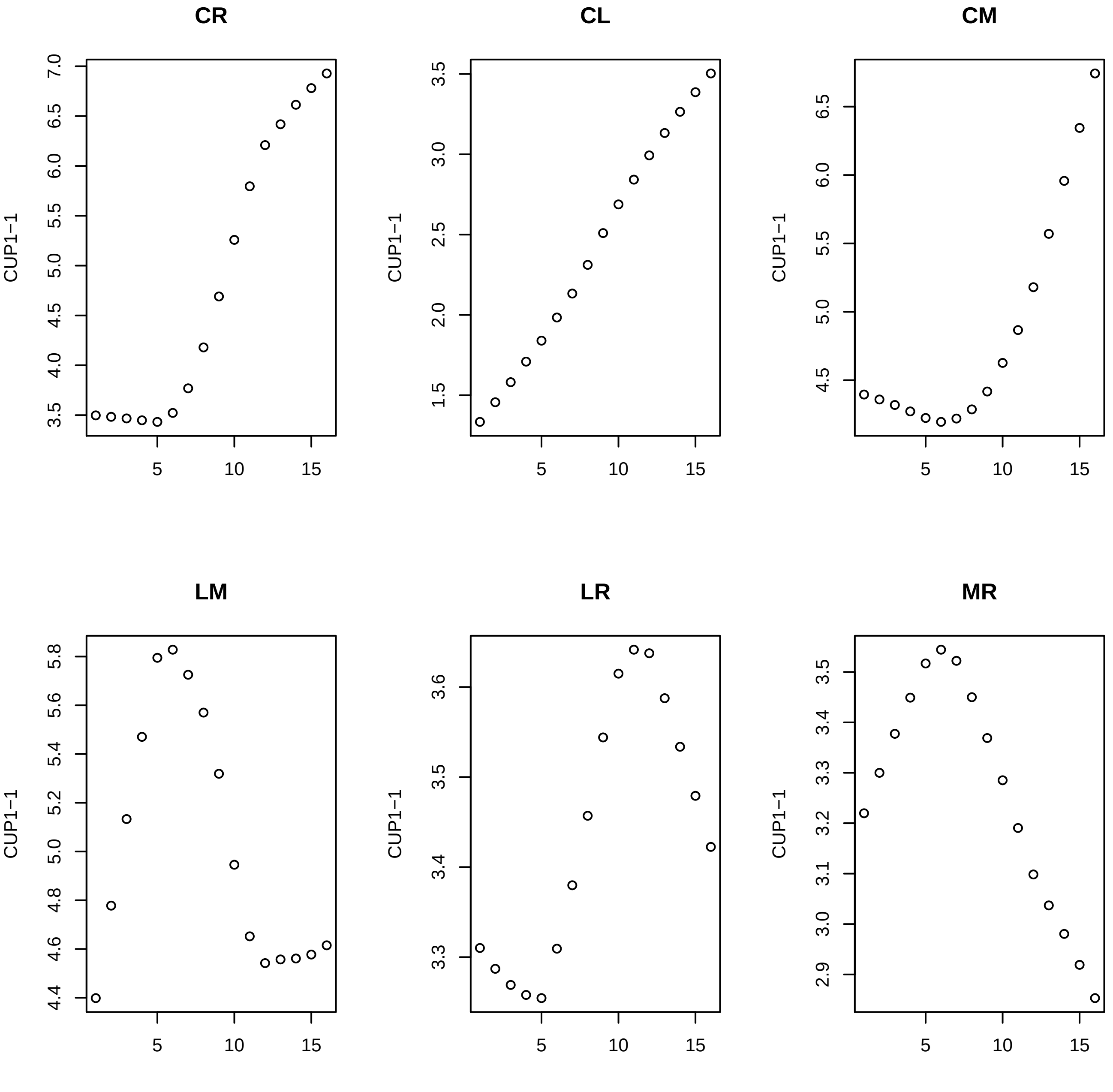

CR
CL
CM
LM
LR
MR
CUP1−1

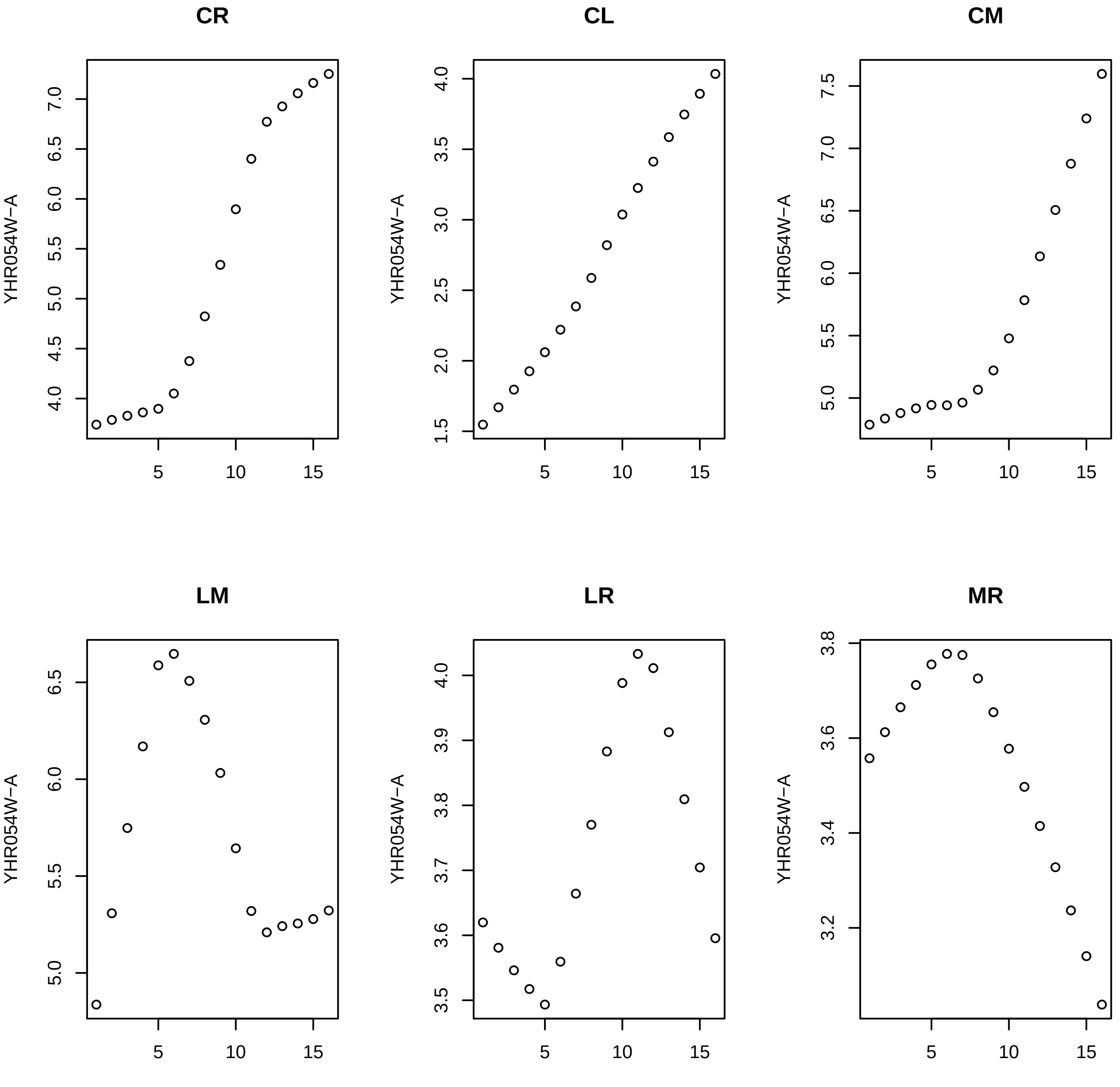
CR
CL
CM
LM
LR
MR
YHR054W−A

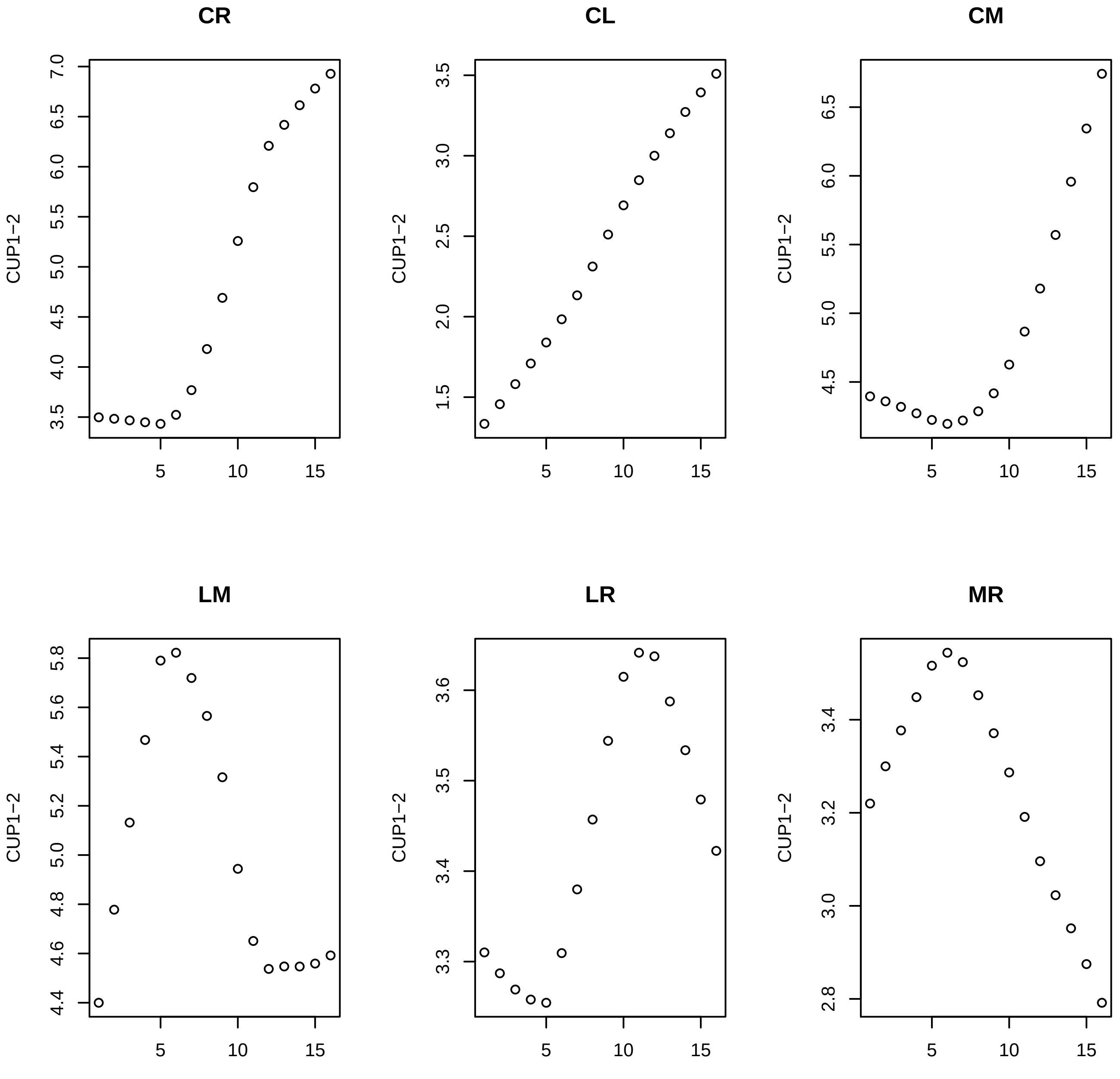
CR
CL
CM
LM
LR
MR
CUP1−2

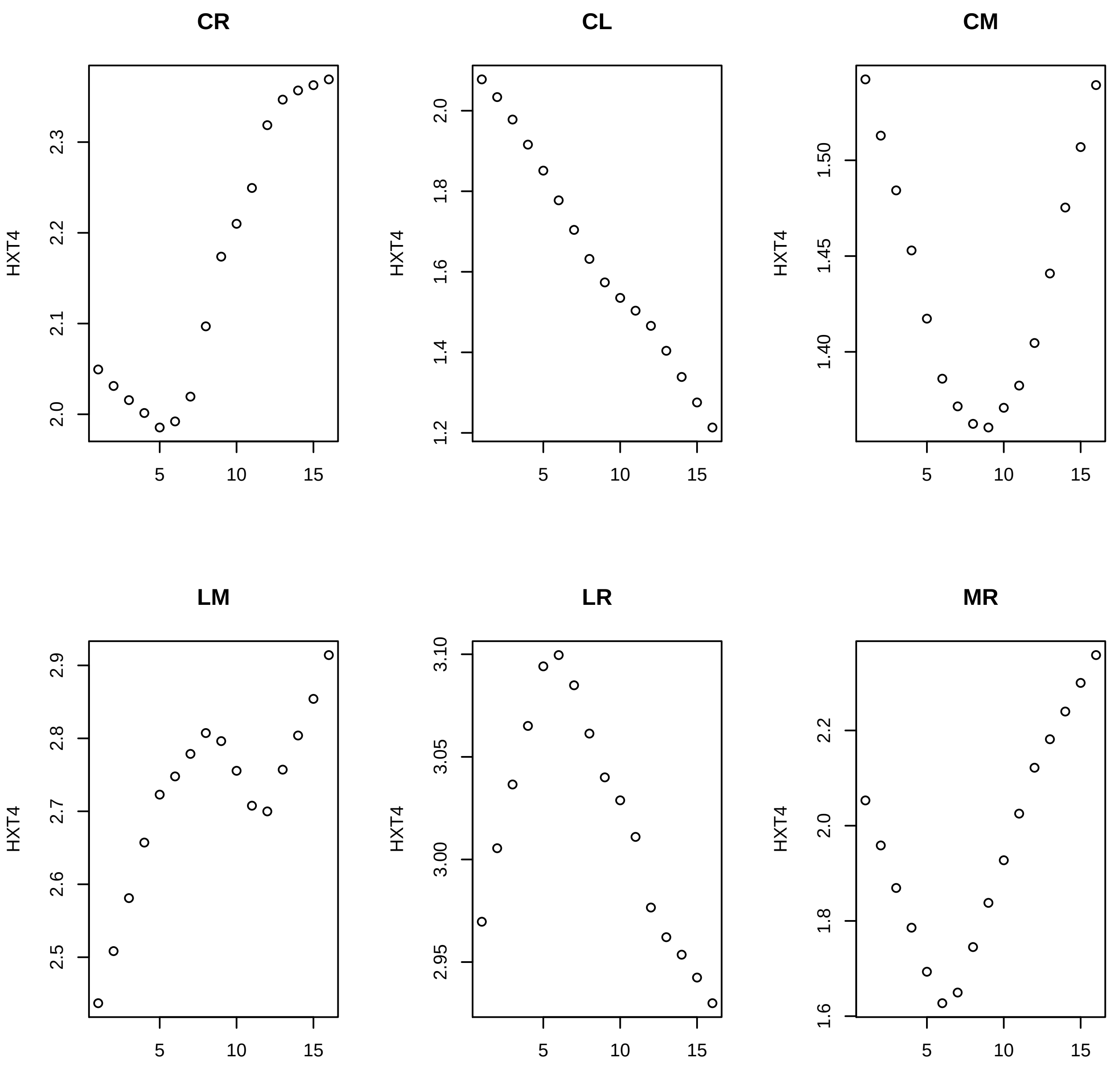
CR
CL
CM
LM
LR
MR
HXT4
5
10
15
2.0
2.1
2.2
2.3
1.2
1.4
1.6
1.8
2.0
1.40
1.45
1.50
2.5
2.6
2.7
2.8
2.9
2.95
3.00
3.05
3.10
1.6
1.8
2.0
2.2

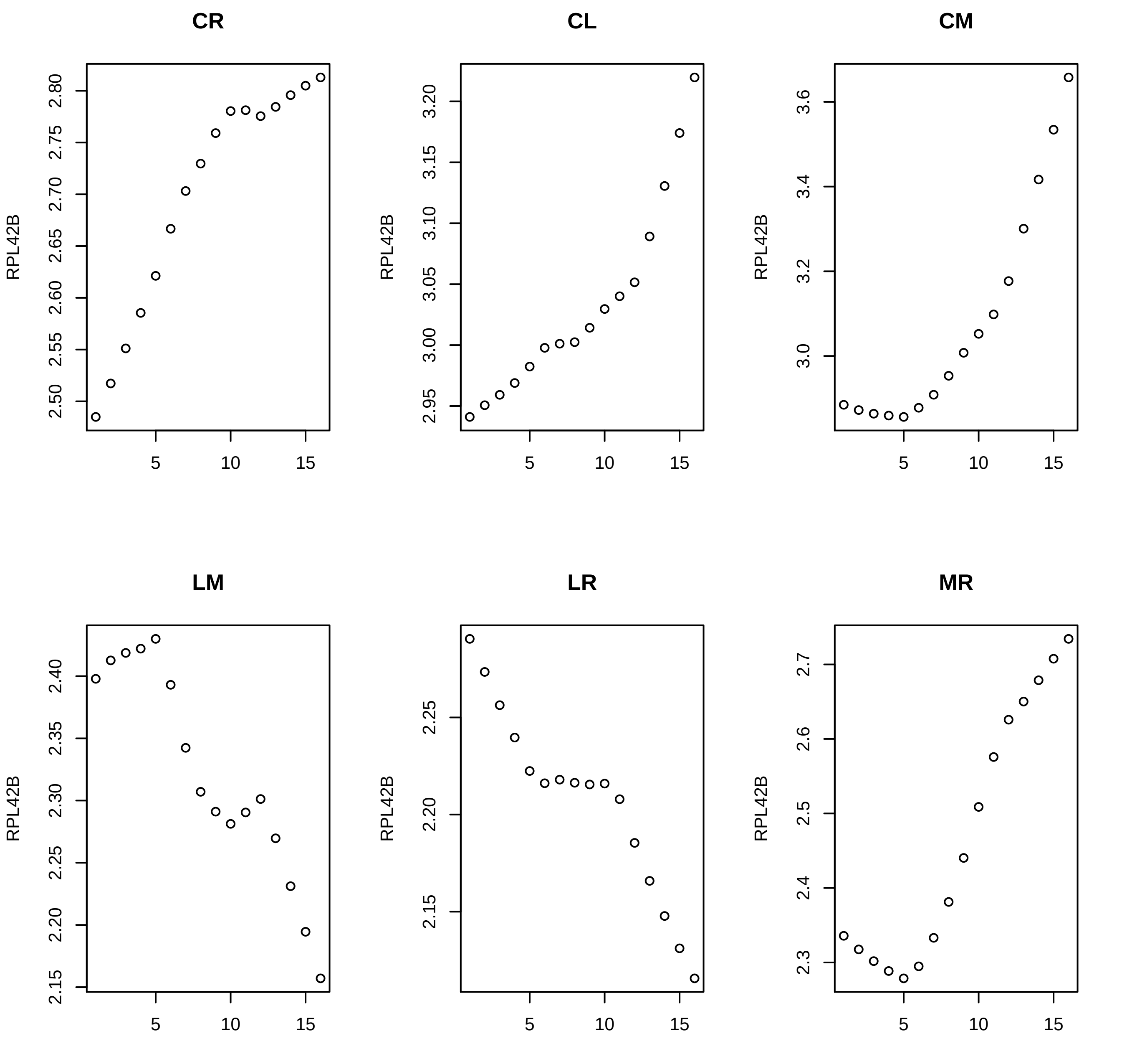
CR
CL
CM
LM
LR
MR
RPL42B
5
10
15
2.50
2.55
2.60
2.65
2.70
2.75
2.80
2.95
3.00
3.05
3.10
3.15
3.20
3.0
3.2
3.4
3.6
2.15
2.20
2.25
2.30
2.35
2.40
2.3
2.4
2.5
2.6
2.7

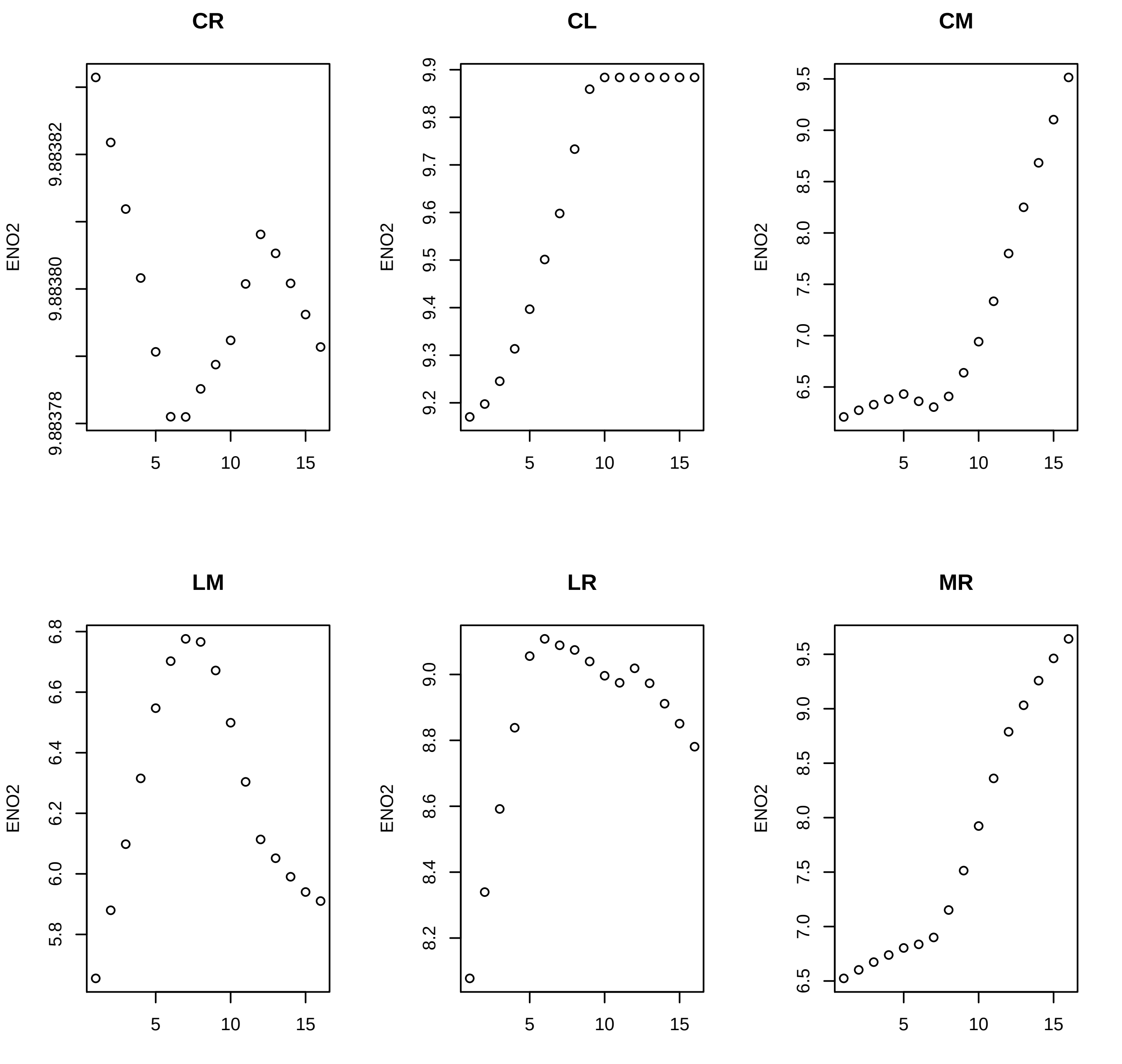
CR
CL
CM
LM
LR
MR
ENO2
9.88378
9.88380
9.88382
9.2
9.3
9.4
9.5
9.6
9.7
9.8
9.9
6.5
7.0
7.5
8.0
8.5
9.0
9.5
5.8
6.0
6.2
6.4
6.6
6.8
8.2
8.4
8.6
8.8
9.0
5
10
15

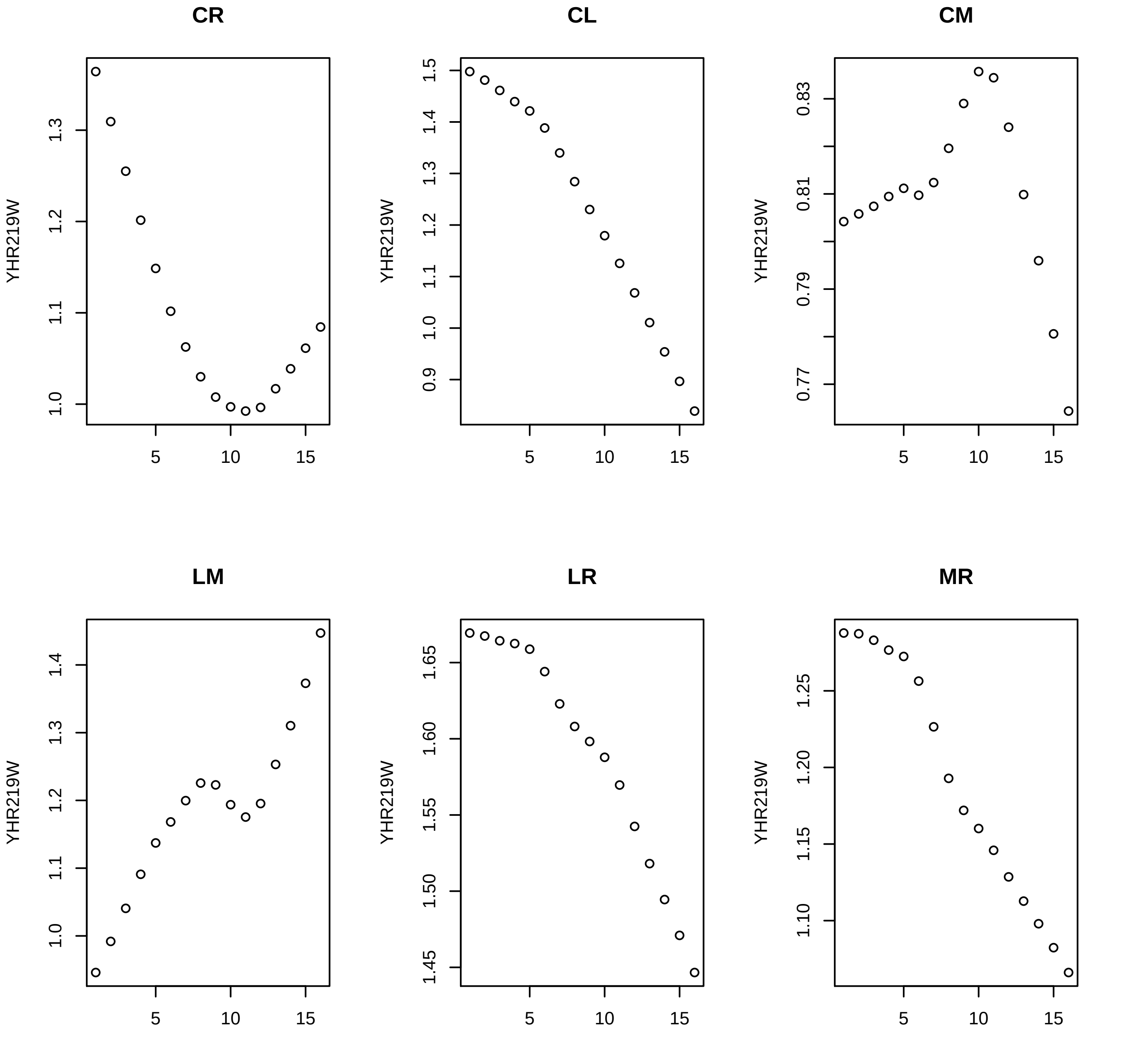

CR
CL
CM
LM
LR
MR
YHR219W
1.0
1.1
1.2
1.3
0.9
1.4
1.5
0.77
0.79
0.81
0.83
1.45
1.50
1.55
1.60
1.65
1.10
1.15
1.20
1.25
5
10
15

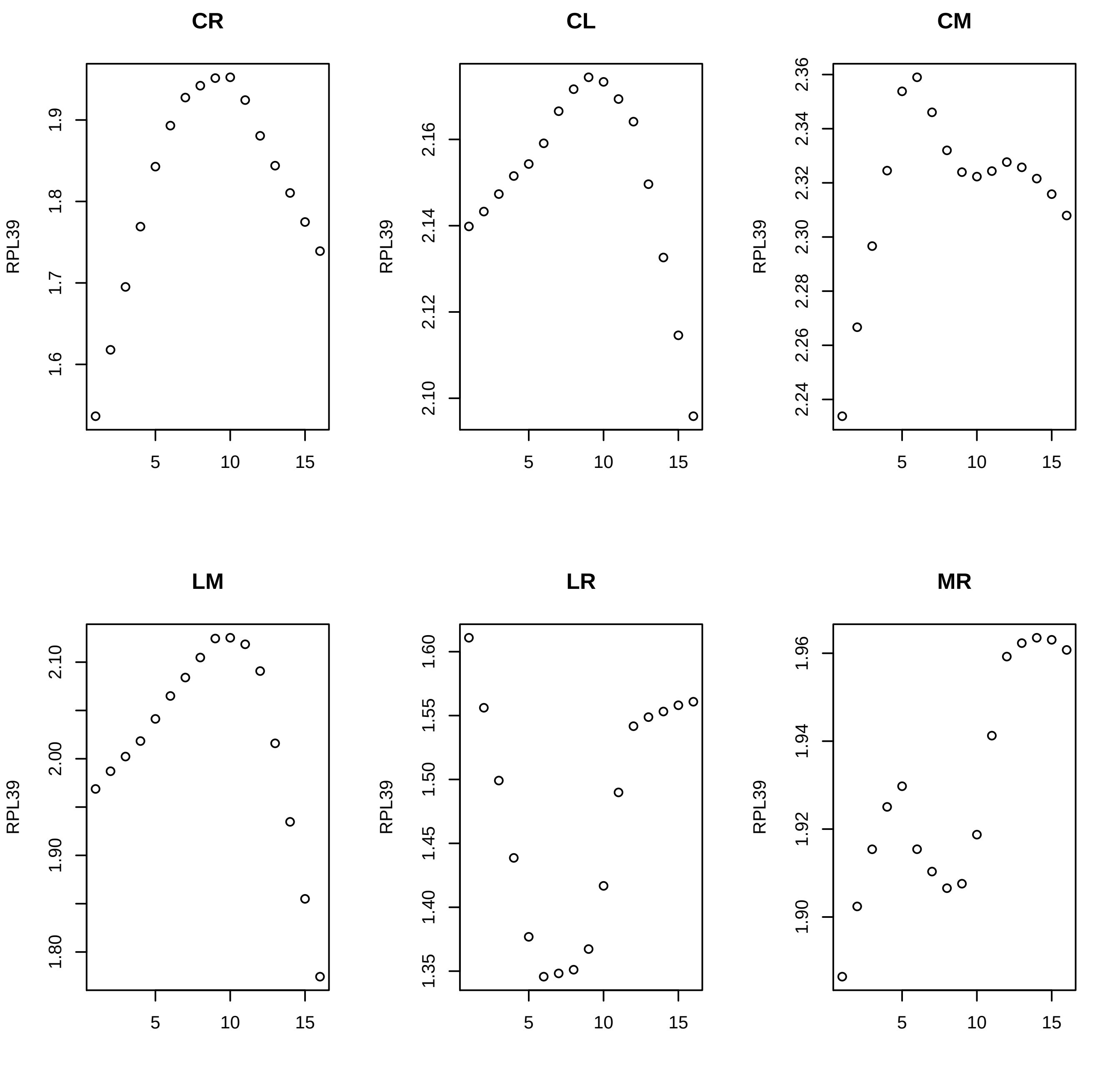
CR
CL
CM
LM
LR
MR
RPL39
5
10
15
1.6
1.7
1.8
1.9
2.10
2.12
2.14
2.16
2.24
2.26
2.28
2.30
2.32
2.34
2.36
1.80
1.90
2.00
2.10
1.35
1.40
1.45
1.50
1.55
1.60
1.90
1.92
1.94
1.96

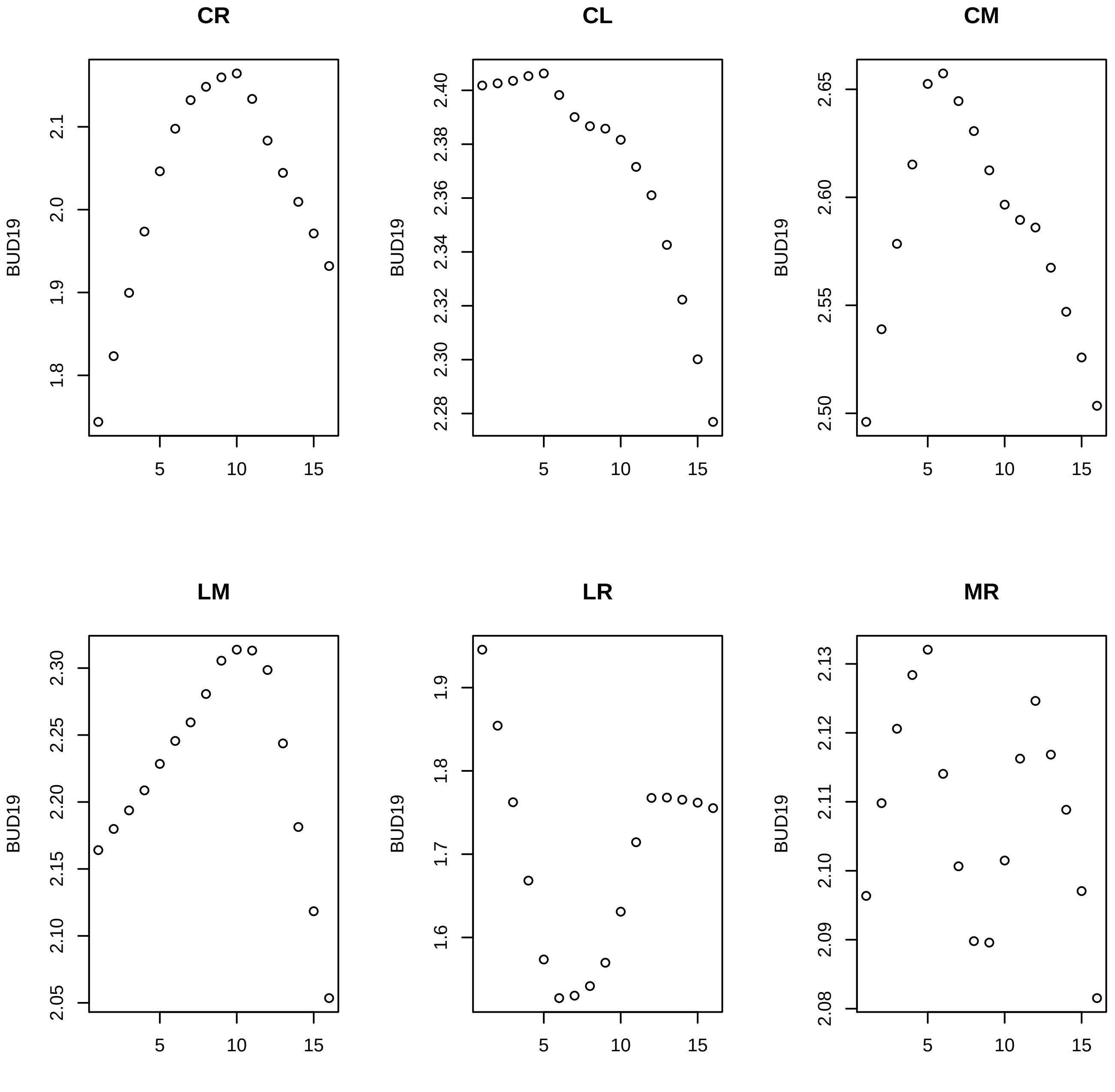
CR
CL
CM
LM
LR
MR
BUD19
5
10
15
1.8
1.9
2.0
2.1
2.28
2.30
2.32
2.34
2.36
2.38
2.40
2.50
2.55
2.60
2.65
2.05
2.10
2.15
2.20
2.25
2.30
1.6
1.7
1.8
1.9
2.08
2.09
2.10
2.11
2.12
2.13

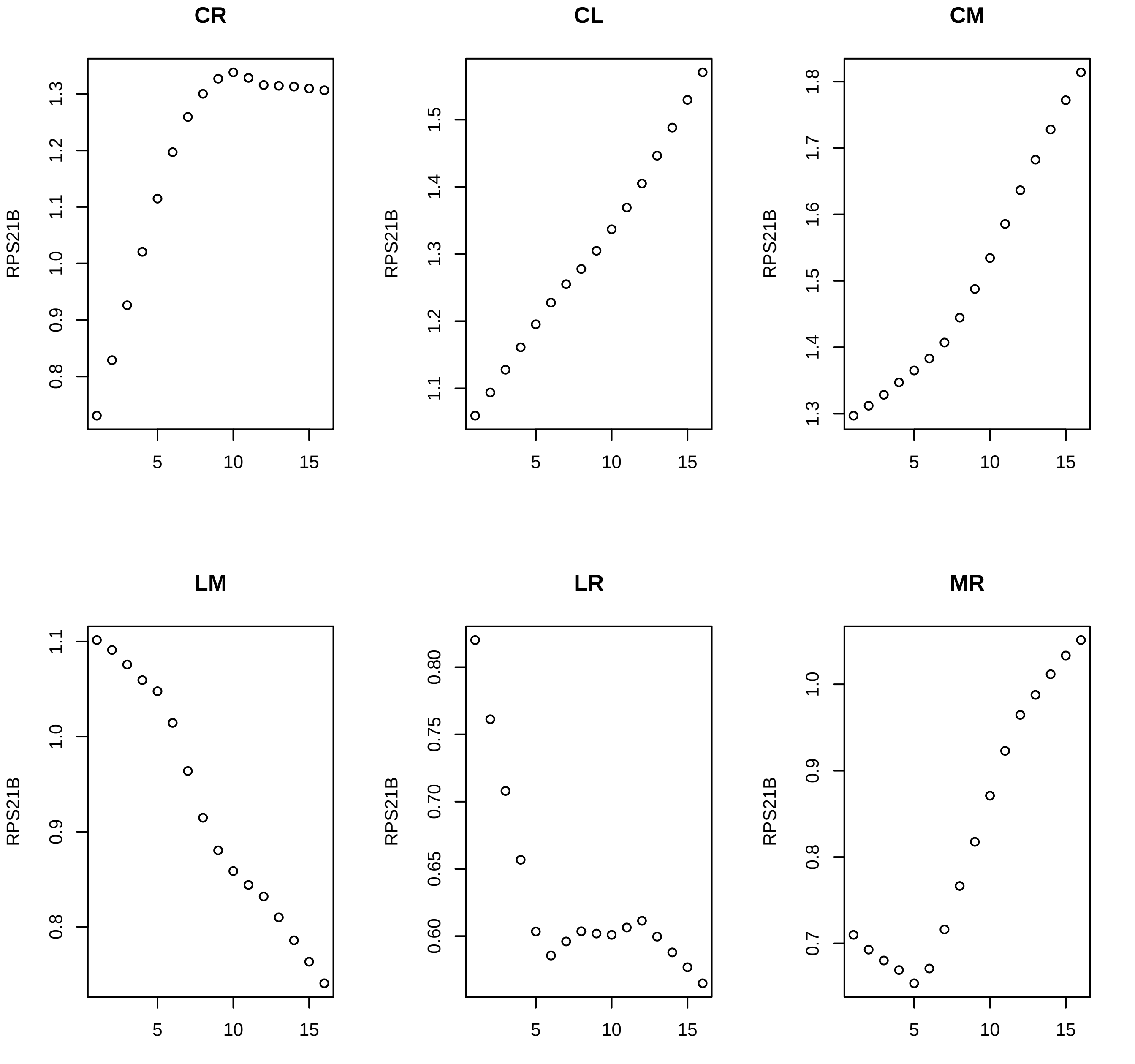

CR
CL
CM
LM
LR
MR
RPS21B

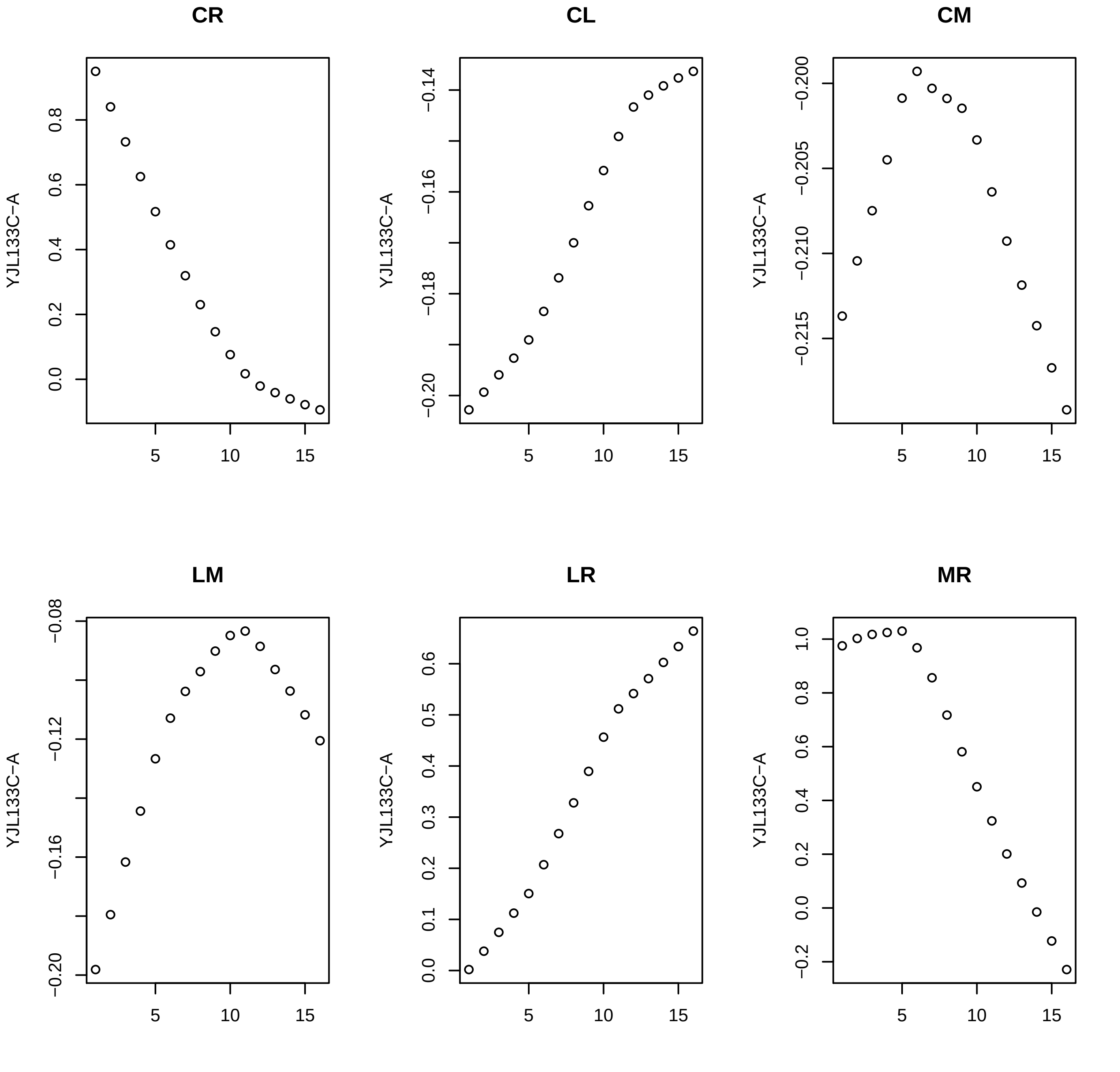
CR
CL
CM
LM
LR
MR
YJL133C−A
0.0
0.2
0.4
0.6
0.8
−0.20
−0.18
−0.16
−0.14
−0.215
−0.210
−0.205
−0.200
−0.20
−0.16
−0.12
−0.08
0.0
0.1
0.2
0.3
0.4
0.5
0.6
−0.2
0.0
0.2
0.4
0.6
0.8
1.0
5
10
15

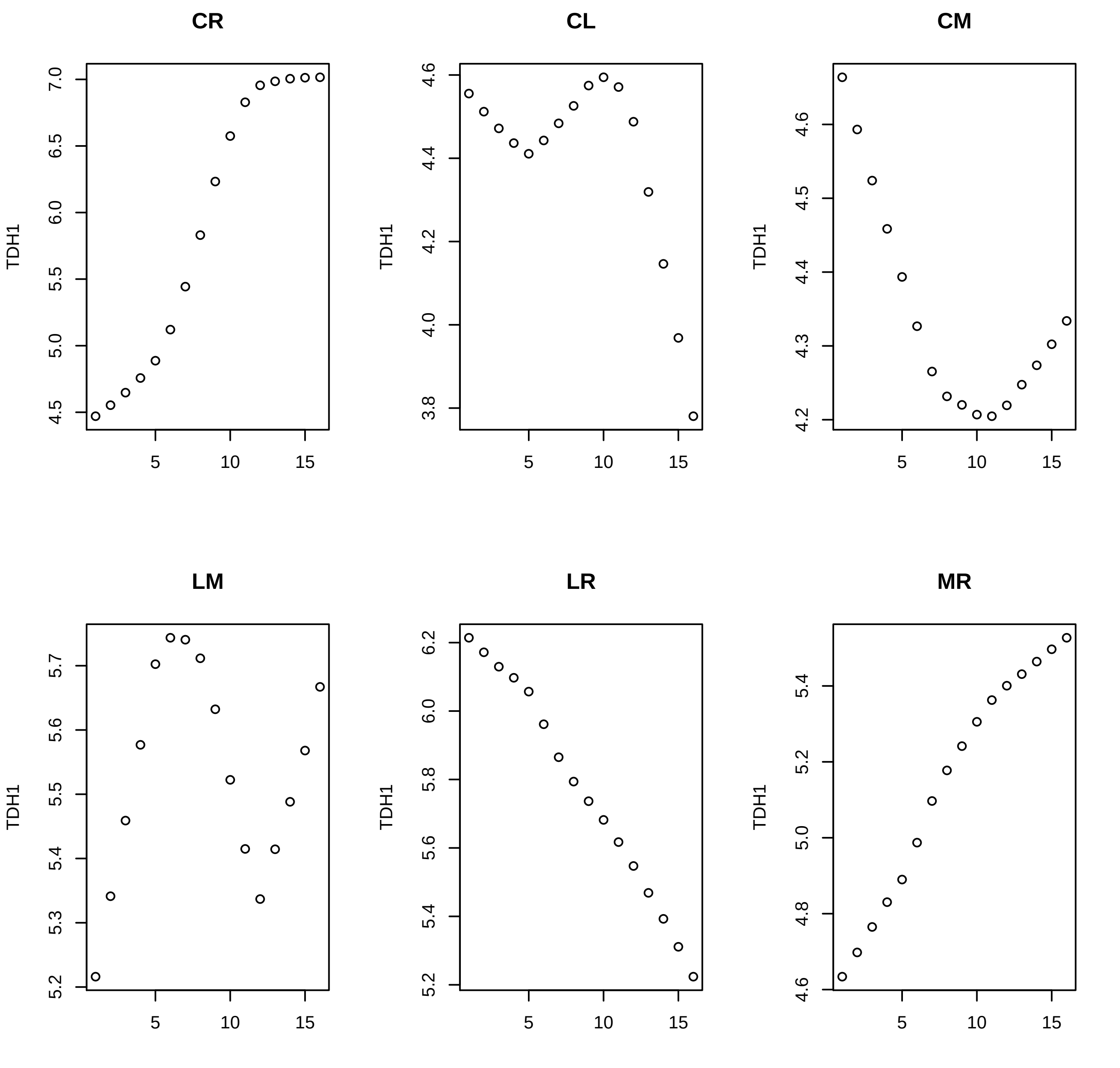
CR
CL
CM
LM
LR
MR
TDH1
5
10
15
7.0
6.5
6.0
5.5
5.0
4.5
4.6
4.4
4.2
4.0
3.8
4.6
4.5
4.4
4.3
4.2
5.7
5.6
5.5
5.4
5.3
5.2
6.2
6.0
5.8
5.6
5.4
5.2
5.4
5.2
5.0
4.8
4.6

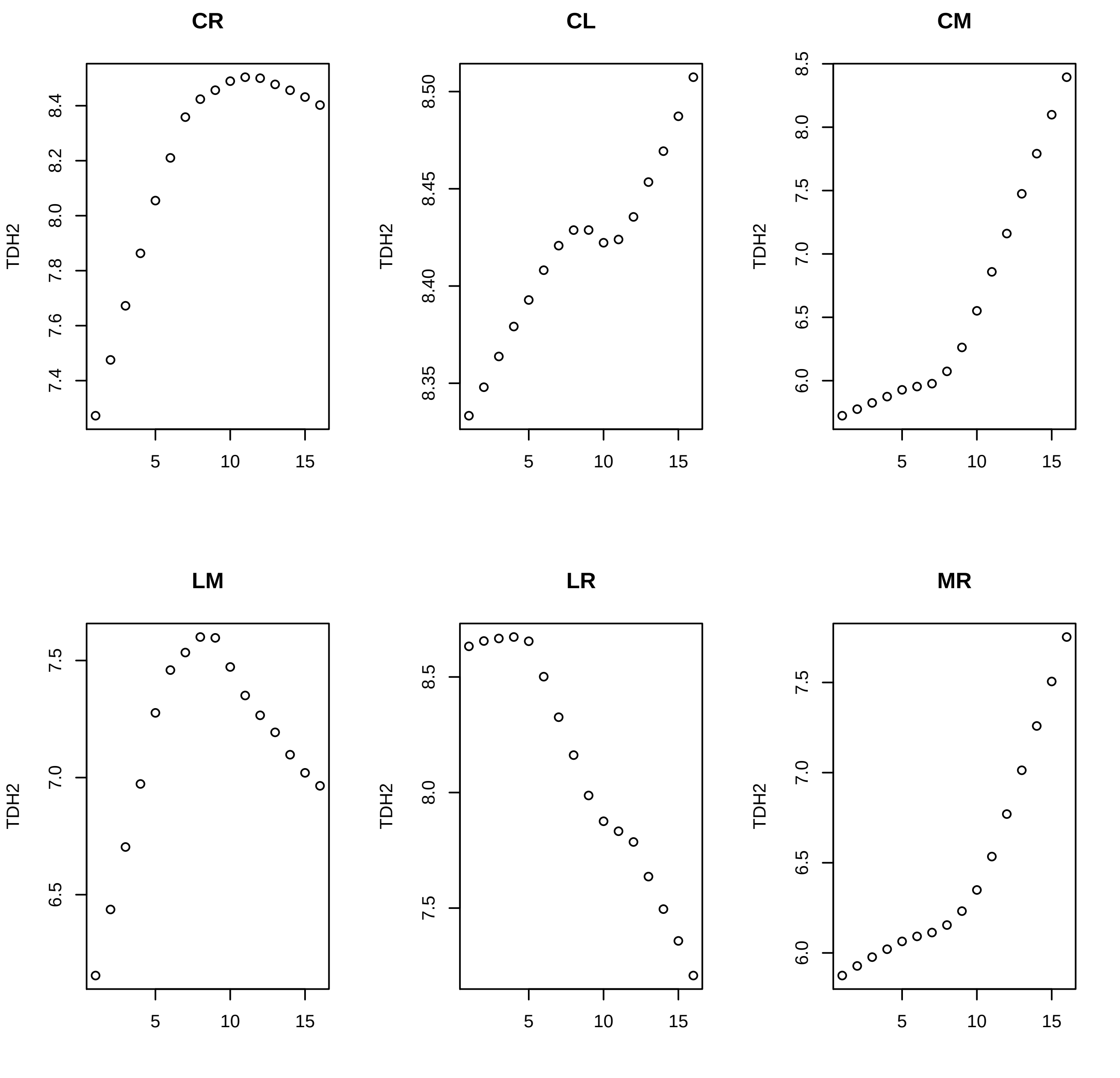
CR
CL
CM
LM
LR
MR
TDH2
5
10
15
7.4
7.6
7.8
8.0
8.2
8.4
8.35
8.40
8.45
8.50
6.0
6.5
7.0
7.5
8.0
8.5

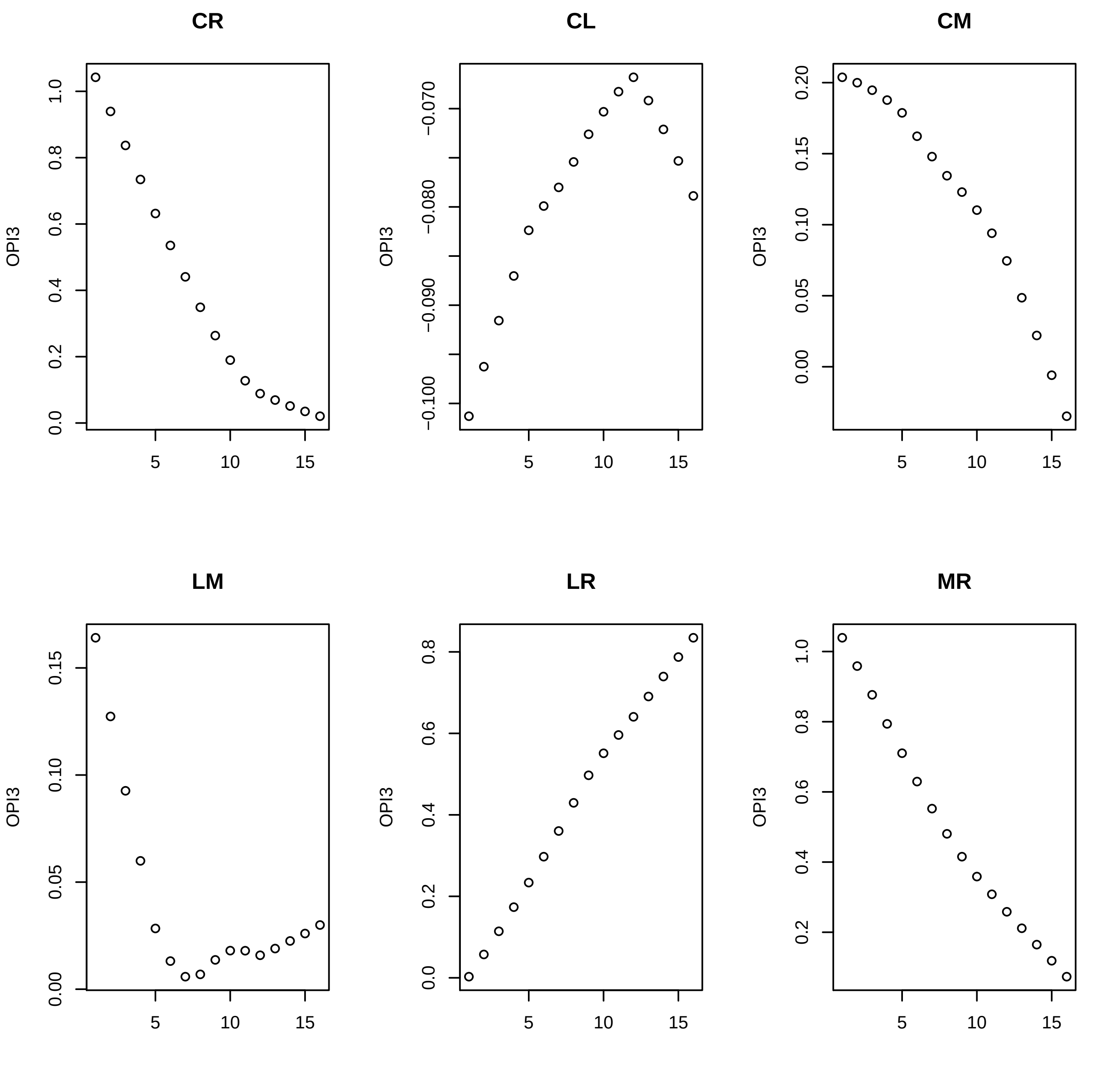

CR
CL
CM
LM
LR
MR
OPI3

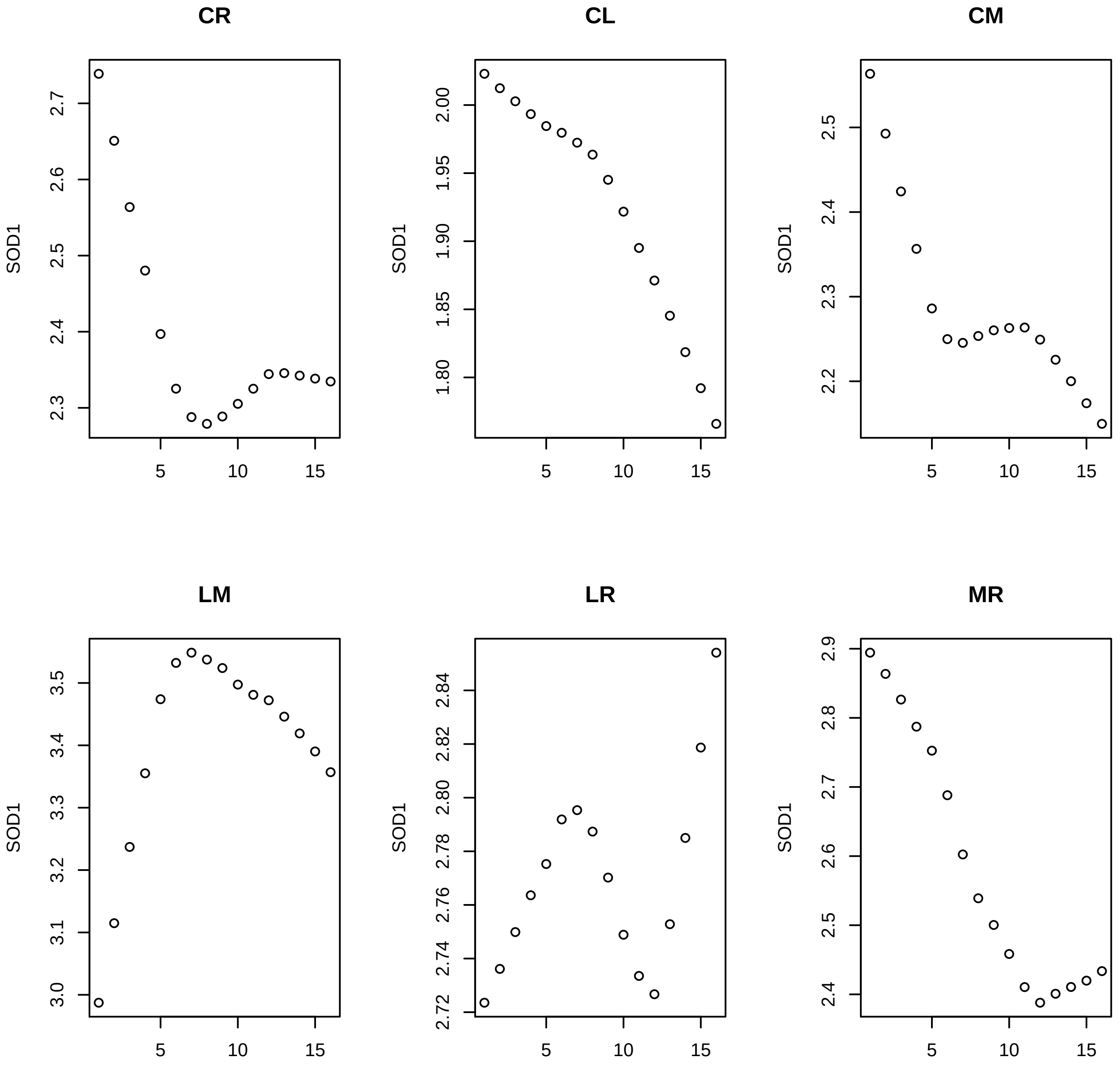
CR
CL
CM
LM
LR
MR
SOD1
5
10
15
2.3
2.4
2.5
2.6
2.7
1.80
1.85
1.90
1.95
2.00
2.2
3.0
3.1
3.2
3.3
3.4
3.5
2.72
2.74
2.76
2.78
2.80
2.82
2.84
2.8
2.9

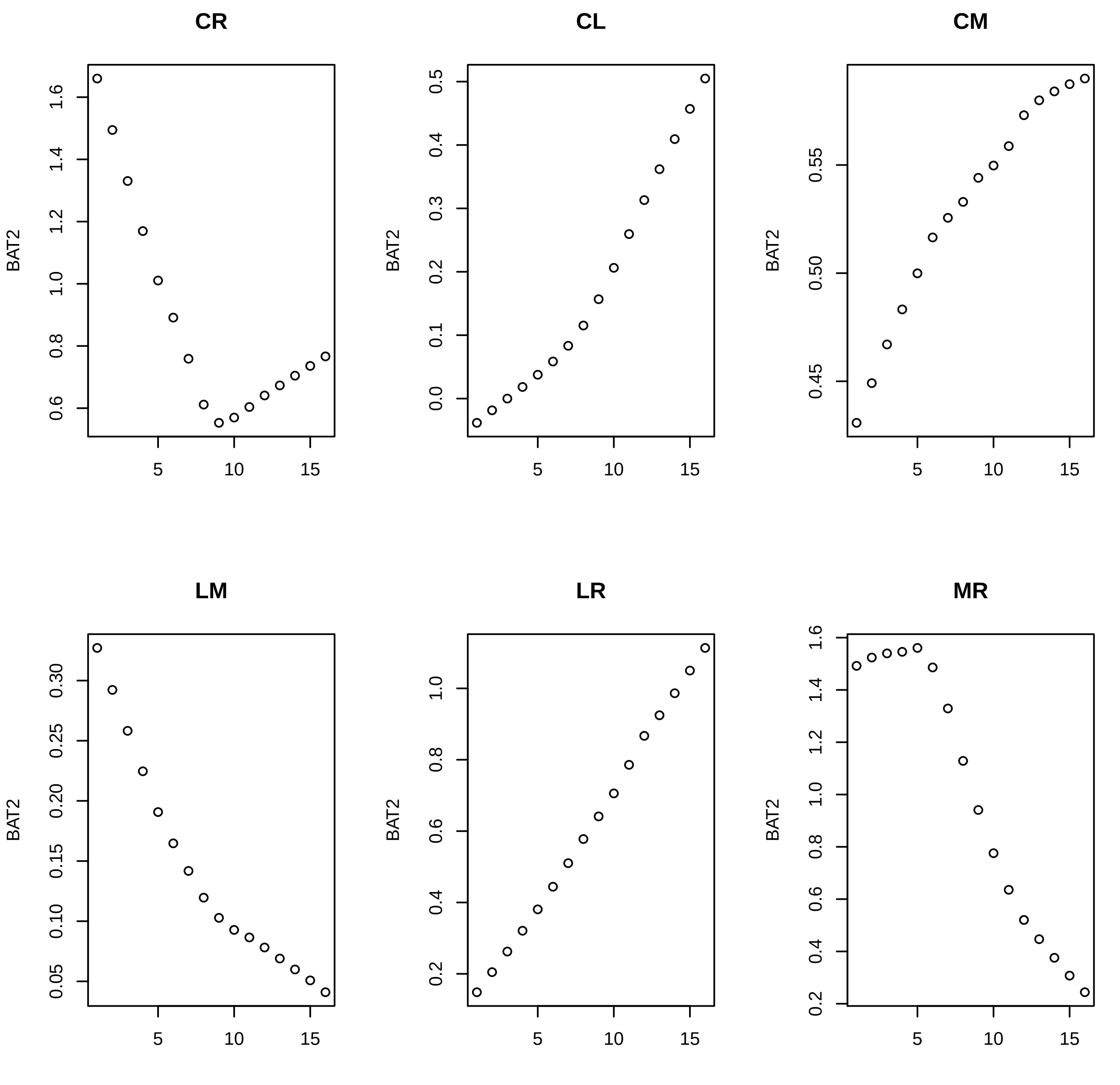
CR
CL
CM
LM
LR
MR
BAT2
5
10
15
1.6
1.4
1.2
1.0
0.8
0.6
0.5
0.4
0.3
0.2
0.1
0.0
0.55
0.50
0.45
0.30
0.25
0.20
0.15
0.10
0.05

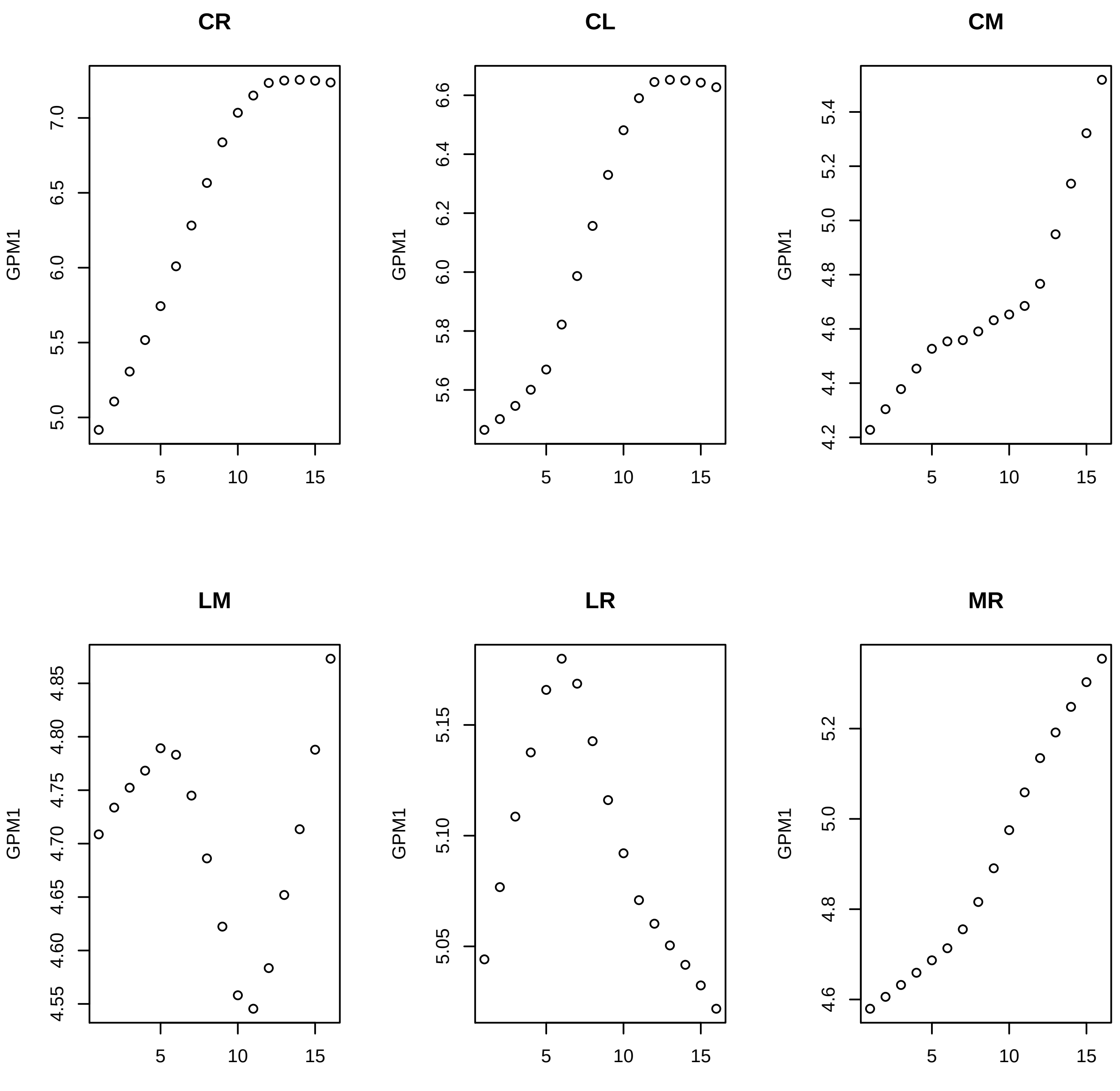
CR
CL
CM
LM
LR
MR
GPM1
5
10
15

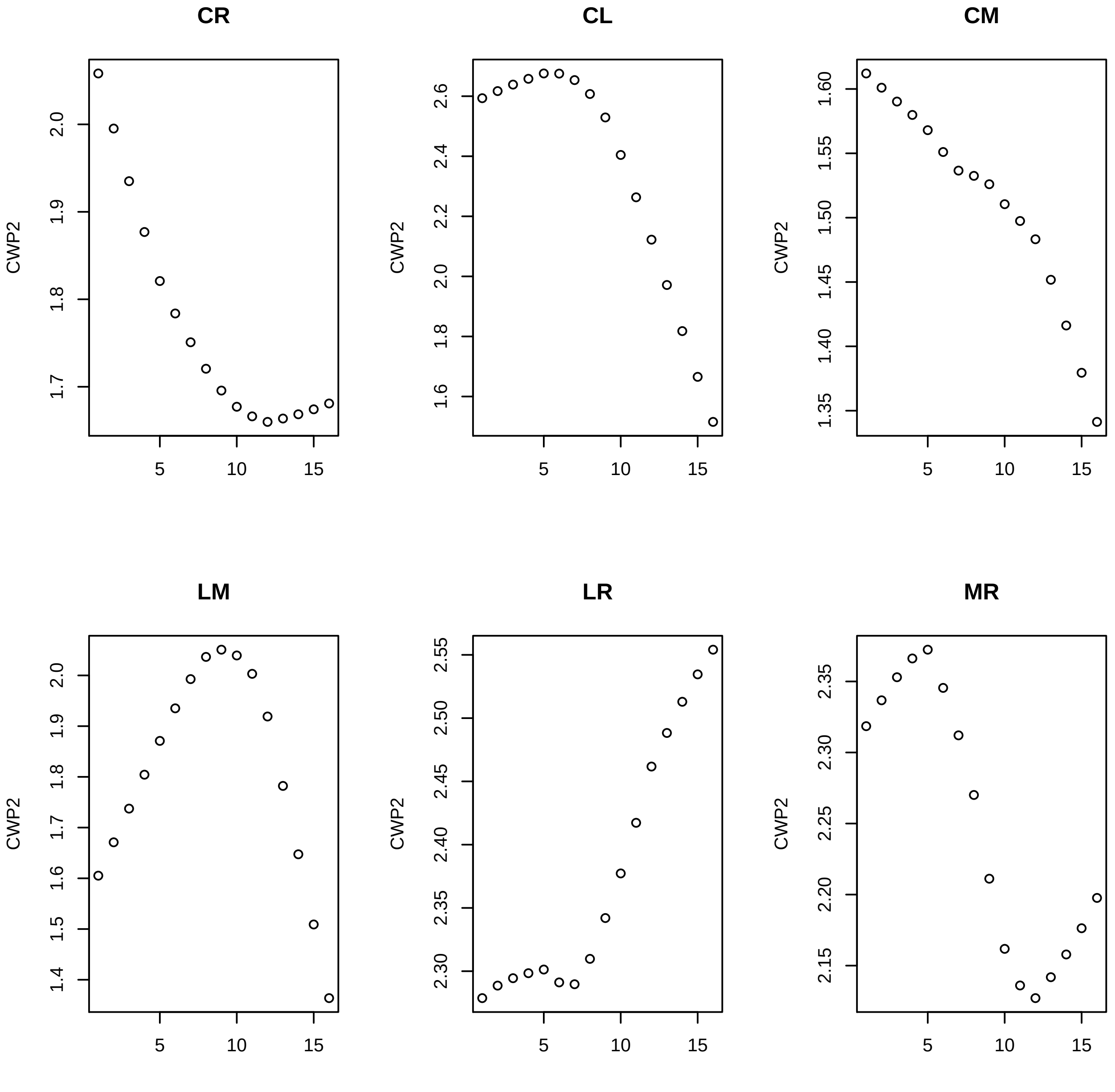
CR
CL
CM
LM
LR
MR
CWP2

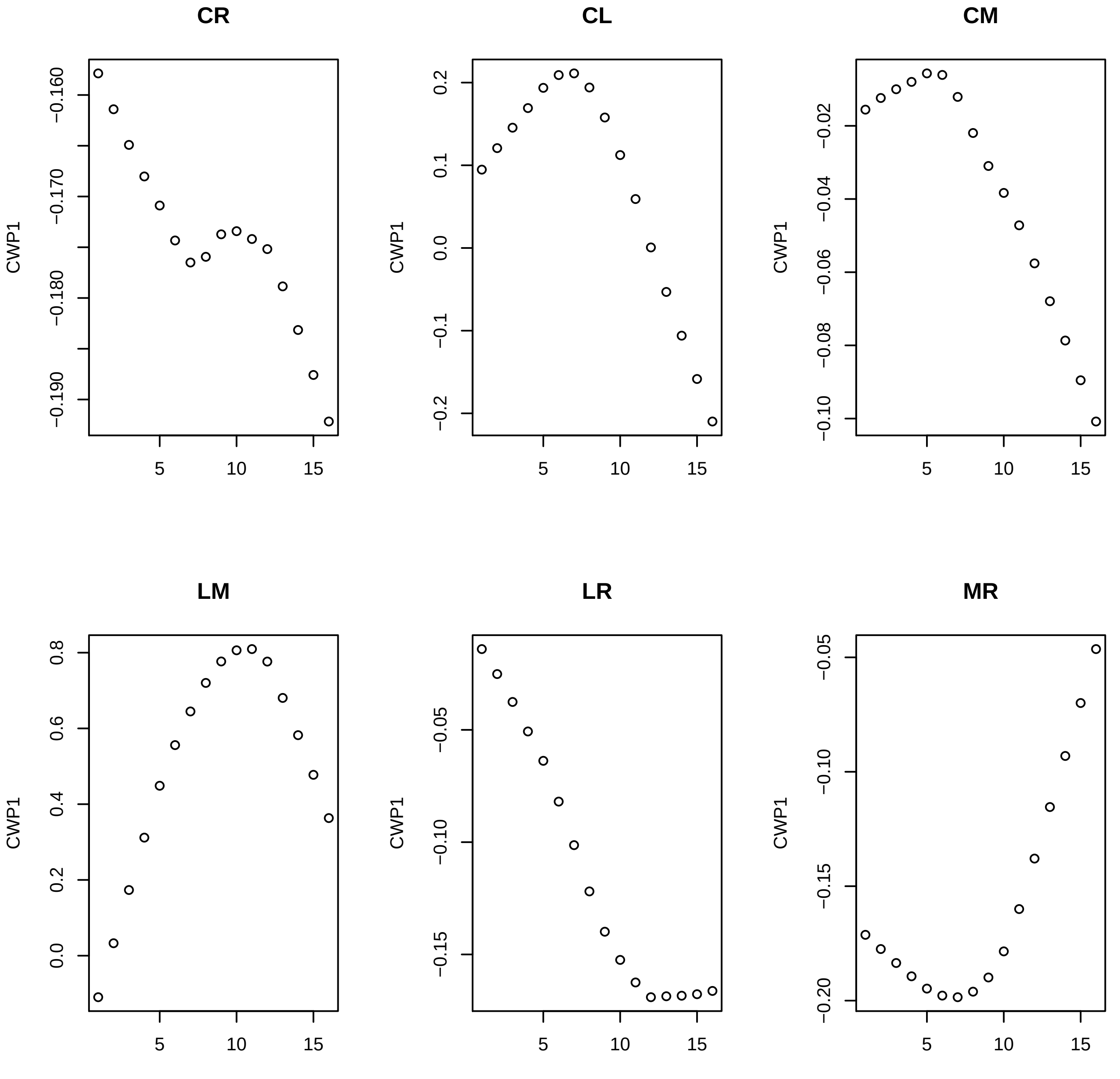
CR
CWP1
−0.160
−0.170
−0.180
−0.190
5
10
15
CL
CWP1
0.2
0.1
0.0
−0.1
−0.2
5
10
15
CM
CWP1
−0.02
−0.04
−0.06
−0.08
−0.10
5
10
15
LM
CWP1
0.8
0.6
0.4
0.2
0.0
5
10
15
LR
CWP1
−0.05
−0.10
−0.15
5
10
15
MR
CWP1
−0.05
−0.10
−0.15
−0.20
5
10
15

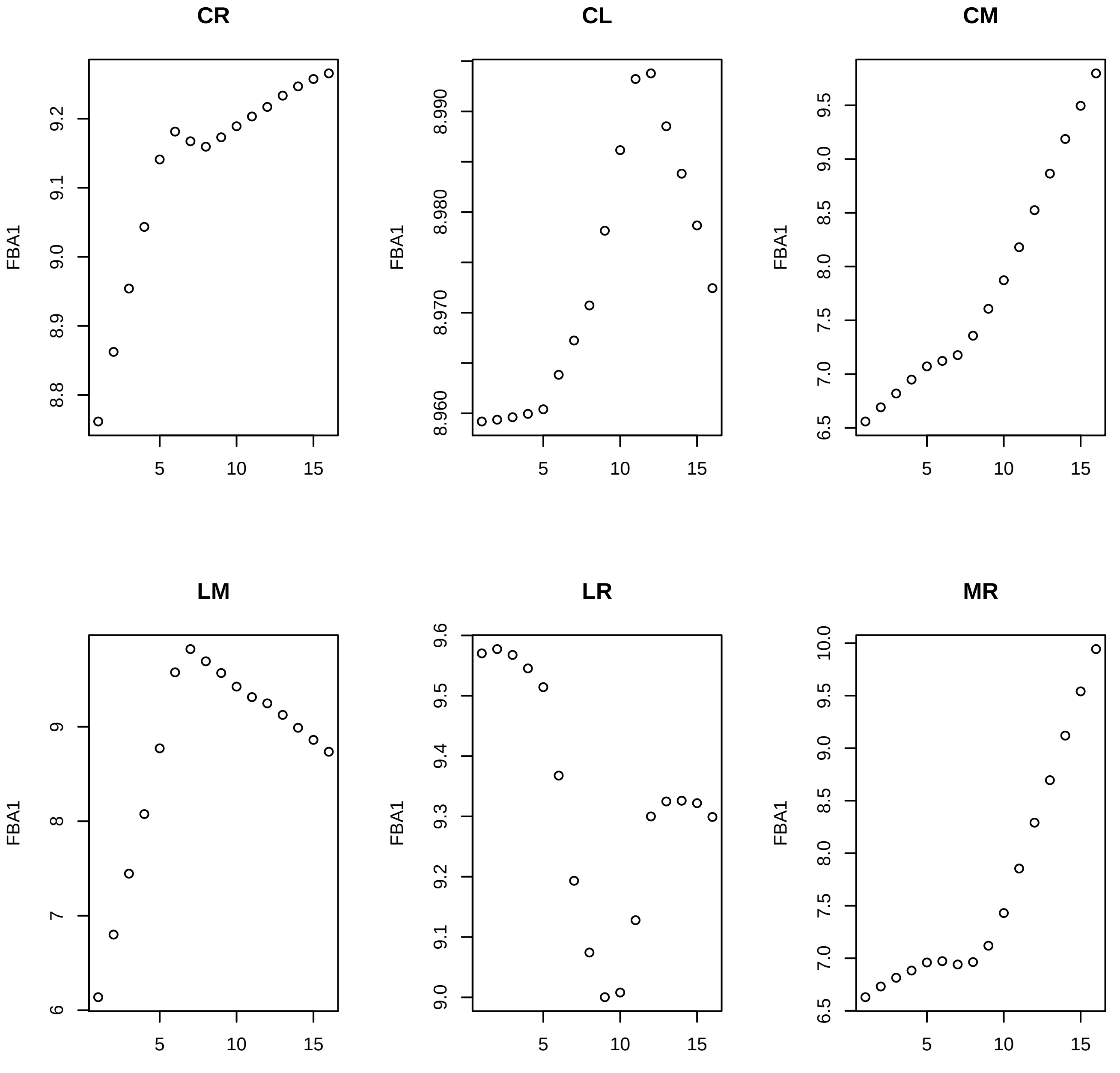

CR
CL
CM
LM
LR
MR
FBA1

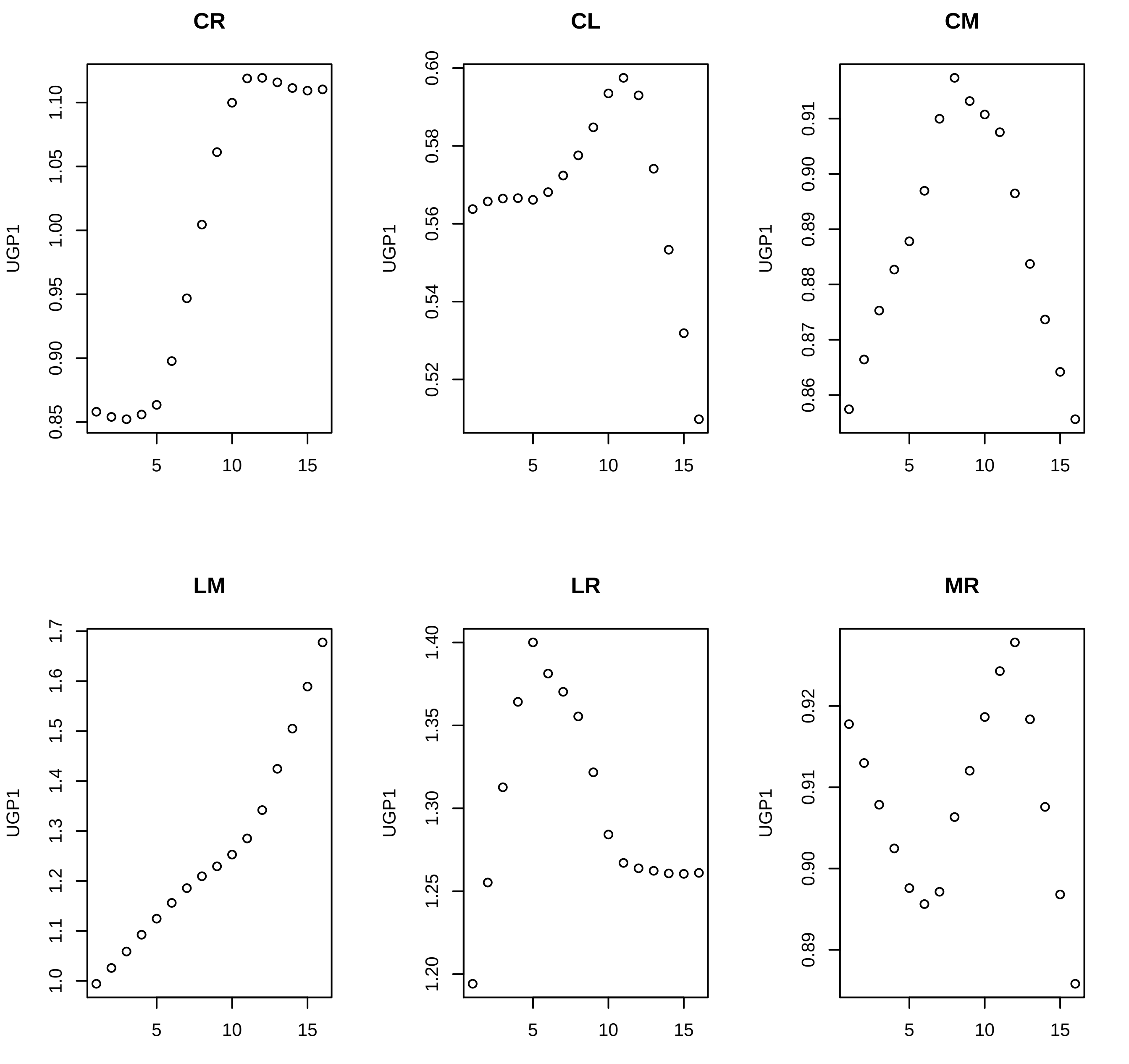

CR
CL
CM
LM
LR
MR
UGP1
5
10
15
0.85
0.90
0.95
1.00
1.05
1.10
0.52
0.54
0.56
0.58
0.60
0.86
0.87
0.88
0.89
0.90
0.91
1.0
1.1
1.2
1.3
1.4
1.5
1.6
1.7
1.20
1.25
1.30
1.35
1.40
0.89
0.90
0.91
0.92

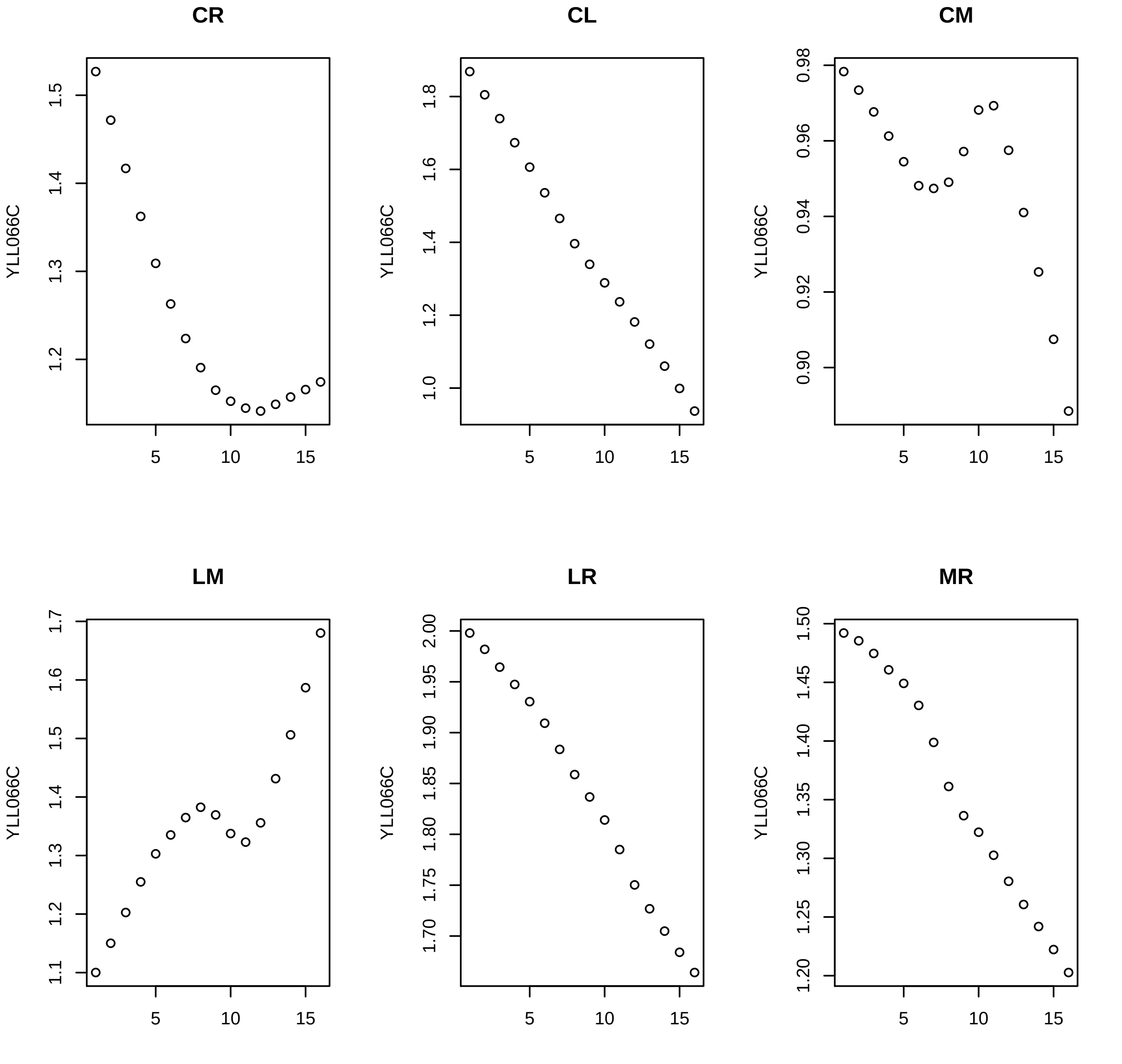
CR
CL
CM
LM
LR
MR
YLL066C
5
10
15
1.2
1.3
1.4
1.5
1.0
1.2
1.4
1.6
1.8
0.90
0.92
0.94
0.96
0.98
1.1
1.2
1.3
1.4
1.5
1.6
1.7
1.70
1.75
1.80
1.85
1.90
1.95
2.00
1.20
1.25
1.30
1.35
1.40
1.45
1.50

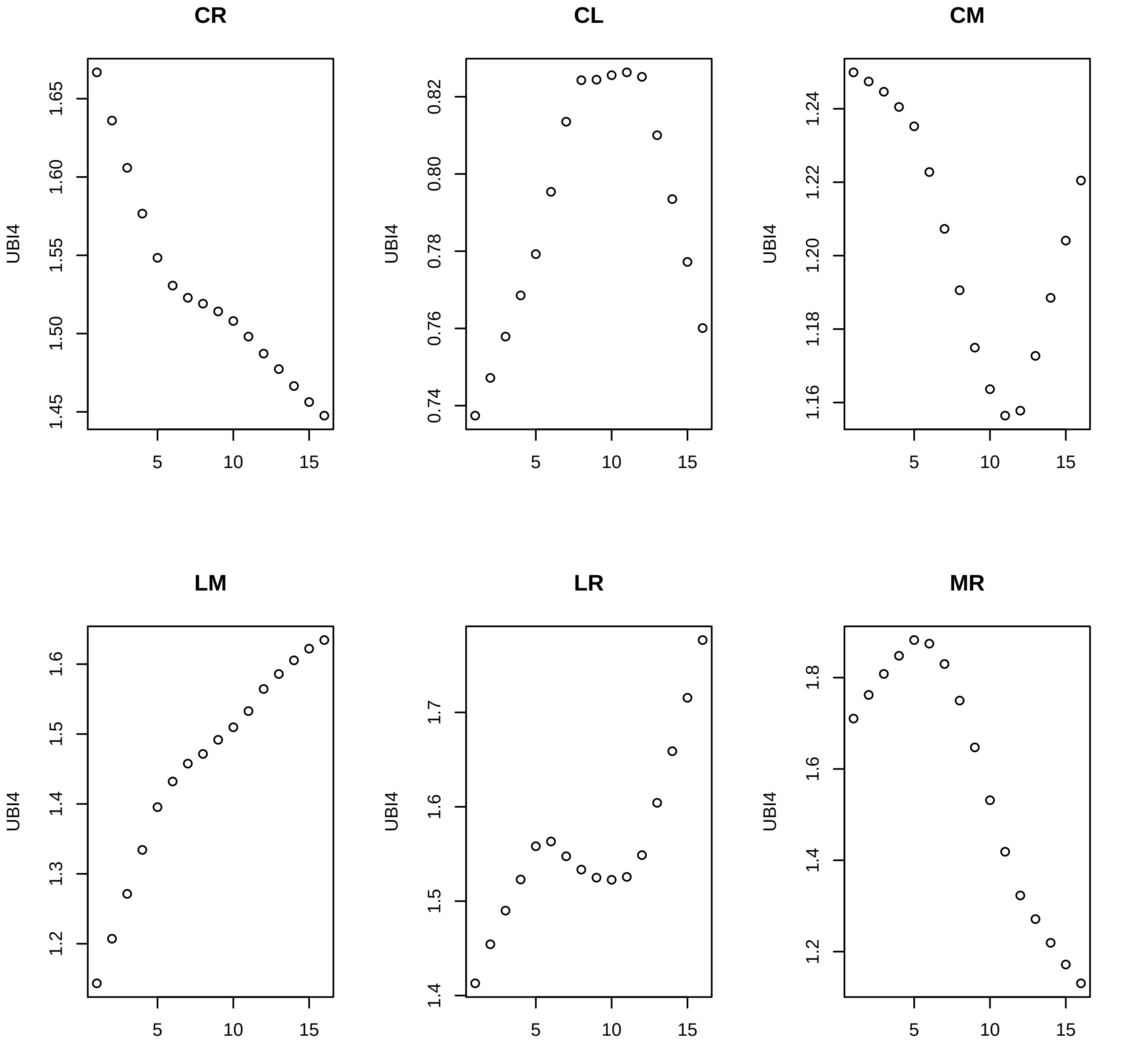
CR
CL
CM
LM
LR
MR
UBI4
UBI4
UBI4
UBI4
UBI4
UBI4
1.65
1.60
1.55
1.50
1.45
0.82
0.80
0.78
0.76
0.74
1.24
1.22
1.20
1.18
1.16
1.6
1.5
1.4
1.3
1.2
1.7
1.6
1.5
1.4
1.8
1.6
1.4
1.2
5
10
15

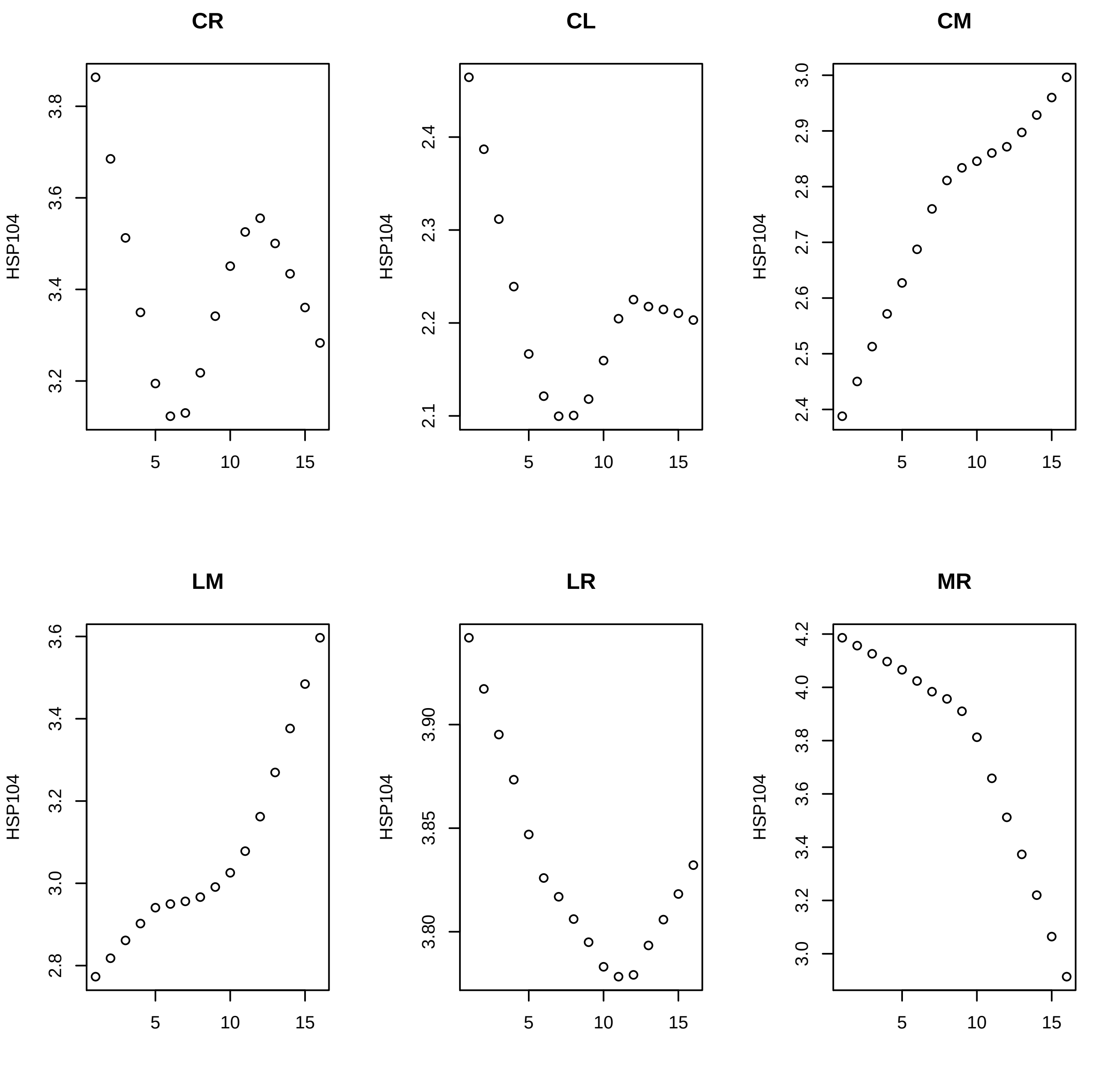
CR
CL
CM
LM
LR
MR
HSP104
3.2
3.4
3.6
3.8
2.1
2.2
2.3
2.4
2.4
2.5
2.6
2.7
2.8
2.9
3.0
2.8
3.0
3.2
3.4
3.6
3.80
3.85
3.90
3.0
3.2
3.4
3.6
3.8
4.0
4.2
5
10
15

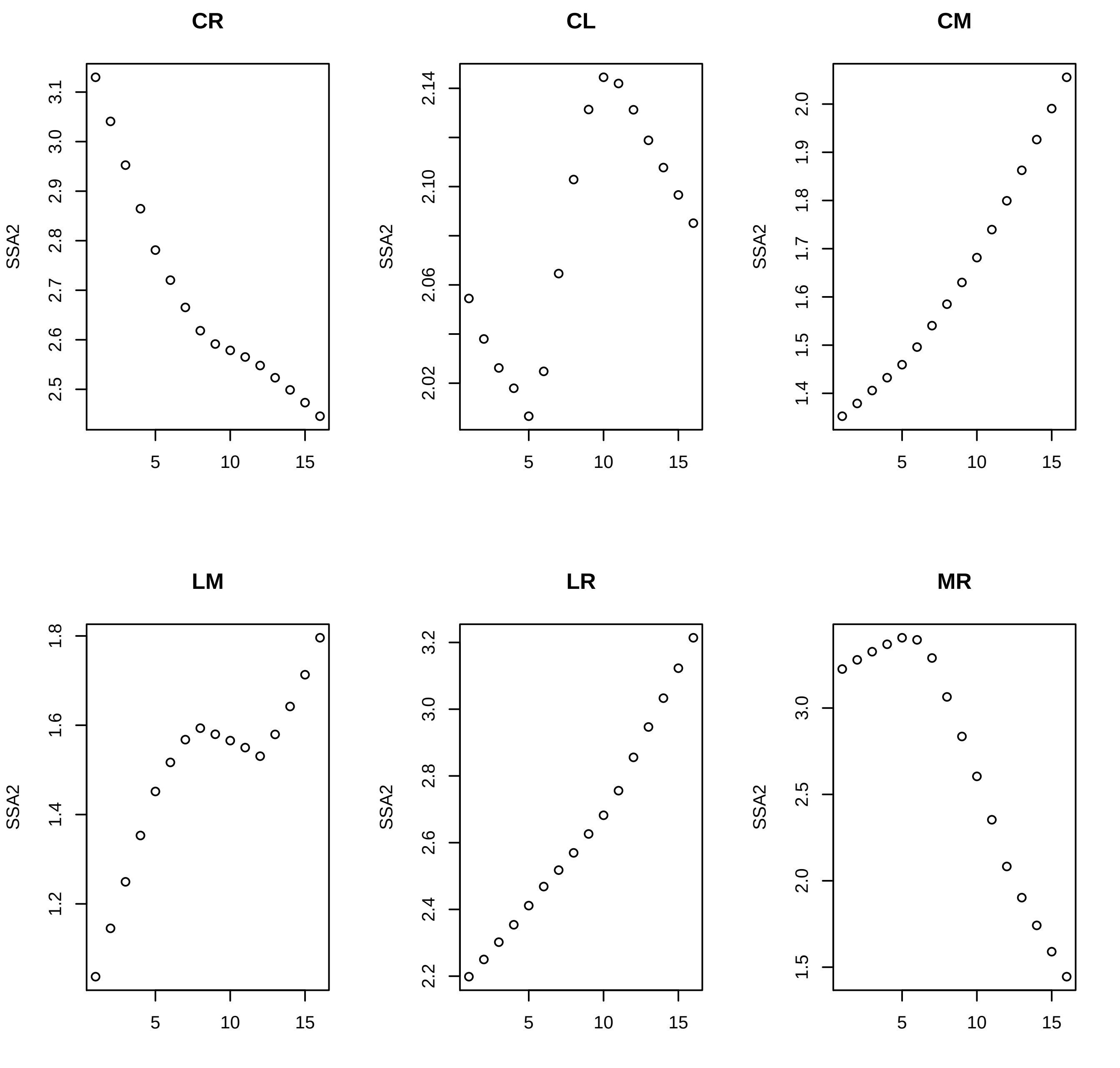
CR
CL
CM
LM
LR
MR
SSA2
5
10
15

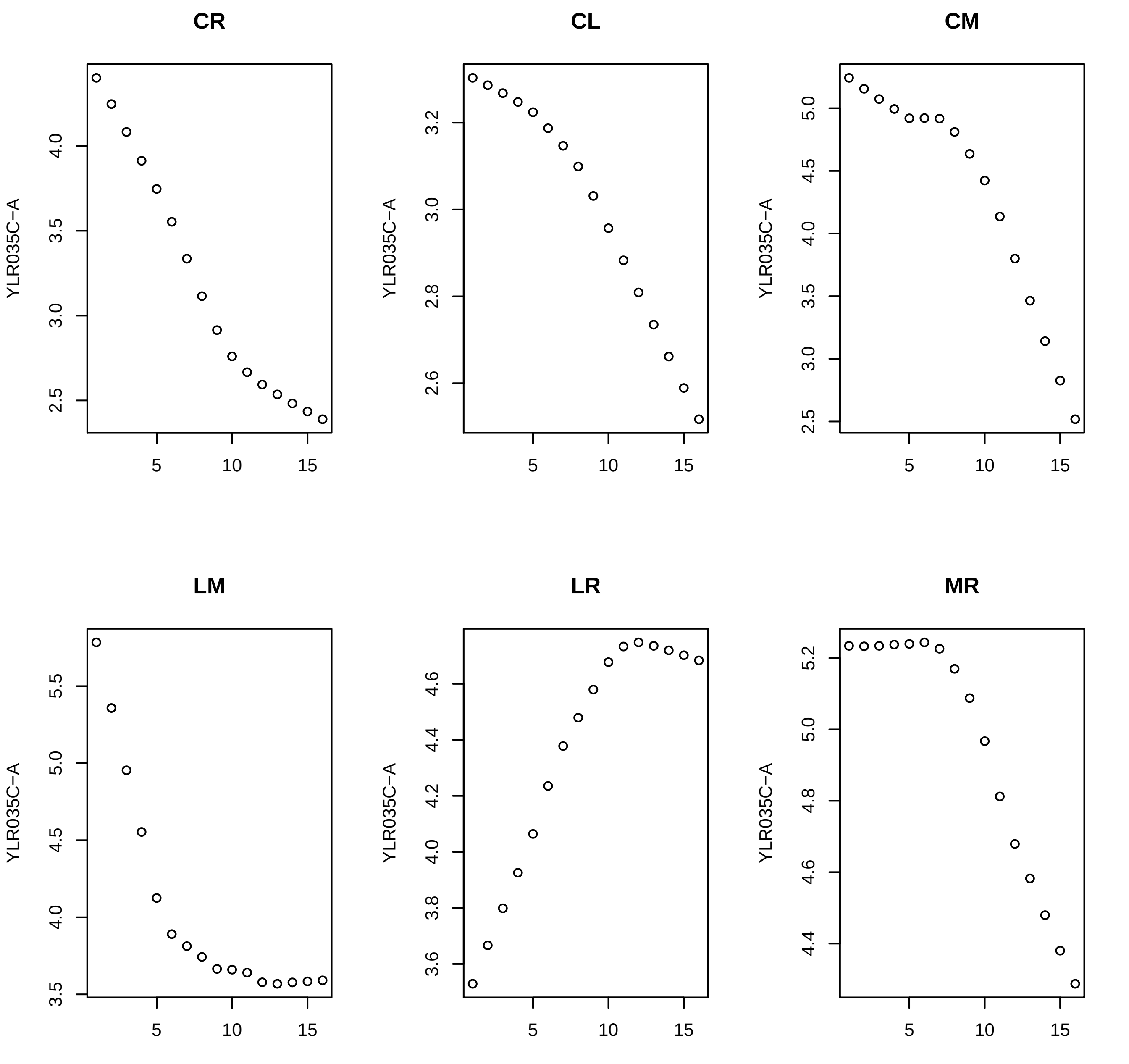
CR
CL
CM
LM
LR
MR
YLR035C−A
5
10
15

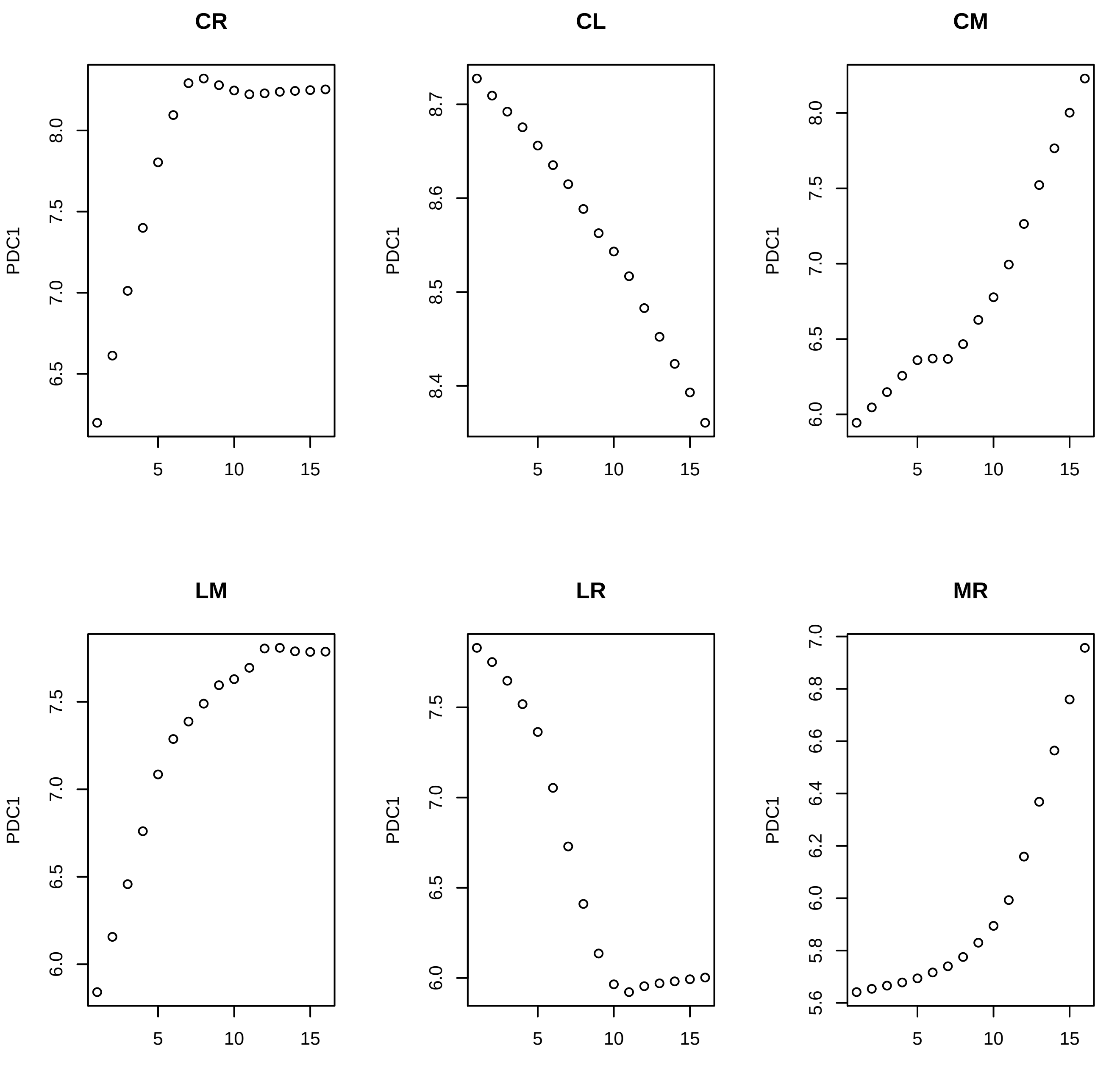
CR
CL
CM
LM
LR
MR
PDC1

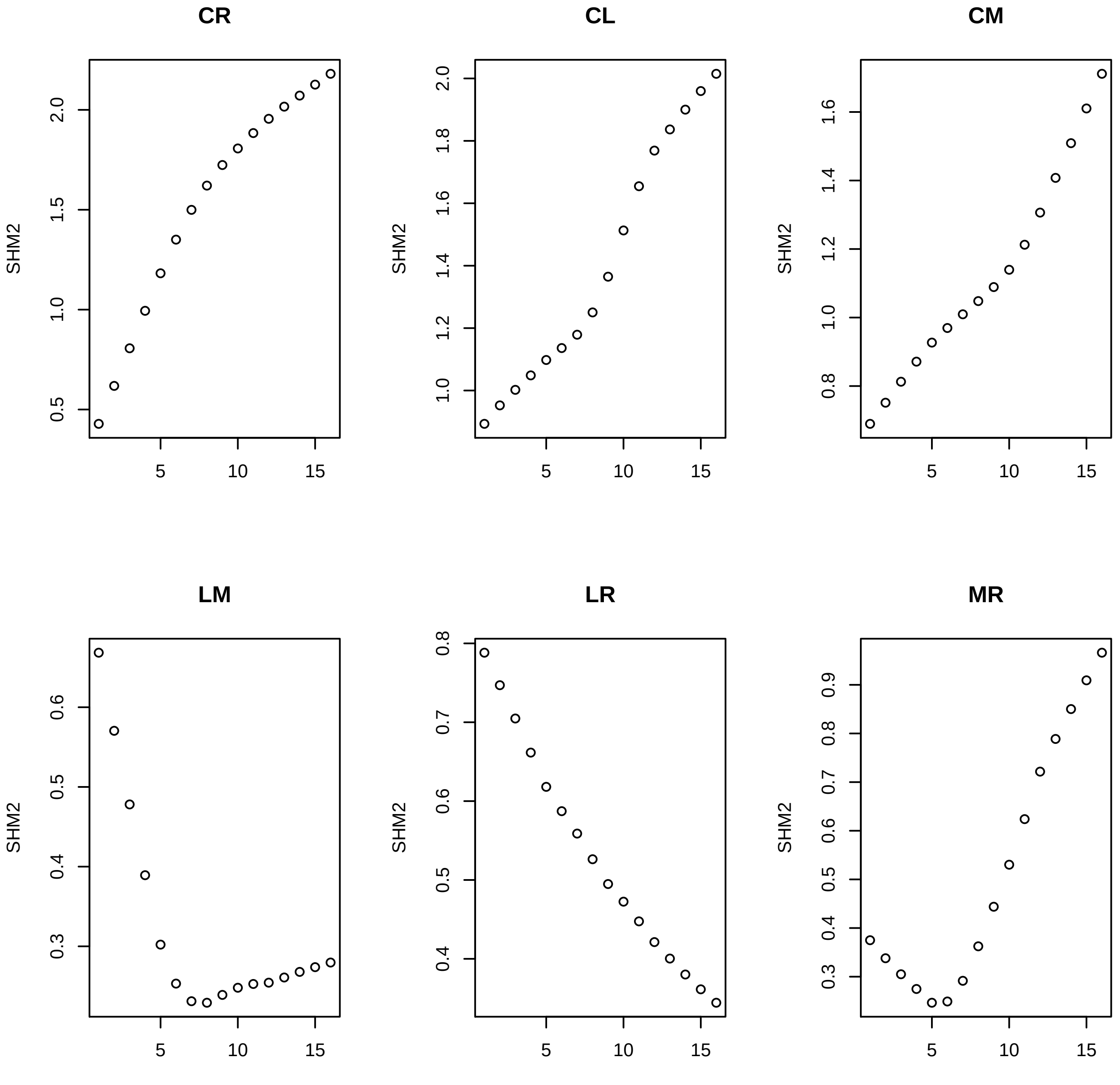
CR
CL
CM
LM
LR
MR
SHM2

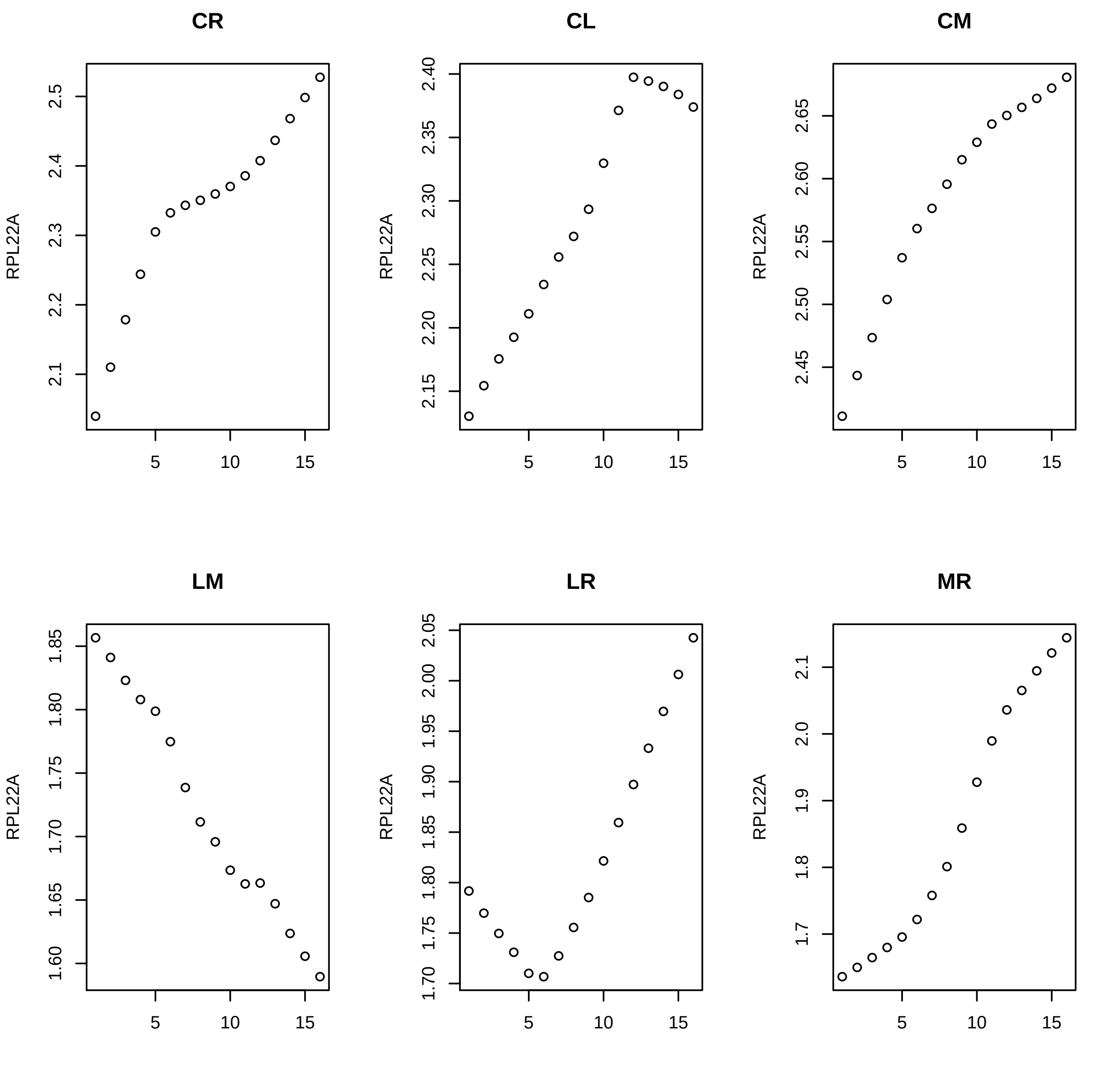

CR
CL
CM
LM
LR
MR
RPL22A
2.1
2.2
2.3
2.4
2.5
2.15
2.20
2.25
2.30
2.35
2.40
2.45
2.50
2.55
2.60
2.65
1.60
1.65
1.70
1.75
1.80
1.85
1.90
1.95
2.00
2.05
1.7
1.8
1.9
2.0
5
10
15

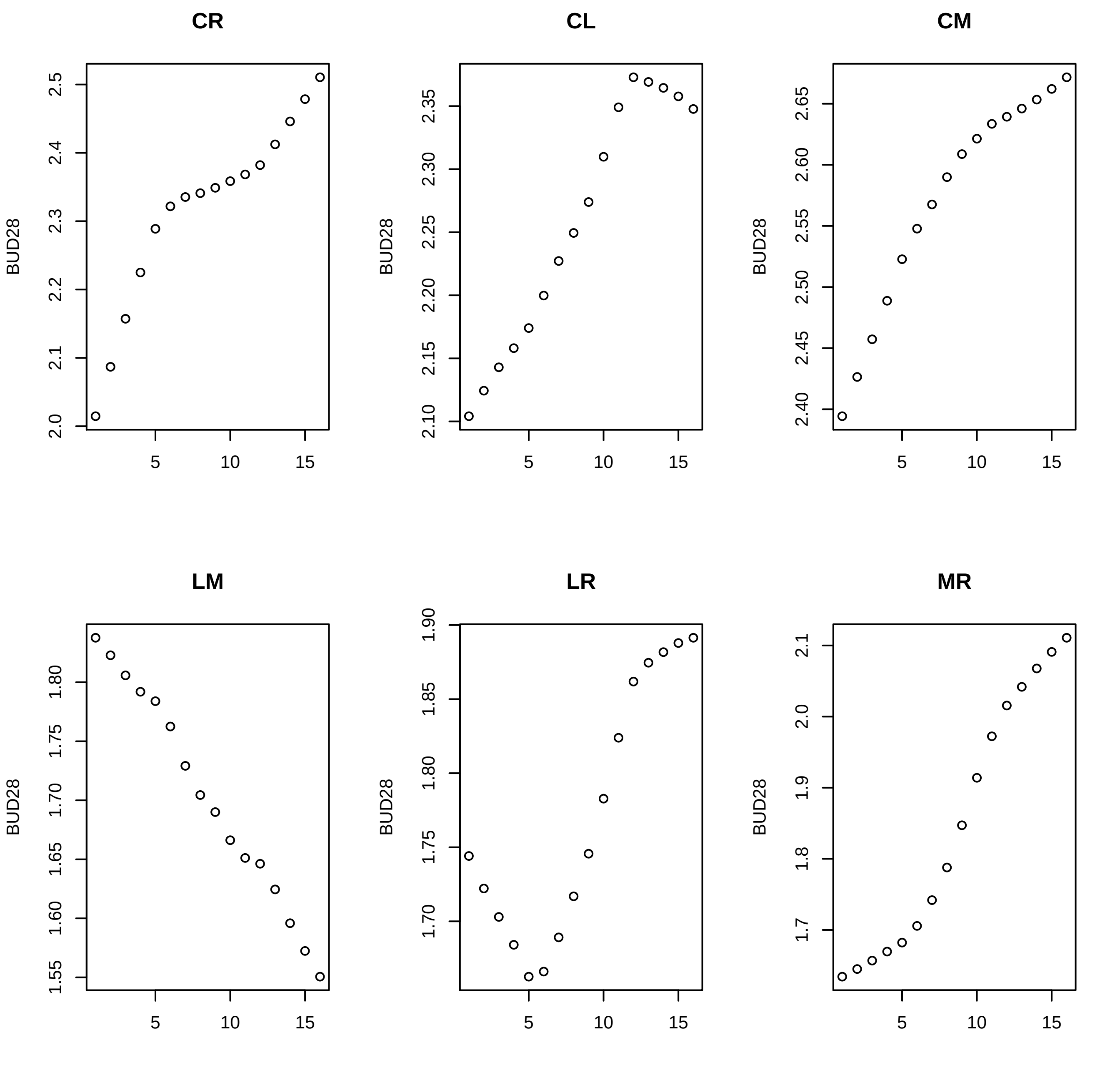
CR
CL
CM
LM
LR
MR
BUD28
5
10
15
2.0
2.1
2.2
2.3
2.4
2.5
2.10
2.15
2.20
2.25
2.30
2.35
2.40
2.45
2.50
2.55
2.60
2.65
1.55
1.60
1.65
1.70
1.75
1.80
1.85
1.90
1.7
1.8
1.9
2.0
2.1

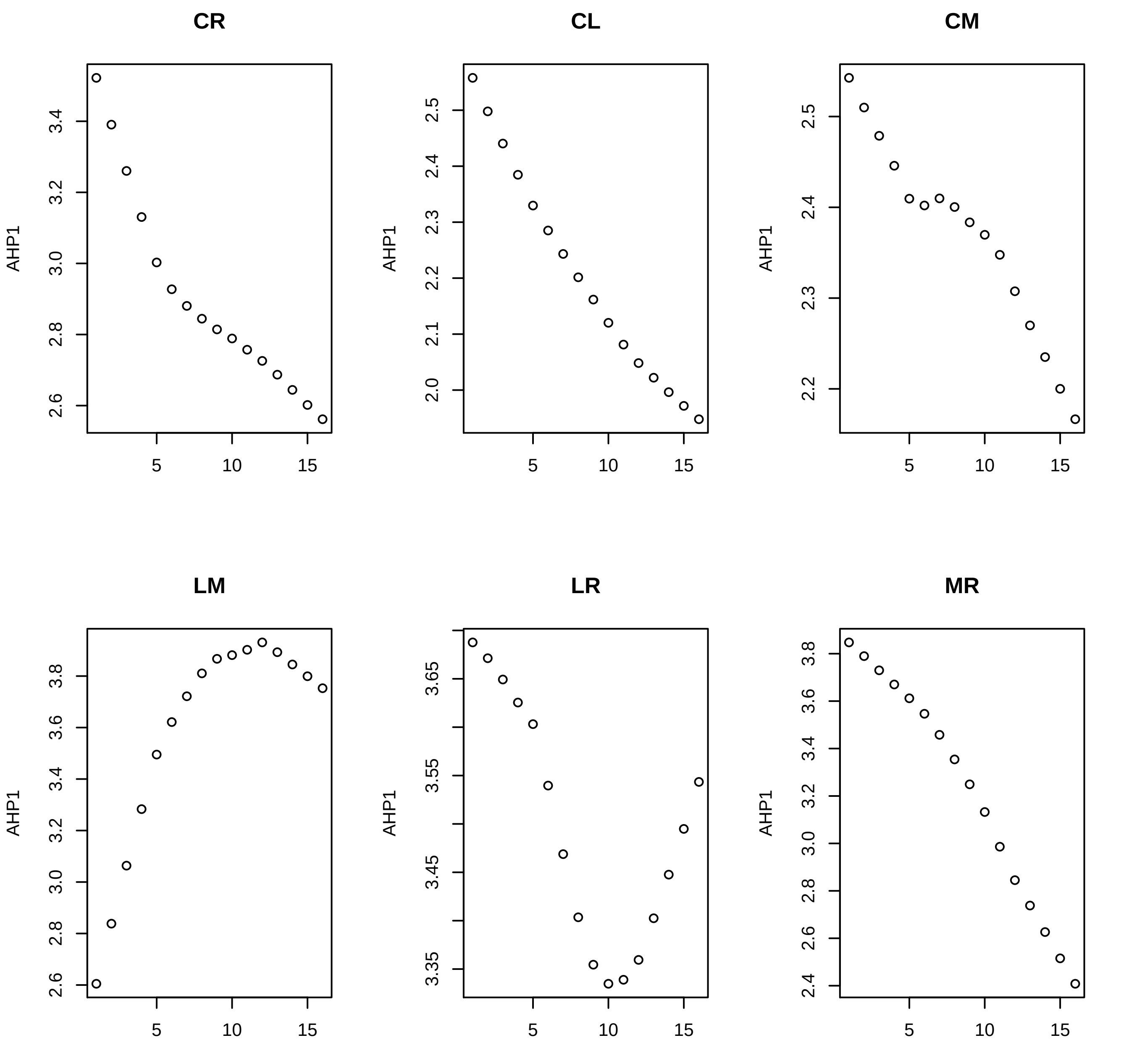
CR
CL
CM
LM
LR
MR
AHP1
5
10
15

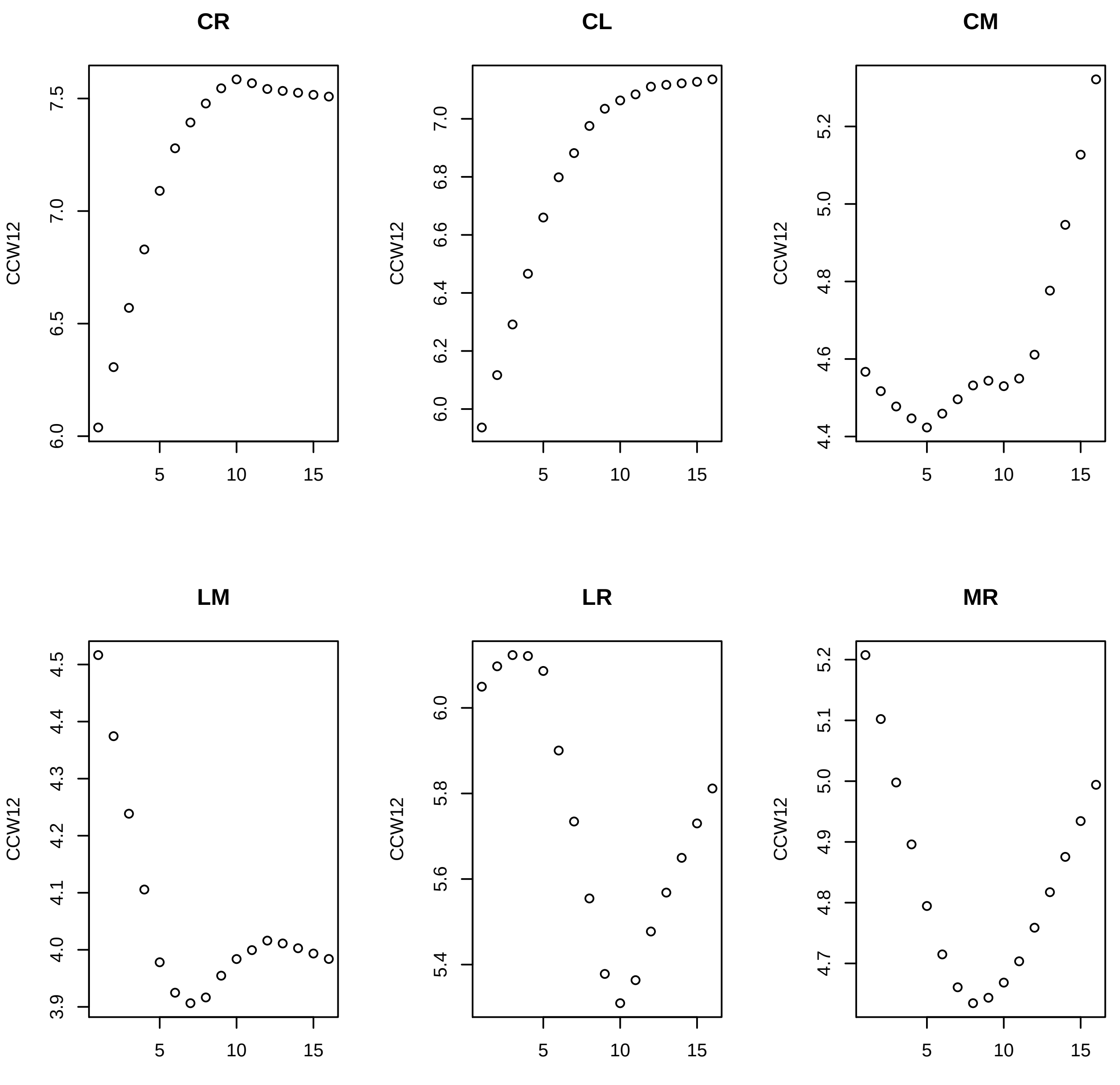

CR
CL
CM
LM
LR
MR
CCW12

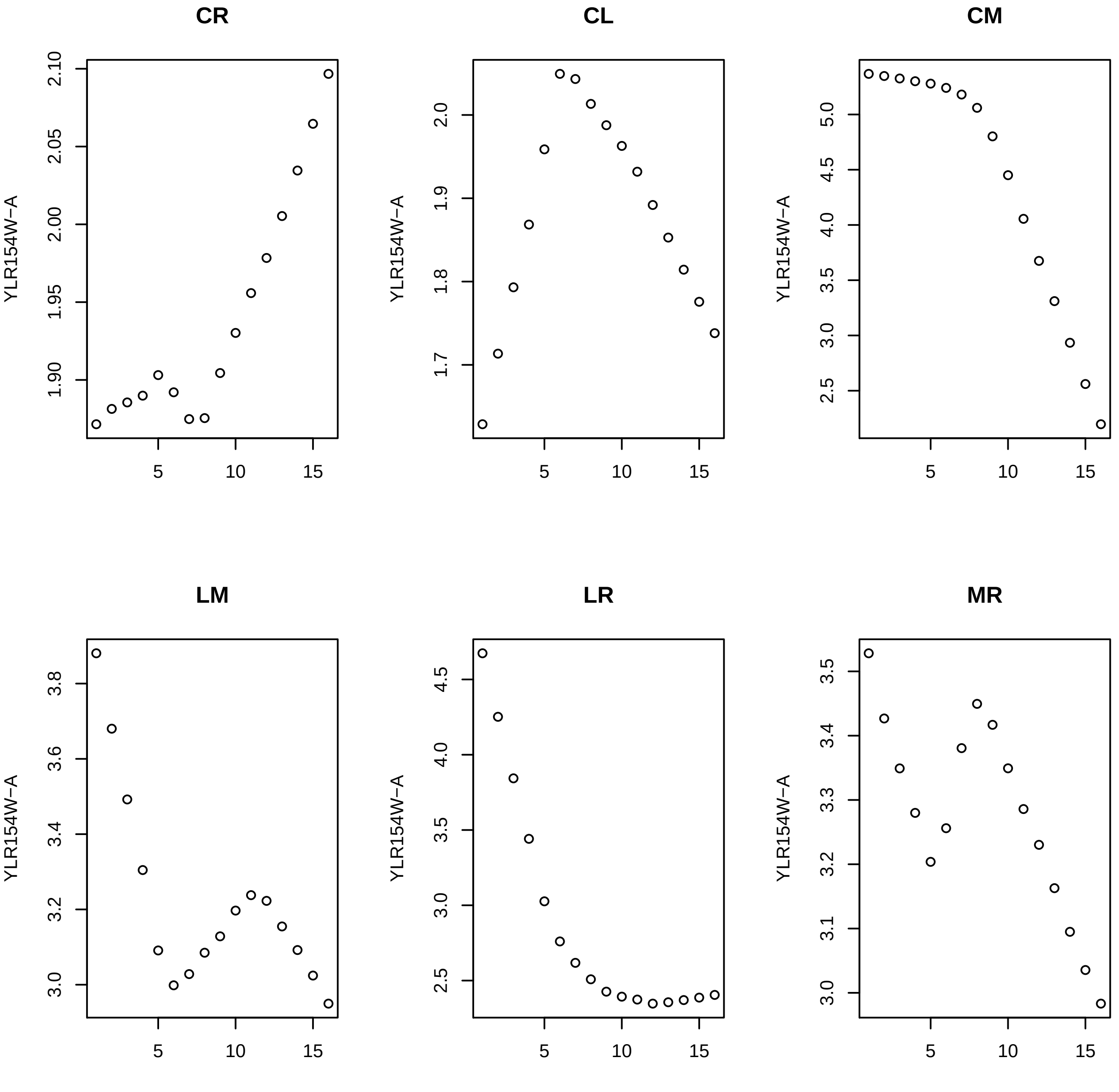
CR
CL
CM
LM
LR
MR
YLR154W−A
2.10
2.05
2.00
1.95
1.90
2.0
1.9
1.8
1.7
5.0
4.5
4.0
3.5
3.0
2.5
3.8
3.6
3.4
3.2
3.0
4.5
4.0
3.5
3.0
2.5
3.5
3.4
3.3
3.2
3.1
3.0
5
10
15

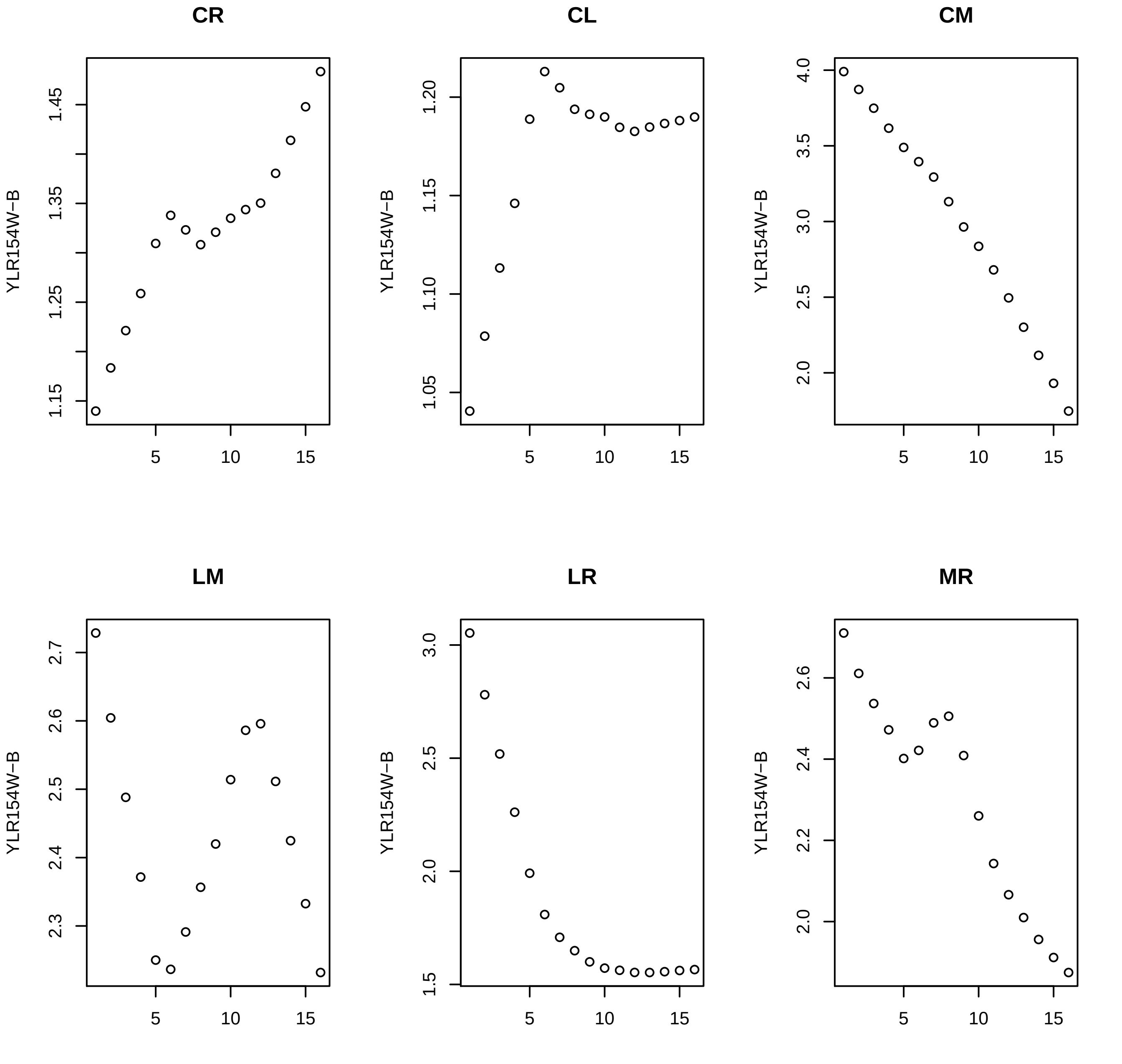
CR
CL
CM
LM
LR
MR
YLR154W−B

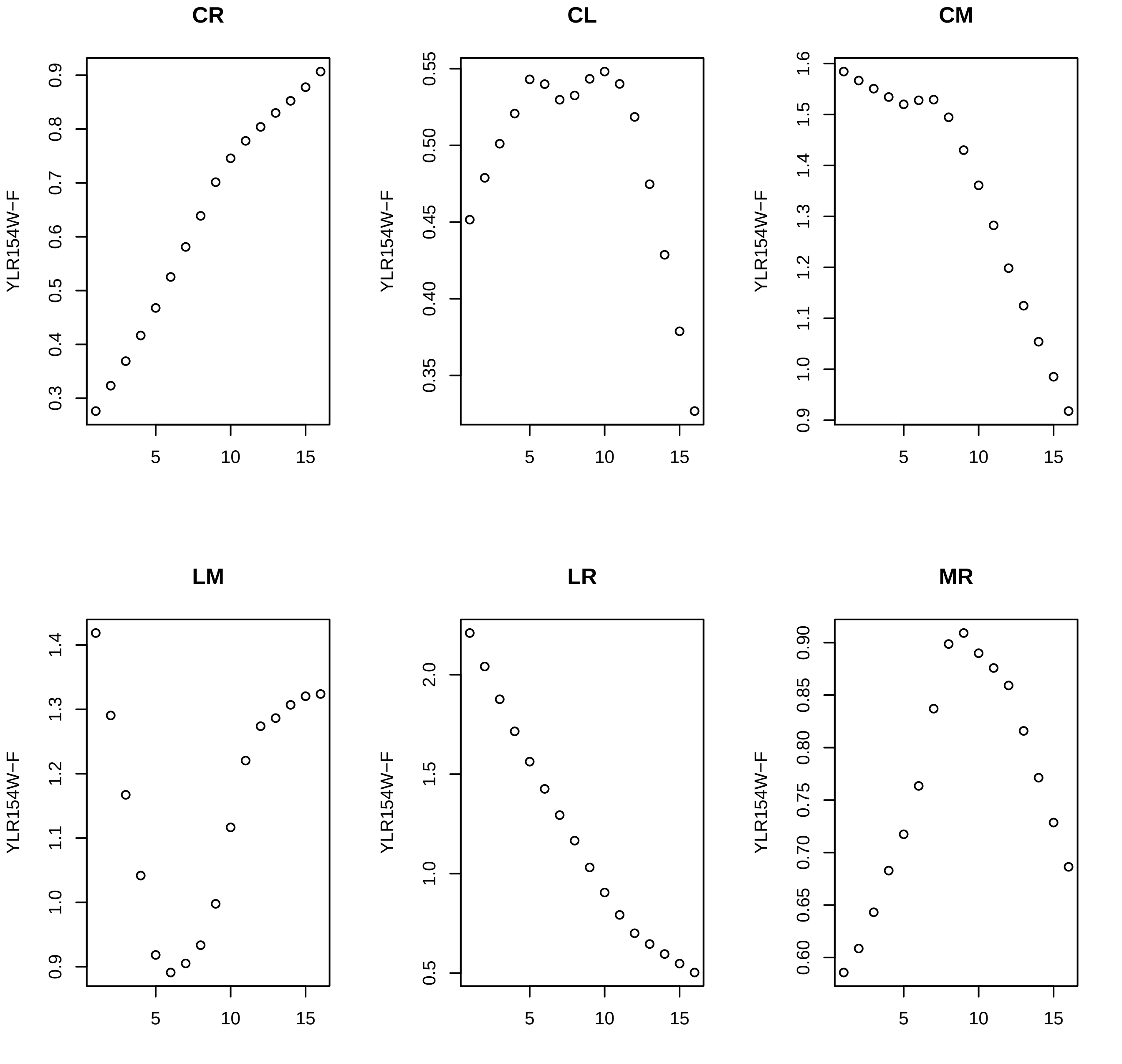
CR
CL
CM
LM
LR
MR
YLR154W−F
5
10
15

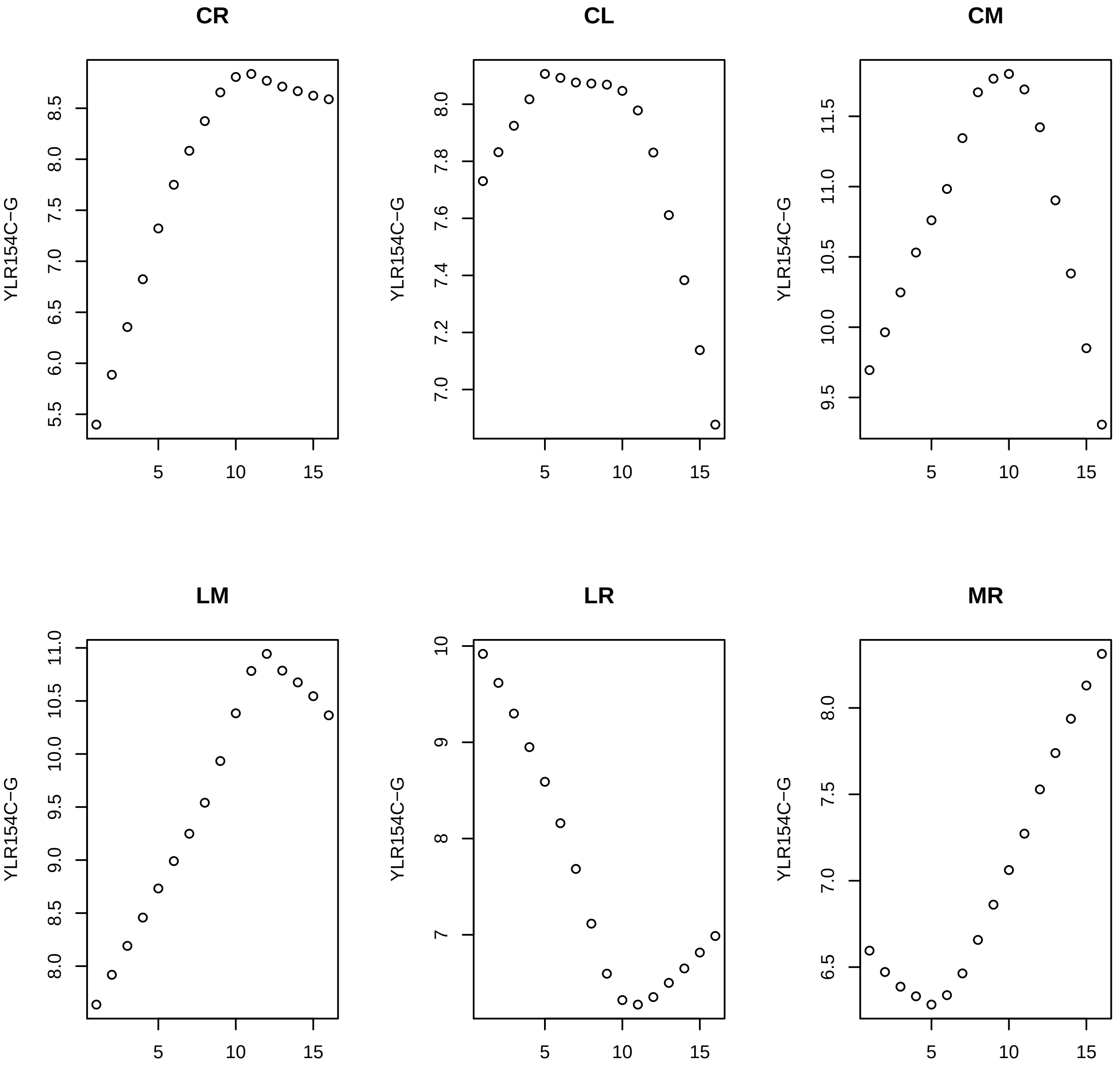
CR
CL
CM
LM
LR
MR
YLR154C−G
5
10
15
5.5
6.0
6.5
7.0
7.5
8.0
8.5
7.0
7.2
7.4
7.6
7.8
8.0
9.5
10.0
10.5
11.0
11.5
9.0
9
10
6.5

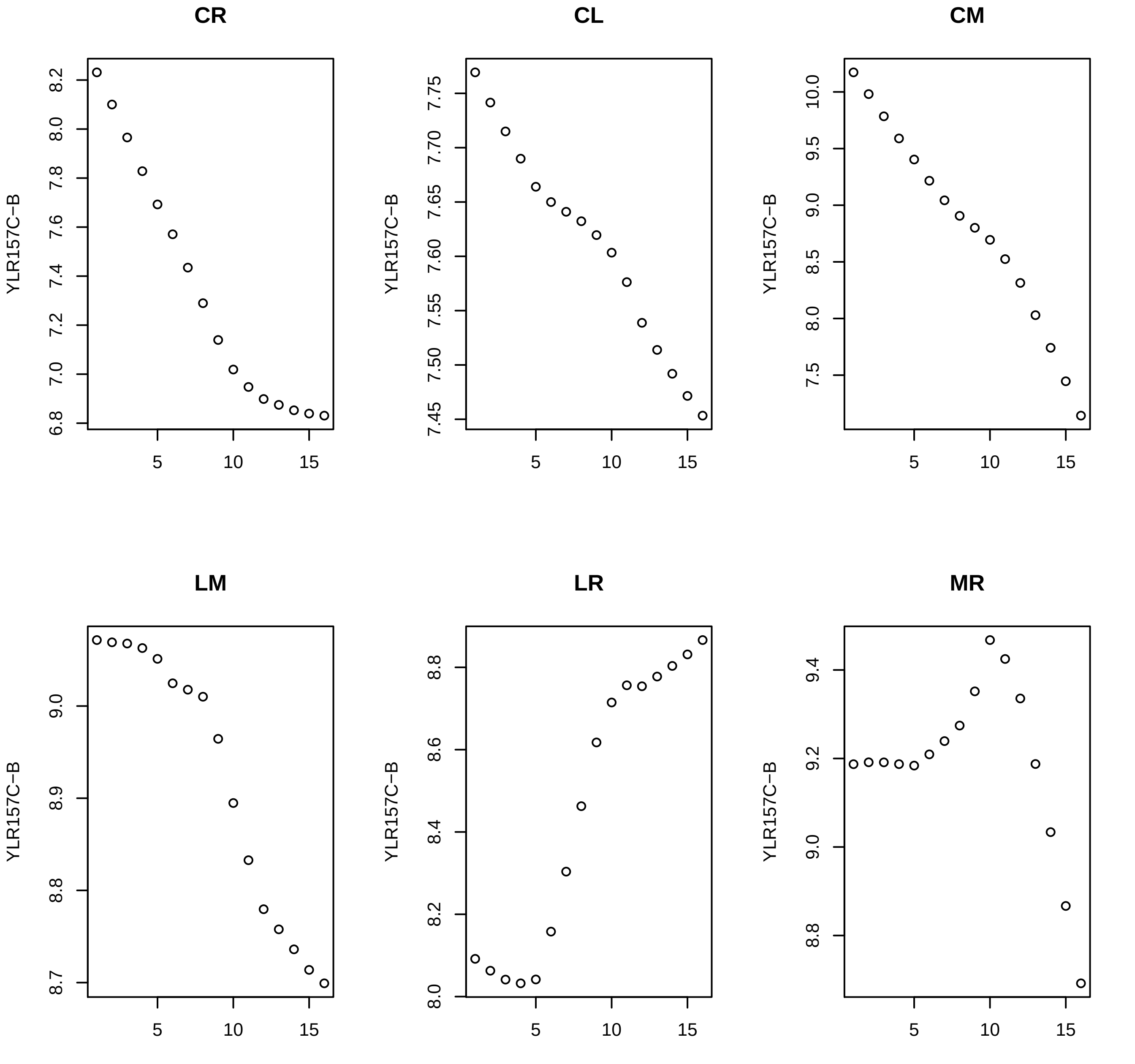
CR
CL
CM
LM
LR
MR
YLR157C−B
5
10
15

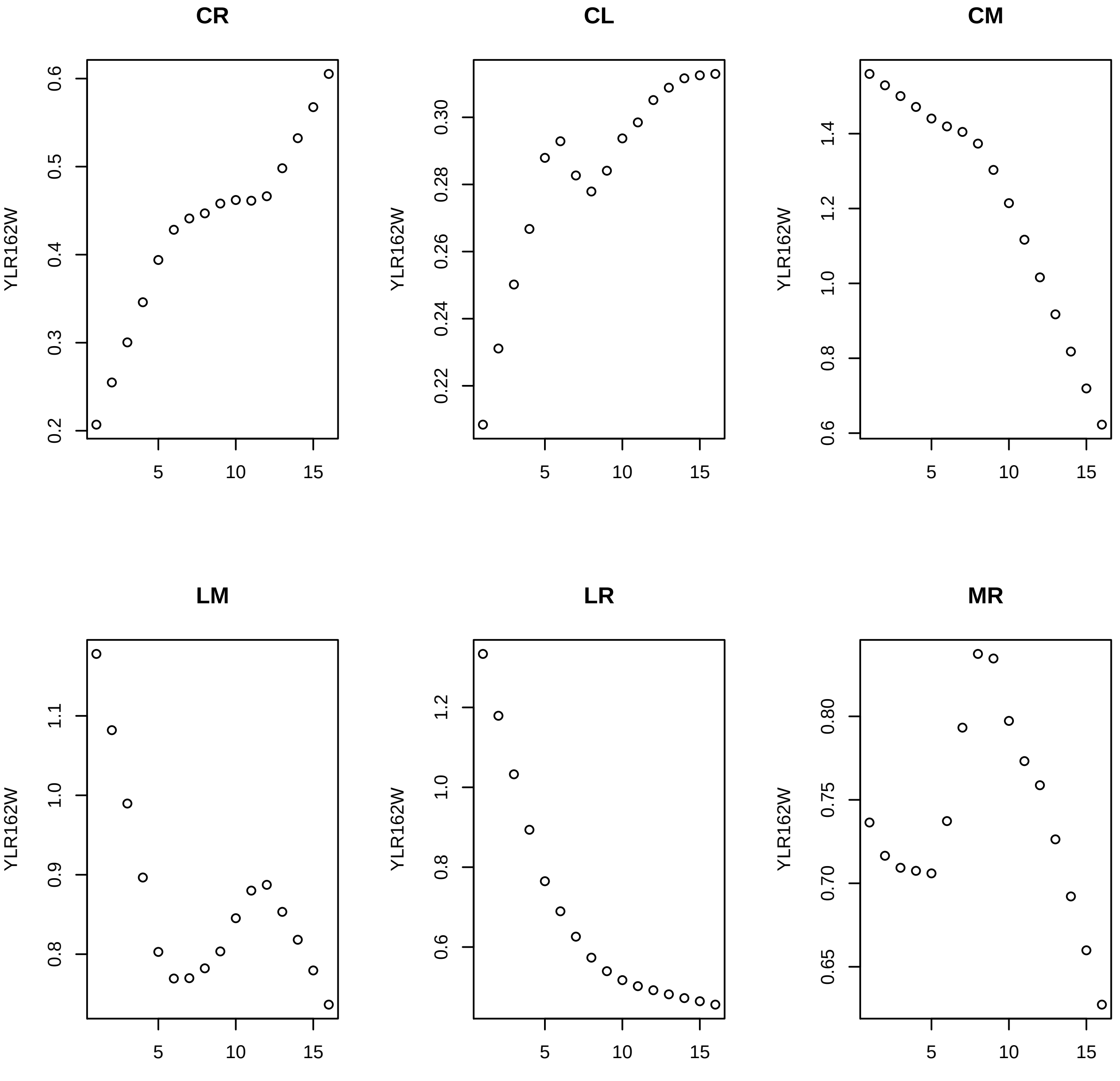
CR
YLR162W
0.2
0.3
0.4
0.5
0.6
5
10
15
CL
YLR162W
0.22
0.24
0.26
0.28
0.30
5
10
15
CM
YLR162W
0.6
0.8
1.0
1.2
1.4
5
10
15
LM
YLR162W
0.8
0.9
1.0
1.1
5
10
15
LR
YLR162W
0.6
0.8
1.0
1.2
5
10
15
MR
YLR162W
0.65
0.70
0.75
0.80
5
10
15

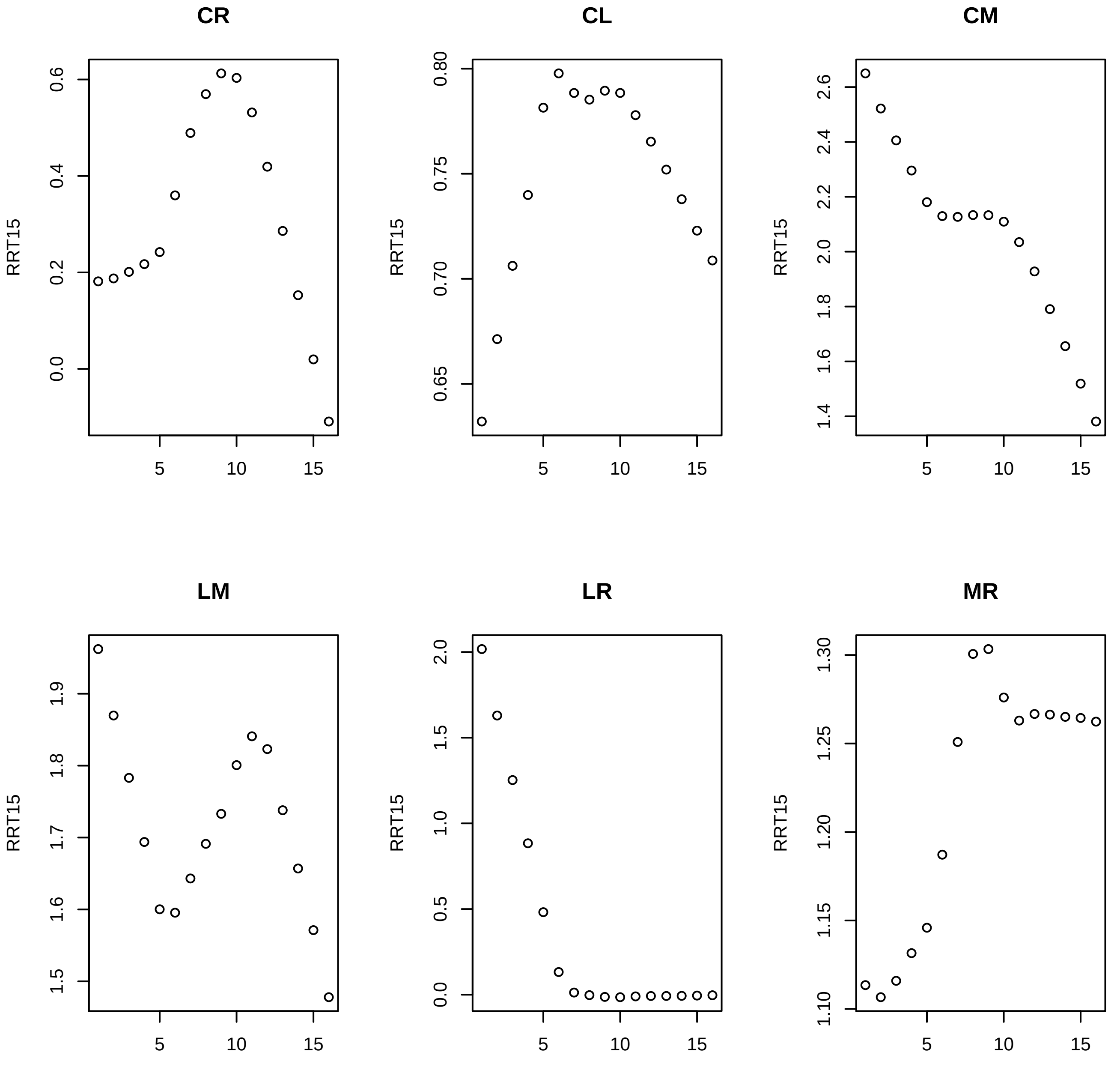

CR
CL
CM
LM
LR
MR
RRT15
0.0
0.2
0.4
0.6
0.65
0.70
0.75
0.80
1.4
1.6
1.8
2.0
2.2
2.4
2.6
1.5
1.6
1.7
1.8
1.9
0.0
0.5
1.0
1.5
2.0
1.10
1.15
1.20
1.25
1.30
5
10
15

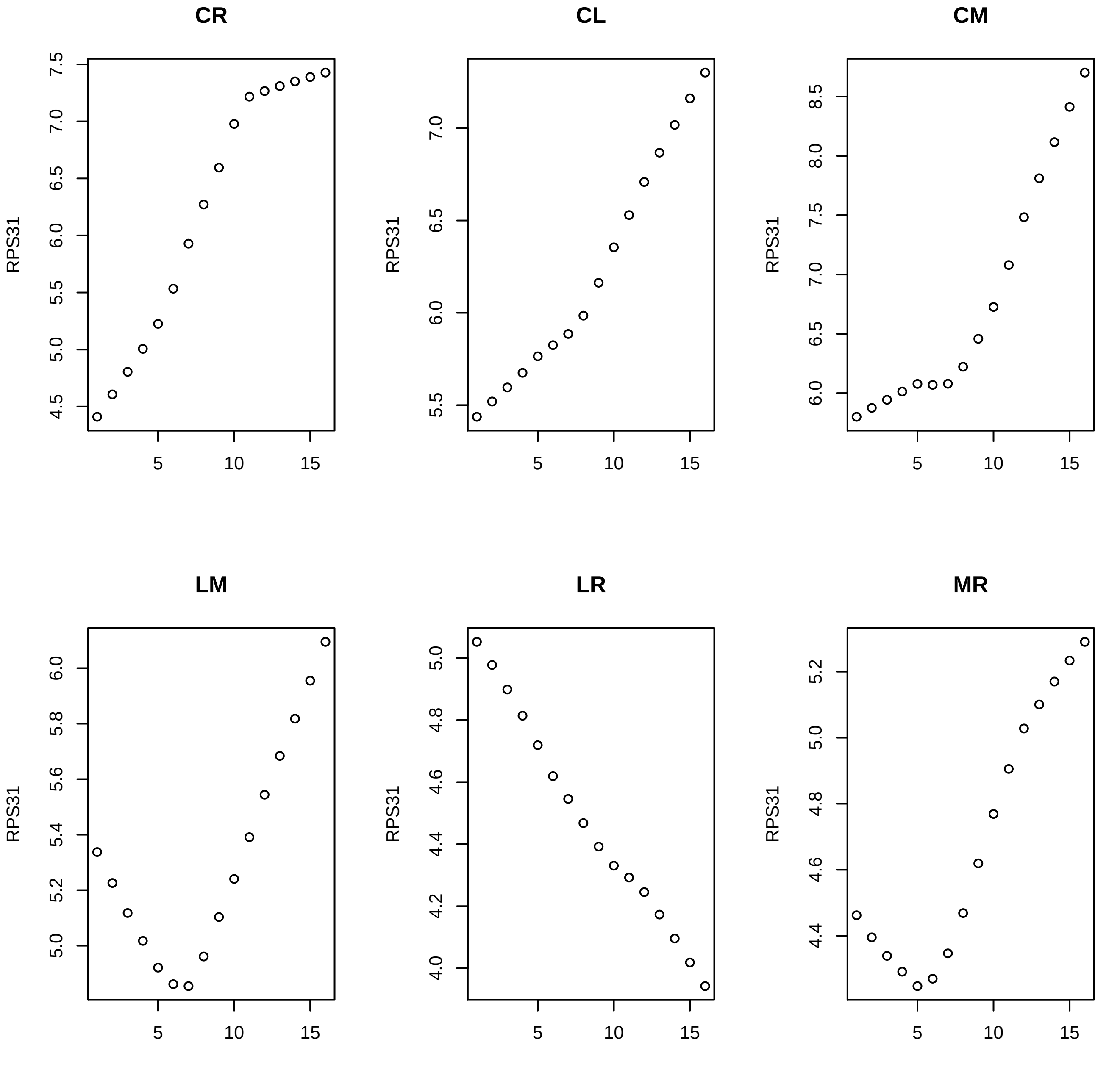
CR
CL
CM
LM
LR
MR
RPS31

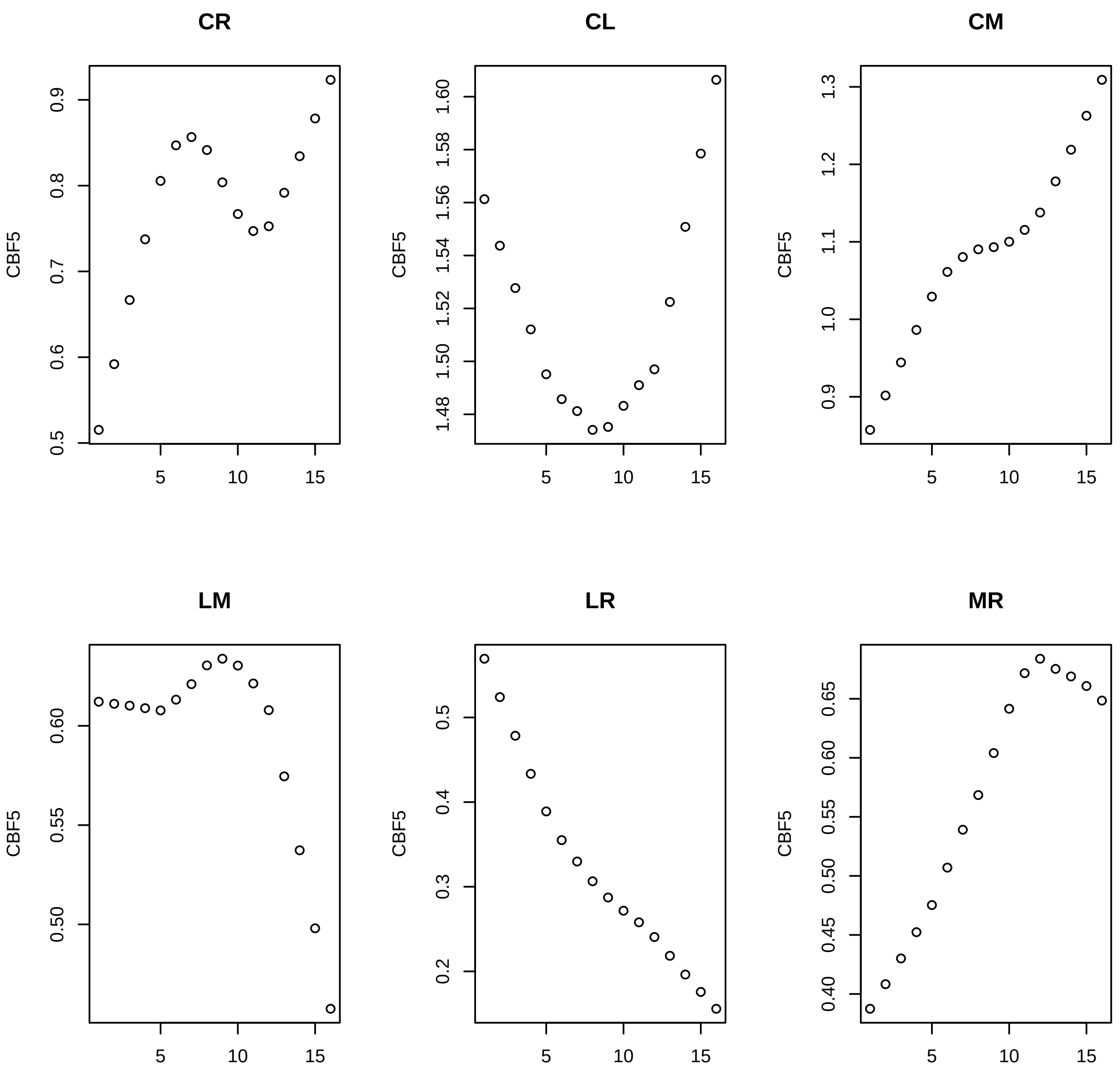
CR
CL
CM
LM
LR
MR
CBF5
5
10
15
0.5
0.6
0.7
0.8
0.9
1.48
1.50
1.52
1.54
1.56
1.58
1.60
0.9
1.0
1.1
1.2
1.3
0.50
0.55
0.60
0.2
0.3
0.4
0.5
0.40
0.45
0.50
0.55
0.60
0.65

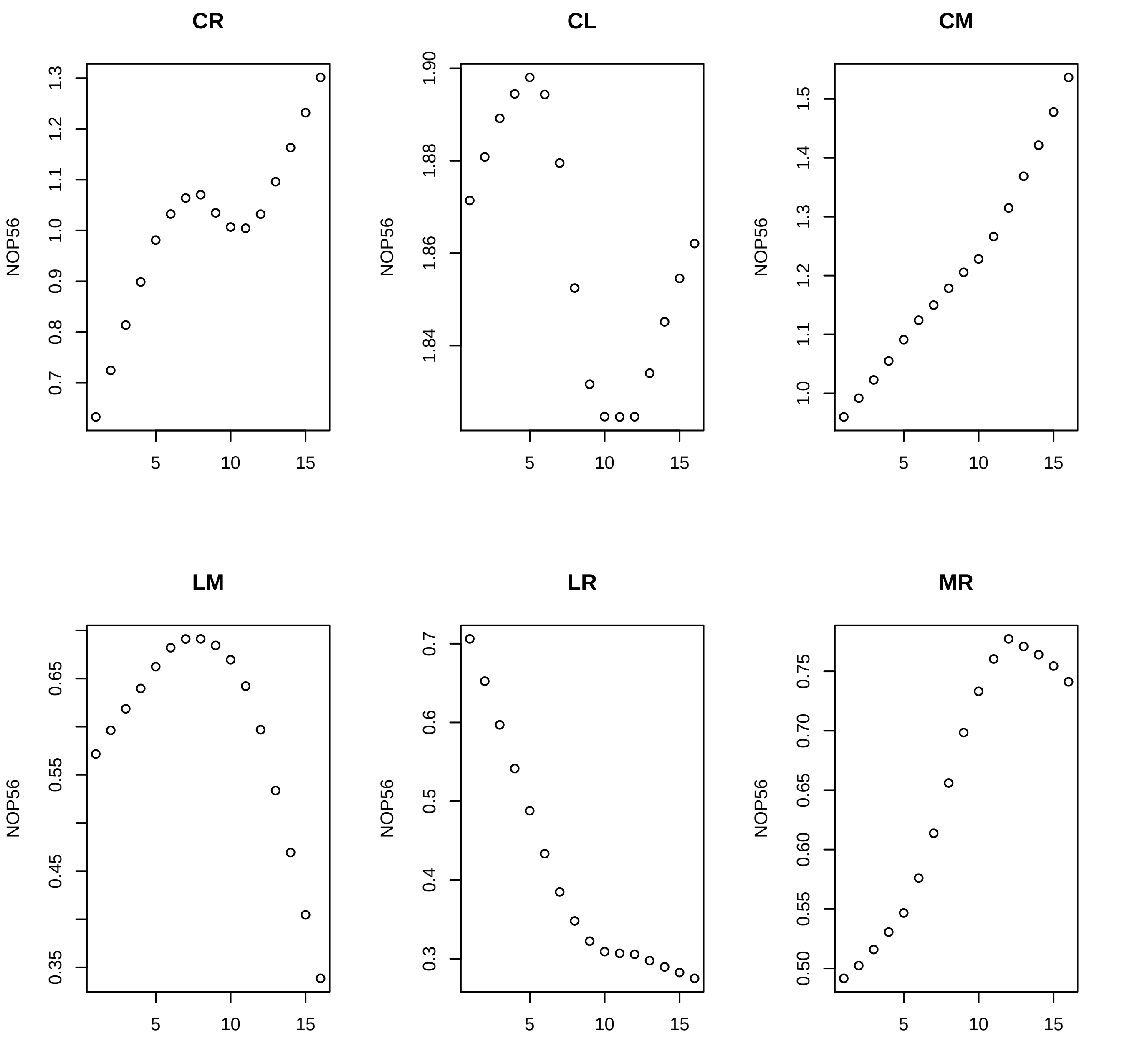

CR
CL
CM
LM
LR
MR
NOP56
1.3
1.2
1.1
1.0
0.9
0.8
0.7
1.90
1.88
1.86
1.84
1.5
1.4
1.3
1.2
1.1
1.0
0.65
0.55
0.45
0.35
0.7
0.6
0.5
0.4
0.3
0.75
0.70
0.65
0.60
0.55
0.50
5
10
15

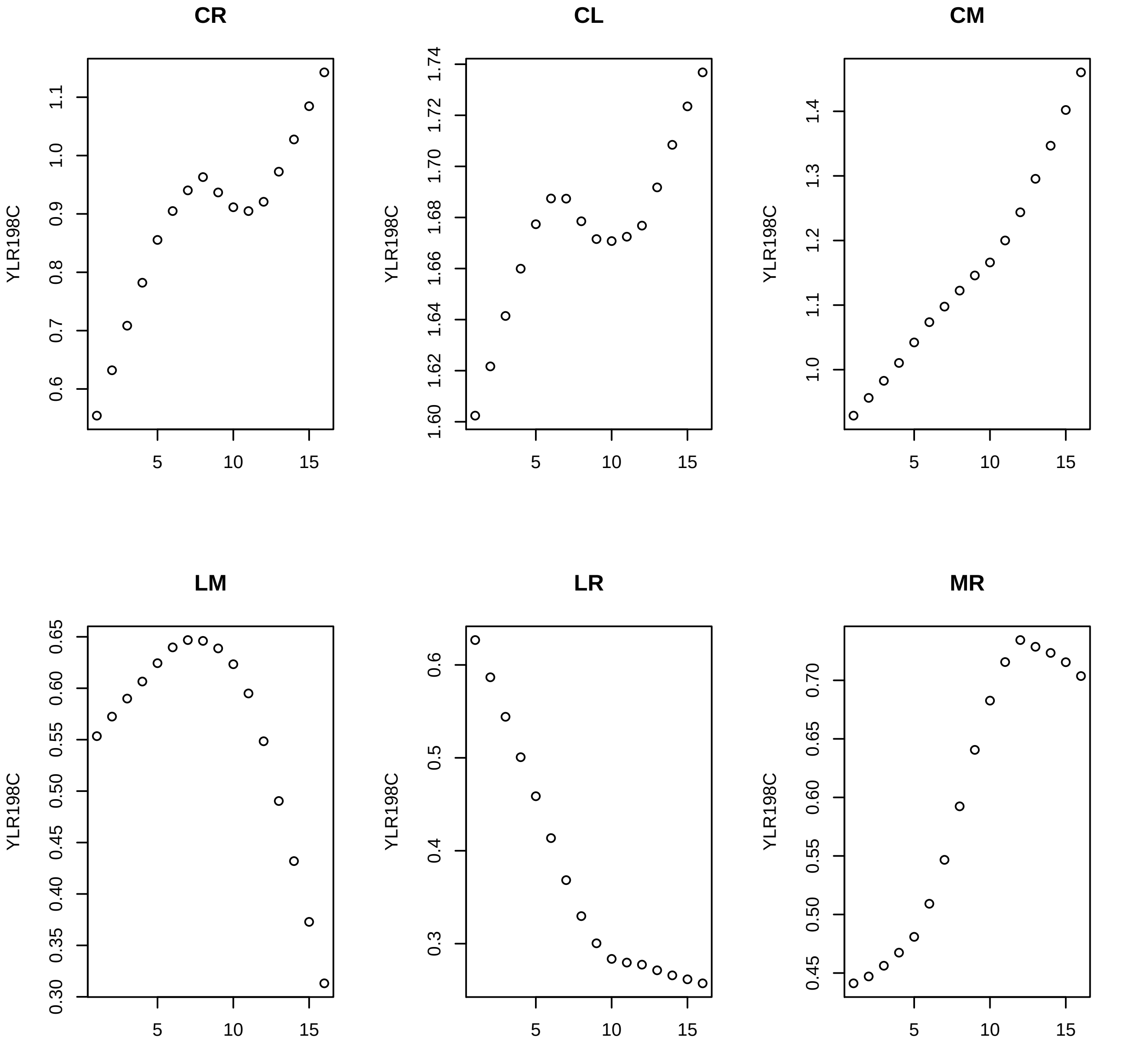

CR
CL
CM
LM
LR
MR
YLR198C
5
10
15

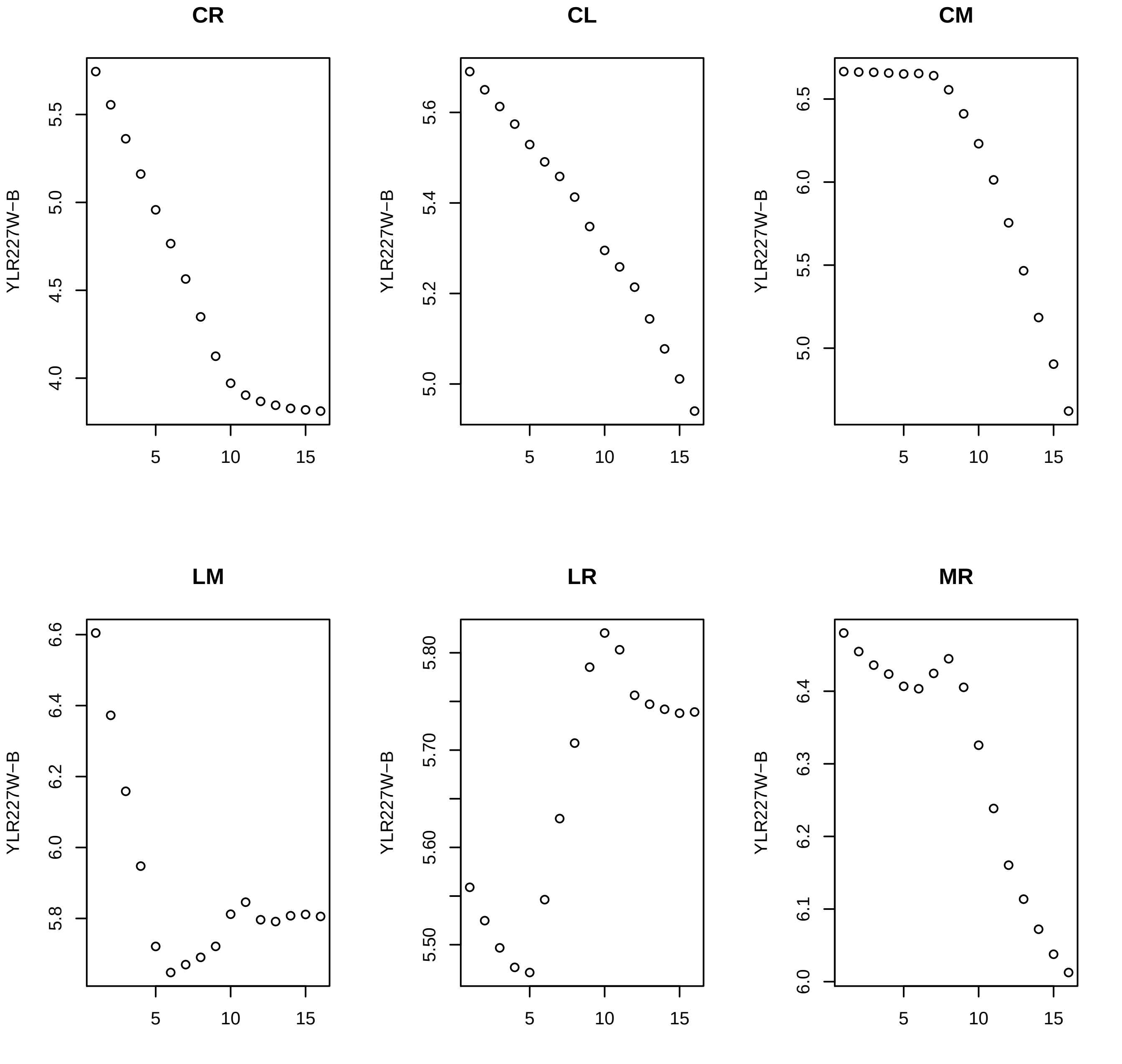
CR
CL
CM
LM
LR
MR
YLR227W−B

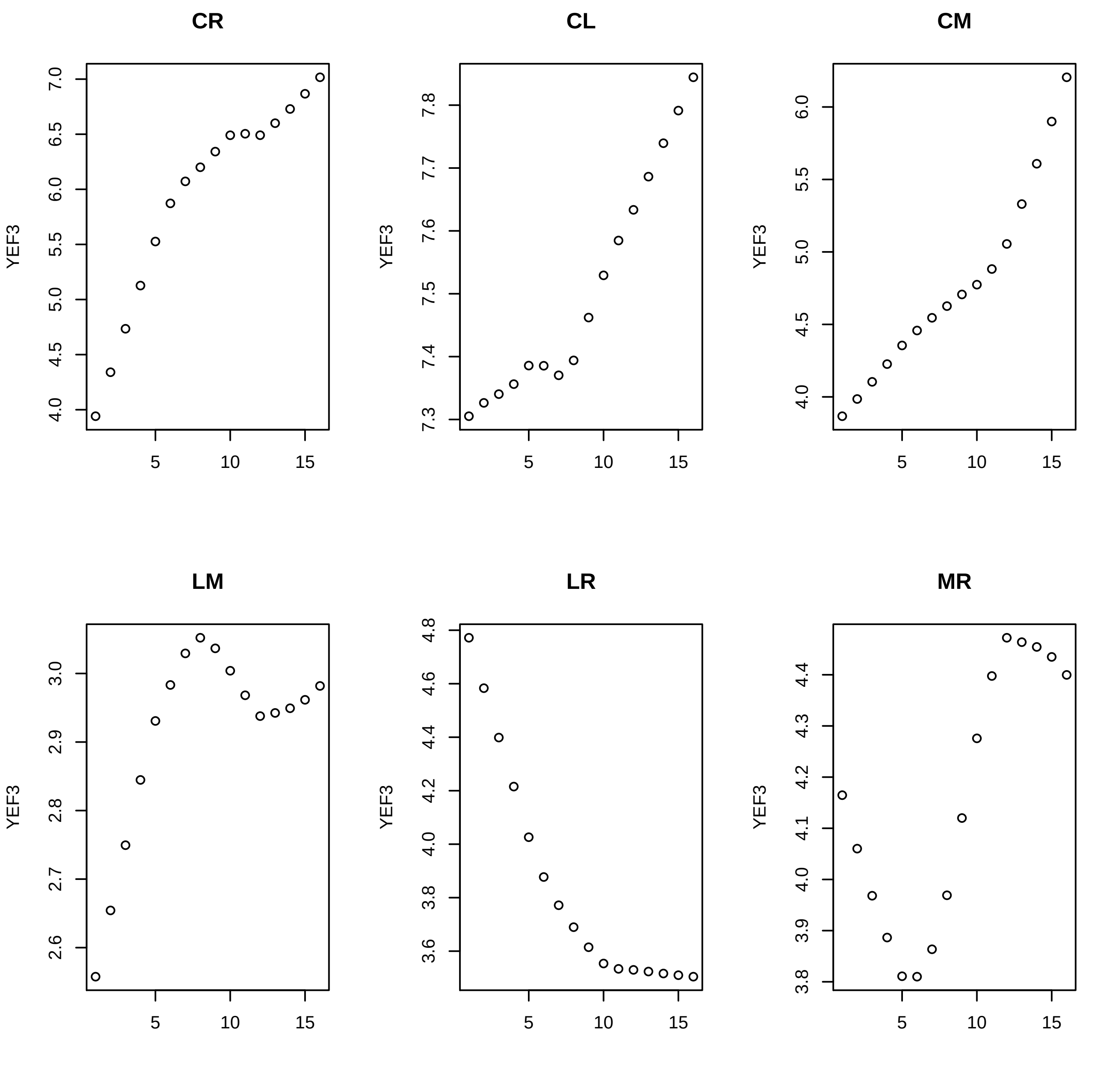
CR
CL
CM
LM
LR
MR
YEF3
5
10
15
4.0
4.5
5.0
5.5
6.0
6.5
7.0
7.3
7.4
7.5
7.6
7.7
7.8
2.6
2.7
2.8
2.9
3.0
3.6
3.8
4.2
4.4
4.6
4.8
3.9
4.1
4.3
4.4

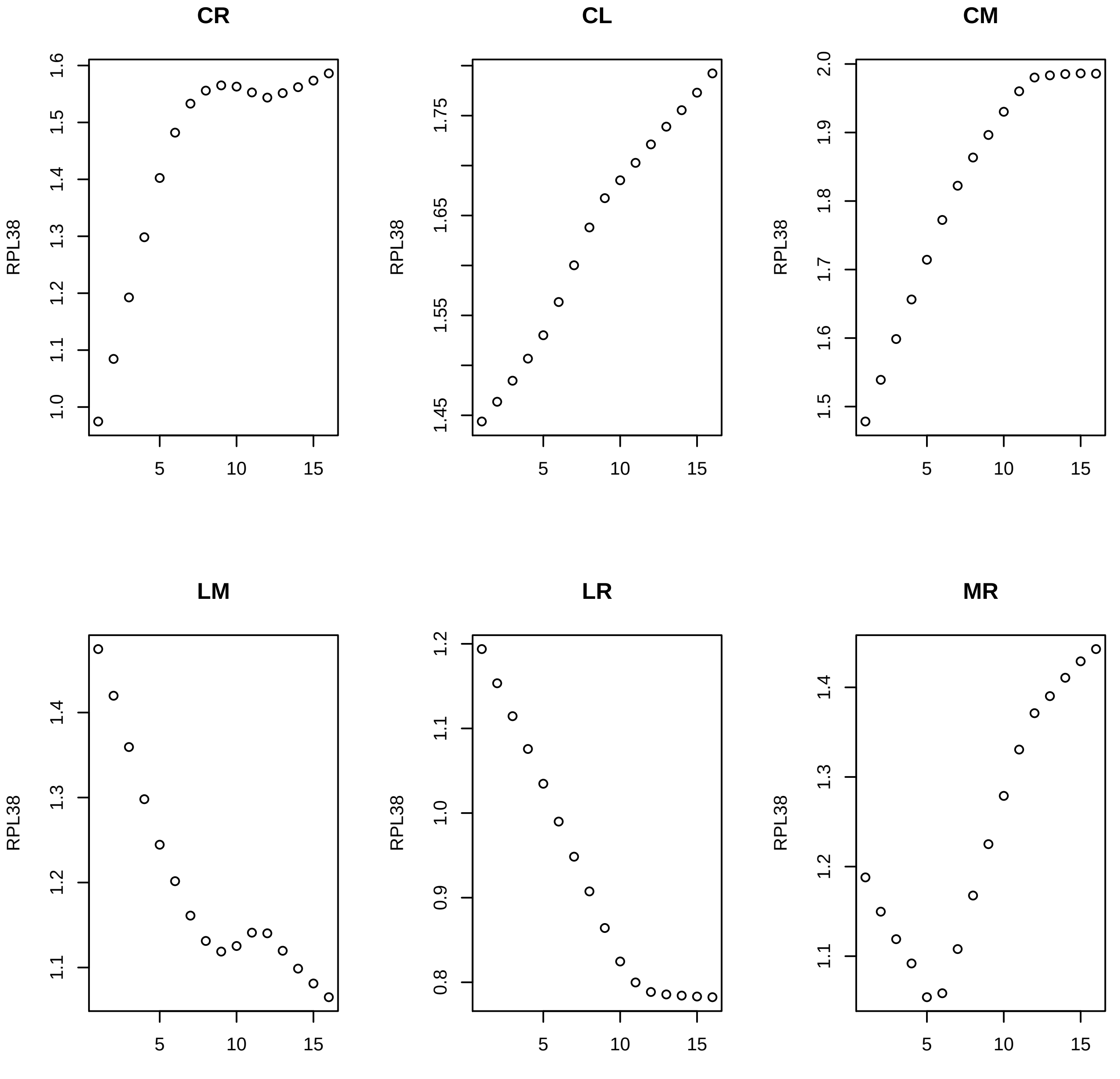
CR
CL
CM
LM
LR
MR
RPL38
1.0
1.1
1.2
1.3
1.4
1.5
1.6
1.45
1.55
1.65
1.75
1.5
1.6
1.7
1.8
1.9
2.0
1.1
1.2
1.3
1.4
0.8
0.9
1.0
1.1
1.2
1.1
1.2
1.3
1.4
5
10
15

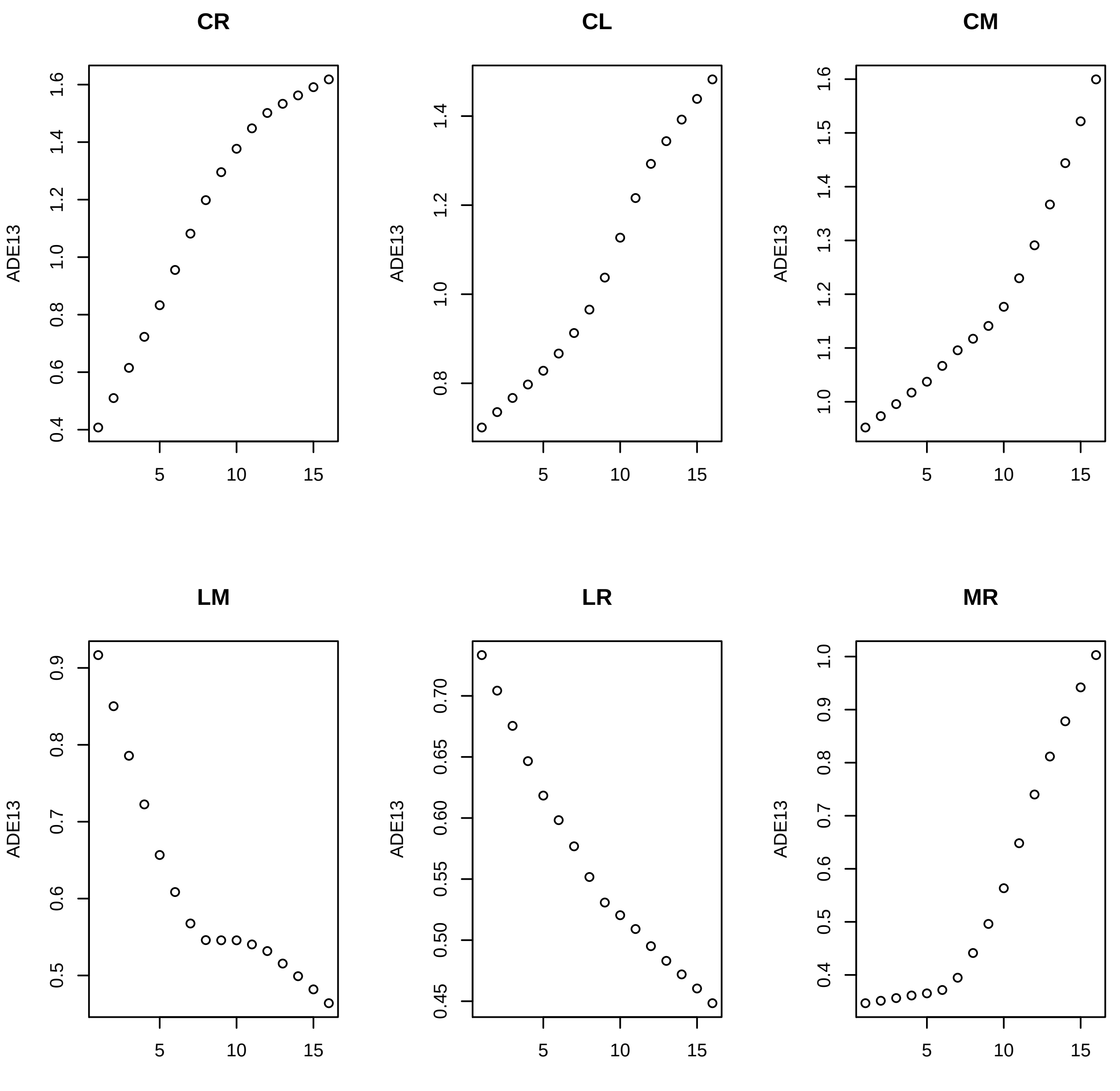

CR
CL
CM
LM
LR
MR
ADE13

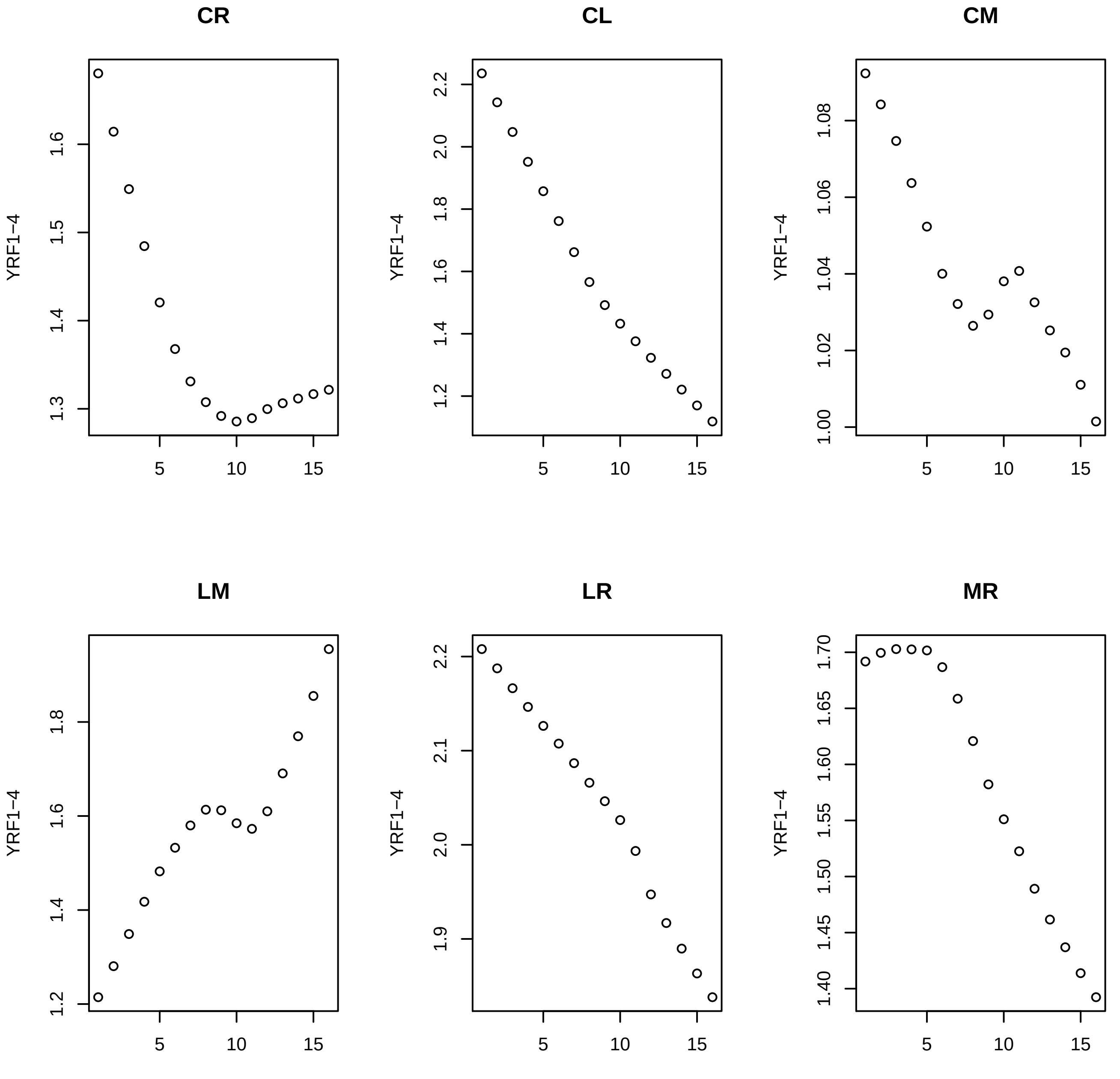

CR
CL
CM
LM
LR
MR
YRF1−4

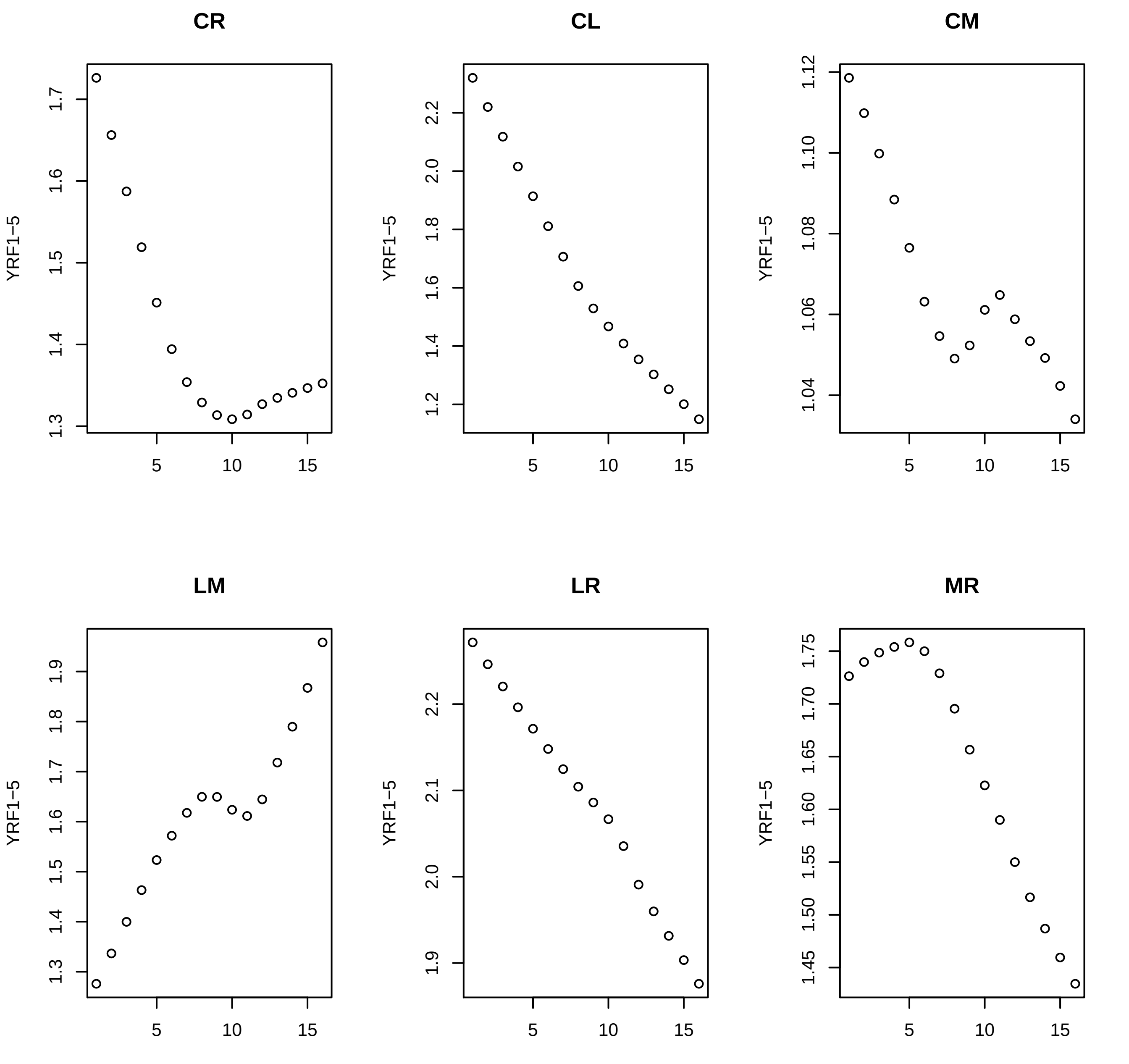
CR
CL
CM
LM
LR
MR
YRF1–5
5
10
15

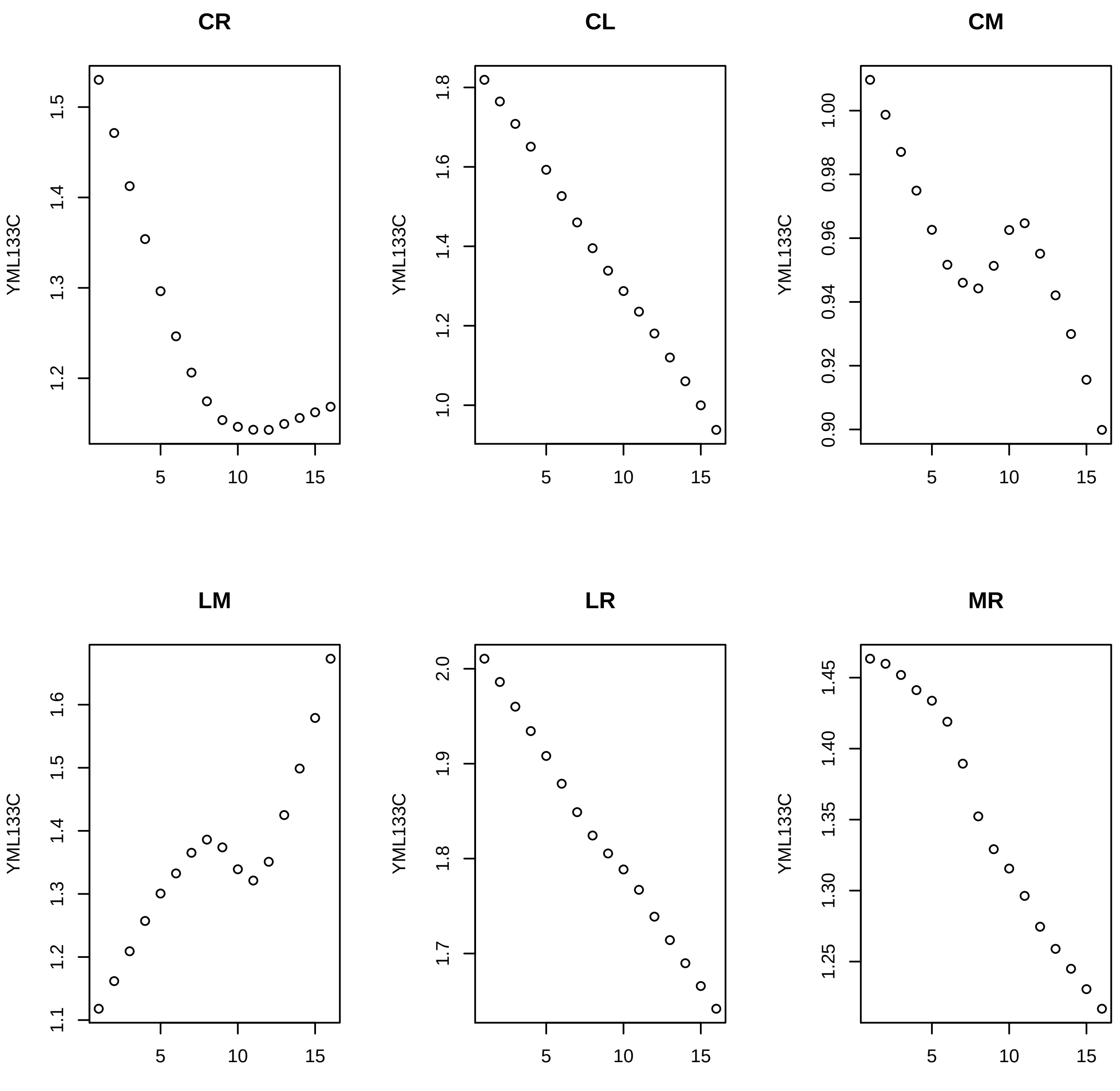
CR
CL
CM
LM
LR
MR
YML133C
5
10
15
1.2
1.3
1.4
1.5
1.0
1.2
1.4
1.6
1.8
0.90
0.92
0.94
0.96
0.98
1.00
1.1
1.6
1.7
1.8
1.9
2.0
1.25
1.30
1.35
1.40
1.45

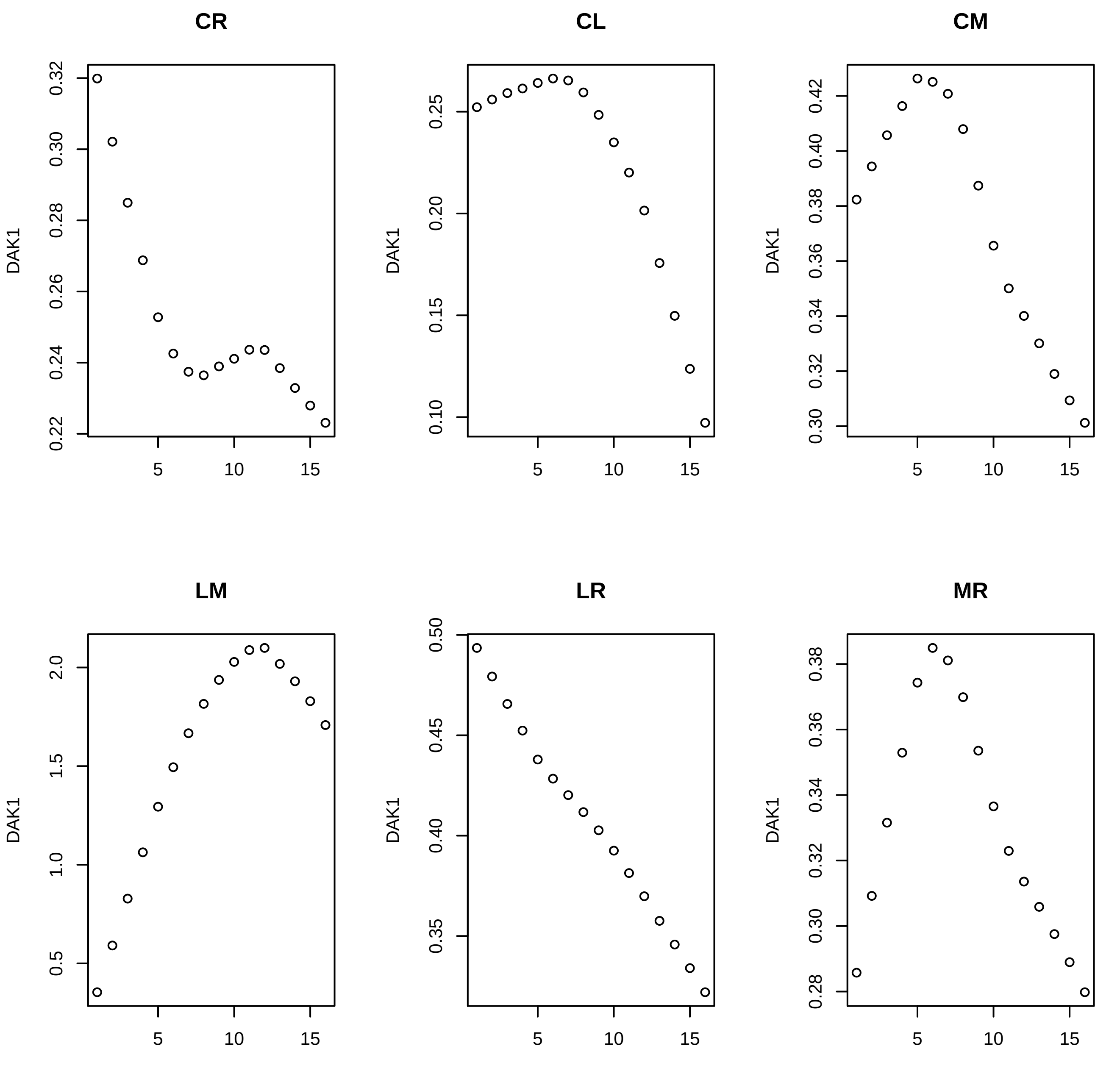
CR
CL
CM
LM
LR
MR
DAK1
0.32
0.30
0.28
0.26
0.24
0.22
0.25
0.20
0.15
0.10
0.42
0.40
0.38
0.36
0.34
0.32
0.30
2.0
1.5
1.0
0.5
0.50
0.45
0.40
0.35
0.38
0.36
0.34
0.32
0.30
0.28
5
10
15

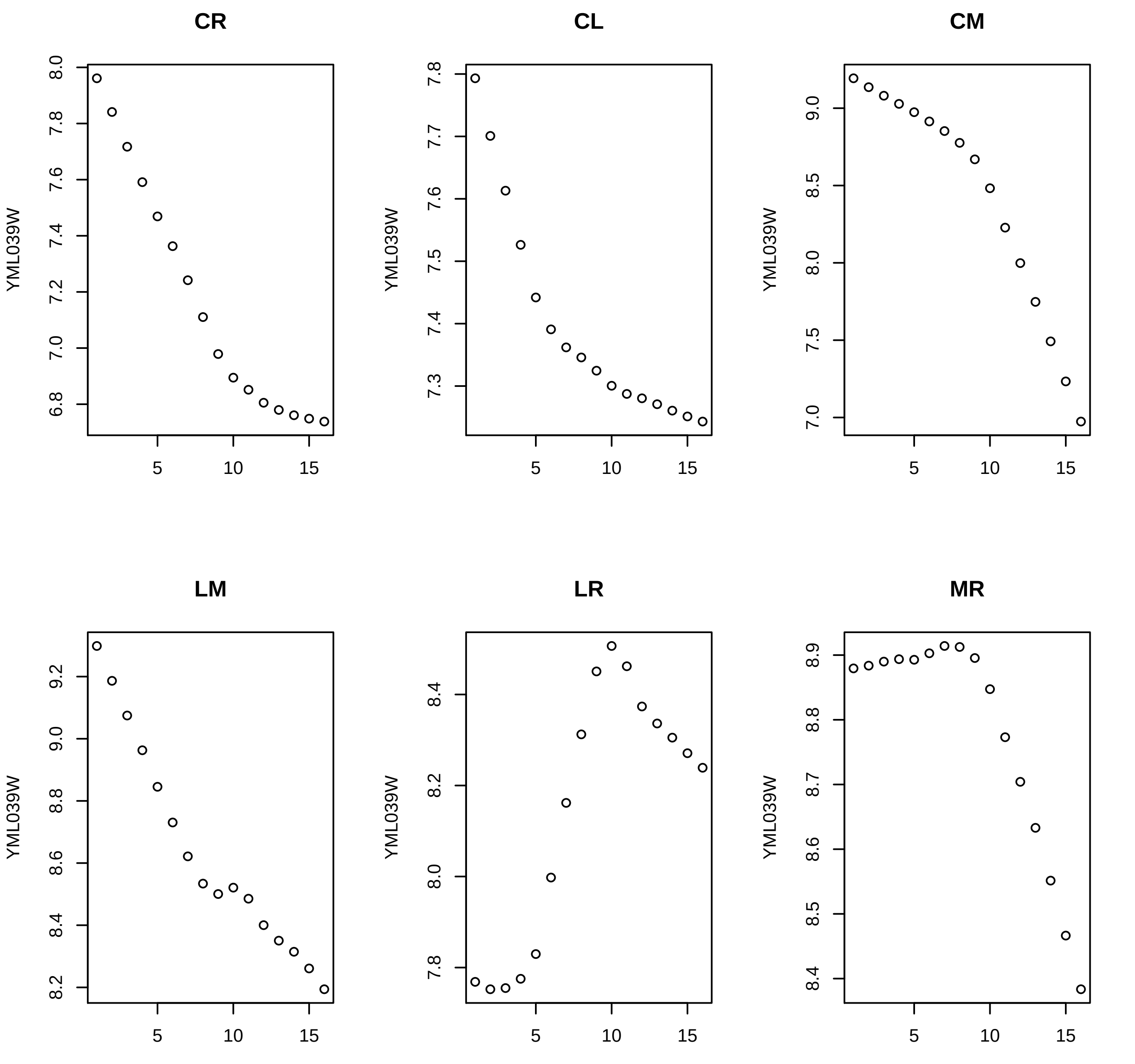
CR
CL
CM
LM
LR
MR
YML039W
5
10
15

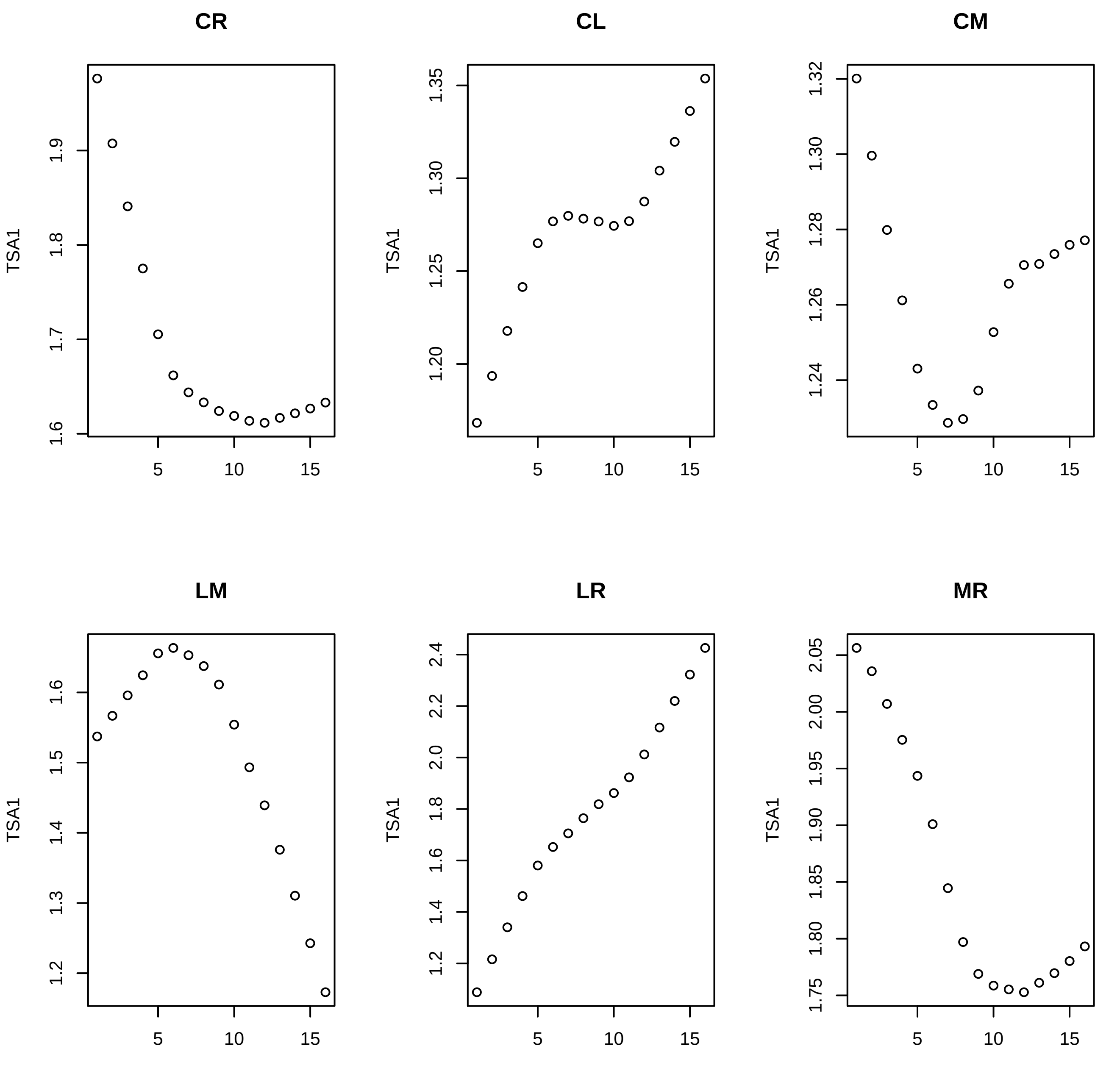
CR
CL
CM
LM
LR
MR
TSA1
1.6
1.7
1.8
1.9
1.20
1.25
1.30
1.35
1.24
1.26
1.28
1.30
1.32
1.2
1.3
1.4
1.5
1.6
1.2
1.4
1.6
1.8
2.0
2.2
2.4
1.75
1.80
1.85
1.90
1.95
2.00
2.05
5
10
15

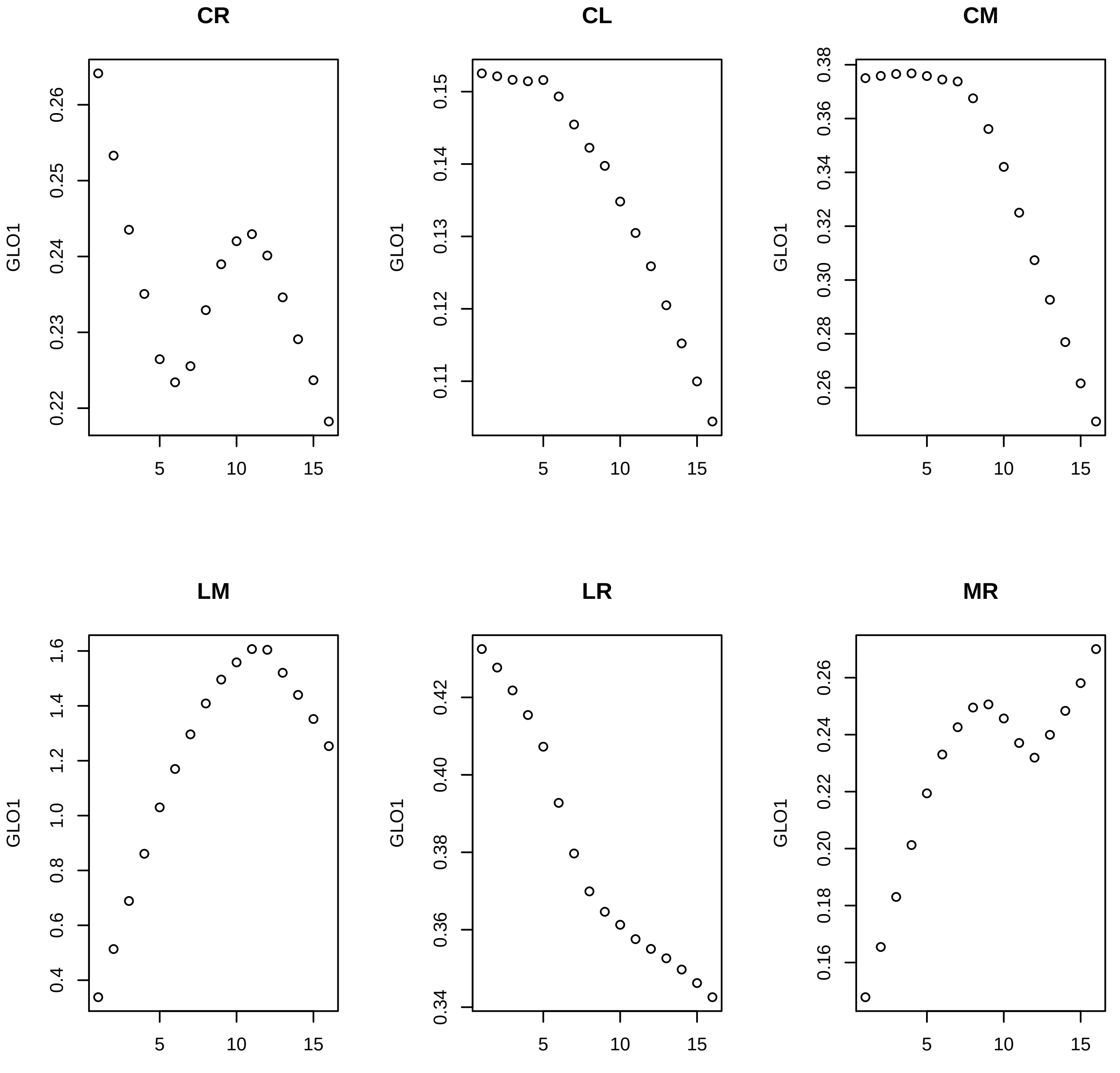
CR
CL
CM
LM
LR
MR
GLO1
0.22
0.23
0.24
0.25
0.26
0.11
0.12
0.13
0.14
0.15
0.26
0.28
0.30
0.32
0.34
0.36
0.38
0.4
0.6
0.8
1.0
1.2
1.4
1.6
0.34
0.36
0.38
0.40
0.42
0.16
0.18
0.20
0.22
0.24
0.26
5
10
15

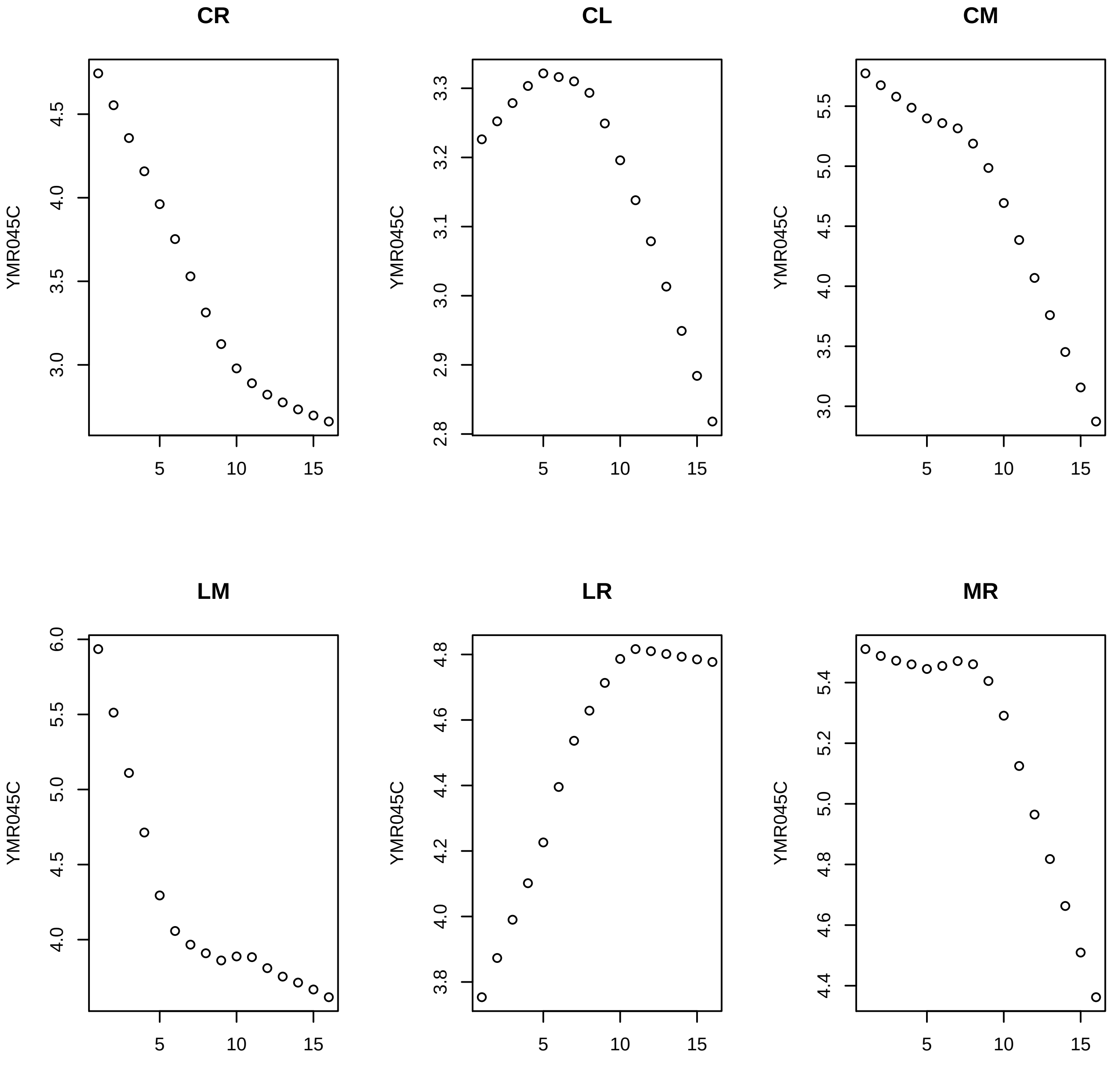

CR
YMR045C
4.5
4.0
3.5
3.0
5
10
15
CL
YMR045C
3.3
3.2
3.1
3.0
2.9
2.8
5
10
15
CM
YMR045C
5.5
5.0
4.5
4.0
3.5
3.0
5
10
15
LM
YMR045C
6.0
5.5
5.0
4.5
4.0
5
10
15
LR
YMR045C
4.8
4.6
4.4
4.2
4.0
3.8
5
10
15
MR
YMR045C
5.4
5.2
5.0
4.8
4.6
4.4
5
10
15

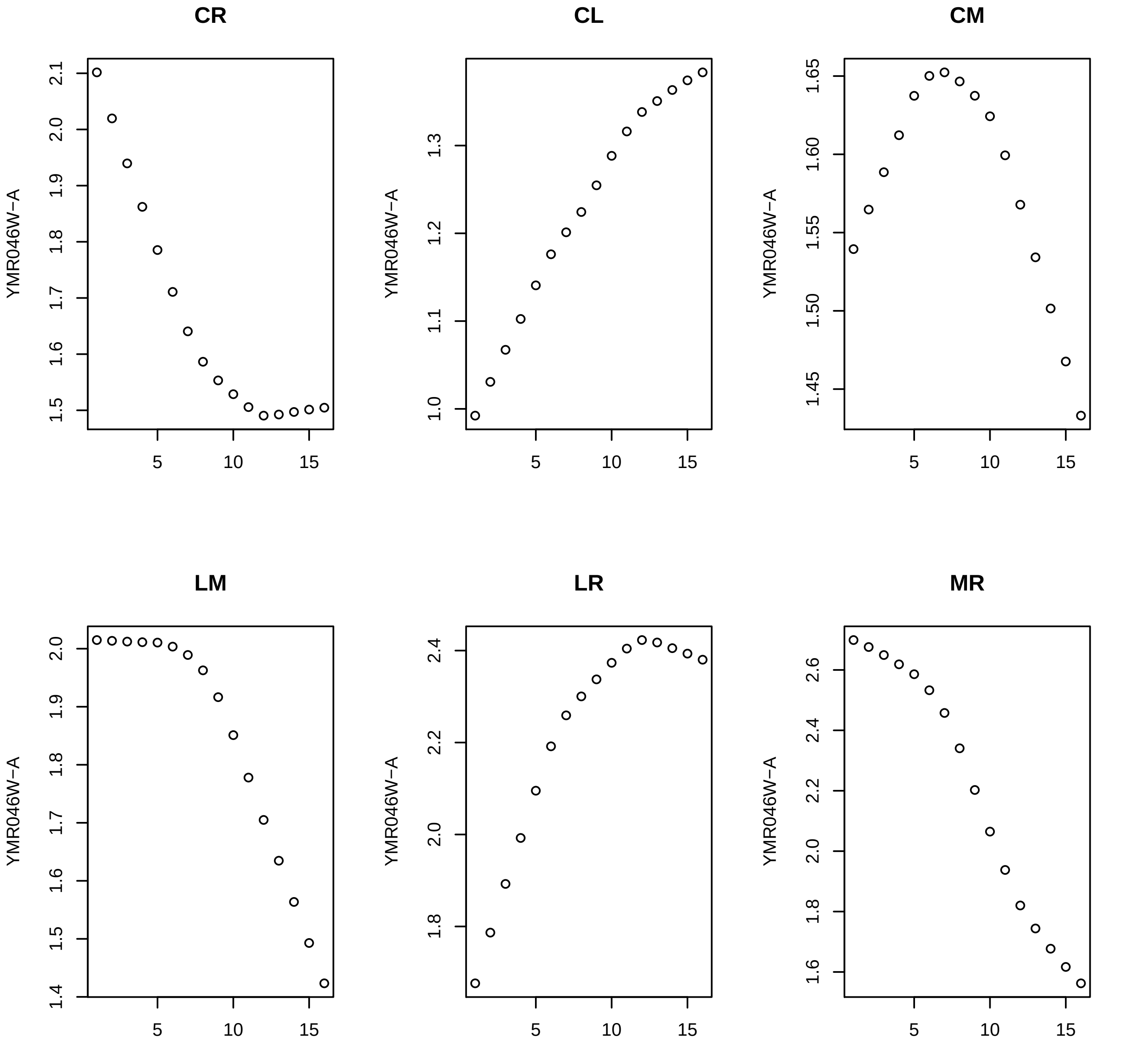
CR
CL
CM
LM
LR
MR
YMR046W−A
5
10
15

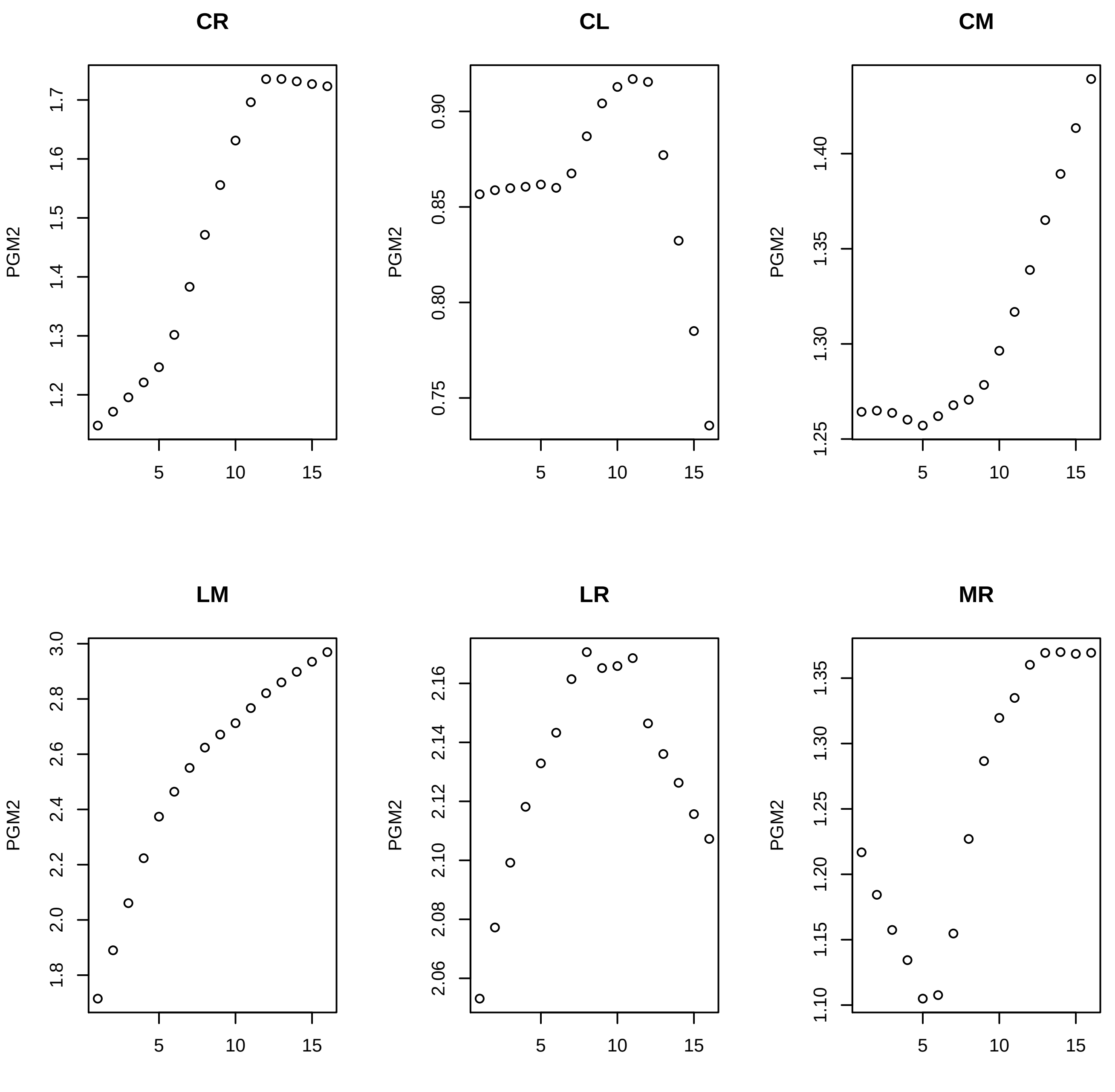
CR
CL
CM
LM
LR
MR
PGM2
1.2
1.3
1.4
1.5
1.6
1.7
0.75
0.80
0.85
0.90
1.25
1.30
1.35
1.40
1.8
2.0
2.2
2.4
2.6
2.8
3.0
2.06
2.08
2.10
2.12
2.14
2.16
1.10
1.15
1.20
1.25
1.30
1.35
5
10
15

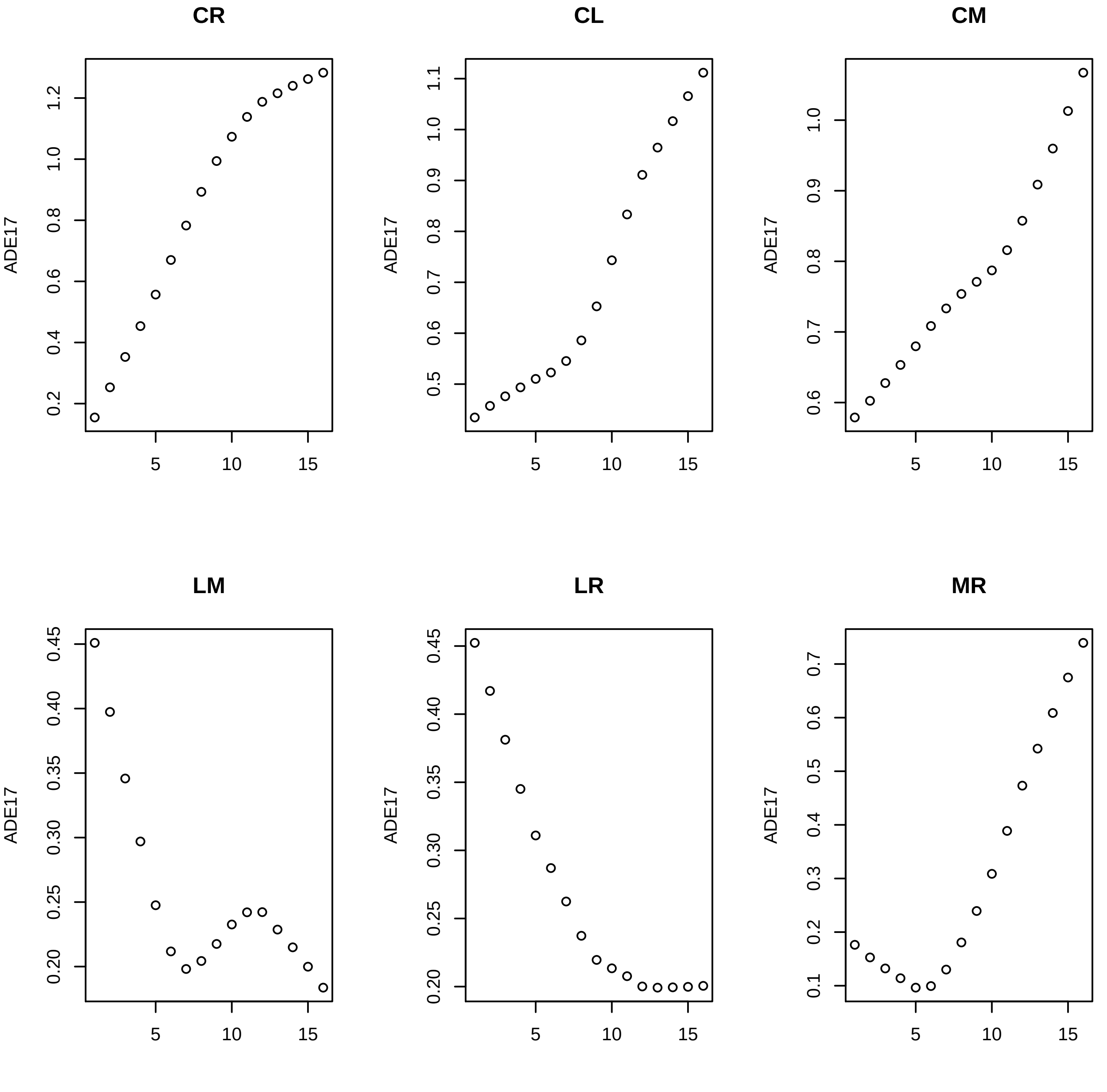
CR
CL
CM
LM
LR
MR
ADE17
5
10
15

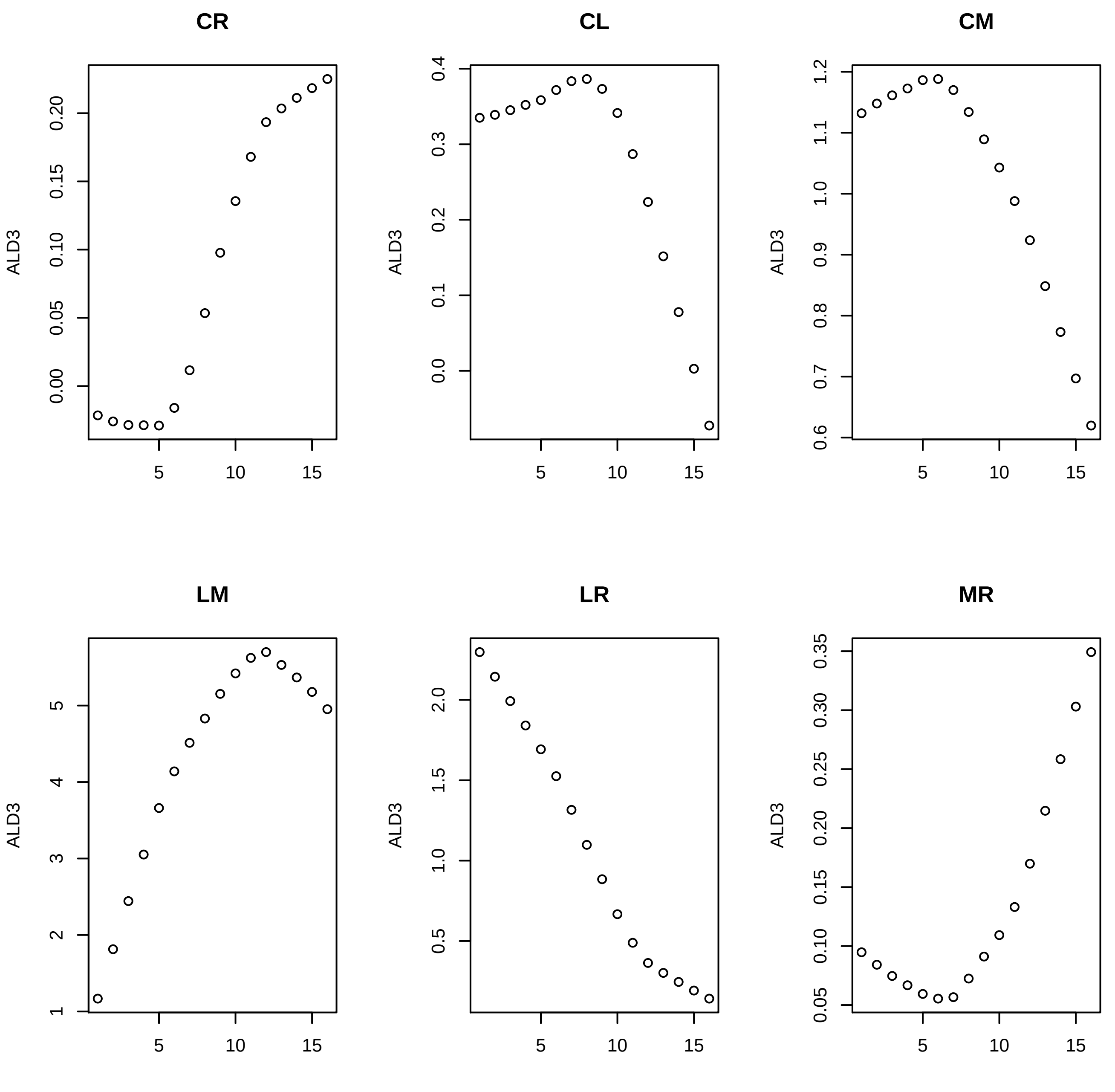
CR
CL
CM
LM
LR
MR
ALD3

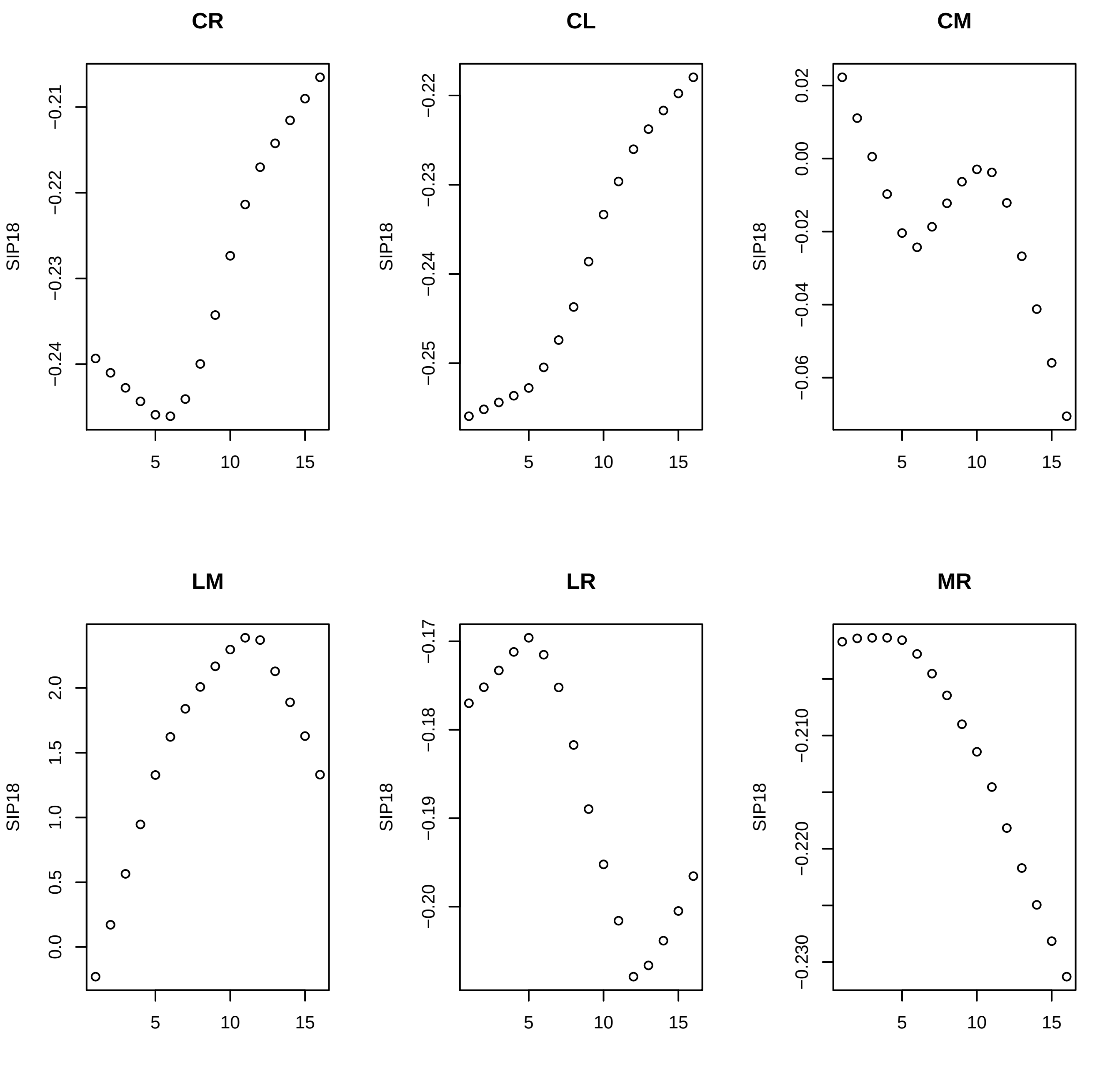
CR
CL
CM
LM
LR
MR
SIP18
−0.21
−0.22
−0.23
−0.24
−0.25
−0.17
−0.18
−0.19
−0.20
0.02
0.00
−0.02
−0.04
−0.06
0.0
0.5
1.0
1.5
2.0
−0.210
−0.220
−0.230
5
10
15

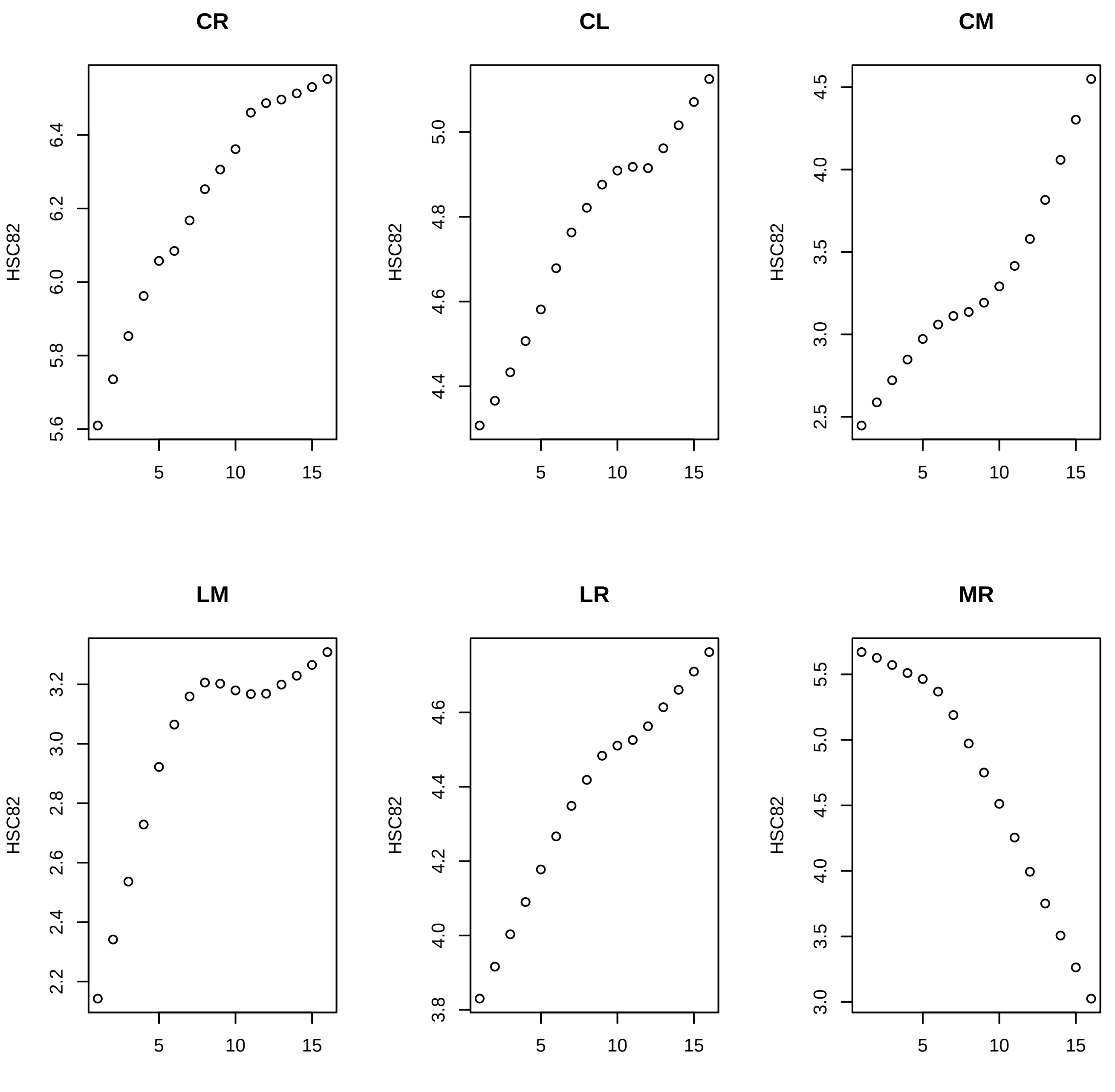
CR
CL
CM
LM
LR
MR
HSC82

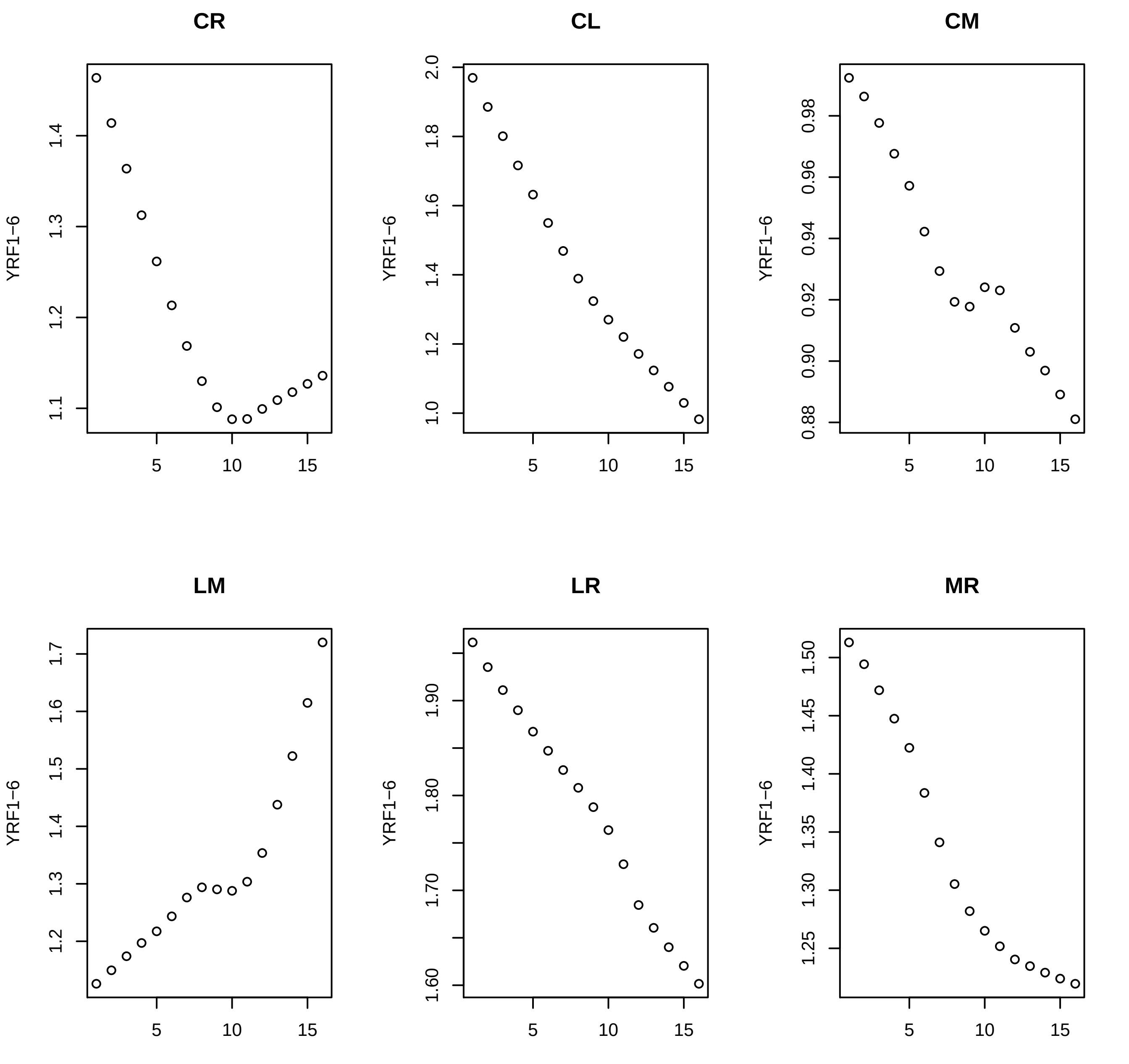

CR
CL
CM
LM
LR
MR
YRF1−6

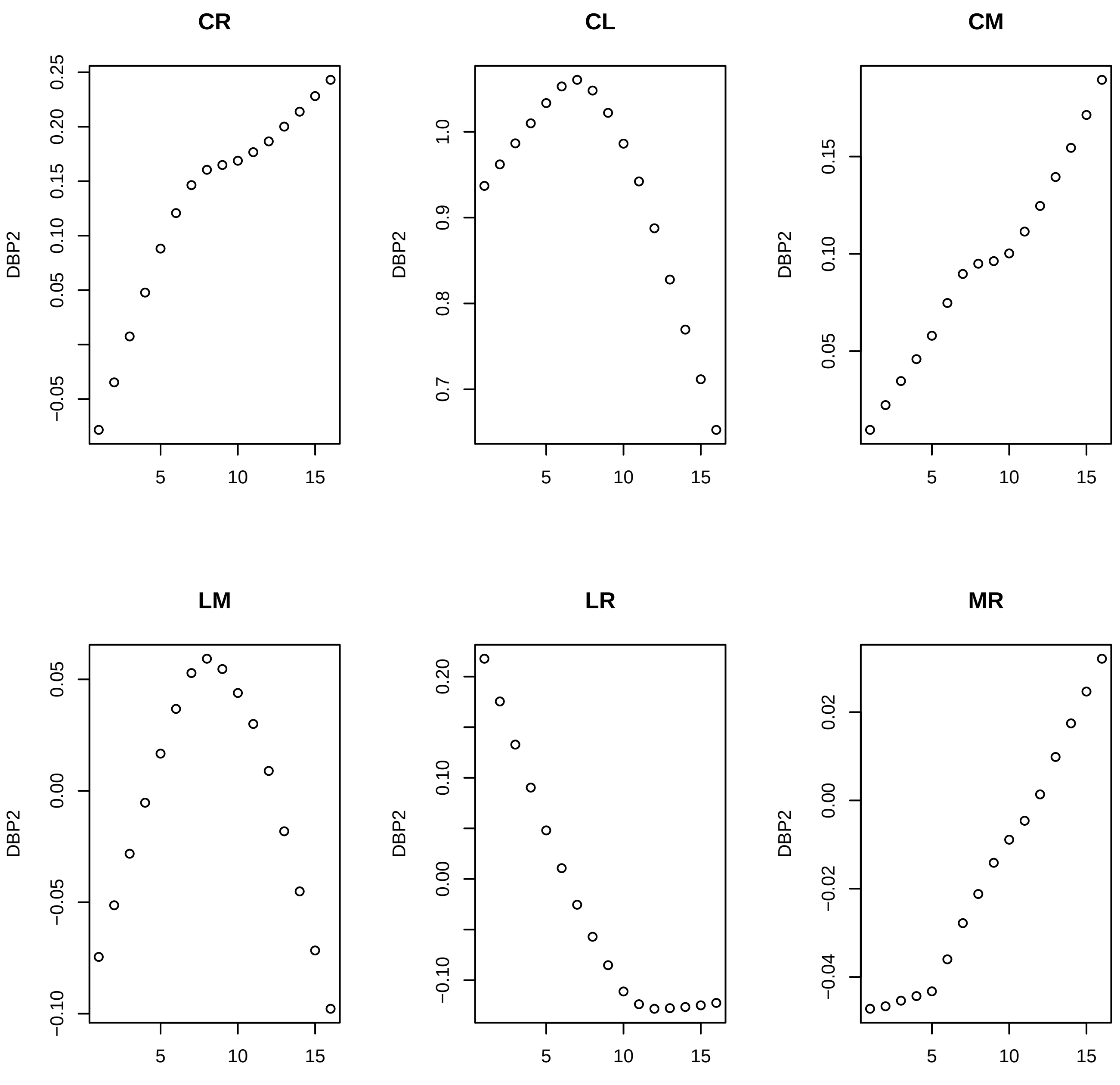
CR
CL
CM
LM
LR
MR
DBP2
−0.05
0.05
0.10
0.15
0.20
0.25
0.7
0.8
0.9
1.0
0.05
0.10
0.15
−0.10
−0.05
0.00
0.05
−0.10
0.00
0.10
0.20
−0.04
−0.02
0.00
0.02
5
10
15

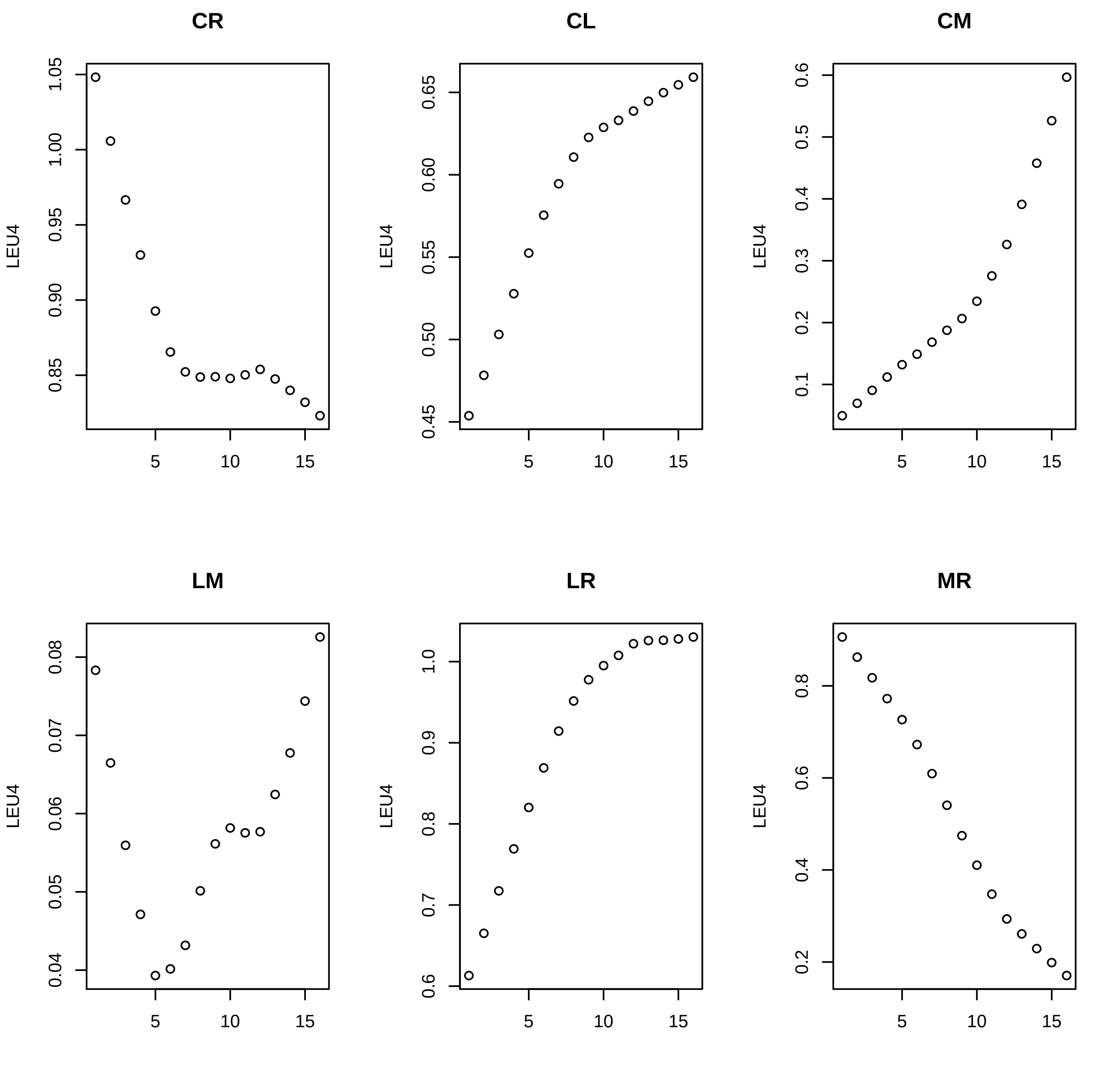

CR
CL
CM
LM
LR
MR
LEU4

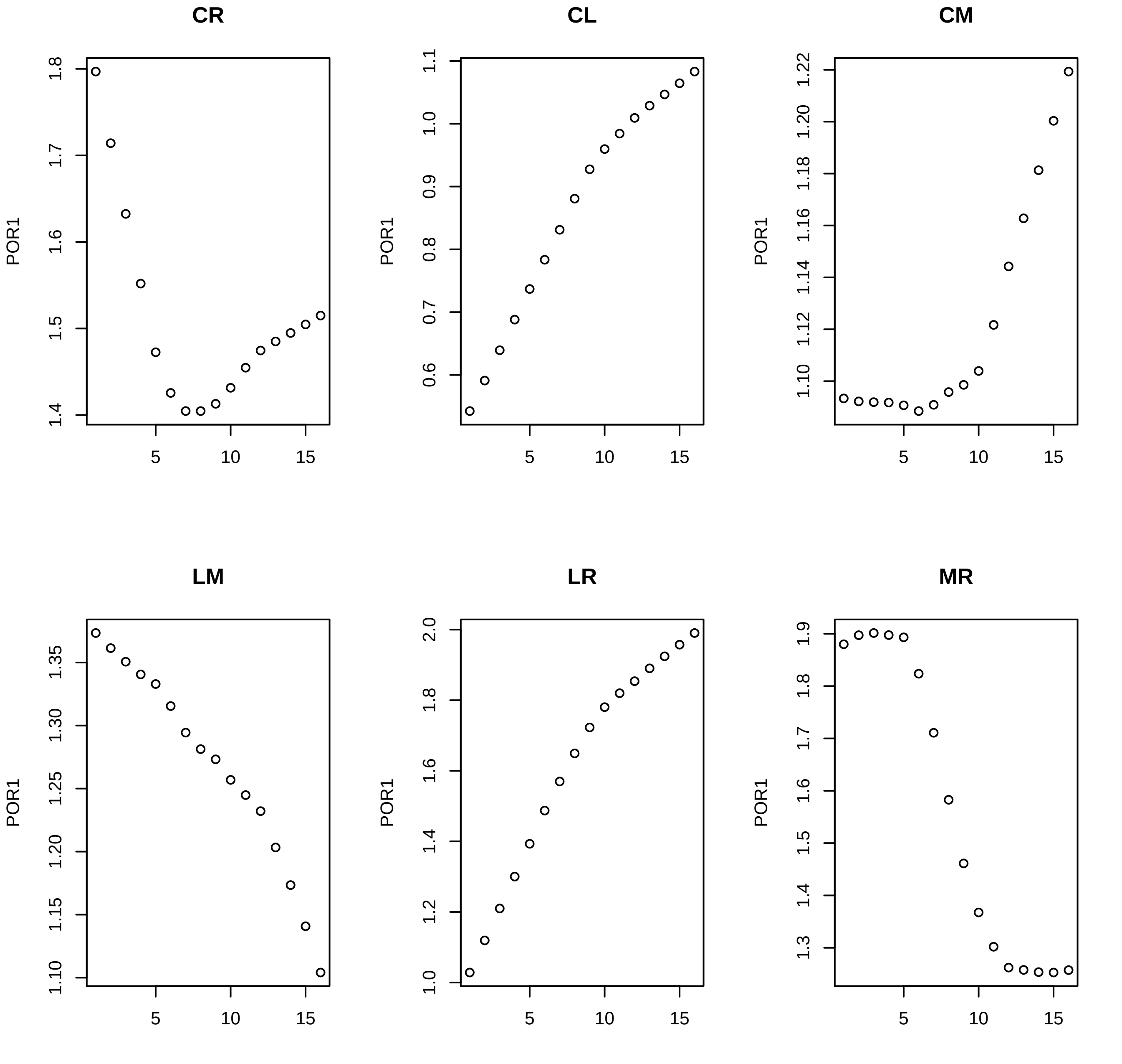
CR
CL
CM
LM
LR
MR
POR1
5
10
15
1.4
1.5
1.6
1.7
1.8
0.6
0.7
0.8
0.9
1.0
1.1
1.10
1.12
1.14
1.16
1.18
1.20
1.22
1.15
1.25
1.30
1.35
1.2
1.3
2.0

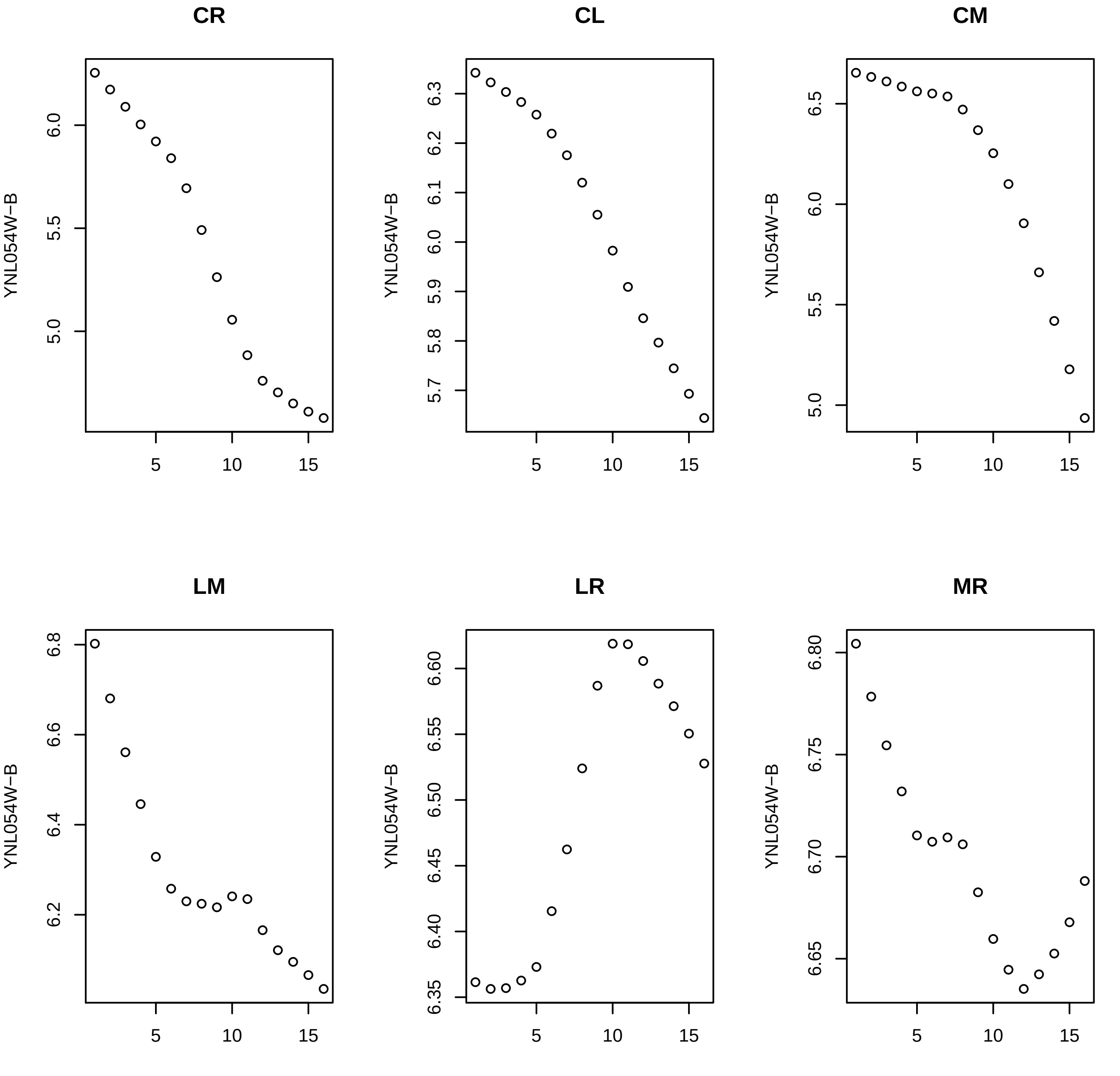
CR
CL
CM
LM
LR
MR
YNL054W−B
6.0
5.5
5.0
6.3
6.2
6.1
6.0
5.9
5.8
5.7
6.5
6.0
5.5
5.0
6.8
6.6
6.4
6.2
6.60
6.55
6.50
6.45
6.40
6.35
6.80
6.75
6.70
6.65
5
10
15

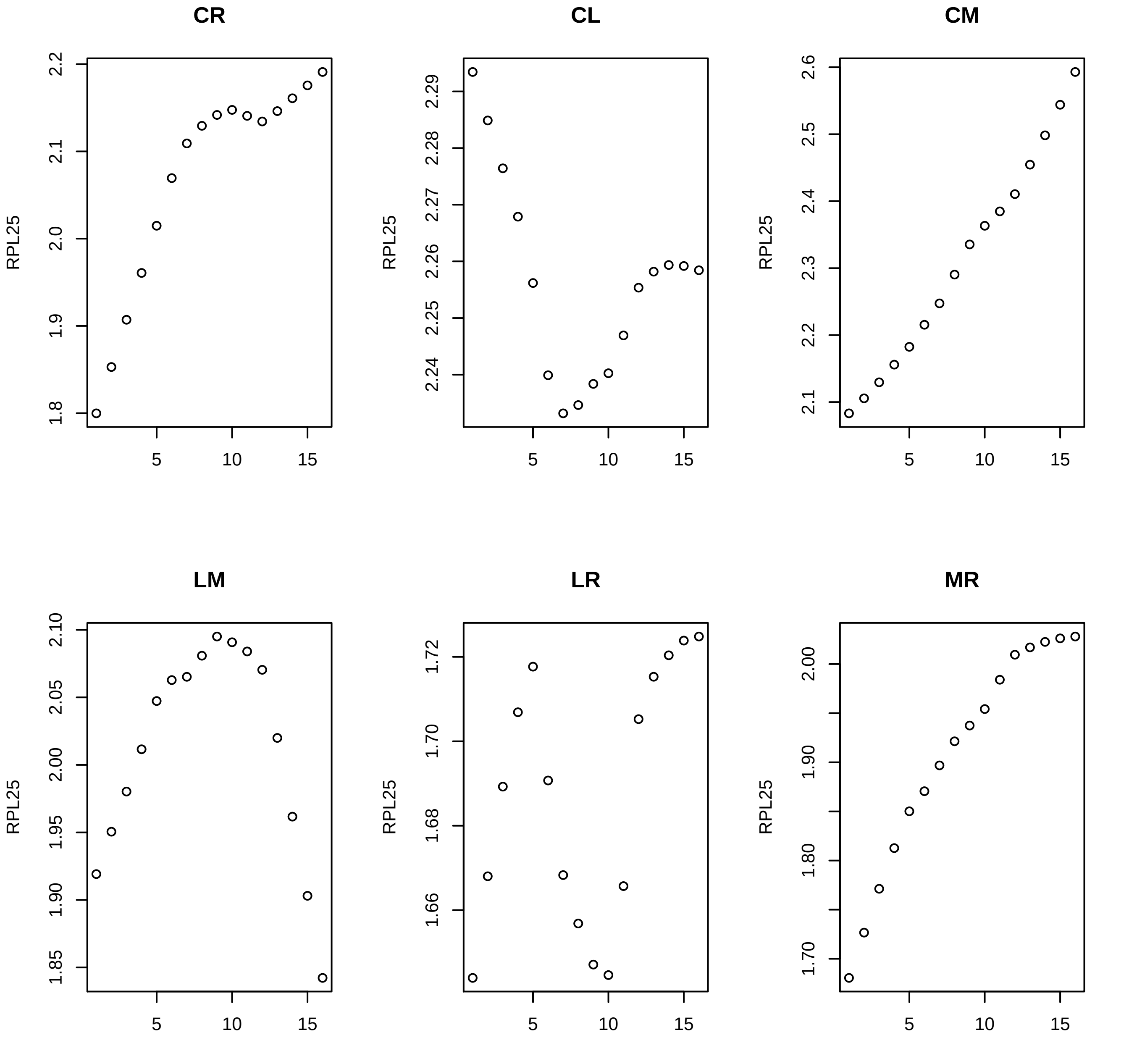
CR
RPL25
1.8 1.9 2.0 2.1 2.2
5 10 15
CL
RPL25
2.24 2.25 2.26 2.27 2.28 2.29
5 10 15
CM
RPL25
2.1 2.2 2.3 2.4 2.5 2.6
5 10 15
LM
RPL25
1.85 1.90 1.95 2.00 2.05 2.10
5 10 15
LR
RPL25
1.66 1.68 1.70 1.72
5 10 15
MR
RPL25
1.70 1.80 1.90 2.00
5 10 15

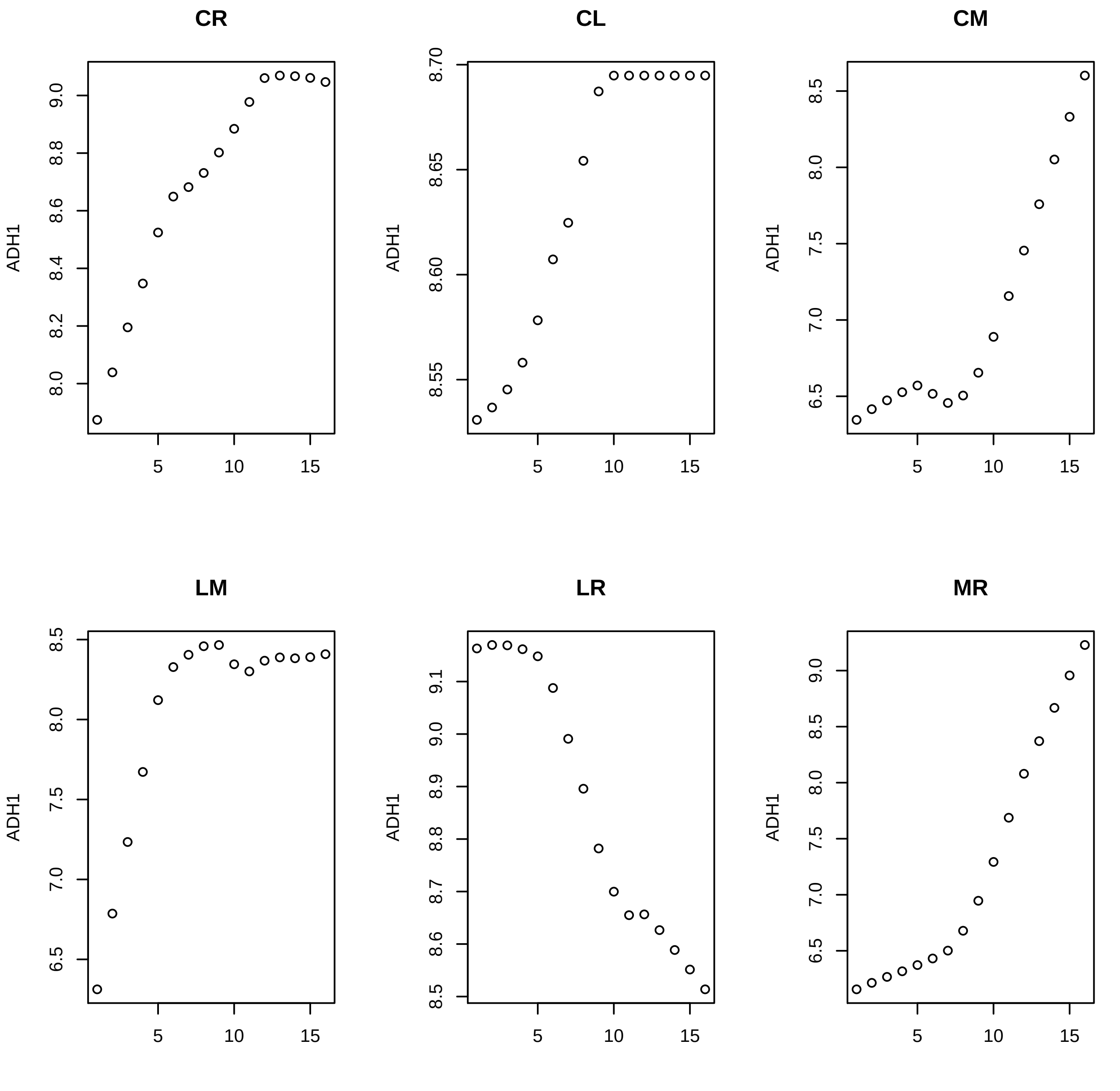
CR
CL
CM
LM
LR
MR
ADH1
5
10
15
8.0
8.2
8.4
8.6
8.8
9.0
8.55
8.60
8.65
8.70
6.5
7.0
7.5
8.0
8.5
9.0
8.5
8.6
8.7
8.8
8.9
9.1

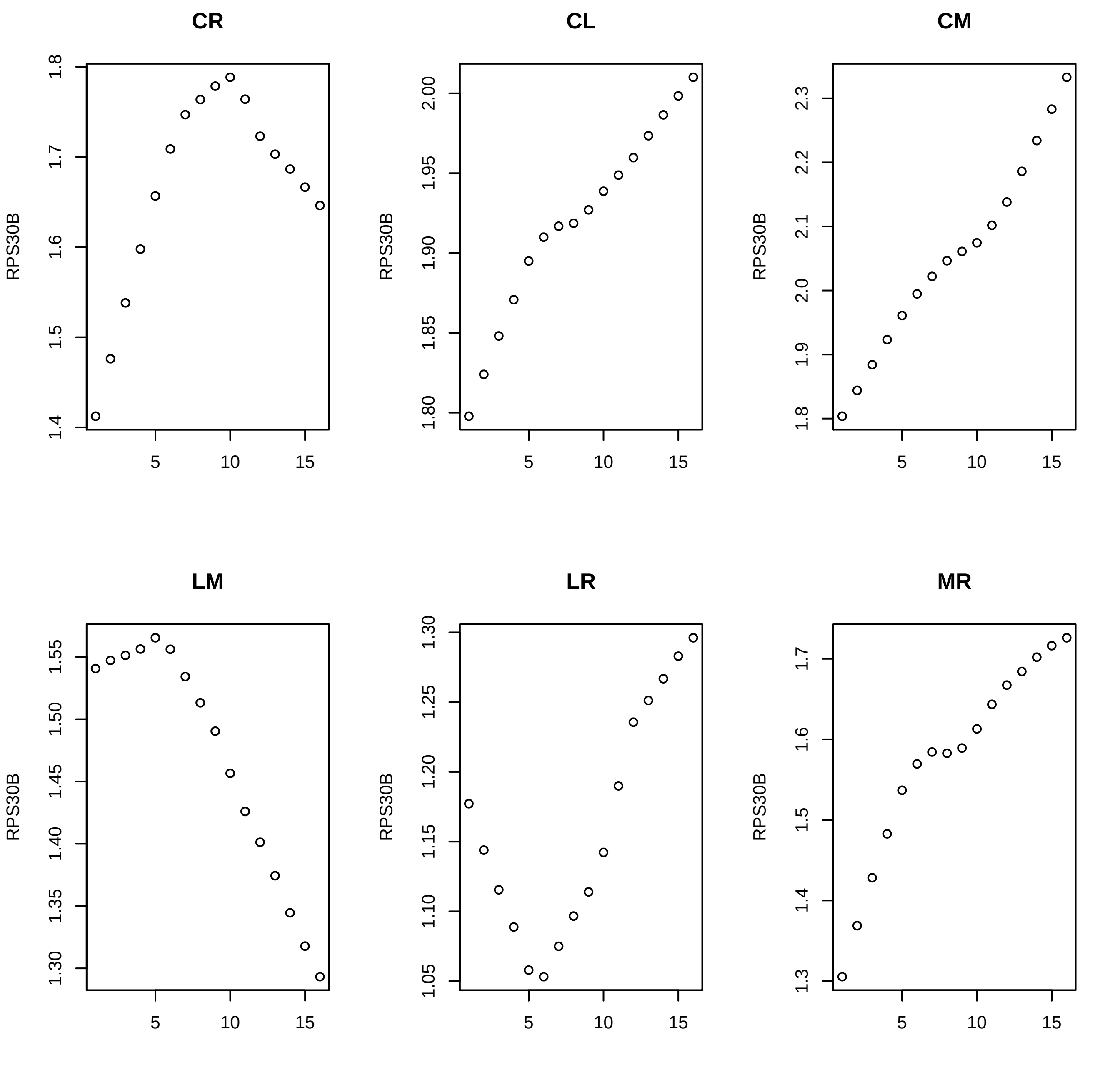
CR
CL
CM
LM
LR
MR
RPS30B
1.4
1.5
1.6
1.7
1.8
1.80
1.85
1.90
1.95
2.00
1.9
2.0
2.1
2.2
2.3
1.30
1.35
1.40
1.45
1.50
1.55
1.05
1.10
1.15
1.20
1.25
1.3
5
10
15

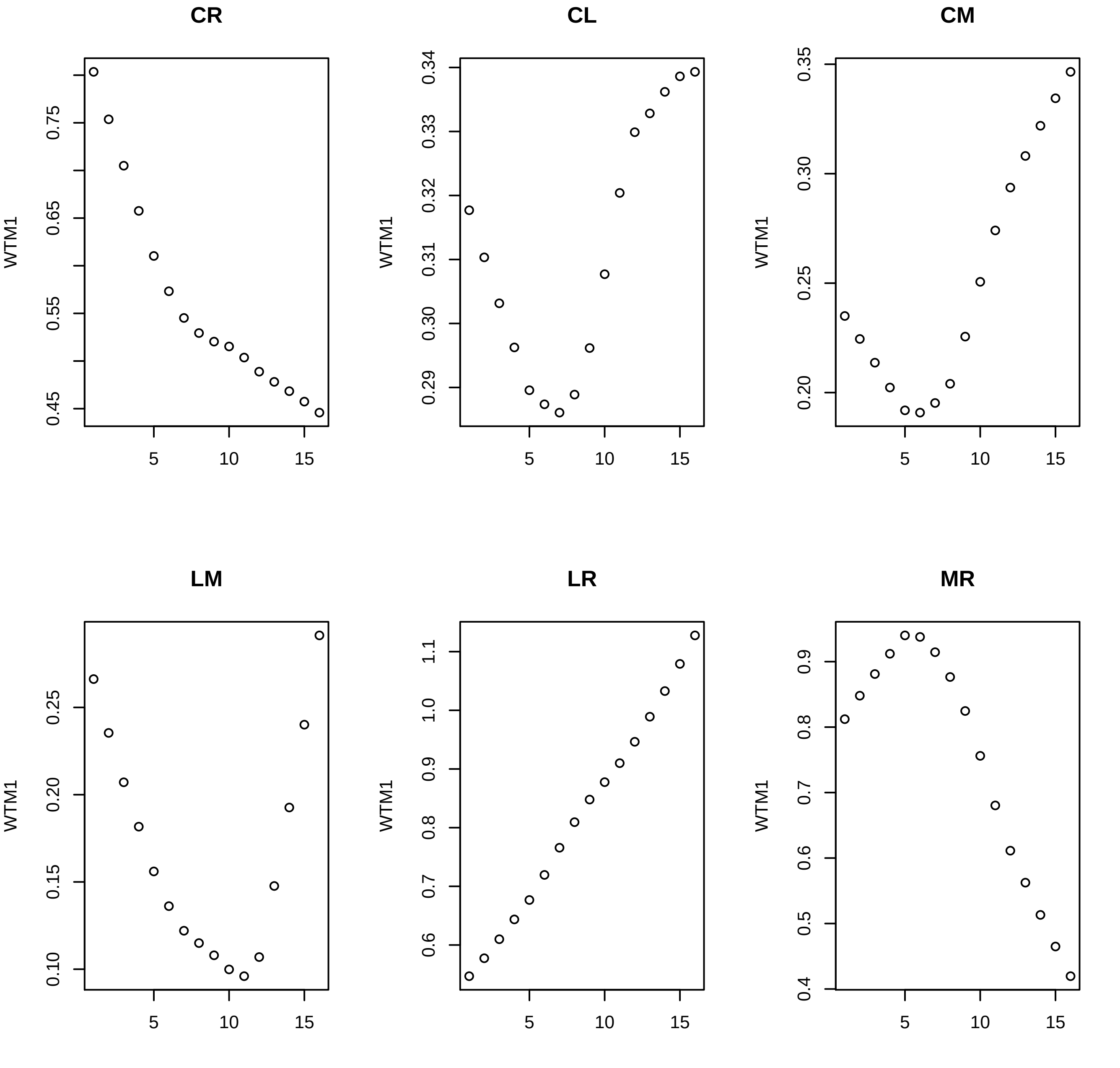
CR
CL
CM
LM
LR
MR
WTM1
0.45
0.55
0.65
0.75
0.29
0.30
0.31
0.32
0.33
0.34
0.20
0.25
0.30
0.35
0.10
0.15
0.20
0.25
0.6
0.7
0.8
0.9
1.0
1.1
0.4
0.5
0.6
0.7
0.8
0.9
5
10
15

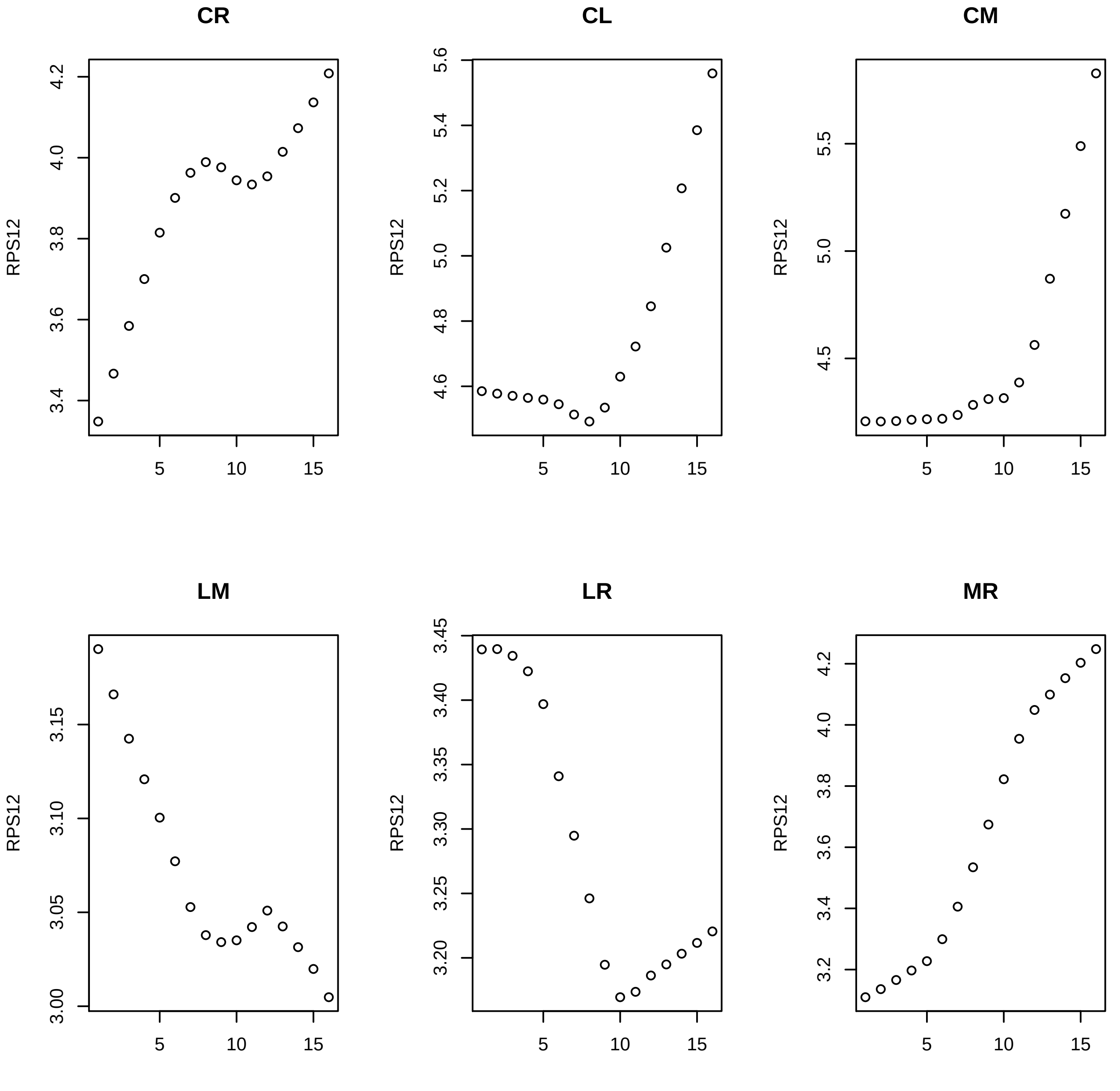
CR
CL
CM
LM
LR
MR
RPS12
5
10
15

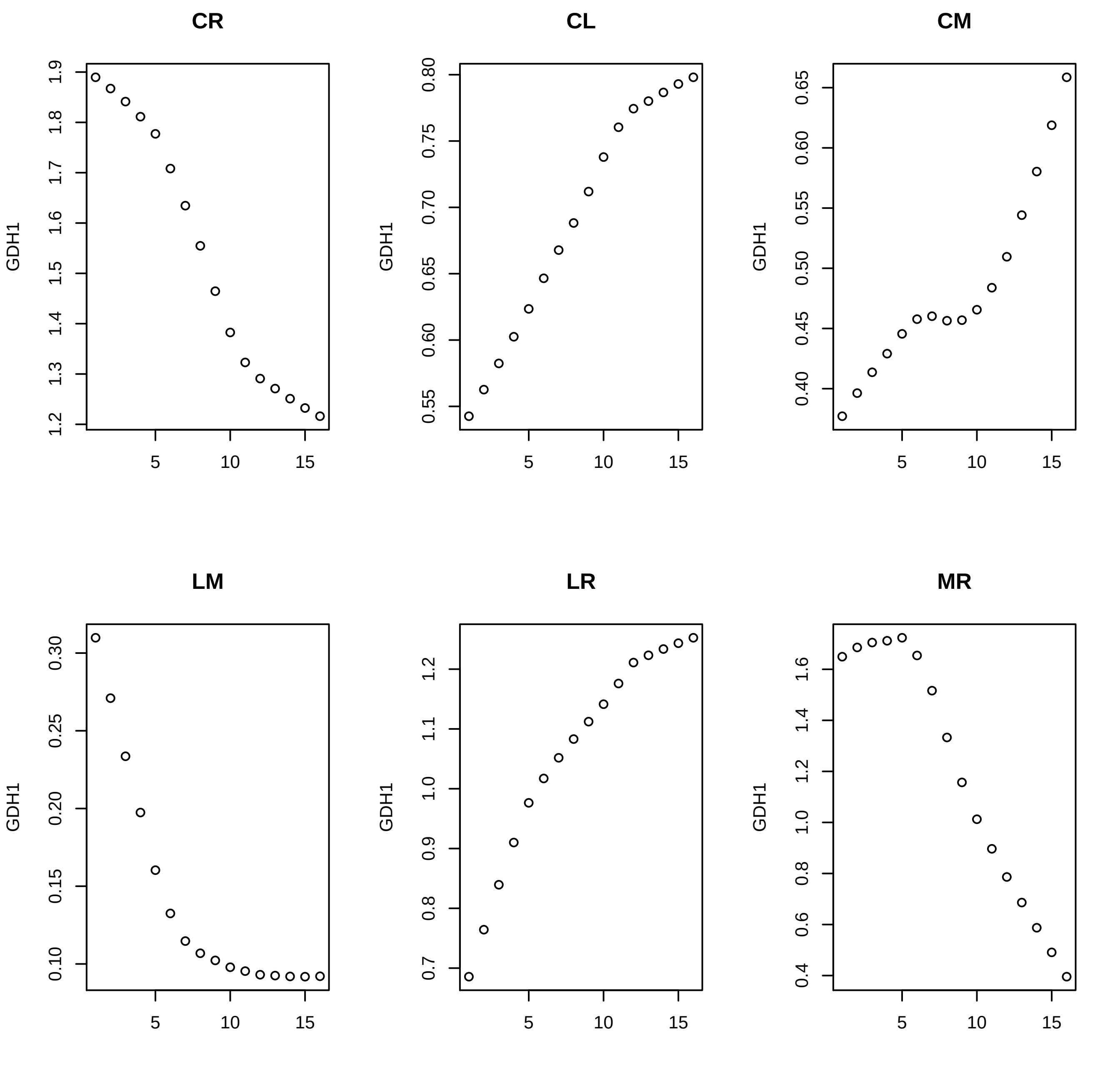
CR
CL
CM
LM
LR
MR
GDH1
1.2
1.3
1.4
1.5
1.6
1.7
1.8
1.9
0.55
0.60
0.65
0.70
0.75
0.80
0.40
0.45
0.50
0.55
0.60
0.65
0.10
0.15
0.20
0.25
0.30
0.7
0.8
0.9
1.0
1.1
1.2
0.4
0.6
0.8
1.0
1.2
1.4
1.6
5
10
15

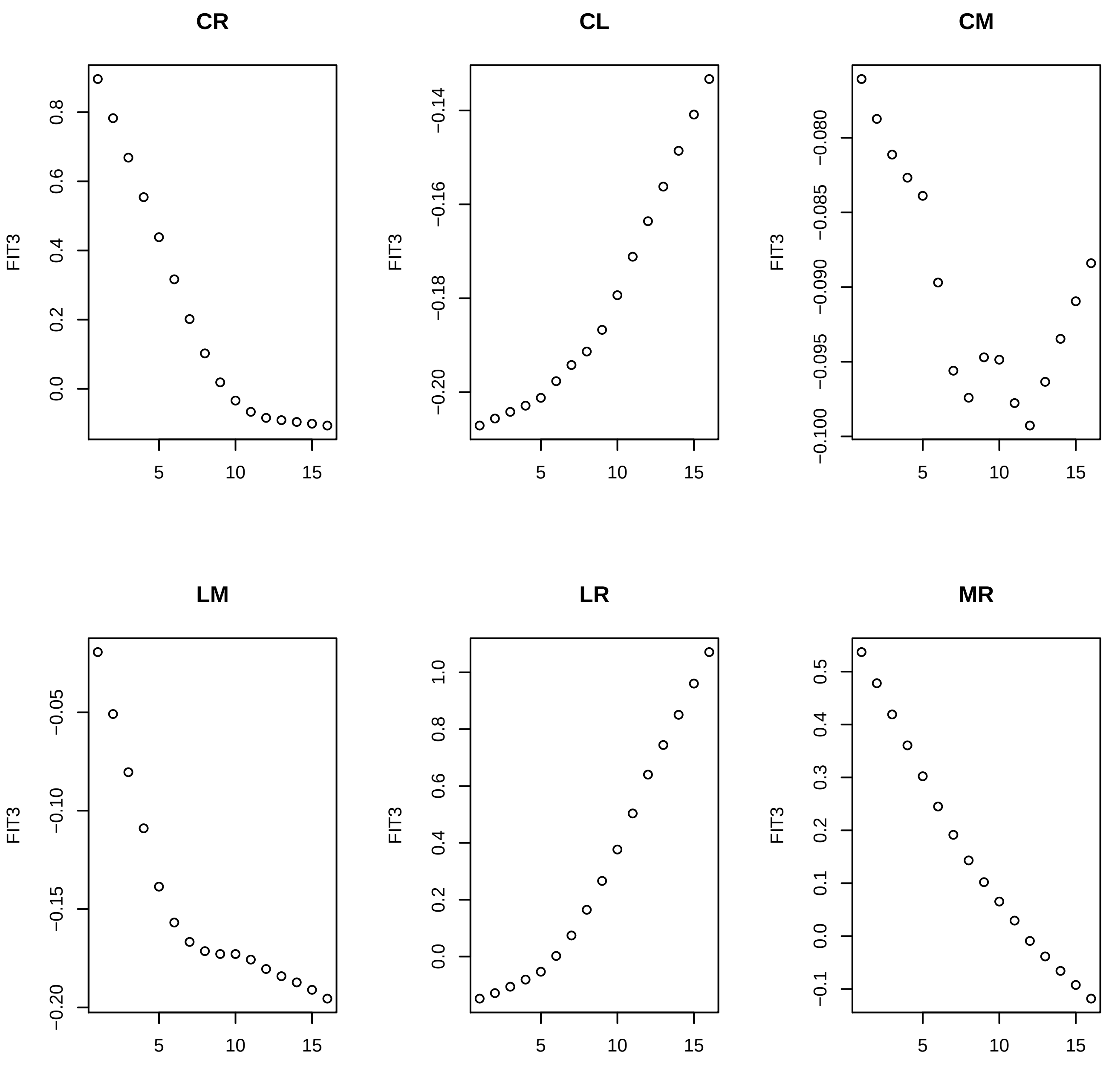
CR
CL
CM
LM
LR
MR
FIT3
5
10
15
0.0
0.2
0.4
0.6
0.8
−0.14
−0.16
−0.18
−0.20
−0.080
−0.085
−0.090
−0.095
−0.100
−0.05
−0.10
−0.15
−0.20
1.0
−0.1
0.1
0.3
0.5

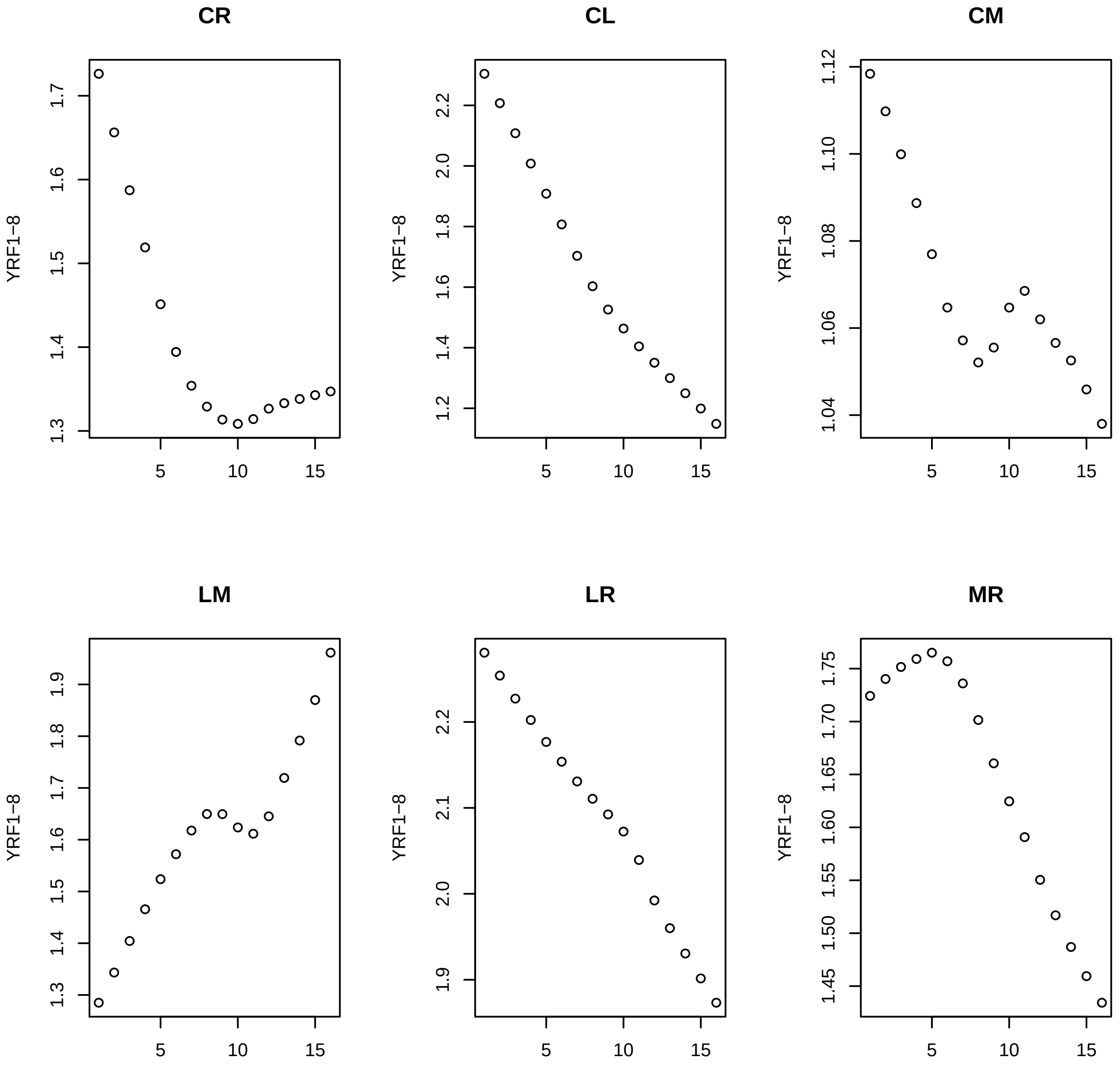
CR
CL
CM
LM
LR
MR
YRF1−8
1.3
1.4
1.5
1.6
1.7
1.2
1.8
2.0
2.2
1.04
1.06
1.08
1.10
1.12
1.9
2.1
1.45
1.50
1.55
1.60
1.65
1.70
1.75
5
10
15

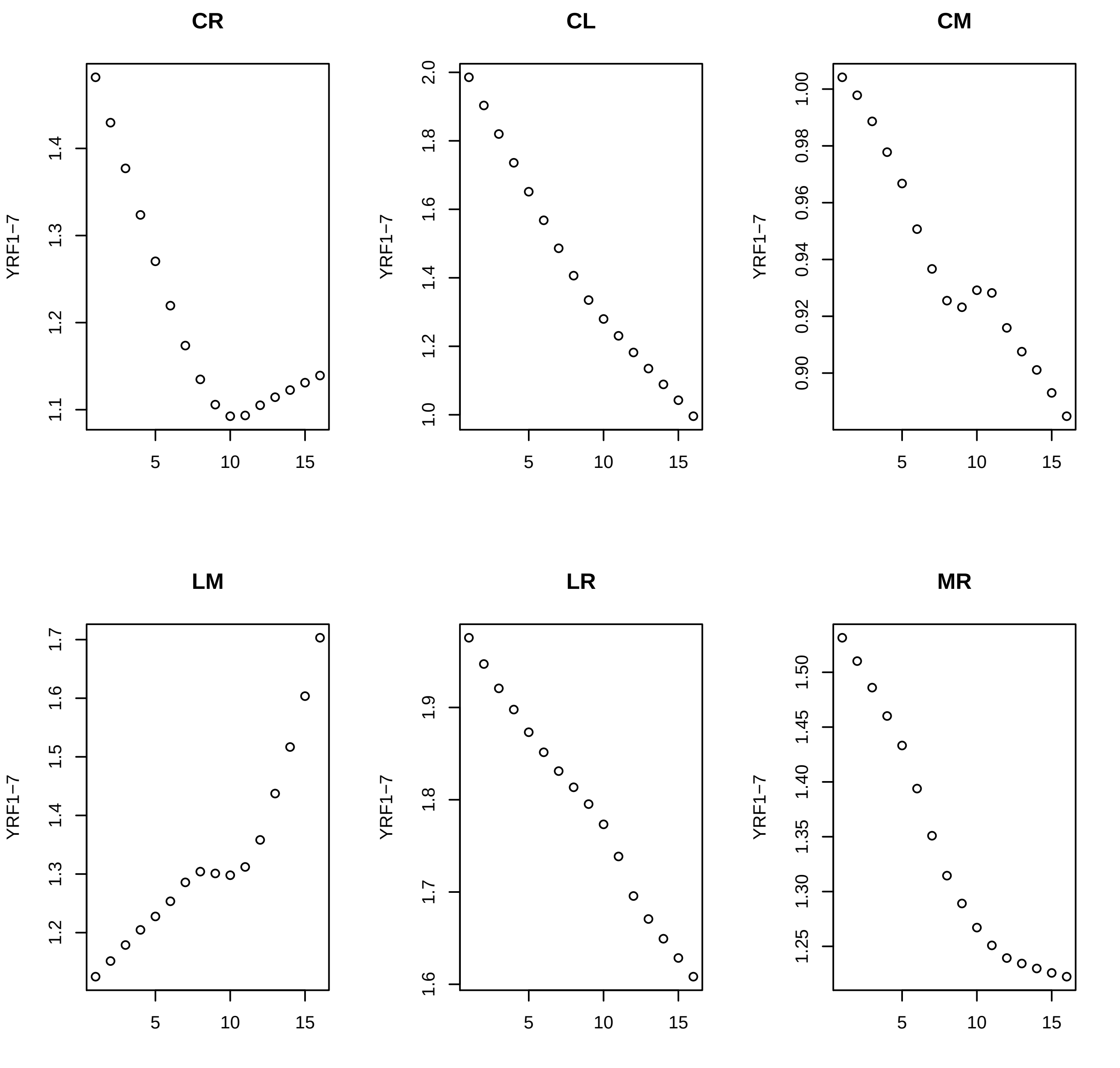
CR
CL
CM
LM
LR
MR
YRF1−7
5
10
15
1.1
1.2
1.3
1.4
1.0
1.2
1.4
1.6
1.8
2.0
0.90
0.92
0.94
0.96
0.98
1.00
1.5
1.6
1.7
1.9
1.25
1.30
1.35
1.40
1.45
1.50

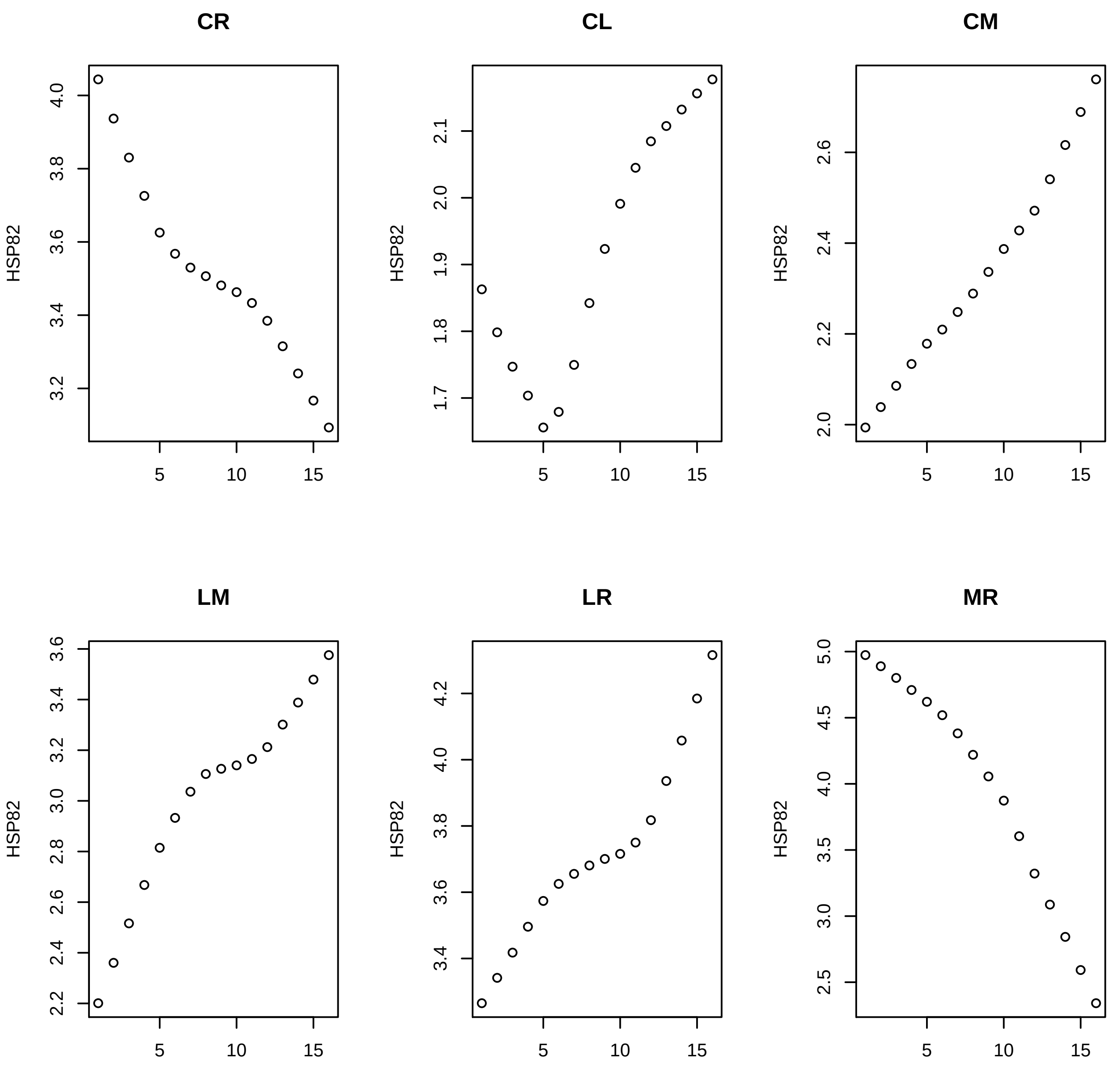
CR
CL
CM
LM
LR
MR
HSP82
5
10
15

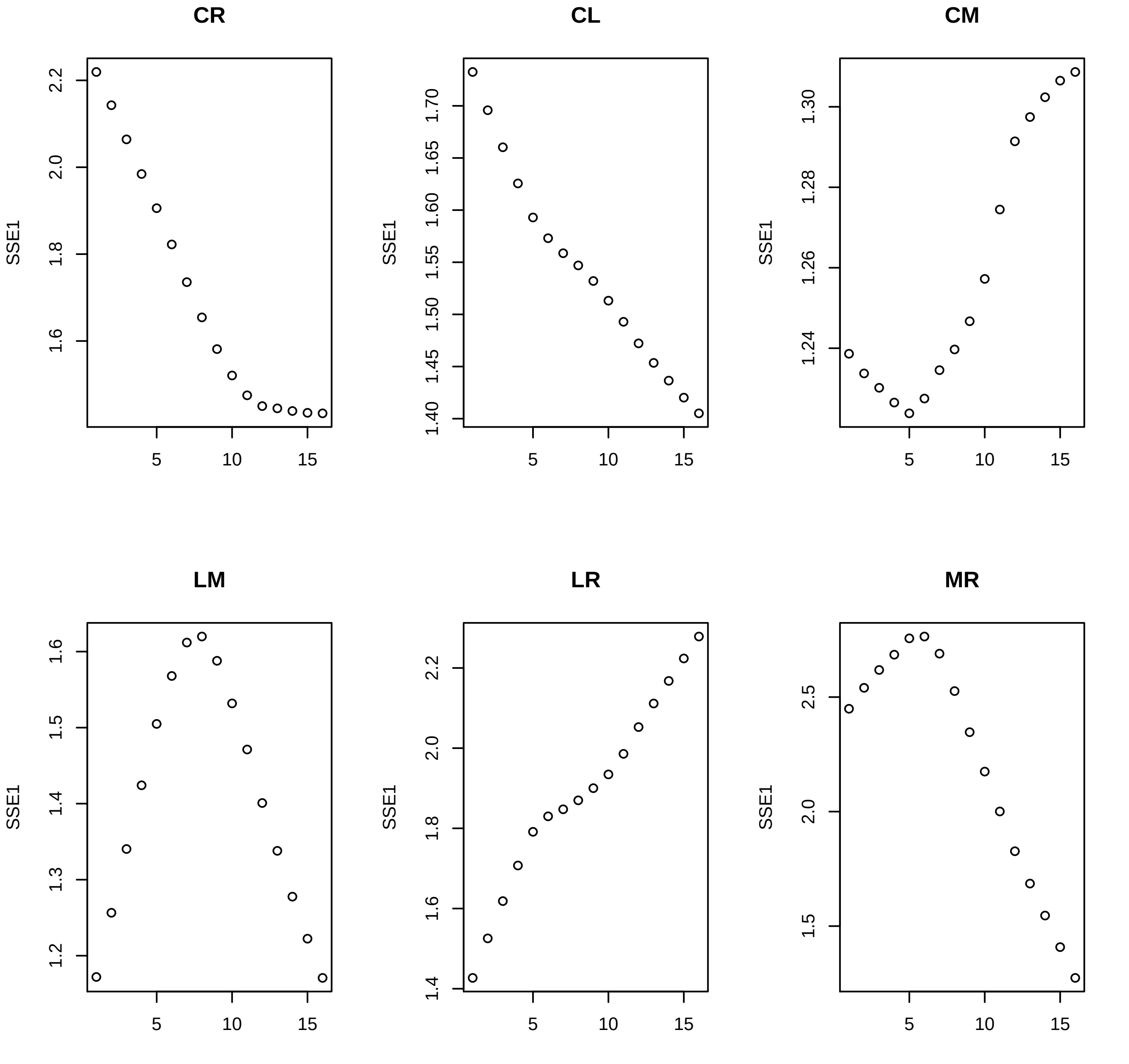
CR
CL
CM
LM
LR
MR
SSE1
2.2
2.0
1.8
1.6
1.70
1.65
1.60
1.55
1.50
1.45
1.40
1.30
1.28
1.26
1.24
1.6
1.5
1.4
1.3
1.2
2.2
2.0
1.8
1.6
1.4
2.5
2.0
1.5
5
10
15

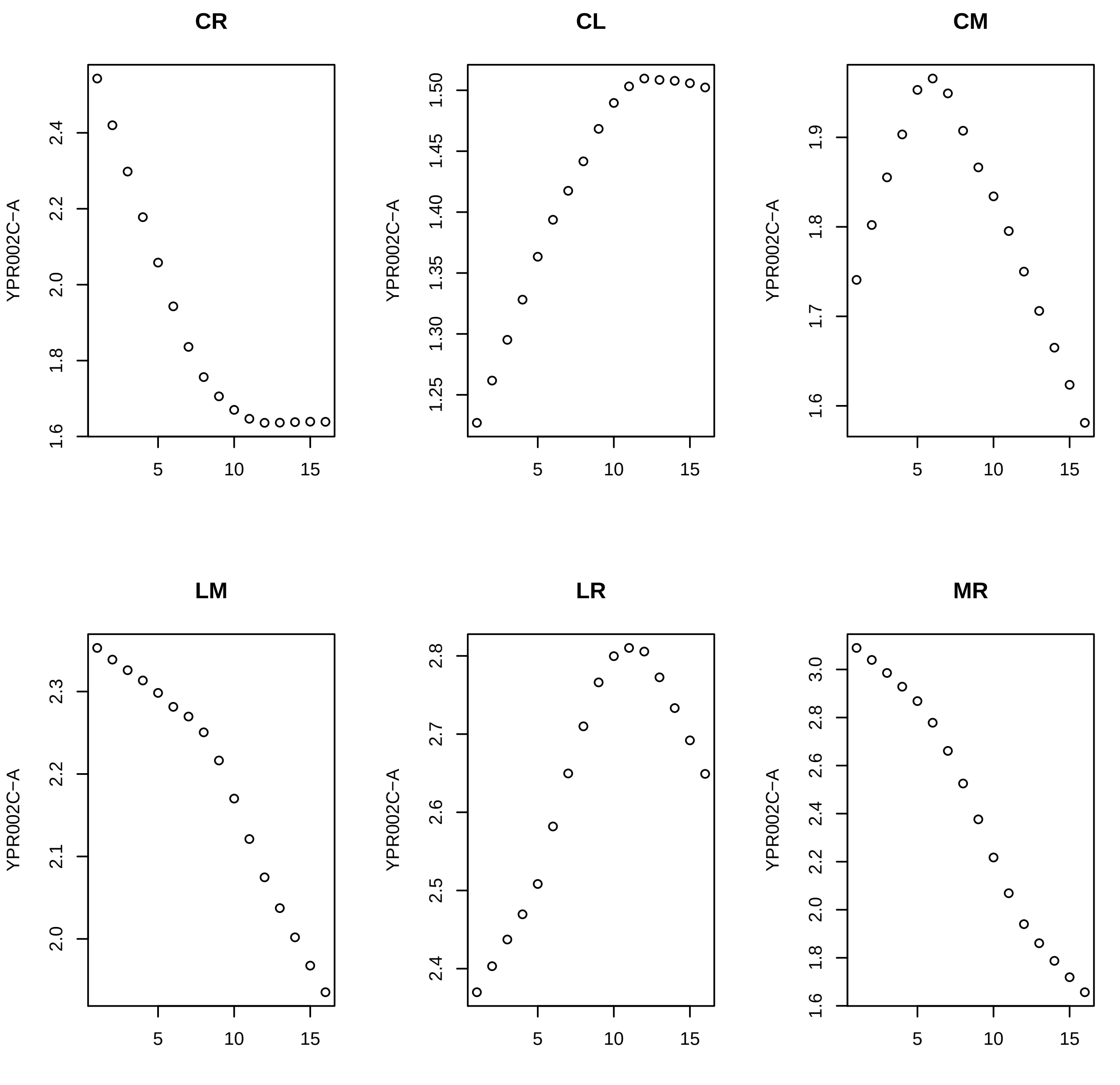
CR
CL
CM
LM
LR
MR
YPR002C−A
5
10
15

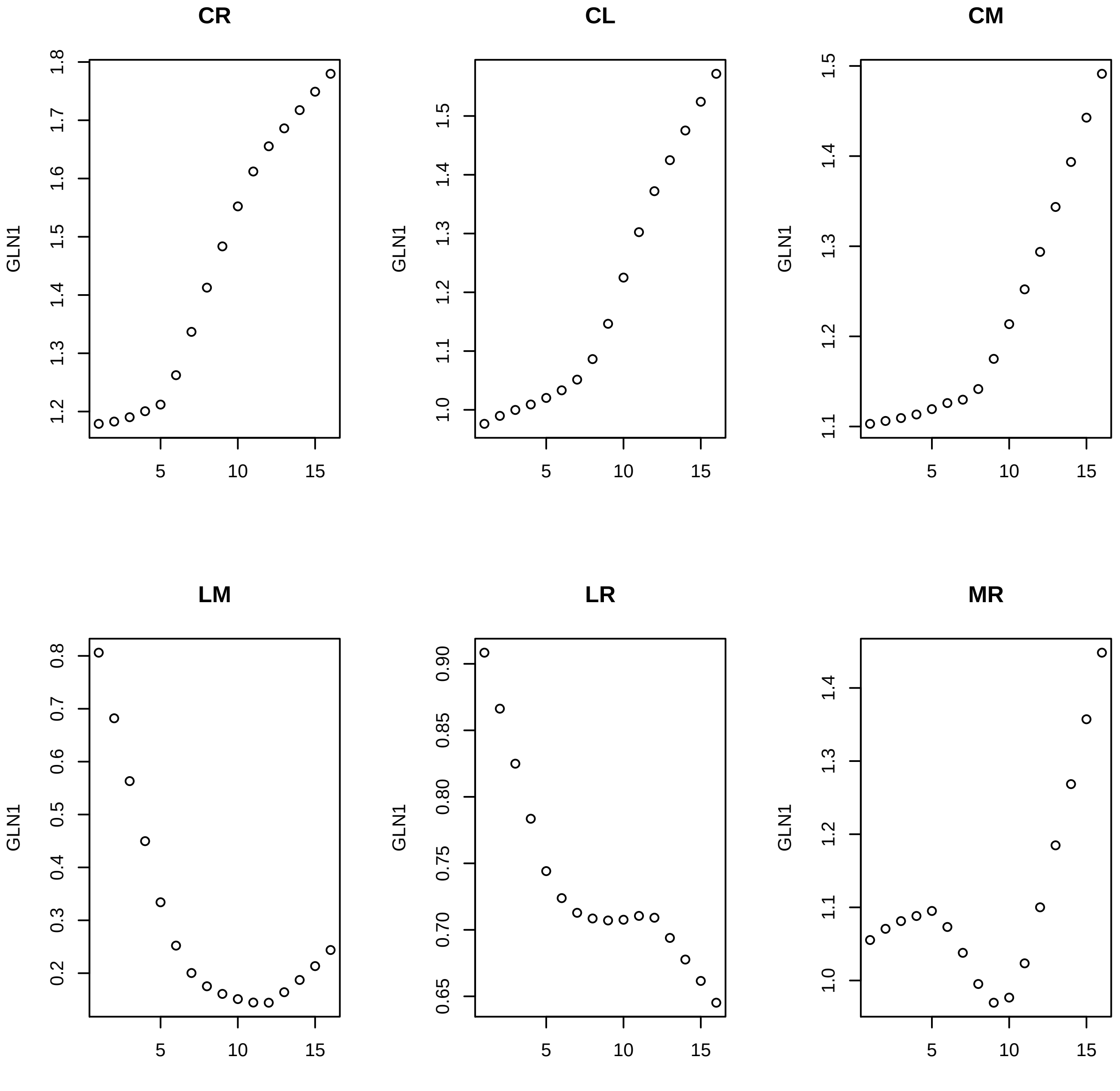

CR
CL
CM
LM
LR
MR
GLN1
1.2
1.3
1.4
1.5
1.6
1.7
1.8
1.0
1.1
1.2
1.3
1.4
1.5
1.1
1.2
1.3
1.4
1.5
0.2
0.3
0.4
0.5
0.6
0.7
0.8
0.65
0.70
0.75
0.80
0.85
0.90
1.0
1.1
1.2
1.3
1.4
5
10
15

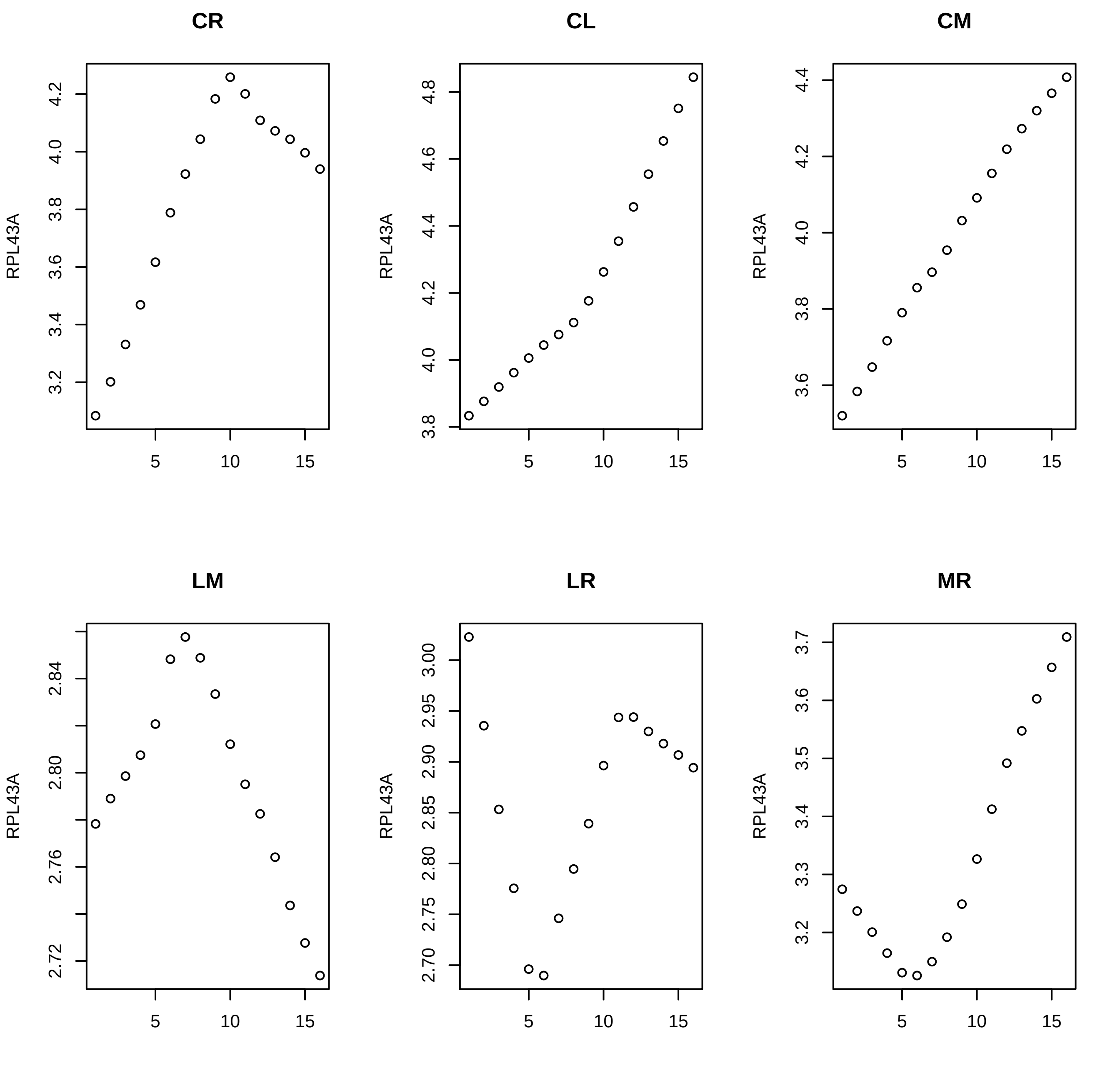
CR
CL
CM
LM
LR
MR
RPL43A
3.2
3.4
3.6
3.8
4.0
4.2
3.8
4.0
4.2
4.4
4.6
4.8
3.6
3.8
4.0
4.2
4.4
2.72
2.76
2.80
2.84
2.70
2.75
2.80
2.85
2.90
2.95
3.00
3.2
3.3
3.4
3.5
3.6
3.7
5
10
15

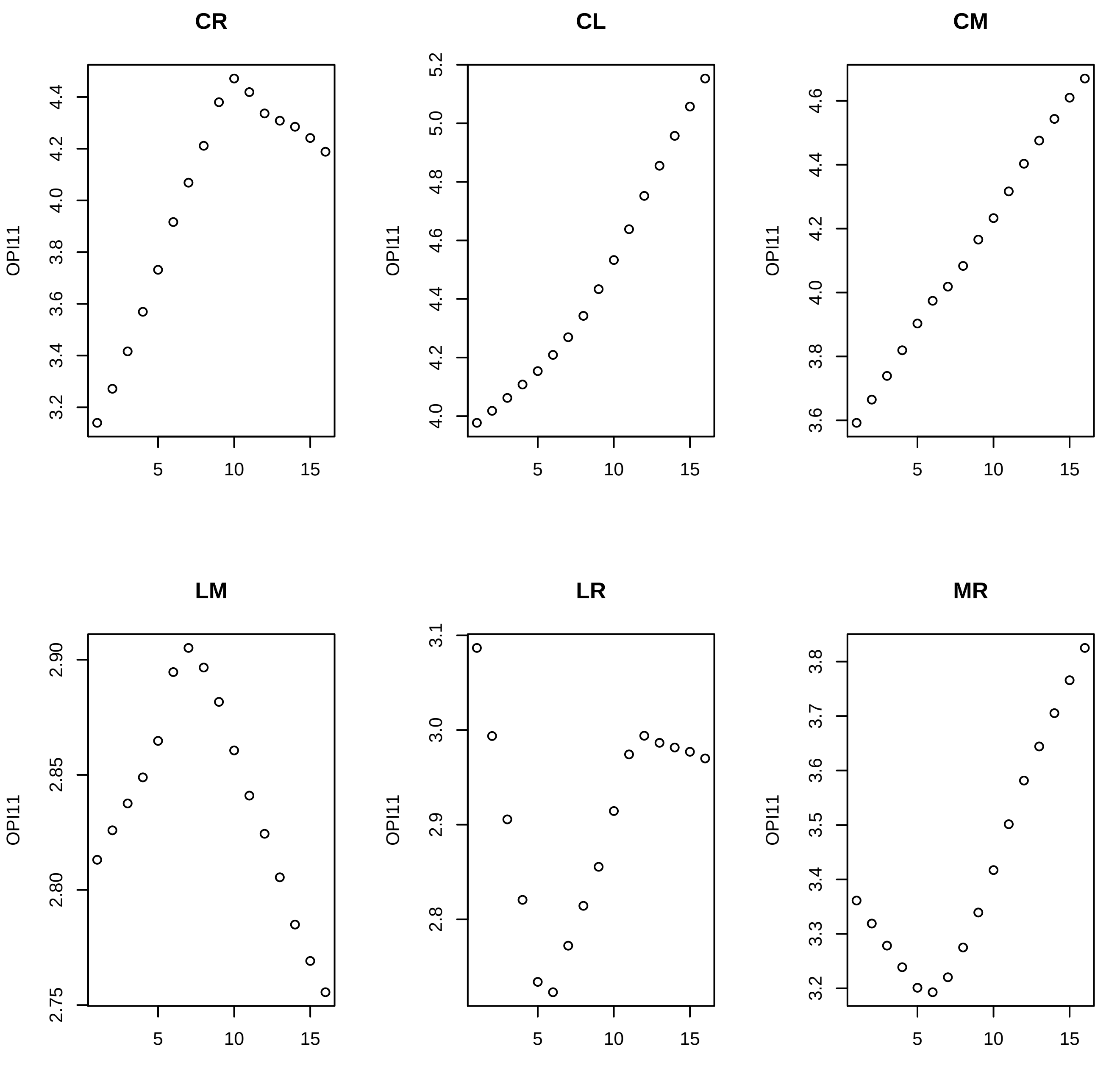

CR
CL
CM
LM
LR
MR
OPI11

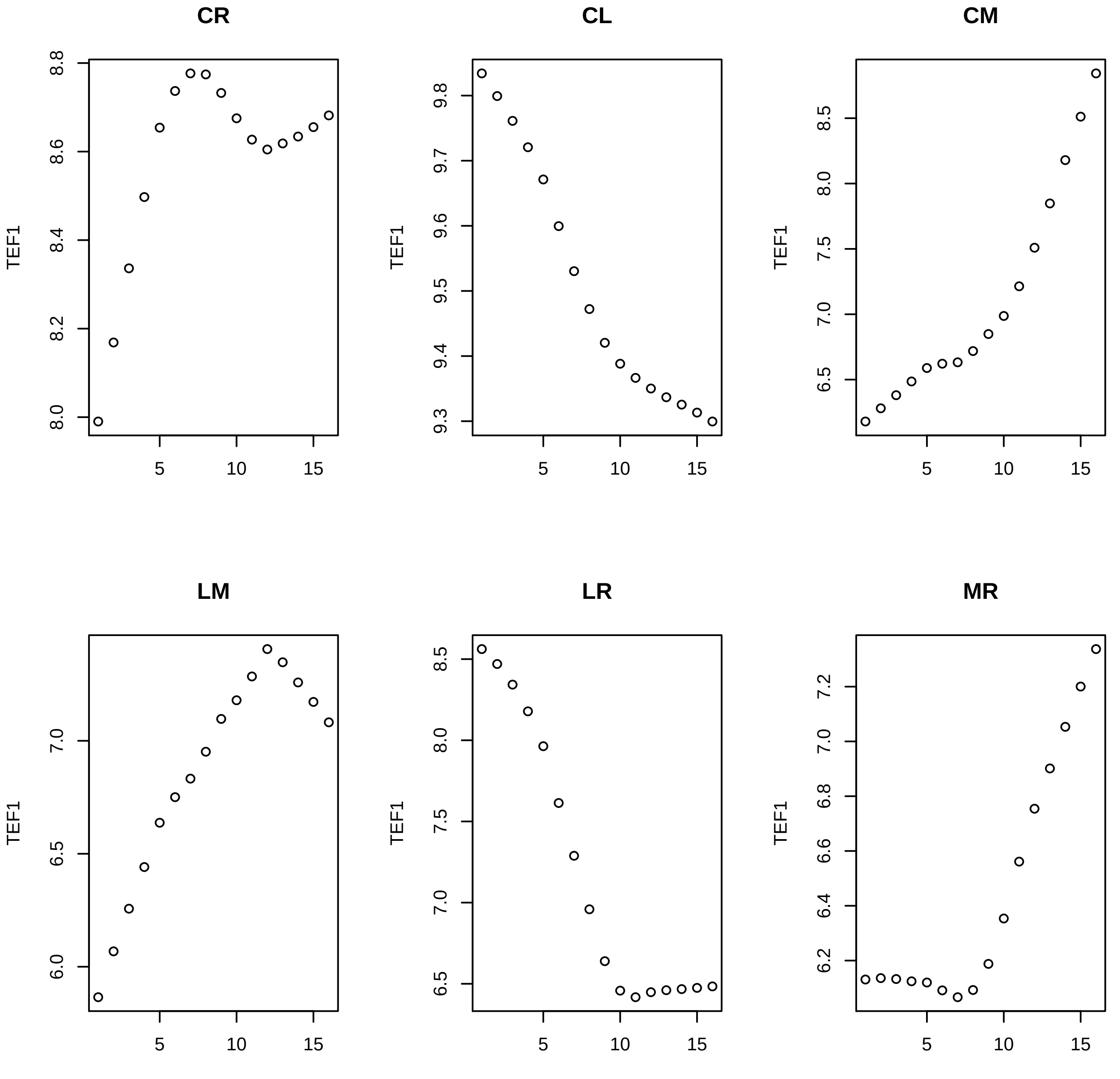
CR
CL
CM
LM
LR
MR
TEF1
5
10
15
8.0
8.2
8.4
8.6
8.8
9.3
9.4
9.5
9.6
9.7
9.8
6.5
7.0
7.5
8.0
8.5
6.0
6.5
7.0
6.5
7.0
7.5
8.0
8.5
6.2
6.4
6.6
6.8
7.0
7.2

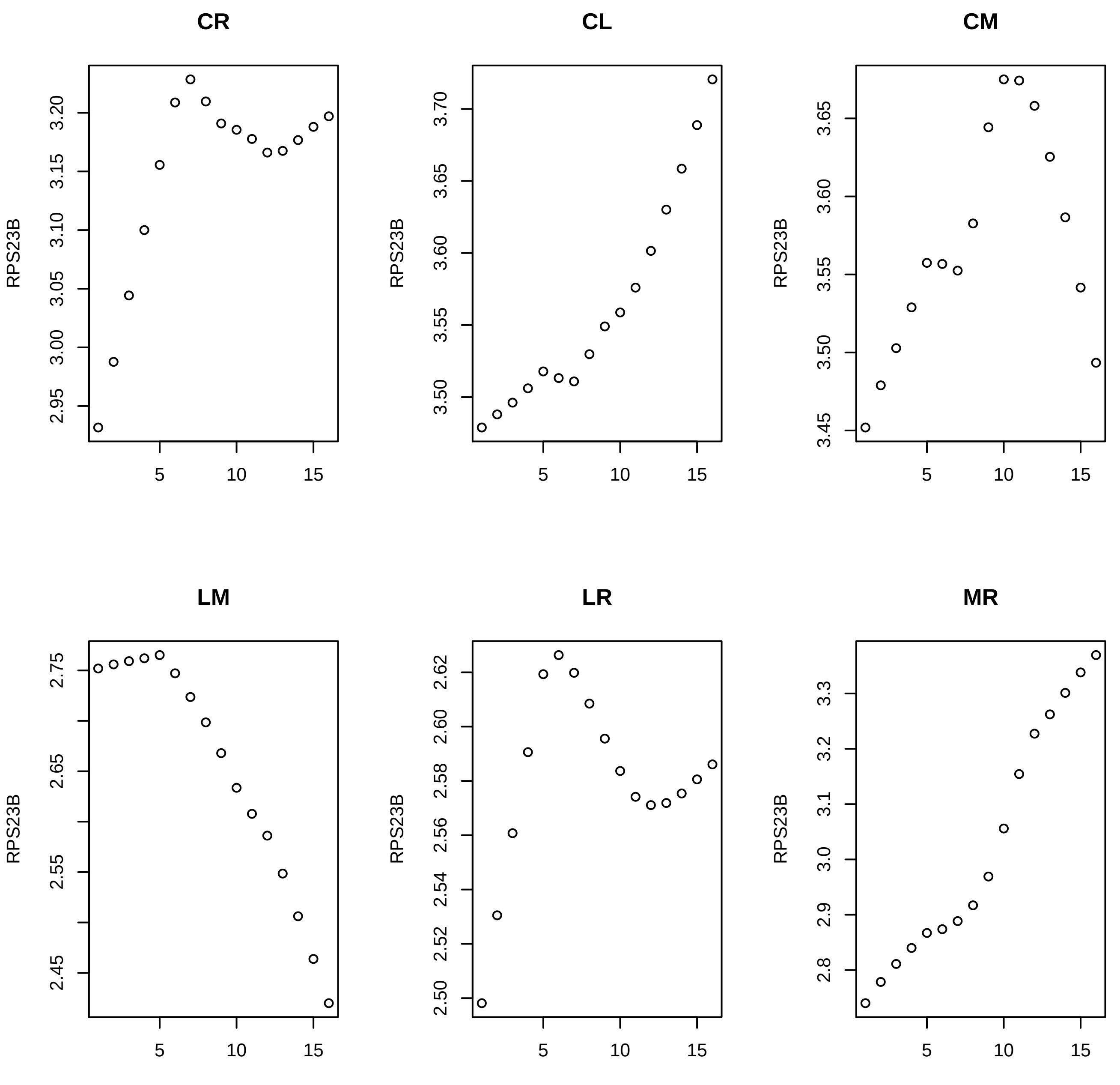
CR
CL
CM
LM
LR
MR
RPS23B
3.20
3.15
3.10
3.05
3.00
2.95
3.70
3.65
3.60
3.55
3.50
3.45
2.75
2.65
2.55
2.45
2.62
2.60
2.58
2.56
2.54
2.52
2.50
3.3
3.2
3.1
3.0
2.9
2.8
5
10
15

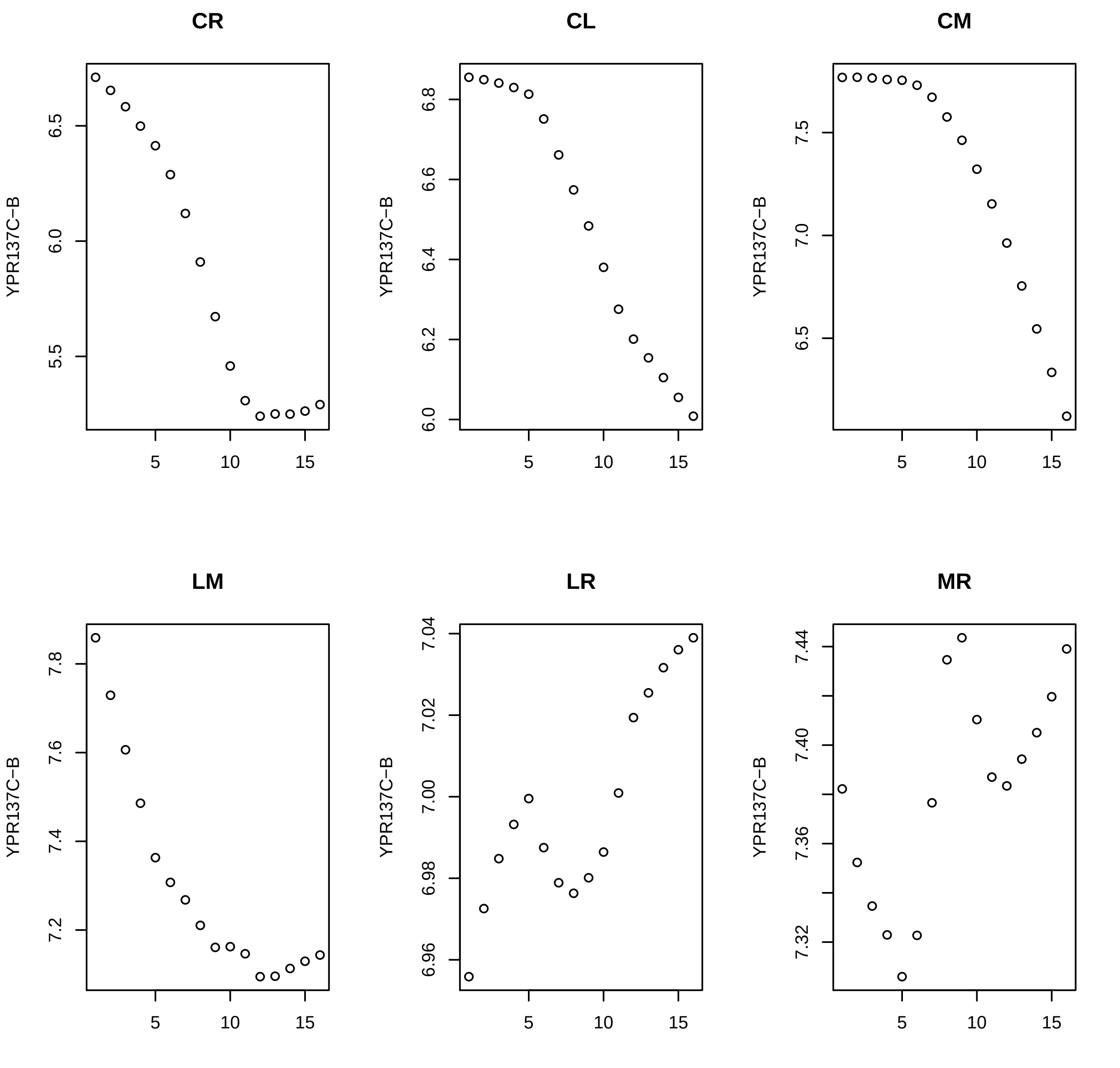
CR
CL
CM
LM
LR
MR
YPR137C−B

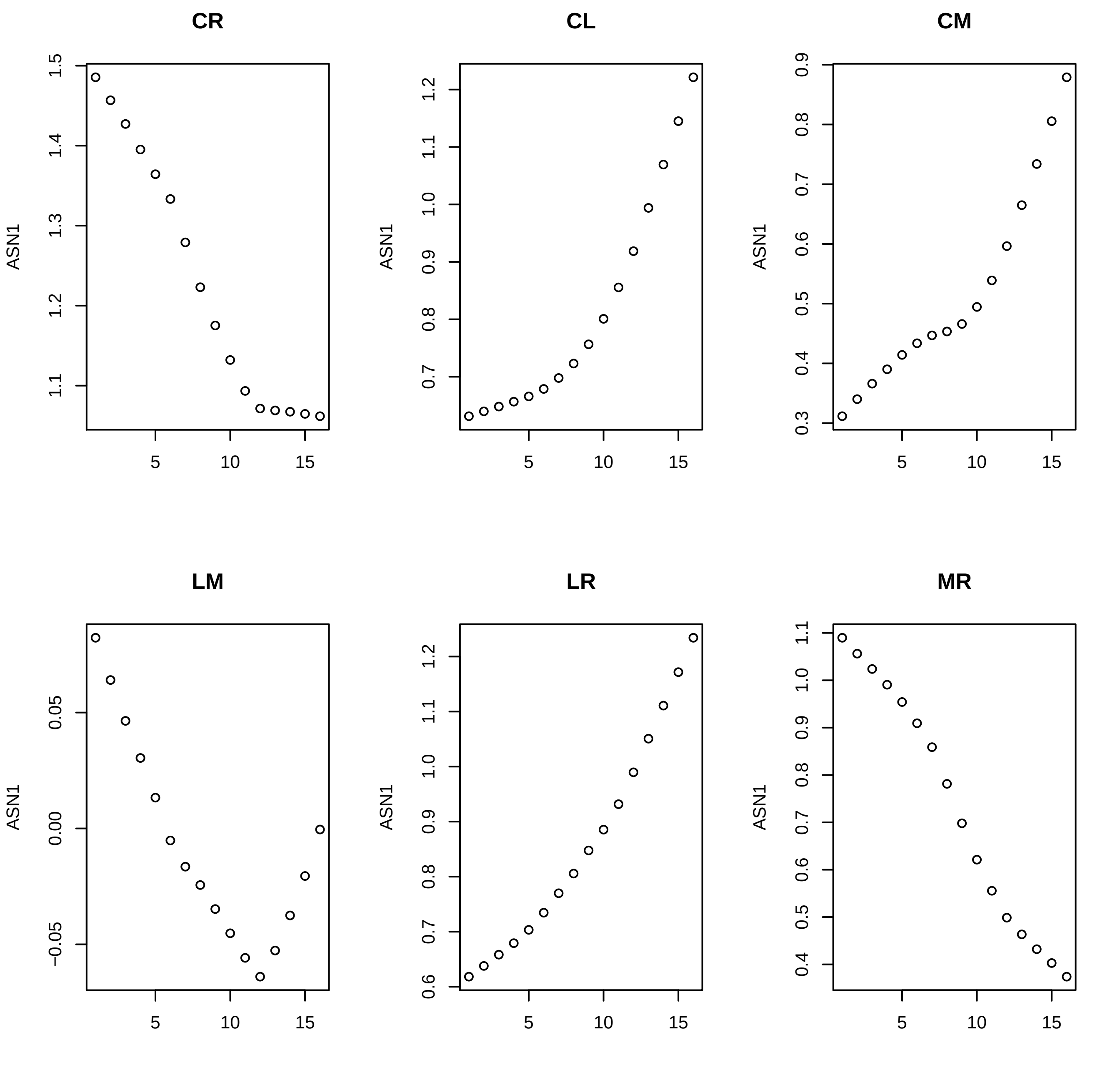

CR
CL
CM
LM
LR
MR
ASN1

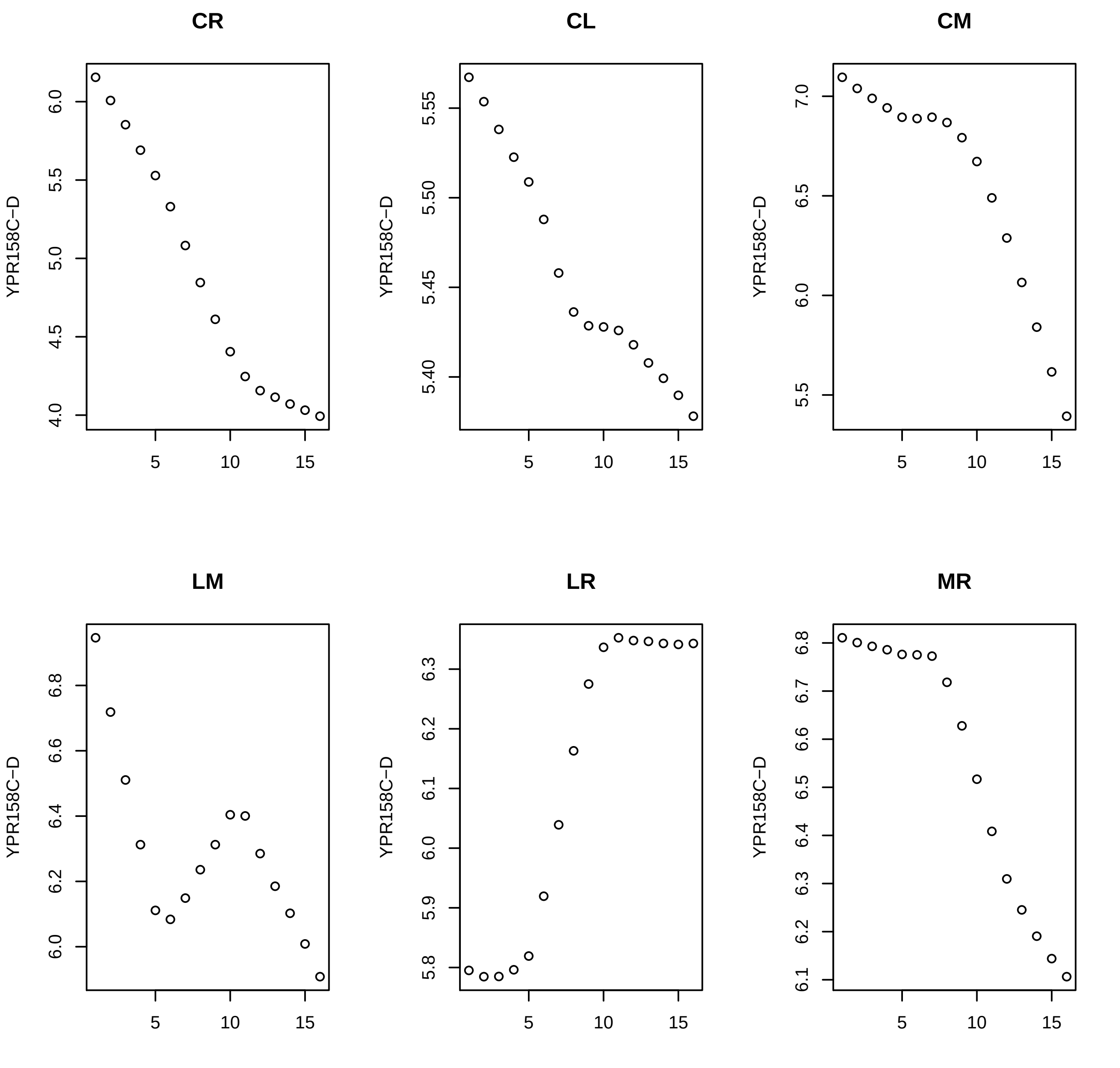

CR
CL
CM
LM
LR
MR
YPR158C−D

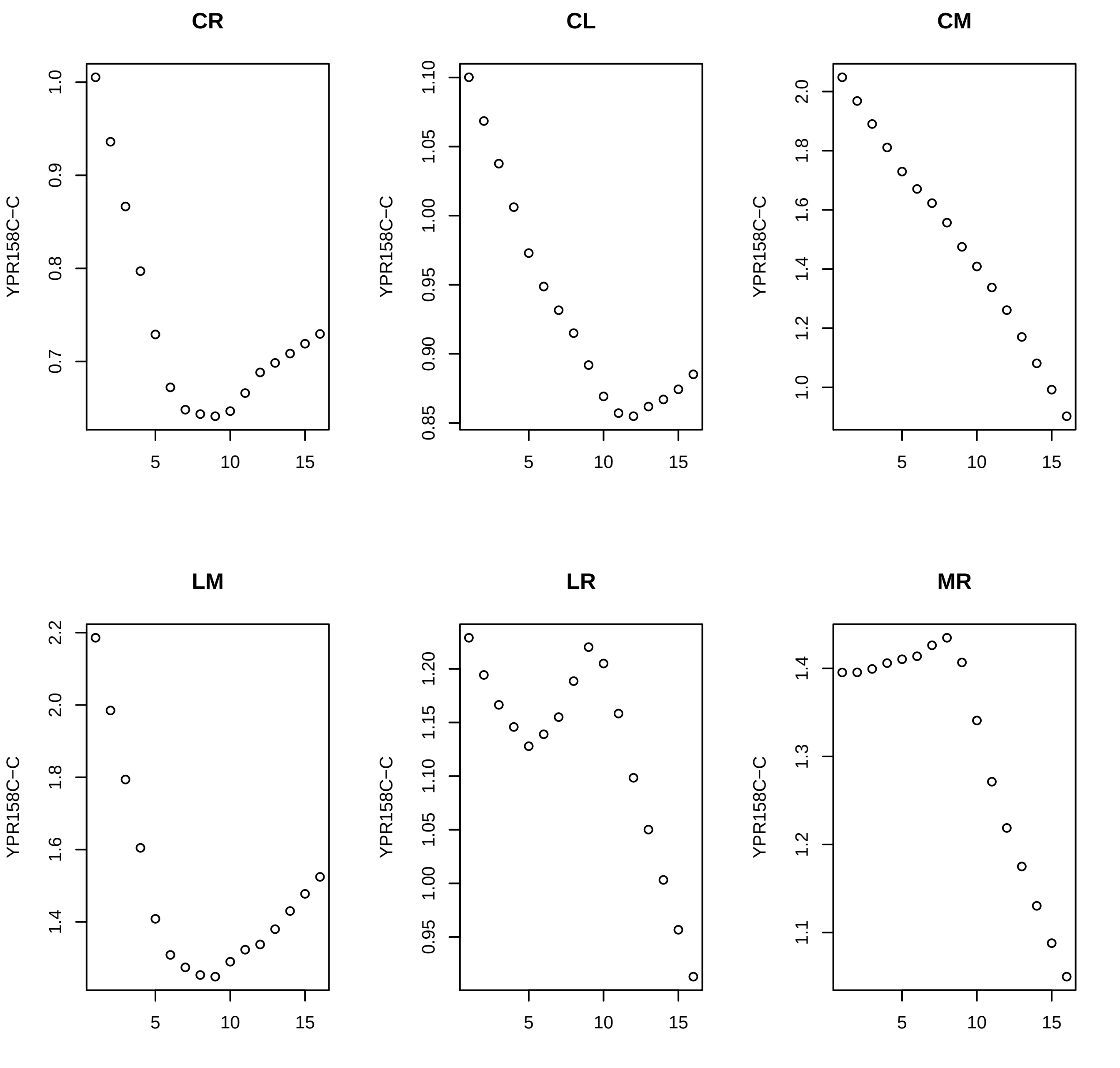
CR
CL
CM
LM
LR
MR
YPR158C−C
1.0
0.9
0.8
0.7
1.10
1.05
1.00
0.95
0.90
0.85
2.0
1.8
1.6
1.4
1.2
1.0
2.2
1.20
1.15
1.10
1.05
0.95
1.3
1.1
5
10
15

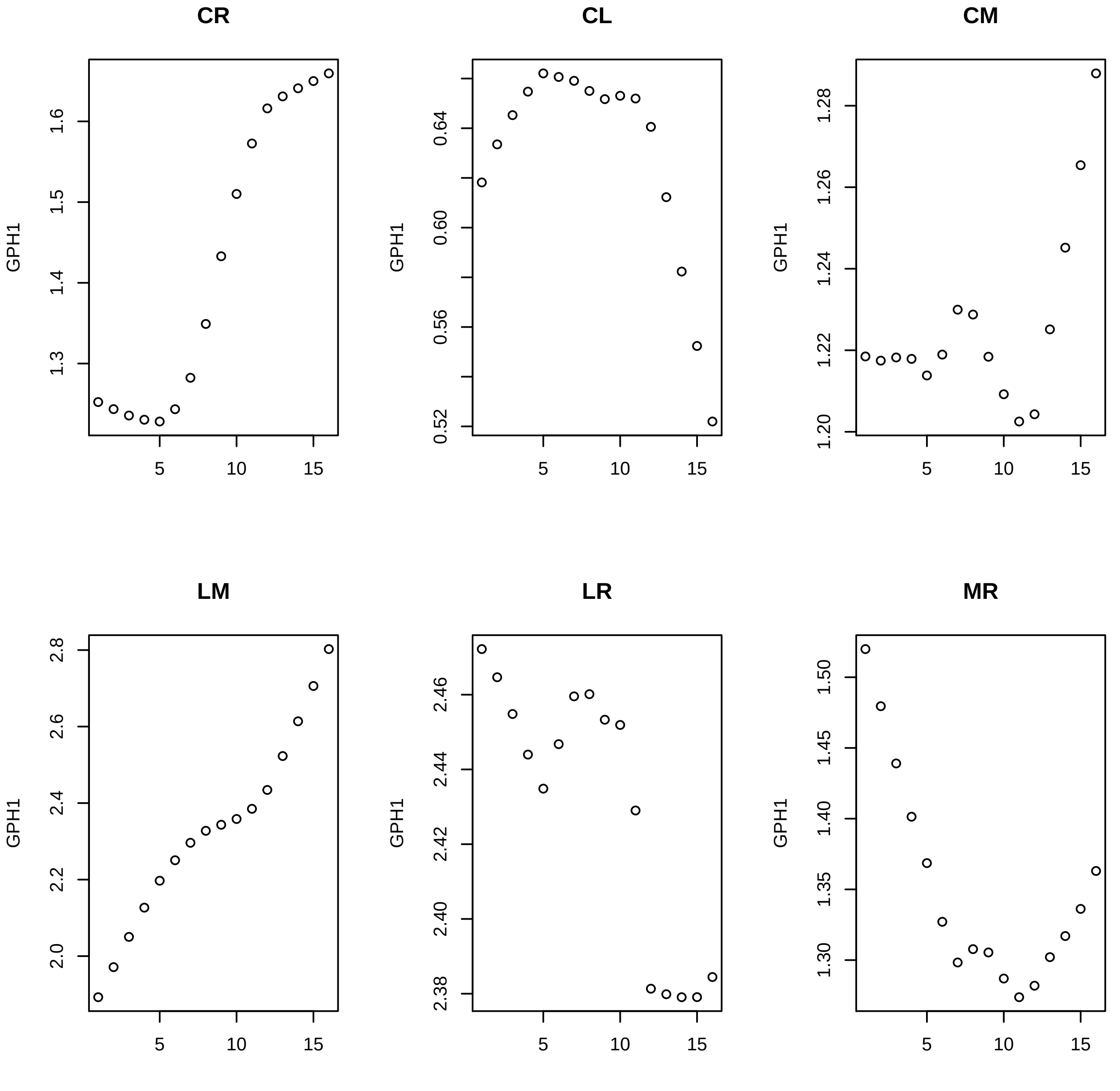

CR
CL
CM
LM
LR
MR
GPH1

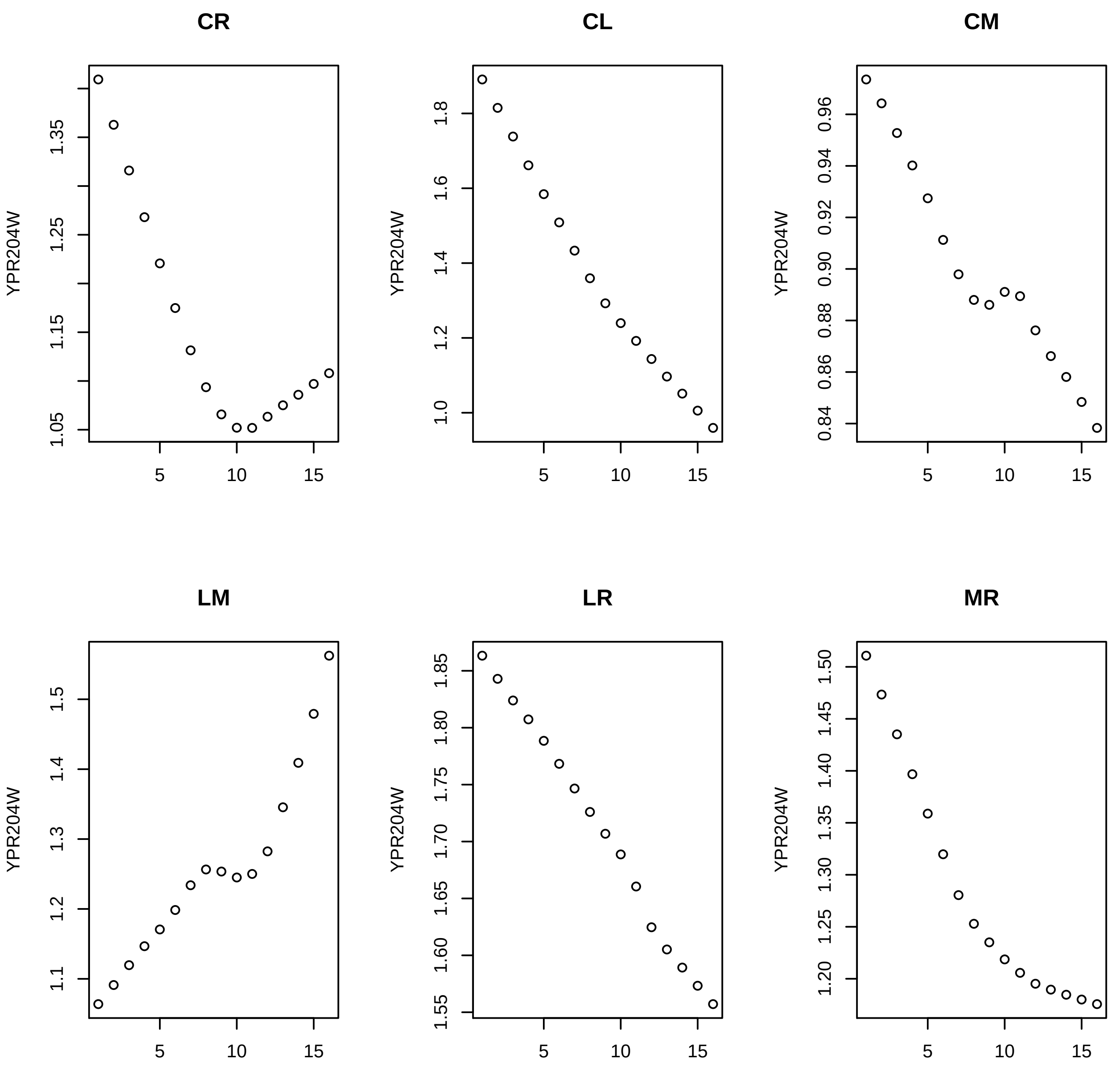
CR
CL
CM
LM
LR
MR
YPR204W
1.05
1.15
1.25
1.35
1.0
1.2
1.4
1.6
1.8
0.84
0.86
0.88
0.90
0.92
0.94
0.96
1.1
1.2
1.3
1.4
1.5
1.55
1.60
1.65
1.70
1.75
1.80
1.85
1.20
1.25
1.30
1.35
1.40
1.45
1.50
5
10
15
