## Supplementary material for "Novel method for the prediction of drug-drug Interaction based on Gene Expression profiles": gene_exp_single.pdf

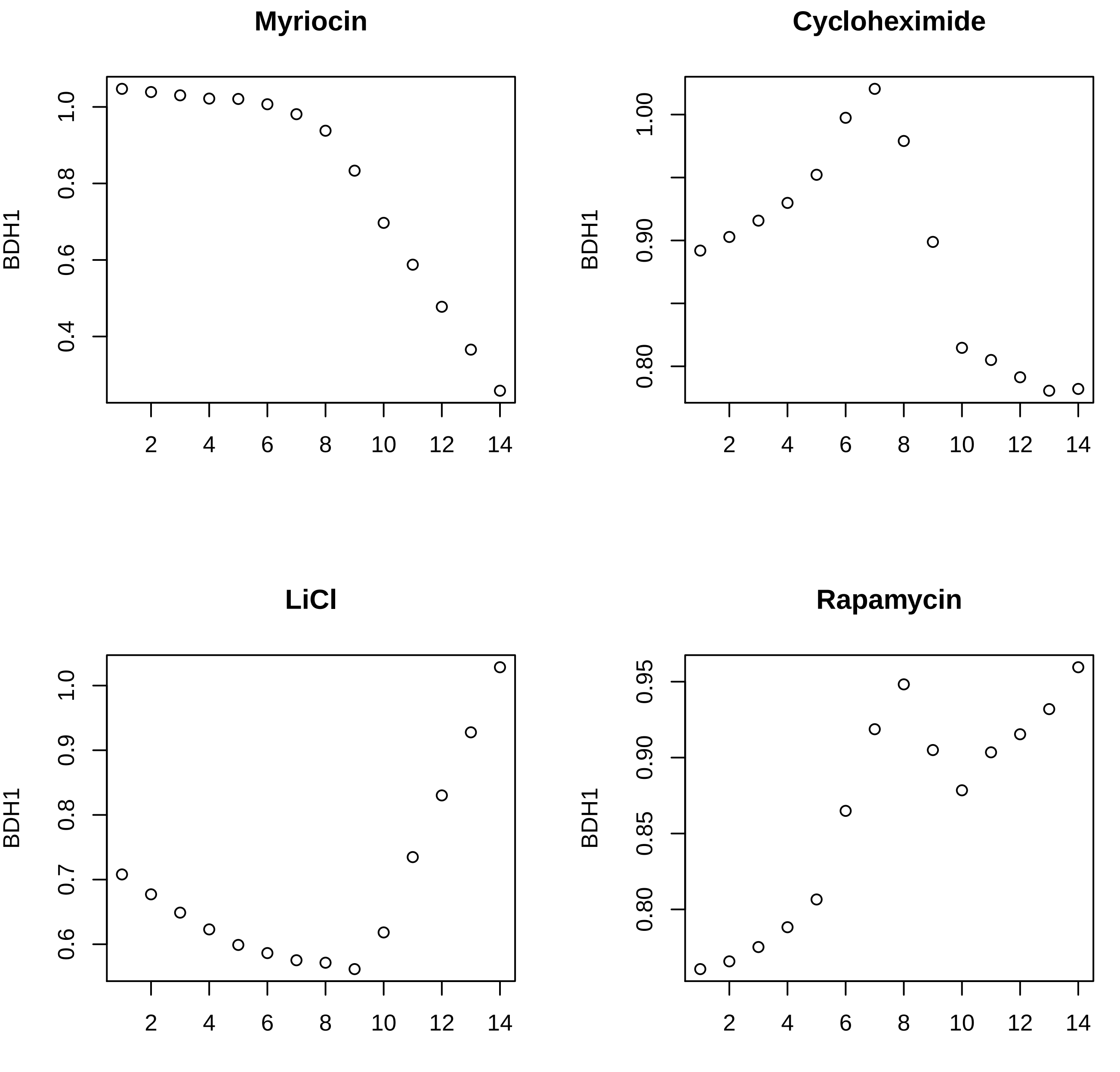
Myriocin
Cycloheximide
LiCl
Rapamycin
BDH1
BDH1
BDH1
BDH1
1.0
0.8
0.6
0.4
1.00
0.90
0.80
1.0
0.9
0.8
0.7
0.6
0.95
0.90
0.85
0.80
2
4
6
8
10
12
14

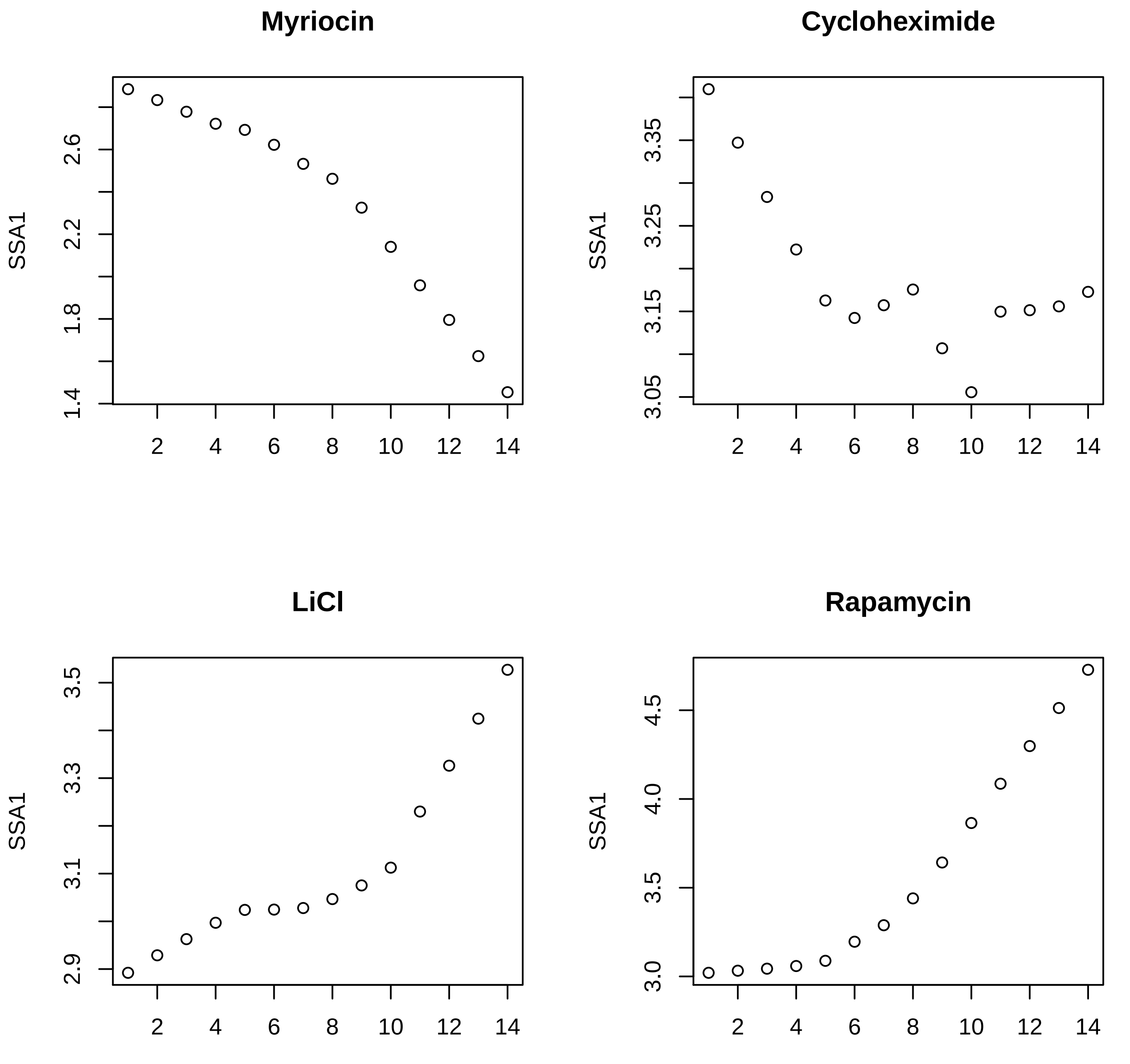
Myriocin
SSA1
2.6
2.2
1.8
1.4
2
4
6
8
10
12
14
Cycloheximide
SSA1
3.35
3.25
3.15
3.05
2
4
6
8
10
12
14
LiCl
SSA1
3.5
3.3
3.1
2.9
2
4
6
8
10
12
14
Rapamycin
SSA1
4.5
4.0
3.5
3.0
2
4
6
8
10
12
14

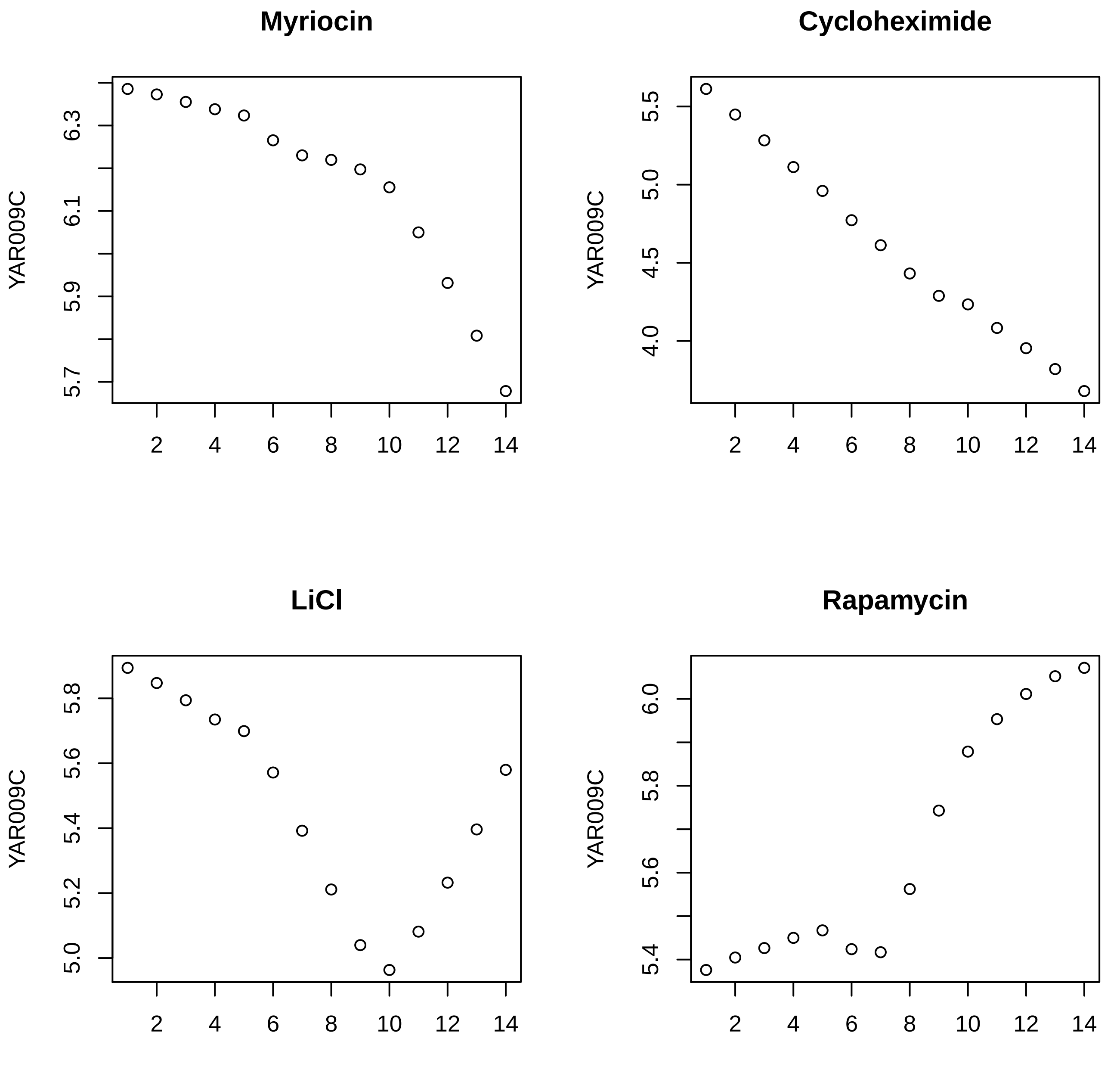
Myriocin
Cycloheximide
LiCl
Rapamycin
YAR009C
YAR009C
YAR009C
YAR009C
5.7
5.9
6.1
6.3
4.0
4.5
5.0
5.5
5.0
5.2
5.4
5.6
5.8
5.4
5.6
5.8
6.0
2
4
6
8
10
12
14

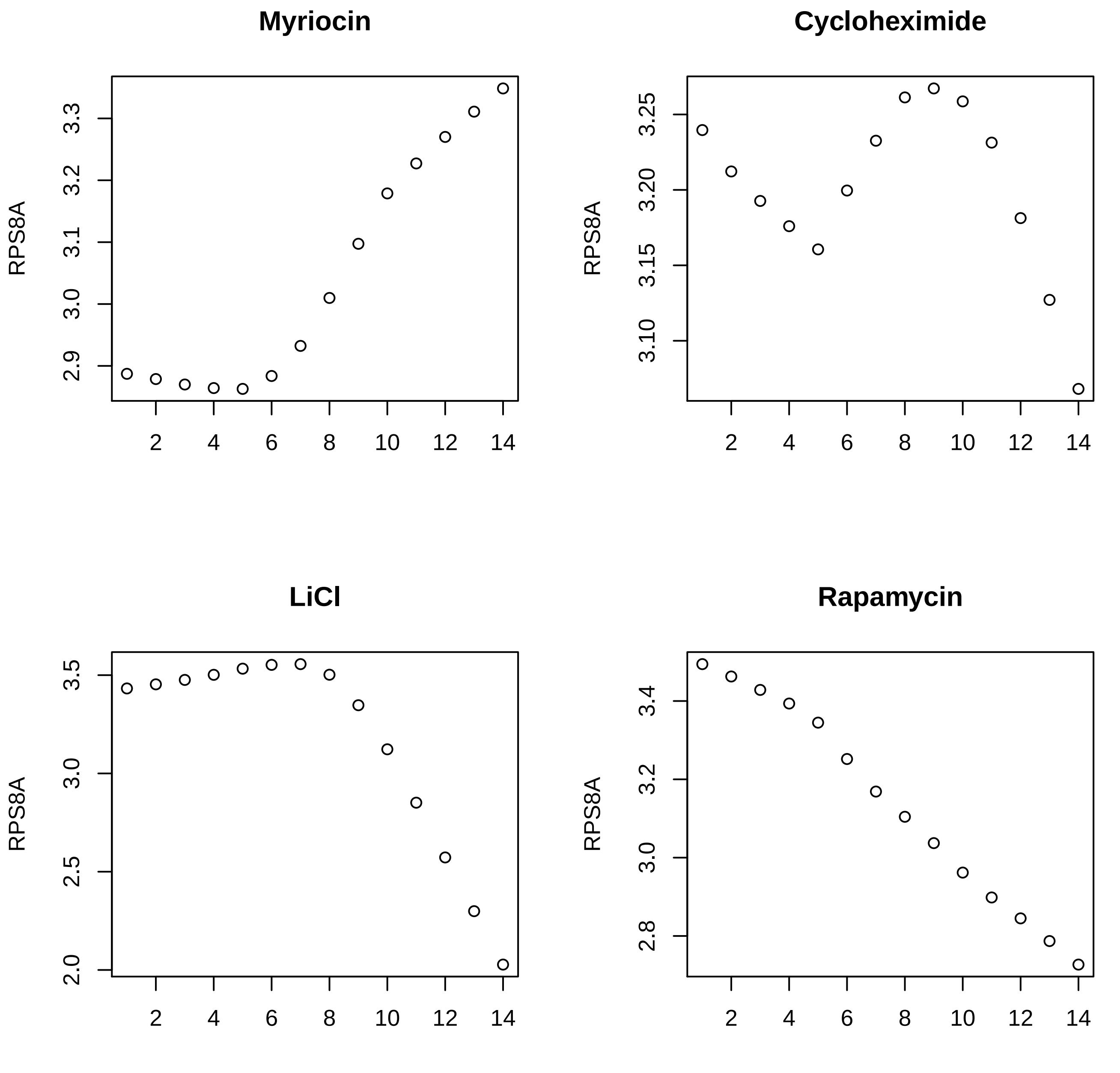
Myriocin
Cycloheximide
LiCl
Rapamycin
RPS8A
2
4
6
8
10
12
14

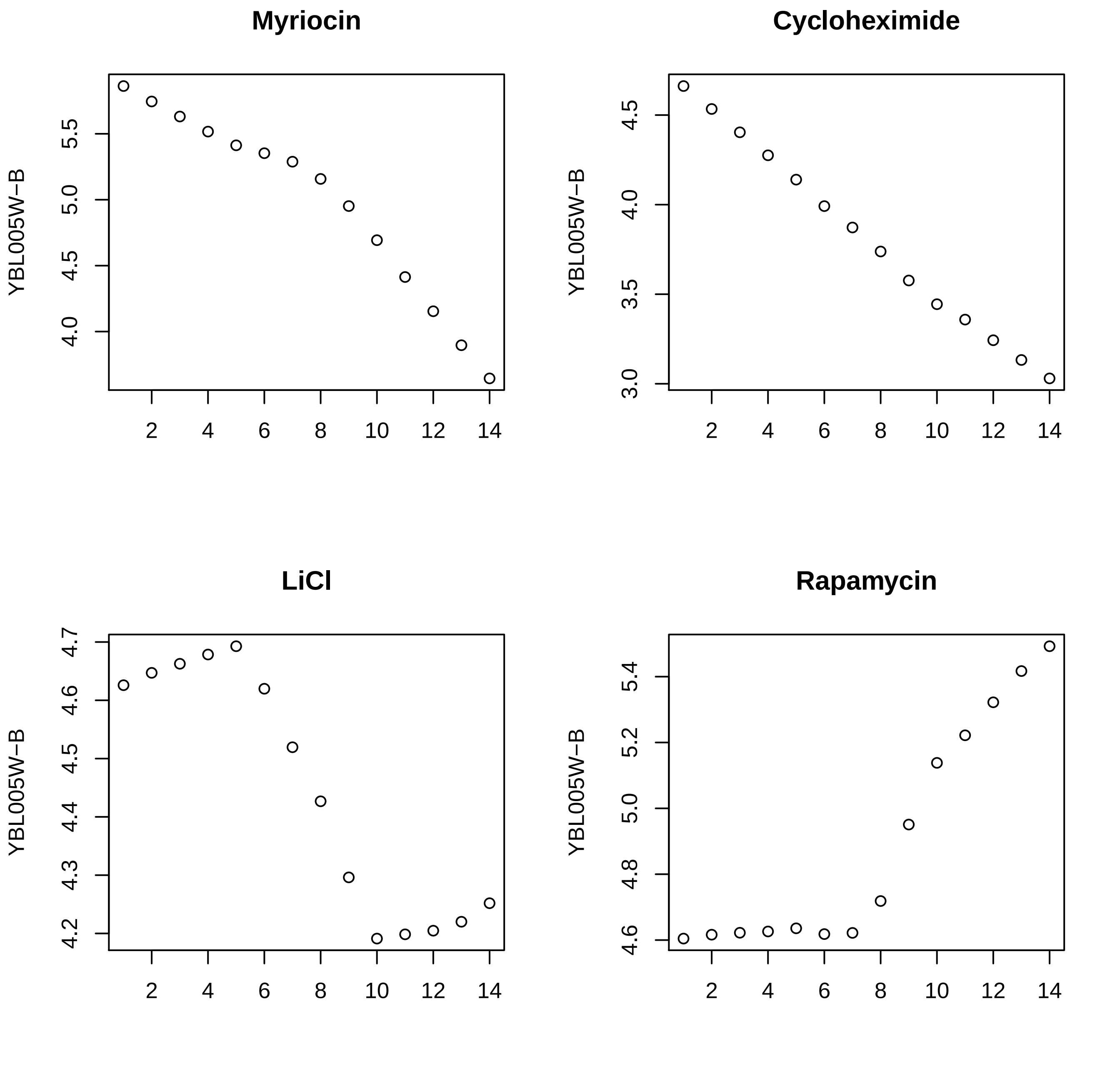

Myriocin
Cycloheximide
LiCl
Rapamycin
YBL005W−B
5.5
5.0
4.5
4.0
3.5
3.0
4.7
4.6
4.4
4.3
4.2
5.4
5.2
4.8
2
4
6
8
10
12
14

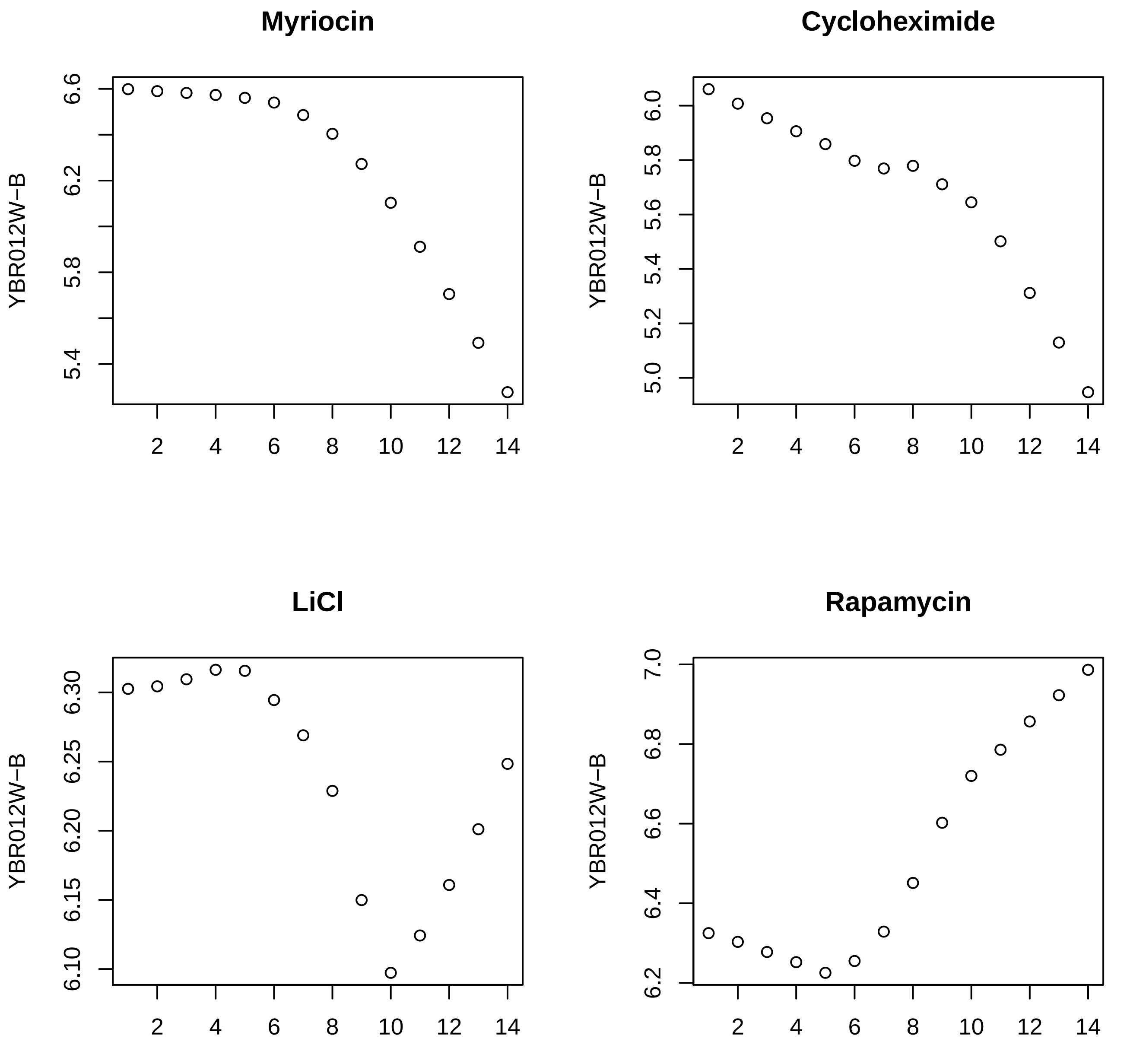
Myriocin
Cycloheximide
LiCl
Rapamycin
YBR012W−B
YBR012W−B
YBR012W−B
YBR012W−B
5.4
5.8
6.2
6.6
5.0
5.2
5.4
5.6
5.8
6.0
6.10
6.15
6.20
6.25
6.30
6.2
6.4
6.6
6.8
7.0
2
4
6
8
10
12
14

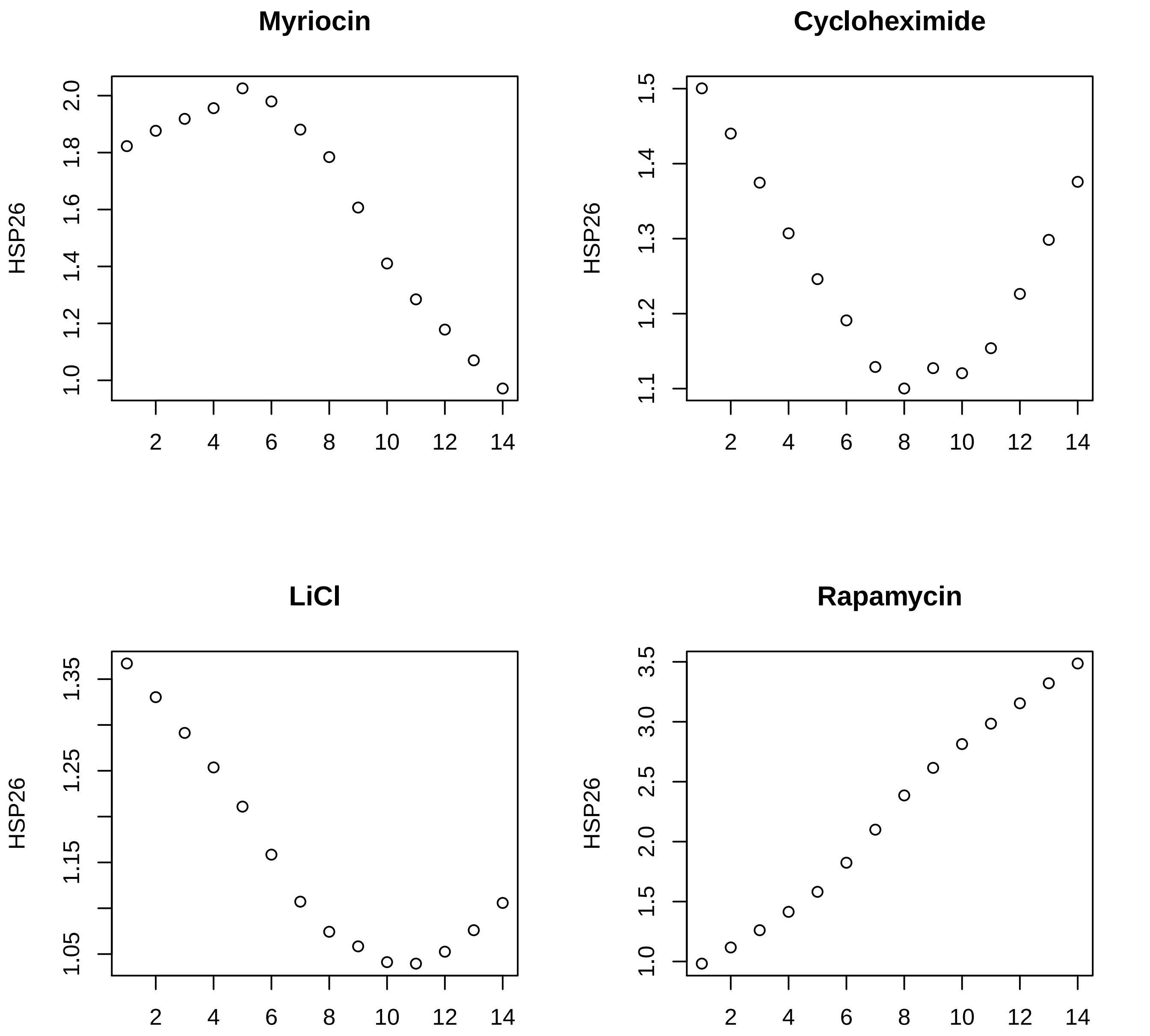
Myriocin
Cycloheximide
LiCl
Rapamycin
HSP26
2.0
1.8
1.6
1.4
1.2
1.0
1.5
1.4
1.3
1.2
1.1
1.35
1.25
1.15
1.05
3.5
3.0
2.5
2.0
1.5
1.0
2
4
6
8
10
12
14

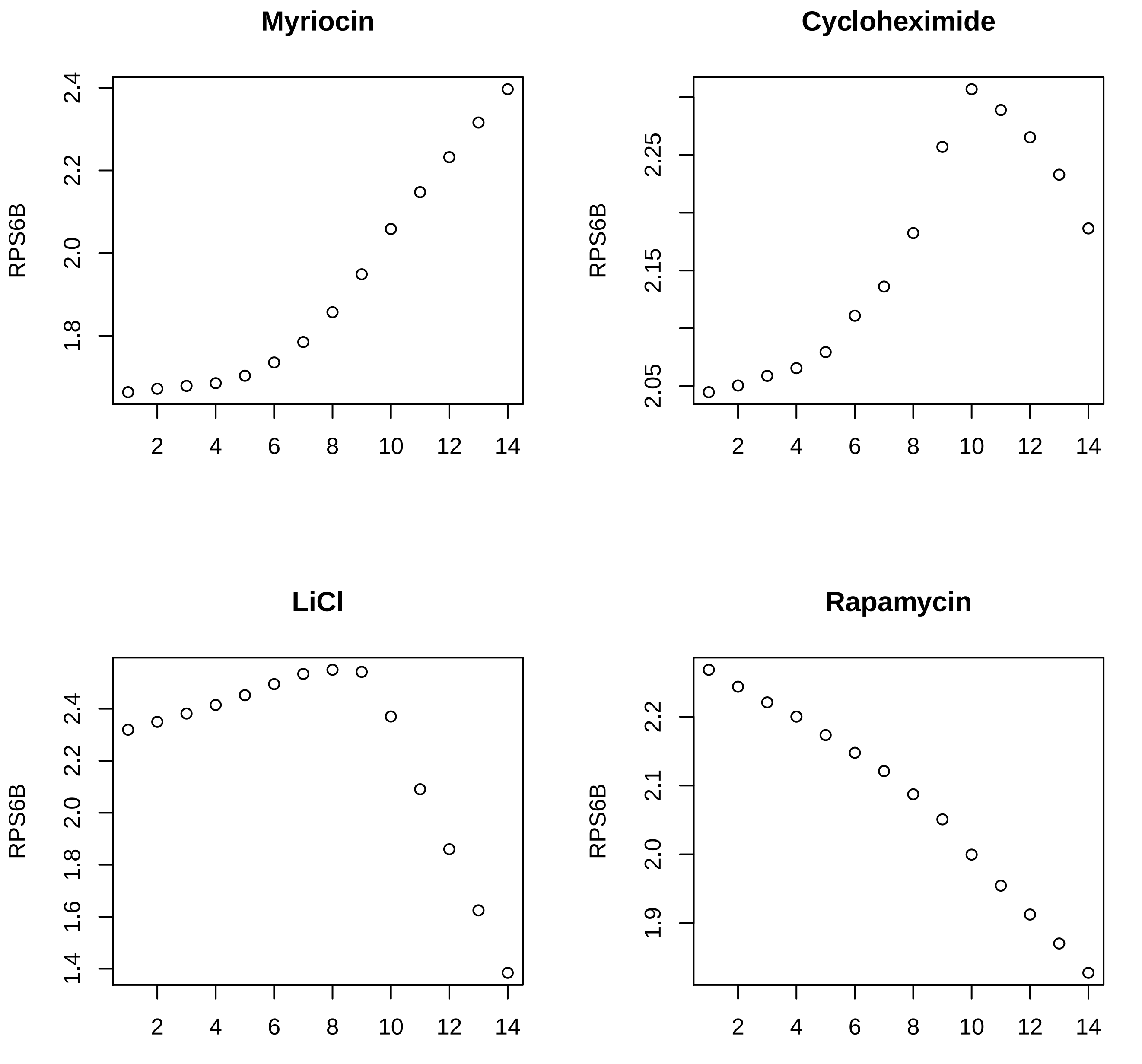

Myriocin
Cycloheximide
LiCl
Rapamycin
RPS6B
RPS6B
RPS6B
RPS6B
2.4
2.2
2.0
1.8
2.25
2.15
2.05
2.4
2.2
2.0
1.8
1.6
1.4
2.2
2.1
2.0
1.9
2
4
6
8
10
12
14

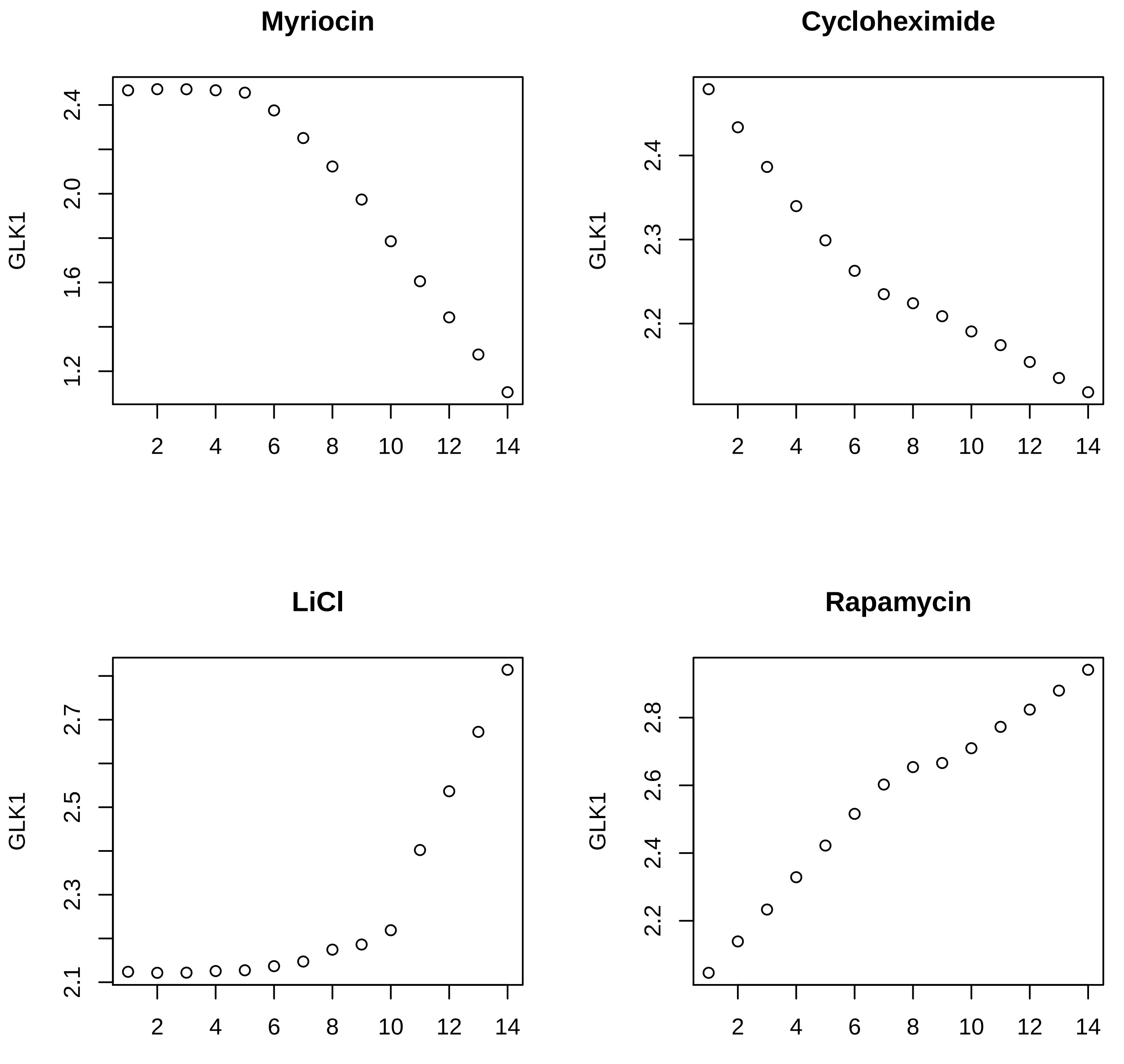

Myriocin
GLK1
2.4
2.0
1.6
1.2
2
4
6
8
10
12
14
Cycloheximide
GLK1
2.4
2.3
2.2
2
4
6
8
10
12
14
LiCl
GLK1
2.7
2.5
2.3
2.1
2
4
6
8
10
12
14
Rapamycin
GLK1
2.8
2.6
2.4
2.2
2
4
6
8
10
12
14

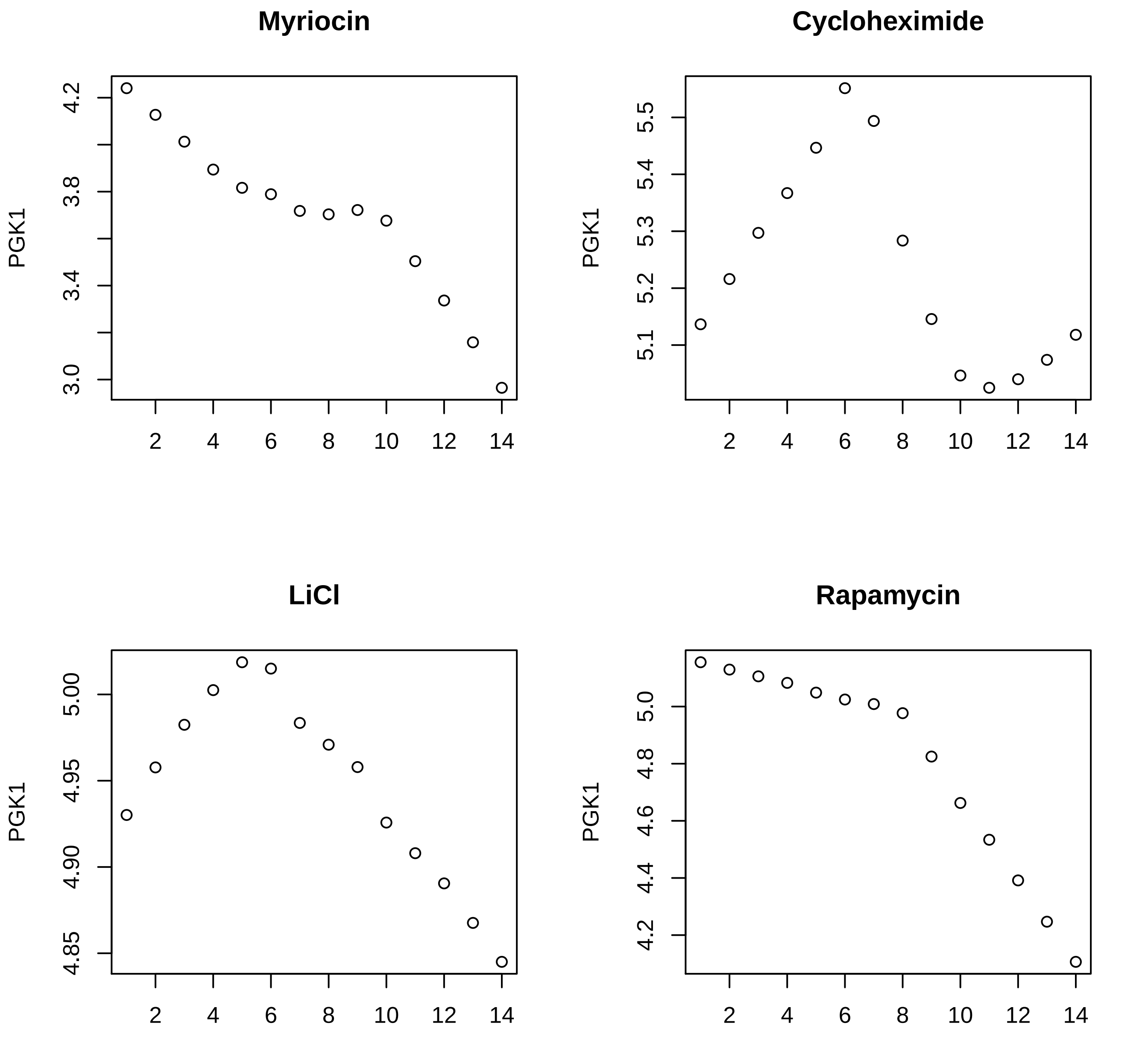
Myriocin
PGK1
4.2
3.8
3.4
3.0
2
4
6
8
10
12
14
Cycloheximide
PGK1
5.5
5.4
5.3
5.2
5.1
2
4
6
8
10
12
14
LiCl
PGK1
5.00
4.95
4.90
4.85
2
4
6
8
10
12
14
Rapamycin
PGK1
5.0
4.8
4.6
4.4
4.2
2
4
6
8
10
12
14

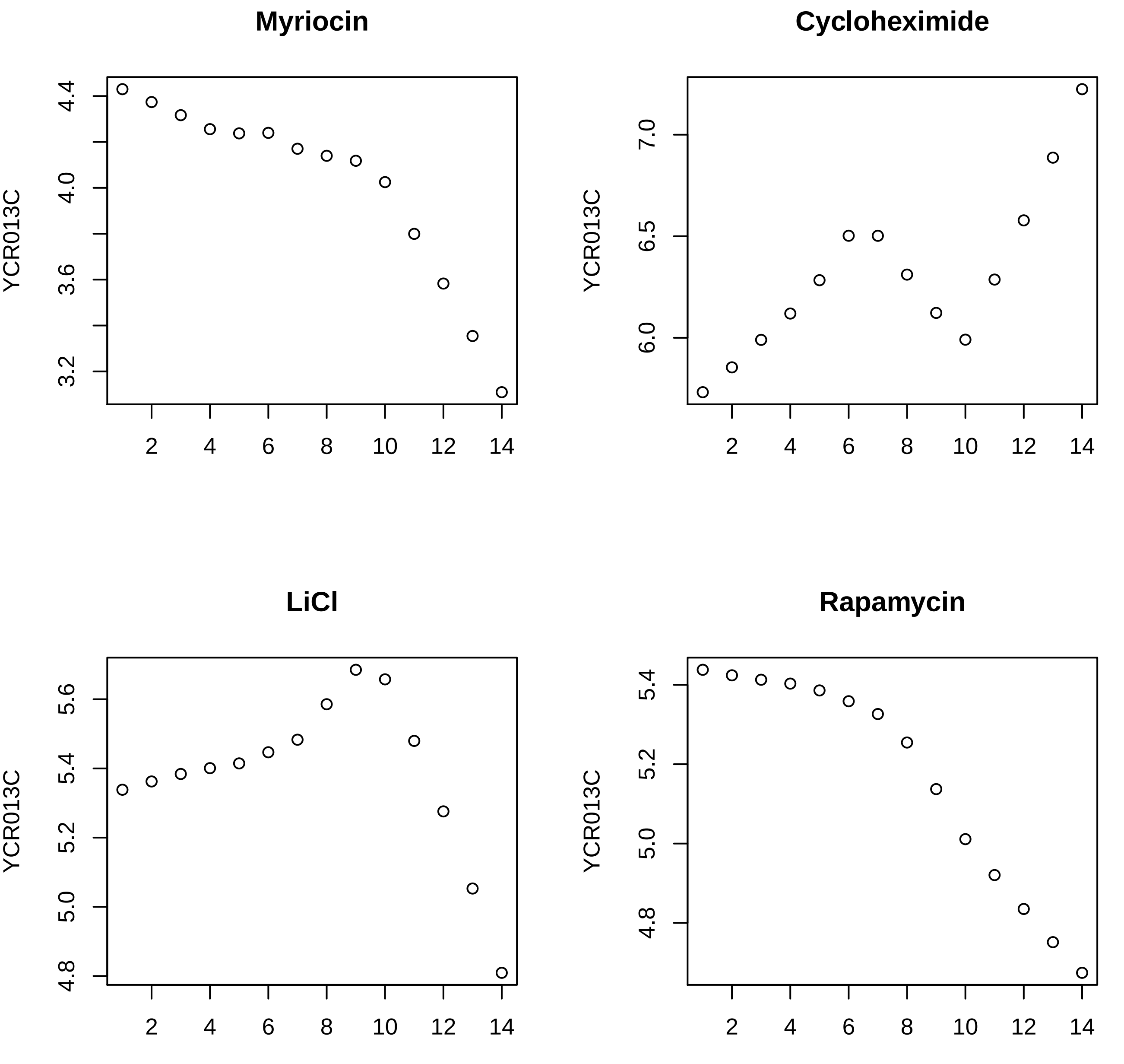
Myriocin
YCR013C
4.4
4.0
3.6
3.2
2
4
6
8
10
12
14
Cycloheximide
YCR013C
7.0
6.5
6.0
2
4
6
8
10
12
14
LiCl
YCR013C
5.6
5.4
5.2
5.0
4.8
2
4
6
8
10
12
14
Rapamycin
YCR013C
5.4
5.2
5.0
4.8
2
4
6
8
10
12
14

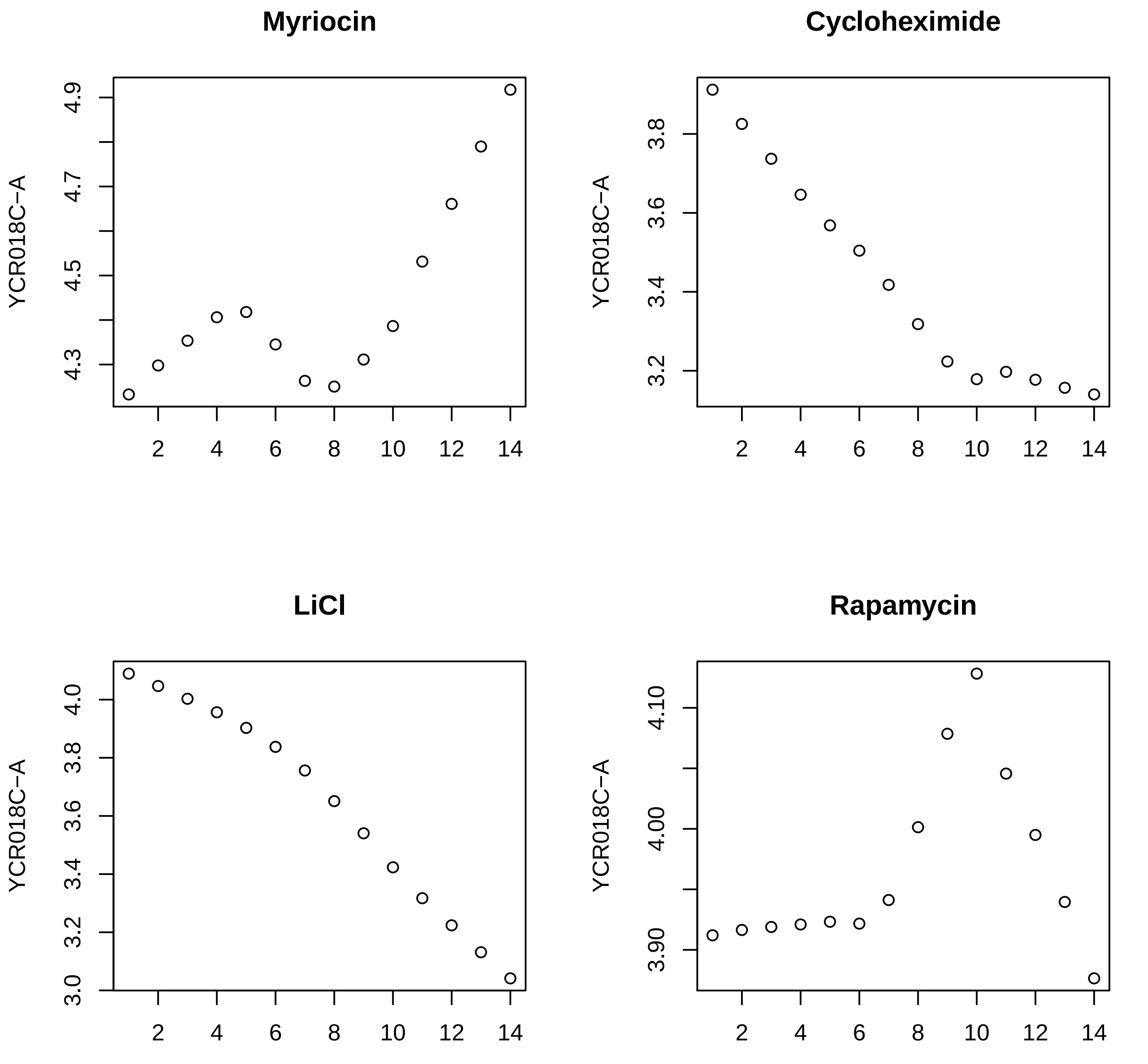

Myriocin
Cycloheximide
LiCl
Rapamycin
YCR018C−A
4.9
4.7
4.5
4.3
3.8
3.6
3.4
3.2
4.0
3.0
4.10
4.00
3.90
2
4
6
8
10
12
14

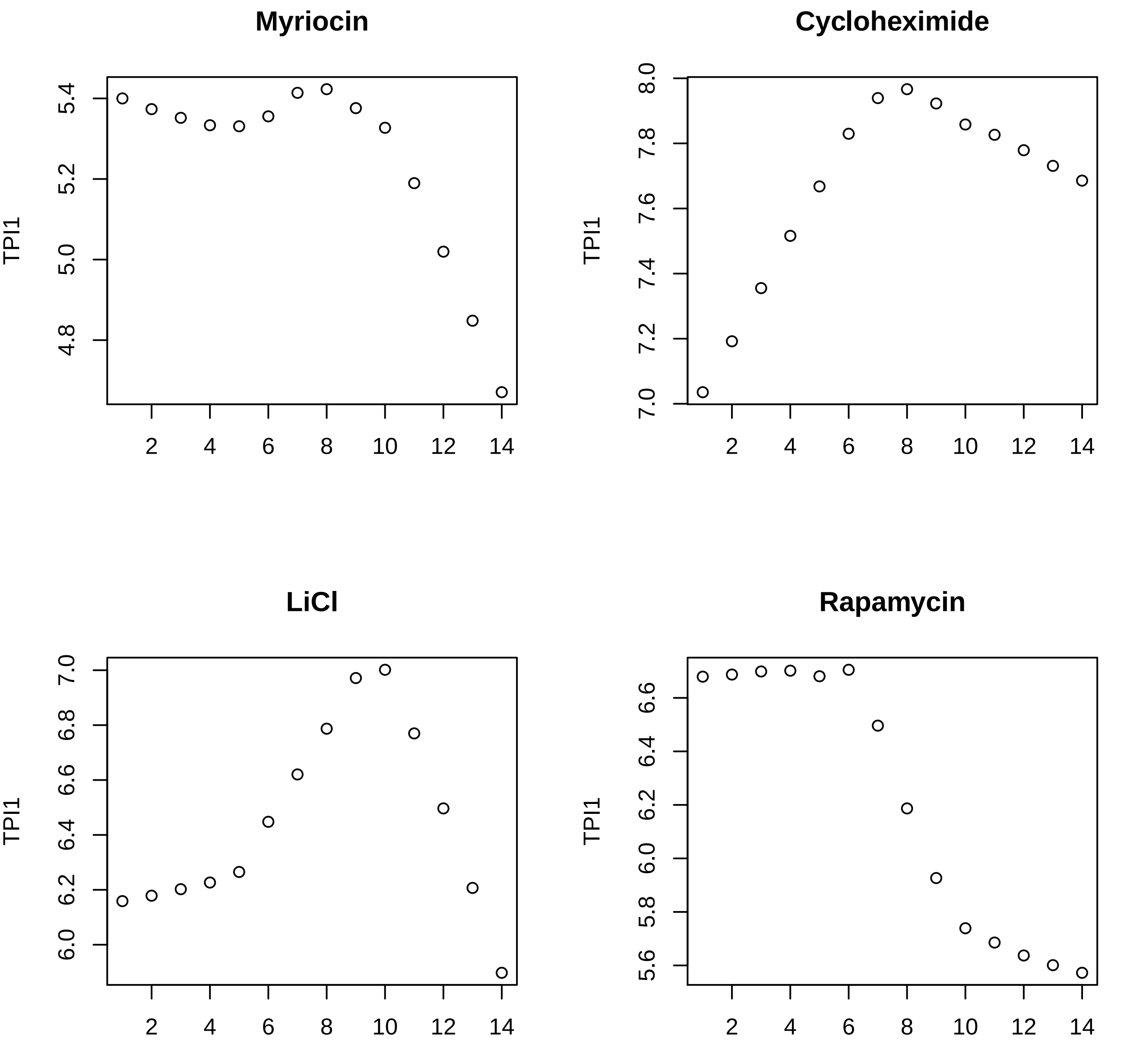
Myriocin
Cyclohexomide
LiCl
Rapamycin
TPI1
5.4
5.2
5.0
4.8
8.0
7.8
7.6
7.4
7.2
7.0
6.8
6.6
6.4
6.2
6.0
5.8
5.6
2
4
6
8
10
12
14

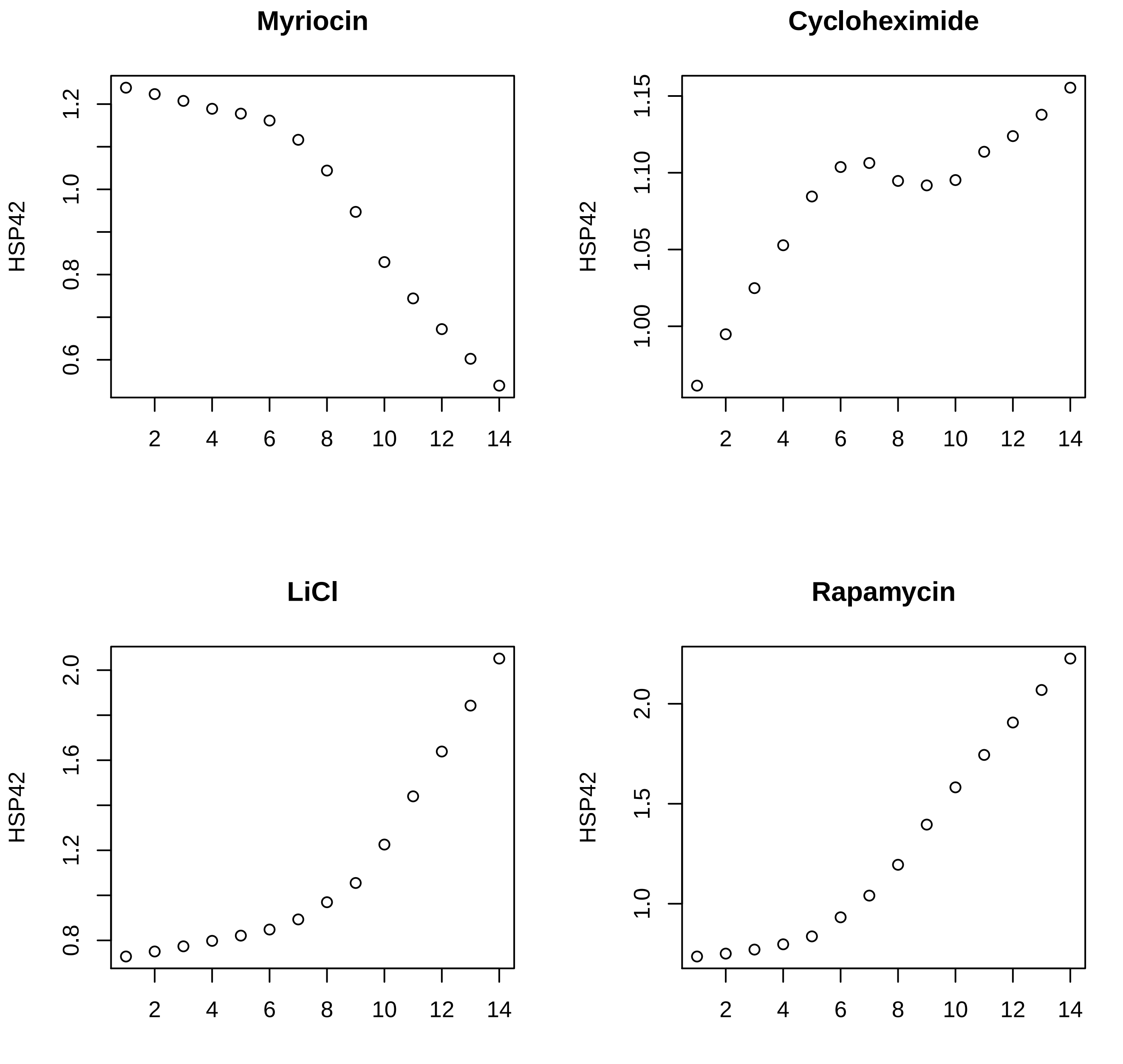
Myriocin
HSP42
0.6
0.8
1.0
1.2
2
4
6
8
10
12
14
Cycloheximide
HSP42
1.00
1.05
1.10
1.15
2
4
6
8
10
12
14
LiCl
HSP42
0.8
1.2
1.6
2.0
2
4
6
8
10
12
14
Rapamycin
HSP42
1.0
1.5
2.0
2
4
6
8
10
12
14

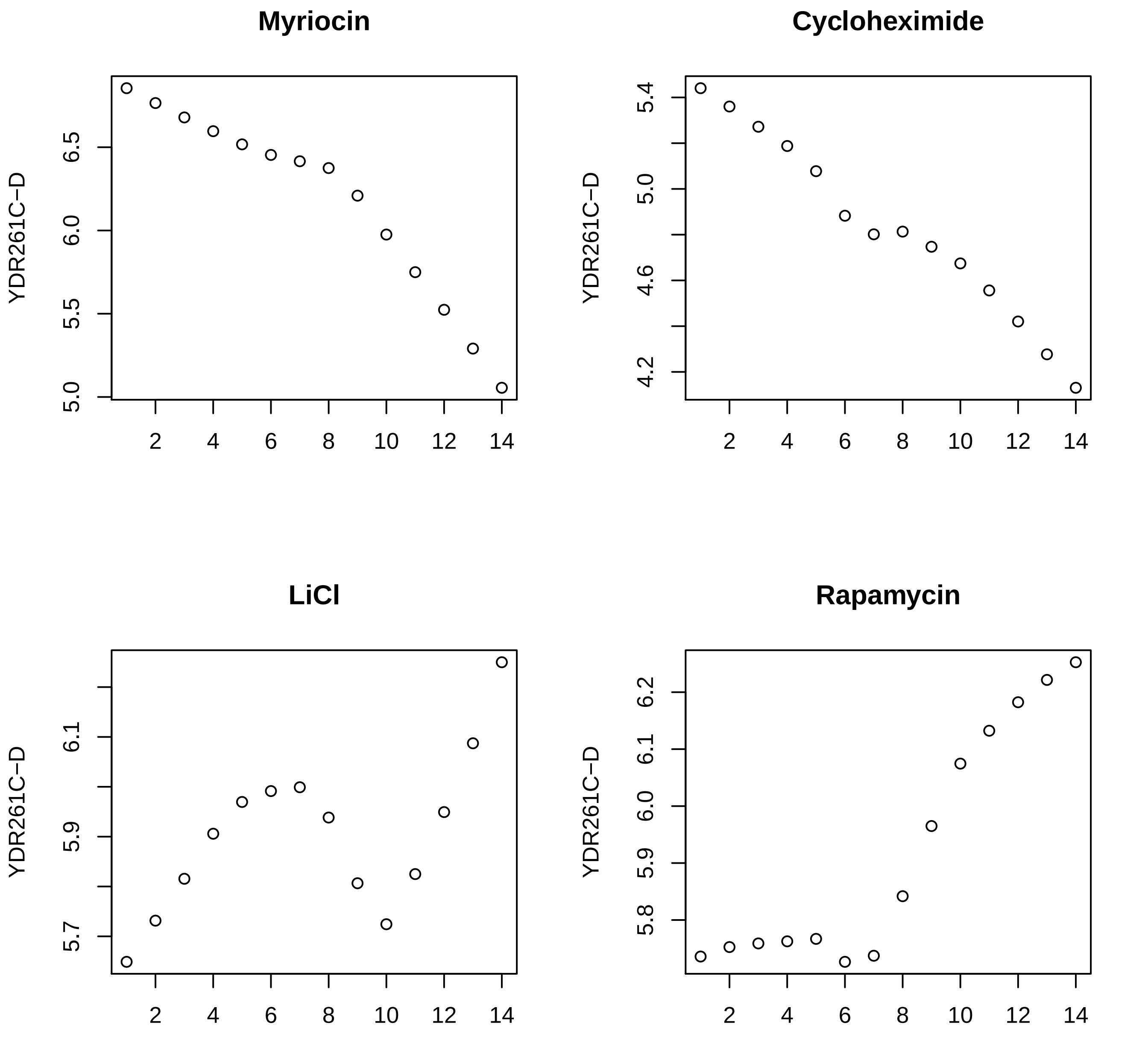

Myriocin
Cycloheximide
LiCl
Rapamycin
YDR261C−D
5.0
5.5
6.0
6.5
5.4
5.0
4.6
4.2
6.1
5.9
5.7
6.2
6.1
6.0
5.9
5.8
2
4
6
8
10
12
14

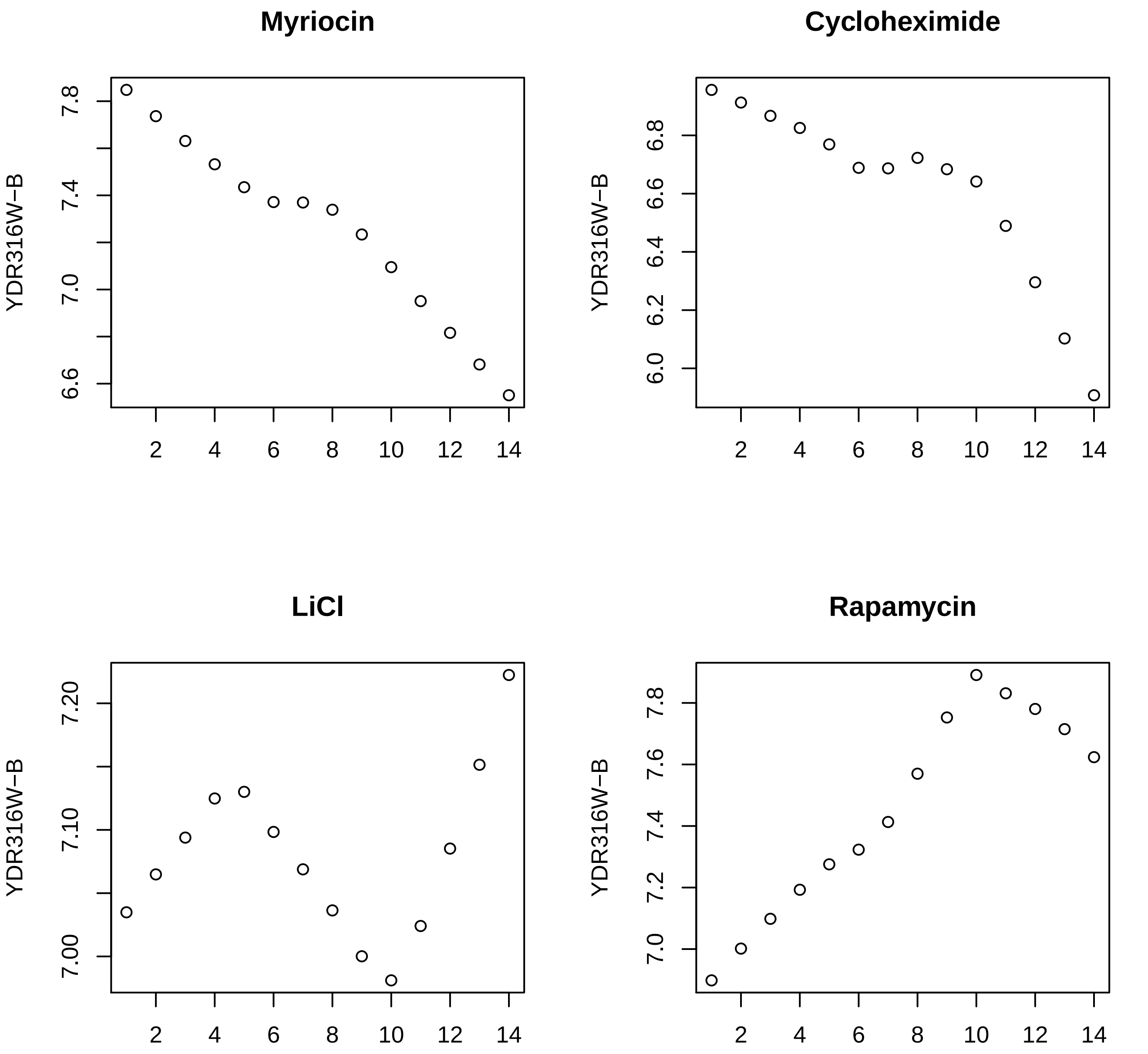
Myriocin
Cycloheximide
LiCl
Rapamycin
YDR316W−B
7.8
7.4
7.0
6.6
6.8
6.6
6.4
6.2
6.0
7.20
7.10
7.00
7.8
7.6
7.4
7.2
7.0
2
4
6
8
10
12
14

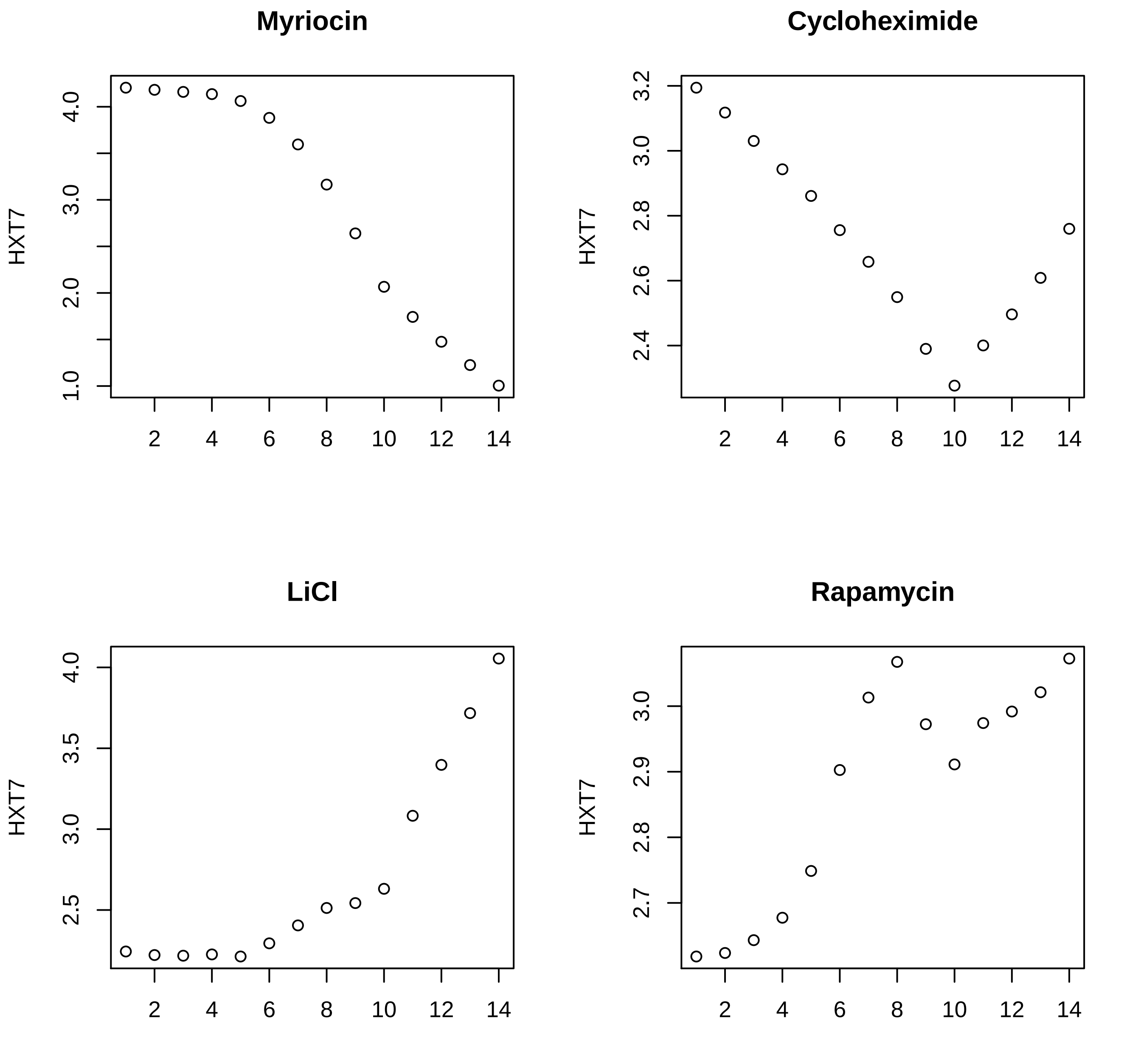
Myriocin
HXT7
1.0
2.0
3.0
4.0
2
4
6
8
10
12
14
Cycloheximide
HXT7
2.4
2.6
2.8
3.0
3.2
2
4
6
8
10
12
14
LiCl
HXT7
2.5
3.0
3.5
4.0
2
4
6
8
10
12
14
Rapamycin
HXT7
2.7
2.8
2.9
3.0
2
4
6
8
10
12
14

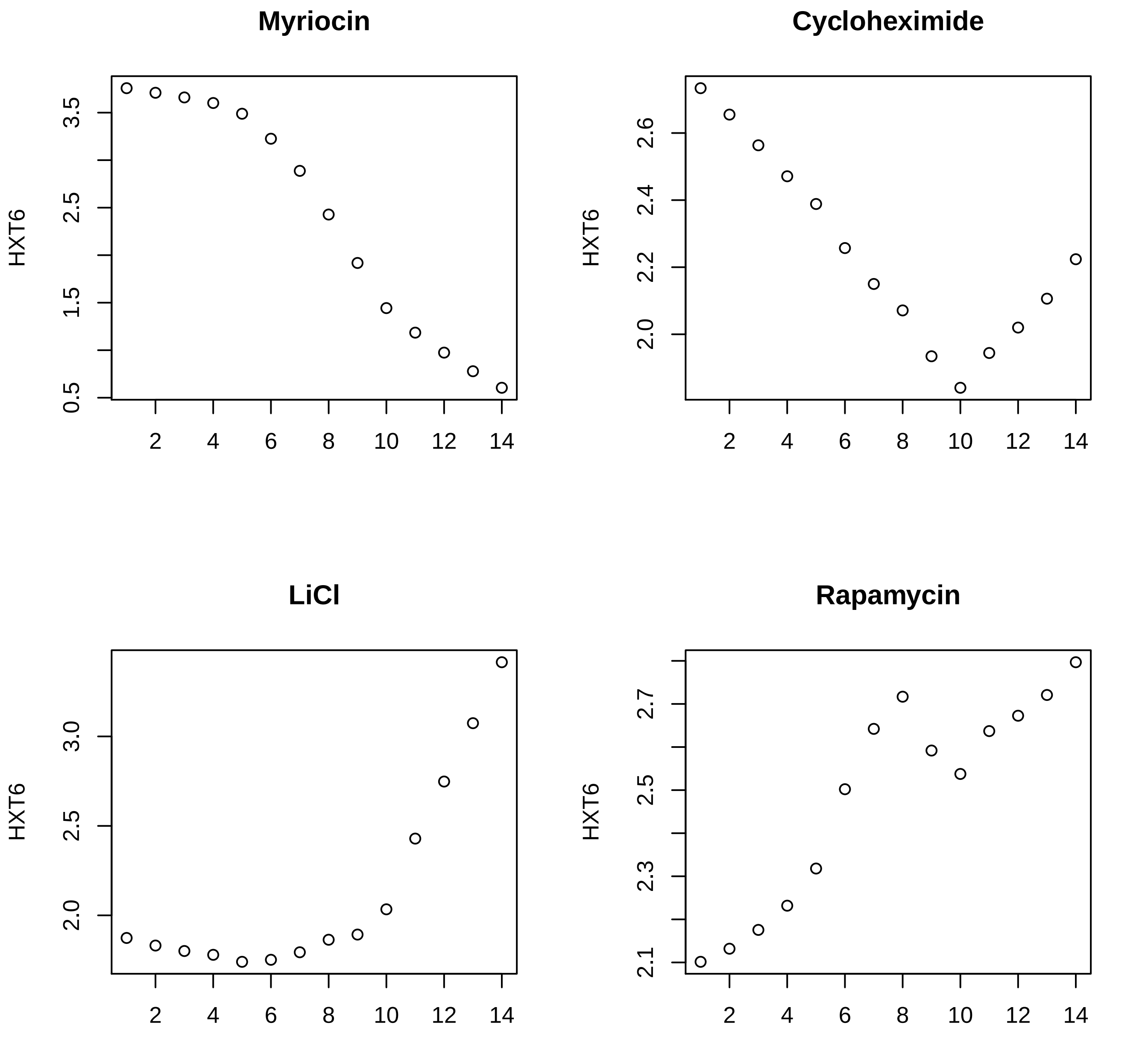
Myriocin
Cycloheximide
LiCl
Rapamycin
HXT6
0.5
1.5
2.5
3.5
2.0
2.2
2.4
2.6
2.0
2.5
3.0
2.1
2.3
2.5
2.7
2
4
6
8
10
12
14

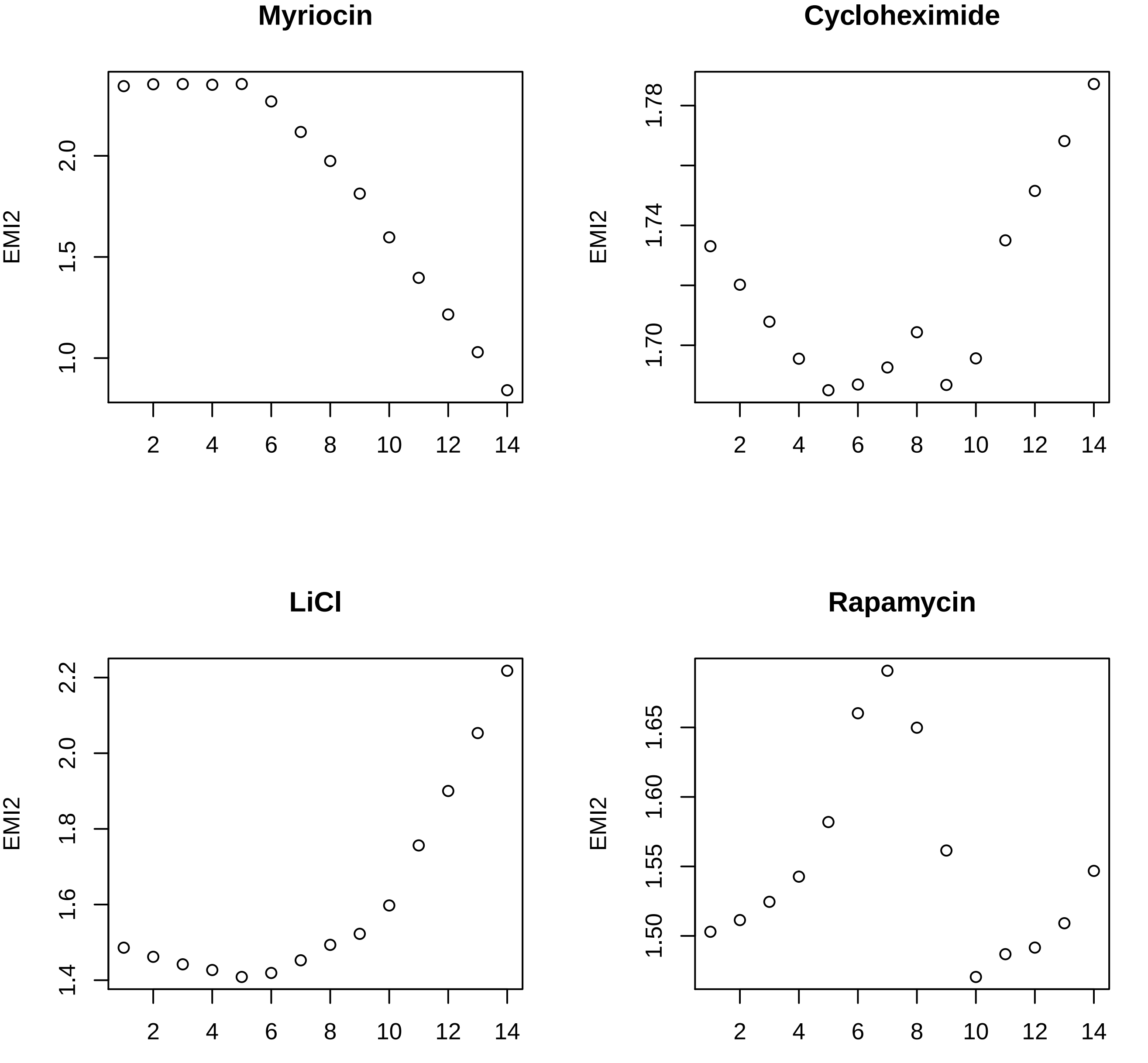

Myriocin
Cycloheximide
LiCl
Rapamycin
EMI2
2.0
1.5
1.0
1.78
1.74
1.70
2.2
2.0
1.8
1.6
1.4
1.65
1.60
1.55
1.50
2
4
6
8
10
12
14

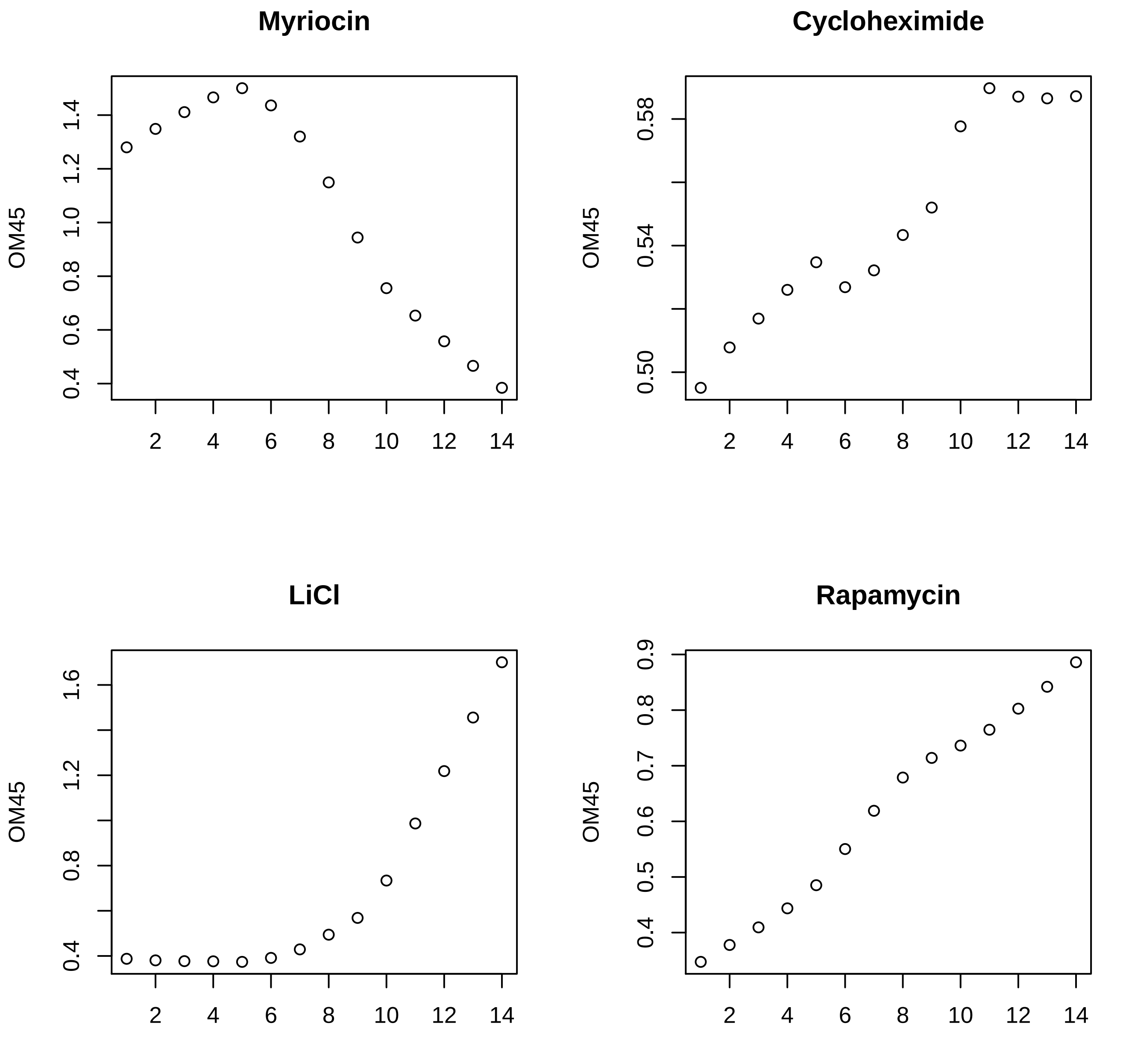

Myriocin
OM45
0.4
0.6
0.8
1.0
1.2
1.4
2
4
6
8
10
12
14
Cycloheximide
OM45
0.50
0.54
0.58
2
4
6
8
10
12
14
LiCl
OM45
0.4
0.8
1.2
1.6
2
4
6
8
10
12
14
Rapamycin
OM45
0.4
0.5
0.6
0.7
0.8
0.9
2
4
6
8
10
12
14

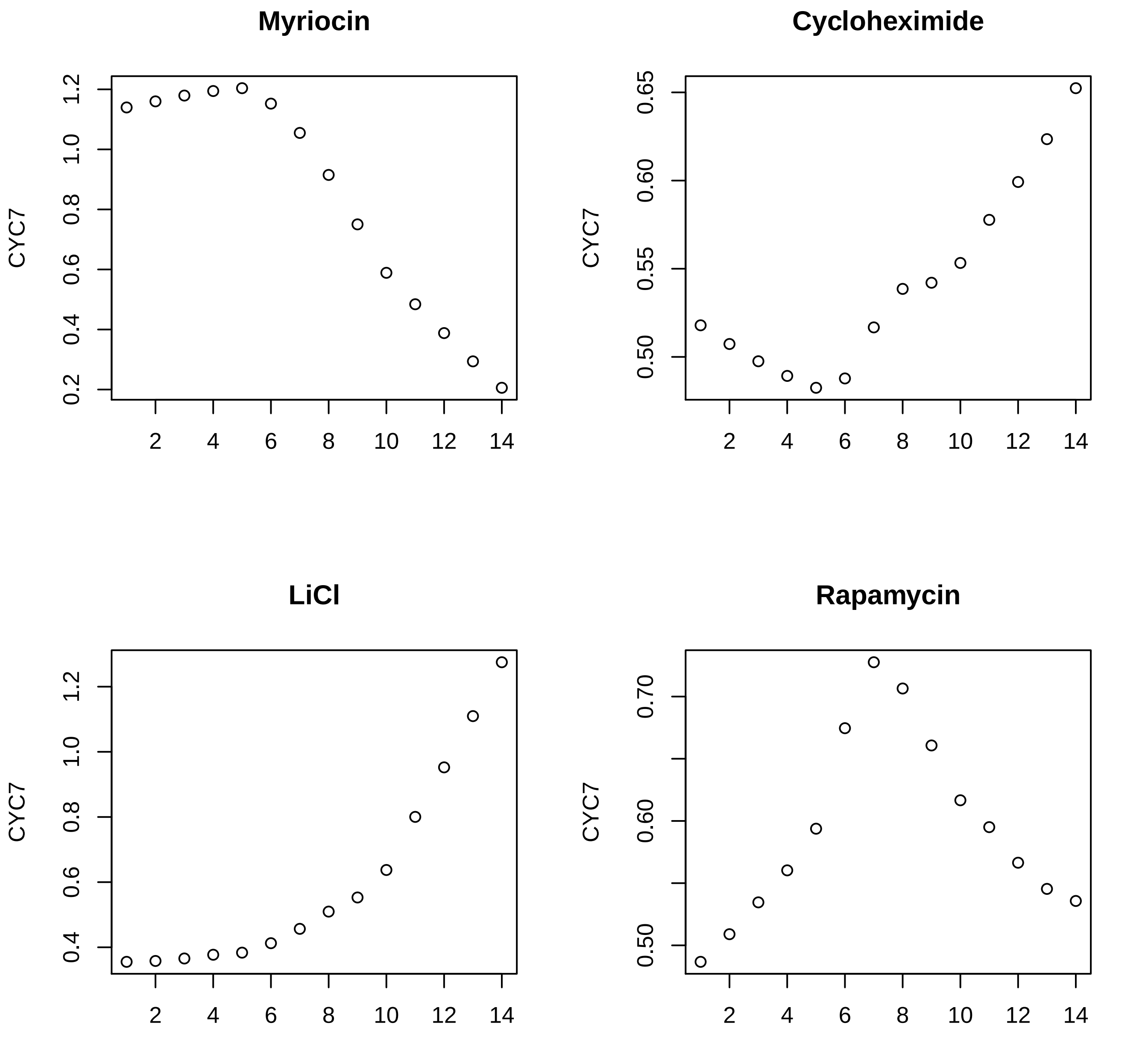
Myriocin
Cycloheximide
LiCl
Rapamycin
CYC7
0.2
0.4
0.6
0.8
1.0
1.2
0.50
0.55
0.60
0.65
0.70
2
4
6
8
10
12
14

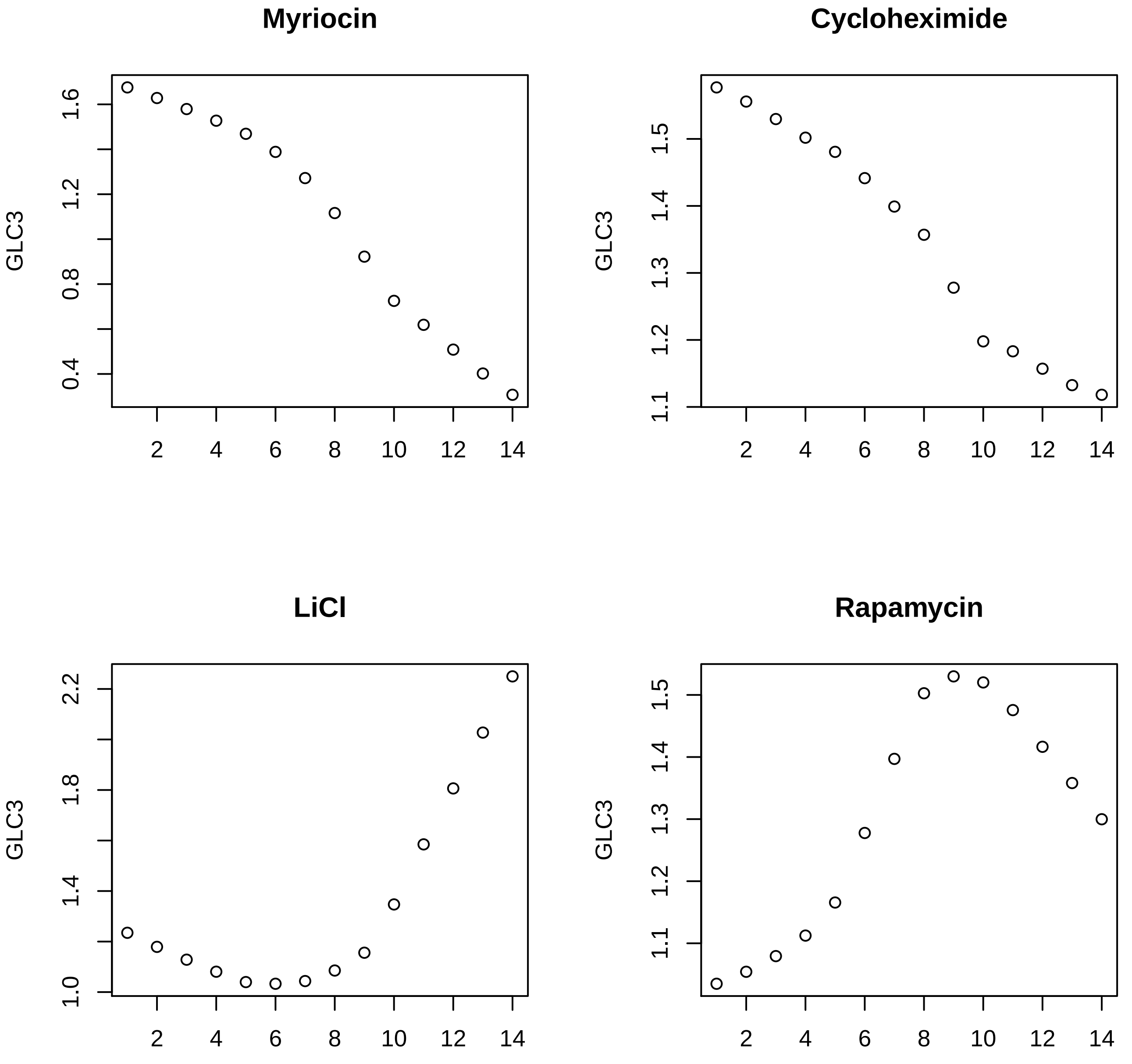
Myriocin
GLC3
0.4
0.8
1.2
1.6
2
4
6
8
10
12
14
Cycloheximide
GLC3
1.1
1.2
1.3
1.4
1.5
2
4
6
8
10
12
14
LiCl
GLC3
1.0
1.4
1.8
2.2
2
4
6
8
10
12
14
Rapamycin
GLC3
1.1
1.2
1.3
1.4
1.5
2
4
6
8
10
12
14

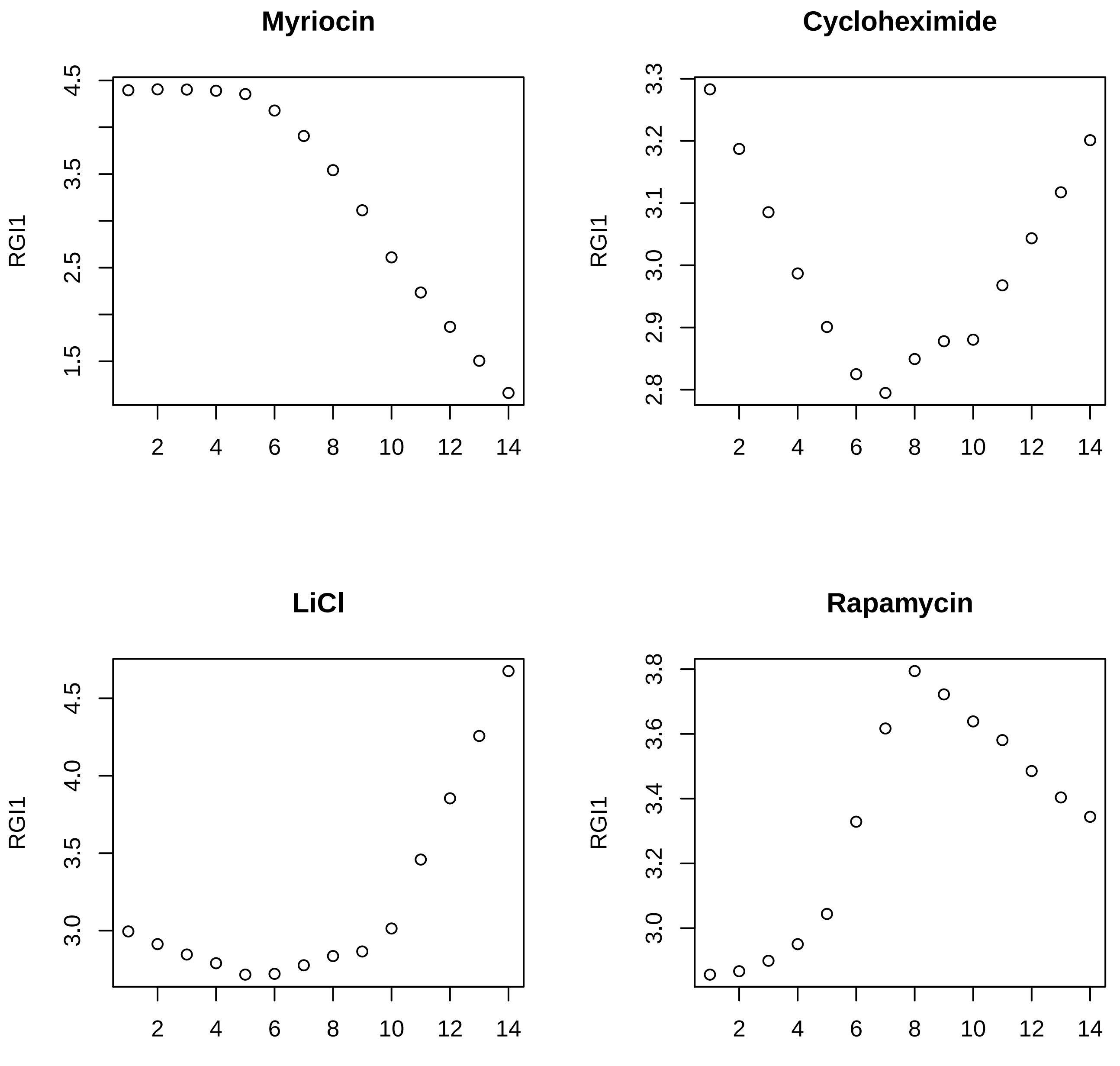
Myriocin
Cycloheximide
LiCl
Rapamycin
RGI1
RGI1
RGI1
RGI1
1.5
2.5
3.5
4.5
2.8
2.9
3.0
3.1
3.2
3.3
3.0
3.5
4.0
4.5
3.0
3.2
3.4
3.6
3.8
2
4
6
8
10
12
14

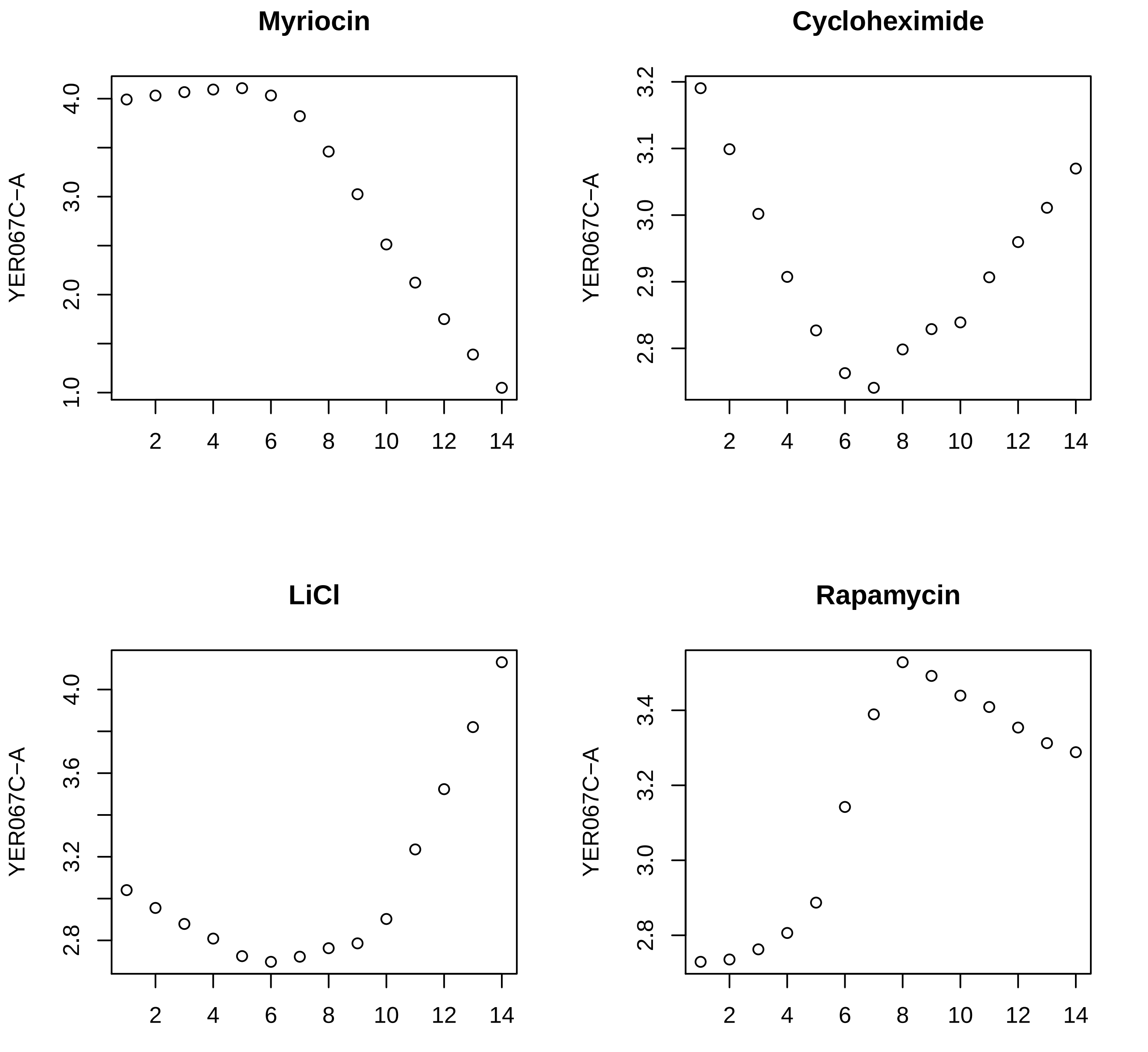
Myriocin
Cycloheximide
LiCl
Rapamycin
YER067C−A
YER067C−A
YER067C−A
YER067C−A
1.0
2.0
3.0
4.0
2.8
2.9
3.0
3.1
3.2
2.8
3.2
3.6
4.0
2.8
3.0
3.2
3.4
2
4
6
8
10
12
14

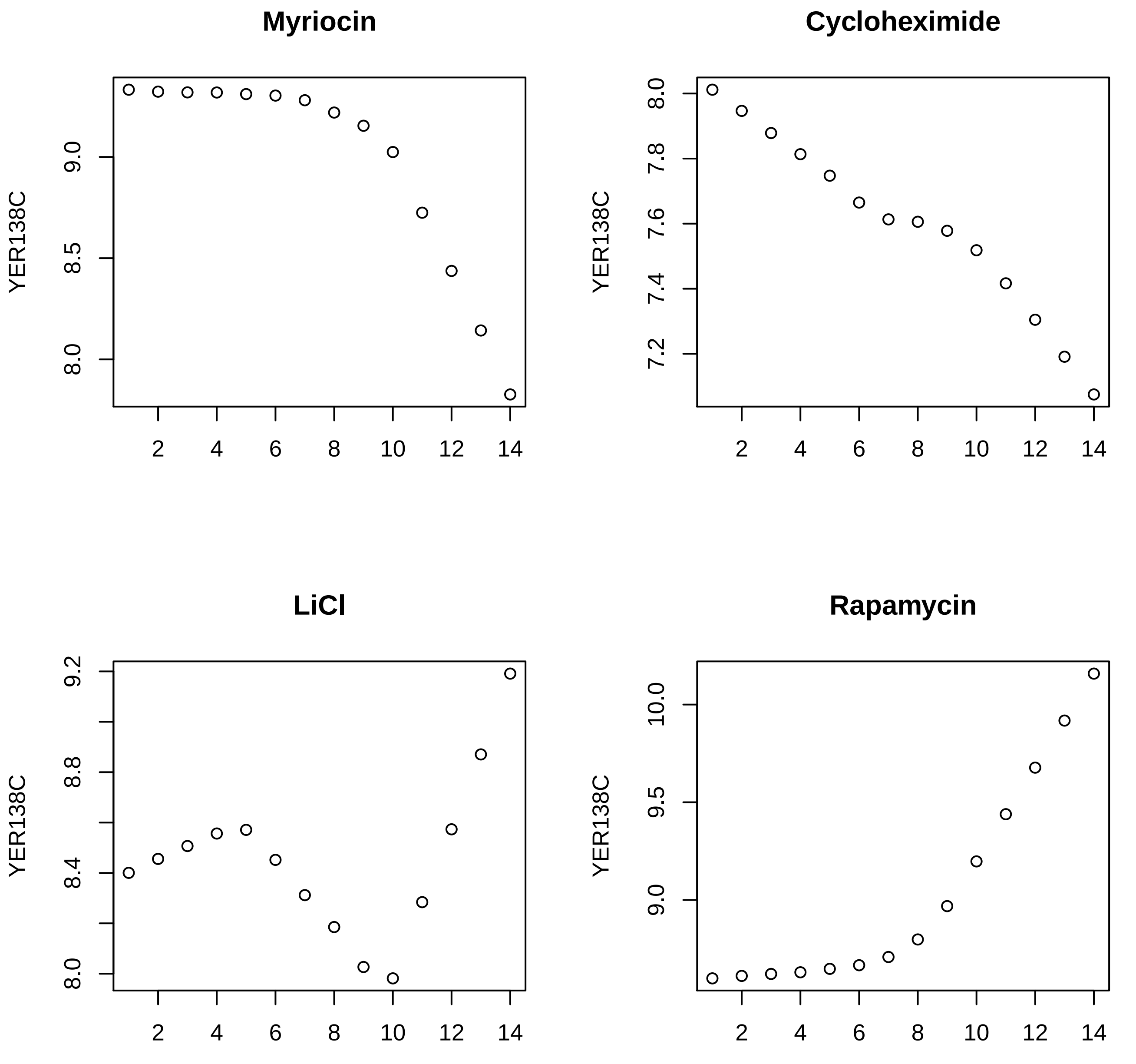
Myriocin
Cycloheximide
LiCl
Rapamycin
YER138C
9.0
8.5
8.0
7.2
7.4
7.6
7.8
9.2
8.8
8.4
10.0
9.5
2
4
6
8
10
12
14

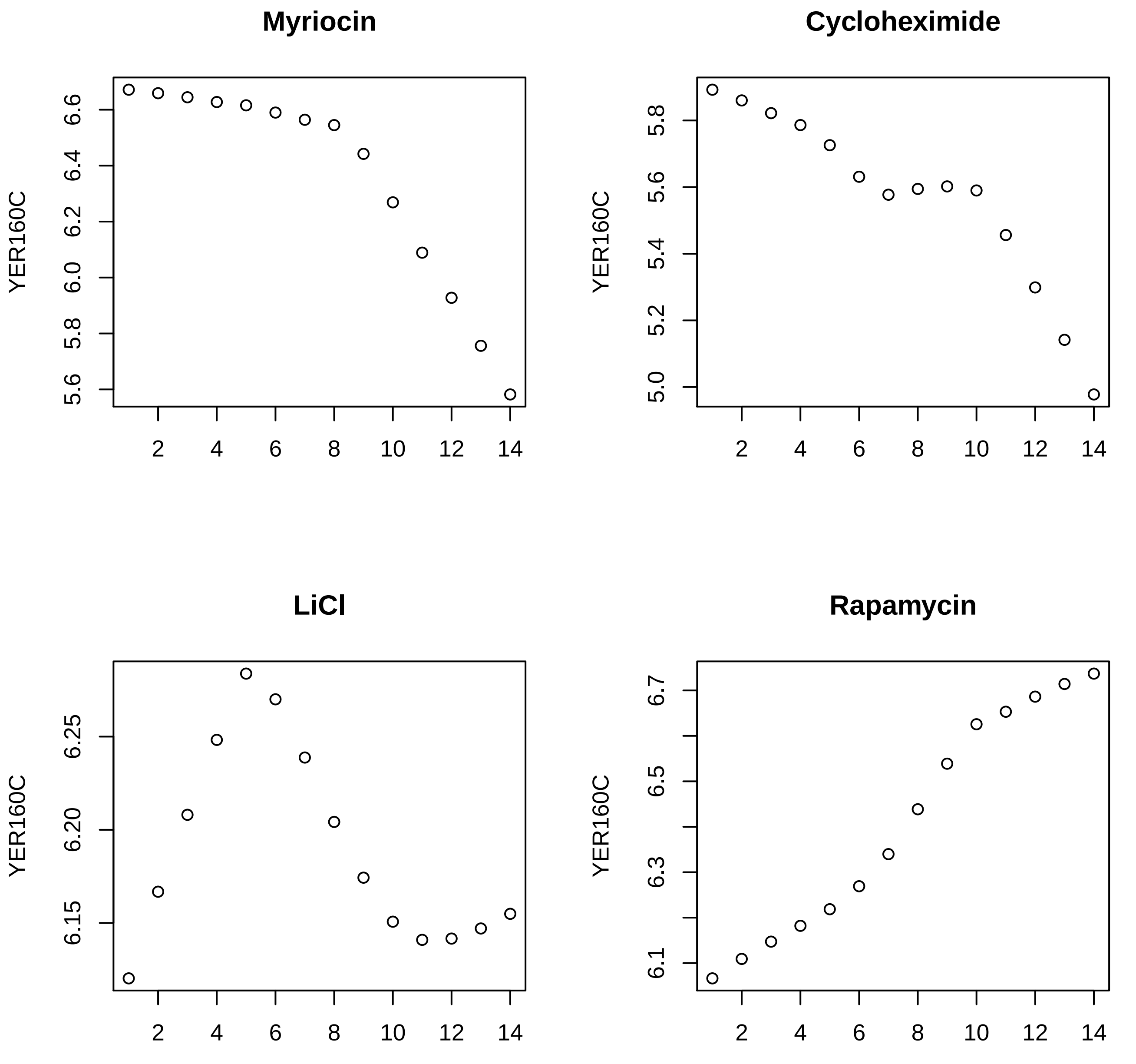

Myriocin
Cycloheximide
LiCl
Rapamycin
YER160C
5.6
5.8
6.0
6.2
6.4
6.6
5.0
5.2
5.4
5.6
5.8
6.15
6.20
6.25
6.1
6.3
6.5
6.7
2
4
6
8
10
12
14

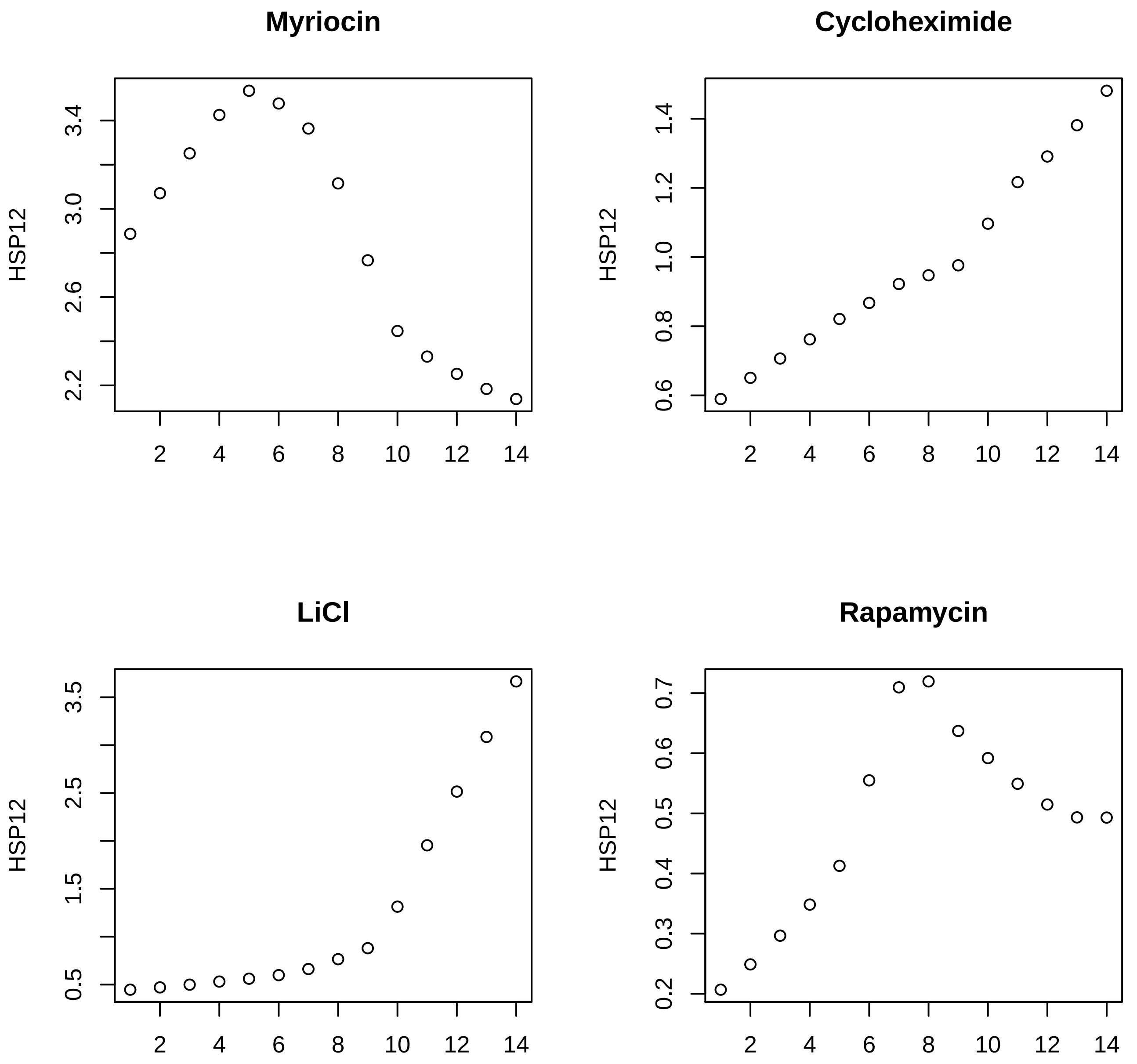

Myriocin
Cycloheximide
LiCl
Rapamycin
HSP12
HSP12
HSP12
HSP12
3.4
3.0
2.6
2.2
1.4
1.2
1.0
0.8
0.6
3.5
2.5
1.5
0.5
0.7
0.6
0.5
0.4
0.3
0.2
2
4
6
8
10
12
14

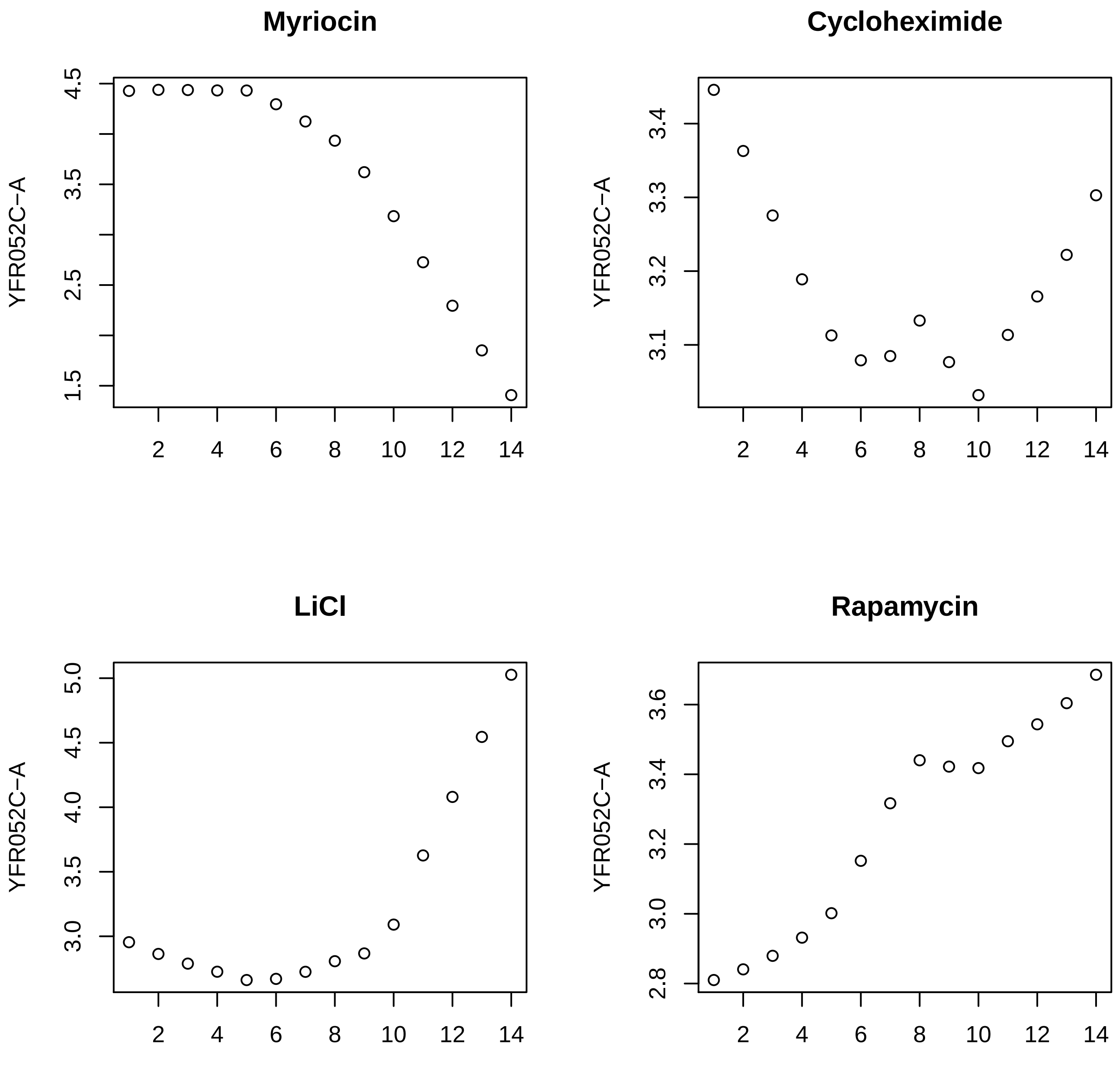

Myriocin
Cycloheximide
LiCl
Rapamycin
YFR052C−A
1.5
2.5
3.5
4.5
3.1
3.2
3.3
3.4
3.0
3.5
4.0
4.5
5.0
2.8
3.0
3.2
3.4
3.6
2
4
6
8
10
12
14

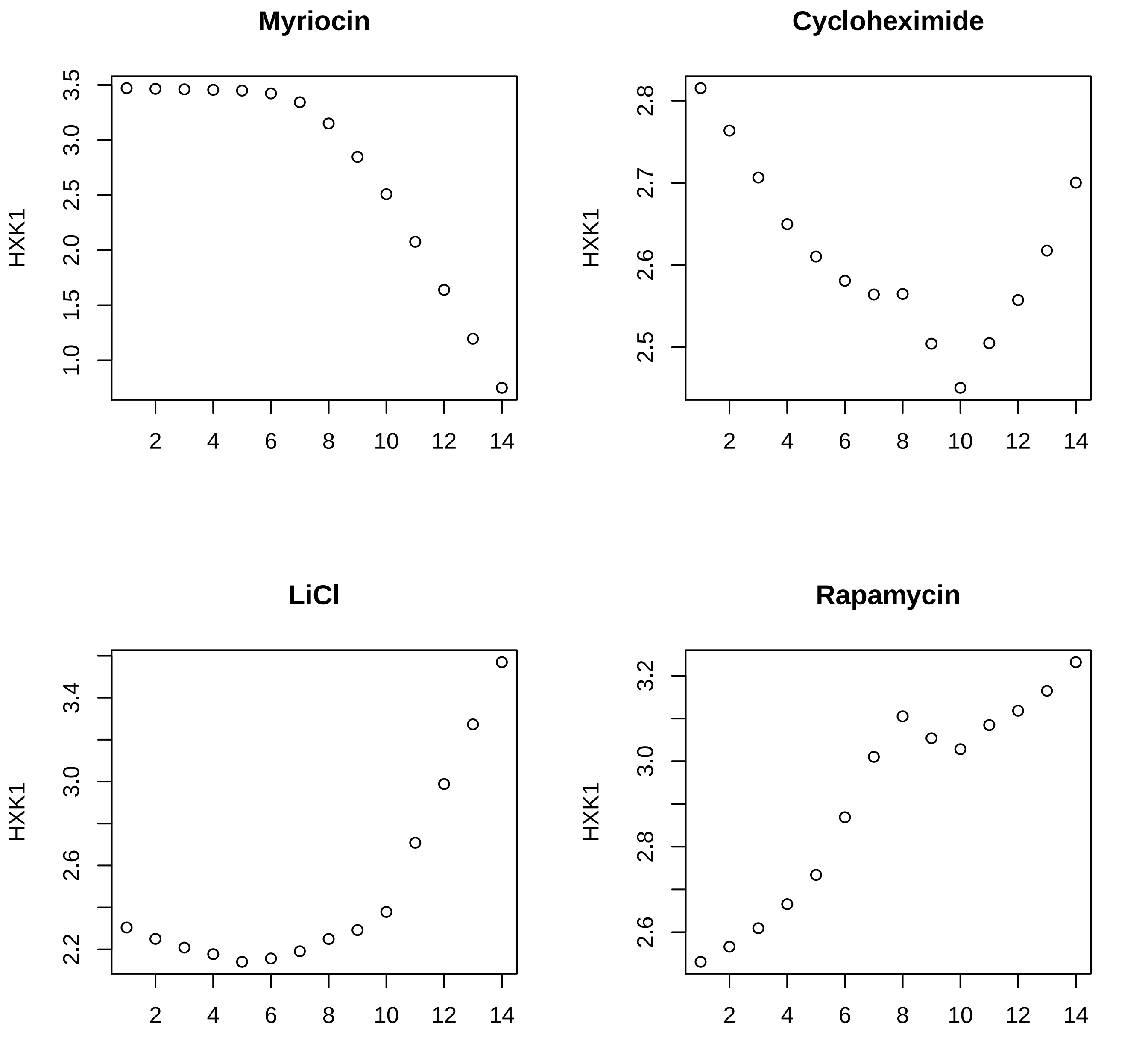
Myriocin
Cycloheximide
LiCl
Rapamycin
HXK1
3.5
3.0
2.5
2.0
1.5
1.0
2.8
2.7
2.6
2.5
3.4
3.0
2.6
2.2
3.2
3.0
2.8
2.6
2
4
6
8
10
12
14

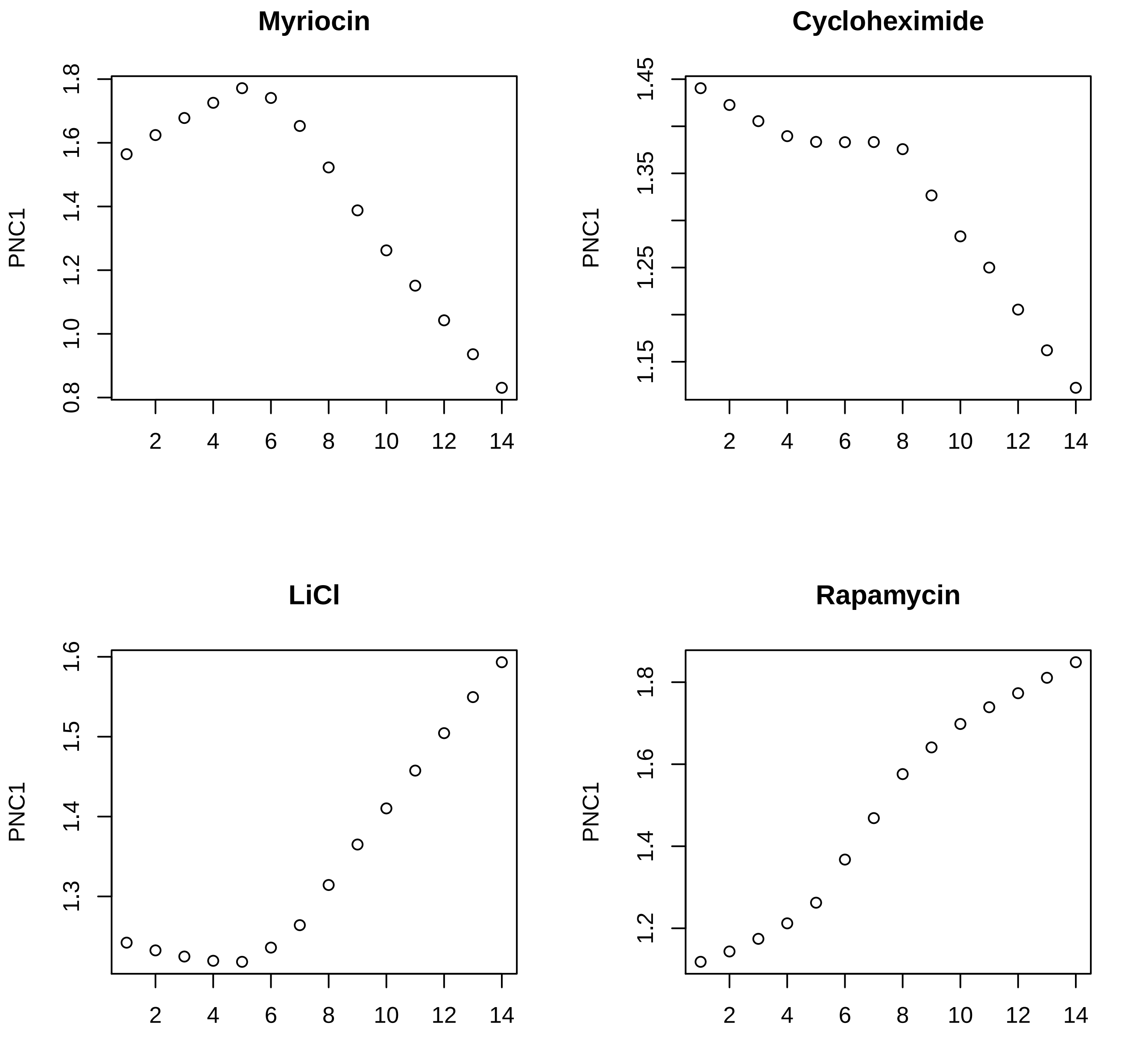
Myriocin
Cycloheximide
LiCl
Rapamycin
PNC1
PNC1
PNC1
PNC1
0.8
1.0
1.2
1.4
1.6
1.8
1.15
1.25
1.35
1.45
1.3
1.4
1.5
1.6
1.2
1.4
1.6
1.8
2
4
6
8
10
12
14

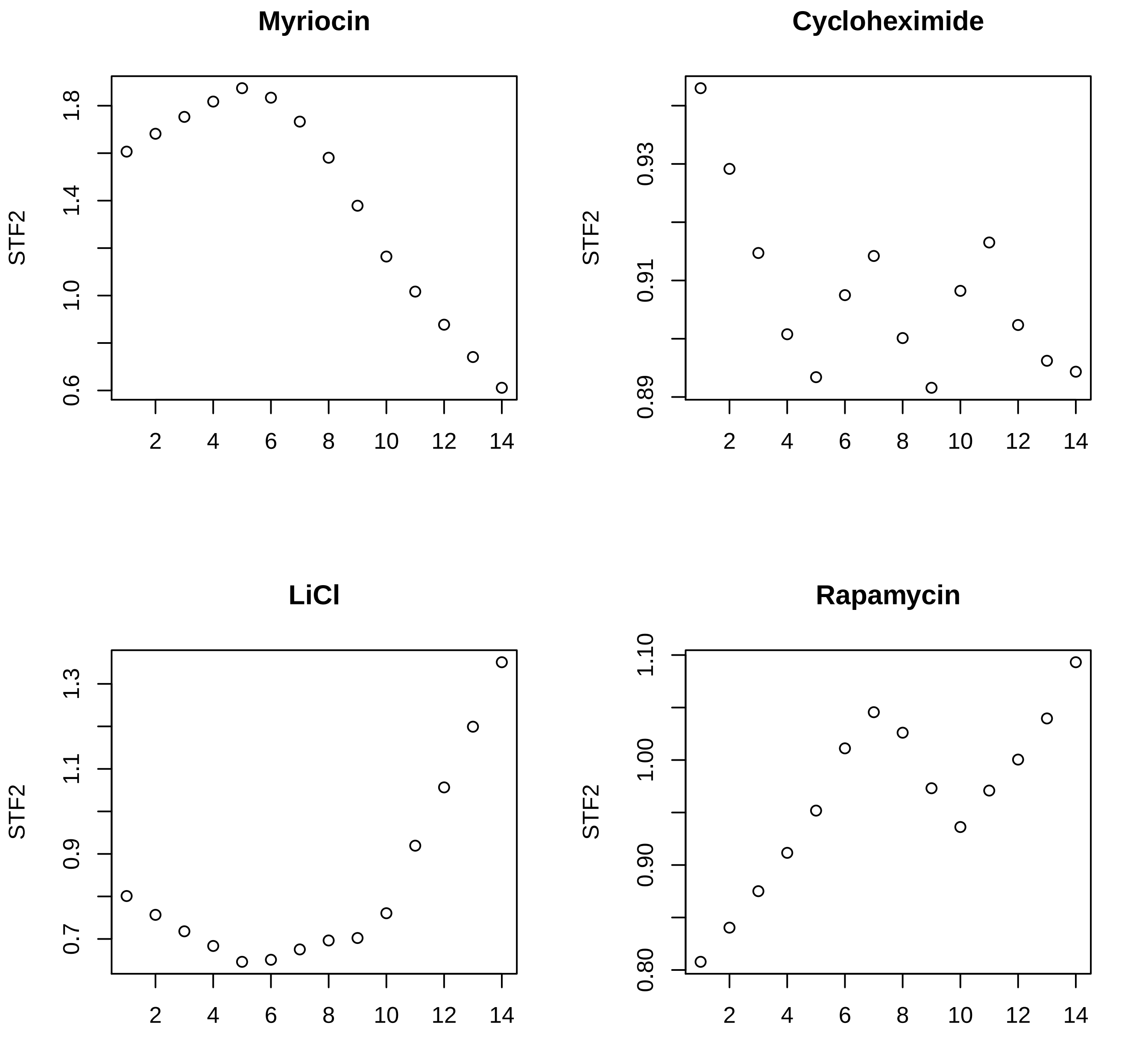
Myriocin
STF2
0.6
1.0
1.4
1.8
2
4
6
8
10
12
14
Cycloheximide
STF2
0.89
0.91
0.93
2
4
6
8
10
12
14
LiCl
STF2
0.7
0.9
1.1
1.3
2
4
6
8
10
12
14
Rapamycin
STF2
0.80
0.90
1.00
1.10
2
4
6
8
10
12
14

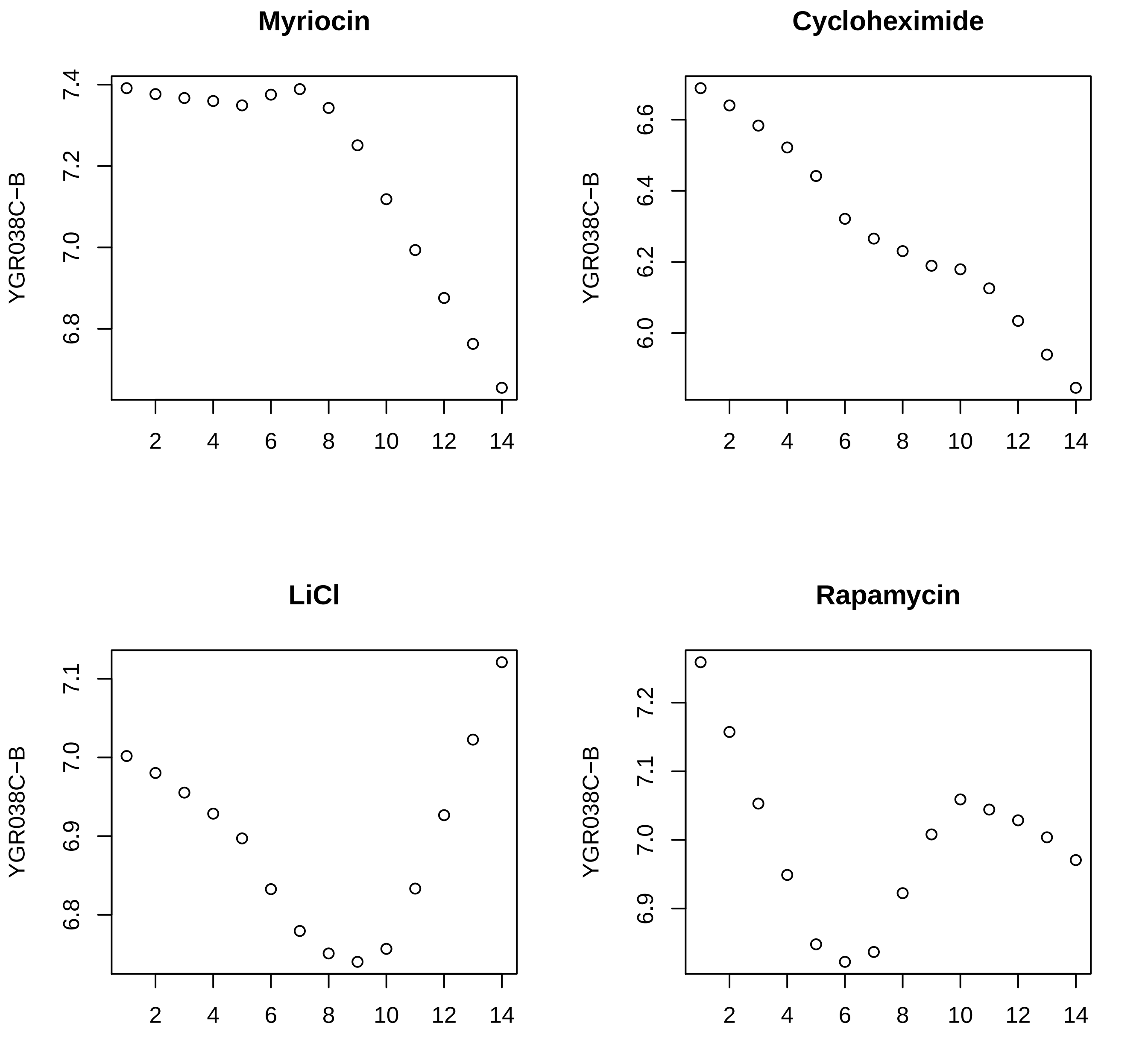
Myriocin
Cycloheximide
LiCl
Rapamycin
YGR038C−B
YGR038C−B
YGR038C−B
YGR038C−B
7.4
7.2
7.0
6.8
6.6
6.4
6.2
6.0
7.1
7.0
6.9
6.8
7.2
7.1
7.0
6.9
2
4
6
8
10
12
14

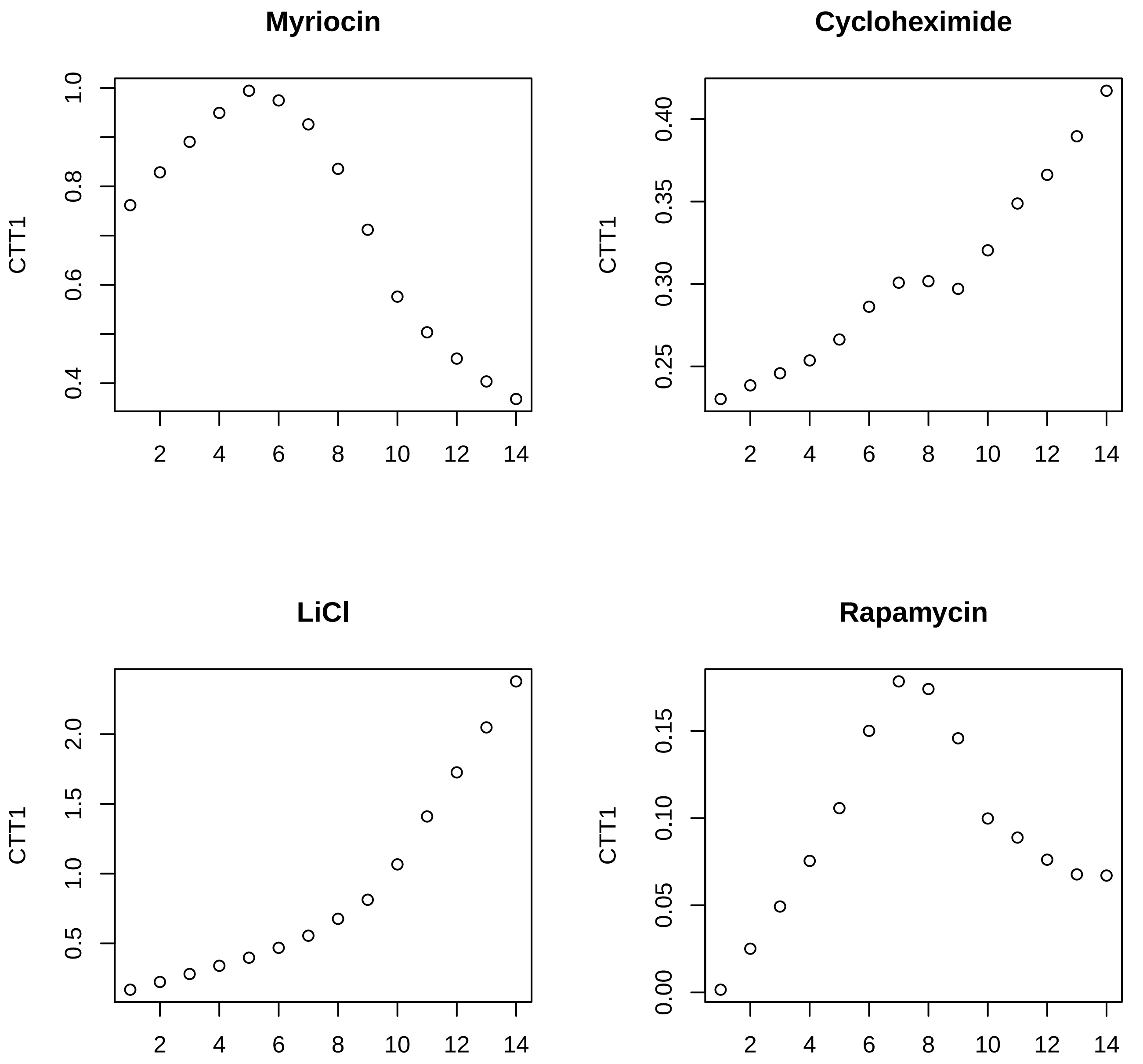

Myriocin
Cycloheximide
LiCl
Rapamycin
CTT1

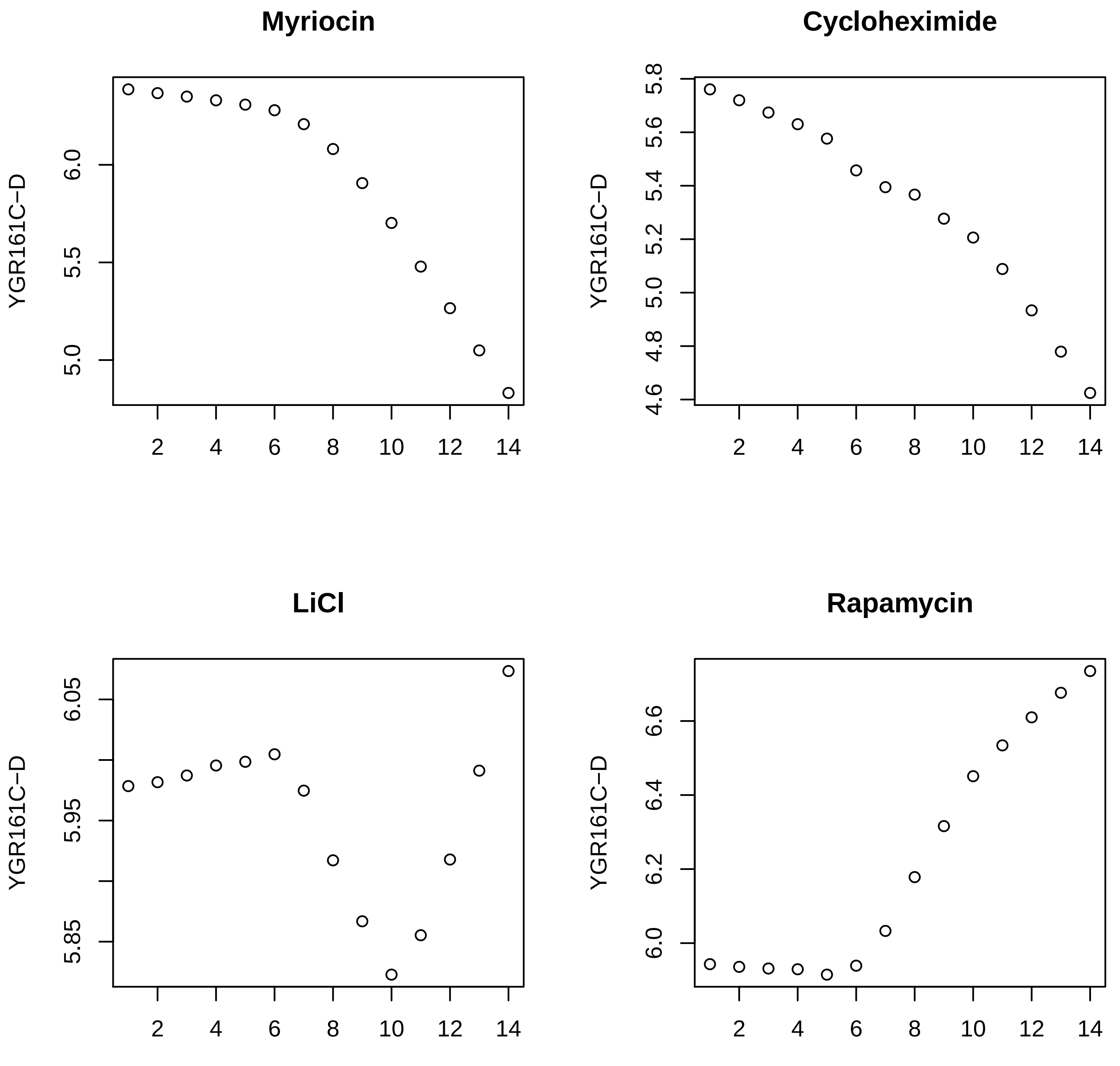
Myriocin
Cycloheximide
LiCl
Rapamycin
YGR161C−D
YGR161C−D
YGR161C−D
YGR161C−D
5.0
5.5
6.0
4.6
4.8
5.0
5.2
5.4
5.6
5.8
5.85
5.95
6.05
6.0
6.2
6.4
6.6
2
4
6
8
10
12
14

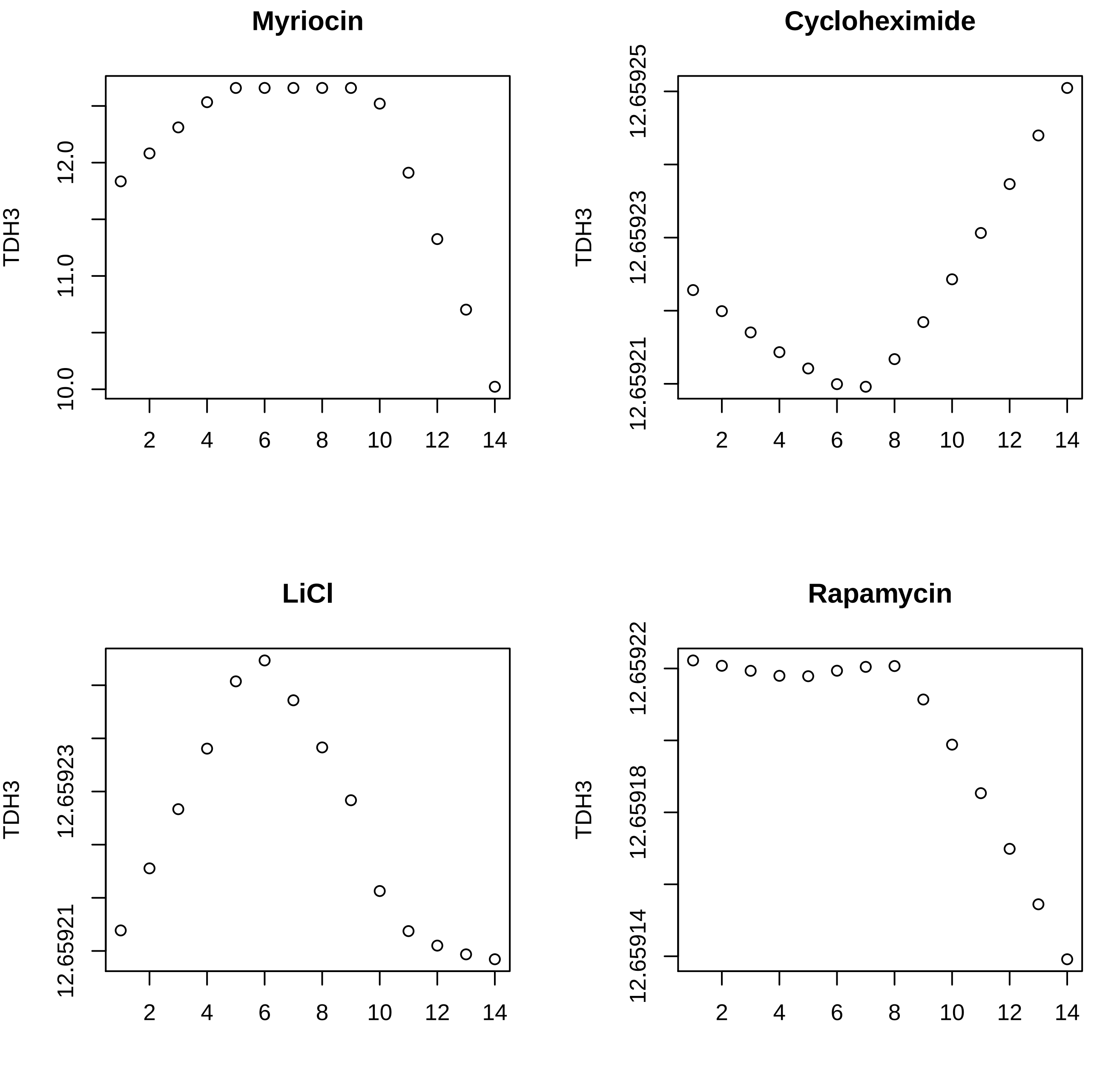

Myriocin
TDH3
10.0
11.0
12.0
2
4
6
8
10
12
14
Cycloheximide
TDH3
12.65921
12.65923
12.65925
LiCl
12.65921
12.65923
Rapamycin
12.65914
12.65918
12.65922

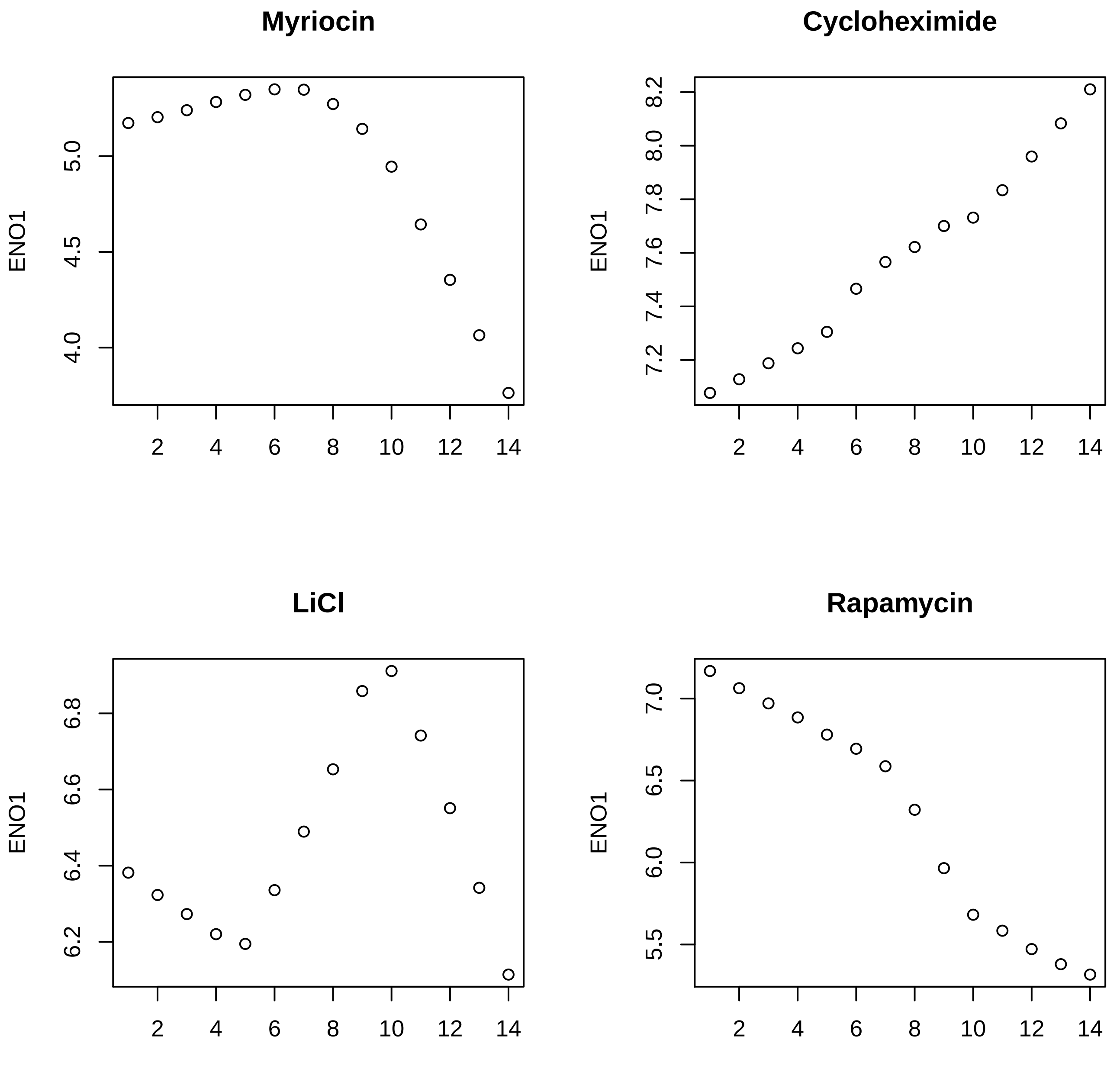
Myriocin
ENO1
5.0
4.5
4.0
2
4
6
8
10
12
14
Cycloheximide
ENO1
8.2
8.0
7.8
7.6
7.4
7.2
2
4
6
8
10
12
14
LiCl
ENO1
6.8
6.6
6.4
6.2
2
4
6
8
10
12
14
Rapamycin
ENO1
7.0
6.5
6.0
5.5
2
4
6
8
10
12
14

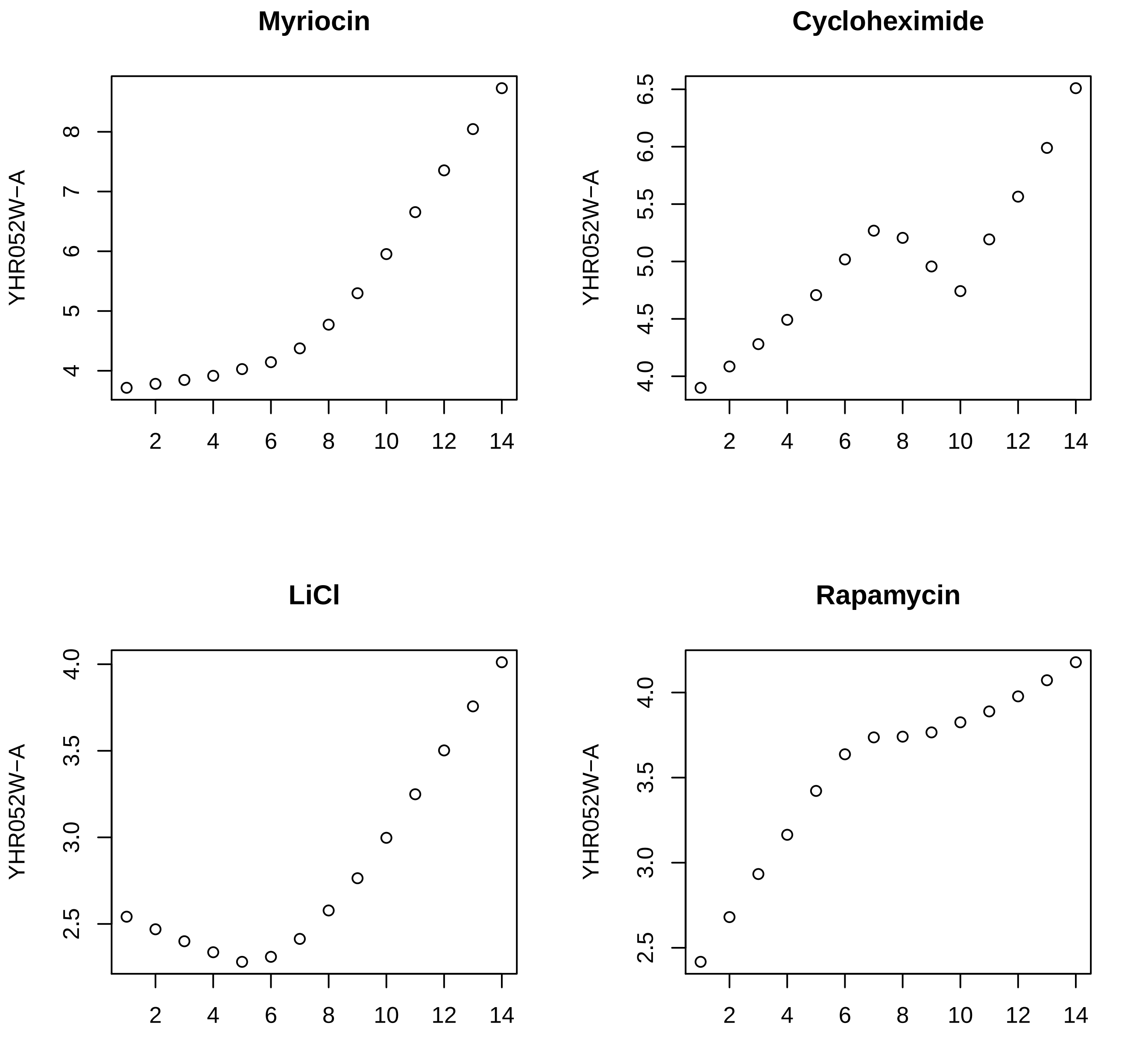

Myriocin
Cycloheximide
LiCl
Rapamycin
YHR052W−A

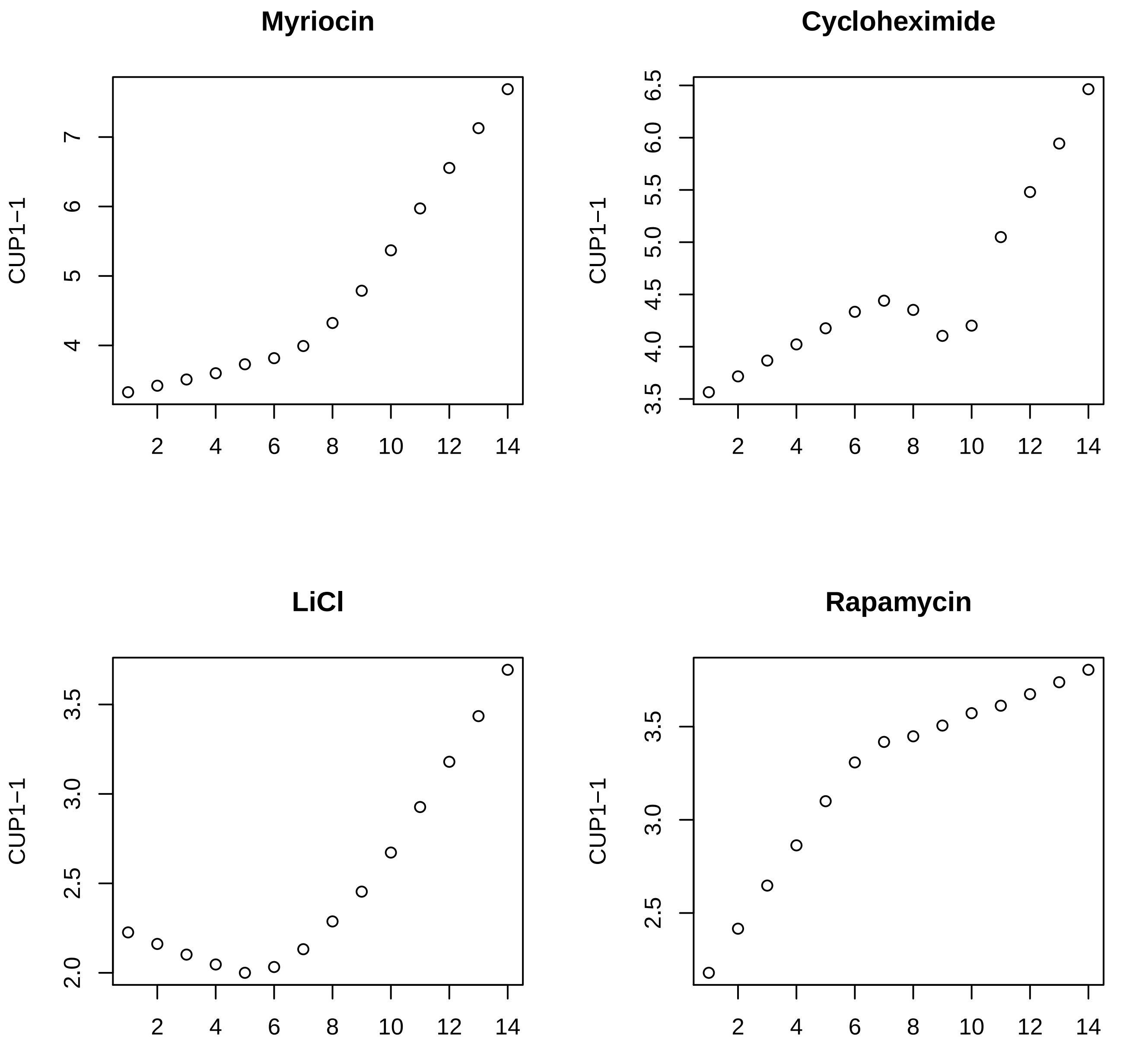
Myriocin
Cycloheximide
LiCl
Rapamycin
CUP1−1
2
4
6
8
10
12
14
4
5
6
7
3.5
4.0
4.5
5.0
5.5
6.0
6.5
2.0
2.5
3.0
3.5

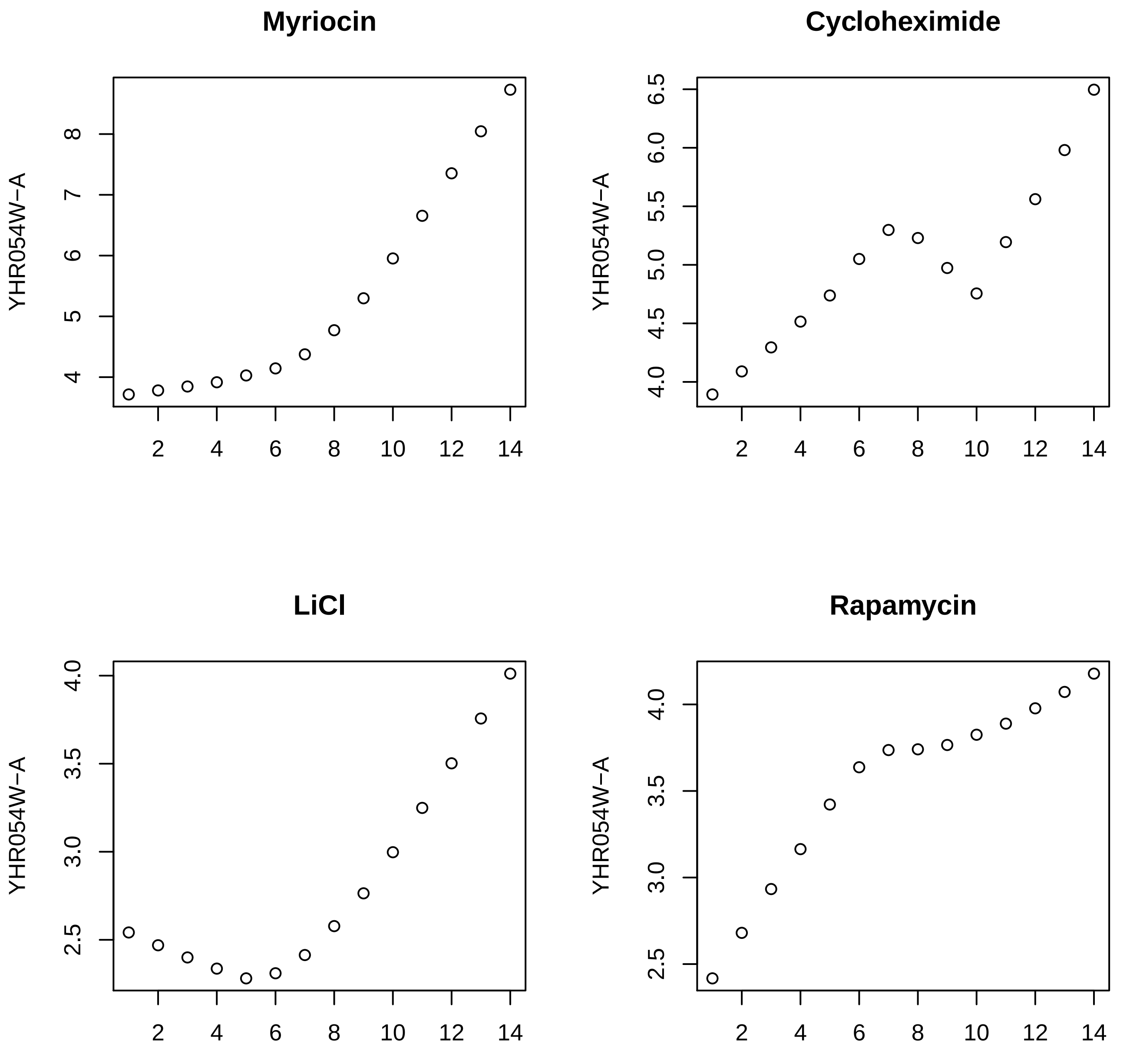

Myriocin
Cycloheximide
LiCl
Rapamycin
YHR054W−A
YHR054W−A
YHR054W−A
YHR054W−A
2
4
6
8
10
12
14

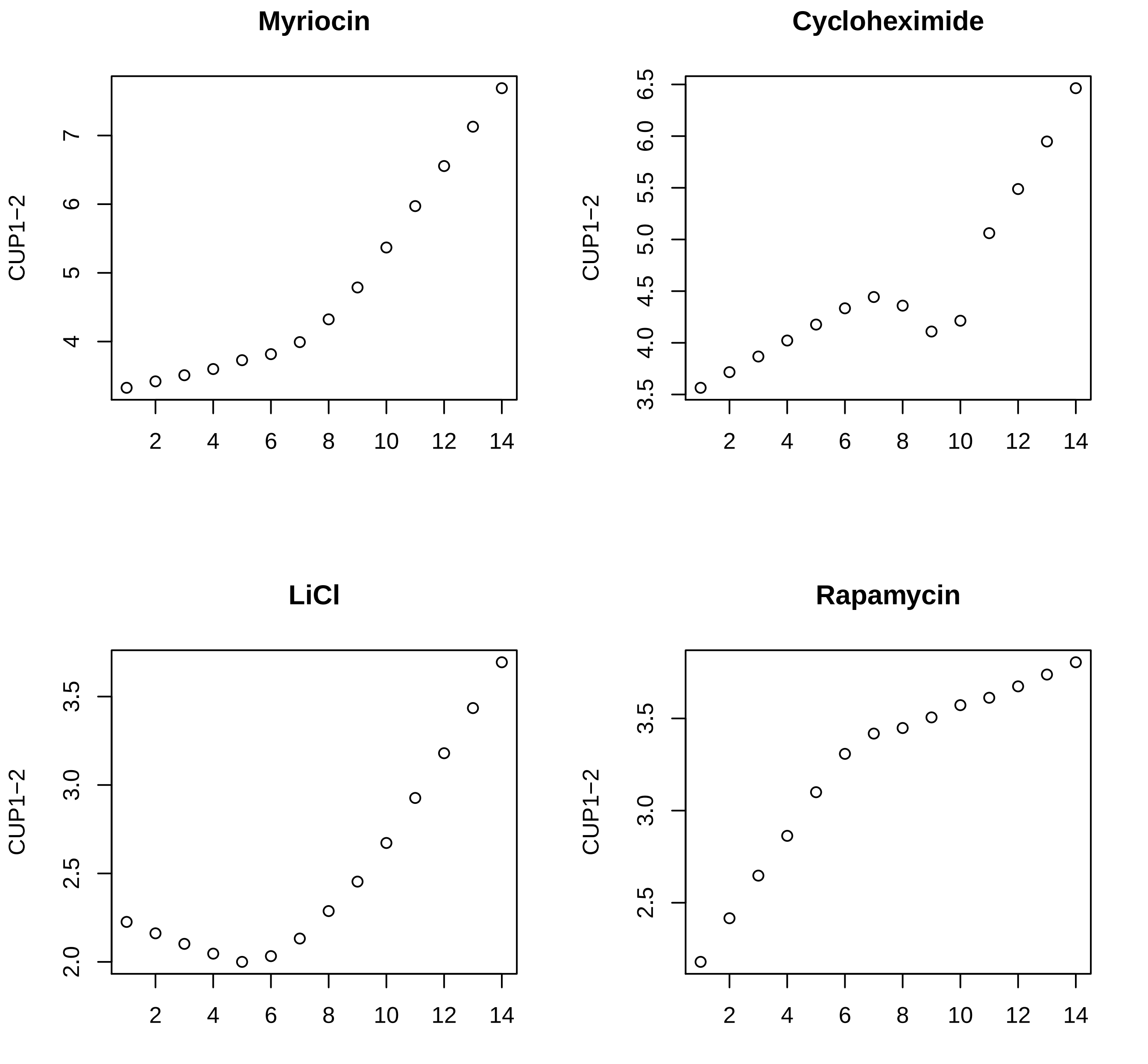
Myriocin
Cycloheximide
LiCl
Rapamycin
CUP1−2
CUP1−2
CUP1−2
CUP1−2
2
4
6
8
10
12
14

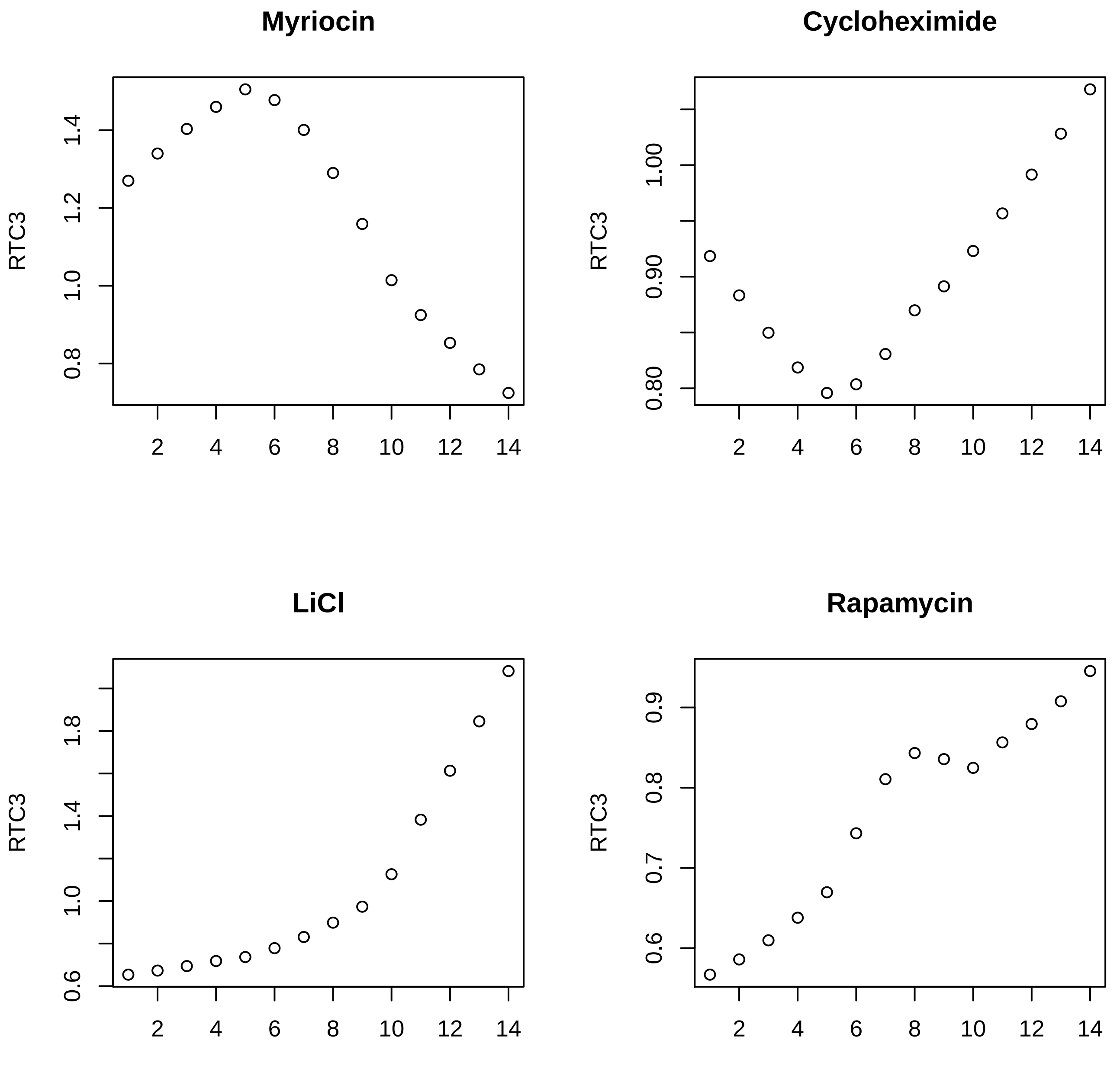

Myriocin
Cycloheximide
LiCl
Rapamycin
RTC3
RTC3
RTC3
RTC3
0.8
1.0
1.2
1.4
0.80
0.90
1.00
0.6
1.0
1.4
1.8
0.6
0.7
0.8
0.9
2
4
6
8
10
12
14

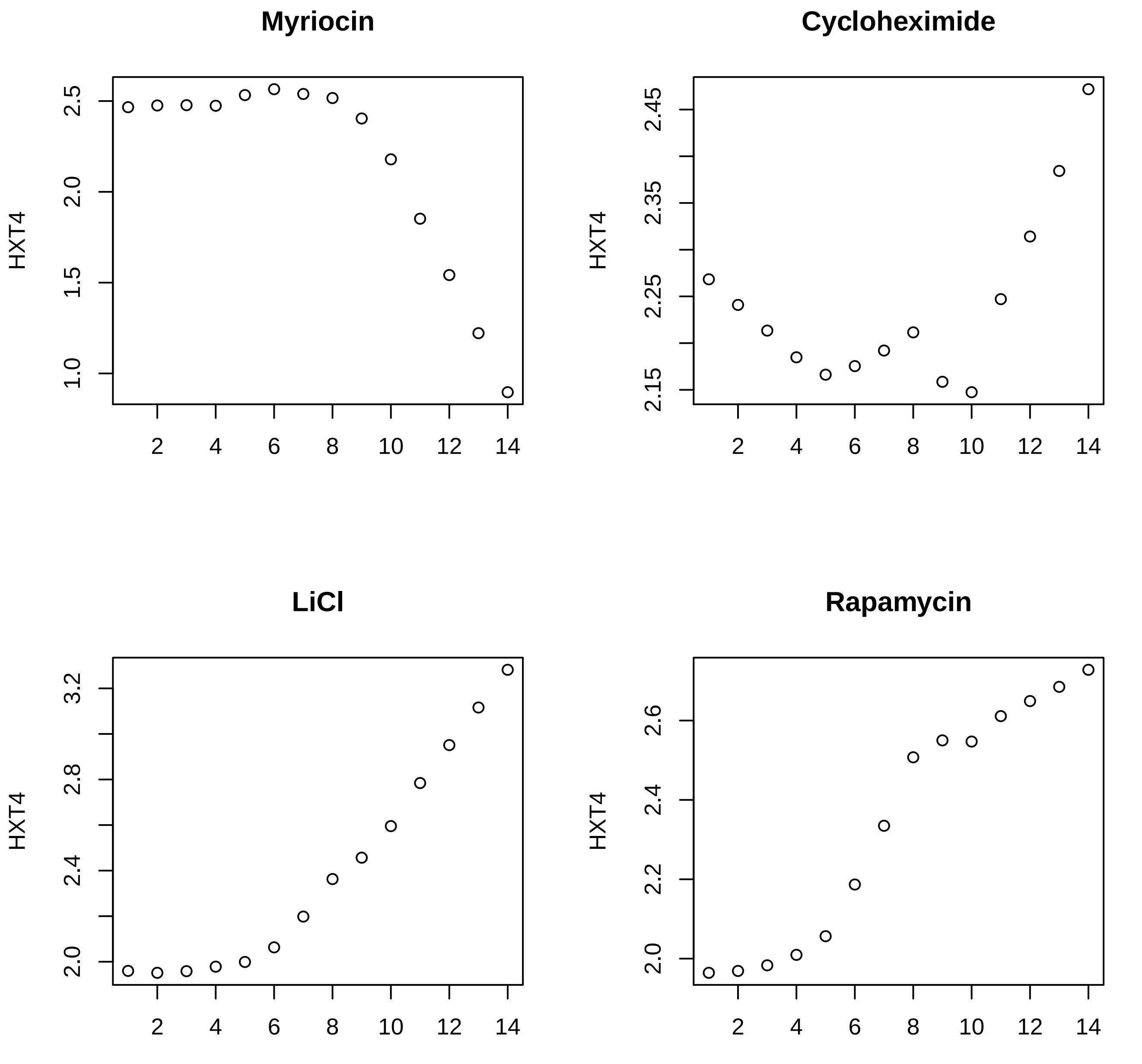

Myriocin
Cycloheximide
LiCl
Rapamycin
HXT4

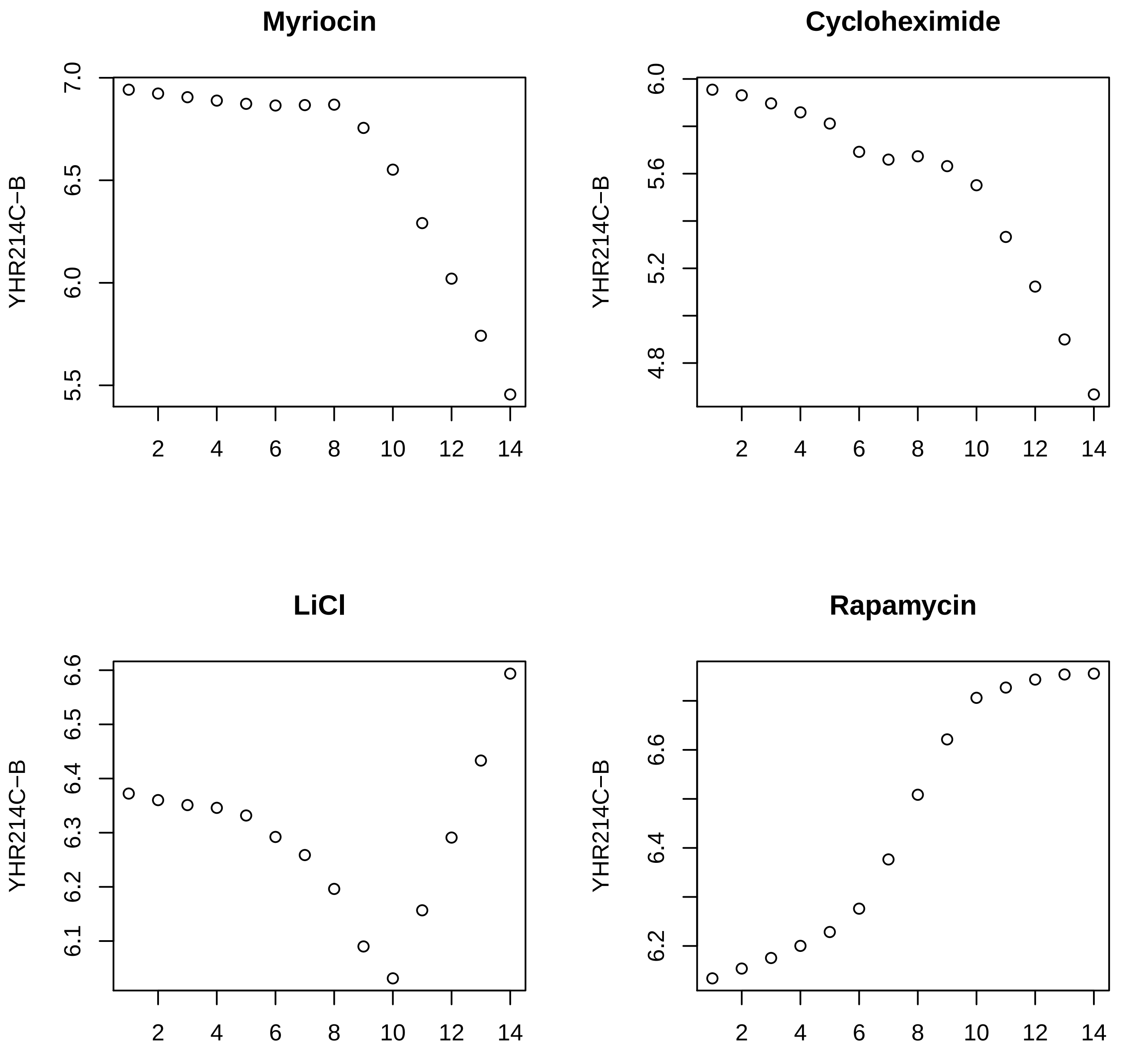

Myriocin
Cycloheximide
LiCl
Rapamycin
YHR214C−B

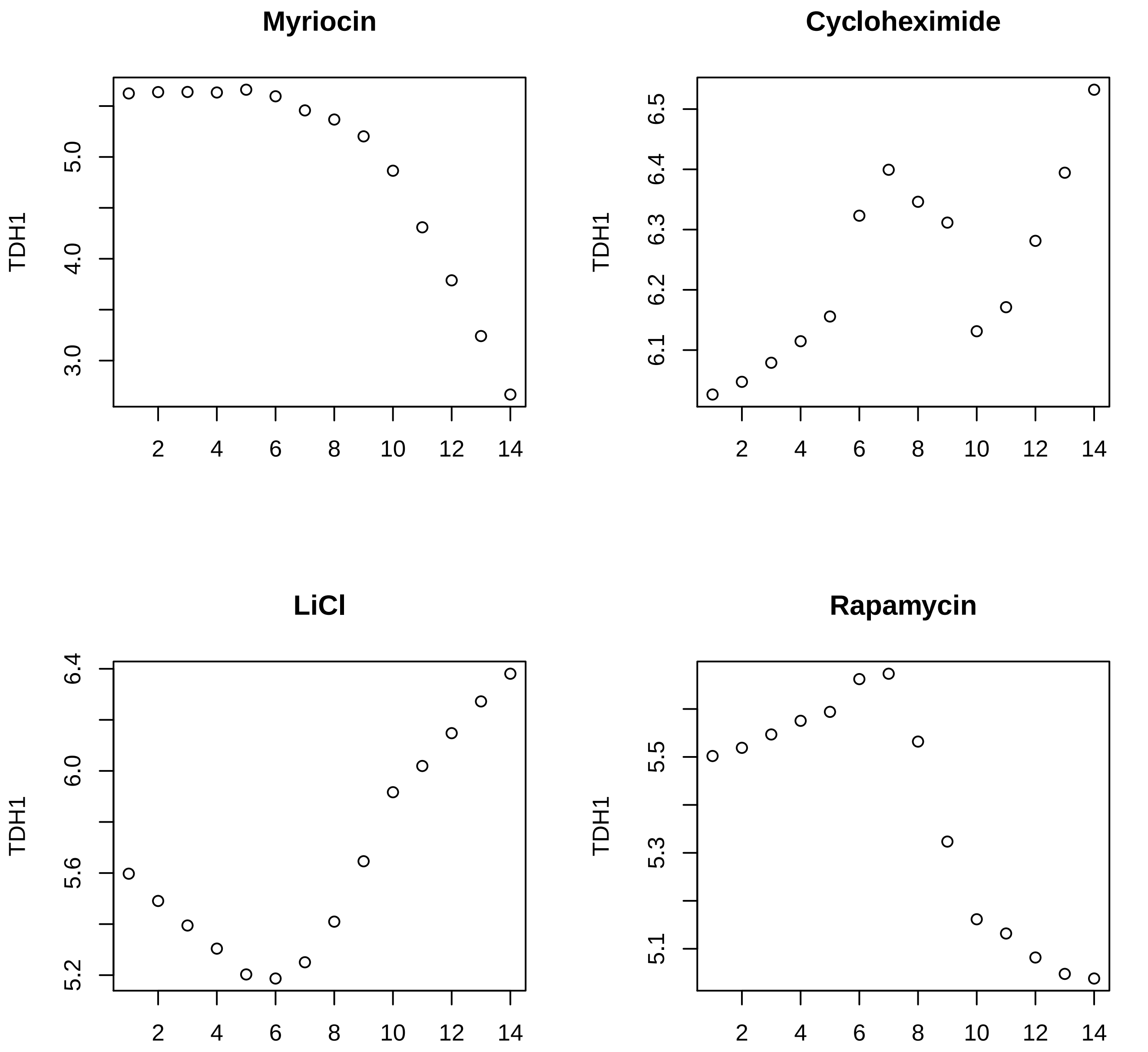
Myriocin
TDH1
3.0
4.0
5.0
2
4
6
8
10
12
14
Cycloheximide
TDH1
6.1
6.2
6.3
6.4
6.5
2
4
6
8
10
12
14
LiCl
TDH1
5.2
5.6
6.0
6.4
2
4
6
8
10
12
14
Rapamycin
TDH1
5.1
5.3
5.5
2
4
6
8
10
12
14

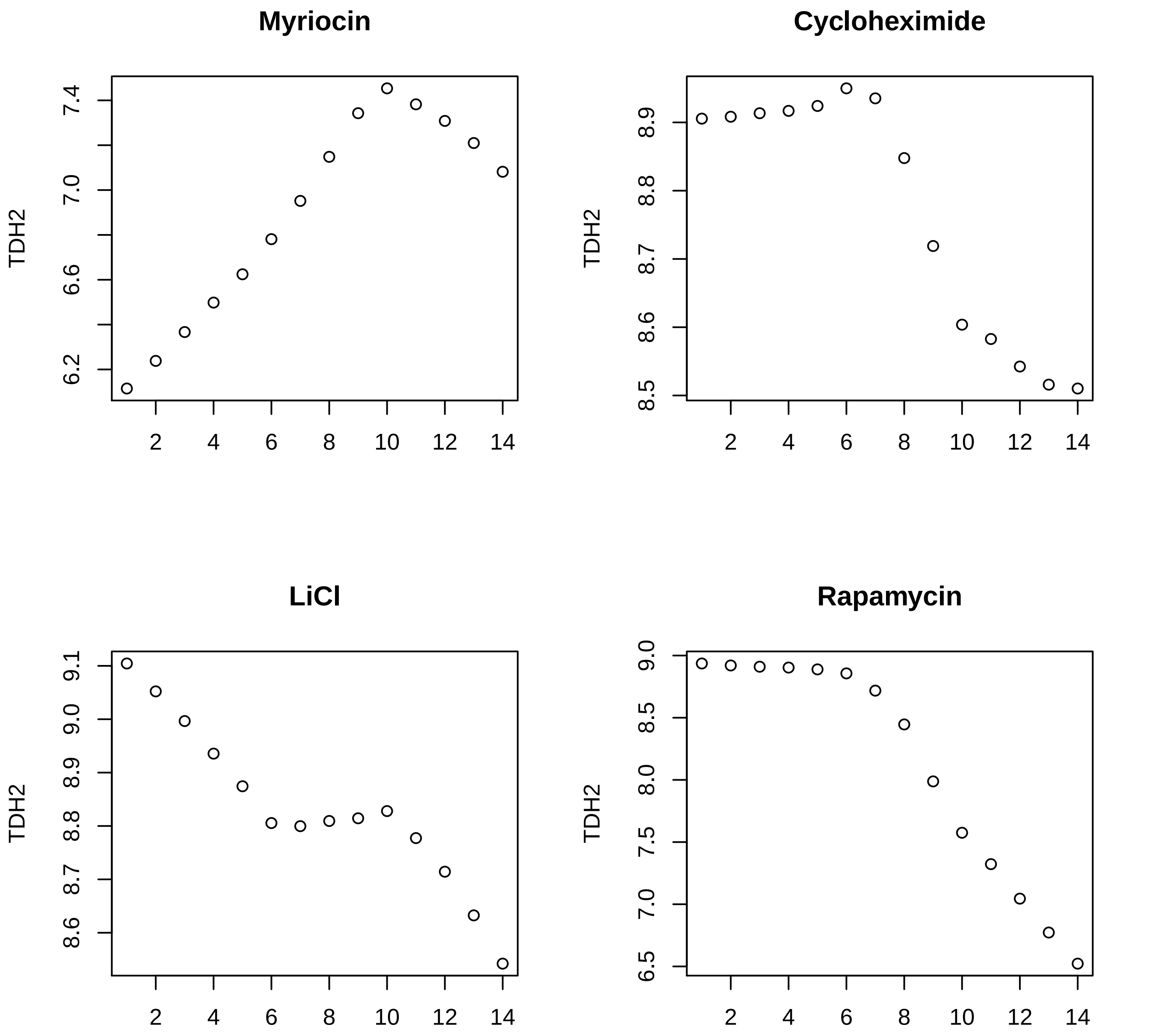
Myriocin
Cycloheximide
LiCl
Rapamycin
TDH2
2
4
6
8
10
12
14
6.2
6.6
7.0
7.4
8.5
8.6
8.7
8.8
8.9
9.0
9.1
6.5
7.0
7.5
8.0
8.5
9.0

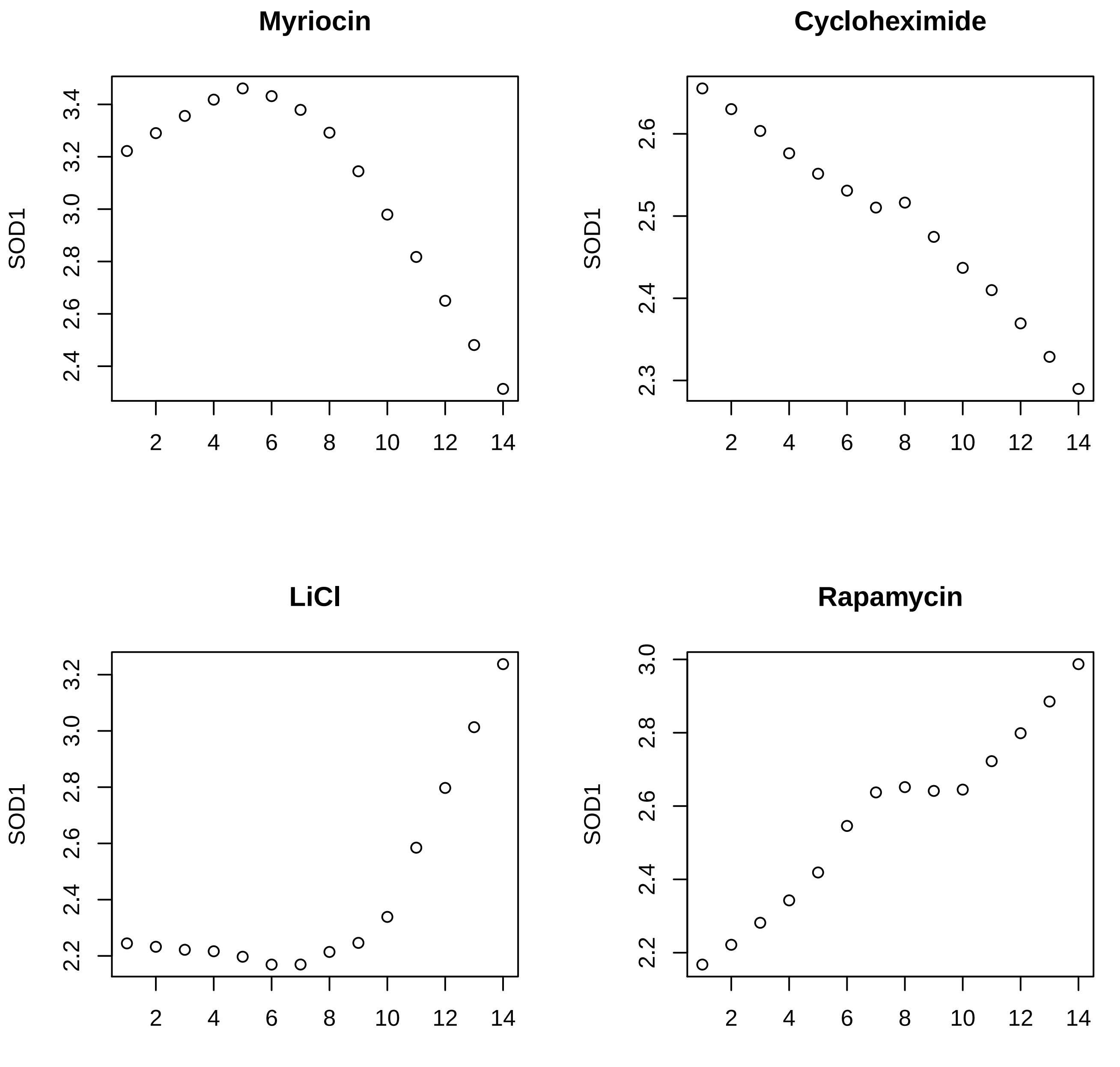
Myriocin
SOD1
2.4
2.6
2.8
3.0
3.2
3.4
2
4
6
8
10
12
14
Cycloheximide
SOD1
2.3
2.4
2.5
2.6
2
4
6
8
10
12
14
LiCl
SOD1
2.2
2.4
2.6
2.8
3.0
3.2
2
4
6
8
10
12
14
Rapamycin
SOD1
2.2
2.4
2.6
2.8
3.0
2
4
6
8
10
12
14

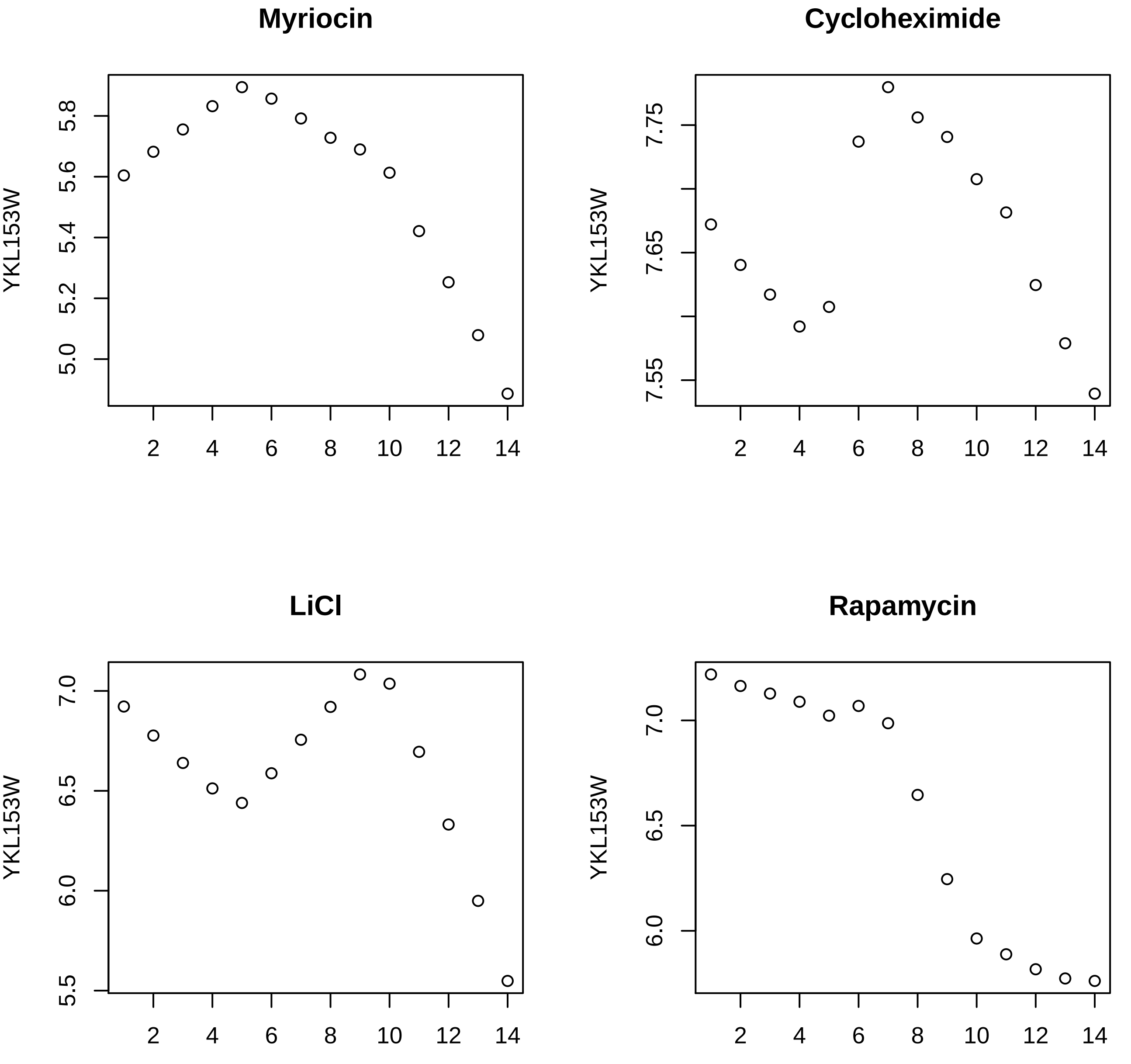
Myriocin
Cycloheximide
LiCl
Rapamycin
YKL153W
YKL153W
YKL153W
YKL153W
5.0
5.2
5.4
5.6
5.8
7.55
7.65
7.75
5.5
6.0
6.5
7.0
6.0
6.5
7.0
2
4
6
8
10
12
14

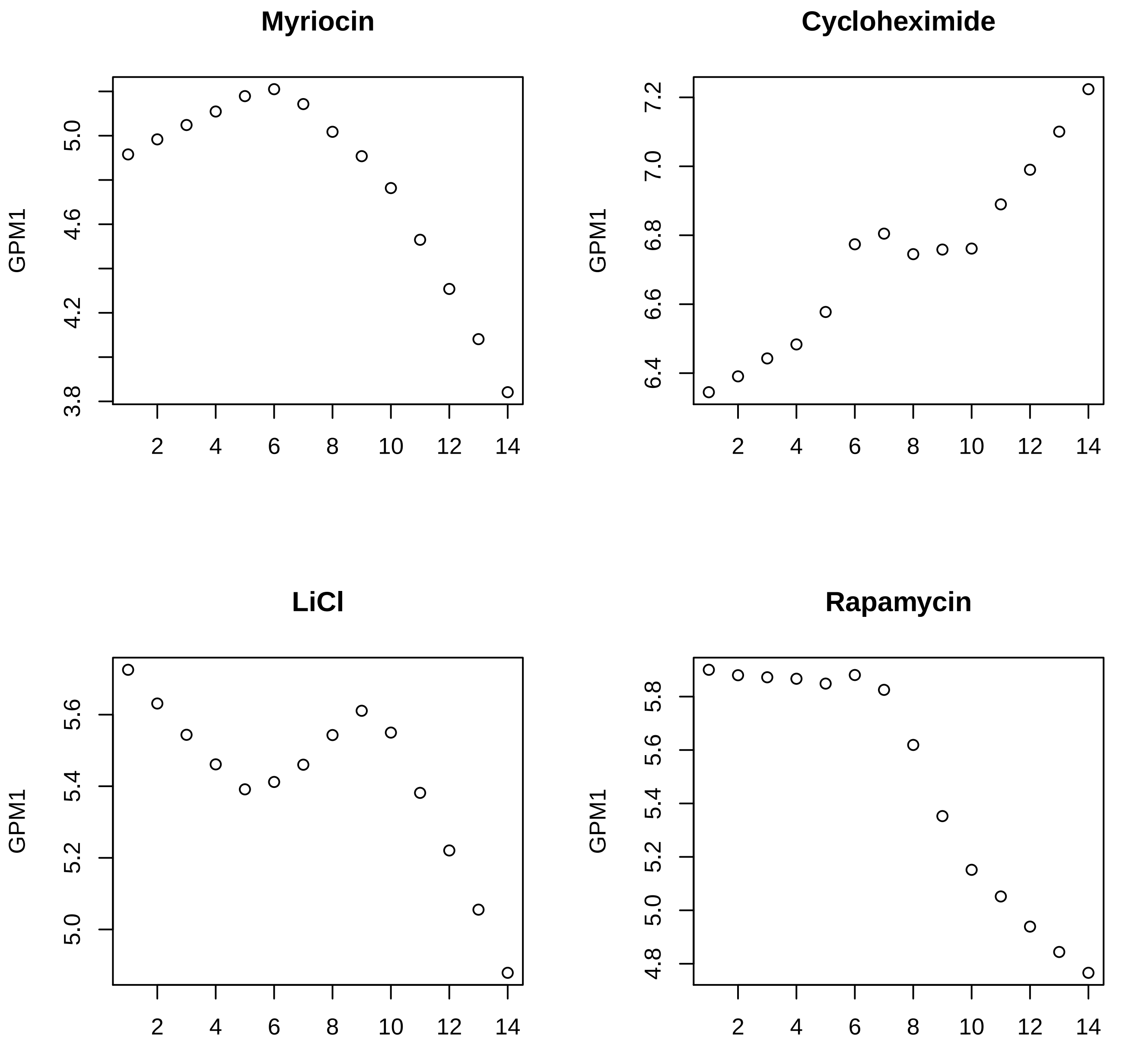

Myriocin
Cycloheximide
LiCl
Rapamycin
GPM1

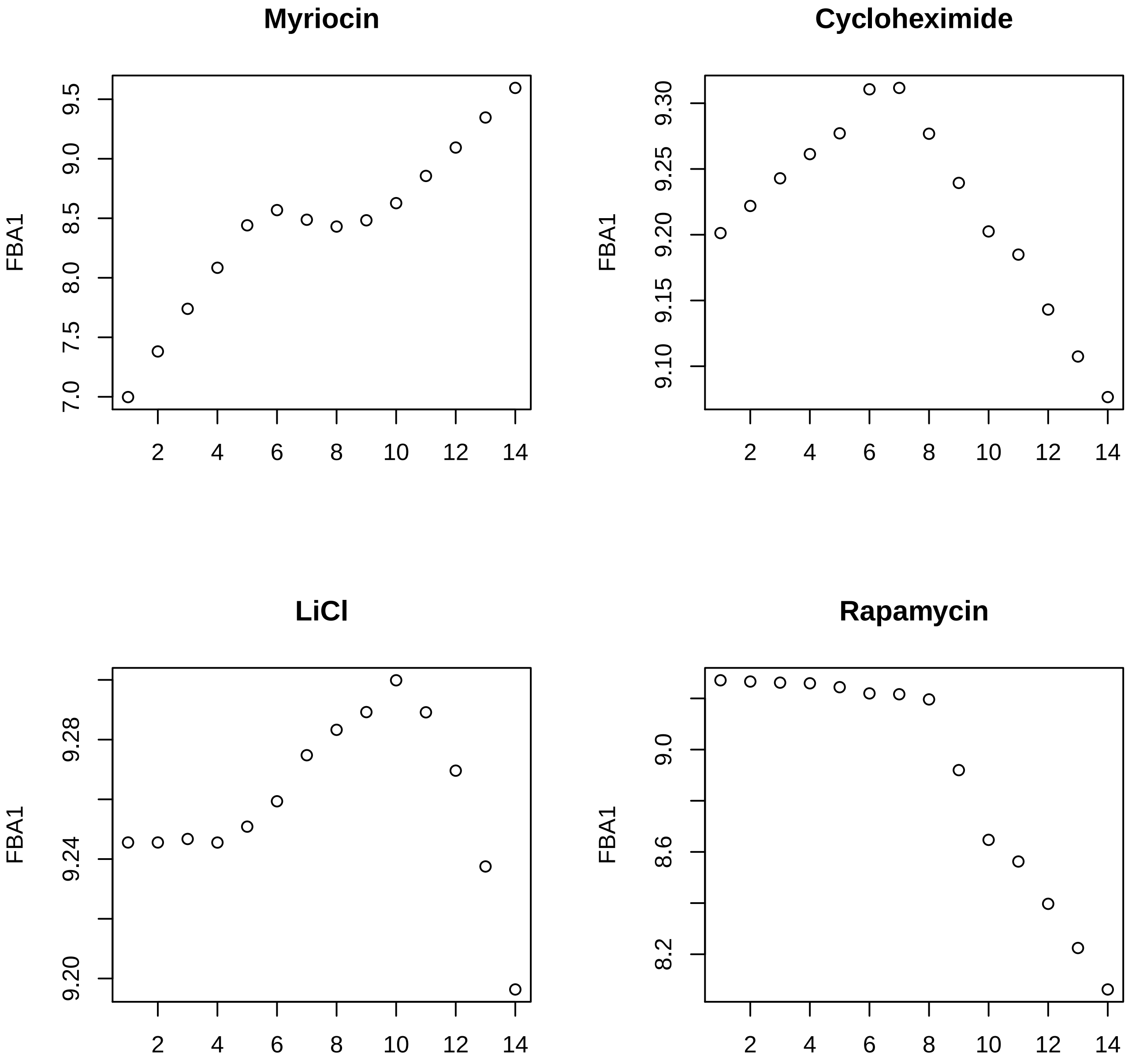
Myriocin
Cycloheximide
LiCl
Rapamycin
FBA1
FBA1
FBA1
FBA1
7.0
7.5
8.0
8.5
9.0
9.5
9.10
9.15
9.20
9.25
9.30
9.20
9.24
9.28
8.2
8.6
9.0
2
4
6
8
10
12
14

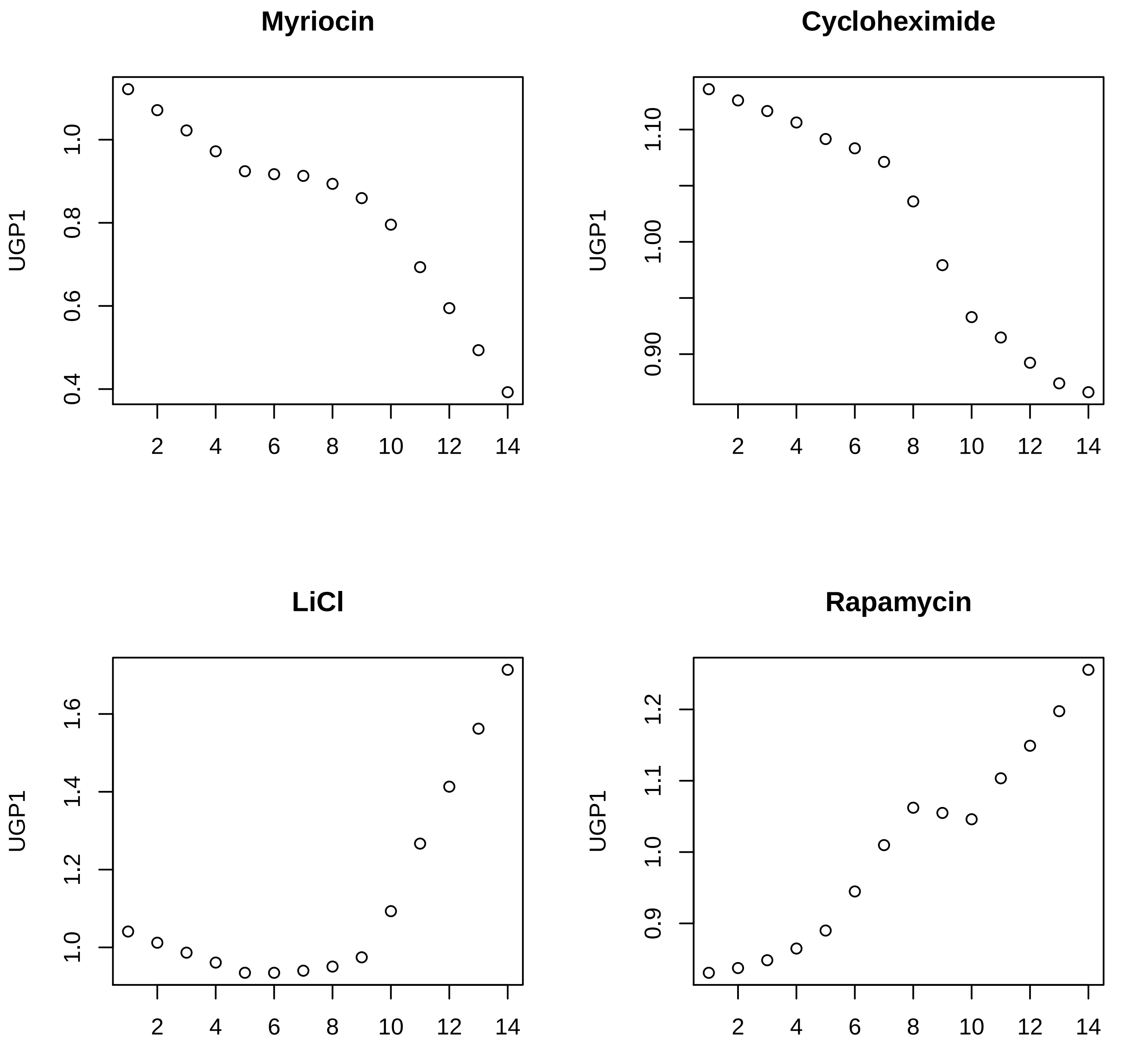

Myriocin
Cycloheximide
LiCl
Rapamycin
UGP1
1.0
0.8
0.6
0.4
1.10
1.00
0.90
1.6
1.4
1.2
1.2
1.1
0.9
2
4
6
8
10
12
14

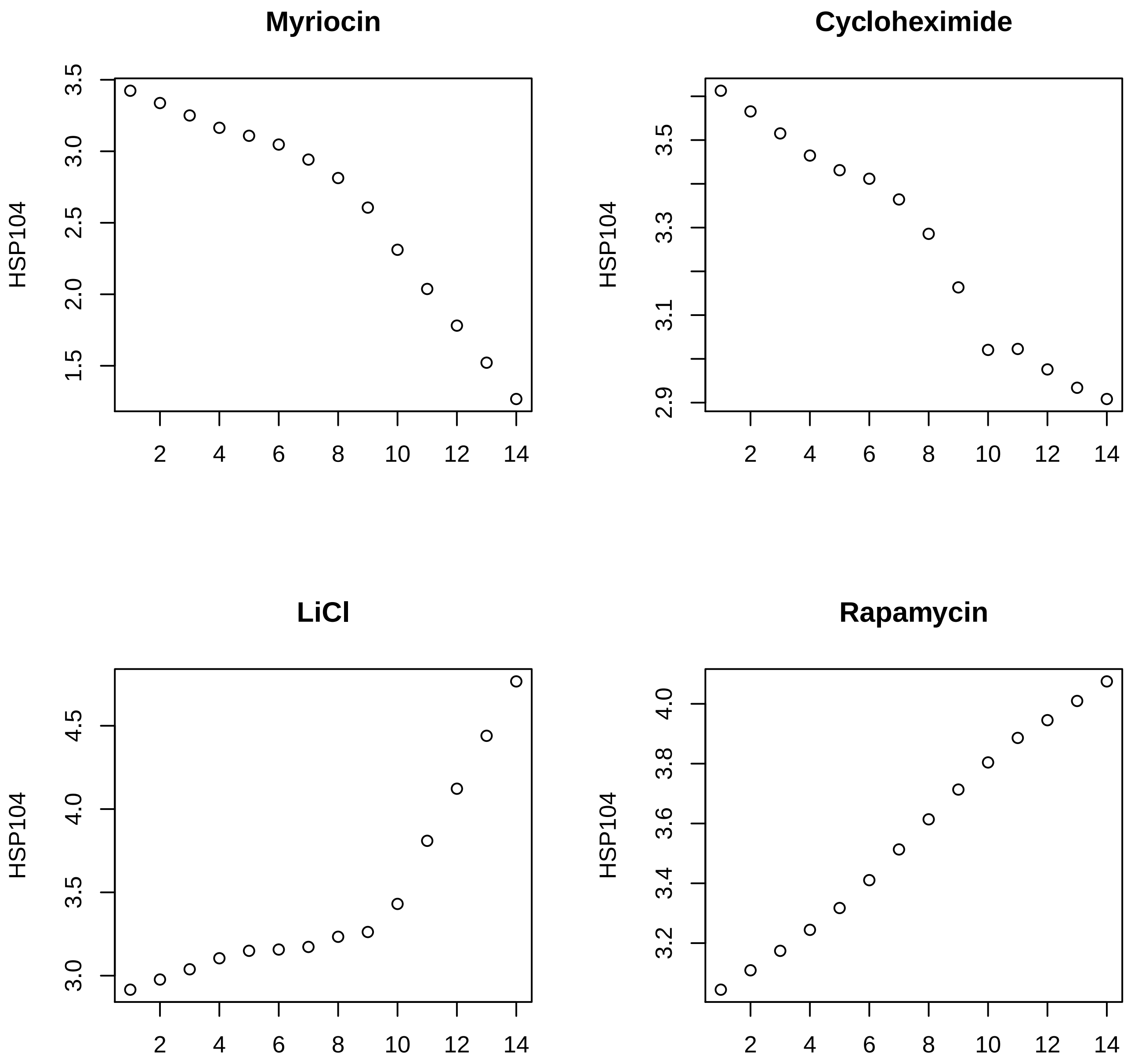
Myriocin
HSP104
3.5
3.0
2.5
2.0
1.5
2 4 6 8 10 12 14
Cycloheximide
HSP104
3.5
3.3
3.1
2.9
2 4 6 8 10 12 14
LiCl
HSP104
4.5
4.0
3.5
3.0
2 4 6 8 10 12 14
Rapamycin
HSP104
4.0
3.8
3.6
3.4
3.2
2 4 6 8 10 12 14

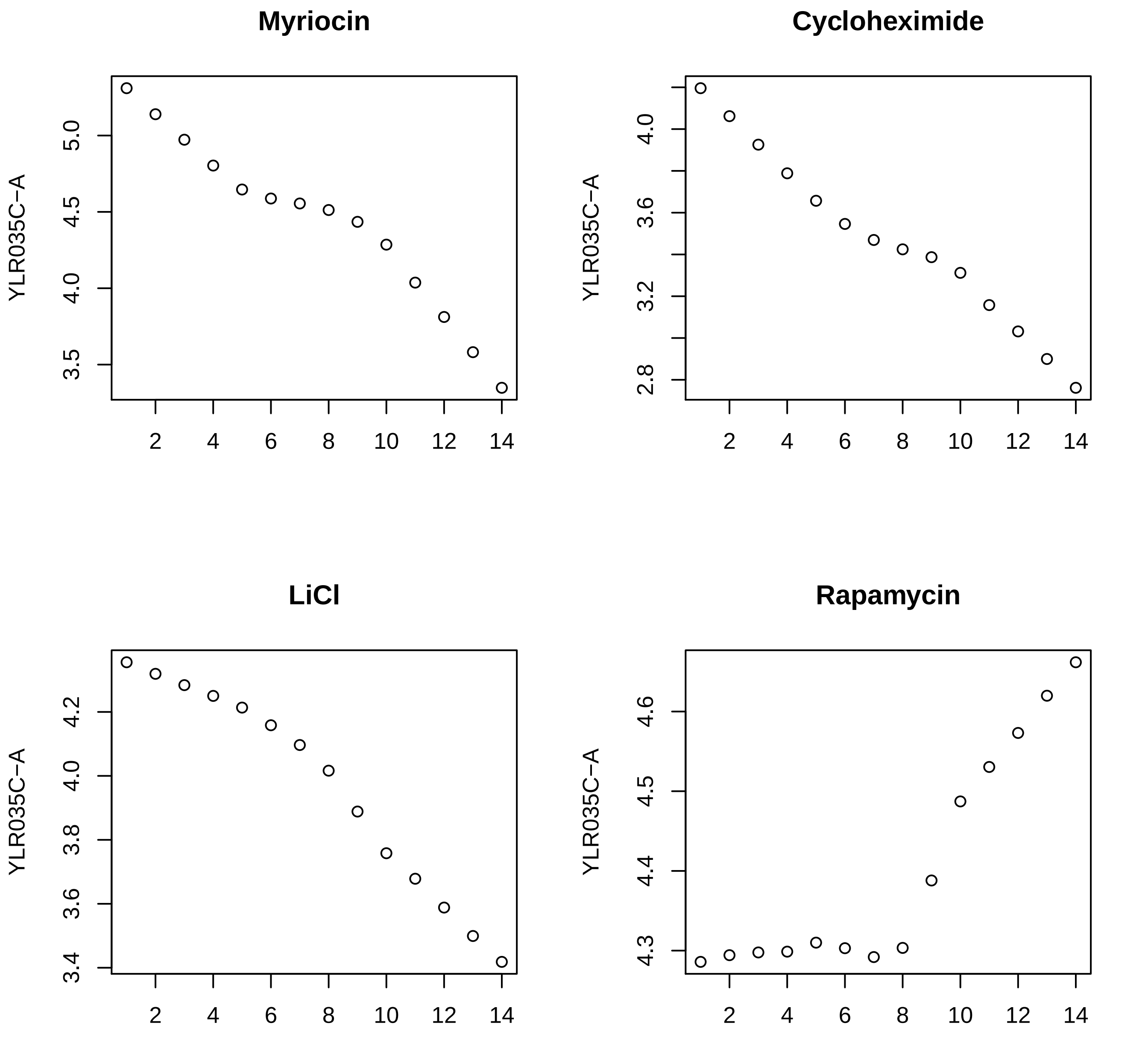
Myriocin
Cyclohexamide
LiCl
Rapamycin
YLR035C−A
5.0
4.5
4.0
3.5
2.8
3.2
3.6
4.2
3.4
3.8
4.3
4.4
4.5
4.6
2
4
6
8
10
12
14

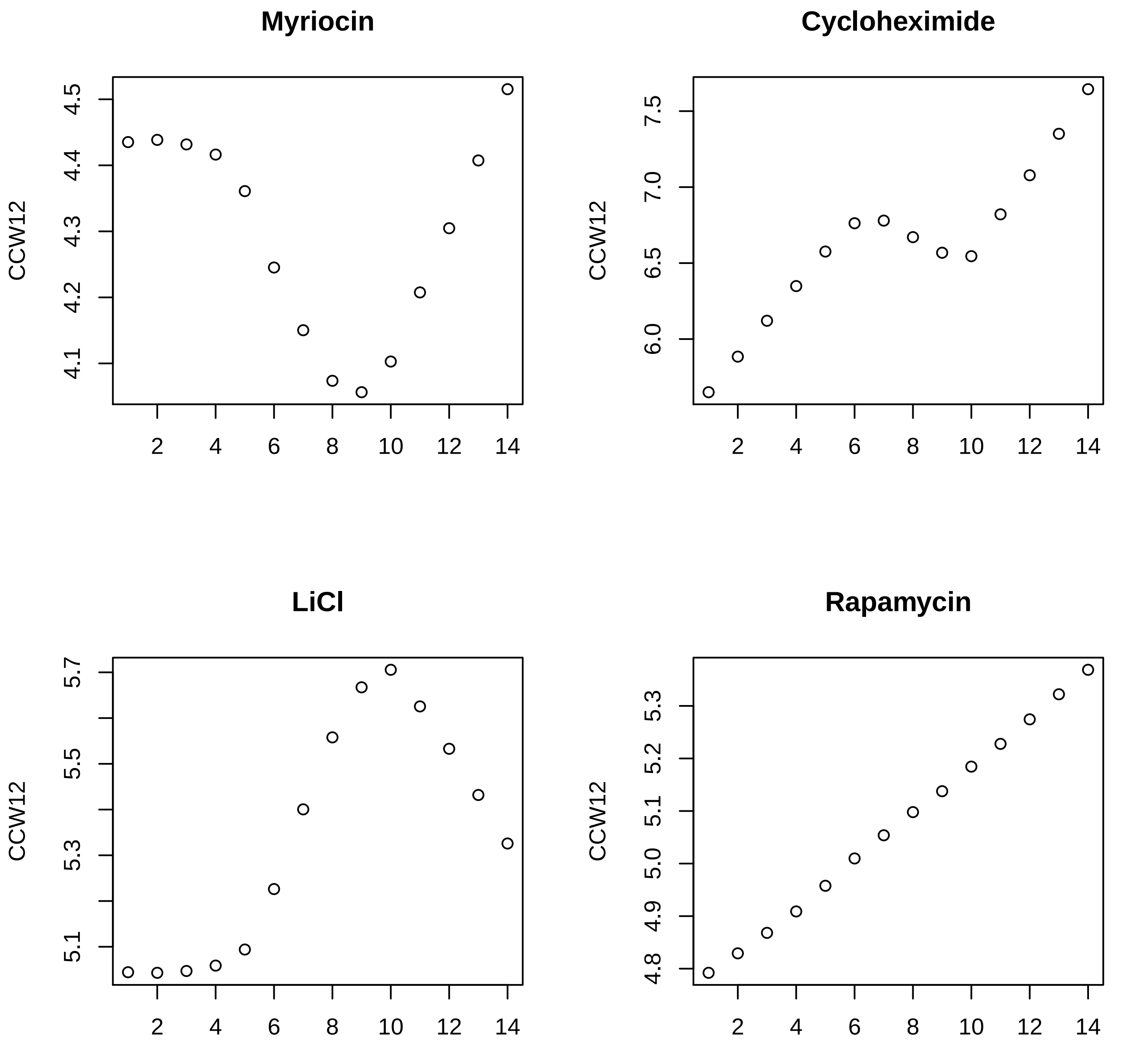
Myriocin
CCW12
4.1
4.2
4.3
4.4
4.5
2
4
6
8
10
12
14
Cycloheximide
CCW12
6.0
6.5
7.0
7.5
2
4
6
8
10
12
14
LiCl
CCW12
5.1
5.3
5.5
5.7
2
4
6
8
10
12
14
Rapamycin
CCW12
4.8
4.9
5.0
5.1
5.2
5.3
2
4
6
8
10
12
14

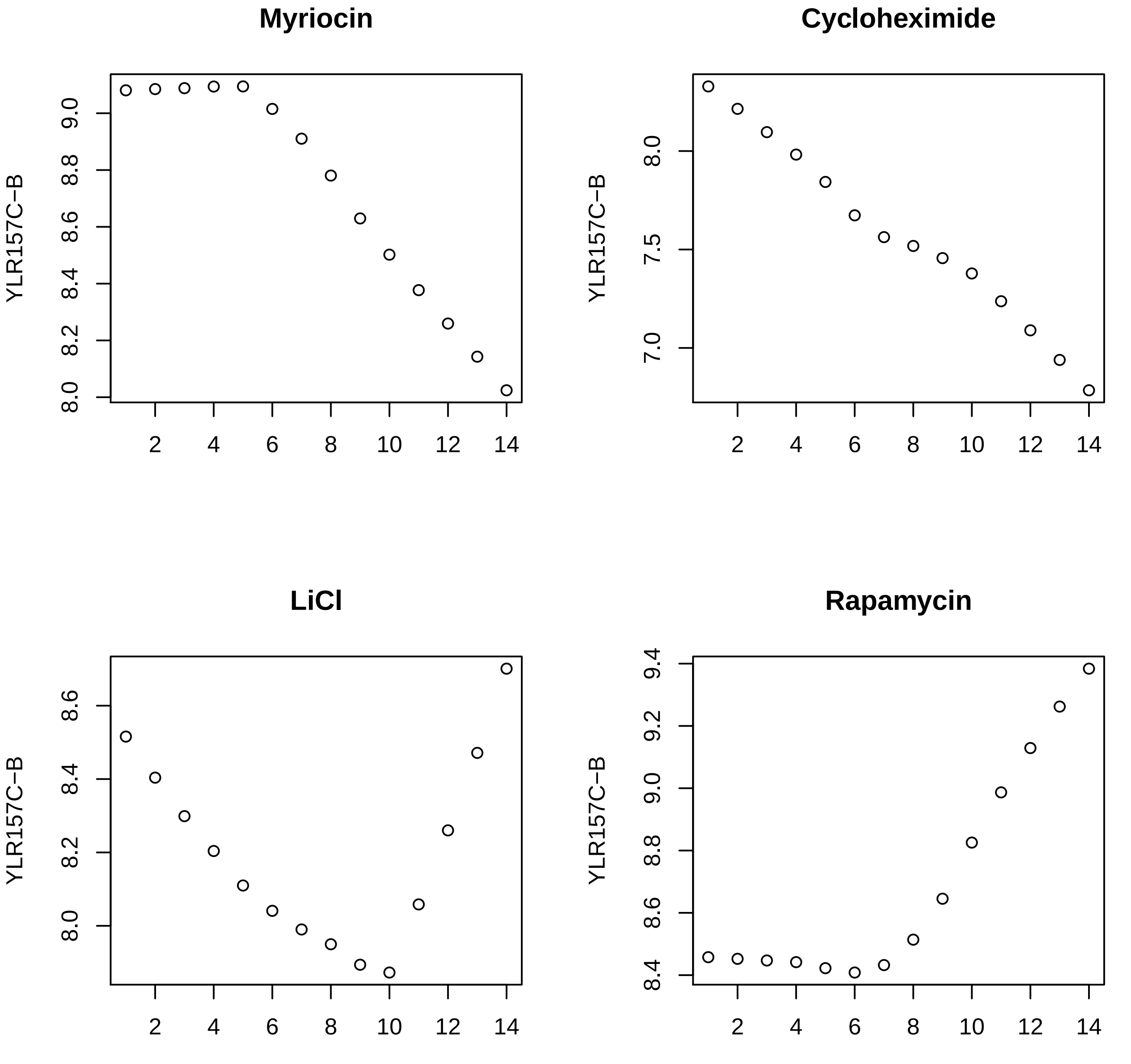
Myriocin
Cycloheximide
LiCl
Rapamycin
YLR157C−B
YLR157C−B
YLR157C−B
YLR157C−B
8.0
8.2
8.4
8.6
8.8
9.0
7.0
7.5
8.0
8.0
8.2
8.4
8.6
8.4
8.6
8.8
9.0
9.2
9.4
2
4
6
8
10
12
14

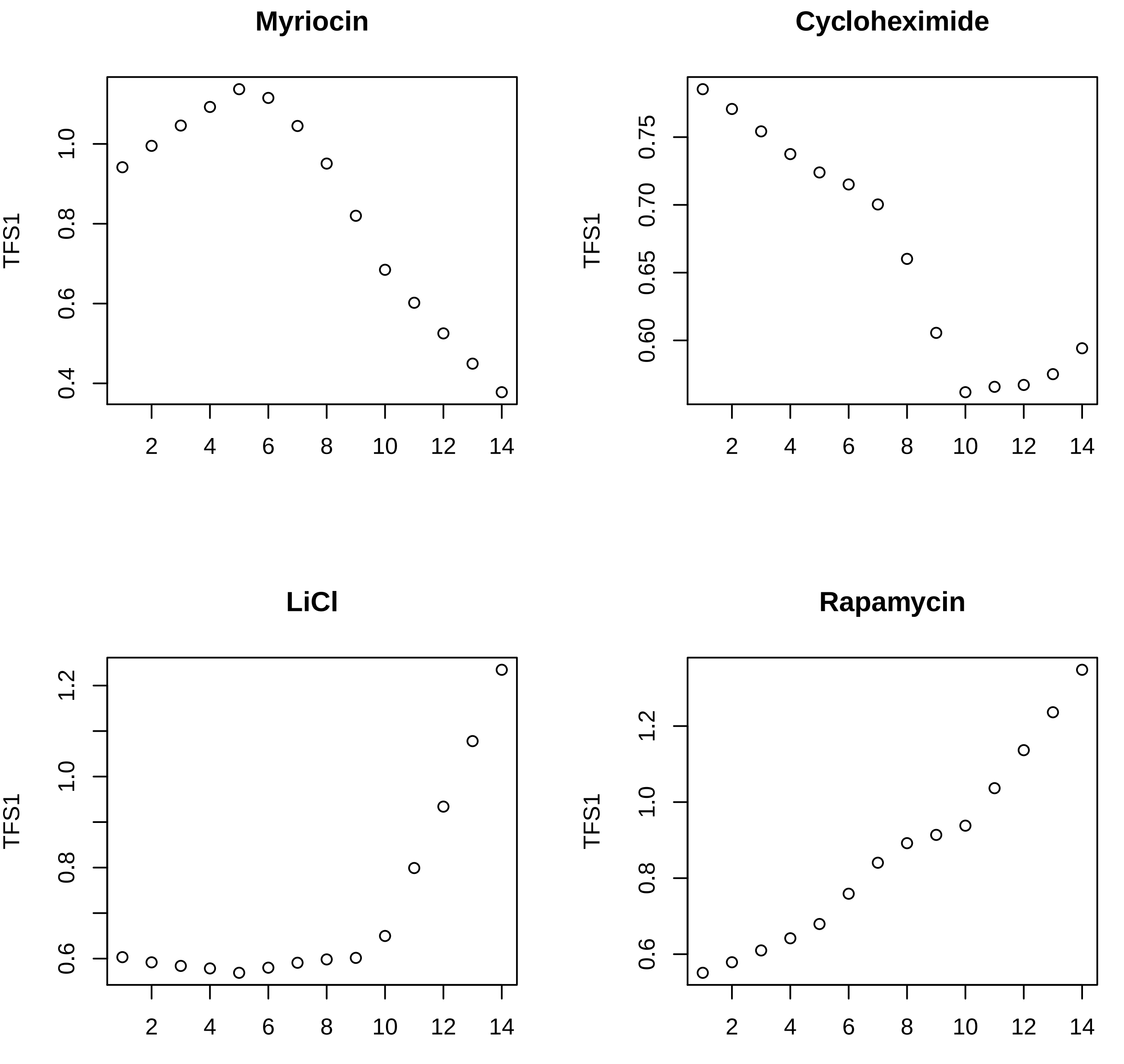
Myriocin
Cyclohexeximide
LiCl
Rapamycin
TFS1
0.4
0.6
0.8
1.0
1.2
0.60
0.65
0.70
0.75
2
4
6
8
10
12
14

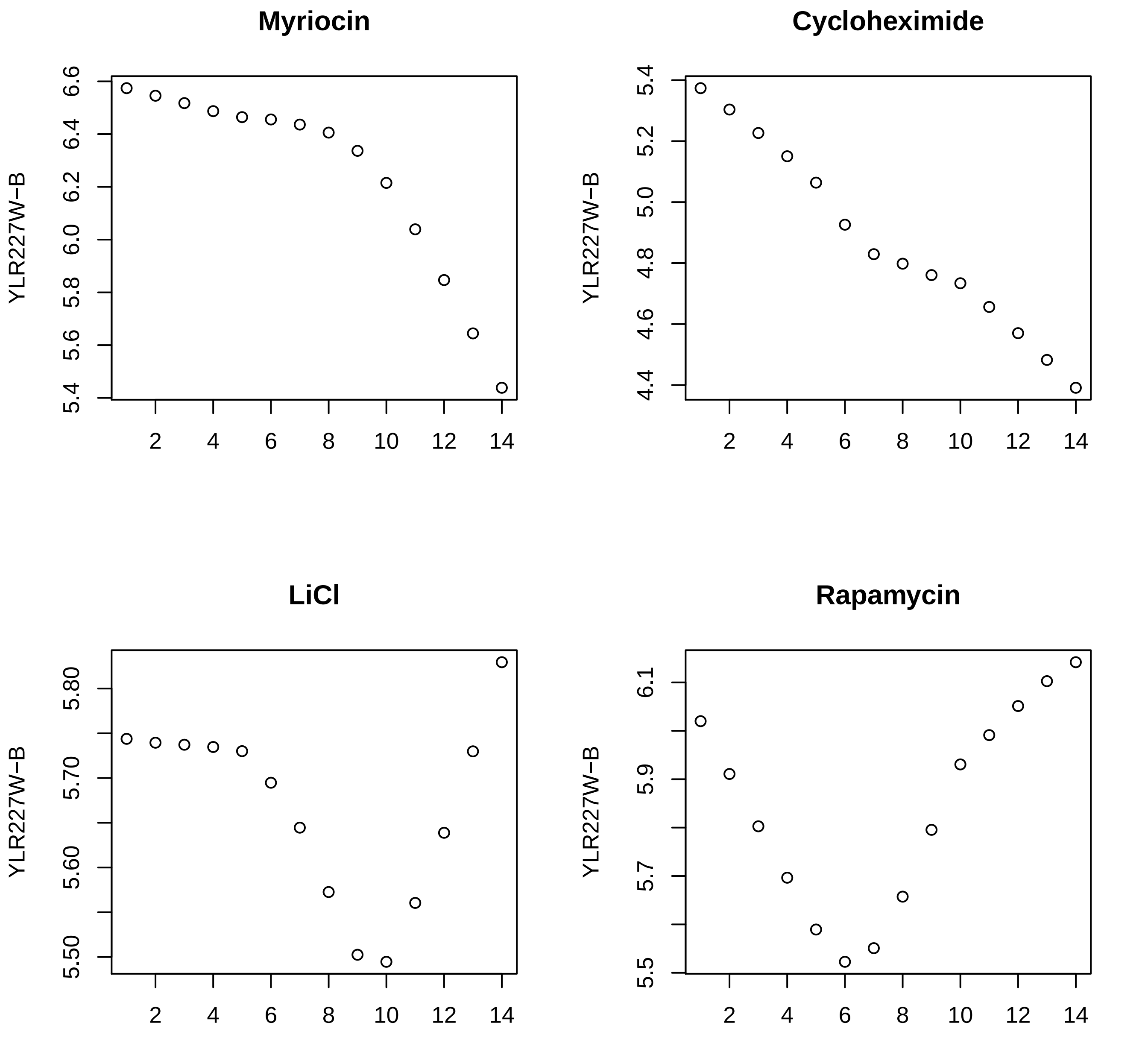

Myriocin
YLR227W−B
6.6
6.4
6.2
6.0
5.8
5.6
5.4
2
4
6
8
10
12
14
Cycloheximide
YLR227W−B
5.4
5.2
5.0
4.8
4.6
4.4
LiCl
YLR227W−B
5.80
5.70
5.60
5.50
Rapamycin
YLR227W−B
6.1
5.9
5.7
5.5

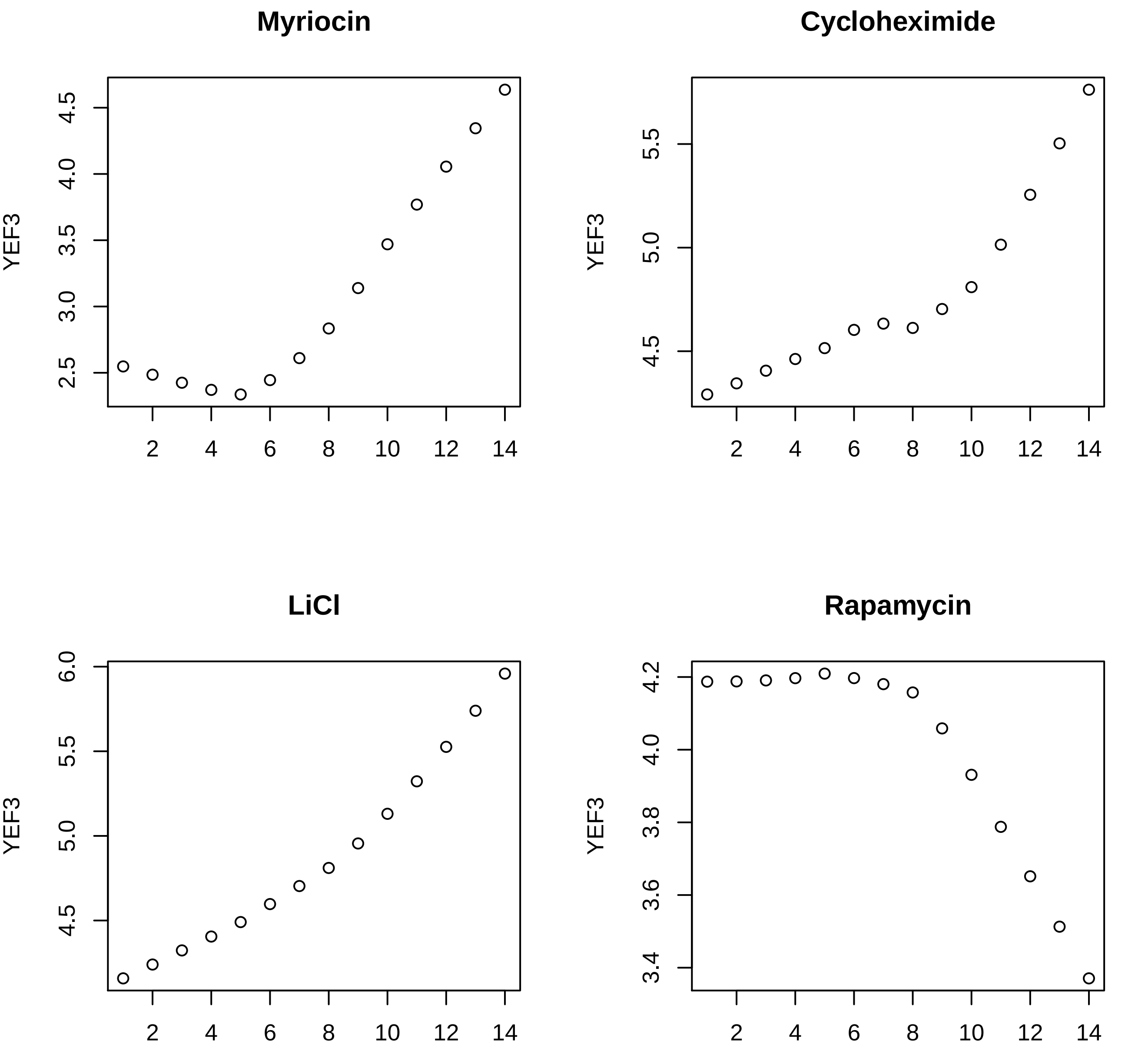
Myriocin
Cycloheximide
LiCl
Rapamycin
YEF3
YEF3
YEF3
YEF3
2.5
3.0
3.5
4.0
4.5
4.5
5.0
5.5
4.5
5.0
5.5
6.0
3.4
3.6
3.8
4.0
4.2
2
4
6
8
10
12
14

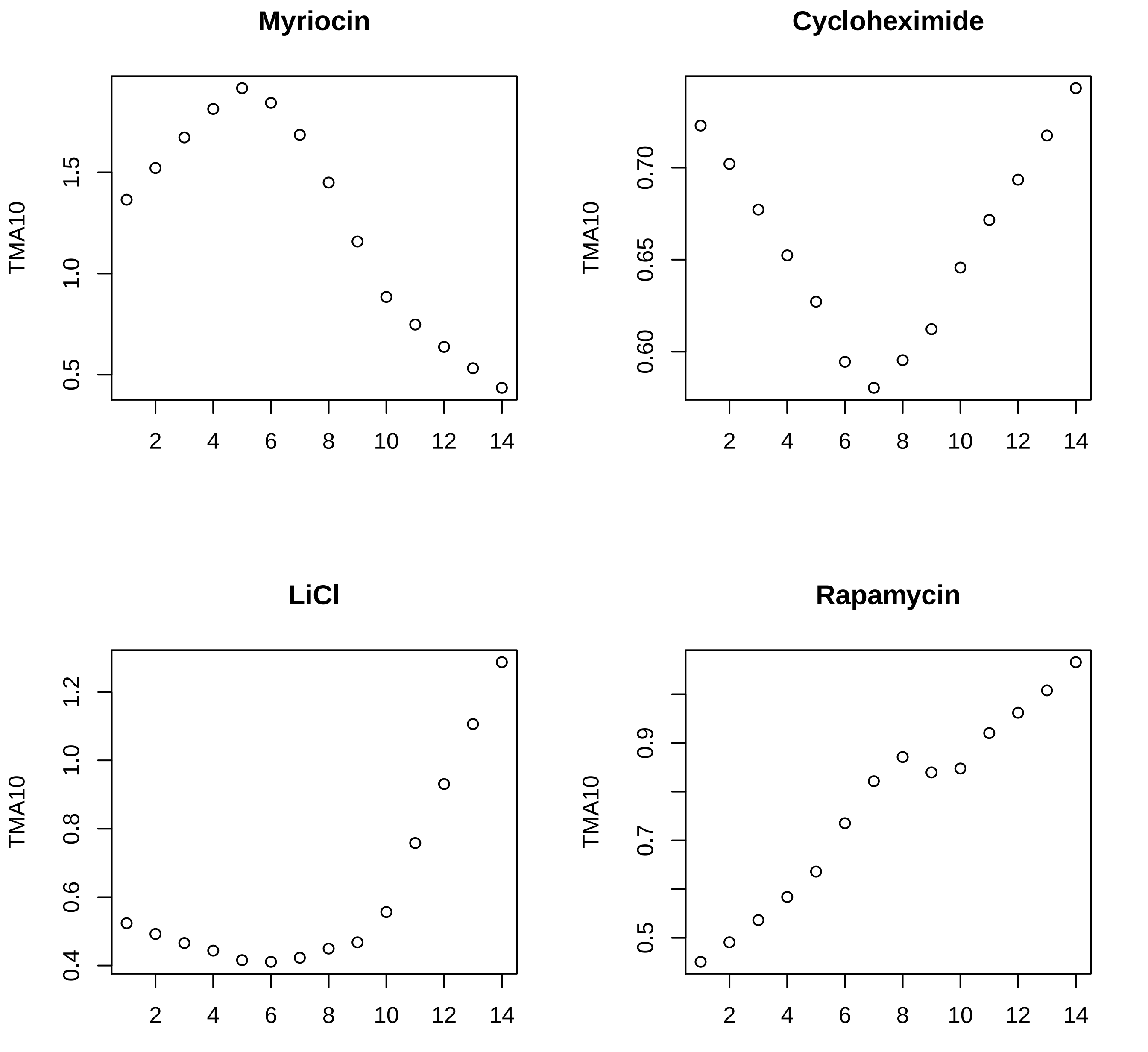
Myriocin
Cycloheximide
LiCl
Rapamycin
TMA10
0.5
1.0
1.5
0.60
0.65
0.70
0.4
0.6
0.8
1.0
1.2
0.5
0.7
0.9
2
4
6
8
10
12
14

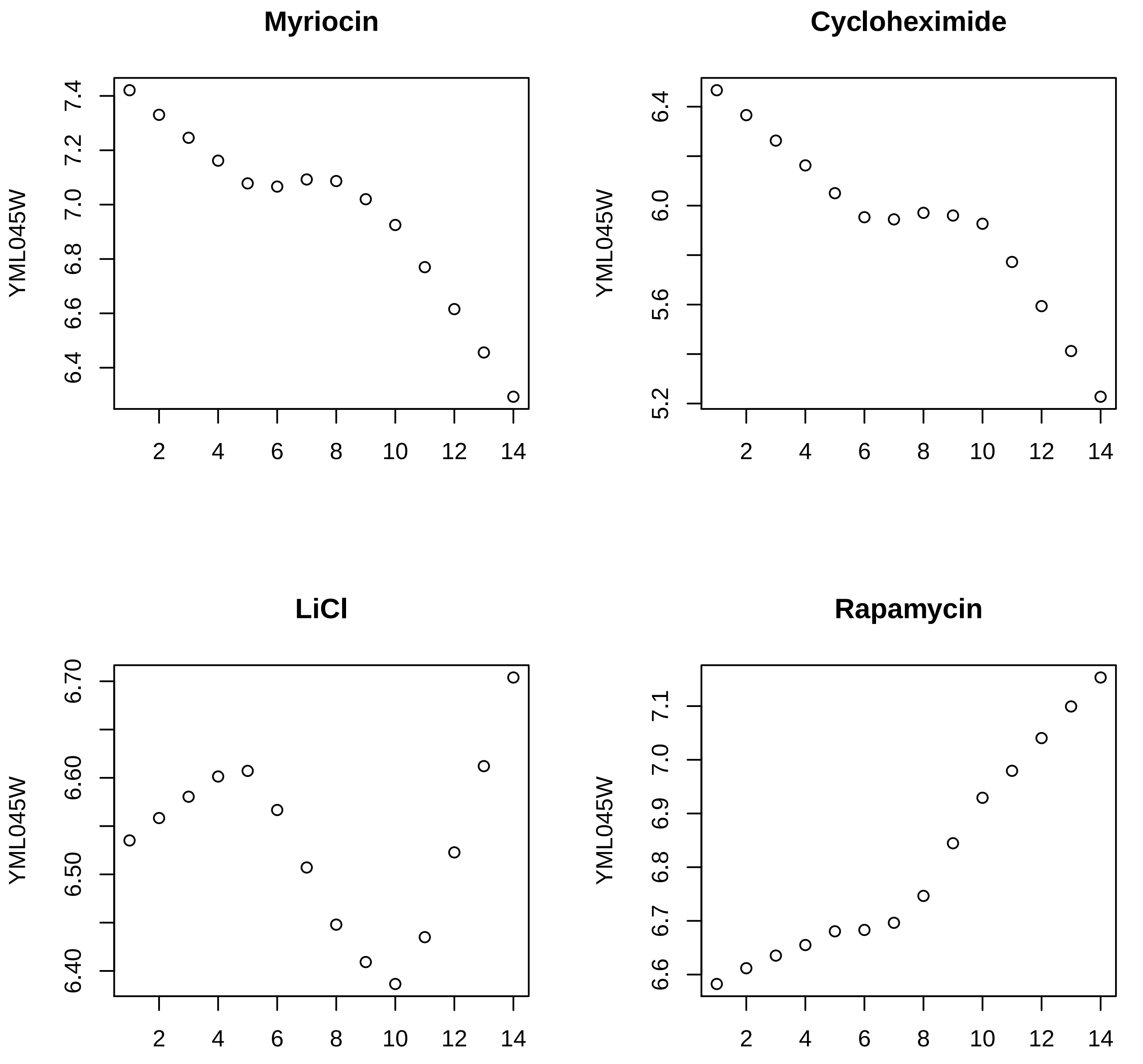
Myriocin
YML045W
7.4
7.2
7.0
6.8
6.6
6.4
2
4
6
8
10
12
14
Cycloheximide
YML045W
6.4
6.0
5.6
5.2
2
4
6
8
10
12
14
LiCl
YML045W
6.70
6.60
6.50
6.40
2
4
6
8
10
12
14
Rapamycin
YML045W
7.1
7.0
6.9
6.8
6.7
6.6
2
4
6
8
10
12
14

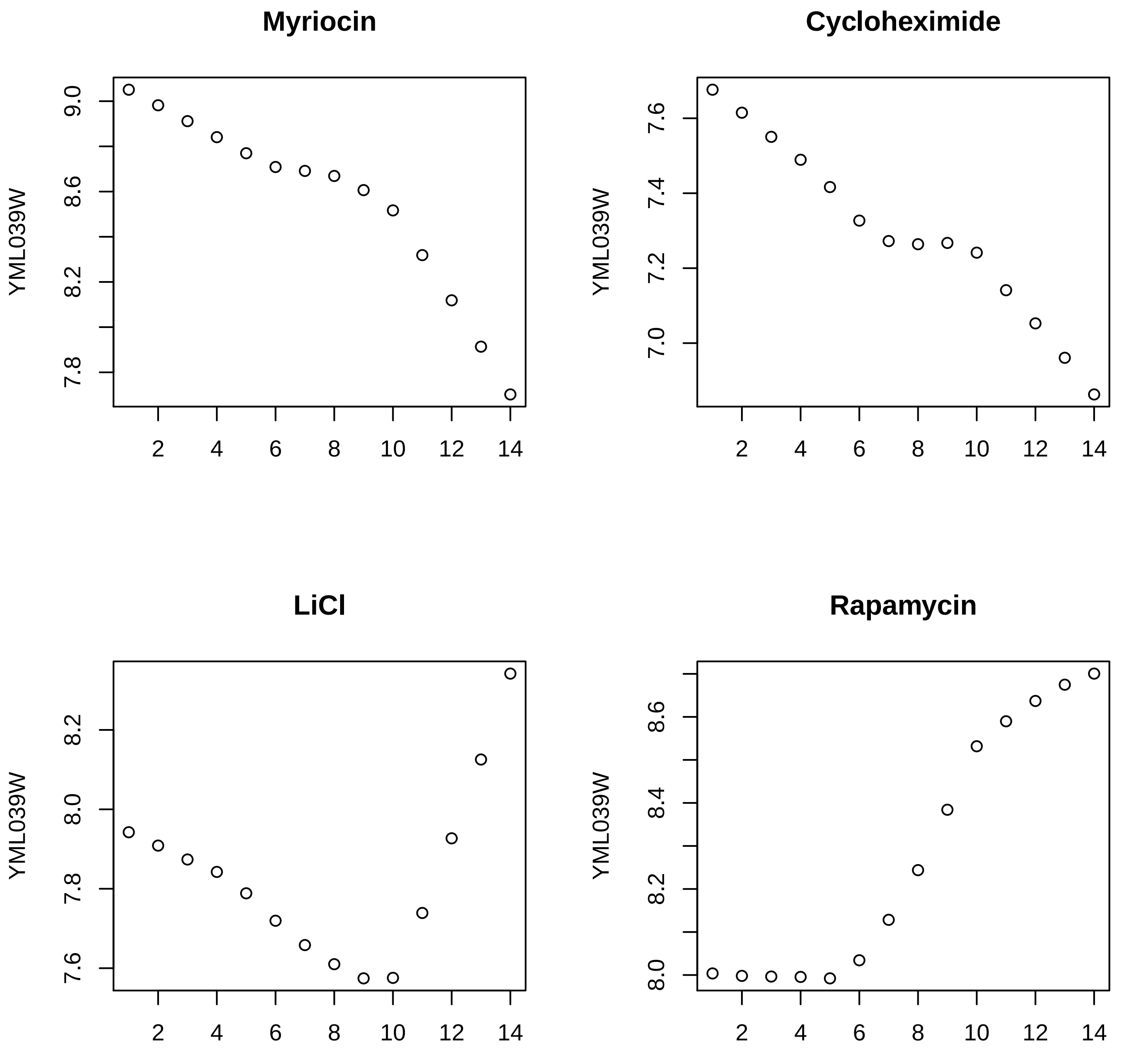
Myriocin
Cycloheximide
LiCl
Rapamycin
YML039W
YML039W
YML039W
YML039W
9.0
8.6
8.2
7.8
7.6
7.4
7.2
7.0
8.2
8.0
7.8
7.6
8.6
8.4
8.2
8.0
2
4
6
8
10
12
14

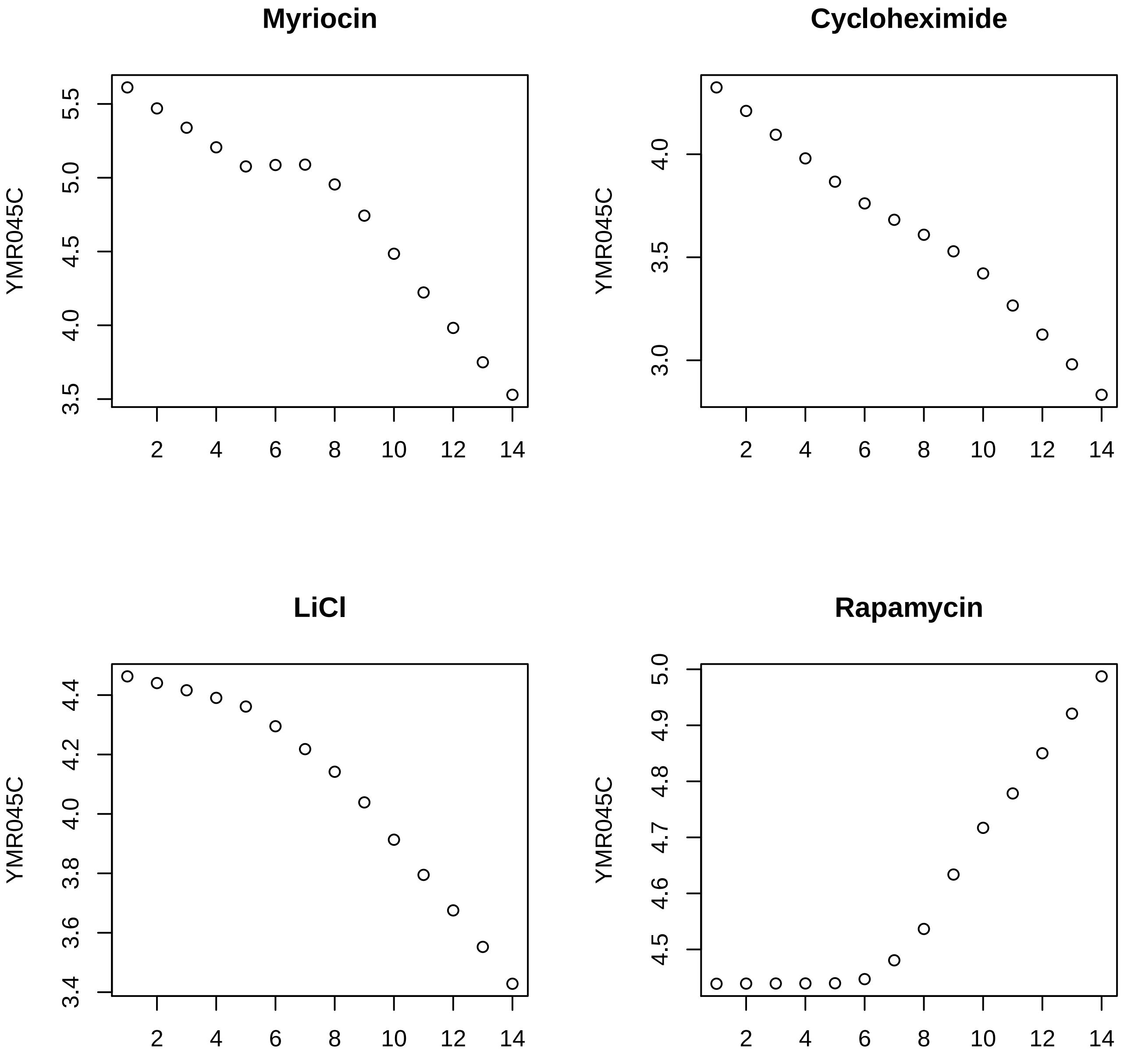
Myriocin
Cycloheximide
LiCl
Rapamycin
YMR045C
YMR045C
YMR045C
YMR045C
3.5
4.0
4.5
5.0
5.5
3.0
3.5
4.0
3.4
3.6
3.8
4.0
4.2
4.4
4.5
4.6
4.7
4.8
4.9
5.0
2
4
6
8
10
12
14

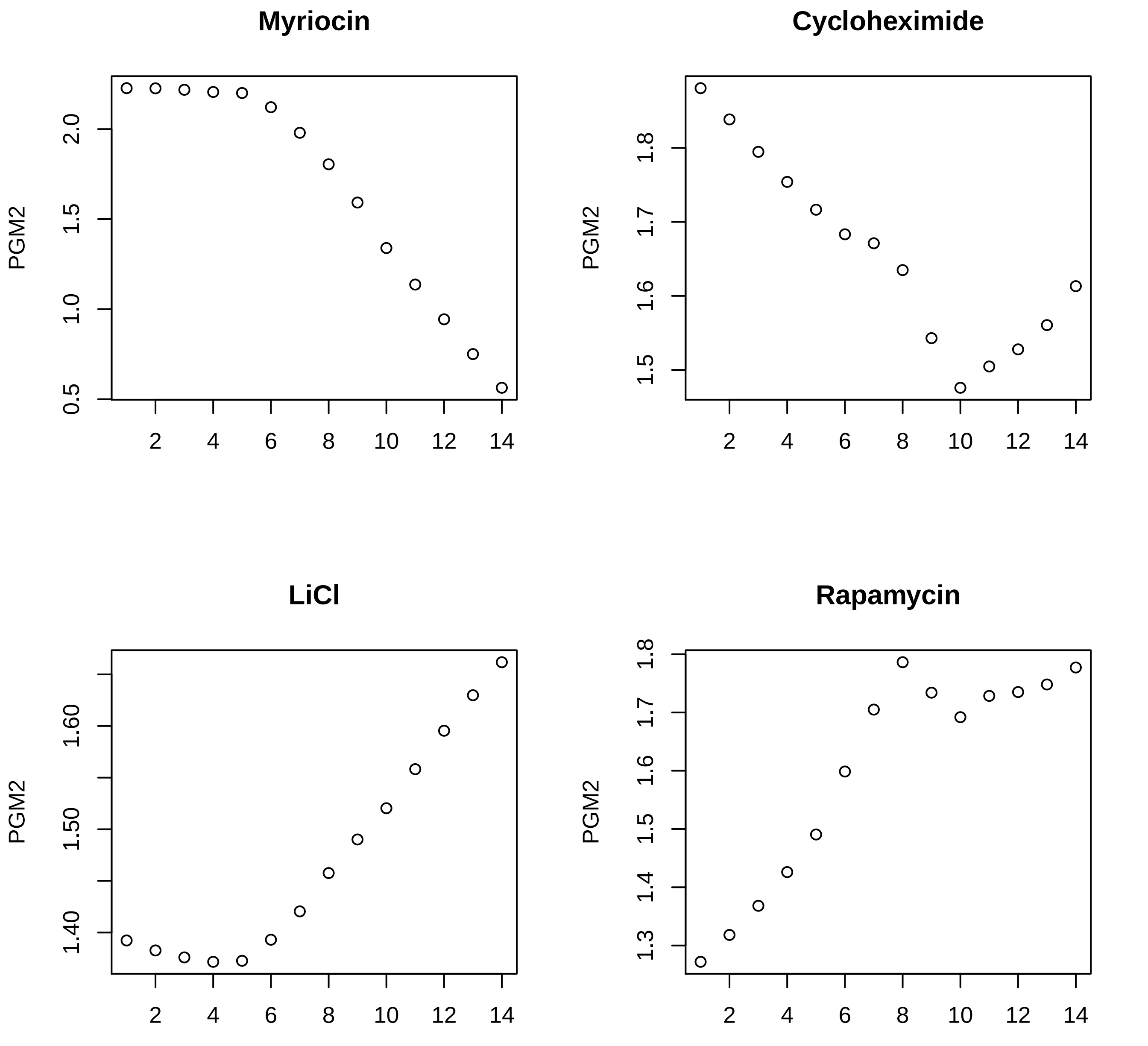
Myriocin
PGM2
0.5
1.0
1.5
2.0
2
4
6
8
10
12
14
Cycloheximide
PGM2
1.5
1.6
1.7
1.8
2
4
6
8
10
12
14
LiCl
PGM2
1.40
1.50
1.60
2
4
6
8
10
12
14
Rapamycin
PGM2
1.3
1.4
1.5
1.6
1.7
1.8
2
4
6
8
10
12
14

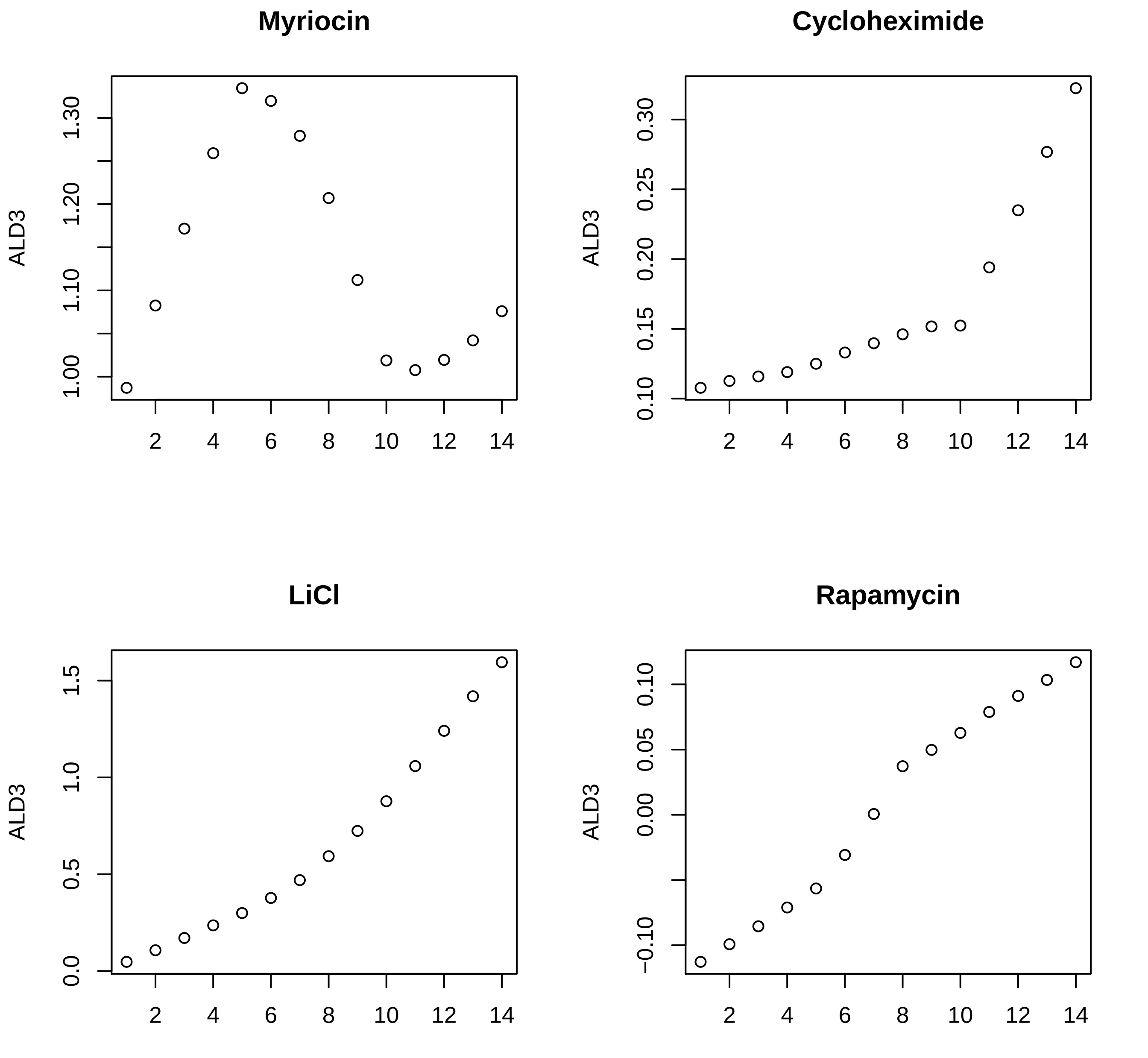

Myriocin
ALD3
1.00
1.10
1.20
1.30
2
4
6
8
10
12
14
Cycloheximide
ALD3
0.10
0.15
0.20
0.25
0.30
2
4
6
8
10
12
14
LiCl
ALD3
0.0
0.5
1.0
1.5
2
4
6
8
10
12
14
Rapamycin
ALD3
−0.10
0.00
0.05
0.10
2
4
6
8
10
12
14

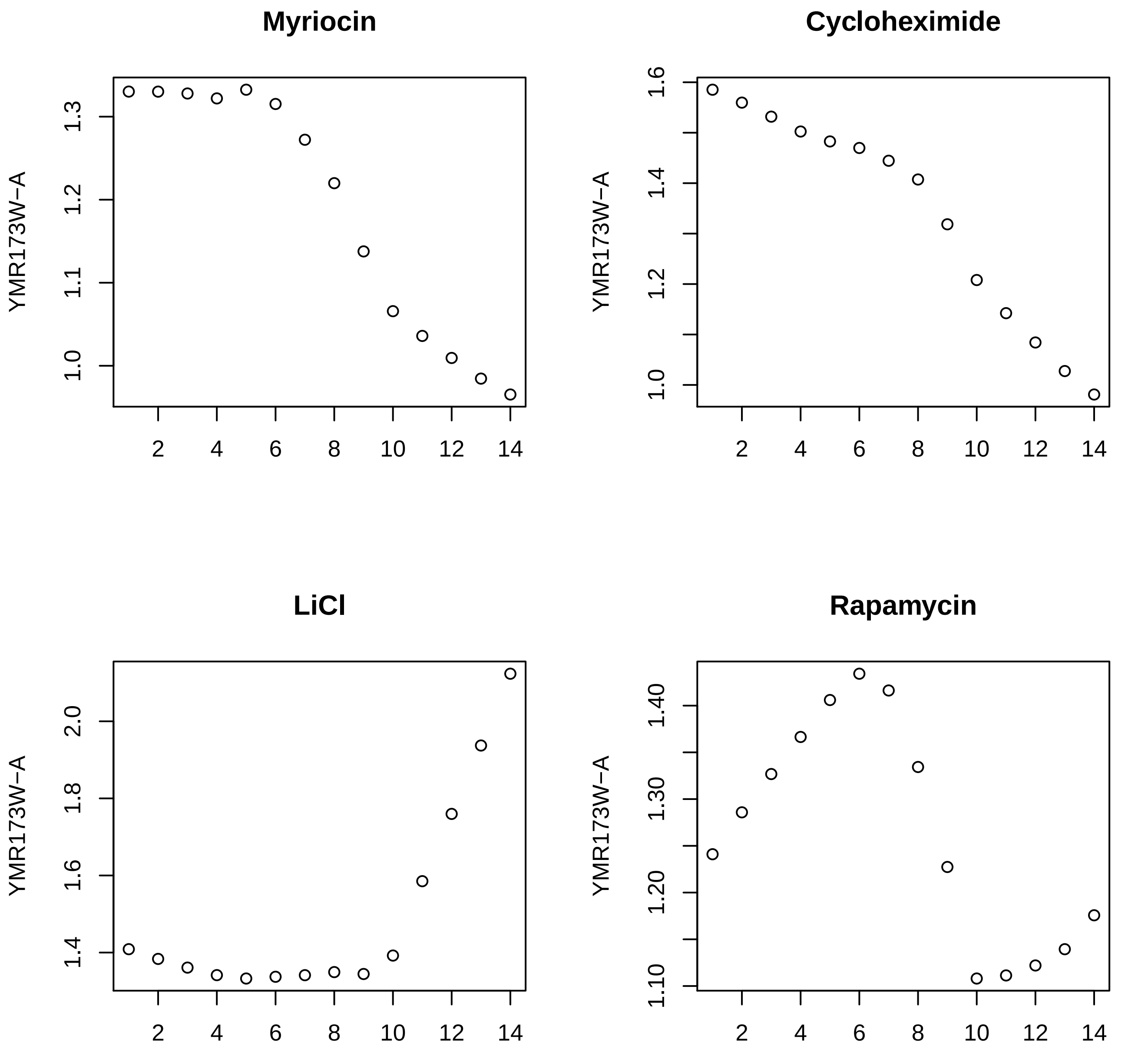

Myriocin
Cycloheximide
LiCl
Rapamycin
YMR173W−A
1.0
1.1
1.2
1.3
1.4
1.6
1.8
2.0
1.10
1.20
1.30
1.40
2
4
6
8
10
12
14

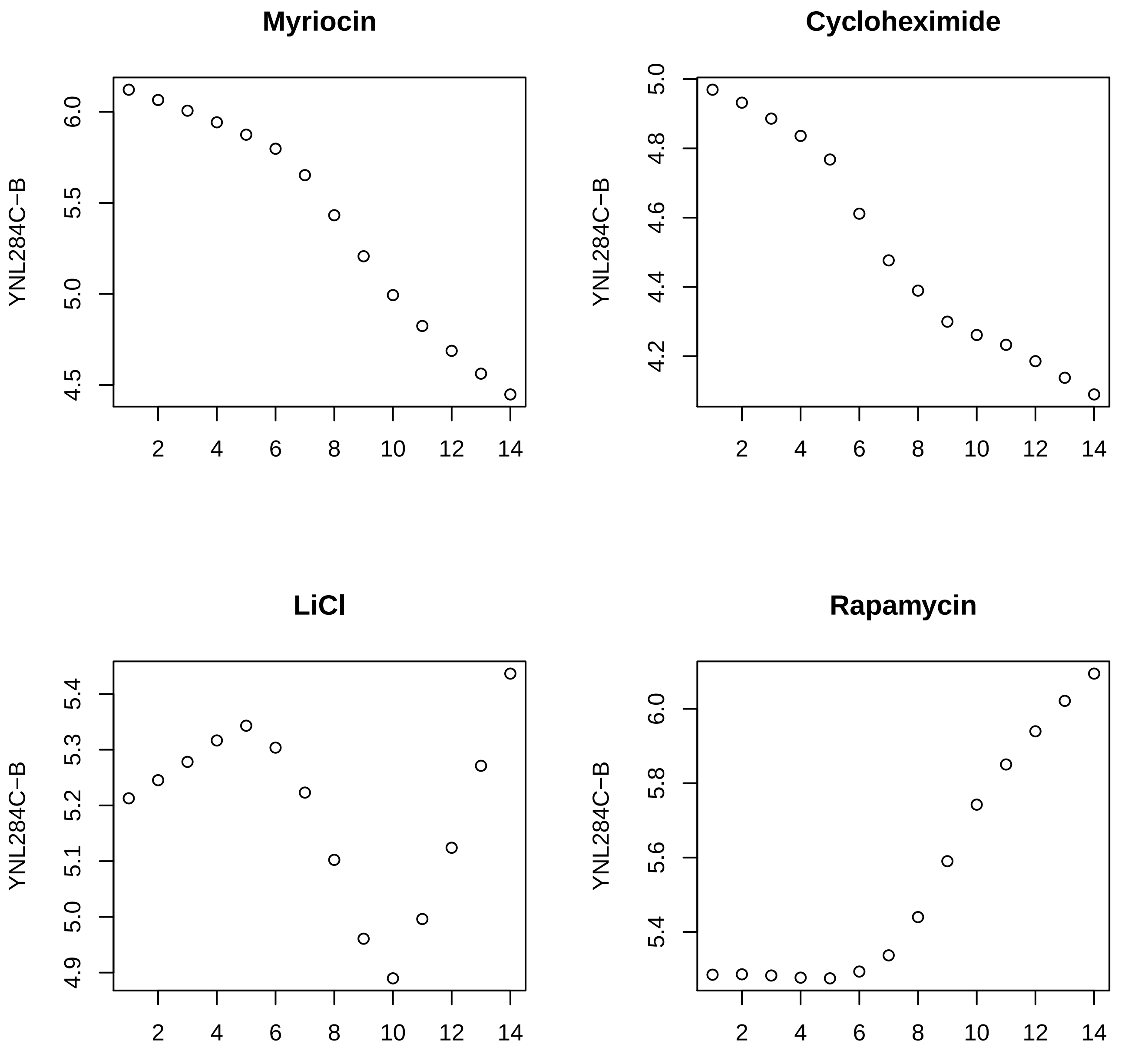

Myriocin
Cycloheximide
LiCl
Rapamycin
YNL284C−B
YNL284C−B
YNL284C−B
YNL284C−B
6.0
5.5
5.0
4.5
5.0
4.8
4.6
4.4
4.2
5.4
5.3
5.2
5.1
5.0
4.9
6.0
5.8
5.6
5.4
2
4
6
8
10
12
14

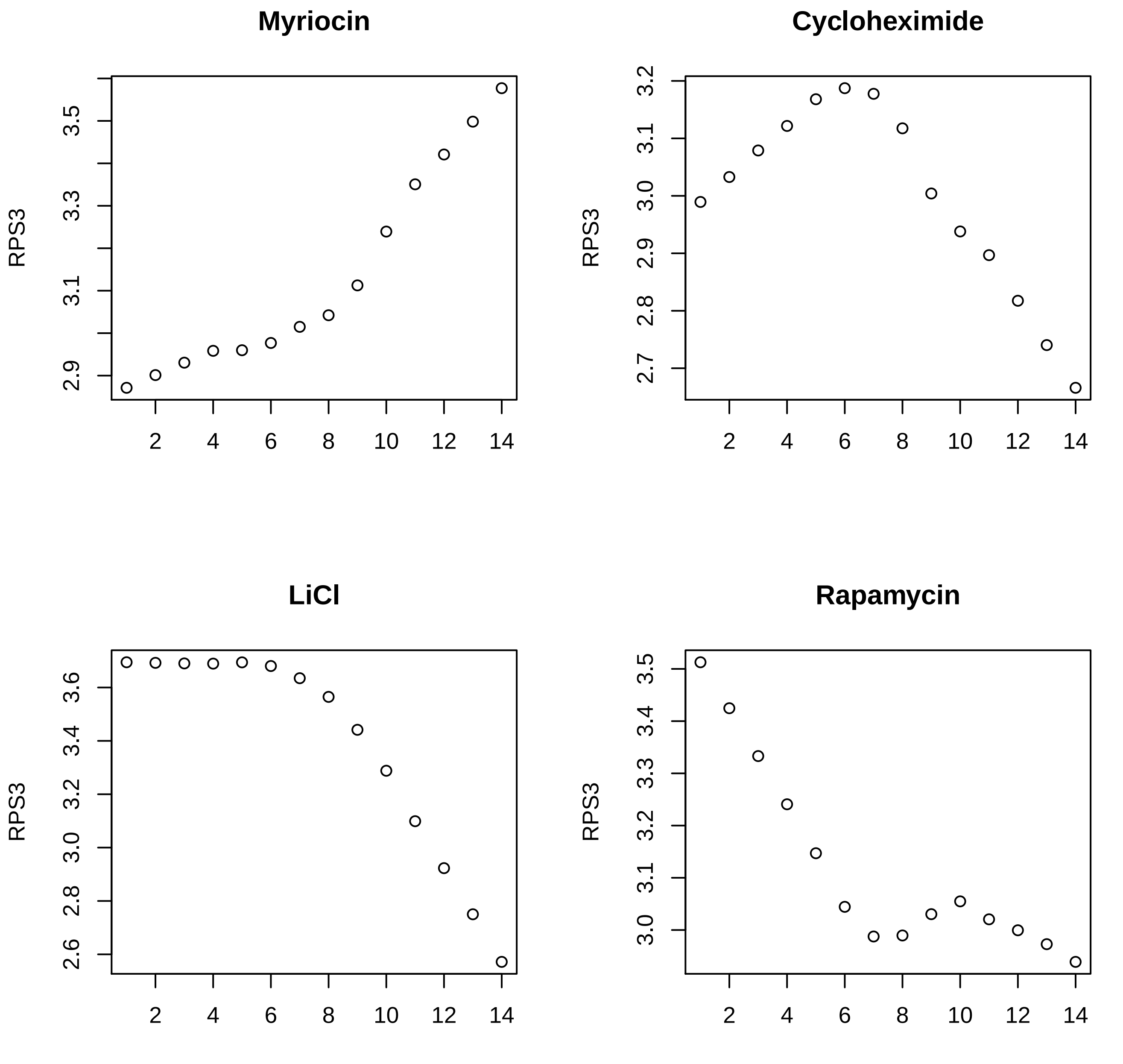
Myriocin
RPS3
2.9
3.1
3.3
3.5
2
4
6
8
10
12
14
Cycloheximide
RPS3
2.7
2.8
2.9
3.0
3.1
3.2
2
4
6
8
10
12
14
LiCl
RPS3
2.6
2.8
3.0
3.2
3.4
3.6
2
4
6
8
10
12
14
Rapamycin
RPS3
3.0
3.1
3.2
3.3
3.4
3.5
2
4
6
8
10
12
14

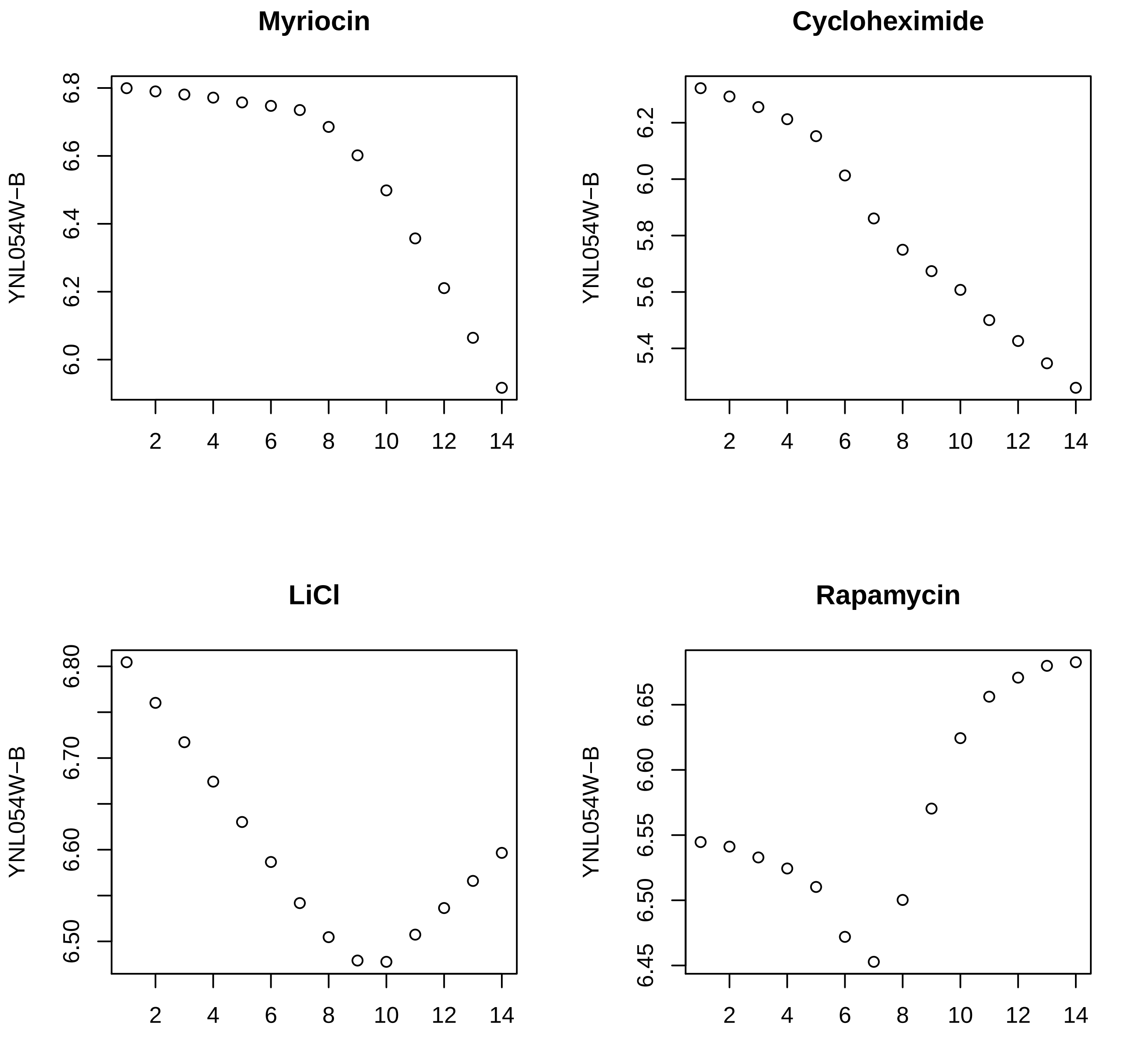
Myriocin
YNL054W−B
6.0
6.2
6.4
6.6
6.8
2
4
6
8
10
12
14
Cycloheximide
YNL054W−B
5.4
5.6
5.8
6.0
6.2
2
4
6
8
10
12
14
LiCl
YNL054W−B
6.50
6.60
6.70
6.80
2
4
6
8
10
12
14
Rapamycin
YNL054W−B
6.45
6.50
6.55
6.60
6.65
2
4
6
8
10
12
14

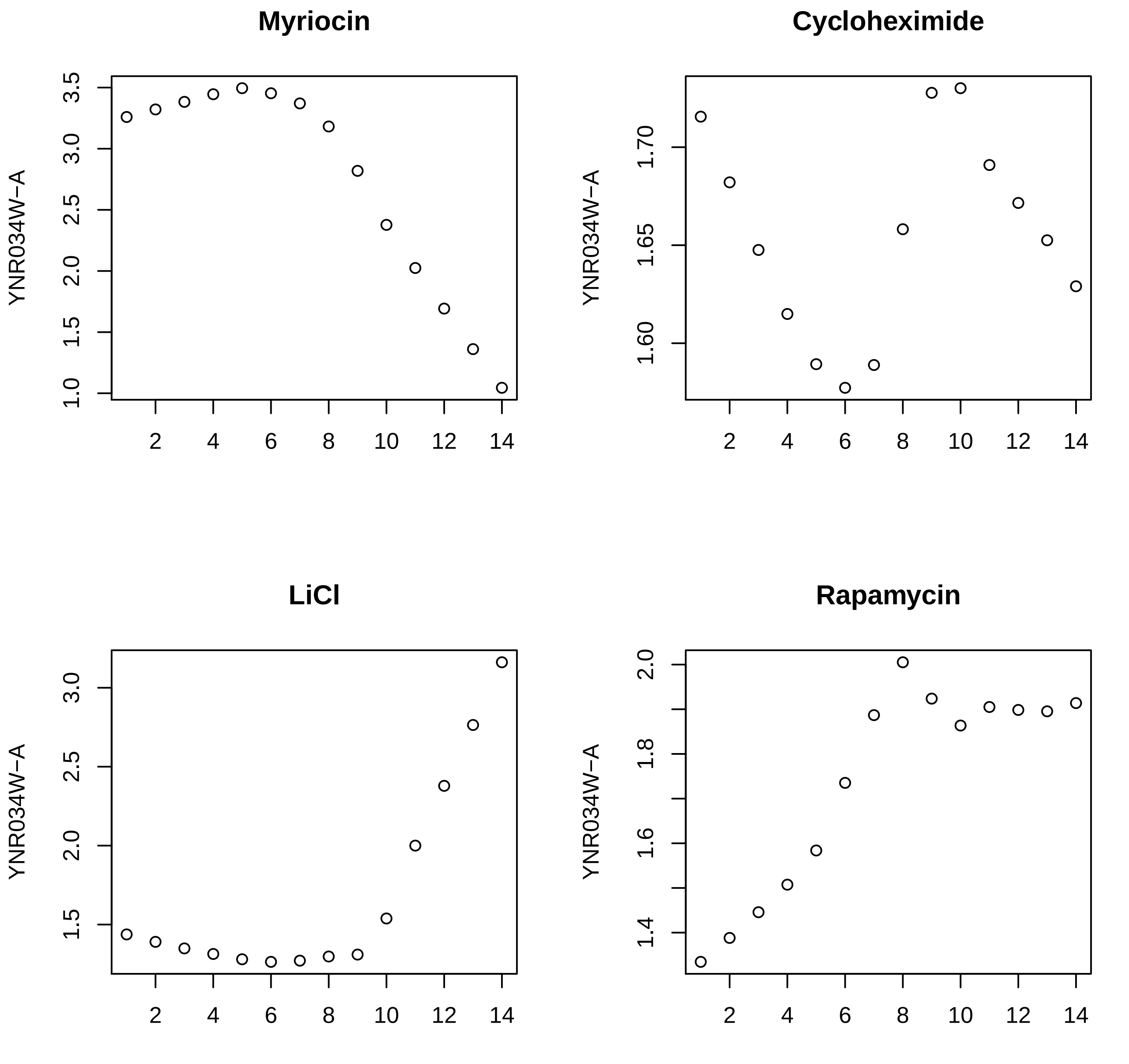
Myriocin
Cycloheximide
LiCl
Rapamycin
YNR034W−A

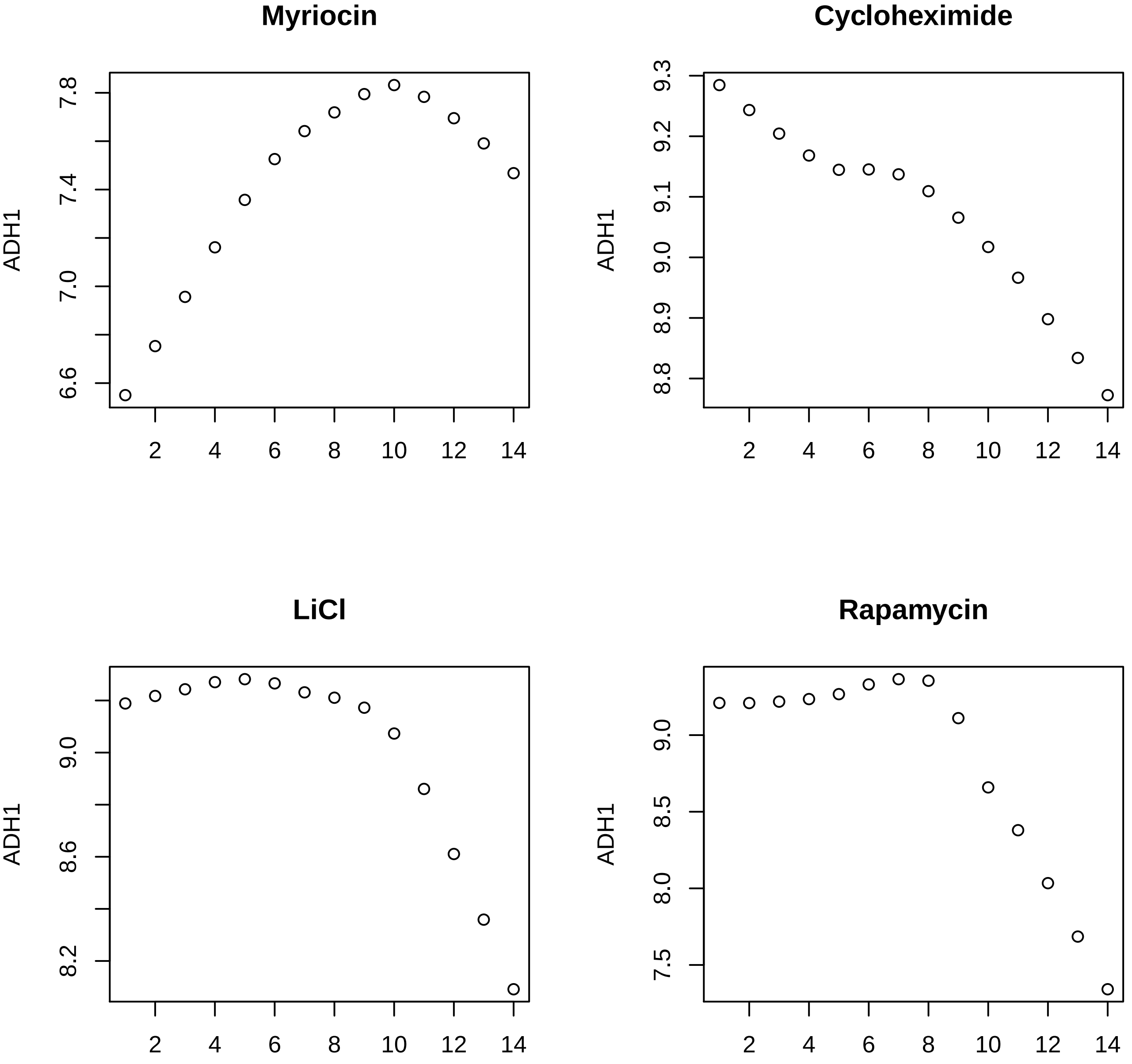

Myriocin
Cycloheximide
LiCl
Rapamycin
ADH1
7.8
7.4
7.0
6.6
9.3
9.2
9.1
9.0
8.9
8.8
8.6
8.2
8.5
8.0
7.5
2
4
6
8
10
12
14

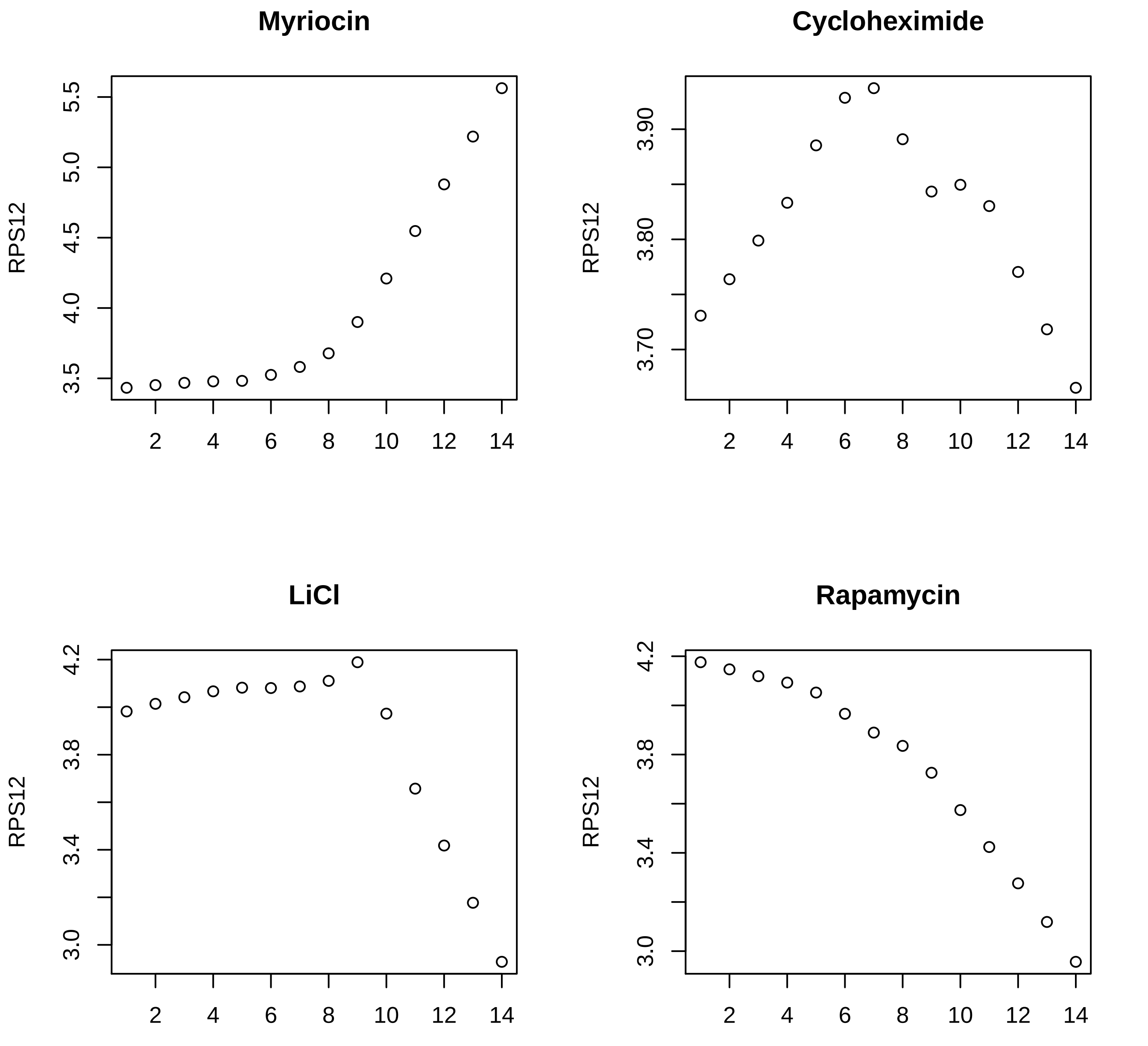
Myriocin
RPS12
3.5
4.0
4.5
5.0
5.5
2
4
6
8
10
12
14
Cycloheximide
RPS12
3.70
3.80
3.90
LiCl
RPS12
3.0
3.4
3.8
4.2
Rapamycin
RPS12
3.0
3.4
3.8
4.2

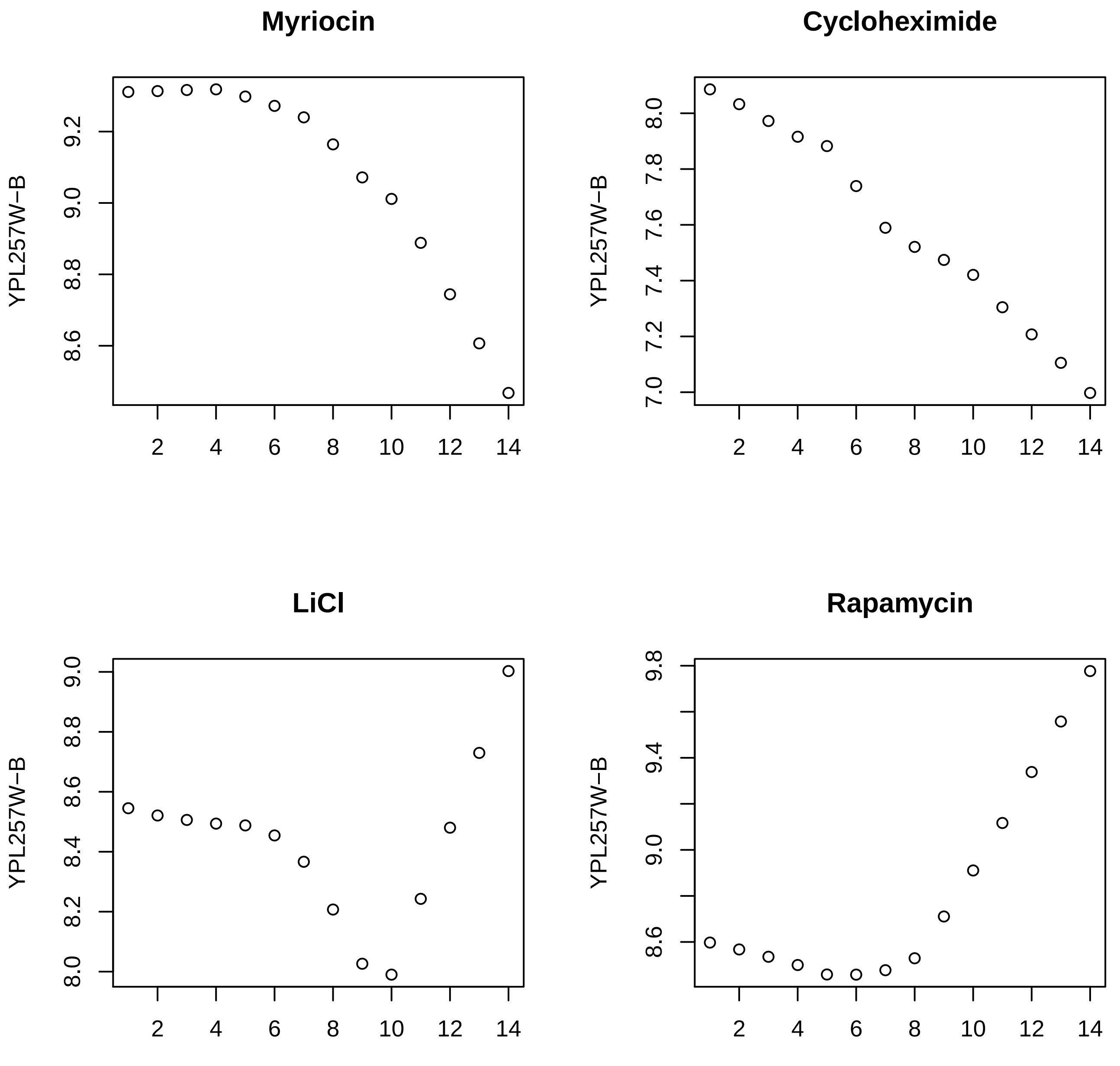
Myriocin
Cycloheximide
LiCl
Rapamycin
YPL257W−B
YPL257W−B
YPL257W−B
YPL257W−B
8.6 8.8 9.0 9.2
7.0 7.2 7.4 7.6 7.8 8.0
8.0 8.2 8.4 8.6 8.8 9.0
8.6 9.0 9.4 9.8
2 4 6 8 10 12 14

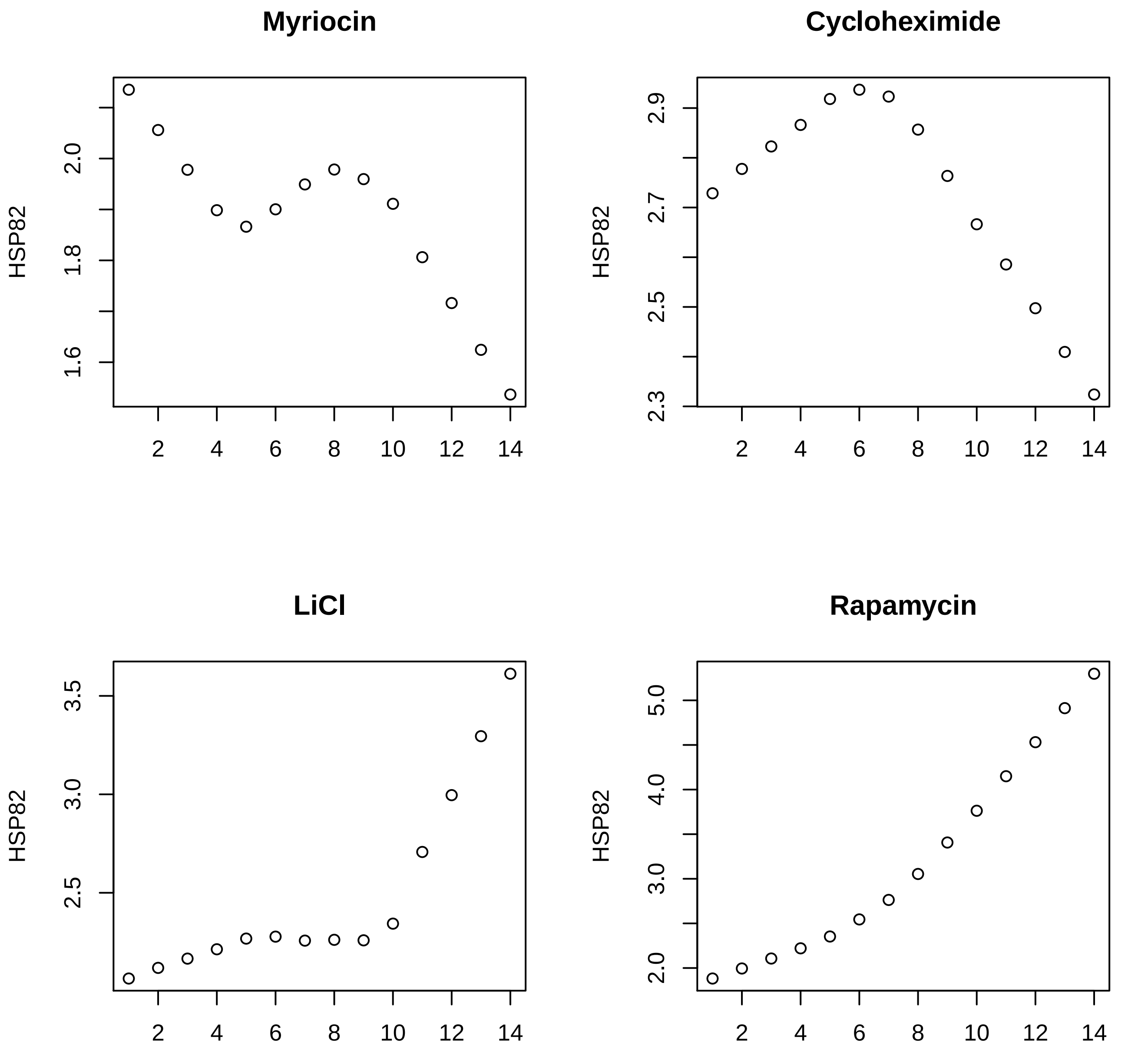
Myriocin
Cycloheximide
LiCl
Rapamycin
HSP82
2.0
1.8
1.6
2.9
2.7
2.5
2.3
3.5
3.0
2.5
5.0
4.0
3.0
2.0
2
4
6
8
10
12
14

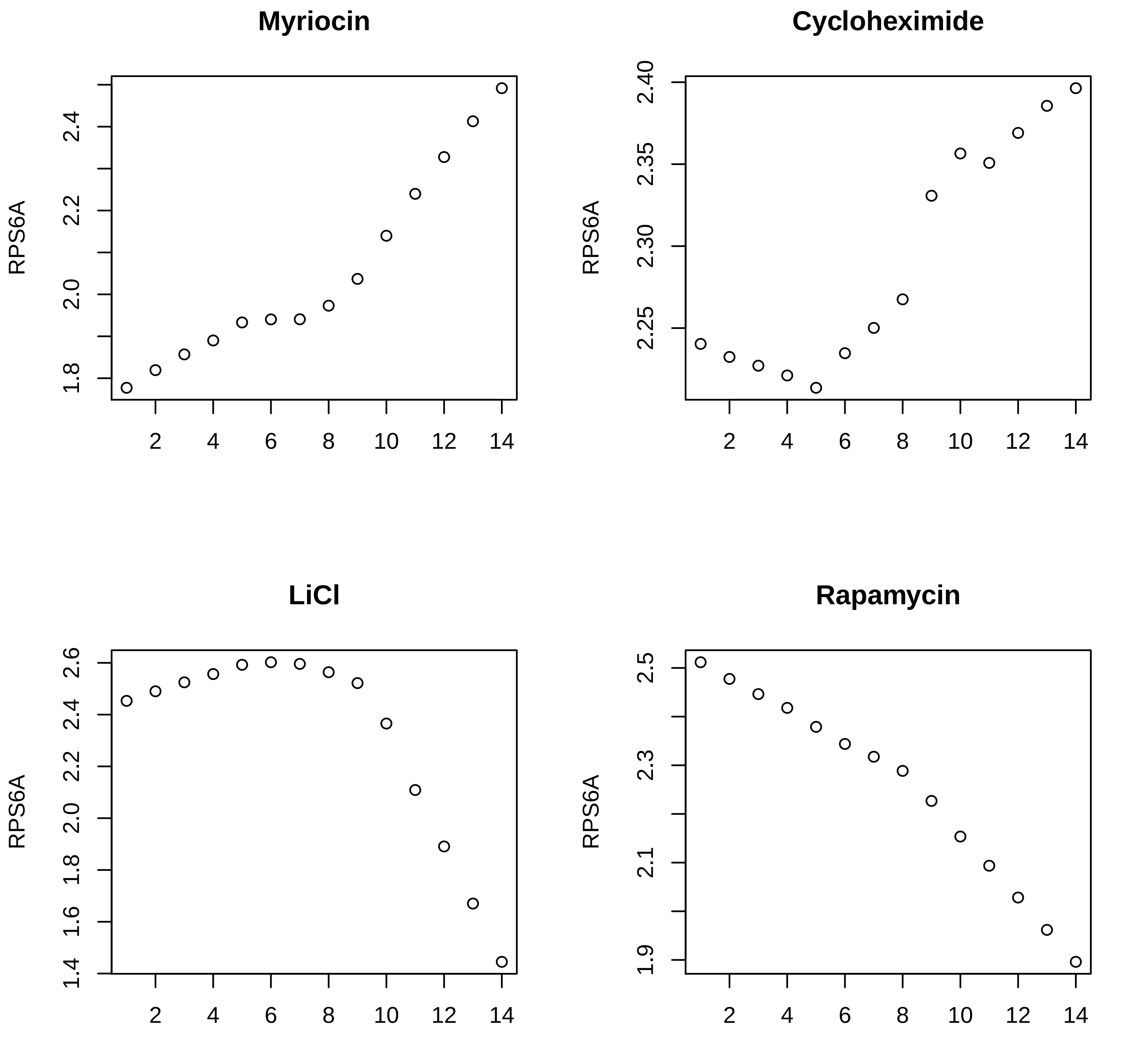
Myriocin
RPS6A
1.8
2.0
2.2
2.4
2
4
6
8
10
12
14
Cycloheximide
RPS6A
2.25
2.30
2.35
2.40
2
4
6
8
10
12
14
LiCl
RPS6A
1.4
1.6
1.8
2.0
2.2
2.4
2.6
2
4
6
8
10
12
14
Rapamycin
RPS6A
1.9
2.1
2.3
2.5
2
4
6
8
10
12
14

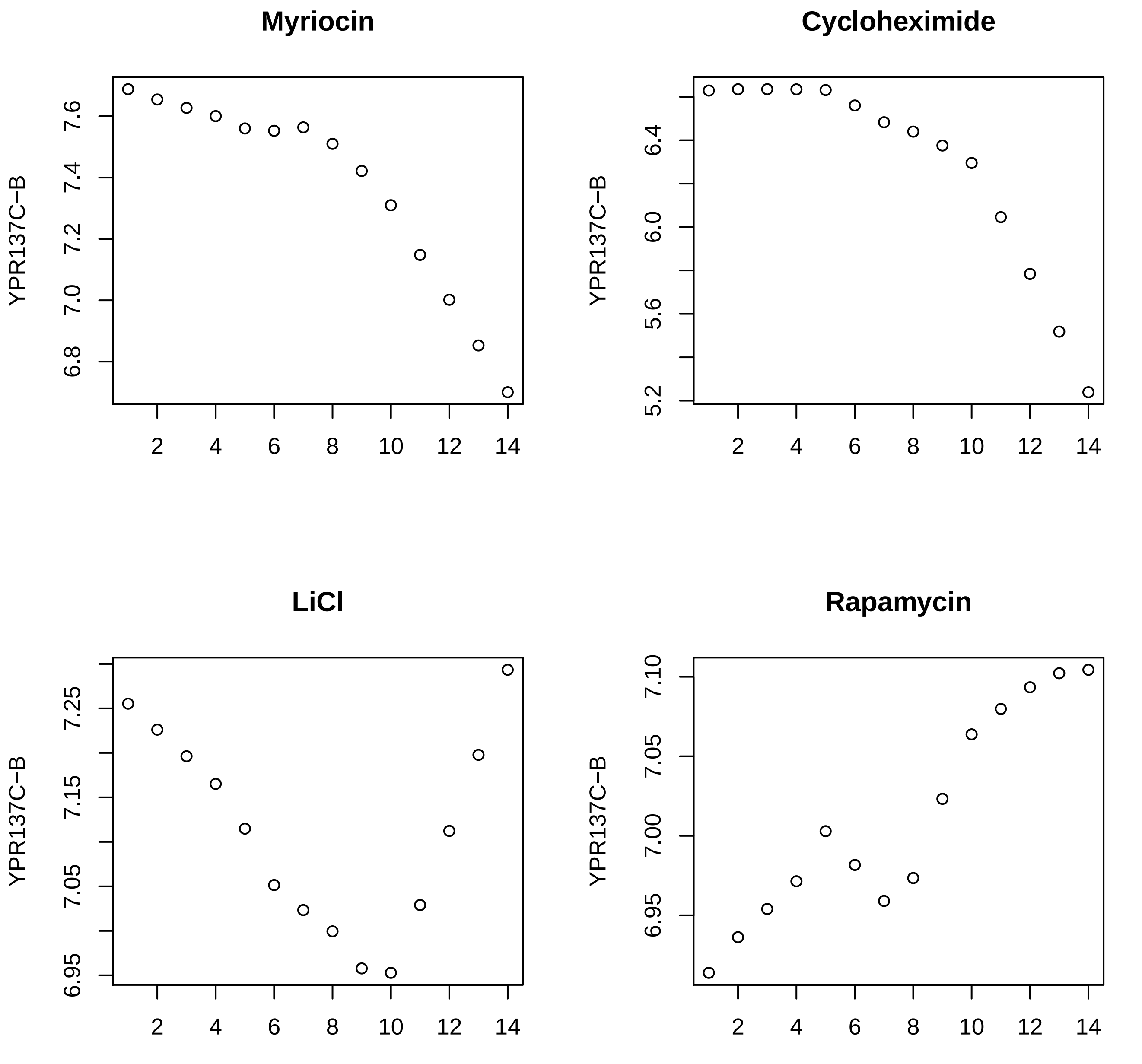
Myriocin
YPR137C−B
7.6
7.4
7.2
7.0
6.8
2
4
6
8
10
12
14
Cycloheximide
YPR137C−B
6.4
6.0
5.6
5.2
2
4
6
8
10
12
14
LiCl
YPR137C−B
7.25
7.15
7.05
6.95
2
4
6
8
10
12
14
Rapamycin
YPR137C−B
7.10
7.05
7.00
6.95
2
4
6
8
10
12
14

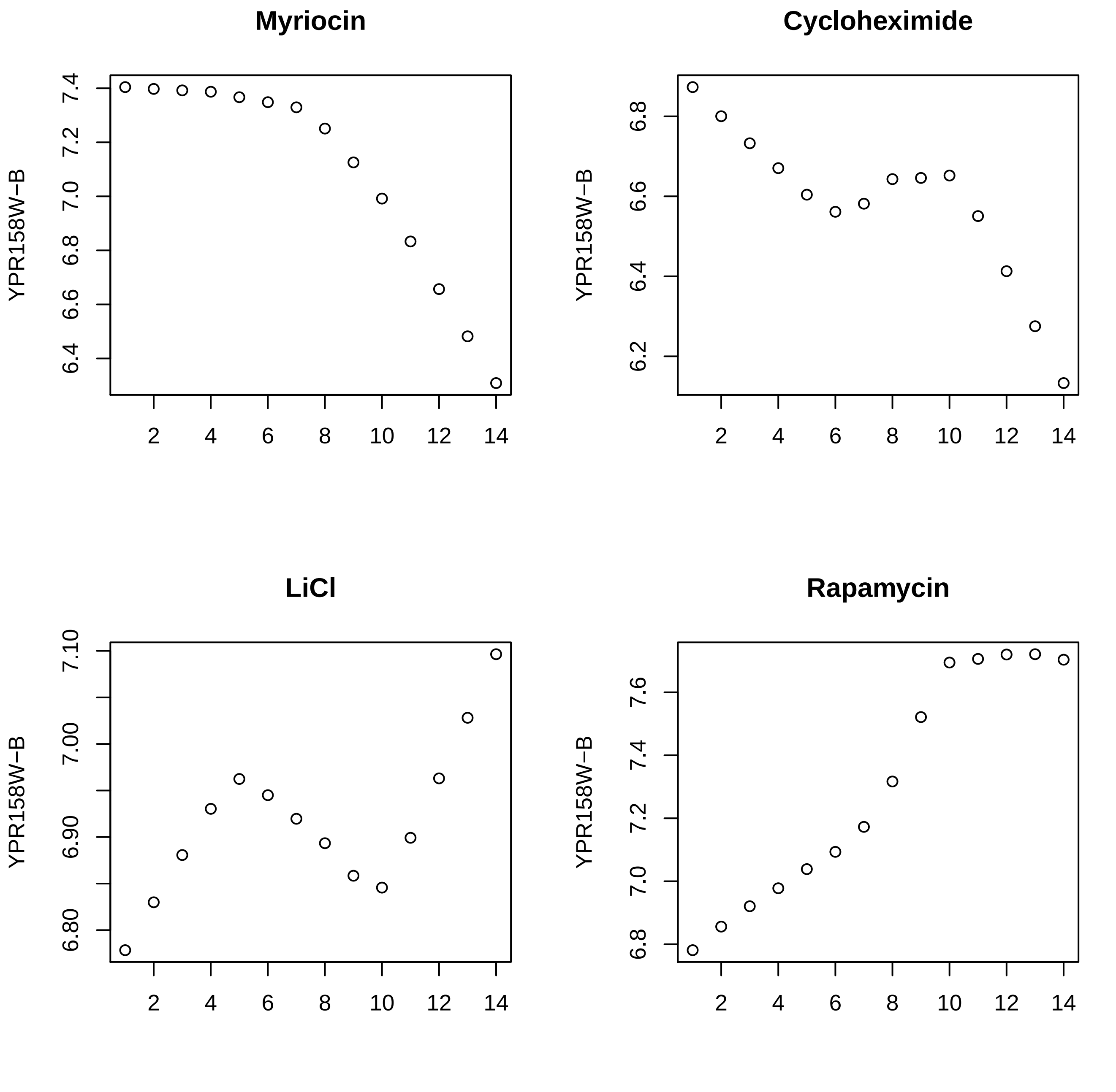
Myriocin
Cycloheximide
LiCl
Rapamycin
YPR158W−B
YPR158W−B
YPR158W−B
YPR158W−B

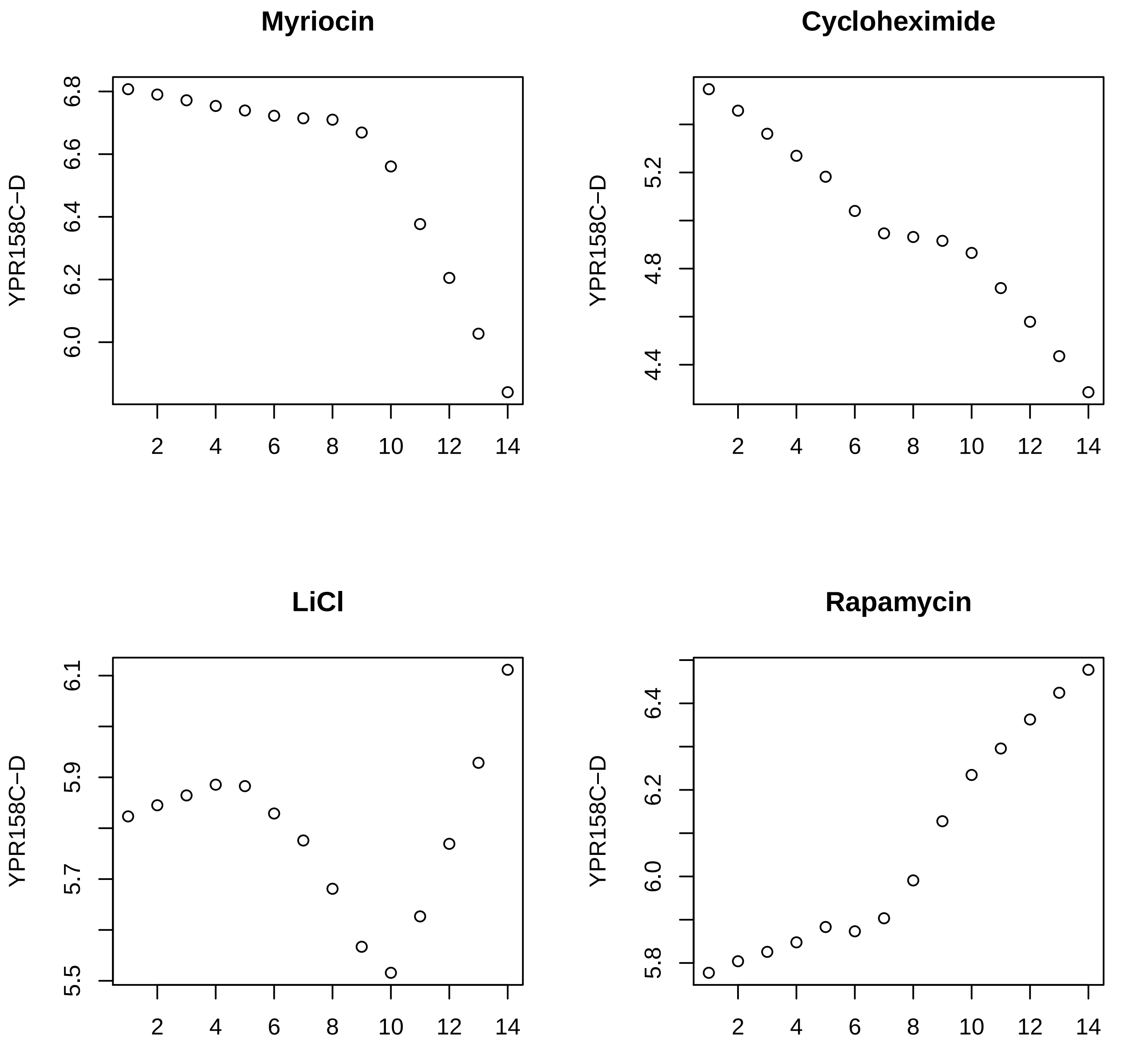
Myriocin
YPR158C−D
6.8
6.6
6.4
6.2
6.0
2
4
6
8
10
12
14
Cycloheximide
YPR158C−D
5.2
4.8
4.4
LiCl
YPR158C−D
6.1
5.9
5.7
5.5
Rapamycin
YPR158C−D
6.4
6.2
6.0
5.8

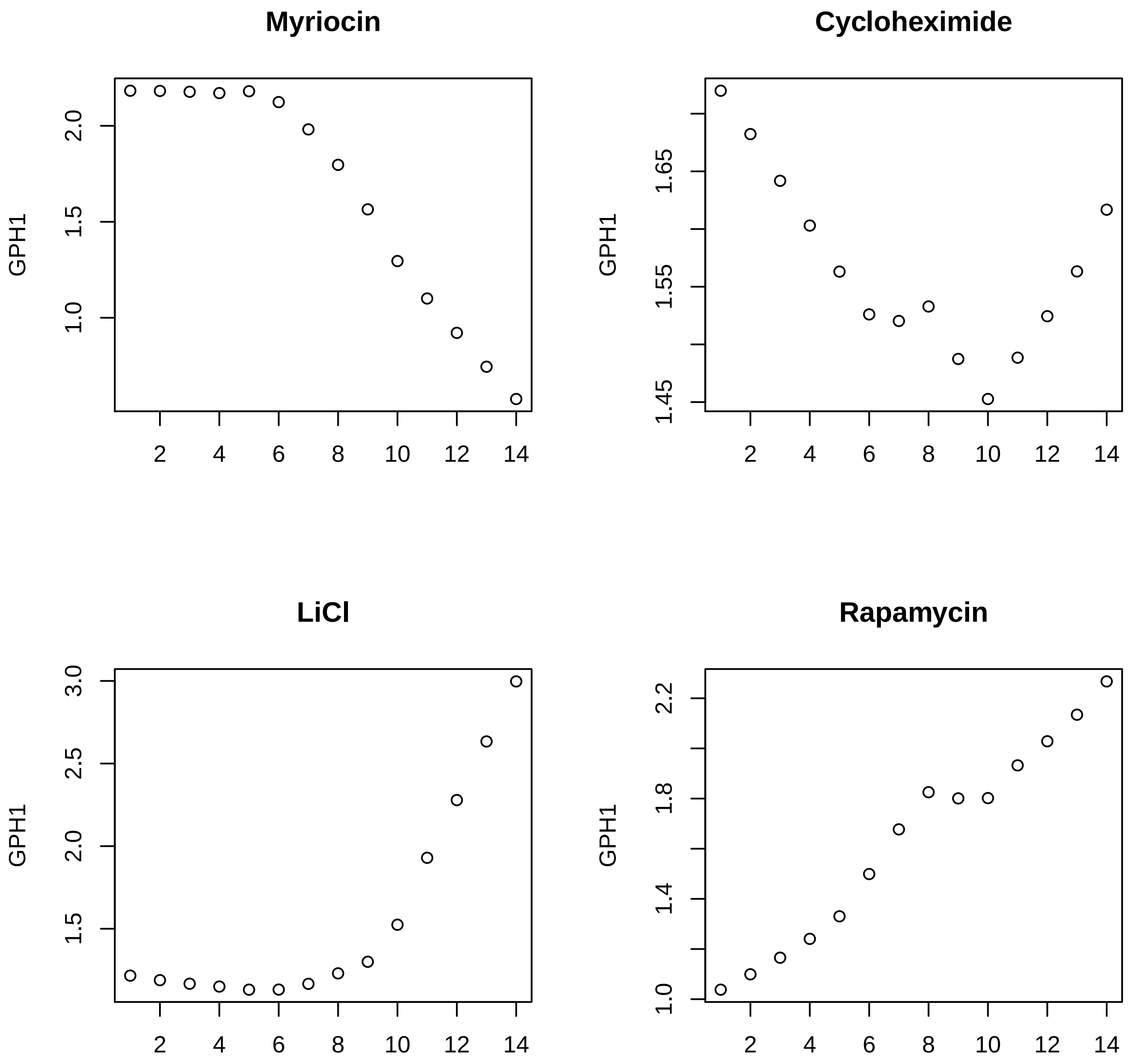
Myriocin
Cycloheximide
LiCl
Rapamycin
GPH1
2.0
1.5
1.0
1.65
1.55
1.45
3.0
2.5
2.0
1.5
2.2
1.8
1.4
1.0
2
4
6
8
10
12
14
