## Supplementary material for "Novel method for the prediction of drug-drug Interaction based on Gene Expression profiles": Supp_fig.pdf

### Supplementary Figures

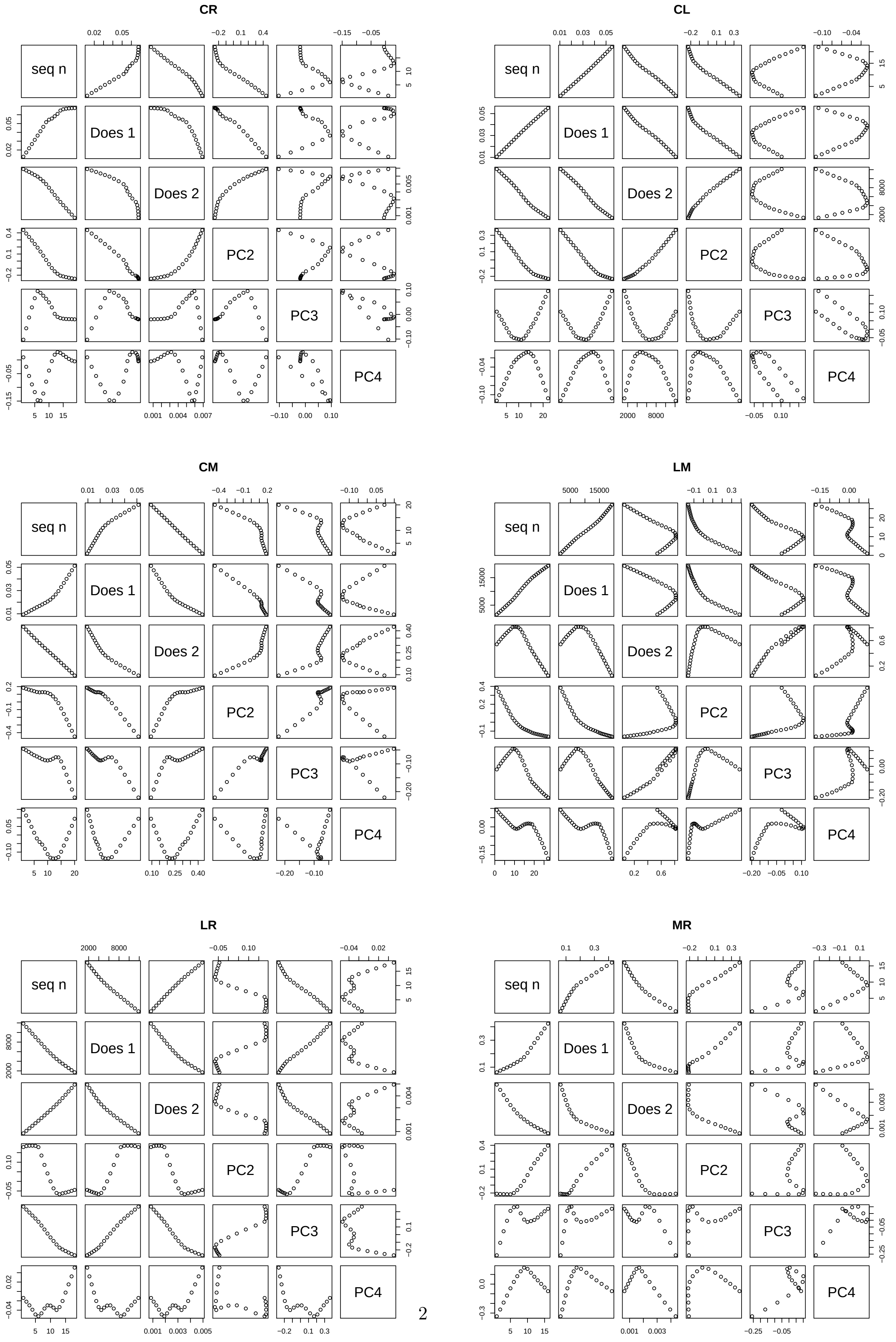


Figure S1: Scatter plots of $j$, dose densities of the first and second drug, and the second to fourth PC loadings. All values are Lowess-smoothed. Sequential numbers are ordered along iso-growth rate lines identified in the previous study [1] as much as possible. Two letters above each panel show the combinations of drugs: M: Myriocin C: Cycloheximide, L: LiCl, R: Rapamycin

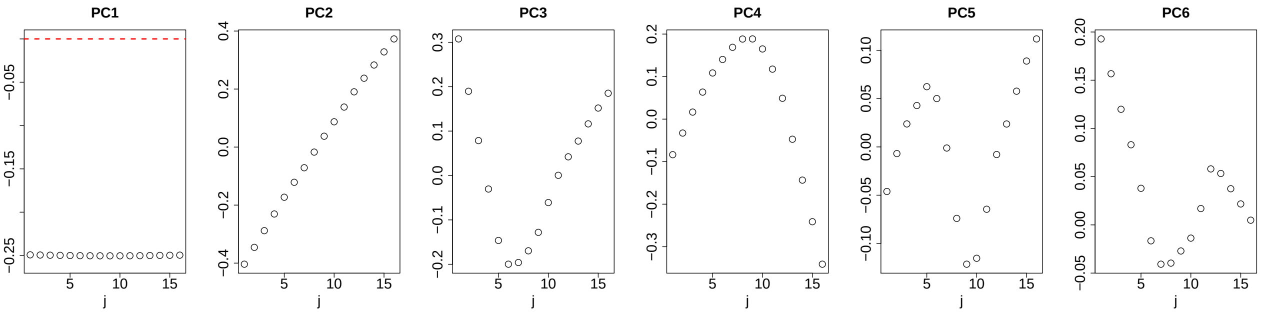


Figure S2: Lowess-smoothed $u_{\ell_1 j}, 1 \leq \ell_1 \leq 6$ for combinatorial drug treatments.

(A)

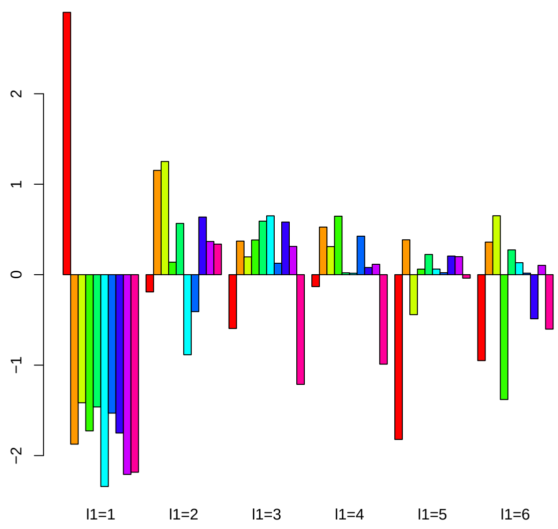


(B)

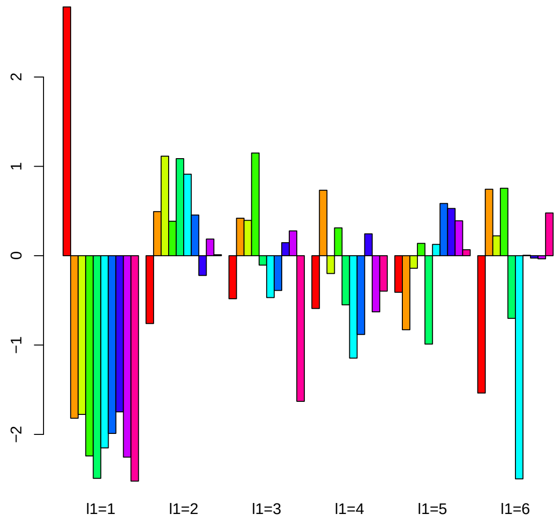


Figure S3: Common logarithmic absolute values of $G(\ell_1\ 1\ \ell_3)$ for combinatorial (A) or single (B) drug treatments. For each $\ell_1$, $G(\ell_1\ 1\ \ell_3)$ values are aligned from left to right in increasing order of $\ell_3$. The same colors correspond to the same $\ell_3$.

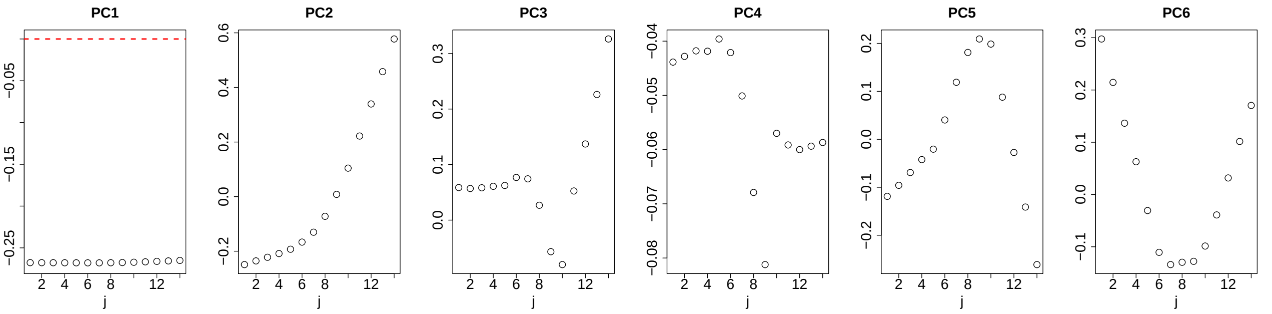


Figure S4: Lowess-smoothed $u_{\ell_1 j}, 1 \leq \ell_1 \leq 6$ for single drug treatments.
